# Mapping fibrotic microenvironments: single-cell and spatial profiling of Schistosoma mansoni-induced tissue fibrosis

**DOI:** 10.64898/2026.04.29.721785

**Authors:** Veronika Tóth, P. Prakrithi, Zherui Xiong, Malcolm Jones, Chaoyi Li, Zhian Chen, Juan C. del Alamo, Yi-Ting Yeh, Grant A. Ramm, Glicia De Almeida, Geoffrey N. Gobert, Quan Nguyen, Hong You

## Abstract

*Schistosoma*-induced tissue fibrosis, driven by immune response to trapped schistosome eggs, is the principal cause of pathology and schistosomiasis-related morbidity. While praziquantel effectively clears adult parasites, no therapies exist to target eggs or egg-induced tissue fibrosis. Here, we utilise single-cell spatial transcriptomics to define the high-resolution molecular architecture of the fibrotic niche in *Schistosoma mansoni*-infected mice at 8 weeks post-infection. Our data reveal that the host executes divergent, organ-specific biological programs: constructing a rigid, concentric fibrotic granuloma for permanent egg sequestration in the liver while building a flexible, discontinuous collagen architecture in the intestine to facilitate egg transit. Spatially resolved cell-cell interaction analysis, unprecedented in schistosome-induced fibrosis, identifies the strongest cellular interactions between macrophages and collagen-producing cells, with TGF-*β*-dominant ligand-receptor pairs in the hepatic niche and integrin-centered pairs in the intestine. Despite this architectural divergence, we detected a core signature of five ligand-receptor pairs shared across both tissues, suggesting shared signalling interactions that could be leveraged for pan-tissue therapeutics for treating schistosome-induced fibrosis. Together, these findings provide the first high-resolution spatial atlas of schistosomiasis-associated fibrosis and provide a foundational framework for discovering novel drug targets capable of reversing established tissue damage, addressing a critical clinical gap where current therapies fail.

## 1. Introduction

Schistosomiasis is a devastating neglected tropical disease caused by parasites of the genus *Schistosoma* that remains a major global health challenge, affecting over 250 million people across 79 countries [1, 2]. The infection cycle begins when infective cercariae penetrate host skin in freshwater and migrate to the mesenteric vasculature, where they develop into mature adult worms [3]. The main immunopathology of intestinal schistosomiasis, primarily caused by *Schistosoma mansoni* and *S*. *japonicum*, is driven by the host immune responses to parasite eggs that are continuously deposited by female worms and become trapped predominantly in the liver and intestine [3–5]. These eggs are highly immunogenic, producing a broad range of excreted-secreted products, including T2 ribonuclease omega-1, IPSE/α1, kappa-5, SmHMGB1, and SmCKBP, which trigger robust granulomatous inflammation [6–11]. Yet, despite the growing number of identified egg-derived antigens, a spatially resolved understanding of the cellular neighbourhoods, ligand-receptor (LR) networks, and fibrogenic programs is limited. Although granulomas initially form to sequester parasite eggs and protect the host from egg-derived toxins, their progression into a chronic fibrotic state drives the ultimate pathology of the disease [12–16]. This progression is defined by the cumulative impact of fibrotic structures, which progressively compromise organ integrity and lead to severe clinical manifestations. Granulomas form around eggs trapped in host tissues and, over time, evolve into collagen-rich fibrotic lesions that cause the organs to become increasingly scarred [15]. In the liver, this structural destruction can obstruct portal blood flow, causing portal hypertension, oesophageal varices, and life-threatening gastrointestinal bleeding [17, 18], while in the intestine, pathological changes can give rise to ulcers, haemorrhoids, polyps, creating a chronic inflammatory state associated with an increased risk of colorectal cancer [19–23].

Current antiparasitic treatment with praziquantel (PZQ), the sole drug used for schistosomiasis, effectively clears mature adult worms but has limited effects on eggs and on mitigating egg-induced pathology, although early use may partially prevent fibrosis [24–27]. Furthermore, no FDA-approved therapies exist for preventing or reversing established hepatic/intestinal fibrosis [27, 28], highlighting a major unmet need in schistosomiasis and other fibrotic diseases. A key barrier to progress is the limited mechanistic understanding of parasite-host interactions that drive fibrosis at the spatial cellular level within tissues. Schistosome-induced fibrosis is a highly microenvironment-driven process in which pathogenic signalling depends on the spatial organisation and interactions among egg antigens, stromal cells, infiltrating immune cells, and damaged parenchyma. Although advanced single-cell RNA sequencing (scRNA-seq) has begun to map the transcriptional atlas of liver granulomas [29–32], these dissociative approaches are intrinsically limited because they disrupt tissue architecture and cannot resolve where specific profibrotic programs occur and which cell populations are spatially adjacent or interacting. Single-cell spatial transcriptomics (SC-ST) overcomes these limitations by measuring gene expression in intact tissues while preserving the spatial location of individual cells and the original tissue structure [33]. This enables the identification of spatially defined cell states, functions, microenvironments, and cell-cell interactions and, with high-resolution platforms, can reach near single-cell or subcellular resolution. In recent years, SC-ST has seen increasing application across biomedicine [34–36], including in the study of parasitic infections such as alveolar echinococcosis [37], the filarial nematode *Brugia malayi* [38], *Trypanosoma brucei* [39], and the *Plasmodium berghei* model of malaria [40]. Thus, enabling high-resolution mapping of both host and parasite transcriptional landscapes within intact tissues. Building on these advances, SC-ST is uniquely positioned to reveal cellular circuits that sustain fibrosis and host-pathogen interactions, enabling identification of antifibrotic targets and biomarkers inaccessible to conventional omics.

In this study, we employed the 10x Genomics Xenium SC-ST platform to investigate the molecular and cellular architecture of *S. mansoni* egg-induced granuloma at 8 weeks post-infection (wpi) across two tissues, livers and intestines, in a murine model. We spatially mapped gene expression within affected tissue sections and validated these patterns at both the transcript and protein levels using quantitative polymerase chain reaction (qPCR) and immunohistochemistry. Finally, we integrated transcriptional profiles with spatial localisation and found specific cellular and molecular interactions in hepatic and intestinal granulomas. By resolving the spatial heterogeneity of the fibrotic niche, this work defines the molecular programs distinguishing hepatic and intestinal pathology while identifying both organ-specific and shared signalling hubs as potential novel targets for schistosome-induced fibrosis.

## 2. Results

To resolve the high-resolution spatial organisation of the fibrotic niche in schistosomiasis at 8 wpi, we employed SC-ST using the 10x Genomics Xenium Mouse Expression Panel targeting 5,006 genes (**Fig. 1A**). The analysis of the single-cell resolved spatial transcriptomic data was performed on selected regions of interest (ROIs) from healthy and infected regions, allowing comparison across the gradient from inflamed to healthy areas. After quality control filtering, leiden clustering of the retained cells identified clusters at a resolution of 0.3 (res=0.3). In total, we obtained 73 million transcripts at the single-cell level from >534k cells obtained from 6 liver sections and 30 million transcripts from >259k cells obtained from 6 intestine sections.

**Figure 1.**
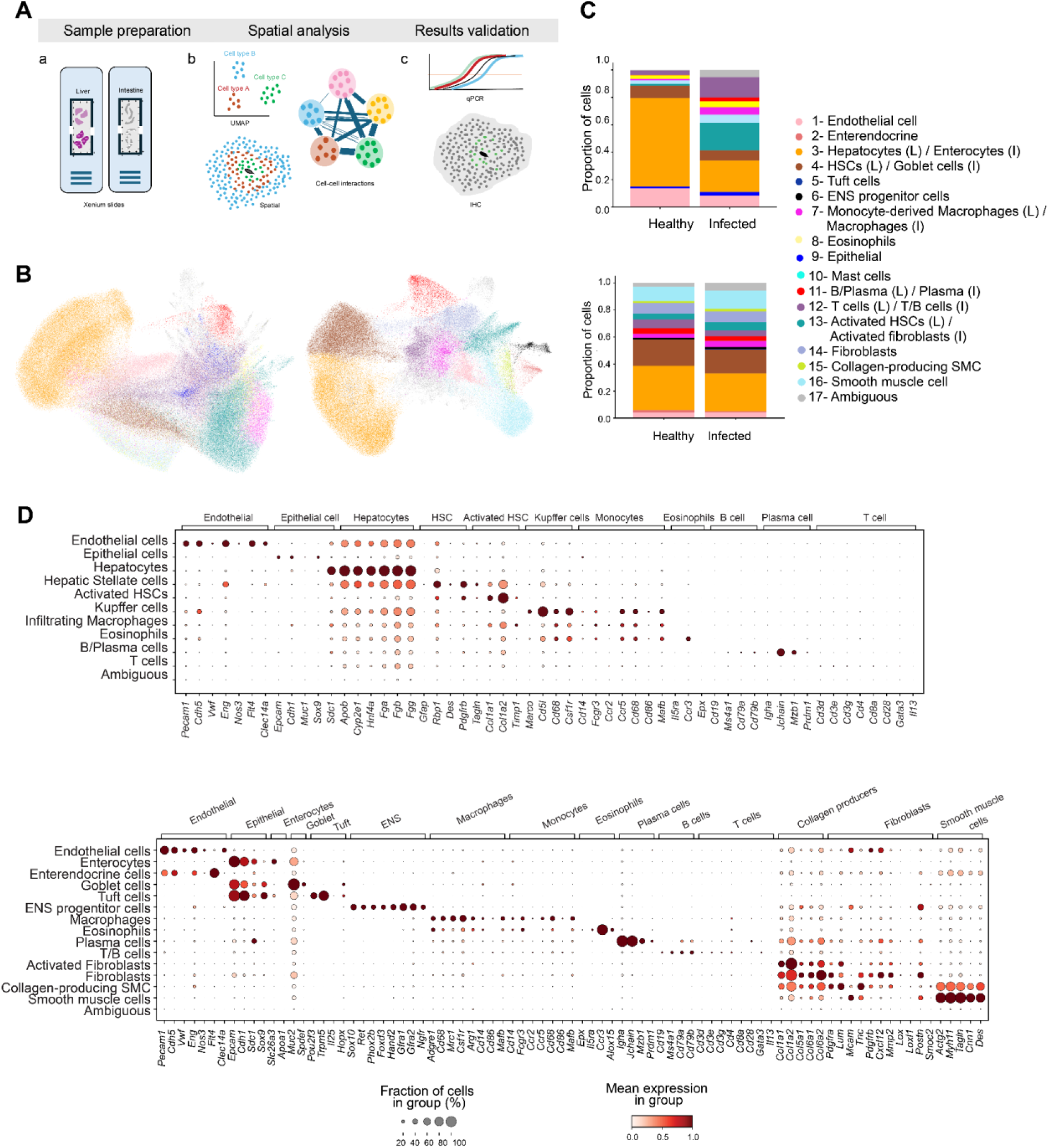
Cell-typing of spatial transcriptomics data from *Schistosoma mansoni-*infected mouse tissues. **(A)** Graphical representation of the workflow involving Xenium *in situ* experiments and analysis of normal and infected tissues from the liver and intestine, followed by experimental validations of genes prioritised from the analysis. **(B)** Cell type UMAPs of the liver (right) and the intestine dataset (left). **(C)** Differences in cell-type proportions between the healthy and infected tissues of the liver (top) and intestine (bottom). **(D)** Dot plots of the distinguishing markers for the cell types identified in the liver (top) and intestine (bottom). The markers are known canonical markers for the cell types shown at the top of the plots.

### 2.1 Spatial architecture of fibrotic niche in hepatic and intestinal granulomas

We next dissected the cellular composition and spatial organisation of granulomas across both organs. In the liver, we identified 10 main cell types, including endothelial cells, epithelial cells, hepatocytes, HSCs, activated HSCs, resident Kupffer cells, monocyte-derived (infiltrating) macrophages, eosinophil-like cells, B/plasma cells, and T cells. Fifteen cell types were identified in the intestine, which included enterocytes, goblet cells, enteric nervous system (ENS) progenitor cells, smooth muscle cells (SMCs), fibroblasts, T/B cells, activated fibroblasts, macrophages, eosinophils, mast cells, endothelial cells, plasma cells, collagen-producing SMCs, enteroendocrine cells, and tuft cells **(Fig. 1B**, **Fig. 1C**, **Fig. 1D).** A small fraction of cells in both tissues showed ambiguous clustering and lacked clear enrichment of canonical marker genes, which were classified as ambiguous.

H&E, Sirius Red staining, and spatial transcriptomic analysis were used to characterise tissues from *S. mansoni*-infected and healthy control (HC) mice. Healthy liver sections showed the expected architecture, with parenchyma composed mainly of hepatocytes, endothelial cells, and quiescent HSCs **(Fig. 1C)**. The portal tracts were thin and free of inflammatory infiltrates **(Fig. 2A)**. Similarly, healthy intestine sections displayed a well-organised lamina propria, mucosa, and submucosa, without pathological changes. Major cell types in healthy intestine regions included enterocytes, goblet cells, smooth muscle cells, and fibroblasts **(Fig. 1C**, **Fig. 2B)**. In contrast, tissues from *S. mansoni*-infected mice displayed markedly altered architecture: H&E staining confirmed the presence of *S. mansoni* eggs and granuloma at 8 wpi, while Sirius Red staining revealed extensive collagen deposition around the eggs in both tissues, indicating persisting fibrosis. In the liver, egg-surrounding granulomas were large and highly cellular, distributed throughout the parenchyma, and exhibited a highly organised architecture consisting of a central core and peripheral regions. The core regions consisted predominantly of monocyte-derived macrophages and activated, collagen-producing HSCs, while the periphery of the granuloma was enriched with T cells and B/plasma cells **(Fig. 2A)**. Intestinal granulomas formed around eggs deposited in the submucosal layer of the large intestine and exhibited a discontinuous, lower-density collagen network **(Fig. 2B)**. The dominant cell types present in these regions were macrophages, collagen-producing fibroblasts, T cells, and B cells **(Fig. 1C**, **Fig. 2B**). Cell type distributions from the remaining tissues used in the study are provided in **Supplementary Fig. 1.**

**Figure 2.**
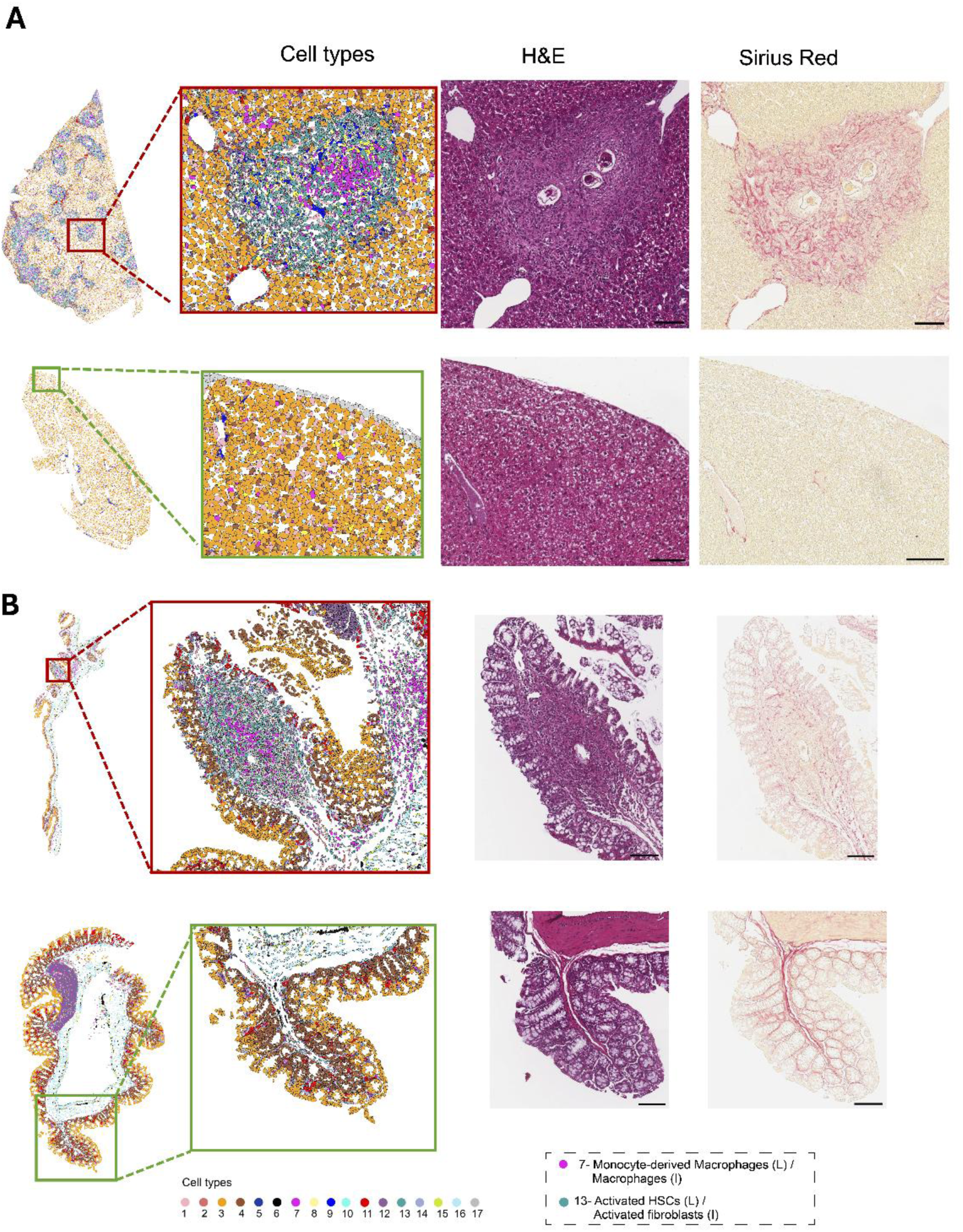
Integrated histological and spatial characterisation of *Schistosoma mansoni-*induced granulomas and fibrosis regions in murine tissues. Healthy and infected. **(A)** liver and **(B)** intestinal sections were collected from BALB/c mice at 8 weeks post-infection with *S. mansoni* cercariae. Tissue architecture is markedly disrupted in infected samples (A, top; B, top) relative to healthy controls (A, bottom; B, bottom). Spatial cell-type maps derived from single-cell spatial transcriptomics illustrate the cellular composition of fibrotic granulomas in hepatic and intestinal tissues. Hepatic granuloma cores are enriched for monocyte-derived macrophages and activated, collagen-producing hepatic stellate cells, with T cells and B/plasma cells predominantly localised to the periphery. Intestinal granulomas are primarily composed of macrophages, collagen-producing fibroblasts, and T cells and B cells. The encoded cell types are 1- Endothelial cells, 2- Enterendocrine cells, 3- Hepatocytes (L) / Enterocytes (I), 4- HSCs (L) / Goblet cells (I), 5- Tuft cells, 6- Enteric Nervous System progenitor cells, 7- Monocyte-derived Macrophages (L) / Macrophages (I), 8- Eosinophils, 9-Epithelial cells, 10- Mast cells, 11- B/Plasma cells (L) / Plasma cells (I), 12- T cells (L) / T/B cells (I), 13- Activated HSCs (L) / Activated fibroblasts (I), 14- Fibroblasts, 15- Collagen-producing smooth muscle cells, 16- SMCs, 17- Ambiguous. (L) and (I) indicate different nomenclatures in the liver and intestine, respectively. The key cell types are named in the dotted box. Hematoxylin and eosin (H&E) staining highlights trapped *S. mansoni* eggs surrounded by concentric granulomatous cell layers and adjacent parenchymal cells. Sirius Red staining of adjacent sections reveals collagen deposition (red), indicating fibrotic regions that spatially overlap with the granuloma.

### 2.2 Spatially resolved cell-cell interaction networks in schistosome-induced fibrosis

Having defined the cellular architecture of granulomas in both organs, we next asked which intercellular communication axes might underlie the divergent fibrotic phenotypes. We performed spatially-informed cell-cell interaction (CCI) analysis on the single-cell-resolved Xenium data, inferring candidate LR axes from co-expression in proximal cells (**Supplementary Table 1**). We dissected these predicted communication networks within hepatic and intestinal granulomas in turn, and then performed and overlap analysis of the two networks to identify LR pairs shared between the two niches.

#### 2.2.1 A TGF-β–centred macrophage–activated HSC axis defines the hepatic fibrotic niche

In uninfected hepatic tissue, the predicted CCI landscape was dominated by interactions between hepatocytes and quiescent HSCs, including homotypic interactions **(Fig. 3)**. The integrin receptor *Itga5* emerged as the most highly connected node, interacting with *Col18a1*, *Vtn* and *Angptl3* in a homeostatic ECM-receptor program (**Supplementary Table 1A**). At 8 wpi, this landscape was substantially remodelled: Liver granulomas exhibited a strong global enrichment of integrin-mediated interactions (16/20 of the top-ranked LR pairs), a feature we found to be conserved between the two organs and addressed in detail in Section 2.2.3. When the analysis was restricted to the most strongly interacting cell pair, the network resolved into a dominant axis between activated HSCs and infiltrating macrophages within the granuloma core, with substantial homotypic interactions within each population (**Fig. 3A**). The highest-scoring predicted LR pairs within the aHSC-macrophage axis were *Col1a2* and *Itgb3*, *Tgfb3* and *Tgfbr2*, and *Fn1* and *Nt5e* (**Supplementary Table 1B**). Notably, a TGF-β pathway-related LR pair (*Tgfb3* and *Tgfbr2*) ranked among the top three interactions, and examination of the remaining high-ranking LR interactions revealed persistent representation of additional TGF-β-associated LR pairs, including *Tgfb3* and its receptors *Tgfbr1*, *Tgfbr3*, and *Eng*, *Tgfb1* and *Itgb8.* In addition, several indirectly related interactions were observed, consistent with the enrichment analysis of the LR-derived gene set. These findings suggest that the TGF-β signalling pathway is highly active between infiltrating macrophages and aHSCs. To strengthen these transcriptomic predictions, we mapped the inferred interacting populations and their associated LR-expressing cells back onto the Xenium coordinate space. The top-scoring LR pairs were spatially co-distributed within the granuloma core, with ligand- and receptor-expressing cells positioned in close proximity (**Fig. 3B**), providing direct spatial support for the predicted LR axes.

**Figure 3.**
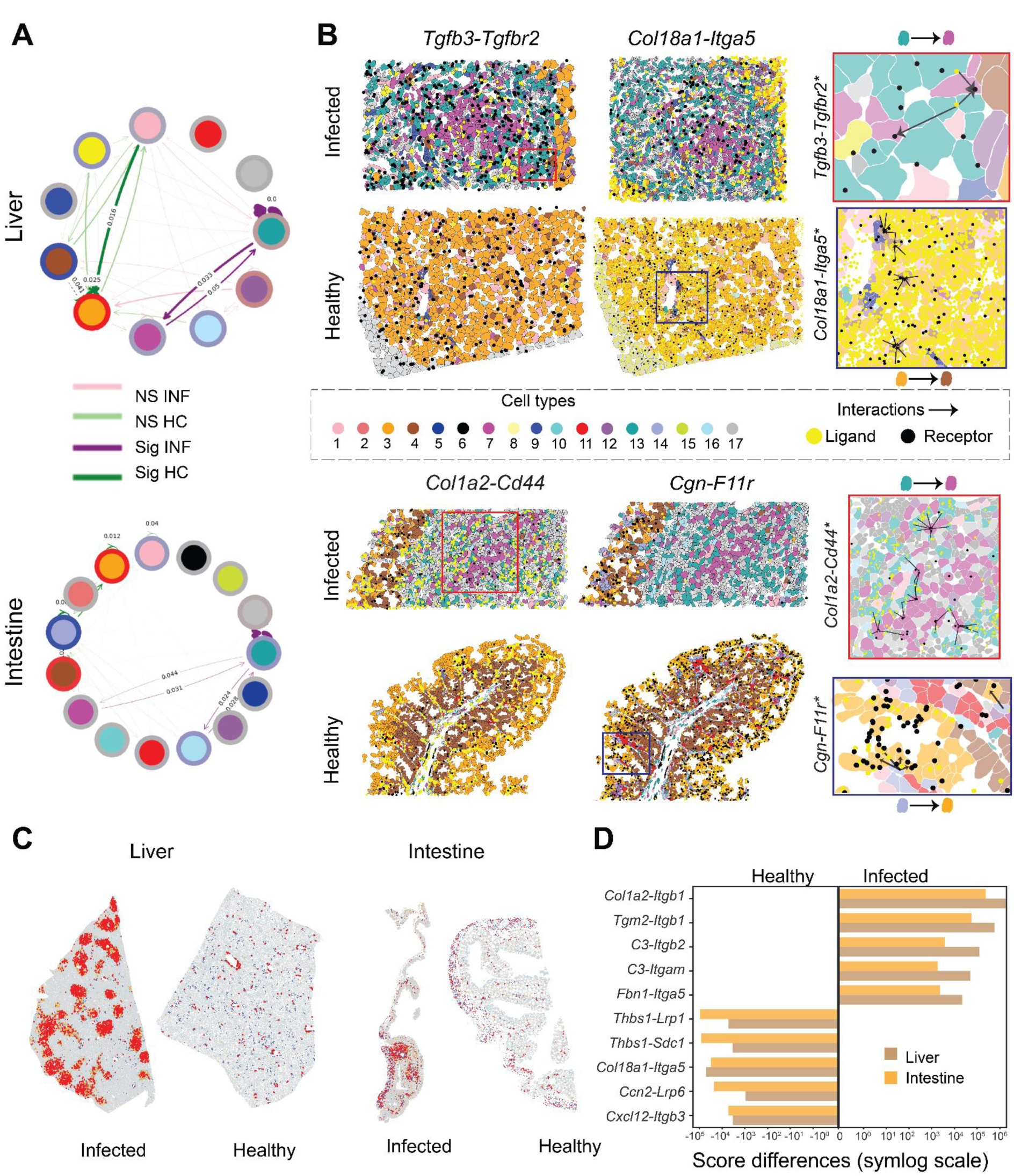
Spatial transcriptomics detects shared spatially coexpressed ligand-receptor (LR) pairs in hepatic and intestinal granulomas. **(A)** Comparing differential interactions at the cell type (nodes) level for infected vs. healthy samples in the liver (top) and intestinal (bottom) datasets using all LR pairs. The purple arrows show more interactions in the infection, and green arrows indicate more interactions in healthy tissues. **(B)** Spatial mapping of the LR pairs on the Xenium data. The polygons are colored by the cell types using the same color scheme as in Figure 2. Black and yellow dots overlaid on the cell types represent the ligand and receptor transcripts, respectively. The L-R* spatial plots shows interactions between top-predicted cell types for zoomed-in regions of corresponding larger regions of interest outlined by color-coded boxes. (*Tgfb3*-*Tgfbr2* for aHSCs-Monocyte derived macrophages in infected liver, *Col18a1*-*Itga5* for Hepatocytes-HSCs in healthy liver, *Col1a2*-*Cd44* for Activated fibroblasts-Macrophages in infected intestine and *Cgn*-*F11r* for Fibroblasts-Enterocytes in normal intestine). (C) *Col1a2-Itgb1* coexpression (red) in healthy and infected liver and intestinal samples. The red dots indicate spatial co-expression of the ligand and the receptor; light blue and orange represent the expression of either the ligand or the receptor; dark blue represents the absence of both; and light gray dots indicate non-significant cells. **(D)** LR pairs with the highest overall interaction in the infected and healthy samples across both the tissues.

#### 2.2.2 An integrin-centred activated fibroblast-macrophage axis defines the intestinal fibrotic niche

In uninfected intestinal tissue, the dominant predicted cell-pair axis linked fibroblasts and enterocytes through LR pairs, including *Cgn*-*F11r*, *Afdn*-*F11r*, and *Efnb1*-*Erbb2* (**Supplementary Table 1B**). Independent of this axis, the integrin receptor *Itgb1* emerged as the most highly connected receptor across all cell types in the healthy tissue, engaging a broad repertoire of basement membrane ligands, including *Lama2*, *Lamb1*, *Col4a1*, *Col18a1*, and *Hspg2*, alongside structural partners *Col1a1*, *Fbn1*, *Vcan*, and *Thbs* (**Supplementary Table 1A**). At 8 wpi, this cellular landscape had undergone extensive remodelling. Similar to hepatic granulomas, intestinal granulomas exhibited a strong global enrichment of integrins, a conserved cross-organ signature - addressed in Section 2.2.3. The *Itgb1* interactions reorganise to *Col1a2* and *Tgm2*, indicating a shift in *Itgb1*-engaged ligands from a homeostatic basement-membrane programme to a fibrotic ECM programme. Furthermore, at the cell-pair level, the network centered around an activated fibroblast-macrophage axis is enriched within the submucosal granuloma core, with substantial homotypic interactions (**Fig. 3B**). Within this axis, the highest-scoring predicted LR pairs were *Col1a2* and *Cd44*, *App* and *Lrp1*, and *Fn1* and *Itga5*, with additional integrin-family receptors also engaged (*Itgb7*, *Itgb3*, and *Itga1*; **Supplementary Table 1B**). To strengthen these predictions, we mapped activated fibroblasts and macrophages and their cognate LR-expressing cells back onto the Xenium coordinate space. The top-scoring LR transcripts were spatially co-distributed within the granuloma core (**Fig. 3B**), providing spatial support for the predicted axes beyond the cell-type co-occurrence already established by the architectural analysis (Section 2.1).

#### 2.2.3 LR analysis identifies shared integrin-centred signalling networks in schistosome-induced granulomas

To identify the molecular features common to both schistosome-induced fibrotic niches, we compared the predicted LR networks of hepatic and intestinal granulomas at 8 wpi at two levels of resolution. At the receptor-family level, integrins were broadly represented among the top-ranked LR pairs of both organs (Sections 2.2.1, 2.2.2), establishing integrin-mediated signalling as a programme-wide feature of the fibrotic communication landscape rather than an organ-restricted finding. At the LR-pair level, intersecting the two networks identified five interactions that were directly shared - *Col1a2*-*Itgb1*, *Tgm2*-*Itgb1*, *C3*-*Itgb2*, *C3*-*Itgam*, and *Fbn1*-*Itga5* (**Fig. 3C**, **Fig. 3D**) - all engaging integrin receptors. Together, these analyses identify integrin-mediated cell-surface interactions as the principal conserved signature of the fibrotic niche, anchored broadly by family-wide integrin engagement and concentrated at the LR-pair level on recurrent *Itgb1*, *Itgb2*, and *Itga5* partnerships.

### 2.3 Differential gene expression and pathway analyses across schistosome-induced fibrotic niches

Having identified the dominant communication axes within each fibrotic niche, we next asked which broader transcriptional programmes underpin the response to schistosome eggs in each organ. We performed differential expression analysis between infected and uninfected tissue in the liver and intestine, prioritising genes by both statistical significance and magnitude of change (logarithm of fold-change, logFC), to define a high-confidence set of transcriptional changes at 8 wpi **(Fig. 4C, Supplementary Table 2)**. Differential gene expression analysis revealed widespread transcriptional remodelling within granulomatous regions in both organs, with 1,061 upregulated and 114 downregulated genes in the liver and 303 upregulated and 504 downregulated genes in the intestine **(Fig. 4A)**.

**Figure 4.**
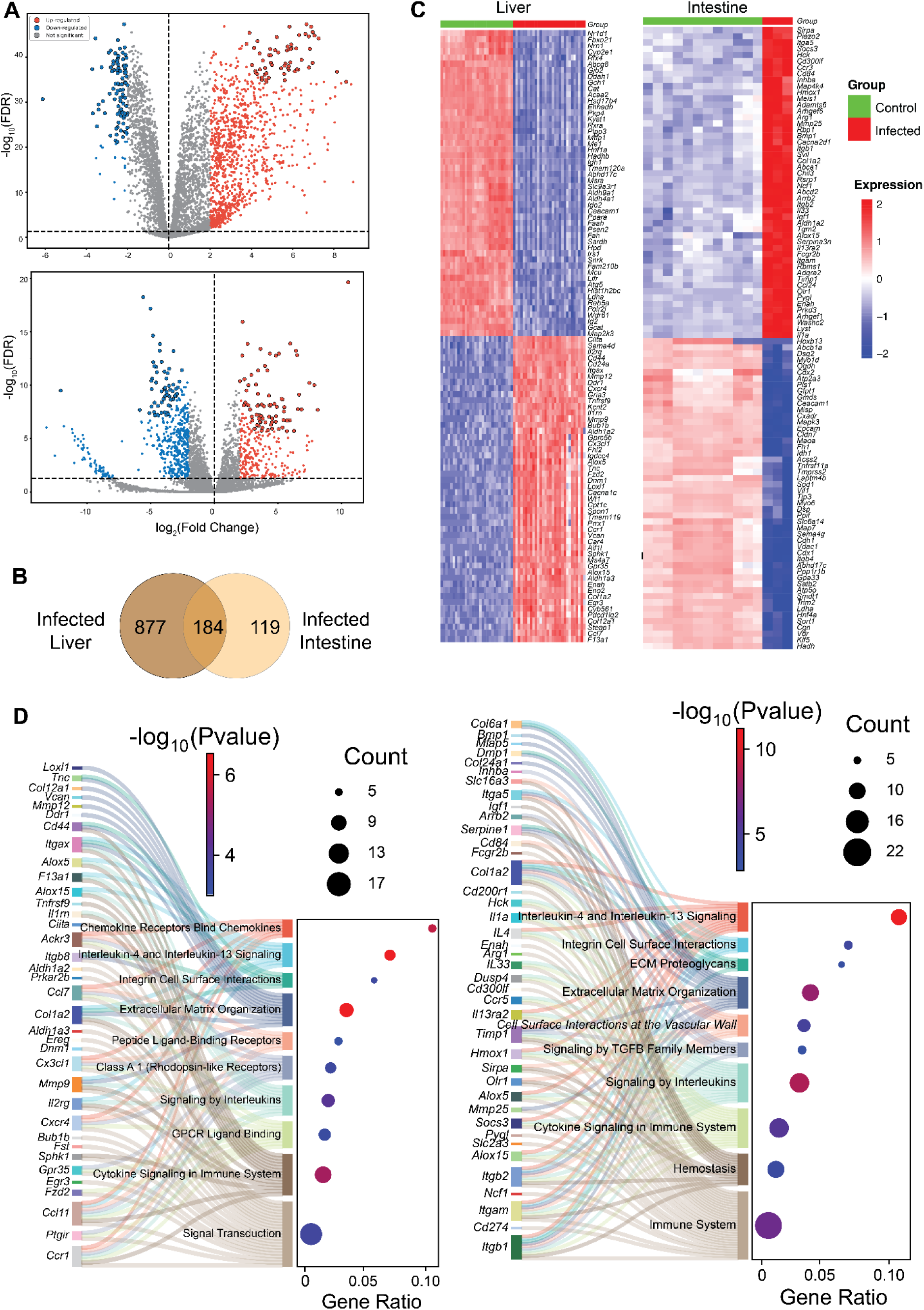
Transcriptomic signatures and pathway enrichment analysis reveal conserved and organ-specific gene expression patterns in schistosome-induced granulomas. **(A)** Volcano plots of genes differentially expressed in infected (red) versus healthy control (blue) samples in the liver (top) and intestine (bottom). Genes outlined in black represent the top candidate genes. **(B)** Venn diagram depicting the overlap of significantly upregulated genes in infected tissues between the liver and intestine. **(C)** Heatmaps showing the expression of differentially expressed genes from **(A)** across conditions. Each column represents a region of interest: granulomas from infected samples or randomly selected regions from healthy controls. **(D)** Reactome pathway analysis of the top differentially expressed genes from infected samples in **(A)** (left-liver, right-intestine). The Sankey-dot plot in panel **(D)** was generated using the SRplot online visualisation platform [86].

#### 2.3.1 Spatial transcriptomics of hepatic granulomas reveals a rigid, liver-specific fibrotic niche associated with permanent egg sequestration

The enrichment analysis of hepatic granulomas indicated 61 top upregulated differentially expressed genes (DEGs) that were strongly associated with extracellular matrix organization, IL-4 and IL-13 signaling, chemokine signaling, and cytokine signaling (**Table 1**, **Fig. 4D, Supplementary Fig. 2**). Among these genes, *Col1a2*, *Mmp9, Itgax*, and *Ccl11* appear the most frequently across these biological pathways, highlighting them as central molecular nodes that integrate the host’s immune recognition of the egg with subsequent pathological tissue remodeling in the liver. However, this tissue remodelling/repair response comes at a high cost to basal tissue homeostasis. Our spatial transcriptomic results suggest that as granulomas and fibrotic scars expand around trapped eggs, the functional metabolic zones of the liver are physically displaced and replaced by regions of inflammation and scarring. By applying strict thresholds, we identified 155 high-priority DEGs that were significantly downregulated in hepatic granulomas compared to HC. These downregulated genes, including *Rxra*, *Ppara*, *Nr1d1*, *Ehhadh,* and *Cyp1a2*, were strongly associated with the parenchymal functional baseline of the healthy liver, specifically involving the metabolism of lipids, amino acids, and fatty acids **(Fig. 4C)**. Additionally, comparison of hepatic and intestinal upregulated DEGs revealed 53 genes uniquely enriched in hepatic granulomas, characterised by a specialised program for immune recruitment (*Ccl11, Cxcl14, Cx3cl1, Ccr1, Cxcr4*) and structural stiffening of the matrix (*Col12a1, Tnc, Vcan, Spon1, Loxl1*). This unique signature of hepatic granulomas provides the molecular basis for the permanent and rigid sequestration of parasite eggs within the liver tissue.

**Table 1.**
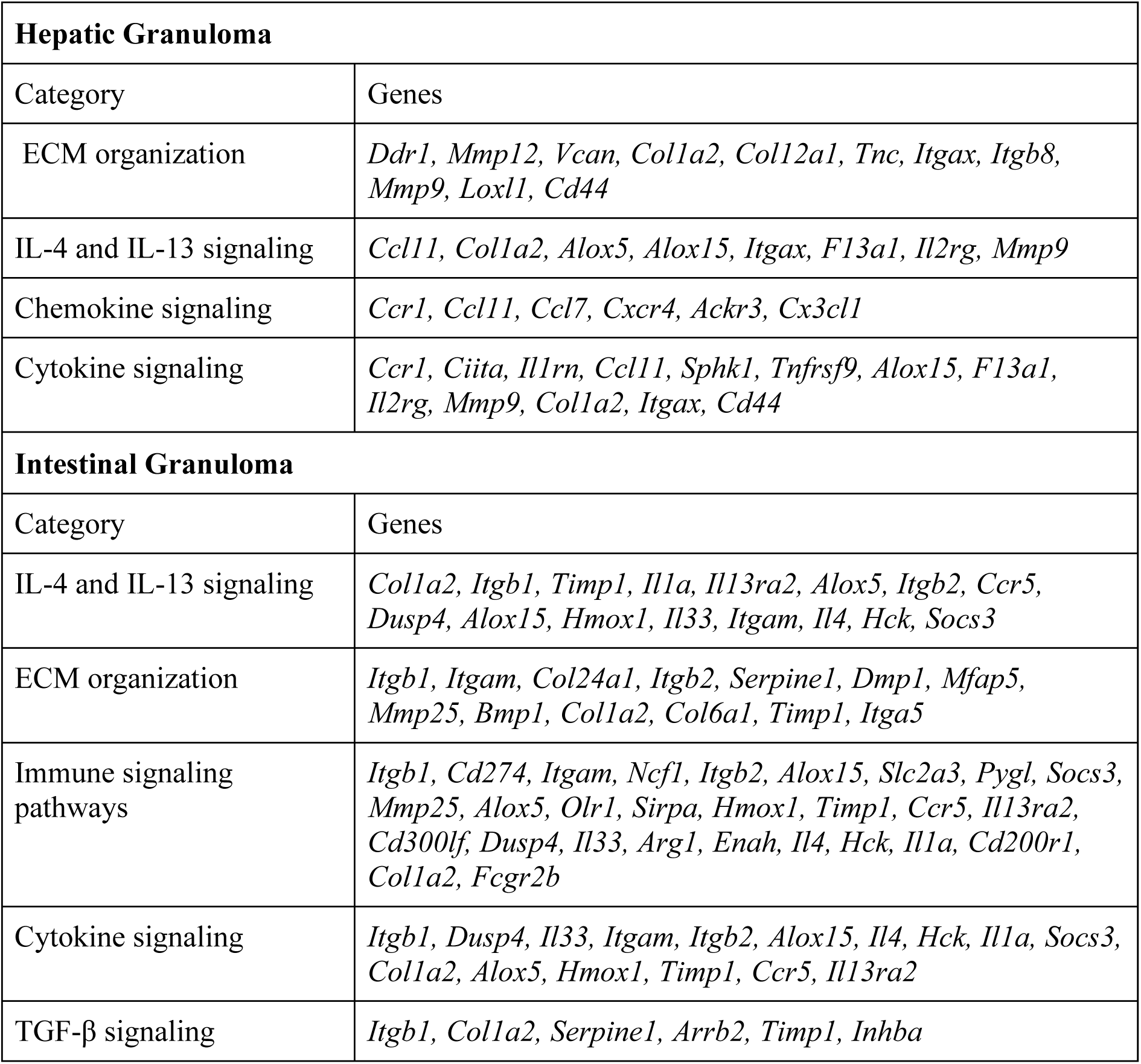
Functional categories associated with top upregulated DEGs in hepatic and intestinal granulomas.

#### 2.3.2 Intestinal granulomas exhibit a specialised transcriptomic and extracellular matrix program supporting egg translocation

The enrichment analysis of intestinal granulomas identified 78 top upregulated DEGs related to interleukin signaling, especially signaling via IL-4 and IL-13, ECM organization, immune signaling pathways, and cytokine signaling (**Table 1**, **Fig. 4C**, **Fig. 4D, Supplementary Fig. 2**). Notably, six of the top upregulated DEGs are members of the TGF-β family (**Table 1**, **Fig. 4D**), a pathway recognised as a master regulator of fibrogenesis across nearly all types of tissue injury and repair [41, 42]. By contrast, the most downregulated DEGs in intestinal granulomas comprised genes involved in maintaining structural architecture, mucosal barrier integrity, and the homeostasis of healthy tissue (*Cdh1*, *Dsp*, *Dsg2*, *Cldn7, Cdx2,* and *Hnf4a*; **Fig. 4C**). The downregulation of these genes suggests a significant disturbance of the intestinal epithelial barrier and a loss of tissue identity in regions affected by schistosome eggs. Furthermore, intestinal granulomas displayed a tissue-specific gene-expression signature comprising 70 uniquely enriched DEGs (**Fig. 4B**). This signature included M2 macrophage markers (*Arg1, Chil3, Cd200r1, Sirpa*); integrin-mediated leukocyte adhesion (*Itgam, Itgb2*); chemokine-mediate recruitment (*Ccl24, Ccr5, Ccr3*); structural genes (*Col6a1* and *Col24a1*); and ECM regulators (*Bmp1*, *Adamts6*, *Mfap5*, *Tgm2*, *Timp1*, and *Serpine1*; **Supplementary Table 2**). Consistent with this transcriptional signature, the corresponding granulomas exhibited a discontinuous, lower-density collagen matrix on Sirius Red staining at 8 wpi (**Fig. 2**).

#### 2.3.3 A core fibrogenic transcriptional programme is shared across hepatic and intestinal granulomas

Of all the significantly upregulated DEGs, 184 were shared between hepatic and intestinal granulomas **(Fig. 4B)**, pointing to a conserved transcriptional response to schistosome egg deposition across these distinct tissues. Reactome pathway enrichment of this shared set was dominated by ECM organization, cytokine and interleukin signalling, and leukocyte signal transduction. *Col1a2* and *Mmp9* recurred across all five top-ranked pathways, underscoring their central role in the shared fibrogenic and tissue-remodelling response; *Timp1,* the leukocyte integrins *Itgax, Itgb2,* and *Itgam,* the chemokines *Ccl2* and *Ccl11*, and the cytokine-signalling regulators *Socs3* and *Socs1* were represented in four. Among the top-ranked DEGs meeting strict significance thresholds in both tissues, eight genes were consistently upregulated, including the lipoxygenases *Alox15* and *Alox5*, *Col1a2*, and *Ccl7*. In contrast, only four genes were shared among the top downregulated DEGs in both tissues (*Abhd17c*, *Ceacam1*, *Idh1*, and *Ldha*). Given the substantial functional differences between hepatic and intestinal tissue, the limited overlap among downregulated genes is not unexpected. Rather than reflecting a coordinated suppression programme, these genes likely represent the convergent depression of baseline metabolic and homeostatic functions.

Taken together, the shared programme integrates ECM remodelling, leukocyte recruitment, and cytokine-signalling regulation as the conserved core of schistosome-induced fibrogenesis regardless of organ context. Notably, the recurrence of *Itgax*, *Itgb2* and *Itgam* within this shared transcriptional signature matches the integrin-centred LR communication identified at the intercellular signalling level (Section 2.2.3), where integrins emerged as dominant mediators of the fibrogenic microenvironment in both organs. The convergence of integrin signalling across both the transcriptional and intercellular communication analyses strengthens the case for this family of molecules as a pan-tissue feature of schistosome-induced fibrosis and a compelling candidate axis for therapeutic targeting.

### 2.4 Validation of spatial transcriptomics findings by qPCR and IHC

To validate the SC-ST findings, we performed complementary assays at both the transcript and protein levels using selected genes. qPCR validation focused on candidate DEGs selected among the most significantly upregulated or downregulated DEGs in the liver (*Lox, Vcan, F13a1*, *Ccl7, Ccl8*, *Cd163, Chil3, Mmp12*, *Trpc5*, and *Bmp10*). Gene expression differences between HC and INF groups were analysed by unpaired two-tailed t-tests, which confirmed significant upregulation of *Ccl7, Ccl8*, *Cd163*, *Chil3*, *F13a1*, *Lox, Mmp12*, and *Vcan* in infected samples, while *Bmp10* showed significantly higher expression in HC. Furthermore, *Trpc5* exhibited the same directional change as observed in spatial transcriptomic data (an upward trend in infected samples), however, the difference between HC and INF samples did not reach statistical significance in qPCR (**Supplementary Fig. 3**). Overall, qPCR analysis supported the spatial transcriptomics results, with consistent directionality of significant gene expression changes.

Furthermore, we validated the transcriptomic predictions at the protein level by immunohistochemistry, which was performed on liver and intestine tissues for two of the top upregulated DEGs identified in granulomatous regions of each tissue. In the liver, *Col1a2* showed strong staining throughout the granulomas, while *Mmp9* displayed a more diffuse pattern. These protein expression patterns were consistent with the spatial transcriptomic maps (**Fig. 5**). In the intestine, *Olr1* and *Chil3* were abundantly expressed across the granuloma. While protein abundance exceeded transcript levels, their overall localisation was generally consistent with the SC-ST results, with minor deviations expected due to differences between mRNA and protein expression.

**Figure 5.**
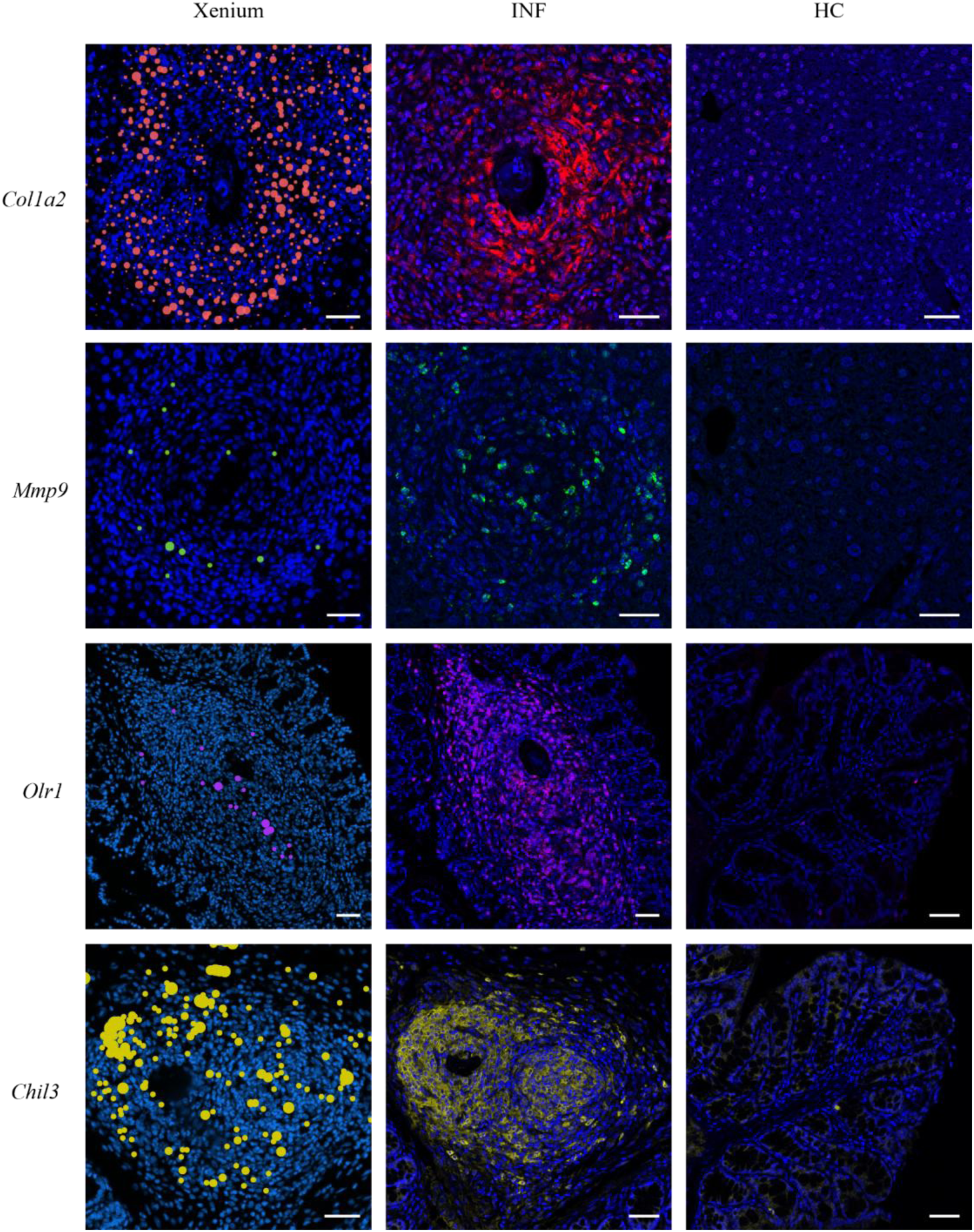
Independent protein-level validation of DEGs identified by single-cell spatial transcriptomics in tissues from healthy and *Schistosoma mansoni*–infected mice via immunohistochemistry. Immunohistochemical staining, visualised by confocal microscopy, confirms the protein expression patterns observed in single-cell spatial transcriptomics. Specifically, infected liver sections were stained for *Col1a2* and *Mmp9*, while infected intestinal sections were stained for *Olr1* and *Chil3*. Scale bars represent 50 µm.

## 3. Discussion

The schistosome-induced granuloma is a complex structure that initially forms to protect the host tissues from toxic egg secretions. While this response is protective in the acute phase, the persistence of granulomas drives chronic inflammation and irreversible tissue remodelling, which represents the main immunopathology of hepatic schistosomiasis [13, 15, 29, 43, 44]. Previous RNA sequencing studies have resolved cellular heterogeneity mainly within hepatic granulomas [29–32]; however, dissociation-based approaches inherently sacrifice spatial context, limiting the interpretation of gene expression within intact tissue architecture. This limitation is particularly relevant in schistosome-induced fibrosis, where pathogenic signalling is governed by the spatial organisation and interactions between egg secretion and host immune cells within the tissue microenvironment. SC-ST uniquely overcome this limitation by both preserving spatial context and enabling the interrogation of cellular co-localisation, niche architecture, and tissue-specific organisation alongside gene expression. To date, spatial transcriptomics has only explored parasite maturation in *S. japonicum* [45], leaving host pathology unexamined. Our study fills this gap by providing the first spatially resolved single-cell map of host fibrotic responses to schistosome eggs trapped in two distinct tissue environments: liver and intestine. By comparing hepatic and intestinal granulomas, we revealed conserved and organ-adapted programs that reflect the divergent demands of permanent hepatic containment versus flexible intestinal translocation, providing a framework to understand how organ-specific fibrotic niches may generate distinct therapeutic vulnerabilities.

SC-ST provided the resolution necessary to reveal schistosome-induced fibrosis as a highly organised yet tissue-specific process. While granuloma formation in both organs is fundamentally initiated to sequester immunogenic parasite eggs and shield the host from toxic factors [13, 15, 21, 46], the resulting architectural programs appear to reflect the distinct physiological demands of each tissue: terminal egg sequestration in the liver versus egg transit through the intestinal wall for excretion [13, 15, 46, 47]. In the intestine, eggs lodged within the vasculature first engage host endothelial cells and initiate extravasation, a process recently shown to involve host-cell encapsulation, contractile force generation, and endothelial junction disruption [48]. Here, we further show that intestinal granulomas are associated with a distinct immune-stromal and ECM-remodelling program by 8 wpi. Our data show that the intestinal environment is defined by a coordinated metabolic and architectural axis designed to build a flexible transit environment for egg excretion rather than a rigid fibrotic scar. The master molecular engine driving this niche is the IL-4/IL-13 signaling pathway, which polarises macrophages towards a metabolic M2 phenotype characterised by high expression of *Arg1* and *Chil3* [47, 49]. Here, *Arg1* converts L-arginine into proline building blocks required by activated fibroblasts for rapid collagen synthesis [50–53]. Our integrated spatial analysis identified a dominant macrophage-activated fibroblast axis as the primary engine of extracellular matrix construction. This cellular interaction is anchored between macrophage-expressed *Cd44*, a multi-functional adhesion receptor and known integrator of fibrosis [54], and the fibroblast-secreted *Col1a2* scaffold. This architecture is further specialised by the upregulation of structural genes *Col6a1* and *Col24a1*, which are associated with the formation of more flexible collagen networks. Such an organisation may better accommodate the mechanical stress of peristaltic movement and is consistent with the discontinuous and less dense collagen organisation observed in intestinal histology [55]. Finally, the localised enrichment of TGF-β family members suggests that this master fibrogenic pathway [41, 42] could be differentially regulated in the gut, potentially creating a matric environment permissive for egg transit. Future studies incorporating direct measurements of collagen turnover, matrix mechanics, or egg movement will be important to more directly test these possibilities.

Hepatic fibrosis in schistosomiasis represents a highly organised host-adaptive strategy designed to permanently sequester immunogenic eggs and prevent the liver parenchyma from potent toxins such as *Omega-1* and *IPSE/alpha-1* [13, 15]. Consistent with this concept, our spatial mapping of hepatic granulomas at 8 wpi identified the molecular machinery required to execute this sequestration strategy. We found a profound enrichment of IL-4 and IL-13 signalling, which acts as the master molecular switch to redirect the host from destructive inflammation towards a Th2-mediated tissue-remodelling program [50, 56]. This cytokine environment orchestrates a dense immune swarm of monocyte-derived macrophages and eosinophils, providing the physical volume necessary for the initial egg encapsulation [50]. Simultaneously, these signals drive the activation of HSCs from a vitamin A-storing quiescent state into collagen-producing cells [13, 57]. Our CCI analysis identified an aHSC-monocyte-derived macrophage axis coordinated by TGF-β signalling. While *Tgfb1* is the canonical driver of fibrogenesis [58, 59], we identified *Tgfb3* as a central interaction hub at 8 wpi. *Tgfb3* uses the same canonical SMAD-dependent signalling pathway as *Tgfb1* [60], yet is associated with more controlled ECM deposition and establishment of defined granuloma boundaries [61]. Thus, we propose that *Tgfb3* acts as a critical architect of spatial organisation, establishing defined granuloma boundaries seen in hepatic granulomas while limiting disorganised scarring. Ultimately, the interactions centred on *Tgfb3* may form a self-reinforcing feed-forward loop. In this model, macrophage-derived ligands (specifically members of the TGF-β family) provide the primary stimulus for HSC transdifferentiation and matrix deposition, while the resulting fibrotic microenvironment provides the critical cues necessary to sustain macrophages in an M2 activation state [62, 63]. This self-sustaining loop ensures the permanent sequestration of parasite antigens while simultaneously maintaining a persistent, non-resolving fibrosis.

Despite marked differences in fibrotic architecture between hepatic and intestinal granulomas, our comparative analyses indicate that granuloma formation in both tissues is driven by shared molecular programs. We identified 184 overlapping genes that define a shared transcriptional blueprint for the host response to *S. mansoni* eggs. Among these, *Col1a2* and *Mmp9* repeatedly emerged as central hubs across major enriched pathways, including cytokine signalling and ECM organisation. *Col1a2* likely provides the structural scaffold required for egg sequestration, whereas *Mmp9* supports active matrix remodelling and inflammatory cell recruitment [13, 64]. This core program is further shaped by *Socs1* and *Socs3*, which constrain excessive Th2 activation, and by chemotactic factors such as *Ccl2* and *Ccl11* that coordinate immune-cell influx [65, 66]. Importantly, all LR interactions shared between hepatic and intestinal granulomas at 8 wpi centred on integrin receptors, identifying integrin signalling as a common communication hub within the granuloma niche. Although integrins have not traditionally been viewed as primary drivers of schistosome-associated fibrosis in the same way as TGF-β or IL-13 [56, 67], they are increasingly recognised as key regulators of matrix sensing, HSC activation, and profibrotic signalling [68, 69]. Among these, *Itgb1* emerged as the dominant integrin node in both healthy and infected tissues. In healthy tissue, β1-integrin interactions likely support normal matrix adhesion and tissue architecture. During infection, however, the *Itgb1-*centred interactome shifted towards ECM ligands associated with remodelling and fibrosis, including *Col1a2*, *Fn1*, *Tgm2*, *Fgg*, *Fga*, *Col4a1*, *Fbn1*, *Lamb1*, *Col18a1*, and *Cxcl12*. The strongest shared interaction in granulomas was *Col1a2*-*Itgb1*, consistent with the known role of β1-containing integrins in binding newly deposited collagen and reinforcing stromal-cell attachment to the fibrotic matrix [69]. Rather than simply indicating the presence of scar tissue, the *Col1a*-*Itgb1* interaction may reflect a self-reinforcing signalling circuit, where egg-stimulated ECM accumulation provides the ligands required to engage β1-integrins on aHSCs and fibroblasts, sustaining their profibrotic state and continuing to drive tissue remodelling and organ destruction even after the primary immunogenic stimulus (egg) has been sequestered. The additional interactions between *Itgb1* and *Fn1* (fibronectin) or *Tgm2* (transglutaminase) support this idea because fibronectin- and transglutaminase-based interactions are known to strengthen integrin signalling and drive myofibroblast differentiation [41]. The fact that similar *Itgb1*-centred interactions were present in both liver and intestine suggests that schistosome eggs trigger a common integrin-dependent stromal response across tissues.

Importantly, this study provides a comprehensive molecular framework for the rational development of anti-fibrotic interventions in schistosomiasis. Among the shared LR pairs, we identify *Itgb1* as a leading pan-tissue candidate, as inhibition of this central hub has the potential to disrupt the self-reinforcing profibrotic feedback loop through which activated myofibroblasts sense and perpetuate ECM deposition across multiple organs. Notably, *Itgb1* has recently emerged as a target in pulmonary fibrosis [70] and diaphragm fibrosis [71], supporting the feasibility of a broad-spectrum anti-fibrotic strategy. While pan-tissue targets represent an attractive strategy to disrupt the core fibrotic machinery shared across multiple organs, our data demonstrate that schistosome-induced fibrosis is not a uniform process. Instead, hepatic and intestinal granulomas adopt highly specialised transcriptional and signalling programs that reflect their distinct functional and pathological roles. Therefore, pan-tissue intervention alone may be insufficient to fully resolve organ-specific pathology. Accordingly, our data support tissue-targeted therapeutic strategies that build on this shared fibrotic backbone. In hepatic granulomas, targeting *Itgb8* or *Tgfbr1* in the liver may attenuate latent TGF-β activation and limit the formation of the rigid, self-reinforcing ring that underpins permanent egg sequestration. In contrast, blockade of *Cd44* and *Itga5* in intestinal tissue may disrupt macrophage anchorage to the fibronectin matrix, destabilising the flexible microfibrillar architecture that facilitates egg migration toward the lumen. These strategies were previously shown to suppress profibrotic signalling in fibrosis of lungs, airways, kidneys, and spinal cord [72–75], underscoring their therapeutic relevance beyond schistosomiasis. The identified targets (*Itgb1*, *Itgb8*, *Tgfbr1*, *Cd44*, and *Itga5*) represent prioritised candidates derived from spatial network analysis and require further experimental validation before therapeutic application. In parallel with the identification of novel targets, we cross-referenced the transcriptomic signatures of liver granuloma against genes modulated by miRNA-25-3p (miR-25), an anti-fibrotic candidate targeting Notch1 signalling and TGF-β-induced collagen expression in HSCs that is currently under preclinical development [76]. The analysis identified 20 genes regulated in opposing directions, involved in Notch and Wnt pathways, both of which interact with TGF-β. These findings provide a hypothesis-generating cross-validation of pathway convergence and suggest that miR-25-associated regulatory networks may intersect with schistosome-driven fibrotic signalling. Ultimately, current schistosomiasis treatment relies exclusively on PZQ, which eliminates adult worms but does not target the eggs and tissue fibrosis [25–27, 77]. Viable eggs continue to release immunogenic excreted-secreted products that sustain granulomatous inflammation and drive progressive fibrosis independently of worm burden. Fully resolving disease pathology will therefore require a combination model that pairs PZQ with agents targeting egg viability and downstream anti-fibrotic therapies to fully disrupt the pathological cascade from infection to fibrosis.

While this SC-ST study reveals shared and tissue-specific features of schistosome-induced fibrosis, it is important to acknowledge several limitations related to experimental scope and interpretation. First, despite eosinophils being a hallmark of the host’s Th2-mediated immune response [78, 79] and dominant cell type of schistosome-induced granulomas [79, 80], they were detected at low levels in our intestinal SC-ST dataset. This is likely attributable to their notoriously low cytoplasmic mRNA levels [81], which can fall below the sensitivity threshold of probe-based spatial platforms. Next, our intestinal analysis identified a cluster of cells with a hybrid transcriptional profile and weak enrichment of canonical markers, preventing confident annotation. Considering the biological landscape of intestinal schistosomiasis, where the entrapment of eggs in the submucosa triggers a state of intense tissue remodelling and chronic inflammatory states [46], these ambiguous cells may represent undifferentiated progenitors in a transitional state that lack the mature marker expression. Finally, our study assesses fibrosis at a single time point and cannot capture the prognosis of fibrogenesis. This stage corresponds to the peak of the egg-driven granulomatous response [13], when hepatic and intestinal lesions display their most distinct structural features, and profibrotic signalling is maximal. Focusing on this time point, therefore, enabled detailed spatial resolution of the developing fibrotic niche and the tissue-specific cellular interactions underlying chronic remodelling. We plan to extend this work to cover two time points, early inflammatory and late fibrotic stages, to track how the spatially resolved cellular interactions identified here evolve throughout granuloma progression and fibrosis development.

To move towards clinical translation, future research should focus on functionally disrupting identified signalling nodes via CRISPR/Cas9 or RNAi to distinguish essential drivers of pathology from secondary inflammatory effects. Furthermore, incorporating a schistosome-specific probe panel will formally delineate the host-parasite transcriptional interface *in situ*, resolving how egg-secreted proteins stimulate the host immune response within the granuloma core and drive myofibroblast transdifferentiation. Ultimately, integrating host and parasite spatial profiles within a single analytical framework will provide a comprehensive picture of the schistosome fibrotic niche, one that simultaneously shows the parasite-derived drivers of pathology and the host effector populations that sustain it.

By pioneering SC-ST in schistosome-induced fibrosis, this study resolves the cellular and molecular architecture of hepatic and intestinal fibrotic niches at unprecedented resolution. Our high-resolution spatial mapping reveals fundamentally distinct architectural programs underpinning rigid egg sequestration in the liver versus permissive egg transit in the intestine. The identification of organ-specific LR hubs, including TGF-*β*-dominant network in hepatic granulomas and the integrin-centered axis in the intestine, provides a foundational framework for discovering novel therapeutic targets to reverse established tissue scarring, addressing a critical clinical gap where PZQ-driven parasite clearance fails to resolve existing damage.

## 4. Methods

### Ethics statement

Ethics approval for animal use was granted by the Animal and Human Ethics Committee of QIMR Berghofer Medical Research Institute (QIMRB; approval number P3705). All experiments were conducted in accordance with the guidelines of the National Health and Medical Research Council of Australia outlined in the Australian Code of Practice for the Care and Use of Animals for Scientific Purposes, 7th edition, 2004 (www.nhmrc.gov.au). The experimental procedures related to live *S. mansoni* parasites were conducted in quarantine-approved facilities of QIMRB.

### Experimental animals, infection, and tissue processing

Female BALB/c mice were housed at the Queensland Medical Research Institute Berghofer in compliance with international and national standards. To establish a mild burden *S. mansoni* infection, ensuring survival of mice for 8 weeks post-infection in all experimental groups, anesthetised mice were exposed to 25 cercariae by abdominal skin penetration [46]. Mouse tissue collection was performed at 8 weeks p.i. with *S. mansoni*, a time point strategically chosen to coincide with the peak of the egg-driven granulomatous response [43]. This timing ensures the robust presence of *S. mansoni* eggs in host tissues, as egg production is well-established by this stage, typically initiating around 5-6 weeks p.i. [25, 27]. All mice were injected intraperitoneally with 100 µL of 5000 IU/5ml heparin sodium (porcine mucous; Pfizer, USA) and allowed to circulate for approximately 3 minutes before proceeding with euthanasia by exposure to CO₂ gas. Portal perfusion was performed to obtain and count the adult worms. Mouse livers and intestines were collected for pathological tissue staining and scST. Individual livers were split into two. The first section was fixed in 10% neutral buffered formalin overnight at room temperature (RT) in preparation for paraffin embedding. The second section was cut into fine pieces, treated with TRIzol reagent (Invitrogen, Thermo Fisher Scientific, USA), and stored at −80°C. The intestines were cut along their entire length and rinsed thoroughly with phosphate-buffered saline (PBS, pH 7.4) to remove contents. The colon and rectum, which harbour the highest egg density [9, 28], were dissected from the intestine and immediately fixed in 10% neutral-buffered formalin for histological processing and scST.

### Pre-Xenium histology

Mouse liver and intestine samples, fixed overnight in 10% neutral buffered formalin, were submitted to the Histology Facility (QIMRB) for processing. At the facility, samples were dehydrated, cleared, and embedded in paraffin according to standard protocols [82]. Afterwards, paraffin-embedded tissues were cut, mounted on glass slides, and stained with H&E for general tissue architecture and identification of *S. mansoni* eggs and egg-induced granuloma and with Picrosirius Red for collagen visualisation (tissue fibrosis). The stained slides were scanned using the Aperio AT turbo brightfield microscope (Leica Biosystems, Nussloch, Germany) at 20x optical magnification. Captured images were processed using Aperio ImageScope software (v.12.4.0.5043). After initial identification of the parasite eggs in the tissues, the existing paraffin-embedded tissue blocks were gently warmed to allow removal from the paraffin matrix. Selected regions containing eggs were excised from the original blocks, and the tissues were re-embedded in fresh paraffin to generate separate blocks for liver and intestine specimens in preparation for Xenium.

### Xenium Data Generation

The formalin-fixed paraffin-embedded (FFPE) blocks containing liver and intestinal tissues, respectively, were processed following the procedure outlined in the Tissue Preparation Handbook (10x Genomics, CG000578, Rev F). In brief, the FFPE blocks were hydrated in ice-cold RNase-free water for 15 - 30 min. After trimming, the FFPE blocks were sectioned at 5 μm, and tissue sections were placed onto equilibrated Xenium slides (10x Genomics, PN# 3000941). After collecting all tissue sections onto the Xenium slides, the tissues were baked in a 42°C oven for 3 h followed by overnight drying in a Falcon tube with desiccant (Merck, PN# 1.01972). The tissue deparaffinization was performed by baking the slides on the Xenium Thermocycler Adapter v2 (10x Genomics, PN# 1000739) on a PCR machine (Bio-Rad, PN# 1851197) at 60°C for 30 min, followed by two Xylene washes, serial ethanol washes (2 × 100%, 2 x 96%, 1 x 70%), and lastly the nuclease-free water wash. The Xenium slides were assembled into Xenium Cassette (10x Genomics, PN# 1000726) before proceeding to the Decrosslinking. After decrosslinking, Xenium slides were processed according to the Xenium Prime In Situ Gene Expression User Guide (10x Genomics, CG000760, Rev B). The Xenium slides underwent Priming Hybridisation with Xenium 5K Mouse PTP Priming Oligos (10x Genomics, PN# 2001224), followed by RNase treatment and Polishing. Xenium 5K Mouse PTP Panel (10x Genomics, PN# 2001225) was hybridised with sample overnight (18 – 19 h) at 50°C. After probe hybridisation, samples underwent a series washes, probe ligation, amplification, cell segmentation staining, and autofluorescence quenching and nuclei staining. Slides were then loaded into Xenium Analyser (10x Genomics, PN# 1000529) with the appropriate buffers and decoding module plates, as outlined in the Xenium Analyser User Guide (10x Genomics, CG000584, Rev K). After Xenium imaging completion, the slides were stored in PBS-T for up to three days at 4°C before the H&E staining and imaging using Leica Aperio AT turbo slide scanner (Leica, Germany).

### Selection of the regions of interest

ROIs were manually defined based on Xenium morphology and DAPI images using the freehand selection tool in Xenium Explorer 4 (10x Genomics) to capture different microenvironments in healthy and parasite-infected tissues. In the infected tissues, each ROIs contained parasite egg(s), a surrounding inflammatory area, and a 3-5 cell-thick layer of morphologically intact tissue directly adjacent to the granuloma, allowing us to include a gradient from inflamed to healthy regions. For comparison, ROIs were also selected from uninfected (healthy) mouse tissues. These were structurally comparable to those selected in the infected tissues but lacked evidence of inflammation or parasite eggs. Ten ROIs were defined for each liver sample across three biological replicates. For intestinal samples, where fewer parasite eggs were detected, all available egg-associated granulomas were included to ensure adequate representation of infected regions. Specifically, we selected three ROIs from the *S. mansoni-*infected mice and nine ROIs from the healthy mice. Coordinates were exported from Xenium Explorer and used to extract transcript-level data for each ROI.

### Preprocessing and Cell Typing

Xenium spatial transcriptomics data were preprocessed using *Scanpy*. During quality control (QC), low-quality genes were filtered by retaining genes expressed in at least 3 cells (*min.cells* = 3), and cells with fewer than 10 total counts (*min.counts* = 10) were removed. Data were subsequently log-normalised, and batch effects across samples were corrected using the Harmony algorithm. Principal component analysis (PCA) was performed, and the top 50 principal components were used for downstream analysis. Cell clustering was performed using the Leiden algorithm with a resolution parameter of 0.3 (res=0.3). Clusters were then annotated based on canonical marker genes using manual curation. The main cell types identified and the main markers used for their identification are in Table 2.

**Table 2.**
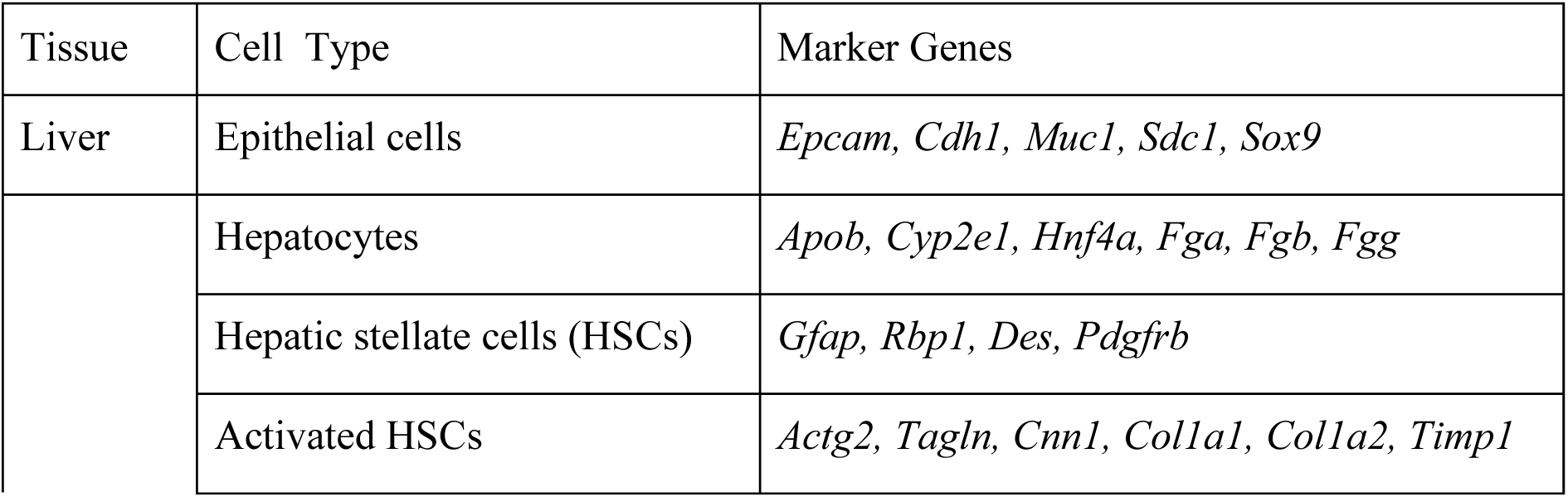
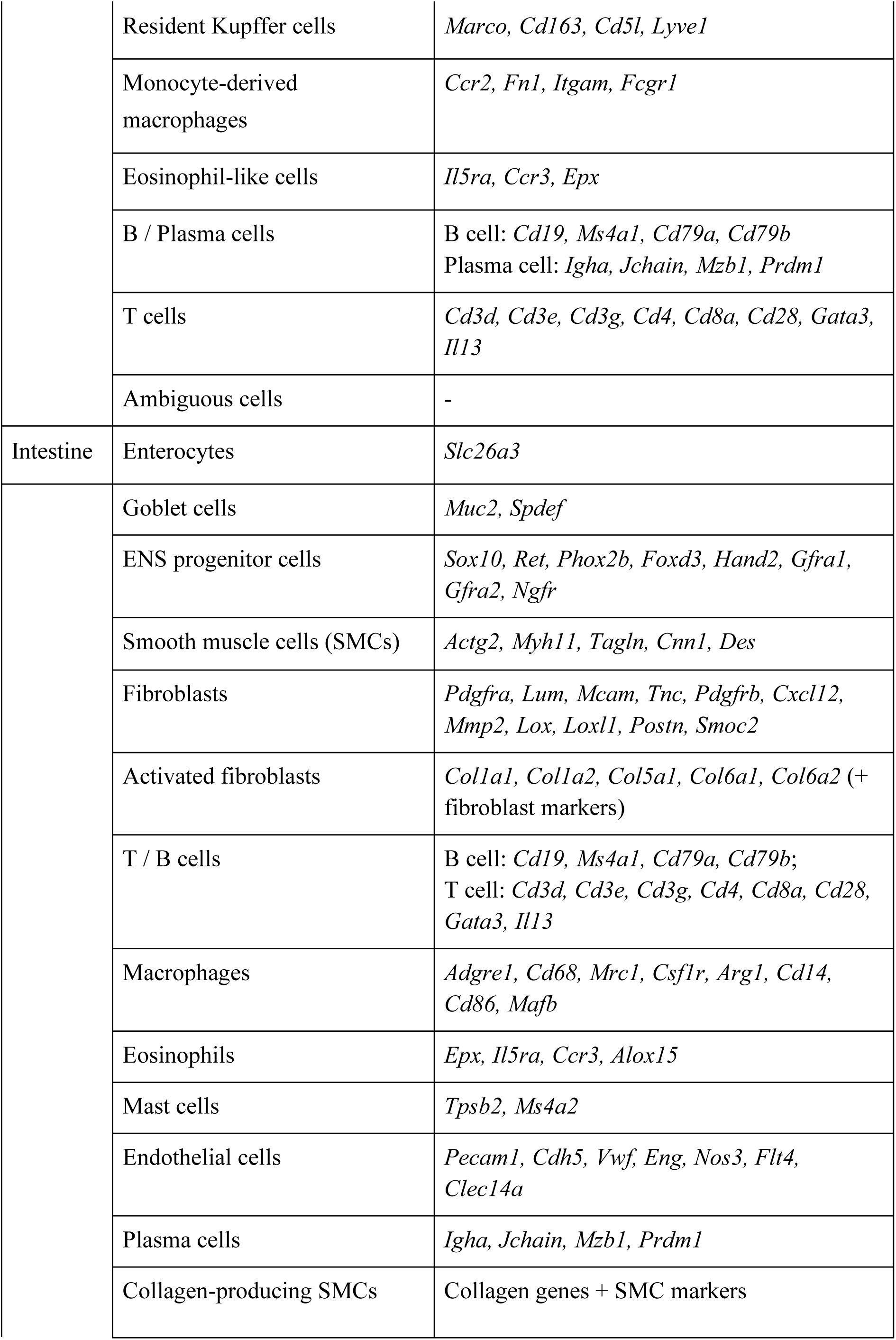
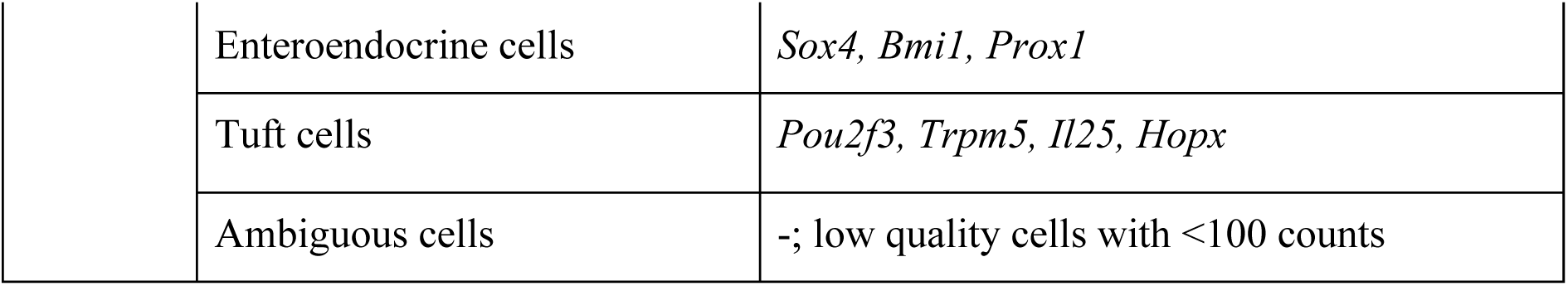
Marker genes used for cell type annotation.

### Transcriptomics and pathways

Differential gene expression analysis between healthy and infected tissues was performed using a pseudo-bulk strategy. ROIs were selected from each tissue type, including thirty ROIs each from healthy and infected liver samples, and eleven healthy and three infected ROIs from intestinal samples. Single-cell expression data were aggregated into pseudo-bulk profiles using the *Seurat2PB* function, grouping cells by ROI and condition to generate sample-level count matrices. Library sizes were normalised using the trimmed mean of M-values (TMM) method implemented in edgeR. To account for inter-individual variability, a paired experimental design was used, incorporating donor identity as a blocking factor *(∼ donor + condition)* [83]. Dispersion estimates were computed using robust methods, and differential expression testing was performed using quasi-likelihood F-tests. Resulting p-values were adjusted for multiple testing using the Benjamini–Hochberg procedure, and genes with a false discovery rate (FDR) < 0.05 were considered statistically significant.

For the selection of top upregulated genes in infected tissue (logFC) > 0), two complementary ranking strategies were applied to the resulting DE table (containing columns: logFC, logCPM, F, PValue, FDR): (i) Statistical ranking: Genes were first sorted by FDR in ascending order, then by logFC in descending order within the same FDR value. (ii) Combined biological-statistical score: A score was calculated as logFC × (−log10(FDR)) to favour genes with both large effect size and high statistical significance. The top 100 genes from each ranking method were extracted. The intersection of these two lists was used as the final high-confidence set of upregulated genes. This approach ensures that selected genes rank highly under both statistical rigour and biological effect size. These genes were used for volcano plot labelling and downstream pathway analysis in EnrichR using the Reactome Pathways 2024 library [84].

For visualisation of top DE genes between the two conditions for selected ROIs, log-transformed counts per million (logCPM) values were used to generate heatmaps.

### Cell-cell interactions

For the further selected regions of interest, including 5 regions each from healthy and infected liver samples and 11 healthy and 3 infected regions from the intestinal samples, cell-cell interactions were inferred using stLearn. To enable cross-sample and cross-tissue comparisons, CCI results were further integrated using a multi-modal cell–cell interaction (MMCCI; [85]) framework. For each tissue, the healthy and the infected granuloma ROIs were grouped as HC and INF, respectively. The network plots of the CCI networks were visualised for HC, INF, and HC vs. INF. The top LR pairs whose interactions were higher in the healthy and the infected groups, and also those conserved in the infected samples across the two tissues, were also identified. The spatial plots of the LRs were also visualised.

### Immunohistochemistry

Formalin-fixed, paraffin-embedded tissue sections that were previously used for the preparation of Xenium slides (containing livers and intestines from healthy control and *S. mansoni*-infected mice) were dewaxed and rehydrated prior to staining. Multiplex immunofluorescence was performed using tyramide signal amplification (Opal fluorophores) with antibody-specific optimisation as detailed below. Although antibodies were stained as defined multiplex panels, only a single marker from each panel was visualised and analysed in the present study (Liver: GFAP-MMP9, LOX1-COL1A2; Intestine: TGFB1-COL1A2, CHIL3-MMP9). Nuclei were counterstained with DAPI.

MMP9 antibody (EP1254; Abcam, ab76003) was applied following antigen retrieval in DAKO pH 6 retrieval buffer. Sections were blocked with Sniper containing 2% BSA and incubated with primary antibody diluted in DaVinci Green antibody diluent. Detection was performed using Biocare Medical MACH 1 Universal Polymer, followed by tyramide signal amplification with Opal 520.

LOX1 (Olr1) antibody (Abcam, ab317689) was applied following heat-induced antigen retrieval in DAKO pH 6 buffer. Sections were blocked with Sniper containing 2% BSA and incubated with primary antibody diluted in Sniper/BSA. Detection was performed using MACH 1 Universal Polymer, followed by tyramide signal amplification with Opal 520.

COL1A2 antibody (Invitrogen, PA5-96426) was applied following antigen retrieval in DAKO pH 6 buffer. Blocking was performed using Sniper supplemented with 2% BSA and 20% goat serum. The primary antibody was diluted in DaVinci Green. Detection was performed using PerkinElmer anti-mouse HRP followed by tyramide signal amplification with Opal 570. Slides were counterstained with DAPI.

CHIL3 antibody (Abcam, ab230610) was applied following heat-induced antigen retrieval in DAKO pH 6 buffer. Sections were blocked with Sniper containing 2% BSA and incubated with primary antibody diluted in Sniper/BSA. Detection was performed using MACH 1 Universal Polymer, followed by tyramide signal amplification with Opal 570.

Confocal images were acquired using a Leica Stellaris 5 (Leica Microsystems, Germany) controlled by LAS X (version 4.8.2). Imaging was performed using a 20× HC PL APO CS2 objective. Images were acquired at a pixel size of 0.45 µm in xyz scan mode with a scan speed of 400 Hz and a pixel dwell time of 1.41 µs. Liver datasets were acquired at 8-bit, and intestine datasets at 16-bit depth. DAPI, FITC, and TRITC channels were acquired using 405 nm, 493 nm, and 553 nm laser lines, respectively, and emission was detected over spectral ranges of 425–501 nm, 501–555 nm, and 558–691 nm. Images were acquired at 1.28× zoom.

Image processing was performed using ImageJ (v1.54r). Samples were imaged using four distinct staining panels, and images within each panel were processed identically. Display settings were optimised for each panel and then applied uniformly to all images within that panel. Brightness and contrast were adjusted linearly by modifying display ranges for visualisation purposes, and channels were merged to generate composite images.

### RNA extraction from mouse liver samples and cDNA synthesis

All steps were performed under RNase-free conditions to preserve RNA integrity. Total RNA was isolated from mouse liver samples using TRIzol extraction method followed by Qiagen affinity column purification as outlined by [29] with some modification. Thawed liver samples were mechanically homogenised in RNAse-free microcentrifuge tubes on ice in 1-minute intervals using a hand-held tissue homogeniser (KIMBLE® PELLET PESTLE® Cordless Motor, DWK Life Sciences, Vineland, New Jersey, USA) with RNase-free pestles (Kimble™ Kontes™ Pellet Pestle™, DWK Life Sciences, Vineland, New Jersey, USA). The homogenised mixture was kept on ice for 2 h. Chloroform (Chem-Supply, Gillman, SA, Australia; 0.1 mL per 0.5 mL TRIzol) was added to the mixture and shaken vigorously for 1 min to induce phase separation. Samples were incubated at RT for 2 min, then centrifuged for 15 min at 4 °C at 12,000 × g. The aqueous phase containing RNA was carefully recovered, mixed with an equal volume of 70% ethanol (Sigma-Aldrich, Darmstadt, Germany), transferred to a RNeasy MinElute spin column (RNeasy® Micro Kit, Qiagen, Hilden, Germany) and centrifuged for 5 min at RT at 5,000 × g. The flow-through was vortexed and spun again at the same settings to maximise RNA capture and ensure complete binding of RNA to the column matrix. The column was washed sequentially with 700 µl of Buffer RW1 and centrifuged for 15 s at RT at 8,000 × g. This was followed by two washes with 500 µL of Buffer RPE, with centrifugation at 8,000 × g at RT after each wash (15 s for the first spin, 2 min for the second). A final 5-min centrifugation with no buffer added was performed to ensure complete drying of the column prior to RNA elution. The total RNA was eluted from the column by applying 20 µl of RNase-free water and centrifugation for 1 min and 15 s at RT at 8,000 × g. To facilitate RNA precipitation, we added 2 µl of 5M NaCl (Chem-Supply, Gillman, SA, Australia) and 50 µl of 100% EtOH (Sigma-Aldrich, Darmstadt, Germany) to the RNA, vortexed, and incubated the sample at −20°C overnight. RNA was recovered by centrifugation at 12,000 × g for 15 min at 4°C. The resulting pellet was washed twice with 75% ethanol under the same centrifugation conditions. After washing, the pellet was air-dried for 10 min and dissolved in 50 µl of RNase- free water. The purity of the isolated RNA (A260/A280) was assessed using a NanoDrop OneC spectrophotometer (Thermo Fisher Scientific, Waltham, MA, USA) and ranged from 2.03 to 2.08. Purified RNA was reverse-transcribed into complementary DNA (cDNA) using the QuantiTect Reverse Transcription Kit (Qiagen, Hilden, Germany) according to the manufacturer’s instructions.

### Quantitative polymerase chain reaction

qPCR was performed using GoTaq® Probe qPCR Master Mix kit (Promega, USA) on a Mic qPCR (Cycler Bio Molecular Systems, Upper Coomera, QLD, Australia) according to the manufacturer’s instructions. Gene IDs were retrieved from the Xenium Mouse 5k Gene Expression Panel (probe sequences, Rev B). For each gene, the corresponding Ensembl ID (e.g., ENSMUSG00000029661 for *Col1a2*) was used to identify the transcript, and primers were subsequently designed using the PubMed Primer Design Tool. Sequences of all primers used for qPCR are listed in **Supplementary Table 5.** Mouse glyceraldehyde-3-phosphate dehydrogenase (*GapDH*) primers were used to amplify the housekeeping reference *GapDH* gene (NCBI Reference Sequence: NM_008084.4) [30]. The qPCR reactions contained 5 µl GoTaq® Probe qPCR Master Mix, 150 ng cDNA, and 0.8 µM of forward and reverse primers (each). Thermal cycling included an initial denaturation at 95°C for 2 min, 40 cycles of denaturation at 95°C for 30 s, annealing at 52-60°C (primer-dependent) for 30 s, and extension at 72°C for 30 s. Optimal annealing temperatures for each primer pair were determined empirically using standard PCR. Ct values were obtained with Mic qPCR software (Bio Molecular Systems), and relative expression levels of each gene were calculated using the 2^−ΔΔCt^ method, relative to the housekeeping GapDH gene.

### Data analysis

The Xenium samples were preprocessed using scanpy. During the QC step, low-quality cells and genes were filtered out using the following criteria: min.cells=3 and min.counts=10. After log-normalisation, sample batch correction was performed using RunHarmony(), followed by PCA using the top 50 principal components. Leiden clustering was performed and clusters were generated using a resolution of 0.3 (res=0.3). Further, the clusters were annotated by using manual canonical marker-based annotations.

### Statistical analysis

Statistical analyses were performed in Python (v3.11.6). For comparisons between two groups, unpaired two-tailed *t*-tests were used unless otherwise specified. For differential expression analyses, statistical testing was performed using a quasi-likelihood framework, and p-values were adjusted for multiple testing using the Benjamini–Hochberg method to control the FDR. For cell–cell interaction analyses, ligand–receptor interaction scores were compared across conditions, and significance was assessed using permutation-based or model-derived p-values as implemented in the respective tools. Unless otherwise stated, statistical significance was defined as *p* < 0.05. Asterisks indicate significance levels as follows: *, p < 0.05; **, p < 0.01; ***, p < 0.001.

## Acknowledgements

We are grateful to Mary Duke (QIMRB) for performing mouse perfusions and maintaining the parasite life cycle, and Dr William C. Dougall, whose insights and support in the early phases helped shape the foundation of this project. We thank the NIAID Schistosomiasis Resource Center at the Biomedical Research Institute (Rockville, MD, USA) for providing *Biomphalaria glabrata* (NMRI strain) snails infected with *S. mansoni* for this work. We also thank the staff of the QIMRB Histology and Microscopy Facilities for their technical assistance and imaging support. An AI-based language model was used to assist with grammar and language editing. This work was funded by the QIMRB 2025 Seed Funding Grant (428140 - Toth) and the National Health and Medical Research Council (NHMRC) Ideas Grant (APP2028650 - You).

## Author contributions

V.T., H.Y., and Q.N. designed the study.

V.T. performed and analysed the pre-Xenium experiments.

Z.X. conducted the Xenium spatial transcriptomic experiments.

P.P. analysed the spatial transcriptomic data.

V.T. performed the validation experiments and acquired and analysed the microscopy data.

G.D.A. contributed to histological evaluation of liver tissue and provided expert input on the miRNA-25 findings.

V.T, H.Y, P.P, Z.X., C.L, and Q.N. interpreted the data.

V.T. and P.P. drafted the manuscript.

All authors reviewed and approved the final manuscript.

## Supplementary

**Supplementary Figure 1.** Cell type distributions in healthy and *S. mansoni-*infected liver and intestinal tissues.

**Supplementary Figure 2.** Spatial expression of differentially expressed genes in the infection group overlaid on representative liver and intestinal tissues.

**Supplementary Figure 3.** Validation of differentially expressed genes identified by single-cell spatial transcriptomics in tissues from healthy and *Schistosoma mansoni*–infected mice using quantitative PCR.

**Supplementary Table 1.** Top ligand–receptor pairs identified in hepatic and intestinal granulomas.

**Supplementary Table 2.** Differentially expressed genes in hepatic and intestinal granulomas.

**Supplementary Table 3.** Reactome pathway enrichment analysis of hepatic, intestinal, and shared granuloma-associated differentially expressed genes.

**Supplementary Table 4.** Statistical summary of qPCR gene expression analysis.

**Supplementary Table 5.** List of primers used for qPCR.

## Abbreviations

CCI: cell-cell interaction
cDNA: complementary DNA
CPM: counts per million
DEGs: differentially expressed genes
ENS: enteric nervous system
FDR: false discovery rate
FFPE: formalin-fixed paraffin-embedded
H&E: hematoxylin and eosin
HC: healthy controls
HSCs: hepatic stellate cells
INF: infected samples
logFC: logarithm of fold change
LR: ligand-receptor
MMCCI: multi-modal cell-cell interaction
miR-25: miRNA-25-3p
NHMRC: The National Health and Medical Research Council
PBS: phosphate-buffered saline
PCA: principal component analysis
PZQ: praziquantel
SEA: soluble egg antigens
QC: quality control
QIMRB: QIMR Berghofer Medical Research Institute
qPCR: quantitative polymerase chain reaction
ROIs: regions of interest
RT: room temperature
scRNA-seq: single-cell RNA sequencing
SC-ST: Single-Cell Spatial Transcriptomics
SMCs: smooth muscle cells
TMM: trimmed mean of M-values
Wpi: weeks post-infection

