## Supplementary Figure 1 for "Mapping fibrotic microenvironments: single-cell and spatial profiling of Schistosoma mansoni-induced tissue fibrosis"

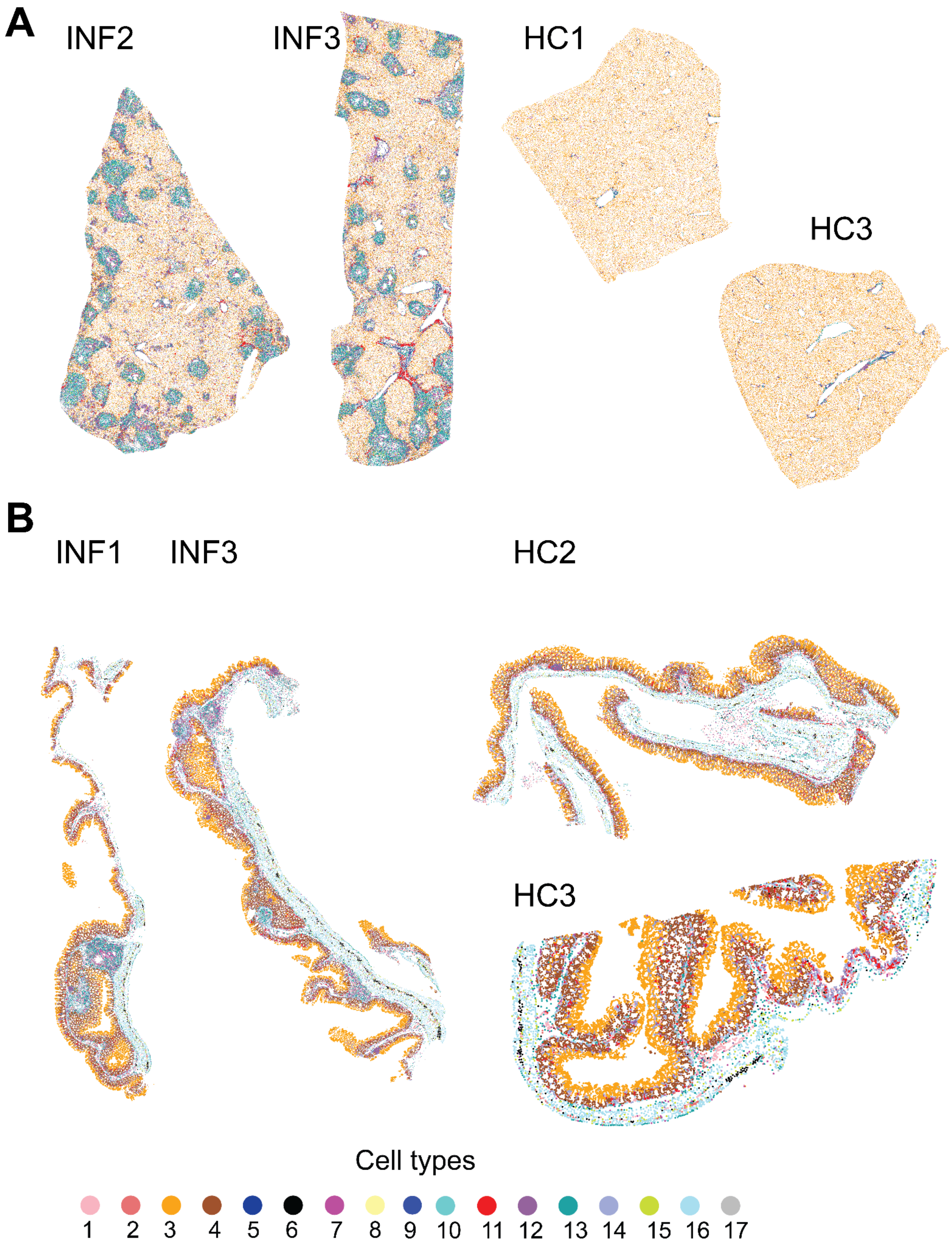


**Supplementary Figure 1. Cell type distributions in healthy and *S. mansoni*-infected (A) liver and (B) intestinal tissues.** They are encoded as: 1- Endothelial cells, 2- Enterendocrine cells, 3- Hepatocytes (L) / Enterocytes (I), 4- HSCs (L) / Goblet cells (I), 5- Tuft cells, 6- Enteric Nervous System progenitor cells, 7- Monocyte-derived Macrophages (L) /

Macrophages (I), 8- Eosinophils, 9- Epithelial cells, 10- Mast cells, 11- B/Plasma cells (L) / Plasma cells (I), 12- T cells (L) / T/B cells (I), 13- Activated HSCs (L) / Activated fibroblasts (I), 14- Fibroblasts, 15- Collagen-producing smooth muscle cells, 16- SMCs, 17- Ambiguous.
