## Supplementary Figure 2 for "Mapping fibrotic microenvironments: single-cell and spatial profiling of Schistosoma mansoni-induced tissue fibrosis"

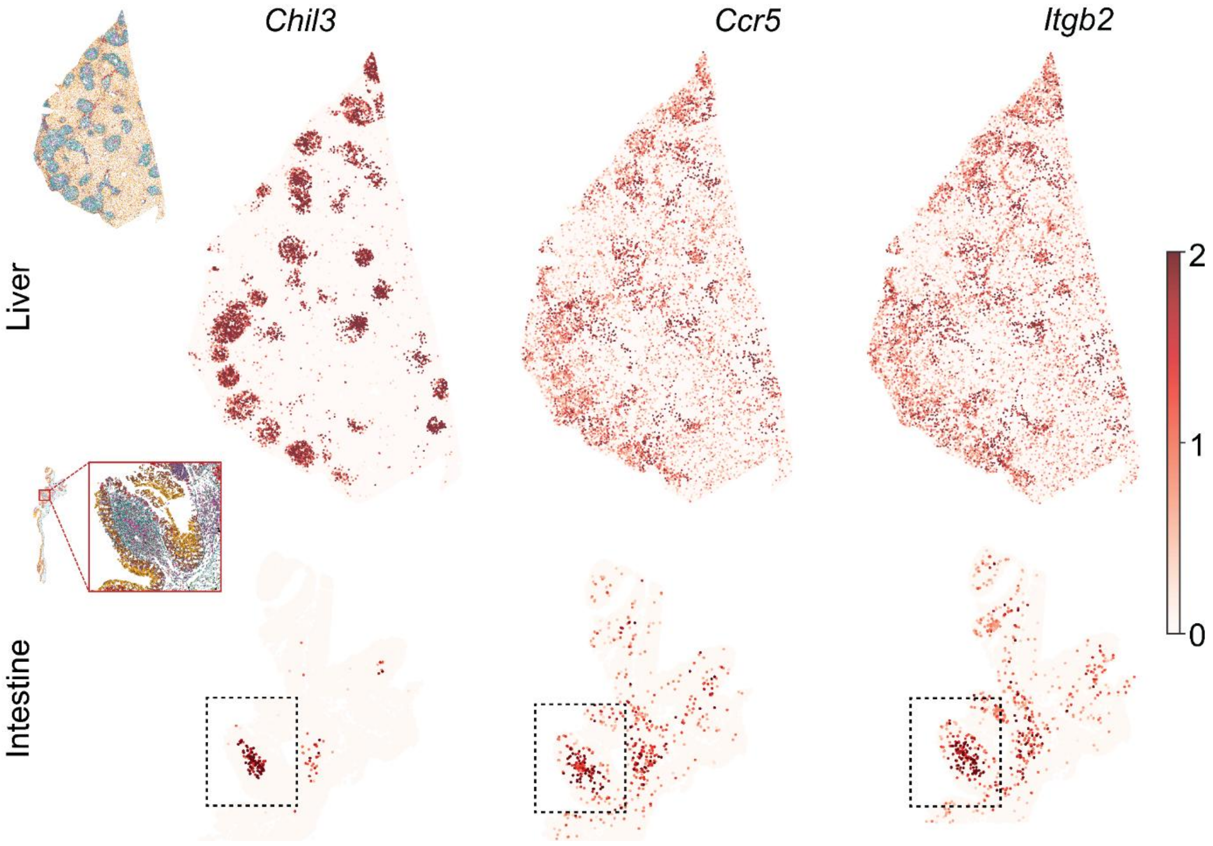


**Supplementary Figure 2. Spatial expression of differentially expressed genes in the infection group overlaid on representative liver and intestinal tissues.** The granuloma region of the intestine is highlighted with a dotted box. The whole tissue samples are shown on the top left of each row.
