## Supplementary Figure 3 for "Mapping fibrotic microenvironments: single-cell and spatial profiling of Schistosoma mansoni-induced tissue fibrosis"

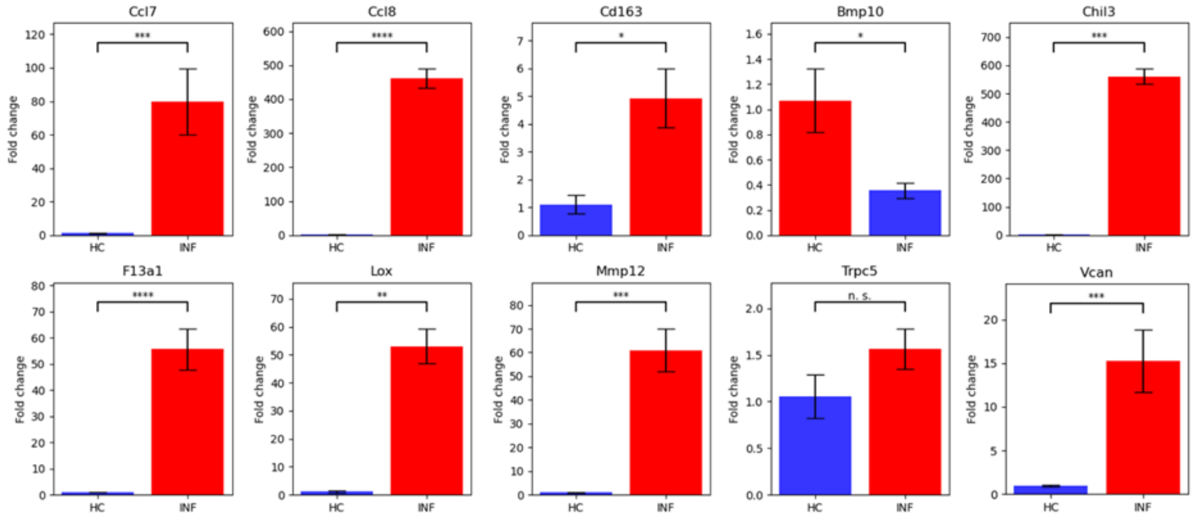


**Supplementary Figure 3. Validation of differentially expressed genes identified by single-cell spatial transcriptomics in tissues from healthy and *Schistosoma mansoni*–infected mice using quantitative PCR.** Relative expression of 10 differentially expressed genes identified by spatial transcriptomics, assessed by qPCR in liver tissue from *S. mansoni*–infected (INF, n = 3) and uninfected control (HC, n = 3) mice. Expression levels were normalised to the housekeeping gene (GapDH) and are presented as fold change (mean ± SEM). Asterisks (*) indicate statistical significance: *p ≤ 0.05, **p ≤ 0.01, ***p ≤ 0.001, ****p ≤ 0.0001. Full statistical details for all genes are provided in Supplementary Table 5.
