## Supplementary Table 2 for "Mapping fibrotic microenvironments: single-cell and spatial profiling of Schistosoma mansoni-induced tissue fibrosis"

| Gene | logFC | logCPM | F | PValue | FDR | Condition | Tissue |
|---|---|---|---|---|---|---|---|
| A1cf | 2 | 8.786462 | 466.0305 | 6.50E-31 | 7.07E-30 | Infection | Liver |
| Tpsb2 | 8.941758 | 5.507067 | 144.4141 | 2.17E-13 | 5.26E-13 | Infection | Liver |
| Aatf | 0.345293 | 7.847972 | 39.08056 | 3.87E-08 | 7.11E-08 | Infection | Liver |
| Abca1 | 0.408655 | 9.487488 | 61.95325 | 5.77E-11 | 1.23E-10 | Infection | Liver |
| Chil3 | 8.832273 | 8.993587 | 535.59 | 4.61E-36 | 1.05E-34 | Infection | Liver |
| Abca3 | -1.72322 | 10.06334 | 402.381 | 3.78E-29 | 3.37E-28 | Healthy control | Liver |
| Abca4 | 1.140342 | 2.224489 | 18.45848 | 3.82E-05 | 6.01E-05 | Infection | Liver |
| Abca7 | 0.839735 | 6.406034 | 105.6905 | 3.95E-15 | 1.05E-14 | Infection | Liver |
| Abca8a | -0.47431 | 9.408108 | 36.54799 | 8.72E-08 | 1.58E-07 | Healthy control | Liver |
| Abca8b | -0.5686 | 8.298937 | 66.38446 | 1.89E-11 | 4.12E-11 | Healthy control | Liver |
| Abcb1a | -0.69996 | 5.354276 | 35.90554 | 1.09E-07 | 1.96E-07 | Healthy control | Liver |
| Ccl11 | 8.599525 | 6.337597 | 688.2592 | 1.14E-36 | 3.00E-35 | Infection | Liver |
| Abcb4 | -0.41483 | 9.437477 | 15.15059 | 2.43E-04 | 3.62E-04 | Healthy control | Liver |
| Abcb8 | -0.47023 | 5.93795 | 33.59121 | 2.33E-07 | 4.13E-07 | Healthy control | Liver |
| Ccl8 | 8.356911 | 8.226459 | 479.467 | 2.69E-36 | 6.64E-35 | Infection | Liver |
| Abcc6 | -1.7826 | 9.709289 | 377.2089 | 2.21E-28 | 1.82E-27 | Healthy control | Liver |
| Abcc8 | 1.495524 | 1.230415 | 7.976948 | 8.74E-03 | 1.15E-02 | Infection | Liver |
| Abcc9 | 0.350465 | 7.341036 | 10.98557 | 1.52E-03 | 2.14E-03 | Infection | Liver |
| Abcd2 | -0.16445 | 7.945776 | 5.265266 | 2.51E-02 | 3.15E-02 | Healthy control | Liver |
| Abcd3 | -1.91718 | 11.17281 | 634.2471 | 1.01E-34 | 1.83E-33 | Healthy control | Liver |
| Abcf1 | -0.40365 | 7.776325 | 84.46085 | 2.88E-13 | 6.95E-13 | Healthy control | Liver |
| Abcg2 | -0.45428 | 6.950134 | 68.28218 | 1.19E-11 | 2.62E-11 | Healthy control | Liver |
| Abcg8 | -2.22141 | 8.966266 | 637.0431 | 8.98E-35 | 1.65E-33 | Healthy control | Liver |
| Abhd17c | -2.06991 | 9.148816 | 1005.815 | 1.33E-40 | 7.57E-39 | Healthy control | Liver |
| Abhd4 | -0.51957 | 7.543697 | 73.54874 | 3.37E-12 | 7.61E-12 | Healthy control | Liver |
| Abhd5 | -1.37263 | 6.889348 | 515.3924 | 3.94E-32 | 4.90E-31 | Healthy control | Liver |
| Abi1 | 1.145337 | 8.683023 | 647.2242 | 5.66E-35 | 1.07E-33 | Infection | Liver |
| Abi3bp | -1.77885 | 5.416383 | 226.6163 | 1.49E-22 | 6.74E-22 | Healthy control | Liver |
| Abitram | 0.079121 | 4.499563 | 0.452663 | 5.04E-01 | 5.31E-01 | Infection | Liver |
| Abl1 | -0.50813 | 7.157417 | 79.84617 | 7.96E-13 | 1.87E-12 | Healthy control | Liver |
| Abracl | 0.72435 | 4.977938 | 61.15717 | 7.32E-11 | 1.55E-10 | Infection | Liver |
| Abraxas1 | 0.41055 | 4.768678 | 16.93995 | 1.14E-04 | 1.74E-04 | Infection | Liver |
| Abraxas2 | -0.70829 | 7.620431 | 196.0205 | 4.61E-21 | 1.84E-20 | Healthy control | Liver |
| Acaa2 | -2.54524 | 11.66752 | 590.5136 | 7.92E-34 | 1.21E-32 | Healthy control | Liver |
| Acaca | -0.8699 | 8.83451 | 100.2526 | 1.12E-14 | 2.89E-14 | Healthy control | Liver |
| Acacb | -1.17743 | 8.095109 | 127.747 | 8.19E-17 | 2.42E-16 | Healthy control | Liver |
| Acan | 1.426398 | 1.275051 | 6.105257 | 1.72E-02 | 2.21E-02 | Infection | Liver |
| Car4 | 8.153352 | 6.319994 | 545.5264 | 4.69E-39 | 1.78E-37 | Infection | Liver |
| Ace2 | 0.697372 | 2.266701 | 4.308942 | 4.04E-02 | 4.96E-02 | Infection | Liver |
| Acer1 | 0.302624 | 1.265425 | 0.272345 | 6.04E-01 | 6.29E-01 | Infection | Liver |
| Ache | 0.426846 | 2.877425 | 3.815618 | 5.47E-02 | 6.58E-02 | Infection | Liver |
| Ackr1 | -1.25476 | 4.547221 | 105.2996 | 4.11E-15 | 1.09E-14 | Healthy control | Liver |
| Ackr2 | 0.742437 | 2.752538 | 17.64578 | 7.27E-05 | 1.12E-04 | Infection | Liver |
| Slit3 | 8.071809 | 5.876051 | 544.4665 | 2.35E-35 | 4.89E-34 | Infection | Liver |
| Ackr4 | -1.05966 | 3.945891 | 39.95171 | 2.73E-08 | 5.06E-08 | Healthy control | Liver |
| Acly | 0.230618 | 10.95789 | 5.153799 | 2.66E-02 | 3.33E-02 | Infection | Liver |
| Thbs4 | 8.053009 | 6.264777 | 373.5047 | 3.68E-33 | 5.19E-32 | Infection | Liver |
| Acot6 | -0.21518 | 4.388046 | 3.59036 | 6.27E-02 | 7.48E-02 | Healthy control | Liver |
| Acox1 | -0.79201 | 12.43654 | 59.17396 | 1.19E-10 | 2.49E-10 | Healthy control | Liver |

| | | | | | | | |
|---|---|---|---|---|---|---|---|
| Acp2 | -0.86127 | 8.168676 | 332.8092 | 6.48E-27 | 4.49E-26 | Healthy control | Liver |
| Acp5 | -1.25905 | 8.259365 | 237.4724 | 4.11E-23 | 1.95E-22 | Healthy control | Liver |
| Acr | 0.412959 | 1.448591 | 0.853051 | 3.58E-01 | 3.84E-01 | Infection | Liver |
| Acsl1 | -2.14488 | 10.72165 | 398.9001 | 5.23E-29 | 4.57E-28 | Healthy control | Liver |
| Acsl4 | -0.23525 | 8.280486 | 22.86026 | 1.08E-05 | 1.75E-05 | Healthy control | Liver |
| Acsl6 | 1.235264 | 1.532582 | 8.013095 | 5.54E-03 | 7.46E-03 | Infection | Liver |
| Col1a1 | 8.036435 | 9.821671 | 376.3393 | 4.42E-27 | 3.11E-26 | Infection | Liver |
| Acss2 | -1.03727 | 9.332614 | 80.89513 | 7.41E-13 | 1.75E-12 | Healthy control | Liver |
| Actc1 | 0.074754 | 1.347376 | 0.016754 | 8.97E-01 | 9.07E-01 | Infection | Liver |
| Timp1 | 7.832942 | 8.310127 | 470.9425 | 1.01E-32 | 1.35E-31 | Infection | Liver |
| Actl6a | -0.15581 | 6.940247 | 11.27318 | 1.33E-03 | 1.89E-03 | Healthy control | Liver |
| Actl6b | 0.566497 | 1.185196 | 1.918096 | 1.87E-01 | 2.10E-01 | Infection | Liver |
| Actn1 | 1.145096 | 8.046558 | 454.0892 | 1.34E-30 | 1.41E-29 | Infection | Liver |
| Gpnmb | 7.720919 | 5.794108 | 197.978 | 4.48E-22 | 1.96E-21 | Infection | Liver |
| Actr1a | 0.073219 | 6.959145 | 1.752242 | 1.90E-01 | 2.13E-01 | Infection | Liver |
| Actr1b | 0.013252 | 7.700986 | 0.057912 | 8.11E-01 | 8.27E-01 | Infection | Liver |
| Actr2 | -0.30242 | 9.671336 | 99.35249 | 1.33E-14 | 3.42E-14 | Healthy control | Liver |
| Acvr1 | -0.2047 | 6.700209 | 7.799081 | 6.90E-03 | 9.21E-03 | Healthy control | Liver |
| Acvr1b | 0.960462 | 6.328788 | 213.0013 | 6.19E-22 | 2.67E-21 | Infection | Liver |
| Acvr1c | -1.21909 | 1.522424 | 6.761401 | 1.05E-02 | 1.38E-02 | Healthy control | Liver |
| Acvr2a | -0.46737 | 7.221979 | 58.98417 | 1.25E-10 | 2.61E-10 | Healthy control | Liver |
| Acvr2b | -0.0772 | 4.58639 | 0.519851 | 4.74E-01 | 5.01E-01 | Healthy control | Liver |
| Acvrl1 | 0.976044 | 5.604922 | 67.14833 | 1.58E-11 | 3.47E-11 | Infection | Liver |
| Ada | -0.50557 | 4.556248 | 16.28056 | 1.50E-04 | 2.27E-04 | Healthy control | Liver |
| Adad1 | 1.959851 | 1.399152 | 12.07156 | 9.17E-04 | 1.31E-03 | Infection | Liver |
| Adad2 | 0.564597 | 1.229354 | 1.064184 | 3.11E-01 | 3.37E-01 | Infection | Liver |
| Adam10 | 0.568444 | 8.185251 | 96.26954 | 2.46E-14 | 6.25E-14 | Infection | Liver |
| Adam17 | 0.023737 | 5.690806 | 0.082441 | 7.75E-01 | 7.94E-01 | Infection | Liver |
| Adam18 | 1.252293 | 1.688858 | 6.921296 | 9.57E-03 | 1.26E-02 | Infection | Liver |
| Ch25h | 7.6953 | 4.778893 | 281.8134 | 9.69E-21 | 3.79E-20 | Infection | Liver |
| Adamdec1 | 0.948209 | 4.992587 | 42.90731 | 1.21E-08 | 2.28E-08 | Infection | Liver |
| Adamts1 | 0.463541 | 7.008604 | 21.82708 | 1.60E-05 | 2.57E-05 | Infection | Liver |
| Adamts13 | -1.18405 | 6.24032 | 128.0875 | 7.57E-17 | 2.24E-16 | Healthy control | Liver |
| Adamts14 | 1.646749 | 4.795105 | 137.7157 | 1.79E-17 | 5.53E-17 | Infection | Liver |
| Adamts18 | 0.657004 | 1.398 | 1.479032 | 2.28E-01 | 2.52E-01 | Infection | Liver |
| Ntrk1 | 7.615111 | 5.518639 | 409.9625 | 2.38E-31 | 2.74E-30 | Infection | Liver |
| Adamts3 | 1.879469 | 3.082586 | 95.96347 | 4.57E-15 | 1.21E-14 | Infection | Liver |
| Tac2 | 7.55571 | 4.766066 | 105.6666 | 1.71E-13 | 4.19E-13 | Infection | Liver |
| Slc4a1 | 7.545879 | 6.144108 | 183.9843 | 1.35E-23 | 6.72E-23 | Infection | Liver |
| Adar | 0.206877 | 7.230242 | 14.71638 | 2.90E-04 | 4.31E-04 | Infection | Liver |
| Adarb1 | 0.362967 | 5.845922 | 27.72807 | 1.78E-06 | 3.00E-06 | Infection | Liver |
| Lox | 7.513307 | 5.852015 | 432.4477 | 1.25E-36 | 3.23E-35 | Infection | Liver |
| F13a1 | 7.262984 | 6.502912 | 705.8456 | 2.02E-47 | 6.73E-45 | Infection | Liver |
| Adcy3 | 1.818684 | 5.104324 | 122.2382 | 2.19E-16 | 6.24E-16 | Infection | Liver |
| Adcy8 | 0.795544 | 1.228976 | 1.940143 | 1.74E-01 | 1.96E-01 | Infection | Liver |
| H19 | 7.158147 | 7.215356 | 460.7011 | 4.08E-36 | 9.51E-35 | Infection | Liver |
| Wnt11 | 7.065911 | 4.163137 | 189.9395 | 1.41E-17 | 4.38E-17 | Infection | Liver |
| Vcan | 7.055162 | 6.130057 | 698.8983 | 3.56E-48 | 1.62E-45 | Infection | Liver |
| C7 | 7.010657 | 4.933666 | 362.1222 | 4.68E-30 | 4.63E-29 | Infection | Liver |
| Adgra3 | -0.26856 | 6.413886 | 21.04518 | 2.17E-05 | 3.47E-05 | Healthy control | Liver |

| | | | | | | | |
|---|---|---|---|---|---|---|---|
| Adgrb1 | 0.379697 | 1.445511 | 0.618968 | 4.33E-01 | 4.61E-01 | Infection | Liver |
| Adgre1 | 0.529096 | 6.809618 | 31.88103 | 4.14E-07 | 7.23E-07 | Infection | Liver |
| Adgre5 | 0.660266 | 6.882063 | 82.77256 | 4.17E-13 | 9.97E-13 | Infection | Liver |
| Adgrf1 | -0.03835 | 3.612904 | 0.074579 | 7.86E-01 | 8.03E-01 | Healthy control | Liver |
| Adgrf5 | -1.09989 | 7.479159 | 135.9211 | 2.05E-17 | 6.30E-17 | Healthy control | Liver |
| Adgrg1 | 1.084421 | 4.606011 | 70.83682 | 6.87E-12 | 1.53E-11 | Infection | Liver |
| Olr1 | 6.977001 | 4.715587 | 378.6668 | 1.95E-29 | 1.80E-28 | Infection | Liver |
| Mmp12 | 6.970231 | 6.378677 | 809.3684 | 3.33E-49 | 2.77E-46 | Infection | Liver |
| Adgrg6 | -0.12956 | 2.218283 | 0.161859 | 6.88E-01 | 7.10E-01 | Healthy control | Liver |
| Adgrg7 | 1.338423 | 1.260954 | 5.273822 | 2.72E-02 | 3.40E-02 | Infection | Liver |
| Adgrl1 | 1.247635 | 5.869017 | 84.3178 | 3.00E-13 | 7.24E-13 | Infection | Liver |
| Adgrl2 | -0.20563 | 6.794052 | 16.56276 | 1.33E-04 | 2.02E-04 | Healthy control | Liver |
| Spon1 | 6.966822 | 5.899364 | 488.4705 | 2.92E-41 | 1.91E-39 | Infection | Liver |
| Adgrv1 | -0.6306 | 2.816991 | 7.870159 | 6.36E-03 | 8.52E-03 | Healthy control | Liver |
| Adh5 | -1.48475 | 10.8025 | 435.9709 | 4.14E-30 | 4.15E-29 | Healthy control | Liver |
| Adipoq | 1.505527 | 1.390132 | 5.856587 | 1.80E-02 | 2.31E-02 | Infection | Liver |
| Adipor1 | 0.546322 | 9.797907 | 82.73993 | 4.18E-13 | 1.00E-12 | Infection | Liver |
| Col6a2 | 6.95473 | 11.74509 | 586.3799 | 4.59E-32 | 5.68E-31 | Infection | Liver |
| Adora1 | -1.21096 | 5.882278 | 90.98946 | 7.44E-14 | 1.84E-13 | Healthy control | Liver |
| Adora2a | 1.195863 | 5.045705 | 114.865 | 7.74E-16 | 2.14E-15 | Infection | Liver |
| Ccl7 | 6.895097 | 5.753989 | 761.6006 | 1.75E-50 | 1.78E-47 | Infection | Liver |
| Adprhl1 | 0.166967 | 1.288693 | 0.07861 | 7.80E-01 | 7.98E-01 | Infection | Liver |
| Adra1a | -2.227 | 5.19835 | 408.8956 | 2.05E-29 | 1.89E-28 | Healthy control | Liver |
| Adra1b | -1.54298 | 8.053303 | 159.1247 | 8.91E-19 | 3.01E-18 | Healthy control | Liver |
| Gpr88 | 6.888465 | 5.042096 | 466.2237 | 3.39E-36 | 8.20E-35 | Infection | Liver |
| Adrb1 | 0.528551 | 4.783881 | 6.47723 | 1.34E-02 | 1.74E-02 | Infection | Liver |
| Adrb2 | 1.36629 | 4.199505 | 84.76115 | 3.01E-13 | 7.25E-13 | Infection | Liver |
| Adrb3 | -1.18453 | 2.107996 | 15.60272 | 1.45E-04 | 2.19E-04 | Healthy control | Liver |
| Adtrp | -2.34864 | 9.007578 | 437.4494 | 5.40E-30 | 5.28E-29 | Healthy control | Liver |
| Aebp2 | 0.543563 | 7.364824 | 95.68032 | 2.77E-14 | 7.03E-14 | Infection | Liver |
| Afap1l2 | 0.917556 | 5.689981 | 86.67132 | 1.81E-13 | 4.42E-13 | Infection | Liver |
| Afdn | -0.14801 | 7.60604 | 6.512452 | 1.31E-02 | 1.70E-02 | Healthy control | Liver |
| Afg3l2 | -0.91513 | 8.671488 | 593.1106 | 7.00E-34 | 1.08E-32 | Healthy control | Liver |
| Afp | 0.99743 | 1.957808 | 6.4409 | 1.23E-02 | 1.61E-02 | Infection | Liver |
| Pgf | 6.867874 | 4.659905 | 395.782 | 4.53E-30 | 4.50E-29 | Infection | Liver |
| Agl | 0.273393 | 8.294803 | 23.70865 | 7.80E-06 | 1.28E-05 | Infection | Liver |
| Ago1 | -0.62466 | 6.616386 | 111.1085 | 1.44E-15 | 3.92E-15 | Healthy control | Liver |
| Ago2 | 0.239742 | 6.553674 | 12.19703 | 8.78E-04 | 1.26E-03 | Infection | Liver |
| Ago3 | -0.10823 | 6.044278 | 2.110372 | 1.51E-01 | 1.71E-01 | Healthy control | Liver |
| Agpat3 | -1.37425 | 10.30871 | 602.9985 | 4.35E-34 | 6.93E-33 | Healthy control | Liver |
| Agpat4 | 1.238976 | 5.089521 | 67.71909 | 1.41E-11 | 3.10E-11 | Infection | Liver |
| Agps | -0.04734 | 7.01491 | 0.660991 | 4.19E-01 | 4.46E-01 | Healthy control | Liver |
| Agr3 | 1.539801 | 1.616113 | 9.260561 | 2.93E-03 | 4.03E-03 | Infection | Liver |
| Agrn | 0.56178 | 6.520334 | 84.61155 | 2.79E-13 | 6.75E-13 | Infection | Liver |
| Dnm1 | 6.836542 | 6.386933 | 656.0742 | 9.44E-45 | 1.48E-42 | Infection | Liver |
| Agt | -2.01269 | 9.717033 | 404.3453 | 3.31E-29 | 2.98E-28 | Healthy control | Liver |
| Agtr1a | -1.33949 | 7.902725 | 254.8939 | 6.87E-24 | 3.53E-23 | Healthy control | Liver |
| Ror2 | 6.835711 | 4.020447 | 244.8006 | 7.43E-20 | 2.75E-19 | Infection | Liver |
| Agtr2 | 0.873743 | 1.234038 | 2.009538 | 1.67E-01 | 1.88E-01 | Infection | Liver |
| Agtrap | 1.757898 | 5.448189 | 242.5959 | 2.65E-23 | 1.28E-22 | Infection | Liver |

| | | | | | | | |
|---|---|---|---|---|---|---|---|
| Agxt2 | -0.44532 | 8.360423 | 17.93534 | 7.60E-05 | 1.17E-04 | Healthy control | Liver |
| Ahi1 | 0.70023 | 3.957356 | 23.66168 | 8.11E-06 | 1.32E-05 | Infection | Liver |
| Ahr | -1.14954 | 6.937327 | 186.5991 | 1.51E-20 | 5.84E-20 | Healthy control | Liver |
| Aicda | 1.839852 | 1.428474 | 9.092508 | 3.56E-03 | 4.86E-03 | Infection | Liver |
| Aida | -0.7849 | 4.784201 | 54.10696 | 4.78E-10 | 9.73E-10 | Healthy control | Liver |
| Aif1 | 1.371182 | 7.776458 | 208.7623 | 1.00E-21 | 4.23E-21 | Infection | Liver |
| Pdpn | 6.791542 | 4.124427 | 154.0611 | 1.24E-16 | 3.60E-16 | Infection | Liver |
| Aifm1 | -1.44634 | 7.567532 | 343.7886 | 2.74E-27 | 1.99E-26 | Healthy control | Liver |
| Cd24a | 6.790412 | 7.57347 | 893.3469 | 4.09E-40 | 1.97E-38 | Infection | Liver |
| Aip | -0.90806 | 6.251568 | 126.9232 | 9.11E-17 | 2.68E-16 | Healthy control | Liver |
| Wt1 | 6.782263 | 6.38148 | 784.6809 | 6.02E-48 | 2.51E-45 | Infection | Liver |
| Ajuba | 1.307366 | 6.163106 | 306.2902 | 6.02E-26 | 3.79E-25 | Infection | Liver |
| Ak4 | -0.62738 | 7.240243 | 13.45272 | 5.27E-04 | 7.69E-04 | Healthy control | Liver |
| Akap5 | -0.61112 | 3.196252 | 13.41824 | 4.81E-04 | 7.04E-04 | Healthy control | Liver |
| Akap9 | 0.066529 | 7.387668 | 2.010138 | 1.61E-01 | 1.82E-01 | Infection | Liver |
| Akip1 | -0.68135 | 6.63715 | 111.7865 | 1.27E-15 | 3.48E-15 | Healthy control | Liver |
| Akr1b3 | 1.833679 | 7.020632 | 913.8617 | 2.35E-39 | 9.58E-38 | Infection | Liver |
| Akt1 | -0.77416 | 8.334335 | 117.5333 | 4.51E-16 | 1.26E-15 | Healthy control | Liver |
| Akt1s1 | -1.73659 | 7.827881 | 1132.375 | 3.88E-42 | 3.08E-40 | Healthy control | Liver |
| Akt2 | -1.57887 | 6.802549 | 719.9035 | 2.72E-36 | 6.68E-35 | Healthy control | Liver |
| Il1rl1 | 6.736331 | 4.282918 | 247.3419 | 5.12E-22 | 2.22E-21 | Infection | Liver |
| Cdh6 | 6.734776 | 4.701461 | 264.0726 | 1.33E-26 | 8.94E-26 | Infection | Liver |
| Gpbar1 | 6.690367 | 4.069781 | 387.806 | 2.85E-25 | 1.66E-24 | Infection | Liver |
| Aldh1l1 | -2.20881 | 10.328 | 336.3809 | 8.41E-27 | 5.75E-26 | Healthy control | Liver |
| Epx | 6.675446 | 3.346677 | 33.4694 | 1.91E-06 | 3.22E-06 | Infection | Liver |
| Aldh2 | -1.51938 | 10.00072 | 260.7025 | 3.84E-24 | 2.05E-23 | Healthy control | Liver |
| Aldh3a1 | 0.712135 | 1.284391 | 1.672325 | 2.02E-01 | 2.25E-01 | Infection | Liver |
| Aldh4a1 | -2.24259 | 10.70447 | 785.6691 | 1.99E-37 | 5.86E-36 | Healthy control | Liver |
| Aldh9a1 | -2.46889 | 9.237121 | 1154.359 | 2.15E-42 | 1.74E-40 | Healthy control | Liver |
| Aldoa | 0.93155 | 9.304939 | 209.9378 | 8.70E-22 | 3.69E-21 | Infection | Liver |
| Aldoc | 0.026477 | 5.314779 | 0.045868 | 8.31E-01 | 8.45E-01 | Infection | Liver |
| Alg13 | -0.24114 | 5.001953 | 7.11693 | 9.69E-03 | 1.28E-02 | Healthy control | Liver |
| Alk | 0.9608 | 1.301455 | 2.723554 | 1.05E-01 | 1.22E-01 | Infection | Liver |
| Alkbh5 | -0.35678 | 7.3706 | 51.50839 | 9.54E-10 | 1.91E-09 | Healthy control | Liver |
| Alox12 | -0.28796 | 2.920017 | 1.718625 | 1.94E-01 | 2.16E-01 | Healthy control | Liver |
| Alox12b | 1.239578 | 1.259812 | 3.234163 | 8.07E-02 | 9.49E-02 | Infection | Liver |
| Tnfrsf9 | 6.674782 | 5.218421 | 845.1825 | 1.94E-51 | 4.85E-48 | Infection | Liver |
| Eno2 | 6.622048 | 6.327501 | 853.0073 | 1.32E-48 | 7.34E-46 | Infection | Liver |
| Cdh11 | 6.573332 | 3.949192 | 201.0182 | 4.40E-19 | 1.52E-18 | Infection | Liver |
| Alpl | -0.1041 | 6.438743 | 1.900215 | 1.73E-01 | 1.94E-01 | Healthy control | Liver |
| Als2 | -0.89108 | 9.479644 | 146.4 | 3.99E-18 | 1.28E-17 | Healthy control | Liver |
| Tph1 | 6.570608 | 3.603233 | 75.66164 | 2.05E-10 | 4.25E-10 | Infection | Liver |
| Alx4 | 1.435559 | 1.363994 | 5.011335 | 2.88E-02 | 3.58E-02 | Infection | Liver |
| Ambra1 | 0.064703 | 7.874711 | 1.814739 | 1.83E-01 | 2.05E-01 | Infection | Liver |
| Amd1 | -0.31505 | 7.107134 | 21.21448 | 2.03E-05 | 3.25E-05 | Healthy control | Liver |
| Mmp7 | 6.498149 | 4.192178 | 240.8289 | 6.58E-23 | 3.07E-22 | Infection | Liver |
| Amer1 | 0.681229 | 5.140816 | 68.90642 | 1.05E-11 | 2.32E-11 | Infection | Liver |
| Amfr | -0.6221 | 11.37931 | 103.2917 | 6.19E-15 | 1.63E-14 | Healthy control | Liver |
| Amh | -0.60108 | 2.240289 | 5.379006 | 2.25E-02 | 2.84E-02 | Healthy control | Liver |
| Amhr2 | -0.033 | 1.610644 | 0.007929 | 9.29E-01 | 9.37E-01 | Healthy control | Liver |

| | | | | | | | |
|---|---|---|---|---|---|---|---|
| Amigo3 | 0.233754 | 2.865245 | 1.442732 | 2.34E-01 | 2.59E-01 | Infection | Liver |
| Amy1 | -1.81747 | 8.287235 | 239.2063 | 5.24E-23 | 2.46E-22 | Healthy control | Liver |
| Gipr | 6.478602 | 3.581681 | 89.6398 | 1.71E-11 | 3.75E-11 | Infection | Liver |
| Anapc11 | -0.87278 | 4.482435 | 79.13591 | 9.58E-13 | 2.24E-12 | Healthy control | Liver |
| Anapc15 | 0.13014 | 2.754755 | 0.379482 | 5.40E-01 | 5.67E-01 | Infection | Liver |
| Anapc16 | -0.03113 | 7.167938 | 0.30781 | 5.81E-01 | 6.07E-01 | Healthy control | Liver |
| Anapc2 | -0.58193 | 8.45248 | 149.5461 | 2.48E-18 | 8.10E-18 | Healthy control | Liver |
| Anapc4 | -0.05565 | 7.399515 | 1.50584 | 2.24E-01 | 2.49E-01 | Healthy control | Liver |
| Anapc5 | 0.30753 | 8.73202 | 51.2962 | 1.01E-09 | 2.02E-09 | Infection | Liver |
| Ang | 1.109677 | 4.034417 | 62.23855 | 5.88E-11 | 1.25E-10 | Infection | Liver |
| Angpt1 | 1.432888 | 3.063604 | 54.85048 | 2.07E-10 | 4.30E-10 | Infection | Liver |
| Angpt2 | 1.642657 | 3.166717 | 56.00311 | 1.60E-10 | 3.33E-10 | Infection | Liver |
| Col1a2 | 6.434035 | 12.37844 | 1207.82 | 5.05E-42 | 3.89E-40 | Infection | Liver |
| Angptl3 | -1.9921 | 10.49996 | 503.407 | 7.42E-32 | 8.97E-31 | Healthy control | Liver |
| Angptl4 | -1.06401 | 7.851235 | 205.6208 | 1.45E-21 | 6.03E-21 | Healthy control | Liver |
| Angptl8 | -0.72709 | 5.117466 | 32.48149 | 3.42E-07 | 6.00E-07 | Healthy control | Liver |
| Ank | -1.03459 | 8.214946 | 341.9293 | 3.15E-27 | 2.26E-26 | Healthy control | Liver |
| Fbln1 | 6.409743 | 6.23281 | 337.5184 | 2.02E-31 | 2.35E-30 | Infection | Liver |
| Nts | 6.406845 | 4.804505 | 344.0422 | 2.14E-33 | 3.11E-32 | Infection | Liver |
| Ankle2 | -0.04558 | 8.105904 | 0.872568 | 3.54E-01 | 3.80E-01 | Healthy control | Liver |
| Ankrd11 | 0.407189 | 6.831369 | 101.2566 | 9.20E-15 | 2.40E-14 | Infection | Liver |
| Ankrd13a | 0.408185 | 7.622701 | 65.72543 | 2.23E-11 | 4.83E-11 | Infection | Liver |
| Ankrd13b | 0.437242 | 5.591878 | 19.50866 | 4.00E-05 | 6.27E-05 | Infection | Liver |
| Ankrd13d | 1.484059 | 2.638901 | 37.02295 | 3.06E-08 | 5.65E-08 | Infection | Liver |
| Anks6 | 1.162234 | 3.015714 | 48.5897 | 1.34E-09 | 2.66E-09 | Infection | Liver |
| Mdk | 6.385165 | 6.521285 | 462.679 | 1.22E-34 | 2.15E-33 | Infection | Liver |
| Ano1 | -0.40607 | 3.945859 | 4.112764 | 4.68E-02 | 5.68E-02 | Healthy control | Liver |
| Cspg4 | 6.370769 | 4.486438 | 286.3782 | 1.38E-28 | 1.16E-27 | Infection | Liver |
| Anp32e | 0.694876 | 5.727599 | 60.93948 | 7.57E-11 | 1.60E-10 | Infection | Liver |
| Anxa2 | 0.785378 | 8.176194 | 23.79426 | 8.51E-06 | 1.39E-05 | Infection | Liver |
| Anxa6 | -0.49223 | 9.023001 | 99.92481 | 1.19E-14 | 3.07E-14 | Healthy control | Liver |
| Aox3 | -1.28308 | 10.3369 | 182.4778 | 2.53E-20 | 9.63E-20 | Healthy control | Liver |
| Ap1b1 | 0.538107 | 8.877054 | 145.883 | 4.32E-18 | 1.38E-17 | Infection | Liver |
| Ap1g1 | -0.4359 | 8.56419 | 52.88136 | 6.50E-10 | 1.31E-09 | Healthy control | Liver |
| Ap1s1 | -0.24103 | 6.799634 | 13.26127 | 5.47E-04 | 7.96E-04 | Healthy control | Liver |
| Ap2a1 | 0.733516 | 6.53717 | 120.0053 | 2.94E-16 | 8.30E-16 | Infection | Liver |
| Ap2a2 | -0.49592 | 8.756187 | 106.0195 | 3.68E-15 | 9.78E-15 | Healthy control | Liver |
| Ap2b1 | 0.077687 | 8.748408 | 3.949572 | 5.12E-02 | 6.18E-02 | Infection | Liver |
| Ap3b1 | -0.44761 | 7.723781 | 142.7446 | 7.00E-18 | 2.21E-17 | Healthy control | Liver |
| Ap3b2 | 0.358936 | 1.45611 | 0.485473 | 4.88E-01 | 5.15E-01 | Infection | Liver |
| Ap3m2 | 0.860955 | 5.441672 | 107.8641 | 2.68E-15 | 7.17E-15 | Infection | Liver |
| Ap3s2 | -1.30638 | 6.042013 | 378.7098 | 2.19E-28 | 1.81E-27 | Healthy control | Liver |
| Ap4s1 | -0.21279 | 5.81261 | 7.399499 | 8.41E-03 | 1.11E-02 | Healthy control | Liver |
| Ap5z1 | 0.118065 | 5.523327 | 1.982265 | 1.64E-01 | 1.85E-01 | Infection | Liver |
| Apaf1 | 1.079868 | 5.707946 | 162.646 | 3.81E-19 | 1.32E-18 | Infection | Liver |
| Dmp1 | 6.370501 | 4.287822 | 332.0241 | 5.28E-29 | 4.61E-28 | Infection | Liver |
| Apba3 | -1.30789 | 7.889772 | 735.8225 | 1.37E-36 | 3.52E-35 | Healthy control | Liver |
| Apc | 0.355201 | 7.541652 | 48.27569 | 2.40E-09 | 4.70E-09 | Infection | Liver |
| Apc2 | 0.310455 | 2.820319 | 2.217934 | 1.41E-01 | 1.60E-01 | Infection | Liver |
| Apcs | -0.00228 | 11.37665 | 0.000541 | 9.82E-01 | 9.83E-01 | Healthy control | Liver |

| | | | | | | | |
|---|---|---|---|---|---|---|---|
| Apex1 | 0.348054 | 3.679338 | 5.888271 | 1.81E-02 | 2.32E-02 | Infection | Liver |
| Aph1c | 1.628603 | 2.676032 | 51.87496 | 2.07E-10 | 4.29E-10 | Infection | Liver |
| Apip | -1.27026 | 6.248934 | 289.8868 | 2.64E-25 | 1.54E-24 | Healthy control | Liver |
| Mmp9 | 6.314181 | 5.551715 | 699.4461 | 2.53E-48 | 1.27E-45 | Infection | Liver |
| Aplnr | 1.698014 | 2.666505 | 25.37254 | 2.49E-06 | 4.17E-06 | Infection | Liver |
| Aplp2 | -0.4676 | 9.23423 | 38.26144 | 5.02E-08 | 9.17E-08 | Healthy control | Liver |
| Apmap | -0.38799 | 8.0984 | 36.87864 | 7.83E-08 | 1.42E-07 | Healthy control | Liver |
| Apoa1 | 0.33448 | 15.38935 | 8.972941 | 3.91E-03 | 5.32E-03 | Infection | Liver |
| Apoa4 | -1.718 | 9.524099 | 79.66159 | 1.69E-12 | 3.88E-12 | Healthy control | Liver |
| Apob | -1.10468 | 13.7366 | 82.01155 | 4.91E-13 | 1.17E-12 | Healthy control | Liver |
| Apobec1 | 0.201263 | 7.568245 | 5.924154 | 1.78E-02 | 2.28E-02 | Infection | Liver |
| Apobec3 | 1.650862 | 6.413205 | 477.2637 | 3.43E-31 | 3.86E-30 | Infection | Liver |
| Prrx1 | 6.268076 | 5.31153 | 450.4995 | 1.49E-39 | 6.36E-38 | Infection | Liver |
| Appbp2 | -0.29503 | 6.613374 | 29.25006 | 1.03E-06 | 1.76E-06 | Healthy control | Liver |
| Appl1 | 0.862924 | 6.988211 | 426.7514 | 7.57E-30 | 7.26E-29 | Infection | Liver |
| Appl2 | -0.45612 | 6.728858 | 45.14365 | 6.02E-09 | 1.16E-08 | Healthy control | Liver |
| Aqp2 | -0.57848 | 1.397285 | 1.12431 | 2.92E-01 | 3.18E-01 | Healthy control | Liver |
| Tmem119 | 6.218512 | 5.832682 | 505.1994 | 1.24E-39 | 5.38E-38 | Infection | Liver |
| Aqp4 | -1.99544 | 2.083993 | 26.33628 | 1.50E-06 | 2.54E-06 | Healthy control | Liver |
| Gdnf | 6.209974 | 3.507848 | 179.3529 | 2.45E-17 | 7.48E-17 | Infection | Liver |
| Aqp6 | 1.458417 | 1.289997 | 6.218809 | 1.63E-02 | 2.11E-02 | Infection | Liver |
| Ar | -1.49706 | 5.653836 | 219.1462 | 3.43E-22 | 1.51E-21 | Healthy control | Liver |
| Araf | -0.50024 | 8.08865 | 98.41441 | 1.60E-14 | 4.10E-14 | Healthy control | Liver |
| Arap1 | 0.467091 | 8.757192 | 52.33838 | 7.56E-10 | 1.52E-09 | Infection | Liver |
| Arc | 1.710618 | 2.044427 | 19.44822 | 2.20E-05 | 3.51E-05 | Infection | Liver |
| Arf2 | 0.792962 | 6.245666 | 109.2452 | 2.03E-15 | 5.48E-15 | Infection | Liver |
| Arf6 | 0.898435 | 6.746096 | 290.7078 | 2.33E-25 | 1.36E-24 | Infection | Liver |
| Arfip2 | -0.03863 | 6.287006 | 0.219953 | 6.41E-01 | 6.64E-01 | Healthy control | Liver |
| Arg1 | -0.46163 | 13.09184 | 23.43284 | 8.65E-06 | 1.41E-05 | Healthy control | Liver |
| Ank2 | 6.205188 | 3.853226 | 521.6194 | 6.39E-30 | 6.19E-29 | Infection | Liver |
| Arhgap17 | -0.81609 | 5.088547 | 90.0864 | 9.22E-14 | 2.27E-13 | Healthy control | Liver |
| Arhgap31 | 0.515721 | 6.417031 | 51.87894 | 8.61E-10 | 1.73E-09 | Infection | Liver |
| Arhgap35 | -0.83348 | 7.517489 | 183.3961 | 2.26E-20 | 8.64E-20 | Healthy control | Liver |
| Arhgap44 | 1.467119 | 3.410793 | 63.32423 | 3.21E-11 | 6.92E-11 | Infection | Liver |
| Arhgef1 | 1.067953 | 9.053051 | 618.2309 | 2.12E-34 | 3.55E-33 | Infection | Liver |
| Arhgef11 | -1.17677 | 6.697865 | 219.1174 | 3.10E-22 | 1.37E-21 | Healthy control | Liver |
| Arhgef12 | -0.57652 | 9.311267 | 228.0607 | 1.13E-22 | 5.17E-22 | Healthy control | Liver |
| Arhgef15 | -0.94948 | 6.112217 | 140.5915 | 1.01E-17 | 3.15E-17 | Healthy control | Liver |
| Fhl2 | 6.170603 | 4.979699 | 568.0878 | 6.66E-45 | 1.19E-42 | Infection | Liver |
| Arhgef3 | -0.47529 | 7.24103 | 56.5427 | 2.39E-10 | 4.95E-10 | Healthy control | Liver |
| Col6a1 | 6.148155 | 10.08017 | 757.0333 | 3.41E-35 | 6.85E-34 | Infection | Liver |
| Arhgef7 | -0.0803 | 7.233055 | 2.325819 | 1.32E-01 | 1.51E-01 | Healthy control | Liver |
| Arid1a | -1.21539 | 7.089685 | 608.3646 | 3.46E-34 | 5.62E-33 | Healthy control | Liver |
| Arid1b | -1.23274 | 6.417178 | 230.9088 | 8.55E-23 | 3.95E-22 | Healthy control | Liver |
| Arid2 | 0.014148 | 6.692556 | 0.042623 | 8.37E-01 | 8.51E-01 | Infection | Liver |
| Arid3a | 1.688436 | 3.840206 | 99.94446 | 1.24E-14 | 3.21E-14 | Infection | Liver |
| Arid3b | 0.152958 | 4.865013 | 2.442833 | 1.23E-01 | 1.42E-01 | Infection | Liver |
| Arid5b | -0.85841 | 7.136025 | 273.4617 | 1.14E-24 | 6.34E-24 | Healthy control | Liver |
| Arl13b | 0.623091 | 6.124937 | 67.5089 | 1.44E-11 | 3.16E-11 | Infection | Liver |
| Armc12 | 1.278219 | 1.359702 | 6.668209 | 1.19E-02 | 1.55E-02 | Infection | Liver |

| | | | | | | | |
|---|---|---|---|---|---|---|---|
| Arnt | 0.264047 | 7.999971 | 38.45811 | 4.72E-08 | 8.63E-08 | Infection | Liver |
| Sphk1 | 6.106988 | 4.419347 | 520.4449 | 3.68E-39 | 1.43E-37 | Infection | Liver |
| Arntl | 0.024355 | 7.727221 | 0.256325 | 6.14E-01 | 6.39E-01 | Infection | Liver |
| Arr3 | 1.57049 | 1.35439 | 7.86182 | 6.81E-03 | 9.10E-03 | Infection | Liver |
| Arrb1 | 1.063509 | 7.09406 | 169.2496 | 1.48E-19 | 5.35E-19 | Infection | Liver |
| Ace | 6.095676 | 6.555938 | 267.1482 | 7.23E-26 | 4.51E-25 | Infection | Liver |
| Arrdc3 | -1.28359 | 10.07619 | 66.65851 | 2.70E-11 | 5.83E-11 | Healthy control | Liver |
| Arx | 1.312276 | 1.397296 | 5.103825 | 2.66E-02 | 3.33E-02 | Infection | Liver |
| Asap1 | 0.011656 | 7.715299 | 0.047098 | 8.29E-01 | 8.43E-01 | Infection | Liver |
| Asap2 | -0.16618 | 6.797556 | 5.187479 | 2.61E-02 | 3.27E-02 | Healthy control | Liver |
| Asb9 | 1.172332 | 1.291128 | 3.219381 | 7.90E-02 | 9.30E-02 | Infection | Liver |
| Ascc2 | 0.705441 | 6.85802 | 146.8219 | 3.76E-18 | 1.21E-17 | Infection | Liver |
| Ascl1 | 0.577667 | 1.243722 | 1.013604 | 3.21E-01 | 3.47E-01 | Infection | Liver |
| Zbtb8b | 6.090589 | 4.105713 | 259.6712 | 7.65E-26 | 4.76E-25 | Infection | Liver |
| Asf1a | -0.7933 | 5.602878 | 69.94646 | 8.12E-12 | 1.81E-11 | Healthy control | Liver |
| Ash1l | -0.64511 | 7.540573 | 241.32 | 2.75E-23 | 1.32E-22 | Healthy control | Liver |
| Ash2l | -0.56944 | 7.610814 | 122.9997 | 1.74E-16 | 5.02E-16 | Healthy control | Liver |
| Asic1 | 1.64543 | 1.299366 | 6.099343 | 1.70E-02 | 2.19E-02 | Infection | Liver |
| Asic3 | -0.243 | 2.231018 | 1.060921 | 3.06E-01 | 3.32E-01 | Healthy control | Liver |
| Asna1 | -0.96 | 7.50519 | 439.7344 | 3.29E-30 | 3.35E-29 | Healthy control | Liver |
| Aspa | -2.04422 | 5.534466 | 266.5141 | 2.48E-24 | 1.34E-23 | Healthy control | Liver |
| Clmp | 6.084722 | 4.336078 | 292.0031 | 1.11E-29 | 1.05E-28 | Infection | Liver |
| Atad3a | -0.96947 | 7.046481 | 167.9371 | 1.78E-19 | 6.37E-19 | Healthy control | Liver |
| Atf1 | -0.61434 | 7.099513 | 126.4342 | 9.75E-17 | 2.86E-16 | Healthy control | Liver |
| Atf2 | -0.02333 | 6.243026 | 0.14583 | 7.04E-01 | 7.25E-01 | Healthy control | Liver |
| Atf4 | 0.483414 | 10.05812 | 29.59893 | 9.12E-07 | 1.56E-06 | Infection | Liver |
| Atf5 | -0.82077 | 12.4934 | 134.0225 | 2.78E-17 | 8.46E-17 | Healthy control | Liver |
| Atf6 | -0.28535 | 6.545269 | 22.37983 | 1.29E-05 | 2.09E-05 | Healthy control | Liver |
| Atf6b | -0.25501 | 7.492007 | 39.27372 | 3.64E-08 | 6.70E-08 | Healthy control | Liver |
| Atg101 | -1.49592 | 8.391038 | 686.8187 | 1.02E-35 | 2.21E-34 | Healthy control | Liver |
| Atg13 | -0.4895 | 8.368252 | 123.5179 | 1.59E-16 | 4.60E-16 | Healthy control | Liver |
| Atg14 | 0.064232 | 5.60105 | 0.498868 | 4.83E-01 | 5.10E-01 | Infection | Liver |
| Atg16l1 | -1.64767 | 6.431085 | 548.1277 | 7.42E-33 | 1.00E-31 | Healthy control | Liver |
| Atg3 | -1.56634 | 8.285915 | 727.3154 | 1.92E-36 | 4.88E-35 | Healthy control | Liver |
| Atg5 | -2.05316 | 7.972724 | 1161.505 | 1.81E-42 | 1.51E-40 | Healthy control | Liver |
| Atg7 | -0.9922 | 7.639799 | 342.9528 | 2.91E-27 | 2.11E-26 | Healthy control | Liver |
| Atg9a | -1.54375 | 7.21441 | 471.8133 | 4.68E-31 | 5.17E-30 | Healthy control | Liver |
| Atl1 | 0.074754 | 3.788391 | 0.320218 | 5.73E-01 | 5.99E-01 | Infection | Liver |
| Atm | -0.06905 | 5.023603 | 0.694638 | 4.08E-01 | 4.35E-01 | Healthy control | Liver |
| Atn1 | -1.19319 | 6.335015 | 280.2716 | 6.24E-25 | 3.56E-24 | Healthy control | Liver |
| Atoh1 | 0.491215 | 1.256672 | 0.5492 | 4.63E-01 | 4.91E-01 | Infection | Liver |
| Atoh7 | 1.072516 | 1.388229 | 3.211478 | 7.72E-02 | 9.09E-02 | Infection | Liver |
| Lypd1 | 6.072817 | 3.687413 | 265.7582 | 7.38E-23 | 3.43E-22 | Infection | Liver |
| Atp2a2 | -0.51113 | 9.830523 | 110.917 | 1.48E-15 | 4.03E-15 | Healthy control | Liver |
| Gbx2 | 6.056485 | 2.784785 | 106.6628 | 2.31E-12 | 5.26E-12 | Infection | Liver |
| Atp2b2 | -2.83446 | 5.43234 | 252.3395 | 4.78E-24 | 2.51E-23 | Healthy control | Liver |
| Atp2b3 | 1.166121 | 1.252081 | 3.989933 | 5.28E-02 | 6.35E-02 | Infection | Liver |
| Atp2b4 | 0.033339 | 5.42619 | 0.124876 | 7.25E-01 | 7.45E-01 | Infection | Liver |
| Atp4a | 1.387083 | 1.624746 | 9.999901 | 2.00E-03 | 2.79E-03 | Infection | Liver |
| Atp4b | 0.378333 | 1.234481 | 0.393486 | 5.35E-01 | 5.62E-01 | Infection | Liver |

| | | | | | | | |
|---|---|---|---|---|---|---|---|
| Atp5f1 | -1.49554 | 10.62582 | 615.9039 | 2.36E-34 | 3.92E-33 | Healthy control | Liver |
| Atp5o | -1.64928 | 9.855877 | 984.2501 | 2.53E-40 | 1.28E-38 | Healthy control | Liver |
| Atp6ap2 | 0.429932 | 8.192753 | 51.07845 | 1.08E-09 | 2.15E-09 | Infection | Liver |
| Atp6v0a1 | -0.01054 | 8.080997 | 0.024248 | 8.77E-01 | 8.89E-01 | Healthy control | Liver |
| Atp6v0d2 | 1.094443 | 6.852343 | 45.41417 | 6.94E-09 | 1.33E-08 | Infection | Liver |
| Atp6v1a | -0.52997 | 8.442493 | 53.24769 | 5.87E-10 | 1.19E-09 | Healthy control | Liver |
| Atp6v1b2 | 0.393798 | 8.414916 | 80.68949 | 6.58E-13 | 1.56E-12 | Infection | Liver |
| Atp7a | 1.076482 | 6.552567 | 209.3878 | 9.38E-22 | 3.97E-21 | Infection | Liver |
| Atp7b | -1.25412 | 6.952007 | 159.748 | 5.64E-19 | 1.93E-18 | Healthy control | Liver |
| Atr | 0.667455 | 5.752203 | 96.63167 | 2.32E-14 | 5.90E-14 | Infection | Liver |
| Atraid | -1.57276 | 7.205807 | 519.2968 | 3.16E-32 | 3.96E-31 | Healthy control | Liver |
| Atrx | 0.628166 | 7.718965 | 170.6537 | 1.22E-19 | 4.46E-19 | Infection | Liver |
| Atxn1 | -0.91408 | 6.314922 | 273.0163 | 1.21E-24 | 6.76E-24 | Healthy control | Liver |
| Atxn10 | -0.18204 | 8.507844 | 11.56216 | 1.17E-03 | 1.66E-03 | Healthy control | Liver |
| Atxn3 | -0.613 | 6.496955 | 163.0333 | 3.55E-19 | 1.24E-18 | Healthy control | Liver |
| Trem2 | 5.996907 | 6.810697 | 588.9041 | 1.06E-34 | 1.90E-33 | Infection | Liver |
| Add2 | 5.994767 | 3.705981 | 136.1636 | 3.68E-17 | 1.11E-16 | Infection | Liver |
| Avil | 1.456249 | 2.070329 | 24.64669 | 2.25E-06 | 3.77E-06 | Infection | Liver |
| Avpr1b | 0.969368 | 1.243249 | 2.959315 | 9.42E-02 | 1.10E-01 | Infection | Liver |
| Avpr2 | 0.987792 | 1.301883 | 2.70241 | 1.06E-01 | 1.23E-01 | Infection | Liver |
| Axin1 | -0.03728 | 5.891138 | 0.253434 | 6.16E-01 | 6.41E-01 | Healthy control | Liver |
| Axin2 | -0.64571 | 6.871651 | 44.77804 | 6.71E-09 | 1.29E-08 | Healthy control | Liver |
| Axl | 1.496414 | 8.084765 | 347.3277 | 2.07E-27 | 1.53E-26 | Infection | Liver |
| Azi2 | 0.512064 | 8.139985 | 71.82457 | 5.05E-12 | 1.13E-11 | Infection | Liver |
| Medag | 5.987568 | 3.473252 | 210.735 | 1.33E-19 | 4.81E-19 | Infection | Liver |
| B3glct | 0.195305 | 2.813944 | 1.26745 | 2.64E-01 | 2.90E-01 | Infection | Liver |
| B3gnt2 | -0.37778 | 7.035804 | 58.09659 | 1.58E-10 | 3.30E-10 | Healthy control | Liver |
| B3gnt6 | 0.881875 | 1.365011 | 2.731733 | 1.03E-01 | 1.20E-01 | Infection | Liver |
| B4galnt1 | -1.25579 | 9.110496 | 645.2775 | 6.18E-35 | 1.16E-33 | Healthy control | Liver |
| B4galnt2 | 1.048496 | 1.287725 | 3.808872 | 5.75E-02 | 6.89E-02 | Infection | Liver |
| B9d2 | 0.402671 | 4.406941 | 19.84609 | 3.53E-05 | 5.56E-05 | Infection | Liver |
| Babam2 | -1.8395 | 6.335207 | 571.4809 | 2.34E-33 | 3.38E-32 | Healthy control | Liver |
| Bace1 | -0.68416 | 6.199366 | 69.96824 | 7.94E-12 | 1.77E-11 | Healthy control | Liver |
| Bace2 | 0.3778 | 5.688766 | 21.92655 | 1.54E-05 | 2.48E-05 | Infection | Liver |
| Bach1 | 0.392192 | 7.747542 | 53.14349 | 6.04E-10 | 1.22E-09 | Infection | Liver |
| Bach2 | -0.30478 | 6.679731 | 17.30186 | 9.78E-05 | 1.50E-04 | Healthy control | Liver |
| Bad | 0.546175 | 7.492059 | 132.6221 | 3.49E-17 | 1.06E-16 | Infection | Liver |
| Bag2 | 0.497214 | 2.628521 | 6.400465 | 1.34E-02 | 1.74E-02 | Infection | Liver |
| Baiap2 | -0.80551 | 9.111532 | 188.368 | 1.19E-20 | 4.64E-20 | Healthy control | Liver |
| Bak1 | 1.691465 | 6.86502 | 1280.281 | 9.73E-44 | 1.04E-41 | Infection | Liver |
| Banp | 0.055155 | 6.056214 | 0.276474 | 6.01E-01 | 6.26E-01 | Infection | Liver |
| Bap1 | -1.26447 | 6.51252 | 352.7094 | 1.42E-27 | 1.08E-26 | Healthy control | Liver |
| Serpine2 | 5.962033 | 7.268034 | 205.2716 | 1.77E-21 | 7.32E-21 | Infection | Liver |
| Barhl1 | -1.49228 | 1.751286 | 13.15638 | 4.37E-04 | 6.41E-04 | Healthy control | Liver |
| Barhl2 | 1.603208 | 1.42186 | 5.980917 | 1.68E-02 | 2.16E-02 | Infection | Liver |
| Barx1 | 0.774958 | 1.254091 | 1.736081 | 1.96E-01 | 2.18E-01 | Infection | Liver |
| Aldh1a3 | 5.946984 | 4.773505 | 564.9914 | 5.27E-45 | 1.01E-42 | Infection | Liver |
| Alox15 | 5.930497 | 5.075088 | 585.0569 | 6.22E-46 | 1.64E-43 | Infection | Liver |
| Bax | 0.730088 | 8.4245 | 222.3859 | 2.11E-22 | 9.47E-22 | Infection | Liver |
| Baz1a | 1.009761 | 7.497073 | 239.5242 | 3.31E-23 | 1.58E-22 | Infection | Liver |

| | | | | | | | |
|---|---|---|---|---|---|---|---|
| Baz1b | 0.424665 | 8.094868 | 80.41057 | 7.01E-13 | 1.65E-12 | Infection | Liver |
| Bbc3 | -0.74619 | 5.844252 | 19.62034 | 3.90E-05 | 6.13E-05 | Healthy control | Liver |
| Bbox1 | -1.93785 | 9.274121 | 309.4038 | 5.94E-26 | 3.74E-25 | Healthy control | Liver |
| Bbs1 | 0.108257 | 4.021791 | 0.755892 | 3.88E-01 | 4.15E-01 | Infection | Liver |
| Bbs2 | 0.530733 | 4.596973 | 31.60642 | 4.64E-07 | 8.09E-07 | Infection | Liver |
| Bbs4 | -0.35353 | 4.312122 | 10.43686 | 1.97E-03 | 2.74E-03 | Healthy control | Liver |
| Bbs5 | 0.020271 | 4.149512 | 0.036357 | 8.49E-01 | 8.62E-01 | Infection | Liver |
| Bbs7 | 0.351302 | 5.157884 | 14.84591 | 2.75E-04 | 4.09E-04 | Infection | Liver |
| Bbs9 | -0.52365 | 5.392036 | 50.54394 | 1.28E-09 | 2.54E-09 | Healthy control | Liver |
| Bcam | 1.3932 | 4.595531 | 87.62103 | 1.63E-13 | 3.98E-13 | Infection | Liver |
| Bcar1 | -0.03457 | 8.158582 | 0.659616 | 4.20E-01 | 4.47E-01 | Healthy control | Liver |
| Bcl10 | -0.14663 | 6.717404 | 4.706555 | 3.38E-02 | 4.18E-02 | Healthy control | Liver |
| Pdcd1lg2 | 5.906504 | 5.205472 | 581.0385 | 1.08E-44 | 1.63E-42 | Infection | Liver |
| Mmp2 | 5.900496 | 8.787397 | 358.5775 | 1.86E-26 | 1.23E-25 | Infection | Liver |
| Loxl1 | 5.888636 | 5.767687 | 603.4073 | 2.27E-41 | 1.53E-39 | Infection | Liver |
| Tgfb3 | 5.868999 | 5.938194 | 357.133 | 2.19E-31 | 2.54E-30 | Infection | Liver |
| Bcl2l1 | 0.22387 | 8.171443 | 20.39772 | 2.80E-05 | 4.43E-05 | Infection | Liver |
| Bcl2l11 | 1.746047 | 6.044879 | 208.7012 | 1.04E-21 | 4.38E-21 | Infection | Liver |
| Bcl2l12 | -0.97113 | 5.679456 | 130.5087 | 5.21E-17 | 1.55E-16 | Healthy control | Liver |
| Bcl2l2 | -0.02076 | 7.356594 | 0.134408 | 7.15E-01 | 7.36E-01 | Healthy control | Liver |
| Bcl3 | -0.35158 | 6.470053 | 22.18453 | 1.40E-05 | 2.25E-05 | Healthy control | Liver |
| Bcl6 | 0.951163 | 6.937873 | 95.61035 | 2.81E-14 | 7.13E-14 | Infection | Liver |
| Bcl7a | -0.94973 | 5.946947 | 167.0182 | 2.11E-19 | 7.49E-19 | Healthy control | Liver |
| Bcl7b | 0.462777 | 7.984683 | 91.78457 | 6.11E-14 | 1.52E-13 | Infection | Liver |
| Bcl7c | -0.0998 | 7.144796 | 2.154577 | 1.47E-01 | 1.67E-01 | Healthy control | Liver |
| Bclaf1 | 0.666888 | 8.306104 | 299.0999 | 1.09E-25 | 6.65E-25 | Infection | Liver |
| Bco1 | -1.04367 | 4.199155 | 68.82666 | 9.70E-12 | 2.15E-11 | Healthy control | Liver |
| Bcor | 0.345505 | 6.723527 | 25.26897 | 4.35E-06 | 7.21E-06 | Infection | Liver |
| Bcr | -0.62341 | 7.640968 | 153.7136 | 1.34E-18 | 4.49E-18 | Healthy control | Liver |
| Bdh1 | -1.98366 | 10.75007 | 787.0222 | 1.89E-37 | 5.60E-36 | Healthy control | Liver |
| Bdkrb1 | 1.878288 | 1.328886 | 10.86294 | 1.70E-03 | 2.38E-03 | Infection | Liver |
| Galnt17 | 5.850529 | 4.413269 | 333.5267 | 8.59E-34 | 1.31E-32 | Infection | Liver |
| Bdnf | 1.412604 | 3.879079 | 80.10144 | 7.73E-13 | 1.82E-12 | Infection | Liver |
| Becn1 | -0.10779 | 7.703952 | 6.038759 | 1.67E-02 | 2.16E-02 | Healthy control | Liver |
| Itgax | 5.84706 | 5.094516 | 775.3692 | 1.78E-50 | 1.78E-47 | Infection | Liver |
| Sfrp1 | 5.84296 | 5.204482 | 332.0772 | 1.56E-33 | 2.29E-32 | Infection | Liver |
| Bglap | -0.61205 | 2.147634 | 1.775477 | 1.85E-01 | 2.08E-01 | Healthy control | Liver |
| Bglap2 | 0.069749 | 1.778696 | 0.028427 | 8.66E-01 | 8.79E-01 | Infection | Liver |
| Bhlha15 | 1.687133 | 2.054626 | 24.56414 | 2.24E-06 | 3.76E-06 | Infection | Liver |
| Aldh1l2 | 5.832804 | 4.388326 | 332.1517 | 1.16E-33 | 1.73E-32 | Infection | Liver |
| Bhlhe40 | -0.01029 | 8.144434 | 0.02302 | 8.80E-01 | 8.91E-01 | Healthy control | Liver |
| Bhmt | -2.48729 | 14.16075 | 309.6047 | 5.82E-26 | 3.67E-25 | Healthy control | Liver |
| Bicd2 | 0.222592 | 7.633916 | 13.47825 | 4.97E-04 | 7.27E-04 | Infection | Liver |
| Bid | 1.476245 | 6.245926 | 336.0327 | 5.15E-27 | 3.59E-26 | Infection | Liver |
| Bin1 | 0.975432 | 7.125255 | 282.9935 | 4.67E-25 | 2.68E-24 | Infection | Liver |
| Birc2 | -0.1422 | 5.461688 | 5.057708 | 2.80E-02 | 3.50E-02 | Healthy control | Liver |
| Birc3 | 0.271455 | 6.510598 | 15.98099 | 1.70E-04 | 2.55E-04 | Infection | Liver |
| Egr3 | 5.828511 | 4.332816 | 642.3434 | 1.17E-44 | 1.68E-42 | Infection | Liver |
| Ccl17 | 5.825689 | 3.295791 | 153.8892 | 4.64E-17 | 1.39E-16 | Infection | Liver |
| Blmh | 0.165932 | 7.171312 | 9.832142 | 2.60E-03 | 3.59E-03 | Infection | Liver |

| | | | | | | | |
|---|---|---|---|---|---|---|---|
| Aif1l | 5.813934 | 4.152745 | 532.3953 | 1.59E-39 | 6.69E-38 | Infection | Liver |
| Bloc1s5 | -1.2562 | 6.015172 | 296.342 | 1.52E-25 | 9.14E-25 | Healthy control | Liver |
| Bloc1s6 | 0.395618 | 5.705777 | 28.69269 | 1.26E-06 | 2.14E-06 | Infection | Liver |
| Bmf | 0.48668 | 5.372613 | 6.591495 | 1.27E-02 | 1.65E-02 | Infection | Liver |
| Bmi1 | 0.594404 | 8.17817 | 106.757 | 3.21E-15 | 8.55E-15 | Infection | Liver |
| Bmp1 | -0.50012 | 10.28292 | 79.1278 | 9.33E-13 | 2.18E-12 | Healthy control | Liver |
| Bmp10 | -3.48702 | 5.095683 | 274.64 | 3.33E-26 | 2.17E-25 | Healthy control | Liver |
| Bmp15 | 0.244696 | 1.275865 | 0.183365 | 6.70E-01 | 6.93E-01 | Infection | Liver |
| Bmp2 | -1.86367 | 6.677287 | 349.1439 | 1.93E-27 | 1.44E-26 | Healthy control | Liver |
| Bmp3 | 1.361187 | 1.276093 | 4.83113 | 3.32E-02 | 4.11E-02 | Infection | Liver |
| Bmp4 | 0.799574 | 4.143394 | 28.11262 | 1.59E-06 | 2.69E-06 | Infection | Liver |
| Bmp5 | -1.39421 | 5.663674 | 173.7785 | 8.80E-20 | 3.24E-19 | Healthy control | Liver |
| Bmp6 | -2.12124 | 5.066848 | 224.34 | 1.30E-22 | 5.93E-22 | Healthy control | Liver |
| Bmp7 | 0.7172 | 1.451537 | 2.274099 | 1.35E-01 | 1.54E-01 | Infection | Liver |
| Igdcc4 | 5.812023 | 4.828069 | 453.8911 | 2.03E-40 | 1.07E-38 | Infection | Liver |
| Bmpr1a | 0.667516 | 6.326593 | 140.4841 | 1.00E-17 | 3.14E-17 | Infection | Liver |
| Bmpr1b | 1.048463 | 1.311427 | 2.901287 | 9.41E-02 | 1.10E-01 | Infection | Liver |
| Bmpr2 | 0.149013 | 8.654333 | 7.869044 | 6.67E-03 | 8.92E-03 | Infection | Liver |
| Bnc1 | 1.435794 | 1.301385 | 6.218706 | 1.64E-02 | 2.11E-02 | Infection | Liver |
| Gprc5b | 5.791177 | 4.865819 | 451.4709 | 6.29E-40 | 2.86E-38 | Infection | Liver |
| Bnip3 | -0.46958 | 9.006268 | 29.50943 | 9.42E-07 | 1.61E-06 | Healthy control | Liver |
| Fzd2 | 5.760949 | 4.400452 | 621.2746 | 1.96E-45 | 4.46E-43 | Infection | Liver |
| Borcs8 | -1.32345 | 7.1226 | 298.4841 | 1.17E-25 | 7.11E-25 | Healthy control | Liver |
| Sulf1 | 5.754175 | 4.650669 | 253.2332 | 6.63E-29 | 5.76E-28 | Infection | Liver |
| Braf | -0.5349 | 5.307736 | 44.3312 | 7.80E-09 | 1.49E-08 | Healthy control | Liver |
| Brca1 | 1.409947 | 3.663721 | 92.50017 | 5.30E-14 | 1.33E-13 | Infection | Liver |
| Brca2 | 0.303338 | 3.972284 | 4.611093 | 3.56E-02 | 4.40E-02 | Infection | Liver |
| Brcc3 | 0.001776 | 5.740673 | 0.000311 | 9.86E-01 | 9.87E-01 | Infection | Liver |
| Brd2 | -0.09193 | 8.75708 | 5.051285 | 2.81E-02 | 3.51E-02 | Healthy control | Liver |
| Brd4 | -0.52818 | 8.395382 | 199.2779 | 3.09E-21 | 1.25E-20 | Healthy control | Liver |
| Brd7 | -0.8044 | 7.784092 | 257.3254 | 5.39E-24 | 2.81E-23 | Healthy control | Liver |
| Brd8 | 0.582051 | 7.837264 | 229.301 | 9.86E-23 | 4.55E-22 | Infection | Liver |
| Brip1 | 0.293533 | 4.758866 | 6.305492 | 1.46E-02 | 1.89E-02 | Infection | Liver |
| Brpf1 | 0.08813 | 6.871751 | 1.928845 | 1.70E-01 | 1.91E-01 | Infection | Liver |
| Brpf3 | -0.81514 | 6.028879 | 138.2469 | 1.45E-17 | 4.51E-17 | Healthy control | Liver |
| Brs3 | 0.636573 | 1.28026 | 1.02378 | 3.17E-01 | 3.43E-01 | Infection | Liver |
| Brsk2 | 0.580323 | 1.298528 | 0.900174 | 3.47E-01 | 3.73E-01 | Infection | Liver |
| Brwd3 | -0.78112 | 5.462269 | 158.8909 | 6.80E-19 | 2.32E-18 | Healthy control | Liver |
| Bsn | 0.286705 | 1.403223 | 0.346816 | 5.57E-01 | 5.84E-01 | Infection | Liver |
| Bsnd | 0.187317 | 1.27867 | 0.088974 | 7.67E-01 | 7.85E-01 | Infection | Liver |
| Bsx | -0.09897 | 1.207576 | 0.028838 | 8.67E-01 | 8.79E-01 | Healthy control | Liver |
| Btbd2 | -1.83586 | 4.882812 | 350.0807 | 1.50E-27 | 1.14E-26 | Healthy control | Liver |
| Btbd9 | -0.07208 | 6.741437 | 2.00504 | 1.62E-01 | 1.82E-01 | Healthy control | Liver |
| Btc | -0.13021 | 3.070455 | 0.480276 | 4.91E-01 | 5.18E-01 | Healthy control | Liver |
| Btk | 1.971896 | 4.733907 | 286.2291 | 4.99E-25 | 2.86E-24 | Infection | Liver |
| Il13 | 5.734723 | 2.979715 | 111.5048 | 2.60E-13 | 6.29E-13 | Infection | Liver |
| Btrc | 0.015317 | 6.169765 | 0.063828 | 8.01E-01 | 8.18E-01 | Infection | Liver |
| Fcrls | 5.730133 | 3.389568 | 226.9986 | 2.06E-21 | 8.48E-21 | Infection | Liver |
| Ccr1 | 5.72284 | 5.458557 | 662.6039 | 2.45E-44 | 3.27E-42 | Infection | Liver |
| C1ql1 | 0.907061 | 1.440712 | 3.446343 | 6.64E-02 | 7.88E-02 | Infection | Liver |

| | | | | | | | |
|---|---|---|---|---|---|---|---|
| C1qtnf12 | 0.917012 | 3.993695 | 53.23042 | 6.35E-10 | 1.28E-09 | Infection | Liver |
| Col12a1 | 5.65061 | 5.198125 | 529.4029 | 4.49E-41 | 2.85E-39 | Infection | Liver |
| C2 | 0.455065 | 9.517026 | 33.44674 | 2.43E-07 | 4.31E-07 | Infection | Liver |
| C2cd6 | -0.05922 | 1.233459 | 0.010999 | 9.17E-01 | 9.25E-01 | Healthy control | Liver |
| C3 | -0.41135 | 14.55566 | 28.83625 | 1.19E-06 | 2.03E-06 | Healthy control | Liver |
| C3ar1 | 1.584665 | 6.035392 | 61.6293 | 9.27E-11 | 1.95E-10 | Infection | Liver |
| C4b | 0.045165 | 12.35373 | 0.230535 | 6.33E-01 | 6.57E-01 | Infection | Liver |
| C4bp | 0.018445 | 11.87337 | 0.043918 | 8.35E-01 | 8.48E-01 | Infection | Liver |
| C5ar1 | 1.482176 | 4.189118 | 80.07913 | 8.55E-13 | 2.00E-12 | Infection | Liver |
| C5ar2 | -0.02325 | 3.740622 | 0.032483 | 8.58E-01 | 8.70E-01 | Healthy control | Liver |
| C6 | 1.496018 | 7.884235 | 195.531 | 5.53E-21 | 2.20E-20 | Infection | Liver |
| Cxcl14 | 5.622978 | 6.020222 | 572.8548 | 4.95E-37 | 1.38E-35 | Infection | Liver |
| Cab39l | -1.81498 | 7.658501 | 1090.843 | 1.20E-41 | 8.70E-40 | Healthy control | Liver |
| Ctsk | 5.614475 | 6.682741 | 291.178 | 4.56E-25 | 2.62E-24 | Infection | Liver |
| Cacna1a | -1.0474 | 3.479423 | 41.92111 | 1.17E-08 | 2.20E-08 | Healthy control | Liver |
| Tnc | 5.612009 | 5.643582 | 588.2265 | 1.44E-40 | 7.94E-39 | Infection | Liver |
| Ackr3 | 5.58794 | 5.694376 | 551.4726 | 1.56E-38 | 5.46E-37 | Infection | Liver |
| Cacna1d | 1.384503 | 3.822676 | 73.20234 | 3.89E-12 | 8.77E-12 | Infection | Liver |
| Ccl12 | 5.585209 | 4.622693 | 345.249 | 8.33E-35 | 1.54E-33 | Infection | Liver |
| Adcy2 | 5.577687 | 3.333752 | 222.5608 | 3.91E-21 | 1.57E-20 | Infection | Liver |
| Padi4 | 5.545056 | 3.215586 | 175.6602 | 3.49E-18 | 1.12E-17 | Infection | Liver |
| Itga11 | 5.537786 | 3.297247 | 155.5834 | 8.62E-18 | 2.71E-17 | Infection | Liver |
| Ncam1 | 5.530388 | 4.586846 | 243.61 | 8.61E-29 | 7.42E-28 | Infection | Liver |
| Slc16a3 | 5.499933 | 6.253915 | 548.3772 | 1.53E-34 | 2.66E-33 | Infection | Liver |
| Cacng1 | 1.39142 | 1.304463 | 4.952046 | 3.02E-02 | 3.75E-02 | Infection | Liver |
| Cacng2 | 0.200416 | 1.264475 | 0.116791 | 7.34E-01 | 7.54E-01 | Infection | Liver |
| Insrr | 5.474301 | 2.638133 | 181.7228 | 1.14E-16 | 3.34E-16 | Infection | Liver |
| Cadm1 | -1.21061 | 7.569466 | 424.8789 | 8.54E-30 | 8.14E-29 | Healthy control | Liver |
| Cadm3 | -0.56288 | 4.540336 | 18.92786 | 5.07E-05 | 7.90E-05 | Healthy control | Liver |
| Cadps2 | -1.0812 | 7.243987 | 164.1875 | 3.00E-19 | 1.05E-18 | Healthy control | Liver |
| Calb1 | 0.650038 | 1.300359 | 1.079099 | 3.03E-01 | 3.30E-01 | Infection | Liver |
| Calb2 | 1.307625 | 1.332331 | 4.559806 | 3.68E-02 | 4.53E-02 | Infection | Liver |
| Calcr | -0.10963 | 1.245396 | 0.035975 | 8.51E-01 | 8.63E-01 | Healthy control | Liver |
| Calcrl | -0.4931 | 7.113322 | 33.25869 | 2.59E-07 | 4.59E-07 | Healthy control | Liver |
| Calu | 0.93317 | 8.851742 | 213.6047 | 5.69E-22 | 2.46E-21 | Infection | Liver |
| Camk1d | -1.0887 | 5.851752 | 120.4652 | 2.81E-16 | 7.94E-16 | Healthy control | Liver |
| Jph2 | 5.462511 | 3.029796 | 169.8257 | 7.84E-18 | 2.47E-17 | Infection | Liver |
| Camk2b | -0.71091 | 2.14015 | 4.23644 | 4.21E-02 | 5.15E-02 | Healthy control | Liver |
| Serpine1 | 5.458326 | 4.735257 | 367.3161 | 1.49E-35 | 3.17E-34 | Infection | Liver |
| Camk2g | -1.41954 | 5.042955 | 167.6519 | 1.96E-19 | 6.99E-19 | Healthy control | Liver |
| Camk4 | 1.641596 | 2.291161 | 39.85904 | 5.94E-09 | 1.14E-08 | Infection | Liver |
| Camkv | -0.12338 | 1.266527 | 0.048252 | 8.27E-01 | 8.42E-01 | Healthy control | Liver |
| Camsap1 | 0.736341 | 6.174608 | 99.9876 | 1.18E-14 | 3.06E-14 | Infection | Liver |
| Camsap2 | -0.3958 | 6.776291 | 40.23877 | 2.69E-08 | 4.99E-08 | Healthy control | Liver |
| Camsap3 | -1.03497 | 6.737145 | 164.3202 | 2.96E-19 | 1.04E-18 | Healthy control | Liver |
| Camta2 | 0.013308 | 7.359186 | 0.080658 | 7.77E-01 | 7.95E-01 | Infection | Liver |
| Canx | 0.216633 | 10.512 | 18.06514 | 7.14E-05 | 1.10E-04 | Infection | Liver |
| Cap2 | 0.994325 | 1.782934 | 7.467125 | 7.12E-03 | 9.48E-03 | Infection | Liver |
| Capn1 | 0.086401 | 6.680906 | 1.033813 | 3.13E-01 | 3.40E-01 | Infection | Liver |
| Capn10 | -0.75663 | 7.796496 | 152.7233 | 1.55E-18 | 5.16E-18 | Healthy control | Liver |

| | | | | | | | |
|---|---|---|---|---|---|---|---|
| Capn2 | -0.24597 | 8.732933 | 20.69958 | 2.49E-05 | 3.95E-05 | Healthy control | Liver |
| Caprin1 | -0.30714 | 10.06781 | 46.8607 | 3.62E-09 | 7.04E-09 | Healthy control | Liver |
| Ntf5 | 5.450619 | 3.337987 | 172.4244 | 1.63E-19 | 5.86E-19 | Infection | Liver |
| Car7 | -0.02514 | 3.303839 | 0.022991 | 8.80E-01 | 8.91E-01 | Healthy control | Liver |
| Aldh1a2 | 5.44321 | 4.701409 | 453.5566 | 6.50E-40 | 2.93E-38 | Infection | Liver |
| Carm1 | -0.73812 | 7.107254 | 131.9046 | 3.94E-17 | 1.19E-16 | Healthy control | Liver |
| Carmil1 | -0.75491 | 6.103738 | 27.95321 | 1.66E-06 | 2.81E-06 | Healthy control | Liver |
| Cd34 | 5.415556 | 7.184412 | 486.4586 | 4.75E-30 | 4.69E-29 | Infection | Liver |
| Cartpt | 1.528912 | 1.438748 | 8.645407 | 4.27E-03 | 5.79E-03 | Infection | Liver |
| Casc4 | 1.978492 | 4.464908 | 125.5645 | 1.32E-16 | 3.83E-16 | Infection | Liver |
| Caskin1 | -0.66924 | 1.626264 | 2.208236 | 1.40E-01 | 1.59E-01 | Healthy control | Liver |
| Casp1 | 1.417081 | 5.758568 | 248.8089 | 1.32E-23 | 6.57E-23 | Infection | Liver |
| Casp12 | 1.441678 | 4.043806 | 78.94284 | 1.11E-12 | 2.58E-12 | Infection | Liver |
| Casp2 | 0.109258 | 4.086809 | 0.830108 | 3.66E-01 | 3.93E-01 | Infection | Liver |
| Casp3 | 0.563444 | 6.718557 | 68.58731 | 1.10E-11 | 2.44E-11 | Infection | Liver |
| Cldn7 | 5.401056 | 3.332619 | 213.4331 | 1.02E-21 | 4.30E-21 | Infection | Liver |
| Casp7 | -0.58056 | 7.161073 | 79.26831 | 9.07E-13 | 2.12E-12 | Healthy control | Liver |
| Casp8 | 0.37359 | 7.744293 | 92.35463 | 5.44E-14 | 1.36E-13 | Infection | Liver |
| Casp9 | -0.77288 | 5.614687 | 111.6997 | 1.34E-15 | 3.65E-15 | Healthy control | Liver |
| Actn2 | 5.380634 | 3.295741 | 141.5203 | 1.31E-17 | 4.09E-17 | Infection | Liver |
| Cd177 | 5.353491 | 3.759754 | 125.6479 | 4.59E-18 | 1.47E-17 | Infection | Liver |
| Rxfp3 | 5.340496 | 3.842477 | 331.7872 | 1.36E-32 | 1.78E-31 | Infection | Liver |
| Casz1 | 1.19195 | 3.464457 | 49.93178 | 1.41E-09 | 2.78E-09 | Infection | Liver |
| Cat | -2.69358 | 12.41694 | 648.2916 | 5.39E-35 | 1.03E-33 | Healthy control | Liver |
| Cx3cl1 | 5.302549 | 4.533126 | 549.7205 | 7.98E-44 | 9.08E-42 | Infection | Liver |
| Cav2 | -0.02054 | 5.832939 | 0.048121 | 8.27E-01 | 8.42E-01 | Healthy control | Liver |
| Igkc | 5.266127 | 9.32132 | 229.1611 | 7.95E-22 | 3.39E-21 | Infection | Liver |
| Tpsab1 | 5.265779 | 3.283522 | 138.4144 | 4.97E-18 | 1.58E-17 | Infection | Liver |
| Cbfb | 0.576853 | 5.77265 | 62.92159 | 4.55E-11 | 9.74E-11 | Infection | Liver |
| Ms4a7 | 5.230824 | 4.959288 | 790.2936 | 9.26E-48 | 3.57E-45 | Infection | Liver |
| Cblb | 1.108888 | 5.972433 | 99.6952 | 1.26E-14 | 3.24E-14 | Infection | Liver |
| Cblc | -1.02903 | 6.903371 | 156.9628 | 8.41E-19 | 2.85E-18 | Healthy control | Liver |
| Cblif | 0.782191 | 1.313286 | 1.588988 | 2.13E-01 | 2.36E-01 | Infection | Liver |
| Cbln1 | 1.336796 | 1.341092 | 5.046907 | 2.81E-02 | 3.51E-02 | Infection | Liver |
| Cbln2 | 1.916103 | 1.378616 | 11.66456 | 1.11E-03 | 1.58E-03 | Infection | Liver |
| Cbln3 | -0.8967 | 2.7411 | 23.14351 | 6.81E-06 | 1.12E-05 | Healthy control | Liver |
| Cbln4 | 1.424172 | 1.317293 | 6.142427 | 1.62E-02 | 2.09E-02 | Infection | Liver |
| Cbs | -2.18404 | 11.29442 | 372.1809 | 3.90E-28 | 3.15E-27 | Healthy control | Liver |
| Cbx1 | -0.37856 | 5.099023 | 13.302 | 5.39E-04 | 7.85E-04 | Healthy control | Liver |
| Cbx2 | -1.78085 | 3.394655 | 105.5256 | 4.55E-16 | 1.27E-15 | Healthy control | Liver |
| Cbx3 | 0.818957 | 9.897745 | 150.9373 | 2.02E-18 | 6.63E-18 | Infection | Liver |
| Cbx5 | 0.583437 | 6.531014 | 104.8379 | 4.63E-15 | 1.22E-14 | Infection | Liver |
| Cbx7 | -0.95651 | 4.988442 | 76.19362 | 1.91E-12 | 4.37E-12 | Healthy control | Liver |
| Ccar2 | 0.009859 | 7.53178 | 0.036677 | 8.49E-01 | 8.61E-01 | Infection | Liver |
| Ccdc130 | -1.28047 | 4.71682 | 165.9605 | 2.32E-19 | 8.23E-19 | Healthy control | Liver |
| Ccdc184 | 0.920461 | 1.265706 | 2.77992 | 1.03E-01 | 1.19E-01 | Infection | Liver |
| Cckbr | 0.56421 | 1.183671 | 1.979637 | 1.81E-01 | 2.03E-01 | Infection | Liver |
| Ffar2 | 5.213896 | 2.945786 | 202.1228 | 1.66E-19 | 5.97E-19 | Infection | Liver |
| Egr2 | 5.204353 | 3.919318 | 336.7875 | 4.65E-34 | 7.35E-33 | Infection | Liver |
| Mmp25 | 5.165467 | 3.56314 | 278.5816 | 2.14E-28 | 1.78E-27 | Infection | Liver |

| | | | | | | | |
|---|---|---|---|---|---|---|---|
| Htr1b | 5.154562 | 2.143925 | 106.4074 | 3.18E-13 | 7.65E-13 | Infection | Liver |
| Cacna1c | 5.152773 | 4.541881 | 592.3921 | 2.19E-44 | 3.04E-42 | Infection | Liver |
| Ccl21b | 5.132093 | 3.207001 | 158.3503 | 2.05E-19 | 7.31E-19 | Infection | Liver |
| Eln | 5.129087 | 6.761423 | 350.7329 | 1.92E-26 | 1.27E-25 | Infection | Liver |
| Fbln2 | 5.108905 | 6.144123 | 269.602 | 1.59E-24 | 8.72E-24 | Infection | Liver |
| Col6a3 | 5.096136 | 9.577431 | 484.3837 | 6.80E-30 | 6.58E-29 | Infection | Liver |
| Acod1 | 5.094384 | 3.118591 | 156.6499 | 9.72E-19 | 3.28E-18 | Infection | Liver |
| Ccl25 | -0.85295 | 4.297175 | 50.51122 | 1.27E-09 | 2.53E-09 | Healthy control | Liver |
| Ccl3 | 1.875149 | 2.982103 | 57.27051 | 5.99E-11 | 1.28E-10 | Infection | Liver |
| Smoc2 | 5.080523 | 7.525378 | 370.9437 | 9.06E-27 | 6.18E-26 | Infection | Liver |
| Itgam | 5.078821 | 6.325953 | 595.7788 | 3.89E-34 | 6.31E-33 | Infection | Liver |
| Ccn1 | -0.13213 | 7.329356 | 1.239583 | 2.70E-01 | 2.96E-01 | Healthy control | Liver |
| Thbs1 | 5.07488 | 6.770504 | 487.0835 | 1.45E-30 | 1.51E-29 | Infection | Liver |
| Ccna1 | 0.907586 | 1.265163 | 2.211622 | 1.44E-01 | 1.64E-01 | Infection | Liver |
| Fgf18 | 5.019626 | 2.892979 | 130.4509 | 2.52E-16 | 7.18E-16 | Infection | Liver |
| Enah | 5.000476 | 5.246563 | 665.7695 | 9.20E-41 | 5.42E-39 | Infection | Liver |
| Cacna1g | 4.993917 | 2.654884 | 95.30053 | 5.33E-13 | 1.27E-12 | Infection | Liver |
| Ccnd1 | -0.04609 | 8.453159 | 0.151693 | 6.98E-01 | 7.20E-01 | Healthy control | Liver |
| Ccnd2 | 0.859795 | 7.421585 | 83.21963 | 3.77E-13 | 9.04E-13 | Infection | Liver |
| Ccnd3 | -1.38286 | 7.724239 | 122.9358 | 2.36E-16 | 6.73E-16 | Healthy control | Liver |
| Igf2 | 4.992965 | 4.420572 | 196.7949 | 1.43E-24 | 7.91E-24 | Infection | Liver |
| Ccnt1 | -0.63058 | 6.552903 | 159.3668 | 5.97E-19 | 2.04E-18 | Healthy control | Liver |
| Ccp110 | -0.3962 | 3.379794 | 7.921943 | 6.44E-03 | 8.62E-03 | Healthy control | Liver |
| Klf5 | 4.980146 | 3.88674 | 281.6279 | 2.03E-31 | 2.35E-30 | Infection | Liver |
| Ccr10 | 1.692576 | 1.710682 | 14.31995 | 2.47E-04 | 3.68E-04 | Infection | Liver |
| Ms4a2 | 4.972492 | 2.782312 | 113.5567 | 7.75E-15 | 2.02E-14 | Infection | Liver |
| Ccr3 | -0.27584 | 5.176491 | 3.957364 | 5.10E-02 | 6.16E-02 | Healthy control | Liver |
| Ccr4 | 1.970368 | 1.719145 | 26.6969 | 1.17E-06 | 1.99E-06 | Infection | Liver |
| Tmem108 | 4.971446 | 2.60919 | 151.5688 | 1.76E-16 | 5.05E-16 | Infection | Liver |
| Alox5 | 4.946001 | 4.314085 | 474.2095 | 2.94E-41 | 1.91E-39 | Infection | Liver |
| Steap1 | 4.937479 | 4.607476 | 597.4883 | 4.31E-43 | 3.92E-41 | Infection | Liver |
| Ccr9 | 1.538623 | 1.965892 | 17.98683 | 4.24E-05 | 6.64E-05 | Infection | Liver |
| Ccser2 | -0.07911 | 5.339849 | 1.19025 | 2.79E-01 | 3.06E-01 | Healthy control | Liver |
| Cct3 | -0.31838 | 8.658752 | 25.51188 | 3.98E-06 | 6.60E-06 | Healthy control | Liver |
| Cct5 | -0.59326 | 9.639004 | 301.3288 | 8.99E-26 | 5.51E-25 | Healthy control | Liver |
| Cct7 | -0.38716 | 8.498078 | 77.74926 | 1.28E-12 | 2.95E-12 | Healthy control | Liver |
| Cct8 | -0.60104 | 9.565611 | 167.9027 | 1.78E-19 | 6.36E-19 | Healthy control | Liver |
| Chil4 | 4.920661 | 2.314879 | 60.45668 | 4.58E-10 | 9.33E-10 | Infection | Liver |
| Cd14 | 1.653331 | 5.700276 | 251.7887 | 1.00E-23 | 5.07E-23 | Infection | Liver |
| Cd151 | 0.086457 | 7.816851 | 3.715887 | 5.84E-02 | 6.99E-02 | Infection | Liver |
| Ccl21a | 4.906963 | 4.584637 | 237.4971 | 1.01E-27 | 7.86E-27 | Infection | Liver |
| Cd163 | 1.280721 | 8.526677 | 73.45946 | 5.51E-12 | 1.23E-11 | Infection | Liver |
| Cyb561 | 4.90575 | 4.310505 | 609.7339 | 1.11E-44 | 1.63E-42 | Infection | Liver |
| Crabp2 | 4.902854 | 3.61548 | 202.8625 | 2.06E-25 | 1.22E-24 | Infection | Liver |
| Apln | 4.889584 | 3.540699 | 187.3198 | 2.30E-24 | 1.24E-23 | Infection | Liver |
| Cd1d1 | -2.62779 | 8.86978 | 537.4656 | 1.75E-32 | 2.27E-31 | Healthy control | Liver |
| Cd2 | 1.923911 | 3.432528 | 82.83613 | 2.26E-13 | 5.48E-13 | Infection | Liver |
| Mfap5 | 4.887767 | 2.62428 | 78.70177 | 6.89E-12 | 1.54E-11 | Infection | Liver |
| Bnc2 | 4.876201 | 2.717982 | 132.3943 | 4.56E-16 | 1.27E-15 | Infection | Liver |
| Cd207 | -0.62434 | 4.919455 | 4.630408 | 3.55E-02 | 4.38E-02 | Healthy control | Liver |

| | | | | | | | |
|---|---|---|---|---|---|---|---|
| Gpr35 | 4.863191 | 4.835872 | 675.4656 | 3.88E-43 | 3.60E-41 | Infection | Liver |
| Cd209b | -1.8636 | 2.385686 | 42.52632 | 3.92E-09 | 7.61E-09 | Healthy control | Liver |
| Dspp | 4.850199 | 1.976104 | 55.32927 | 1.56E-09 | 3.08E-09 | Infection | Liver |
| Myo5a | 4.843175 | 6.126874 | 762.2323 | 1.25E-37 | 3.81E-36 | Infection | Liver |
| Cd226 | 0.802905 | 2.801244 | 13.61675 | 4.23E-04 | 6.22E-04 | Infection | Liver |
| Tubb3 | 4.814904 | 2.764297 | 220.2873 | 2.70E-21 | 1.10E-20 | Infection | Liver |
| Unc5c | 4.796215 | 2.249513 | 95.06815 | 6.38E-13 | 1.51E-12 | Infection | Liver |
| Clic6 | 4.775041 | 2.326561 | 105.6038 | 1.15E-13 | 2.84E-13 | Infection | Liver |
| Cyp1b1 | 4.764806 | 4.698829 | 274.0163 | 5.17E-29 | 4.52E-28 | Infection | Liver |
| Gata1 | 4.762824 | 2.774957 | 119.5677 | 3.29E-16 | 9.29E-16 | Infection | Liver |
| Spta1 | 4.752334 | 3.351182 | 108.0155 | 5.90E-17 | 1.76E-16 | Infection | Liver |
| Cd276 | 1.364395 | 3.358735 | 43.11438 | 8.66E-09 | 1.65E-08 | Infection | Liver |
| Runx2 | 4.751602 | 5.878266 | 540.3968 | 1.86E-33 | 2.71E-32 | Infection | Liver |
| Cd2ap | 0.317751 | 7.613678 | 28.40933 | 1.39E-06 | 2.36E-06 | Infection | Liver |
| Cd2bp2 | 0.003682 | 6.029332 | 0.004659 | 9.46E-01 | 9.52E-01 | Infection | Liver |
| Cd300a | 0.907392 | 4.915465 | 51.21275 | 1.07E-09 | 2.13E-09 | Infection | Liver |
| Postn | 4.749442 | 8.707207 | 380.7853 | 3.44E-27 | 2.45E-26 | Infection | Liver |
| Cd300ld | 0.219096 | 2.890952 | 0.908533 | 3.44E-01 | 3.70E-01 | Infection | Liver |
| Ank1 | 4.745283 | 3.628273 | 118.4443 | 1.86E-18 | 6.14E-18 | Infection | Liver |
| Cd320 | 0.525185 | 1.778619 | 2.050895 | 1.55E-01 | 1.75E-01 | Infection | Liver |
| Ereg | 4.739706 | 3.829 | 379.0161 | 1.09E-36 | 2.87E-35 | Infection | Liver |
| Ptchd1 | 4.736777 | 2.154233 | 139.7168 | 8.03E-16 | 2.22E-15 | Infection | Liver |
| Cd36 | -0.57631 | 8.741049 | 48.46145 | 2.27E-09 | 4.46E-09 | Healthy control | Liver |
| Ighm | 4.735699 | 9.958083 | 214.5504 | 3.57E-21 | 1.44E-20 | Infection | Liver |
| Cd38 | 0.770514 | 7.998875 | 124.0185 | 1.46E-16 | 4.24E-16 | Infection | Liver |
| Ptgir | 4.728897 | 3.826611 | 406.5969 | 1.11E-38 | 3.95E-37 | Infection | Liver |
| Glra1 | 4.720394 | 2.373778 | 93.98234 | 2.65E-13 | 6.41E-13 | Infection | Liver |
| Cd3eap | 0.610065 | 3.733441 | 20.73043 | 2.49E-05 | 3.96E-05 | Infection | Liver |
| Cd3g | 1.890608 | 4.292073 | 100.1015 | 1.35E-14 | 3.47E-14 | Infection | Liver |
| Cd4 | 1.840072 | 4.002334 | 139.8409 | 1.18E-17 | 3.67E-17 | Infection | Liver |
| Itgb8 | 4.711651 | 4.354986 | 420.6542 | 4.10E-37 | 1.17E-35 | Infection | Liver |
| Cd40lg | 1.967142 | 2.658736 | 55.55013 | 4.36E-11 | 9.36E-11 | Infection | Liver |
| Tbx1 | 4.708199 | 2.609512 | 104.0588 | 3.41E-14 | 8.62E-14 | Infection | Liver |
| Cd47 | 0.480821 | 9.667687 | 170.409 | 1.26E-19 | 4.58E-19 | Infection | Liver |
| Ugt8a | 4.707221 | 2.808515 | 90.54569 | 8.10E-14 | 2.00E-13 | Infection | Liver |
| Tnnt2 | 4.705501 | 3.48818 | 154.0021 | 4.57E-22 | 1.99E-21 | Infection | Liver |
| Cd53 | 1.725175 | 6.32436 | 180.9293 | 3.16E-20 | 1.19E-19 | Infection | Liver |
| Cd55 | -0.78326 | 7.348144 | 64.39099 | 3.11E-11 | 6.71E-11 | Healthy control | Liver |
| Cd59a | -0.69128 | 7.82038 | 60.6299 | 8.13E-11 | 1.72E-10 | Healthy control | Liver |
| Cd59b | -1.68328 | 6.151466 | 349.1276 | 2.03E-27 | 1.51E-26 | Healthy control | Liver |
| Cd5l | 0.749216 | 10.11266 | 13.42697 | 5.38E-04 | 7.84E-04 | Infection | Liver |
| Boc | 4.68626 | 2.959456 | 208.6648 | 2.82E-23 | 1.35E-22 | Infection | Liver |
| Cd68 | 1.584797 | 7.538008 | 160.8288 | 7.01E-19 | 2.39E-18 | Infection | Liver |
| Nkain3 | 4.684396 | 2.403877 | 134.4884 | 6.60E-16 | 1.84E-15 | Infection | Liver |
| Cd70 | 1.104746 | 1.206252 | 5.32349 | 3.10E-02 | 3.85E-02 | Infection | Liver |
| Fermt1 | 4.684279 | 2.906417 | 184.539 | 3.20E-21 | 1.30E-20 | Infection | Liver |
| Cd79a | 1.61395 | 3.39277 | 52.66601 | 5.34E-10 | 1.08E-09 | Infection | Liver |
| Cd79b | 1.935625 | 4.719823 | 95.1947 | 3.48E-14 | 8.78E-14 | Infection | Liver |
| Cd80 | 1.607768 | 3.380286 | 79.87703 | 5.27E-13 | 1.25E-12 | Infection | Liver |
| Cd81 | -1.29875 | 9.719921 | 248.6794 | 1.28E-23 | 6.36E-23 | Healthy control | Liver |

| | | | | | | | |
|---|---|---|---|---|---|---|---|
| Cd82 | -0.38966 | 7.315008 | 32.32914 | 3.55E-07 | 6.22E-07 | Healthy control | Liver |
| Myb | 4.672001 | 4.445088 | 229.8323 | 3.03E-27 | 2.19E-26 | Infection | Liver |
| Ptgs2 | 4.668166 | 2.823244 | 136.7705 | 1.46E-17 | 4.53E-17 | Infection | Liver |
| Cd86 | 1.276918 | 4.566058 | 114.5778 | 8.83E-16 | 2.44E-15 | Infection | Liver |
| Bub1b | 4.667133 | 6.157283 | 940.5199 | 5.55E-40 | 2.60E-38 | Infection | Liver |
| Htr2a | 4.66461 | 3.461611 | 254.2471 | 1.79E-29 | 1.67E-28 | Infection | Liver |
| Cd9 | 0.899571 | 5.697142 | 85.06086 | 2.56E-13 | 6.21E-13 | Infection | Liver |
| Cd93 | 1.772094 | 6.025718 | 331.4345 | 7.62E-27 | 5.24E-26 | Infection | Liver |
| Fst | 4.657608 | 5.410779 | 662.5204 | 2.15E-38 | 7.31E-37 | Infection | Liver |
| Cdc16 | -0.1964 | 7.119097 | 10.11826 | 2.27E-03 | 3.15E-03 | Healthy control | Liver |
| Epcam | 4.645234 | 5.117991 | 261.5876 | 1.48E-26 | 9.91E-26 | Infection | Liver |
| Cdc23 | 0.347261 | 7.965729 | 74.42341 | 2.74E-12 | 6.21E-12 | Infection | Liver |
| Cdc25a | 1.327126 | 5.456993 | 246.7398 | 1.68E-23 | 8.23E-23 | Infection | Liver |
| Cdc27 | -0.01294 | 6.842437 | 0.069057 | 7.94E-01 | 8.10E-01 | Healthy control | Liver |
| Cdc37 | -0.96257 | 8.839784 | 528.5519 | 1.87E-32 | 2.40E-31 | Healthy control | Liver |
| Cdc42 | -0.09773 | 6.091248 | 2.109144 | 1.51E-01 | 1.72E-01 | Healthy control | Liver |
| Cdc42bpb | 0.421825 | 8.229209 | 86.81618 | 1.73E-13 | 4.23E-13 | Infection | Liver |
| Cdc42ep4 | -0.65838 | 7.629937 | 192.2716 | 7.33E-21 | 2.88E-20 | Healthy control | Liver |
| Cdc5l | 0.625551 | 6.223919 | 73.41874 | 3.48E-12 | 7.87E-12 | Infection | Liver |
| Sox9 | 4.641613 | 5.99006 | 676.3101 | 3.70E-36 | 8.70E-35 | Infection | Liver |
| Cdc73 | -0.31805 | 6.851078 | 56.60159 | 2.36E-10 | 4.88E-10 | Healthy control | Liver |
| Nos2 | 4.62976 | 2.956546 | 129.9441 | 5.22E-18 | 1.66E-17 | Infection | Liver |
| Kcnq3 | 4.611073 | 3.215404 | 191.8276 | 4.12E-24 | 2.18E-23 | Infection | Liver |
| Cdcp1 | 0.078376 | 3.417242 | 0.188033 | 6.66E-01 | 6.89E-01 | Infection | Liver |
| Cdh1 | 1.369001 | 7.362562 | 38.48861 | 6.21E-08 | 1.13E-07 | Infection | Liver |
| Cdh10 | 1.90544 | 1.279167 | 12.22548 | 1.08E-03 | 1.54E-03 | Infection | Liver |
| Adgrl3 | 4.605818 | 3.643325 | 313.7341 | 1.76E-33 | 2.57E-32 | Infection | Liver |
| Cdh12 | 1.022657 | 1.519045 | 4.196467 | 4.29E-02 | 5.24E-02 | Infection | Liver |
| Cdh13 | -0.85464 | 4.12926 | 20.35655 | 2.82E-05 | 4.46E-05 | Healthy control | Liver |
| Cdh2 | -1.31012 | 8.404301 | 584.7149 | 1.06E-33 | 1.58E-32 | Healthy control | Liver |
| Cdh20 | 1.604901 | 1.380737 | 7.491479 | 7.89E-03 | 1.05E-02 | Infection | Liver |
| Hkdc1 | 4.603267 | 2.872582 | 143.6273 | 1.34E-18 | 4.49E-18 | Infection | Liver |
| Gpc3 | 4.5997 | 3.691121 | 155.6468 | 1.21E-22 | 5.56E-22 | Infection | Liver |
| Foxc2 | 4.594328 | 3.219958 | 269.9422 | 2.49E-29 | 2.27E-28 | Infection | Liver |
| Cdh4 | -1.61201 | 3.49843 | 70.00776 | 2.73E-12 | 6.19E-12 | Healthy control | Liver |
| Cdh5 | -0.32056 | 8.015573 | 18.3899 | 6.26E-05 | 9.70E-05 | Healthy control | Liver |
| Etv4 | 4.593181 | 2.867092 | 177.334 | 2.23E-21 | 9.15E-21 | Infection | Liver |
| Fosl1 | 4.592504 | 2.927379 | 127.0563 | 8.32E-18 | 2.61E-17 | Infection | Liver |
| Cdh8 | 1.785821 | 1.303588 | 7.339399 | 9.51E-03 | 1.25E-02 | Infection | Liver |
| Pla2g2e | 4.581056 | 2.509855 | 138.806 | 1.41E-16 | 4.09E-16 | Infection | Liver |
| Cercam | 4.580747 | 6.110071 | 647.6365 | 3.42E-35 | 6.85E-34 | Infection | Liver |
| Cdipt | -0.24325 | 7.937479 | 25.49768 | 4.00E-06 | 6.63E-06 | Healthy control | Liver |
| Cacna1h | 4.576853 | 3.179503 | 214.3061 | 1.51E-25 | 9.09E-25 | Infection | Liver |
| Cdk2 | 0.360481 | 5.799599 | 31.16286 | 5.31E-07 | 9.23E-07 | Infection | Liver |
| Cdk4 | 1.286623 | 3.68756 | 95.27187 | 3.14E-14 | 7.94E-14 | Infection | Liver |
| Cdk5 | 0.695937 | 6.545786 | 79.65377 | 8.33E-13 | 1.95E-12 | Infection | Liver |
| Cdk5r1 | 1.078474 | 2.836587 | 35.85863 | 7.13E-08 | 1.29E-07 | Infection | Liver |
| Cdk5r2 | 1.000209 | 1.276197 | 3.198411 | 8.06E-02 | 9.48E-02 | Infection | Liver |
| Cdk5rap2 | 0.653014 | 6.233484 | 131.4293 | 4.28E-17 | 1.29E-16 | Infection | Liver |
| Cdk6 | -0.28429 | 5.994473 | 9.984555 | 2.42E-03 | 3.35E-03 | Healthy control | Liver |

| | | | | | | | |
|---|---|---|---|---|---|---|---|
| Cdk7 | -1.62877 | 4.206479 | 203.7878 | 7.61E-22 | 3.26E-21 | Healthy control | Liver |
| Cdk8 | -0.65792 | 7.336884 | 175.1925 | 6.64E-20 | 2.47E-19 | Healthy control | Liver |
| Cdk9 | 0.507314 | 8.130492 | 109.3323 | 1.98E-15 | 5.36E-15 | Infection | Liver |
| Igha | 4.560773 | 4.92463 | 105.9827 | 3.94E-16 | 1.11E-15 | Infection | Liver |
| Cdkn1b | 0.161521 | 7.980936 | 13.25601 | 5.48E-04 | 7.97E-04 | Infection | Liver |
| Cdkn1c | -0.04747 | 5.282164 | 0.081794 | 7.76E-01 | 7.94E-01 | Healthy control | Liver |
| Cxcr4 | 4.557653 | 5.789464 | 1028.456 | 8.01E-42 | 5.99E-40 | Infection | Liver |
| Col15a1 | 4.553879 | 8.601033 | 280.1297 | 6.83E-24 | 3.52E-23 | Infection | Liver |
| Cdkn2c | 0.779726 | 5.502017 | 70.59944 | 6.87E-12 | 1.53E-11 | Infection | Liver |
| Cdkn2d | 1.018066 | 5.445236 | 68.01142 | 1.29E-11 | 2.84E-11 | Infection | Liver |
| Cdx1 | 1.119224 | 1.312681 | 4.064159 | 4.91E-02 | 5.95E-02 | Infection | Liver |
| Cdx2 | 0.979515 | 1.234148 | 2.251875 | 1.44E-01 | 1.64E-01 | Infection | Liver |
| Ceacam1 | -2.48445 | 8.431968 | 872.0237 | 9.35E-39 | 3.34E-37 | Healthy control | Liver |
| Cebpa | -1.80366 | 9.585391 | 566.4811 | 2.60E-33 | 3.73E-32 | Healthy control | Liver |
| Cebpg | -0.71132 | 7.255315 | 120.2911 | 2.79E-16 | 7.89E-16 | Healthy control | Liver |
| Celf6 | -0.0373 | 1.288181 | 0.004553 | 9.46E-01 | 9.52E-01 | Healthy control | Liver |
| Celsr1 | -1.02199 | 7.371165 | 61.07486 | 8.27E-11 | 1.75E-10 | Healthy control | Liver |
| Celsr2 | 1.337029 | 2.159504 | 22.39994 | 6.48E-06 | 1.07E-05 | Infection | Liver |
| Cyp11a1 | 4.541169 | 2.085367 | 75.01626 | 1.94E-11 | 4.22E-11 | Infection | Liver |
| Fanci | 4.529925 | 2.218492 | 142.1171 | 4.02E-16 | 1.13E-15 | Infection | Liver |
| Cenpa | 0.533777 | 5.230861 | 23.94616 | 7.17E-06 | 1.18E-05 | Infection | Liver |
| Jchain | 4.527819 | 7.740666 | 141.7651 | 3.42E-17 | 1.03E-16 | Infection | Liver |
| Pdcd1 | 4.527397 | 2.209547 | 204.2693 | 4.37E-19 | 1.51E-18 | Infection | Liver |
| Cenps | 1.576534 | 2.549424 | 51.05986 | 2.00E-10 | 4.15E-10 | Infection | Liver |
| Lama2 | 4.514768 | 6.662753 | 418.7072 | 1.71E-28 | 1.42E-27 | Infection | Liver |
| Cep104 | -0.65789 | 4.368063 | 30.00724 | 7.95E-07 | 1.37E-06 | Healthy control | Liver |
| Cep152 | 0.079717 | 5.245271 | 0.962353 | 3.30E-01 | 3.56E-01 | Infection | Liver |
| Cep164 | -0.54186 | 5.108944 | 33.59332 | 2.35E-07 | 4.17E-07 | Healthy control | Liver |
| Cep250 | 0.75611 | 6.519656 | 125.5804 | 1.13E-16 | 3.30E-16 | Infection | Liver |
| Cep290 | 0.590998 | 4.803828 | 41.1466 | 2.07E-08 | 3.87E-08 | Infection | Liver |
| Cer1 | 1.102042 | 1.332964 | 3.776527 | 5.68E-02 | 6.82E-02 | Infection | Liver |
| Prkar2b | 4.513425 | 5.895019 | 773.6872 | 1.05E-37 | 3.24E-36 | Infection | Liver |
| Ndrg4 | 4.506721 | 3.716364 | 380.9829 | 3.74E-37 | 1.08E-35 | Infection | Liver |
| Cers2 | -1.59234 | 9.916171 | 757.3793 | 5.84E-37 | 1.60E-35 | Healthy control | Liver |
| Cers4 | 1.174621 | 6.216271 | 203.4022 | 1.91E-21 | 7.90E-21 | Infection | Liver |
| Cers5 | 1.807969 | 5.048388 | 228.5586 | 1.29E-22 | 5.91E-22 | Infection | Liver |
| Cers6 | -0.10521 | 5.132022 | 1.104968 | 2.97E-01 | 3.23E-01 | Healthy control | Liver |
| Cetn2 | -1.17254 | 5.249733 | 170.9888 | 1.31E-19 | 4.75E-19 | Healthy control | Liver |
| Cetn3 | -0.29018 | 7.219191 | 32.88424 | 2.94E-07 | 5.19E-07 | Healthy control | Liver |
| Cfap299 | 0.546785 | 2.280181 | 4.278884 | 4.12E-02 | 5.04E-02 | Infection | Liver |
| Cfap36 | -0.50594 | 6.828023 | 102.9241 | 6.68E-15 | 1.75E-14 | Healthy control | Liver |
| Cfap44 | 1.875739 | 1.311993 | 10.97103 | 1.74E-03 | 2.43E-03 | Infection | Liver |
| Jag2 | 4.503284 | 3.284734 | 241.6462 | 2.19E-28 | 1.81E-27 | Infection | Liver |
| Cfap52 | 1.431306 | 1.344674 | 5.239077 | 2.55E-02 | 3.20E-02 | Infection | Liver |
| Cfap53 | 0.504365 | 1.507418 | 1.135604 | 2.89E-01 | 3.15E-01 | Infection | Liver |
| Cfap57 | -0.10131 | 1.647363 | 0.06396 | 8.01E-01 | 8.17E-01 | Healthy control | Liver |
| Dgkg | 4.501052 | 3.11565 | 209.4483 | 3.19E-25 | 1.85E-24 | Infection | Liver |
| Cfap65 | 0.740723 | 1.492474 | 2.339403 | 1.29E-01 | 1.48E-01 | Infection | Liver |
| Gfpt2 | 4.493841 | 4.801727 | 292.018 | 1.33E-28 | 1.12E-27 | Infection | Liver |
| Cfb | -0.18424 | 11.7528 | 2.835457 | 9.71E-02 | 1.13E-01 | Healthy control | Liver |

| | | | | | | | |
|---|---|---|---|---|---|---|---|
| Cst7 | 4.489816 | 3.911769 | 235.0887 | 9.86E-29 | 8.44E-28 | Infection | Liver |
| Cflar | -0.81595 | 7.6954 | 329.4962 | 8.49E-27 | 5.80E-26 | Healthy control | Liver |
| Adora2b | 4.454714 | 2.822251 | 165.4752 | 1.15E-20 | 4.50E-20 | Infection | Liver |
| Cga | 0.575022 | 1.323095 | 0.908767 | 3.44E-01 | 3.70E-01 | Infection | Liver |
| Cgas | 1.771031 | 2.256497 | 38.0264 | 1.14E-08 | 2.16E-08 | Infection | Liver |
| Cgn | -0.97264 | 8.314524 | 132.8063 | 3.39E-17 | 1.03E-16 | Healthy control | Liver |
| Itga2 | 4.447873 | 3.045545 | 243.6123 | 2.48E-27 | 1.82E-26 | Infection | Liver |
| Champ1 | -0.21054 | 5.671573 | 10.56558 | 1.85E-03 | 2.58E-03 | Healthy control | Liver |
| Chat | 0.869996 | 1.231053 | 2.37407 | 1.34E-01 | 1.53E-01 | Infection | Liver |
| Chchd3 | -0.819 | 8.470253 | 242.3191 | 2.47E-23 | 1.20E-22 | Healthy control | Liver |
| Chd1 | -0.22049 | 7.820707 | 26.81761 | 2.47E-06 | 4.13E-06 | Healthy control | Liver |
| Chd4 | 0.549863 | 9.752745 | 218.9535 | 3.09E-22 | 1.37E-21 | Infection | Liver |
| Chd6 | 0.974791 | 6.229122 | 194.8823 | 5.37E-21 | 2.14E-20 | Infection | Liver |
| Chd7 | -0.71719 | 5.645888 | 67.39519 | 1.51E-11 | 3.31E-11 | Healthy control | Liver |
| Plppr3 | 4.436715 | 4.524809 | 367.0478 | 5.29E-33 | 7.28E-32 | Infection | Liver |
| Chek2 | 0.866068 | 4.309306 | 55.50192 | 3.40E-10 | 6.98E-10 | Infection | Liver |
| Kcnn4 | 4.434319 | 4.125711 | 360.6677 | 9.94E-35 | 1.80E-33 | Infection | Liver |
| Chic2 | 1.074727 | 6.432554 | 217.603 | 3.66E-22 | 1.61E-21 | Infection | Liver |
| Cdkn2a | 4.432731 | 5.206217 | 375.177 | 9.35E-31 | 9.96E-30 | Infection | Liver |
| Arhgef28 | 4.428572 | 3.044911 | 268.4409 | 9.69E-29 | 8.30E-28 | Infection | Liver |
| Nrg1 | 4.421829 | 3.814576 | 237.904 | 1.17E-28 | 9.95E-28 | Infection | Liver |
| Chka | -0.24154 | 7.880037 | 7.409204 | 8.37E-03 | 1.11E-02 | Healthy control | Liver |
| Chmp3 | -0.63279 | 6.666139 | 128.0656 | 7.45E-17 | 2.21E-16 | Healthy control | Liver |
| Chmp4c | 0.61325 | 4.573985 | 30.94526 | 5.83E-07 | 1.01E-06 | Infection | Liver |
| Chrd | -0.41623 | 4.228252 | 15.47592 | 2.11E-04 | 3.16E-04 | Healthy control | Liver |
| Chrm1 | -2.63944 | 2.298095 | 73.90932 | 2.38E-13 | 5.77E-13 | Healthy control | Liver |
| Cpt1c | 4.413805 | 4.779672 | 631.0588 | 1.34E-39 | 5.80E-38 | Infection | Liver |
| Ptges | 4.413708 | 4.355372 | 282.6611 | 1.17E-29 | 1.11E-28 | Infection | Liver |
| Chrm4 | 0.499554 | 1.256135 | 0.721352 | 4.01E-01 | 4.29E-01 | Infection | Liver |
| Chrm5 | 0.728678 | 1.248243 | 1.185929 | 2.84E-01 | 3.10E-01 | Infection | Liver |
| Chrna1 | 1.974167 | 1.277904 | 10.81001 | 2.07E-03 | 2.89E-03 | Infection | Liver |
| Chrna2 | -3.40566 | 3.711527 | 311.5603 | 2.77E-31 | 3.14E-30 | Healthy control | Liver |
| Ciita | 4.401203 | 6.85255 | 1290.534 | 1.56E-43 | 1.59E-41 | Infection | Liver |
| Chrna6 | 1.121455 | 1.292011 | 3.278498 | 7.64E-02 | 9.01E-02 | Infection | Liver |
| Chrna7 | 1.359117 | 1.385172 | 5.327284 | 2.39E-02 | 3.02E-02 | Infection | Liver |
| Gdf11 | 4.393915 | 3.893102 | 375.7499 | 2.68E-36 | 6.64E-35 | Infection | Liver |
| Chrnb2 | -1.03637 | 2.833242 | 31.32757 | 2.98E-07 | 5.25E-07 | Healthy control | Liver |
| Chrnb3 | 0.212305 | 1.411333 | 0.174699 | 6.77E-01 | 7.00E-01 | Infection | Liver |
| H2-DMb2 | 4.370597 | 4.875692 | 487.7151 | 6.80E-35 | 1.26E-33 | Infection | Liver |
| Chrnd | 0.769168 | 1.198065 | 2.64219 | 1.21E-01 | 1.39E-01 | Infection | Liver |
| Chrng | 1.679806 | 1.253929 | 9.622563 | 3.63E-03 | 4.95E-03 | Infection | Liver |
| Chst10 | 1.786049 | 1.655374 | 17.88407 | 4.91E-05 | 7.66E-05 | Infection | Liver |
| Chst8 | 0.676195 | 1.667225 | 2.731924 | 1.01E-01 | 1.17E-01 | Infection | Liver |
| Chtop | -0.15863 | 7.554778 | 6.815905 | 1.13E-02 | 1.47E-02 | Healthy control | Liver |
| Chuk | -1.26022 | 8.28432 | 234.9645 | 5.36E-23 | 2.51E-22 | Healthy control | Liver |
| Ciart | 1.414588 | 1.693358 | 12.15716 | 6.77E-04 | 9.77E-04 | Infection | Liver |
| Cib2 | -1.54745 | 4.405429 | 76.75534 | 1.36E-12 | 3.14E-12 | Healthy control | Liver |
| Cic | 0.486727 | 7.125233 | 37.27242 | 6.90E-08 | 1.25E-07 | Infection | Liver |
| Itgb4 | 4.369163 | 3.037065 | 158.7583 | 1.44E-21 | 6.00E-21 | Infection | Liver |
| Cideb | -1.81771 | 10.25035 | 463.5034 | 7.53E-31 | 8.13E-30 | Healthy control | Liver |

| | | | | | | | |
|---|---|---|---|---|---|---|---|
| Tnfsf13b | 4.340167 | 5.958288 | 499.7872 | 1.34E-31 | 1.60E-30 | Infection | Liver |
| Cisd2 | -0.75742 | 7.613945 | 179.1007 | 3.95E-20 | 1.48E-19 | Healthy control | Liver |
| Cish | 1.43976 | 6.528408 | 344.5283 | 2.62E-27 | 1.92E-26 | Infection | Liver |
| Cit | 1.354371 | 4.905584 | 90.34036 | 8.87E-14 | 2.19E-13 | Infection | Liver |
| Cited1 | 1.077895 | 1.404686 | 3.969966 | 4.96E-02 | 6.00E-02 | Infection | Liver |
| Cited2 | -0.97411 | 6.615534 | 203.9328 | 1.80E-21 | 7.44E-21 | Healthy control | Liver |
| Cited4 | 0.192072 | 2.697982 | 0.804036 | 3.73E-01 | 4.00E-01 | Infection | Liver |
| Cd244a | 4.327503 | 4.218609 | 454.5842 | 1.14E-37 | 3.49E-36 | Infection | Liver |
| Cklf | 1.475406 | 2.048038 | 19.29797 | 2.36E-05 | 3.76E-05 | Infection | Liver |
| Cks1b | 0.926823 | 5.570962 | 91.13353 | 7.13E-14 | 1.77E-13 | Infection | Liver |
| Clca3a2 | 1.838539 | 2.060798 | 25.86438 | 1.35E-06 | 2.28E-06 | Infection | Liver |
| Clcn4 | -1.39629 | 5.308432 | 197.0758 | 4.65E-21 | 1.86E-20 | Healthy control | Liver |
| Clcn7 | -1.24254 | 7.80807 | 569.2289 | 2.28E-33 | 3.32E-32 | Healthy control | Liver |
| Clcnka | 1.233195 | 1.256891 | 4.353716 | 4.37E-02 | 5.33E-02 | Infection | Liver |
| Clcnkb | 0.327102 | 1.218472 | 0.374268 | 5.46E-01 | 5.72E-01 | Infection | Liver |
| Cldn1 | -0.60637 | 8.563976 | 33.33091 | 2.59E-07 | 4.59E-07 | Healthy control | Liver |
| Mmrn1 | 4.320893 | 2.864272 | 148.0569 | 2.57E-20 | 9.76E-20 | Infection | Liver |
| Cldn12 | -1.18389 | 8.002472 | 251.9367 | 9.23E-24 | 4.69E-23 | Healthy control | Liver |
| Cldn14 | -0.33362 | 6.012834 | 10.7057 | 1.73E-03 | 2.43E-03 | Healthy control | Liver |
| Cldn16 | 1.619114 | 1.292914 | 6.156261 | 1.68E-02 | 2.16E-02 | Infection | Liver |
| Cldn17 | -0.40029 | 1.28908 | 0.392727 | 5.33E-01 | 5.61E-01 | Healthy control | Liver |
| Cldn19 | -0.19936 | 1.196903 | 0.143114 | 7.10E-01 | 7.31E-01 | Healthy control | Liver |
| Cldn2 | -1.0995 | 8.554694 | 193.8663 | 6.00E-21 | 2.37E-20 | Healthy control | Liver |
| Cldn23 | 1.935458 | 1.385754 | 10.52151 | 1.90E-03 | 2.66E-03 | Infection | Liver |
| Cldn6 | 1.213153 | 1.324658 | 4.006268 | 5.01E-02 | 6.06E-02 | Infection | Liver |
| Pou2af1 | 4.303908 | 4.987167 | 259.5293 | 1.61E-25 | 9.59E-25 | Infection | Liver |
| Mcub | 4.302202 | 3.610402 | 245.0967 | 2.75E-29 | 2.49E-28 | Infection | Liver |
| Cldn9 | 0.073935 | 1.333718 | 0.016851 | 8.97E-01 | 9.07E-01 | Infection | Liver |
| Ddr1 | 4.297542 | 4.835073 | 1090.984 | 9.59E-47 | 2.82E-44 | Infection | Liver |
| Clec14a | -1.03449 | 6.885891 | 177.157 | 5.15E-20 | 1.93E-19 | Healthy control | Liver |
| Clec1b | 0.187253 | 5.940495 | 2.461348 | 1.22E-01 | 1.40E-01 | Infection | Liver |
| 3-Sep | 4.293717 | 2.086142 | 125.7785 | 1.77E-15 | 4.79E-15 | Infection | Liver |
| Clec4n | 1.067969 | 4.397151 | 32.68548 | 3.25E-07 | 5.71E-07 | Infection | Liver |
| Clec9a | 1.285759 | 4.587288 | 76.59522 | 1.83E-12 | 4.19E-12 | Infection | Liver |
| Clgn | 1.244368 | 1.346189 | 4.30178 | 4.21E-02 | 5.15E-02 | Infection | Liver |
| Clic3 | -2.657 | 1.877265 | 46.41439 | 1.52E-09 | 3.01E-09 | Healthy control | Liver |
| Clic4 | -0.57702 | 7.784915 | 124.3293 | 1.39E-16 | 4.03E-16 | Healthy control | Liver |
| Clic5 | -0.65249 | 3.965535 | 17.99799 | 7.28E-05 | 1.13E-04 | Healthy control | Liver |
| Clec4e | 4.278057 | 2.177551 | 116.1949 | 4.07E-15 | 1.08E-14 | Infection | Liver |
| Clk1 | 0.245486 | 8.049689 | 18.73791 | 5.44E-05 | 8.45E-05 | Infection | Liver |
| Ttk | 4.276878 | 4.359853 | 519.1164 | 1.61E-38 | 5.59E-37 | Infection | Liver |
| Cln3 | -0.05989 | 4.997568 | 0.619769 | 4.34E-01 | 4.62E-01 | Healthy control | Liver |
| Cln6 | 1.31138 | 5.633695 | 184.2535 | 2.10E-20 | 8.07E-20 | Infection | Liver |
| Cln8 | -0.14421 | 6.02395 | 5.338955 | 2.41E-02 | 3.04E-02 | Healthy control | Liver |
| Clock | -0.35343 | 7.19288 | 48.66346 | 2.14E-09 | 4.22E-09 | Healthy control | Liver |
| Mtcl1 | 4.269228 | 2.425584 | 109.0733 | 3.57E-15 | 9.50E-15 | Infection | Liver |
| Cltb | -0.41076 | 6.637002 | 32.79338 | 3.04E-07 | 5.34E-07 | Healthy control | Liver |
| Cltc | -0.07038 | 10.15472 | 3.490998 | 6.63E-02 | 7.88E-02 | Healthy control | Liver |
| Foxj1 | 4.267366 | 2.703088 | 132.1158 | 2.65E-18 | 8.61E-18 | Infection | Liver |
| Cmah | -1.20571 | 7.59608 | 107.3168 | 3.23E-15 | 8.61E-15 | Healthy control | Liver |

| | | | | | | | |
|---|---|---|---|---|---|---|---|
| Cmip | -1.04957 | 7.756237 | 122.1785 | 2.01E-16 | 5.75E-16 | Healthy control | Liver |
| Cmklr1 | 1.836722 | 5.436117 | 114.0685 | 9.87E-16 | 2.72E-15 | Infection | Liver |
| Cmtm6 | 0.111405 | 8.640827 | 4.262046 | 4.31E-02 | 5.26E-02 | Infection | Liver |
| Cndp1 | 1.143841 | 1.241727 | 3.684821 | 6.40E-02 | 7.62E-02 | Infection | Liver |
| Cnga2 | 0.965681 | 1.375124 | 3.166386 | 7.92E-02 | 9.32E-02 | Infection | Liver |
| Ephb1 | 4.261897 | 1.906226 | 121.3132 | 1.78E-15 | 4.84E-15 | Infection | Liver |
| Cnmd | -2.13311 | 3.397118 | 113.347 | 4.56E-17 | 1.37E-16 | Healthy control | Liver |
| Cnn1 | 0.643948 | 1.297335 | 1.205528 | 2.77E-01 | 3.03E-01 | Infection | Liver |
| Cnot11 | -0.40039 | 5.088202 | 22.11069 | 1.45E-05 | 2.33E-05 | Healthy control | Liver |
| Cnot2 | 0.310568 | 6.580167 | 31.50825 | 4.70E-07 | 8.19E-07 | Infection | Liver |
| Cnot4 | -0.36492 | 4.620095 | 14.08353 | 3.84E-04 | 5.66E-04 | Healthy control | Liver |
| Cnot8 | -0.73922 | 7.021605 | 198.7013 | 3.34E-21 | 1.35E-20 | Healthy control | Liver |
| Cnot9 | -0.22105 | 6.830749 | 15.12249 | 2.44E-04 | 3.64E-04 | Healthy control | Liver |
| Cnp | -0.39888 | 7.053417 | 44.98819 | 6.30E-09 | 1.21E-08 | Healthy control | Liver |
| Cnppd1 | -0.03597 | 8.356591 | 0.37413 | 5.43E-01 | 5.69E-01 | Healthy control | Liver |
| Cnpy1 | 0.946132 | 1.235354 | 2.223704 | 1.47E-01 | 1.67E-01 | Infection | Liver |
| Cnr2 | -0.58209 | 5.737461 | 48.47989 | 2.28E-09 | 4.49E-09 | Healthy control | Liver |
| Cntf | 0.311116 | 3.017044 | 3.177157 | 7.90E-02 | 9.30E-02 | Infection | Liver |
| Cntn1 | 1.247368 | 1.389756 | 5.212065 | 2.52E-02 | 3.17E-02 | Infection | Liver |
| Rhbdl3 | 4.250014 | 3.57857 | 269.5819 | 4.08E-31 | 4.55E-30 | Infection | Liver |
| Cntn3 | 1.140847 | 1.465178 | 4.854354 | 3.00E-02 | 3.73E-02 | Infection | Liver |
| Cntn4 | 1.977813 | 1.440367 | 13.6415 | 4.20E-04 | 6.18E-04 | Infection | Liver |
| Cntn6 | 1.480035 | 1.310775 | 4.577464 | 3.73E-02 | 4.59E-02 | Infection | Liver |
| Cntnap1 | 1.478532 | 3.832625 | 87.84618 | 1.51E-13 | 3.71E-13 | Infection | Liver |
| Lrp8 | 4.242142 | 3.367201 | 236.4851 | 3.83E-29 | 3.40E-28 | Infection | Liver |
| Cntnap4 | 1.475822 | 1.312654 | 4.729979 | 3.44E-02 | 4.25E-02 | Infection | Liver |
| Cntnap5b | 1.258447 | 1.325946 | 4.605053 | 3.64E-02 | 4.48E-02 | Infection | Liver |
| Cntrob | 0.115039 | 4.479882 | 1.060901 | 3.07E-01 | 3.33E-01 | Infection | Liver |
| Coa7 | -1.24646 | 5.913493 | 204.1738 | 1.85E-21 | 7.66E-21 | Healthy control | Liver |
| Col10a1 | 1.157071 | 1.37776 | 4.37716 | 4.01E-02 | 4.91E-02 | Infection | Liver |
| Cdkn1a | 4.234088 | 6.70783 | 523.6029 | 9.59E-32 | 1.15E-30 | Infection | Liver |
| Col13a1 | -1.45963 | 5.581038 | 118.1494 | 4.34E-16 | 1.22E-15 | Healthy control | Liver |
| Col14a1 | 0.715449 | 9.249266 | 40.17838 | 2.92E-08 | 5.39E-08 | Infection | Liver |
| Cers1 | 4.212812 | 2.752235 | 157.7903 | 9.47E-21 | 3.71E-20 | Infection | Liver |
| Col18a1 | -1.05951 | 12.28978 | 130.3904 | 5.02E-17 | 1.50E-16 | Healthy control | Liver |
| Il1rn | 4.212363 | 5.869076 | 873.1404 | 6.97E-39 | 2.58E-37 | Infection | Liver |
| Ly6g | 4.210266 | 1.964871 | 49.8781 | 3.33E-09 | 6.48E-09 | Infection | Liver |
| Cryaa | 4.20817 | 1.929342 | 83.2593 | 1.56E-12 | 3.59E-12 | Infection | Liver |
| Gal | 4.207213 | 2.128018 | 64.64372 | 6.67E-11 | 1.42E-10 | Infection | Liver |
| Gria3 | 4.195108 | 4.229422 | 531.1454 | 3.25E-39 | 1.30E-37 | Infection | Liver |
| Asxl3 | 4.18706 | 2.370759 | 111.6359 | 1.88E-15 | 5.09E-15 | Infection | Liver |
| Col4a5 | 4.185625 | 5.213057 | 281.2141 | 8.69E-26 | 5.36E-25 | Infection | Liver |
| Col4a3 | 1.346645 | 2.206385 | 18.36422 | 3.89E-05 | 6.12E-05 | Infection | Liver |
| Col4a3bp | -0.11685 | 7.546069 | 6.250228 | 1.50E-02 | 1.94E-02 | Healthy control | Liver |
| 5-Sep | 4.182296 | 4.536652 | 282.2203 | 4.10E-28 | 3.31E-27 | Infection | Liver |
| Slc18a2 | 4.172945 | 3.21338 | 124.5761 | 1.35E-19 | 4.91E-19 | Infection | Liver |
| Htr2b | 4.166347 | 1.937736 | 63.30519 | 5.53E-11 | 1.18E-10 | Infection | Liver |
| Cenpe | 4.165183 | 4.402185 | 382.9743 | 7.06E-33 | 9.55E-32 | Infection | Liver |
| Bhlhe22 | 4.164913 | 4.06411 | 334.07 | 1.80E-32 | 2.31E-31 | Infection | Liver |
| Cdk1 | 4.160587 | 5.638344 | 637.5853 | 5.17E-35 | 1.00E-33 | Infection | Liver |

| | | | | | | | |
|---|---|---|---|---|---|---|---|
| Il10 | 4.14712 | 3.144185 | 198.2072 | 1.04E-25 | 6.31E-25 | Infection | Liver |
| Col9a3 | 1.63352 | 2.321285 | 32.06828 | 1.22E-07 | 2.19E-07 | Infection | Liver |
| Colgalt2 | -0.05796 | 2.472061 | 0.051689 | 8.21E-01 | 8.36E-01 | Healthy control | Liver |
| Colq | 0.7767 | 1.831587 | 4.714981 | 3.17E-02 | 3.93E-02 | Infection | Liver |
| Commd1 | -1.2445 | 8.357256 | 472.142 | 4.51E-31 | 5.00E-30 | Healthy control | Liver |
| Commd10 | 0.101012 | 5.309012 | 1.722854 | 1.94E-01 | 2.17E-01 | Infection | Liver |
| Comt | -1.18676 | 11.17816 | 258.5748 | 4.74E-24 | 2.49E-23 | Healthy control | Liver |
| Cop1 | 0.493122 | 5.831441 | 44.95708 | 6.38E-09 | 1.22E-08 | Infection | Liver |
| Copg1 | 0.288119 | 7.594862 | 42.22877 | 1.45E-08 | 2.74E-08 | Infection | Liver |
| Coprs | -0.70976 | 4.609613 | 53.54135 | 5.63E-10 | 1.14E-09 | Healthy control | Liver |
| Cops2 | -0.42115 | 8.249323 | 51.6659 | 9.12E-10 | 1.83E-09 | Healthy control | Liver |
| Cops5 | -0.6979 | 7.590689 | 155.7653 | 9.94E-19 | 3.35E-18 | Healthy control | Liver |
| Cops8 | -0.9143 | 7.378601 | 335.5495 | 5.24E-27 | 3.65E-26 | Healthy control | Liver |
| Cops9 | -0.49081 | 6.827628 | 69.66206 | 8.50E-12 | 1.89E-11 | Healthy control | Liver |
| Coq8a | -2.19074 | 11.6138 | 551.0525 | 5.71E-33 | 7.81E-32 | Healthy control | Liver |
| Coro1c | 1.144092 | 8.032932 | 287.2453 | 3.15E-25 | 1.84E-24 | Infection | Liver |
| Coro2b | 1.30674 | 5.943493 | 129.7986 | 5.65E-17 | 1.68E-16 | Infection | Liver |
| Il11 | 4.147027 | 2.597342 | 133.1306 | 2.14E-18 | 7.03E-18 | Infection | Liver |
| Cort | 1.844401 | 2.235201 | 48.16554 | 2.07E-10 | 4.30E-10 | Infection | Liver |
| Cox7a1 | -0.56321 | 1.208233 | 1.022802 | 3.23E-01 | 3.50E-01 | Healthy control | Liver |
| Cox8b | 1.81E-16 | 1.160624 | 0 | 1.00E+00 | 1.00E+00 | Infection | Liver |
| Nefl | 4.138649 | 2.571798 | 108.1572 | 3.01E-16 | 8.50E-16 | Infection | Liver |
| Cpeb2 | -0.90103 | 7.304678 | 128.2645 | 7.18E-17 | 2.13E-16 | Healthy control | Liver |
| Cpeb4 | -0.46986 | 9.087923 | 72.8604 | 3.95E-12 | 8.90E-12 | Healthy control | Liver |
| Cplx3 | 0.842845 | 1.320759 | 2.346447 | 1.31E-01 | 1.50E-01 | Infection | Liver |
| Cpn1 | -1.27115 | 10.42703 | 301.6149 | 8.76E-26 | 5.40E-25 | Healthy control | Liver |
| Sycp3 | 4.138237 | 2.109123 | 65.45767 | 5.61E-11 | 1.20E-10 | Infection | Liver |
| Cpox | -1.32735 | 9.323557 | 526.543 | 2.08E-32 | 2.67E-31 | Healthy control | Liver |
| Cps1 | -1.76147 | 13.1133 | 151.3842 | 2.70E-18 | 8.77E-18 | Healthy control | Liver |
| Cpsf6 | -0.03564 | 7.674949 | 0.908261 | 3.44E-01 | 3.70E-01 | Healthy control | Liver |
| Cpt1a | -1.91171 | 9.33574 | 1298.509 | 6.17E-44 | 7.53E-42 | Healthy control | Liver |
| Sgo1 | 4.133682 | 3.90857 | 302.3586 | 9.57E-32 | 1.15E-30 | Infection | Liver |
| Cpt2 | -1.87701 | 8.377717 | 557.4121 | 4.15E-33 | 5.82E-32 | Healthy control | Liver |
| Cptp | -1.1893 | 5.762921 | 303.2782 | 8.51E-26 | 5.26E-25 | Healthy control | Liver |
| Cr1l | 0.565808 | 7.074113 | 100.0836 | 1.16E-14 | 2.99E-14 | Infection | Liver |
| Kcnt2 | 4.132593 | 7.397011 | 1425.466 | 4.79E-45 | 9.60E-43 | Infection | Liver |
| Crabp1 | 1.706249 | 1.244662 | 9.845506 | 3.88E-03 | 5.29E-03 | Infection | Liver |
| Runx1 | 4.129795 | 6.256071 | 586.0203 | 2.36E-33 | 3.40E-32 | Infection | Liver |
| Crb1 | 0.148126 | 1.44626 | 0.083342 | 7.73E-01 | 7.92E-01 | Infection | Liver |
| Crb2 | 0.587288 | 1.483058 | 1.347167 | 2.48E-01 | 2.74E-01 | Infection | Liver |
| Crbn | 0.186571 | 7.2127 | 9.649531 | 2.83E-03 | 3.90E-03 | Infection | Liver |
| Creb1 | 0.442766 | 6.963437 | 75.41846 | 2.18E-12 | 4.97E-12 | Infection | Liver |
| Crebbp | -0.36403 | 7.331834 | 32.03879 | 3.92E-07 | 6.85E-07 | Healthy control | Liver |
| Crebrf | 0.569269 | 7.927024 | 42.70617 | 1.25E-08 | 2.37E-08 | Infection | Liver |
| Creg1 | -0.47023 | 7.882843 | 29.46395 | 9.57E-07 | 1.64E-06 | Healthy control | Liver |
| Creld1 | 0.288591 | 7.269355 | 19.04238 | 4.81E-05 | 7.51E-05 | Infection | Liver |
| Crh | 1.393371 | 1.302652 | 5.676974 | 2.13E-02 | 2.70E-02 | Infection | Liver |
| Crhbp | 1.030968 | 1.302722 | 2.687708 | 1.07E-01 | 1.24E-01 | Infection | Liver |
| Crhr1 | 1.232988 | 1.255168 | 3.928472 | 5.48E-02 | 6.58E-02 | Infection | Liver |
| Crhr2 | -0.08188 | 1.328882 | 0.022074 | 8.82E-01 | 8.93E-01 | Healthy control | Liver |

| | | | | | | | |
|---|---|---|---|---|---|---|---|
| Nrcam | 4.129169 | 2.141882 | 137.6893 | 8.50E-17 | 2.50E-16 | Infection | Liver |
| Crk | -0.28945 | 8.585609 | 64.76092 | 2.83E-11 | 6.12E-11 | Healthy control | Liver |
| Crlf2 | 0.937178 | 5.703469 | 48.08073 | 2.55E-09 | 5.00E-09 | Infection | Liver |
| Crmp1 | 1.476053 | 1.602734 | 12.54741 | 5.76E-04 | 8.36E-04 | Infection | Liver |
| Crocc | 0.872939 | 4.621083 | 47.52454 | 3.10E-09 | 6.06E-09 | Infection | Liver |
| Crtc1 | -0.61142 | 1.242354 | 1.157662 | 2.89E-01 | 3.15E-01 | Healthy control | Liver |
| Crtc2 | -0.07841 | 6.31674 | 1.72945 | 1.93E-01 | 2.16E-01 | Healthy control | Liver |
| Crx | 0.645688 | 1.302222 | 1.195954 | 2.79E-01 | 3.05E-01 | Infection | Liver |
| Cd109 | 4.126904 | 2.19892 | 149.1225 | 6.38E-18 | 2.02E-17 | Infection | Liver |
| Cry1 | -0.83984 | 5.132812 | 94.21311 | 3.97E-14 | 1.00E-13 | Healthy control | Liver |
| Cry2 | -0.84655 | 7.028081 | 147.6426 | 3.33E-18 | 1.07E-17 | Healthy control | Liver |
| Tnfrsf18 | 4.121938 | 2.671307 | 204.2196 | 6.77E-24 | 3.49E-23 | Infection | Liver |
| Cryab | 1.026896 | 6.246116 | 55.29405 | 3.54E-10 | 7.25E-10 | Infection | Liver |
| Cryba2 | -0.19714 | 1.19767 | 0.142101 | 7.11E-01 | 7.31E-01 | Healthy control | Liver |
| Cryba4 | 1.242021 | 1.807494 | 13.08038 | 4.23E-04 | 6.21E-04 | Infection | Liver |
| Il7r | 4.119081 | 4.66572 | 463.1155 | 4.40E-34 | 6.98E-33 | Infection | Liver |
| Crygb | 1.331627 | 1.221612 | 5.158801 | 3.23E-02 | 4.00E-02 | Infection | Liver |
| Crygc | 0.588737 | 1.303299 | 0.871232 | 3.55E-01 | 3.81E-01 | Infection | Liver |
| Crygd | 0.536805 | 1.210192 | 0.848671 | 3.67E-01 | 3.94E-01 | Infection | Liver |
| Flt3 | 4.11904 | 4.08042 | 396.6885 | 2.55E-35 | 5.26E-34 | Infection | Liver |
| Crygf | 1.252507 | 1.220756 | 5.381896 | 2.83E-02 | 3.53E-02 | Infection | Liver |
| Crygs | 0.054224 | 1.337713 | 0.009965 | 9.21E-01 | 9.29E-01 | Infection | Liver |
| Crym | -1.88259 | 6.53694 | 364.6111 | 6.06E-28 | 4.81E-27 | Healthy control | Liver |
| Cs | -0.20547 | 9.933562 | 16.21363 | 1.54E-04 | 2.32E-04 | Healthy control | Liver |
| Csde1 | -0.84025 | 9.330927 | 394.508 | 6.51E-29 | 5.65E-28 | Healthy control | Liver |
| Cse1l | -0.57213 | 5.858934 | 58.98495 | 1.26E-10 | 2.64E-10 | Healthy control | Liver |
| Csf1 | 1.077378 | 7.074632 | 307.3977 | 5.36E-26 | 3.40E-25 | Infection | Liver |
| Csf1r | 0.778628 | 8.270928 | 93.21297 | 4.56E-14 | 1.15E-13 | Infection | Liver |
| Csf2 | 1.137368 | 1.437193 | 4.263974 | 4.20E-02 | 5.14E-02 | Infection | Liver |
| Cd27 | 4.114958 | 2.261636 | 84.54266 | 4.28E-13 | 1.02E-12 | Infection | Liver |
| F2rl1 | 4.10994 | 2.509621 | 120.678 | 4.71E-17 | 1.41E-16 | Infection | Liver |
| Csf3 | 0.265671 | 1.322064 | 0.233477 | 6.31E-01 | 6.54E-01 | Infection | Liver |
| Vax2 | 4.102246 | 1.877991 | 105.0835 | 1.34E-14 | 3.45E-14 | Infection | Liver |
| Csk | 0.096677 | 6.750776 | 1.900848 | 1.73E-01 | 1.94E-01 | Infection | Liver |
| Csnk1a1 | 0.051241 | 8.011472 | 1.422814 | 2.37E-01 | 2.63E-01 | Infection | Liver |
| Csnk1d | 0.21519 | 9.147174 | 55.16613 | 3.47E-10 | 7.11E-10 | Infection | Liver |
| Csnk1e | 1.038644 | 6.379964 | 211.4186 | 7.42E-22 | 3.18E-21 | Infection | Liver |
| Csnk2a1 | -0.6731 | 8.048828 | 170.6207 | 1.23E-19 | 4.47E-19 | Healthy control | Liver |
| Csnk2a2 | -0.99328 | 7.036438 | 322.1087 | 1.57E-26 | 1.05E-25 | Healthy control | Liver |
| Map1b | 4.086827 | 4.761843 | 213.7477 | 1.84E-23 | 8.97E-23 | Infection | Liver |
| Cspg5 | 1.367613 | 1.706434 | 13.02603 | 4.40E-04 | 6.45E-04 | Infection | Liver |
| Ska1 | 4.075857 | 2.925422 | 176.6286 | 1.10E-23 | 5.53E-23 | Infection | Liver |
| Jaml | 4.062677 | 4.997565 | 620.2753 | 3.04E-36 | 7.42E-35 | Infection | Liver |
| Ctbp1 | -0.91915 | 6.946715 | 104.8615 | 4.61E-15 | 1.22E-14 | Healthy control | Liver |
| Ctbp2 | 1.567735 | 5.191709 | 191.8898 | 8.55E-21 | 3.36E-20 | Infection | Liver |
| Ctcf | 0.182057 | 7.748289 | 14.59827 | 3.06E-04 | 4.53E-04 | Infection | Liver |
| Ctdsp1 | -1.11315 | 9.814788 | 676.5097 | 1.57E-35 | 3.31E-34 | Healthy control | Liver |
| Ctdspl2 | 0.68696 | 5.839924 | 108.1002 | 2.53E-15 | 6.78E-15 | Infection | Liver |
| Cth | -2.54019 | 10.79051 | 279.2655 | 2.93E-24 | 1.57E-23 | Healthy control | Liver |
| Il2ra | 4.042193 | 3.21881 | 229.6802 | 1.80E-28 | 1.50E-27 | Infection | Liver |

| | | | | | | | |
|---|---|---|---|---|---|---|---|
| Ctnna1 | 0.600405 | 8.429349 | 234.8243 | 5.43E-23 | 2.54E-22 | Infection | Liver |
| Ctnna3 | 1.998631 | 1.529235 | 16.46324 | 1.08E-04 | 1.65E-04 | Infection | Liver |
| Cd209a | 4.035284 | 3.602497 | 150.6996 | 7.56E-22 | 3.24E-21 | Infection | Liver |
| Ctnnb1 | -0.67023 | 9.999957 | 88.32625 | 1.26E-13 | 3.09E-13 | Healthy control | Liver |
| Ctnnbip1 | 0.139016 | 5.478713 | 1.238682 | 2.70E-01 | 2.96E-01 | Infection | Liver |
| Ctnnd1 | 0.267046 | 9.90314 | 42.08786 | 1.51E-08 | 2.85E-08 | Infection | Liver |
| Ctr9 | 0.228544 | 6.934488 | 19.17902 | 4.55E-05 | 7.12E-05 | Infection | Liver |
| Sema4d | 4.031866 | 6.132038 | 1042.062 | 9.70E-41 | 5.65E-39 | Infection | Liver |
| Ctsl | -0.61275 | 10.06831 | 54.66649 | 3.97E-10 | 8.12E-10 | Healthy control | Liver |
| Ctsq | 1.809857 | 1.320257 | 9.834594 | 2.74E-03 | 3.78E-03 | Infection | Liver |
| Cttn | 0.149205 | 8.203876 | 9.995453 | 2.41E-03 | 3.34E-03 | Infection | Liver |
| Cttnbp2 | 1.964397 | 2.111179 | 35.58425 | 2.39E-08 | 4.44E-08 | Infection | Liver |
| Grhl2 | 4.027988 | 1.886947 | 61.21818 | 1.12E-10 | 2.35E-10 | Infection | Liver |
| Cul1 | -0.98276 | 7.472005 | 341.8263 | 3.19E-27 | 2.28E-26 | Healthy control | Liver |
| Cul3 | -0.7255 | 8.566801 | 311.5259 | 3.75E-26 | 2.41E-25 | Healthy control | Liver |
| Cul4a | -1.30427 | 8.114212 | 988.766 | 2.22E-40 | 1.16E-38 | Healthy control | Liver |
| Cux1 | 0.571912 | 6.341808 | 45.90579 | 4.80E-09 | 9.29E-09 | Infection | Liver |
| Cux2 | -1.44721 | 4.856944 | 61.3472 | 6.70E-11 | 1.42E-10 | Healthy control | Liver |
| Cwc27 | -0.27796 | 5.845895 | 19.15885 | 4.60E-05 | 7.18E-05 | Healthy control | Liver |
| Osm | 4.019657 | 2.645556 | 151.7571 | 2.27E-20 | 8.67E-20 | Infection | Liver |
| Cx3cr1 | -0.16337 | 4.583564 | 0.564412 | 4.55E-01 | 4.83E-01 | Healthy control | Liver |
| Cxadr | -0.34423 | 9.21945 | 12.97923 | 6.19E-04 | 8.97E-04 | Healthy control | Liver |
| Arnt2 | 4.014312 | 2.49538 | 144.3844 | 2.58E-19 | 9.12E-19 | Infection | Liver |
| Cav3 | 4.011363 | 1.686239 | 58.89243 | 2.13E-10 | 4.41E-10 | Infection | Liver |
| Cxcl12 | -0.77585 | 10.1631 | 150.3311 | 2.21E-18 | 7.23E-18 | Healthy control | Liver |
| Cxcl13 | 1.230478 | 4.775701 | 16.8642 | 1.26E-04 | 1.91E-04 | Infection | Liver |
| Gata3 | 4.010124 | 4.271978 | 491.3701 | 1.04E-36 | 2.76E-35 | Infection | Liver |
| Cxcl16 | 1.44445 | 7.946273 | 251.3727 | 9.76E-24 | 4.95E-23 | Infection | Liver |
| Cxcl2 | 1.491039 | 1.329931 | 6.842815 | 1.15E-02 | 1.51E-02 | Infection | Liver |
| Cxcl3 | 1.248614 | 1.559053 | 6.139582 | 1.47E-02 | 1.90E-02 | Infection | Liver |
| A2m | 4.004487 | 2.637233 | 119.101 | 1.07E-17 | 3.35E-17 | Infection | Liver |
| Pknox2 | 4.003872 | 5.03307 | 311.7515 | 2.32E-27 | 1.70E-26 | Infection | Liver |
| Cxcr1 | 1.348792 | 1.396251 | 5.065743 | 2.76E-02 | 3.44E-02 | Infection | Liver |
| Rgcc | 3.999716 | 2.693013 | 135.1037 | 2.08E-19 | 7.40E-19 | Infection | Liver |
| Coro6 | 3.999579 | 3.705517 | 205.5258 | 5.16E-26 | 3.28E-25 | Infection | Liver |
| Orc1 | 3.980814 | 2.894526 | 182.8838 | 3.35E-24 | 1.79E-23 | Infection | Liver |
| Adgrg5 | 3.976056 | 3.752286 | 370.0014 | 1.39E-35 | 2.99E-34 | Infection | Liver |
| Cxcr6 | 1.600241 | 4.092137 | 78.50727 | 1.20E-12 | 2.78E-12 | Infection | Liver |
| Cxxc1 | -0.06175 | 7.281335 | 1.479863 | 2.28E-01 | 2.53E-01 | Healthy control | Liver |
| Cxxc5 | -1.16176 | 8.273037 | 436.0618 | 4.13E-30 | 4.15E-29 | Healthy control | Liver |
| Cftr | 3.972529 | 1.855035 | 57.96856 | 2.34E-10 | 4.83E-10 | Infection | Liver |
| Cyb5a | -1.36799 | 12.57902 | 198.3748 | 3.44E-21 | 1.39E-20 | Healthy control | Liver |
| Cyb5r3 | -0.8906 | 10.47975 | 188.4195 | 1.18E-20 | 4.61E-20 | Healthy control | Liver |
| Cybb | 1.690133 | 7.229637 | 290.7218 | 2.32E-25 | 1.36E-24 | Infection | Liver |
| Cybc1 | 0.787564 | 6.093349 | 98.40435 | 1.62E-14 | 4.14E-14 | Infection | Liver |
| Fbn1 | 3.970653 | 8.647591 | 541.0037 | 1.52E-31 | 1.81E-30 | Infection | Liver |
| Cyfip1 | -0.20162 | 7.55031 | 22.07679 | 1.45E-05 | 2.34E-05 | Healthy control | Liver |
| Cyfip2 | 1.366026 | 6.650698 | 275.0376 | 9.84E-25 | 5.52E-24 | Infection | Liver |
| Top2a | 3.957019 | 6.306177 | 507.3259 | 1.83E-31 | 2.15E-30 | Infection | Liver |
| Cyld | 0.753169 | 8.201431 | 301.4103 | 8.94E-26 | 5.50E-25 | Infection | Liver |

| | | | | | | | |
|---|---|---|---|---|---|---|---|
| Tigit | 3.945367 | 2.857225 | 178.2793 | 1.12E-23 | 5.61E-23 | Infection | Liver |
| Cyp11b2 | 1.681459 | 1.254457 | 9.450649 | 3.91E-03 | 5.32E-03 | Infection | Liver |
| Cyp17a1 | -6.12063 | 7.787 | 521.066 | 1.56E-32 | 2.03E-31 | Healthy control | Liver |
| Cyp19a1 | 1.173435 | 1.219495 | 5.681317 | 2.47E-02 | 3.11E-02 | Infection | Liver |
| Cyp1a1 | -0.7462 | 1.494266 | 2.500746 | 1.17E-01 | 1.35E-01 | Healthy control | Liver |
| Cyp1a2 | -2.40888 | 11.12725 | 413.8319 | 2.46E-29 | 2.25E-28 | Healthy control | Liver |
| Trhr | 3.925574 | 1.887168 | 51.79853 | 1.37E-09 | 2.72E-09 | Infection | Liver |
| Cyp21a1 | 0.871029 | 2.187707 | 11.66737 | 8.96E-04 | 1.29E-03 | Infection | Liver |
| Cyp24a1 | 1.023699 | 1.261688 | 3.656189 | 6.30E-02 | 7.51E-02 | Infection | Liver |
| Cyp26a1 | -2.01521 | 5.835464 | 305.4766 | 7.73E-26 | 4.79E-25 | Healthy control | Liver |
| Cyp26b1 | -0.99818 | 4.28129 | 15.49129 | 2.07E-04 | 3.11E-04 | Healthy control | Liver |
| Cyp27a1 | -1.4162 | 10.29999 | 276.0831 | 8.80E-25 | 4.96E-24 | Healthy control | Liver |
| Lhx9 | 3.922124 | 2.103979 | 42.24688 | 1.51E-08 | 2.85E-08 | Infection | Liver |
| Cyp2a5 | -2.58191 | 10.23706 | 248.5015 | 6.87E-23 | 3.20E-22 | Healthy control | Liver |
| Cyp2c29 | -2.49889 | 11.61094 | 239.3038 | 1.40E-22 | 6.37E-22 | Healthy control | Liver |
| Cyp2e1 | -2.4897 | 13.36758 | 921.62 | 1.79E-39 | 7.45E-38 | Healthy control | Liver |
| Cyp2g1 | 0.884176 | 1.274922 | 2.39046 | 1.29E-01 | 1.48E-01 | Infection | Liver |
| Cyp2r1 | -1.06718 | 4.077622 | 58.90868 | 1.16E-10 | 2.42E-10 | Healthy control | Liver |
| Cyp3a13 | -0.29395 | 7.43572 | 6.510438 | 1.32E-02 | 1.71E-02 | Healthy control | Liver |
| Cyp7a1 | -1.75793 | 8.348694 | 62.43064 | 8.81E-11 | 1.86E-10 | Healthy control | Liver |
| Cyp7b1 | -1.16392 | 7.270688 | 163.6706 | 3.22E-19 | 1.13E-18 | Healthy control | Liver |
| Cyp8b1 | -1.30698 | 8.83452 | 67.56539 | 2.33E-11 | 5.06E-11 | Healthy control | Liver |
| Cyren | 0.525638 | 3.503842 | 15.79169 | 1.83E-04 | 2.76E-04 | Infection | Liver |
| Plaur | 3.915526 | 5.211017 | 325.4411 | 4.64E-27 | 3.25E-26 | Infection | Liver |
| D1Ertd622e | 0.044074 | 4.611603 | 0.139688 | 7.10E-01 | 7.31E-01 | Infection | Liver |
| D1Pas1 | 1.492872 | 1.288242 | 5.57922 | 2.23E-02 | 2.82E-02 | Infection | Liver |
| Fndc5 | 3.91095 | 3.562401 | 138.6566 | 1.01E-20 | 3.94E-20 | Infection | Liver |
| Dab2 | 0.375342 | 6.540049 | 10.47469 | 1.93E-03 | 2.69E-03 | Infection | Liver |
| Dach1 | 0.833772 | 3.358303 | 22.32575 | 1.29E-05 | 2.09E-05 | Infection | Liver |
| Dact2 | -0.71794 | 7.00496 | 91.88751 | 6.00E-14 | 1.50E-13 | Healthy control | Liver |
| Dag1 | -0.92762 | 7.839059 | 142.7283 | 7.02E-18 | 2.22E-17 | Healthy control | Liver |
| Dagla | 1.87356 | 3.769915 | 102.2062 | 6.32E-15 | 1.66E-14 | Infection | Liver |
| Daglb | -0.64848 | 6.226615 | 45.27088 | 5.81E-09 | 1.12E-08 | Healthy control | Liver |
| Dao | 1.464648 | 1.327301 | 5.337245 | 2.50E-02 | 3.15E-02 | Infection | Liver |
| Dapk1 | -0.25059 | 8.566166 | 26.40514 | 2.86E-06 | 4.78E-06 | Healthy control | Liver |
| Daxx | 1.430635 | 6.804999 | 389.2993 | 9.50E-29 | 8.17E-28 | Infection | Liver |
| Ppp2r2b | 3.909056 | 1.937609 | 68.23148 | 1.79E-11 | 3.91E-11 | Infection | Liver |
| Dbh | 1.68866 | 1.250318 | 12.76137 | 9.73E-04 | 1.39E-03 | Infection | Liver |
| Cd44 | 3.908367 | 6.602294 | 1930.993 | 6.50E-49 | 4.07E-46 | Infection | Liver |
| Dbnl | 0.194591 | 7.851021 | 13.33561 | 5.29E-04 | 7.72E-04 | Infection | Liver |
| Dbp | -0.31849 | 5.712056 | 3.201019 | 7.85E-02 | 9.24E-02 | Healthy control | Liver |
| Dbx1 | 0.184385 | 1.246633 | 0.091237 | 7.64E-01 | 7.83E-01 | Infection | Liver |
| Sorl1 | 3.90552 | 5.211335 | 798.0943 | 2.55E-38 | 8.40E-37 | Infection | Liver |
| Dcaf1 | -1.02489 | 6.115721 | 174.321 | 7.72E-20 | 2.85E-19 | Healthy control | Liver |
| Dcaf11 | -1.97546 | 8.143266 | 614.324 | 2.57E-34 | 4.24E-33 | Healthy control | Liver |
| Dcaf7 | -0.60398 | 7.687314 | 155.9286 | 9.70E-19 | 3.27E-18 | Healthy control | Liver |
| Dcc | 0.424419 | 1.37794 | 0.528588 | 4.69E-01 | 4.97E-01 | Infection | Liver |
| Dct | -2.60899 | 4.227529 | 143.9491 | 5.29E-19 | 1.81E-18 | Healthy control | Liver |
| Dctn1 | 0.487136 | 7.736511 | 112.0191 | 1.21E-15 | 3.32E-15 | Infection | Liver |
| Dctn2 | -0.5791 | 8.976133 | 204.3152 | 1.69E-21 | 7.00E-21 | Healthy control | Liver |

| | | | | | | | |
|---|---|---|---|---|---|---|---|
| Dctn3 | -0.22707 | 5.900173 | 11.00103 | 1.51E-03 | 2.13E-03 | Healthy control | Liver |
| Dctn4 | 0.448155 | 5.824018 | 45.39632 | 5.60E-09 | 1.08E-08 | Infection | Liver |
| Dcx | 1.869038 | 1.849425 | 24.08974 | 2.96E-06 | 4.93E-06 | Infection | Liver |
| Ddah1 | -2.57966 | 8.458216 | 712.6896 | 3.50E-36 | 8.39E-35 | Healthy control | Liver |
| Ddb1 | 0.194174 | 8.877232 | 25.57883 | 3.88E-06 | 6.44E-06 | Infection | Liver |
| Ddc | -0.87498 | 7.007197 | 46.04214 | 4.95E-09 | 9.57E-09 | Healthy control | Liver |
| Ddit3 | 0.867459 | 6.165028 | 151.2151 | 1.96E-18 | 6.47E-18 | Infection | Liver |
| Ddit4 | 1.251774 | 7.015121 | 152.4869 | 1.61E-18 | 5.35E-18 | Infection | Liver |
| Cd40 | 3.904368 | 4.562851 | 655.0908 | 1.76E-38 | 6.08E-37 | Infection | Liver |
| Ddx19b | 0.35722 | 6.336415 | 17.13088 | 1.05E-04 | 1.61E-04 | Infection | Liver |
| Ddx21 | -0.47298 | 6.765124 | 54.94379 | 3.69E-10 | 7.56E-10 | Healthy control | Liver |
| Ddx4 | 1.857112 | 1.519109 | 10.26063 | 1.93E-03 | 2.70E-03 | Infection | Liver |
| Ddx41 | -0.72368 | 6.100167 | 115.3227 | 6.81E-16 | 1.89E-15 | Healthy control | Liver |
| Ddx42 | -0.67463 | 7.001768 | 163.3174 | 3.39E-19 | 1.18E-18 | Healthy control | Liver |
| Ddx56 | -0.40911 | 7.304326 | 80.8387 | 6.38E-13 | 1.51E-12 | Healthy control | Liver |
| Ddx58 | 0.393322 | 7.420044 | 29.59397 | 9.14E-07 | 1.57E-06 | Infection | Liver |
| Ddx6 | 0.92338 | 9.407051 | 333.4266 | 6.15E-27 | 4.27E-26 | Infection | Liver |
| Decr1 | -1.97881 | 8.402869 | 565.2414 | 2.79E-33 | 3.99E-32 | Healthy control | Liver |
| Mzb1 | 3.895416 | 5.339882 | 118.912 | 7.38E-16 | 2.04E-15 | Infection | Liver |
| Dio2 | 3.890622 | 1.811264 | 67.56019 | 2.97E-11 | 6.40E-11 | Infection | Liver |
| Dek | 0.535025 | 8.107114 | 92.17761 | 5.64E-14 | 1.41E-13 | Infection | Liver |
| Dennd1a | -0.11606 | 6.472468 | 4.157676 | 4.56E-02 | 5.55E-02 | Healthy control | Liver |
| Derl2 | 0.049806 | 8.203061 | 0.41548 | 5.22E-01 | 5.49E-01 | Infection | Liver |
| Des | 0.635109 | 5.800674 | 18.47087 | 6.12E-05 | 9.48E-05 | Infection | Liver |
| Dffb | 0.131232 | 5.359708 | 2.490891 | 1.20E-01 | 1.38E-01 | Infection | Liver |
| Dgat1 | -0.55484 | 6.635776 | 76.55028 | 1.68E-12 | 3.87E-12 | Healthy control | Liver |
| Dgcr2 | -1.37591 | 8.740488 | 918.8871 | 1.96E-39 | 8.04E-38 | Healthy control | Liver |
| Dgcr8 | 0.334484 | 6.020187 | 28.90391 | 1.17E-06 | 1.99E-06 | Infection | Liver |
| Dgka | 1.598465 | 4.877457 | 245.0992 | 2.30E-23 | 1.12E-22 | Infection | Liver |
| Gli2 | 3.86569 | 2.455861 | 117.5209 | 2.66E-17 | 8.11E-17 | Infection | Liver |
| Dgkq | -0.02574 | 7.330597 | 0.236164 | 6.29E-01 | 6.53E-01 | Healthy control | Liver |
| Dglucy | -1.48958 | 7.945382 | 356.336 | 1.04E-27 | 8.04E-27 | Healthy control | Liver |
| Dhfr | -1.98222 | 7.734762 | 622.596 | 1.77E-34 | 3.02E-33 | Healthy control | Liver |
| Dhh | 1.252457 | 2.956635 | 44.29743 | 4.72E-09 | 9.14E-09 | Infection | Liver |
| Dhodh | -0.70976 | 4.541253 | 60.50974 | 8.80E-11 | 1.86E-10 | Healthy control | Liver |
| Dhx33 | 0.265099 | 7.009922 | 13.65656 | 4.60E-04 | 6.73E-04 | Infection | Liver |
| Dhx58 | 0.472484 | 6.652931 | 20.41402 | 2.78E-05 | 4.41E-05 | Infection | Liver |
| Diablo | -0.1156 | 6.294984 | 3.709345 | 5.86E-02 | 7.01E-02 | Healthy control | Liver |
| Diaph1 | 0.013933 | 8.801971 | 0.069304 | 7.93E-01 | 8.10E-01 | Infection | Liver |
| Chrm2 | 3.865477 | 2.767725 | 116.2708 | 3.12E-18 | 1.01E-17 | Infection | Liver |
| Dicer1 | -0.43478 | 7.33536 | 98.25017 | 1.66E-14 | 4.24E-14 | Healthy control | Liver |
| Dio1 | -2.13541 | 5.759324 | 156.7684 | 1.05E-18 | 3.53E-18 | Healthy control | Liver |
| Gap43 | 3.847399 | 2.398401 | 112.0744 | 1.60E-16 | 4.63E-16 | Infection | Liver |
| Dio3 | 0.327022 | 1.250964 | 0.314344 | 5.78E-01 | 6.04E-01 | Infection | Liver |
| Dip2a | 0.912904 | 5.620446 | 107.5377 | 2.83E-15 | 7.56E-15 | Infection | Liver |
| Il21r | 3.845389 | 4.022464 | 324.9575 | 3.96E-31 | 4.42E-30 | Infection | Liver |
| Chrnb1 | 3.840959 | 3.631989 | 286.8381 | 7.97E-31 | 8.56E-30 | Infection | Liver |
| Disp2 | 0.290847 | 1.87056 | 0.6867 | 4.09E-01 | 4.37E-01 | Infection | Liver |
| Dixdc1 | -0.08579 | 4.719352 | 0.511257 | 4.77E-01 | 5.05E-01 | Healthy control | Liver |
| Il4 | 3.840026 | 1.913339 | 97.05501 | 4.50E-14 | 1.13E-13 | Infection | Liver |

| | | | | | | | |
|---|---|---|---|---|---|---|---|
| Msx1 | 3.83898 | 4.260654 | 251.6963 | 1.20E-26 | 8.13E-26 | Infection | Liver |
| Rapsn | 3.837592 | 2.12768 | 86.34847 | 2.09E-13 | 5.09E-13 | Infection | Liver |
| Dkk4 | 1.045332 | 1.279553 | 3.268217 | 7.71E-02 | 9.09E-02 | Infection | Liver |
| Dlg1 | -0.03206 | 8.623122 | 0.522385 | 4.72E-01 | 5.00E-01 | Healthy control | Liver |
| Dlg3 | -0.52288 | 5.791112 | 41.54231 | 1.80E-08 | 3.38E-08 | Healthy control | Liver |
| Cd69 | 3.836927 | 2.886703 | 224.504 | 3.94E-27 | 2.79E-26 | Infection | Liver |
| Dlg5 | 1.193676 | 6.135418 | 168.3158 | 1.71E-19 | 6.12E-19 | Infection | Liver |
| Dlgap4 | 0.351617 | 6.616112 | 23.12977 | 9.72E-06 | 1.58E-05 | Infection | Liver |
| Dlk1 | -0.2883 | 1.19656 | 0.299444 | 5.92E-01 | 6.17E-01 | Healthy control | Liver |
| Dll1 | 0.060937 | 5.242769 | 0.32181 | 5.73E-01 | 5.99E-01 | Infection | Liver |
| Dll3 | 1.920687 | 1.395829 | 11.49497 | 1.20E-03 | 1.71E-03 | Infection | Liver |
| Dll4 | 0.087227 | 5.777249 | 0.668933 | 4.16E-01 | 4.44E-01 | Infection | Liver |
| Dlx1 | 0.93682 | 1.363033 | 2.812675 | 9.77E-02 | 1.14E-01 | Infection | Liver |
| Dlx2 | 1.728382 | 1.399809 | 8.123374 | 5.73E-03 | 7.71E-03 | Infection | Liver |
| Dlx3 | 1.067111 | 1.234282 | 2.965841 | 9.71E-02 | 1.13E-01 | Infection | Liver |
| Dlx4 | 0.611212 | 1.243545 | 0.96778 | 3.32E-01 | 3.58E-01 | Infection | Liver |
| Ccr7 | 3.835721 | 3.553721 | 198.0263 | 6.48E-26 | 4.06E-25 | Infection | Liver |
| Dlx6 | 1.404051 | 1.432441 | 5.19799 | 2.54E-02 | 3.19E-02 | Infection | Liver |
| Dmap1 | -0.14933 | 6.301102 | 4.749931 | 3.30E-02 | 4.09E-02 | Healthy control | Liver |
| Dmbx1 | 1.766164 | 1.439469 | 10.18362 | 2.03E-03 | 2.83E-03 | Infection | Liver |
| Dmc1 | 1.647905 | 1.43339 | 7.967202 | 6.24E-03 | 8.36E-03 | Infection | Liver |
| Dmd | -1.20718 | 5.705103 | 183.3223 | 2.47E-20 | 9.42E-20 | Healthy control | Liver |
| Rad54l | 3.830896 | 4.054671 | 426.9489 | 6.54E-35 | 1.23E-33 | Infection | Liver |
| Dmrt1 | 1.363149 | 1.265927 | 4.940523 | 3.20E-02 | 3.96E-02 | Infection | Liver |
| Dmrt2 | 0.647256 | 1.661465 | 2.344454 | 1.28E-01 | 1.47E-01 | Infection | Liver |
| Dmrta1 | -1.20308 | 1.898424 | 7.472533 | 7.32E-03 | 9.74E-03 | Healthy control | Liver |
| Dmrta2 | 0.28225 | 1.741039 | 0.75824 | 3.85E-01 | 4.13E-01 | Infection | Liver |
| Dnah11 | -0.6084 | 1.937259 | 3.839403 | 5.26E-02 | 6.34E-02 | Healthy control | Liver |
| Dnah12 | 1.844475 | 1.418098 | 10.9164 | 1.48E-03 | 2.09E-03 | Infection | Liver |
| Dnah5 | -0.0804 | 1.731693 | 0.047542 | 8.28E-01 | 8.42E-01 | Healthy control | Liver |
| Dnajb6 | -0.30958 | 7.276521 | 36.91399 | 7.75E-08 | 1.41E-07 | Healthy control | Liver |
| Dnajc5 | 0.301665 | 7.778421 | 45.34517 | 5.66E-09 | 1.09E-08 | Infection | Liver |
| Dnajc7 | 0.008351 | 7.646408 | 0.050796 | 8.22E-01 | 8.38E-01 | Infection | Liver |
| Dner | 1.11577 | 1.288621 | 3.048785 | 8.71E-02 | 1.02E-01 | Infection | Liver |
| Sgcd | 3.826993 | 4.163414 | 277.4868 | 4.69E-28 | 3.77E-27 | Infection | Liver |
| Dnm1l | -1.16828 | 7.344096 | 341.565 | 3.27E-27 | 2.33E-26 | Healthy control | Liver |
| Dnm2 | -0.02469 | 8.776133 | 0.556075 | 4.59E-01 | 4.86E-01 | Healthy control | Liver |
| Dnmt1 | 0.964534 | 7.071001 | 196.3938 | 4.41E-21 | 1.77E-20 | Infection | Liver |
| Dnmt3a | 0.940752 | 6.536102 | 249.1452 | 1.24E-23 | 6.15E-23 | Infection | Liver |
| Dnmt3b | 0.767784 | 4.849034 | 55.38641 | 3.38E-10 | 6.94E-10 | Infection | Liver |
| Cysltr1 | 3.82272 | 5.586229 | 709.0907 | 3.62E-36 | 8.55E-35 | Infection | Liver |
| Dnttip1 | -0.32865 | 6.916757 | 45.66898 | 5.15E-09 | 9.92E-09 | Healthy control | Liver |
| Dnttip2 | -0.88207 | 5.996772 | 136.2392 | 2.01E-17 | 6.18E-17 | Healthy control | Liver |
| Doc2a | 0.894725 | 1.374421 | 2.733701 | 1.02E-01 | 1.19E-01 | Infection | Liver |
| Btla | 3.814329 | 3.492286 | 383.0998 | 1.16E-36 | 3.04E-35 | Infection | Liver |
| Dock3 | 1.649066 | 1.30439 | 6.735187 | 1.25E-02 | 1.62E-02 | Infection | Liver |
| Dock5 | -0.11971 | 6.356541 | 4.099855 | 4.71E-02 | 5.72E-02 | Healthy control | Liver |
| Dock7 | 1.551659 | 6.734522 | 487.7027 | 1.85E-31 | 2.17E-30 | Infection | Liver |
| Dock8 | -0.33815 | 7.437881 | 35.61697 | 1.18E-07 | 2.13E-07 | Healthy control | Liver |
| Dot1l | 0.630842 | 6.399762 | 124.4663 | 1.36E-16 | 3.97E-16 | Infection | Liver |

| | | | | | | | |
|---|---|---|---|---|---|---|---|
| Dpf2 | -0.37086 | 7.605852 | 65.85639 | 2.16E-11 | 4.69E-11 | Healthy control | Liver |
| Dpf3 | -0.95007 | 5.385502 | 55.74532 | 3.05E-10 | 6.27E-10 | Healthy control | Liver |
| Dpm1 | -0.96788 | 7.992733 | 522.5662 | 2.59E-32 | 3.29E-31 | Healthy control | Liver |
| Itgae | 3.812148 | 3.792116 | 260.6299 | 1.06E-28 | 9.01E-28 | Infection | Liver |
| Dpp4 | -1.29152 | 7.968837 | 250.5534 | 1.06E-23 | 5.37E-23 | Healthy control | Liver |
| Dpp9 | -1.47733 | 8.177791 | 726.0964 | 2.02E-36 | 5.10E-35 | Healthy control | Liver |
| Samd14 | 3.806409 | 4.698128 | 329.8349 | 2.50E-28 | 2.05E-27 | Infection | Liver |
| Dpy30 | -1.54472 | 8.199724 | 1073.626 | 1.90E-41 | 1.32E-39 | Healthy control | Liver |
| Dpyd | -1.46571 | 10.26113 | 155.2364 | 1.47E-18 | 4.91E-18 | Healthy control | Liver |
| Dpysl2 | -0.38939 | 6.424633 | 10.50429 | 1.90E-03 | 2.66E-03 | Healthy control | Liver |
| Il12b | 3.790232 | 1.894438 | 60.42842 | 9.72E-11 | 2.05E-10 | Infection | Liver |
| Ptger4 | 3.775396 | 3.93034 | 246.9637 | 2.31E-27 | 1.70E-26 | Infection | Liver |
| Dpysl5 | 1.346762 | 1.306397 | 5.396733 | 2.39E-02 | 3.02E-02 | Infection | Liver |
| Lif | 3.775093 | 4.174564 | 238.4787 | 8.57E-26 | 5.29E-25 | Infection | Liver |
| Drd1 | -0.06411 | 1.182246 | 0.01769 | 8.96E-01 | 9.06E-01 | Healthy control | Liver |
| Drd2 | 0.596622 | 1.242452 | 1.03766 | 3.15E-01 | 3.42E-01 | Infection | Liver |
| Drd3 | 1.388133 | 1.276254 | 5.566871 | 2.26E-02 | 2.86E-02 | Infection | Liver |
| Il2rg | 3.773672 | 6.85329 | 1550.131 | 4.23E-46 | 1.18E-43 | Infection | Liver |
| Drd5 | 1.069799 | 1.298264 | 2.539337 | 1.18E-01 | 1.36E-01 | Infection | Liver |
| Drosha | 0.042582 | 6.126168 | 0.248597 | 6.20E-01 | 6.44E-01 | Infection | Liver |
| Dsc1 | 1.477217 | 1.464258 | 7.087039 | 9.19E-03 | 1.21E-02 | Infection | Liver |
| Dsc3 | 1.148051 | 1.332478 | 3.576618 | 6.34E-02 | 7.55E-02 | Infection | Liver |
| Dscam | 1.903535 | 1.28611 | 13.27837 | 6.96E-04 | 1.00E-03 | Infection | Liver |
| Dscaml1 | 1.696096 | 1.349987 | 9.520684 | 3.03E-03 | 4.16E-03 | Infection | Liver |
| Sdk1 | 3.767379 | 2.412698 | 110.3778 | 1.06E-16 | 3.11E-16 | Infection | Liver |
| Dsg1b | 0.179645 | 4.509598 | 1.4913 | 2.27E-01 | 2.51E-01 | Infection | Liver |
| Dsg2 | -1.09037 | 8.619828 | 152.0979 | 1.70E-18 | 5.62E-18 | Healthy control | Liver |
| Dsp | -0.74345 | 8.119174 | 113.1824 | 9.81E-16 | 2.70E-15 | Healthy control | Liver |
| Trim29 | 3.76662 | 1.949622 | 105.5452 | 4.70E-15 | 1.24E-14 | Infection | Liver |
| Spink1 | 3.765341 | 1.887908 | 40.93298 | 2.66E-08 | 4.93E-08 | Infection | Liver |
| Cd37 | 3.759586 | 6.734745 | 988.678 | 3.20E-40 | 1.59E-38 | Infection | Liver |
| Dtnbp1 | -1.41119 | 7.750075 | 648.4581 | 5.41E-35 | 1.03E-33 | Healthy control | Liver |
| Dusp1 | 0.165454 | 6.58302 | 1.359052 | 2.48E-01 | 2.74E-01 | Infection | Liver |
| Dusp11 | 0.654367 | 7.098901 | 84.16488 | 3.07E-13 | 7.40E-13 | Infection | Liver |
| Dusp15 | -0.33139 | 1.319136 | 0.390554 | 5.34E-01 | 5.61E-01 | Healthy control | Liver |
| Dusp3 | -0.38823 | 6.326111 | 41.78859 | 1.67E-08 | 3.12E-08 | Healthy control | Liver |
| Dusp4 | 1.939257 | 5.382629 | 151.7263 | 1.97E-18 | 6.49E-18 | Infection | Liver |
| Fmod | 3.738461 | 3.691134 | 124.7596 | 3.49E-19 | 1.22E-18 | Infection | Liver |
| Dusp6 | -0.96973 | 8.968319 | 165.5509 | 2.46E-19 | 8.72E-19 | Healthy control | Liver |
| Dusp7 | 0.439834 | 5.500446 | 29.64494 | 9.02E-07 | 1.55E-06 | Infection | Liver |
| Dut | -0.23232 | 3.326121 | 2.118857 | 1.50E-01 | 1.71E-01 | Healthy control | Liver |
| Duxbl1 | 1.173154 | 1.36184 | 3.868658 | 5.35E-02 | 6.44E-02 | Infection | Liver |
| Dvl1 | -1.34941 | 9.137505 | 1253.571 | 1.79E-43 | 1.79E-41 | Healthy control | Liver |
| Dvl2 | 0.79061 | 4.908792 | 52.67914 | 7.07E-10 | 1.43E-09 | Infection | Liver |
| Dvl3 | -0.59788 | 6.625669 | 83.41144 | 3.63E-13 | 8.72E-13 | Healthy control | Liver |
| Dxo | -0.24289 | 6.924456 | 19.88427 | 3.43E-05 | 5.41E-05 | Healthy control | Liver |
| Dydc2 | 0.639862 | 1.318407 | 1.18906 | 2.80E-01 | 3.06E-01 | Infection | Liver |
| Dync1h1 | -0.06182 | 8.45383 | 1.097711 | 2.99E-01 | 3.25E-01 | Healthy control | Liver |
| Dync1i1 | 0.312005 | 1.362255 | 0.403304 | 5.27E-01 | 5.55E-01 | Infection | Liver |
| Dync1i2 | -1.42485 | 6.16334 | 329.5858 | 9.15E-27 | 6.22E-26 | Healthy control | Liver |

| | | | | | | | |
|---|---|---|---|---|---|---|---|
| Dync1li1 | -0.56665 | 6.673581 | 51.92073 | 8.51E-10 | 1.71E-09 | Healthy control | Liver |
| Dync2h1 | 0.588507 | 4.690567 | 51.24872 | 1.06E-09 | 2.13E-09 | Infection | Liver |
| Dyrk1a | -0.6903 | 6.571017 | 167.7259 | 1.84E-19 | 6.59E-19 | Healthy control | Liver |
| Dysf | 0.80528 | 6.3659 | 81.80549 | 5.17E-13 | 1.23E-12 | Infection | Liver |
| E2f1 | 1.289249 | 4.549637 | 95.23016 | 3.41E-14 | 8.62E-14 | Infection | Liver |
| E2f2 | 1.763292 | 5.124443 | 231.532 | 9.11E-23 | 4.21E-22 | Infection | Liver |
| E2f3 | -0.68845 | 3.841716 | 21.2125 | 2.00E-05 | 3.19E-05 | Healthy control | Liver |
| E2f4 | -0.18781 | 6.592876 | 8.969926 | 3.91E-03 | 5.32E-03 | Healthy control | Liver |
| E2f5 | 0.039431 | 5.916782 | 0.382745 | 5.38E-01 | 5.65E-01 | Infection | Liver |
| Peg10 | 3.738041 | 3.178456 | 173.8115 | 4.13E-24 | 2.18E-23 | Infection | Liver |
| Carns1 | 3.736159 | 3.266752 | 228.2591 | 1.21E-28 | 1.02E-27 | Infection | Liver |
| Osmr | 3.728422 | 7.896099 | 647.862 | 2.01E-34 | 3.37E-33 | Infection | Liver |
| Ece1 | -1.06023 | 10.01132 | 305.4614 | 6.28E-26 | 3.94E-25 | Healthy control | Liver |
| Ecel1 | 0.516163 | 1.29406 | 0.844624 | 3.62E-01 | 3.89E-01 | Infection | Liver |
| Echdc1 | -1.17606 | 8.709327 | 150.7558 | 2.07E-18 | 6.81E-18 | Healthy control | Liver |
| Echs1 | -1.54817 | 10.83711 | 359.2167 | 8.31E-28 | 6.50E-27 | Healthy control | Liver |
| Eci3 | -3.33599 | 3.227332 | 90.07474 | 3.17E-15 | 8.47E-15 | Healthy control | Liver |
| Ecpas | -0.07927 | 8.184525 | 2.651661 | 1.08E-01 | 1.26E-01 | Healthy control | Liver |
| Ecscr | 1.010696 | 3.953847 | 42.59088 | 1.37E-08 | 2.59E-08 | Infection | Liver |
| Ect2 | 1.475476 | 4.414301 | 81.96808 | 5.63E-13 | 1.34E-12 | Infection | Liver |
| Eda | 0.196352 | 2.819411 | 1.504474 | 2.24E-01 | 2.49E-01 | Infection | Liver |
| Edar | 1.902215 | 1.527058 | 14.26955 | 3.02E-04 | 4.48E-04 | Infection | Liver |
| Edc4 | -0.03492 | 7.297895 | 0.663673 | 4.18E-01 | 4.46E-01 | Healthy control | Liver |
| Col5a1 | 3.727654 | 8.634828 | 270.5134 | 1.54E-23 | 7.61E-23 | Infection | Liver |
| Tnfrsf13b | 3.720909 | 4.362774 | 350.9338 | 4.67E-30 | 4.63E-29 | Infection | Liver |
| Edn3 | 1.011446 | 1.233351 | 2.472376 | 1.28E-01 | 1.47E-01 | Infection | Liver |
| Ednra | 1.435037 | 5.7941 | 77.2715 | 1.64E-12 | 3.78E-12 | Infection | Liver |
| Ednrb | -0.39962 | 5.339202 | 15.22188 | 2.35E-04 | 3.50E-04 | Healthy control | Liver |
| Eea1 | 0.083978 | 7.942548 | 3.124978 | 8.19E-02 | 9.63E-02 | Infection | Liver |
| Eed | -0.5686 | 7.336221 | 100.8412 | 9.97E-15 | 2.59E-14 | Healthy control | Liver |
| Eepd1 | -0.53618 | 7.612466 | 42.90951 | 1.18E-08 | 2.23E-08 | Healthy control | Liver |
| Efcab14 | -0.68147 | 6.14787 | 120.789 | 2.59E-16 | 7.36E-16 | Healthy control | Liver |
| Bcl2a1a | 3.715226 | 5.397146 | 477.2089 | 3.32E-31 | 3.75E-30 | Infection | Liver |
| Efemp1 | 0.711049 | 5.958487 | 31.8945 | 4.13E-07 | 7.21E-07 | Infection | Liver |
| Efna1 | -0.7359 | 7.869932 | 35.1686 | 1.46E-07 | 2.61E-07 | Healthy control | Liver |
| Efna2 | 1.052881 | 2.12786 | 15.13893 | 1.69E-04 | 2.55E-04 | Infection | Liver |
| Efna3 | 1.464046 | 1.448025 | 7.401701 | 7.96E-03 | 1.06E-02 | Infection | Liver |
| Efna5 | 0.004033 | 5.569154 | 0.001184 | 9.73E-01 | 9.75E-01 | Infection | Liver |
| Efnb1 | -0.44746 | 7.067483 | 32.7678 | 3.06E-07 | 5.39E-07 | Healthy control | Liver |
| Efnb2 | 0.536068 | 3.940247 | 11.56001 | 1.18E-03 | 1.68E-03 | Infection | Liver |
| Abca13 | 3.714509 | 1.714826 | 48.30952 | 2.98E-09 | 5.83E-09 | Infection | Liver |
| Egfl7 | -0.47799 | 8.389496 | 54.97292 | 3.66E-10 | 7.49E-10 | Healthy control | Liver |
| Egfr | -0.13287 | 8.724182 | 1.190187 | 2.80E-01 | 3.06E-01 | Healthy control | Liver |
| Egln1 | -0.36566 | 7.053829 | 30.80834 | 5.99E-07 | 1.04E-06 | Healthy control | Liver |
| Egln2 | -1.70573 | 7.949381 | 1467.185 | 1.54E-45 | 3.67E-43 | Healthy control | Liver |
| Egln3 | 0.857708 | 7.580983 | 22.55538 | 1.36E-05 | 2.19E-05 | Infection | Liver |
| Lpl | 3.713527 | 9.547461 | 481.3071 | 3.61E-30 | 3.65E-29 | Infection | Liver |
| Cd200 | 3.708761 | 4.550381 | 245.7385 | 7.40E-25 | 4.20E-24 | Infection | Liver |
| Xcr1 | 3.706283 | 4.463382 | 318.3372 | 3.09E-28 | 2.51E-27 | Infection | Liver |
| Egr4 | 0.874264 | 1.330593 | 2.008634 | 1.61E-01 | 1.82E-01 | Infection | Liver |

| | | | | | | | |
|---|---|---|---|---|---|---|---|
| Ehd4 | 1.146688 | 5.914924 | 100.3362 | 1.11E-14 | 2.88E-14 | Infection | Liver |
| Ehhadh | -2.8156 | 9.94748 | 970.305 | 3.88E-40 | 1.89E-38 | Healthy control | Liver |
| Ehmt1 | -0.75565 | 5.464247 | 68.70519 | 1.10E-11 | 2.44E-11 | Healthy control | Liver |
| Ehmt2 | -0.02515 | 8.678906 | 0.563048 | 4.56E-01 | 4.84E-01 | Healthy control | Liver |
| Eif2a | 0.153785 | 7.709634 | 6.089902 | 1.63E-02 | 2.10E-02 | Infection | Liver |
| Eif2ak2 | 0.600798 | 7.236912 | 80.1662 | 7.41E-13 | 1.75E-12 | Infection | Liver |
| Eif2ak3 | 0.712915 | 7.221074 | 239.4786 | 3.33E-23 | 1.59E-22 | Infection | Liver |
| Eif2ak4 | -0.3561 | 5.10396 | 14.4823 | 3.22E-04 | 4.77E-04 | Healthy control | Liver |
| Eif2b1 | -0.88543 | 6.657284 | 246.8321 | 1.57E-23 | 7.77E-23 | Healthy control | Liver |
| Eif2s1 | 1.503978 | 4.58445 | 196.1393 | 5.83E-21 | 2.31E-20 | Infection | Liver |
| Eif2s2 | 0.508978 | 6.327777 | 53.50739 | 5.48E-10 | 1.11E-09 | Infection | Liver |
| Eif3i | -0.63399 | 7.209179 | 144.4551 | 5.39E-18 | 1.71E-17 | Healthy control | Liver |
| Eif4a3 | -0.60824 | 8.668419 | 219.2564 | 2.99E-22 | 1.33E-21 | Healthy control | Liver |
| Eif4e | -1.03348 | 6.243437 | 197.1367 | 4.16E-21 | 1.67E-20 | Healthy control | Liver |
| Eif4ebp1 | -0.30315 | 3.283623 | 3.549653 | 6.39E-02 | 7.61E-02 | Healthy control | Liver |
| Eif4ebp2 | -0.74828 | 10.1163 | 184.6481 | 1.91E-20 | 7.36E-20 | Healthy control | Liver |
| Eif4enif1 | -0.7306 | 6.950015 | 210.4105 | 8.31E-22 | 3.53E-21 | Healthy control | Liver |
| Eif4g1 | -1.40239 | 9.79948 | 448.1052 | 1.93E-30 | 1.99E-29 | Healthy control | Liver |
| Eif4g3 | -0.47351 | 8.358578 | 123.535 | 1.59E-16 | 4.59E-16 | Healthy control | Liver |
| Eipr1 | -0.23463 | 4.74518 | 5.935209 | 1.77E-02 | 2.27E-02 | Healthy control | Liver |
| Elac2 | -1.17499 | 6.28495 | 231.9079 | 7.73E-23 | 3.58E-22 | Healthy control | Liver |
| Elavl1 | -1.36121 | 8.540029 | 593.9297 | 6.74E-34 | 1.05E-32 | Healthy control | Liver |
| Elavl2 | 0.227524 | 1.572353 | 0.246884 | 6.20E-01 | 6.44E-01 | Infection | Liver |
| Elavl3 | 0.309931 | 1.274129 | 0.302672 | 5.85E-01 | 6.10E-01 | Infection | Liver |
| Elavl4 | 0.820685 | 1.333654 | 2.017464 | 1.61E-01 | 1.82E-01 | Infection | Liver |
| Elf2 | -0.25021 | 7.405168 | 24.51035 | 5.77E-06 | 9.51E-06 | Healthy control | Liver |
| Mboat1 | 3.701861 | 3.235922 | 238.882 | 5.56E-29 | 4.84E-28 | Infection | Liver |
| Elk1 | 0.95192 | 5.32705 | 146.5779 | 4.07E-18 | 1.31E-17 | Infection | Liver |
| Elmo1 | 1.807492 | 5.540108 | 261.9539 | 3.75E-24 | 2.00E-23 | Infection | Liver |
| Elmo3 | -1.14229 | 7.021603 | 201.0166 | 2.53E-21 | 1.04E-20 | Healthy control | Liver |
| Ticam2 | 3.701791 | 4.865797 | 690.5954 | 5.59E-37 | 1.54E-35 | Infection | Liver |
| Elob | -0.89889 | 6.429508 | 152.2173 | 1.69E-18 | 5.61E-18 | Healthy control | Liver |
| Eloc | -1.45954 | 5.340974 | 215.2241 | 5.40E-22 | 2.34E-21 | Healthy control | Liver |
| Elovl1 | -1.26854 | 6.950987 | 348.3845 | 1.94E-27 | 1.45E-26 | Healthy control | Liver |
| Elovl4 | 1.671175 | 1.514839 | 12.37744 | 6.80E-04 | 9.81E-04 | Infection | Liver |
| Elp2 | -0.23039 | 6.624637 | 19.91085 | 3.40E-05 | 5.36E-05 | Healthy control | Liver |
| Emc1 | -0.58289 | 8.271639 | 197.5966 | 3.79E-21 | 1.53E-20 | Healthy control | Liver |
| Emc10 | -0.26974 | 8.29902 | 24.78492 | 5.21E-06 | 8.60E-06 | Healthy control | Liver |
| Emc2 | -1.06369 | 6.356731 | 224.511 | 1.72E-22 | 7.74E-22 | Healthy control | Liver |
| Emc3 | -0.43938 | 7.655737 | 57.99744 | 1.62E-10 | 3.38E-10 | Healthy control | Liver |
| Emc4 | -0.87817 | 7.301635 | 278.5628 | 7.03E-25 | 4.00E-24 | Healthy control | Liver |
| Emc6 | -1.81375 | 8.90119 | 1321.019 | 3.67E-44 | 4.60E-42 | Healthy control | Liver |
| Emd | -0.57373 | 6.213995 | 82.48617 | 4.47E-13 | 1.07E-12 | Healthy control | Liver |
| Inhbb | 3.677117 | 3.734066 | 322.0824 | 1.28E-31 | 1.52E-30 | Infection | Liver |
| Sema5a | 3.675783 | 3.035681 | 181.5089 | 1.45E-24 | 7.98E-24 | Infection | Liver |
| Emsy | -0.10467 | 6.453771 | 3.01406 | 8.74E-02 | 1.02E-01 | Healthy control | Liver |
| Emx1 | 1.36932 | 1.344277 | 5.180252 | 2.63E-02 | 3.29E-02 | Infection | Liver |
| Emx2 | 1.25322 | 1.347997 | 3.867158 | 5.41E-02 | 6.51E-02 | Infection | Liver |
| En1 | 1.64315 | 1.432514 | 8.054153 | 5.79E-03 | 7.78E-03 | Infection | Liver |
| Col9a2 | 3.671376 | 1.851873 | 40.12287 | 3.40E-08 | 6.26E-08 | Infection | Liver |

| | | | | | | | |
|---|---|---|---|---|---|---|---|
| Ctla4 | 3.66544 | 2.816161 | 199.3142 | 4.52E-26 | 2.90E-25 | Infection | Liver |
| Eng | -1.01008 | 9.179306 | 136.8014 | 1.78E-17 | 5.49E-17 | Healthy control | Liver |
| Eno1 | -1.46577 | 7.456947 | 496.1244 | 1.14E-31 | 1.36E-30 | Healthy control | Liver |
| Il21 | 3.664383 | 1.616171 | 56.40997 | 4.97E-10 | 1.01E-09 | Infection | Liver |
| Enpp1 | 0.619118 | 6.537522 | 37.6603 | 6.09E-08 | 1.11E-07 | Infection | Liver |
| Enpp6 | 1.152805 | 1.551187 | 6.228352 | 1.41E-02 | 1.82E-02 | Infection | Liver |
| Entpd1 | 0.898096 | 6.170685 | 109.7293 | 1.86E-15 | 5.03E-15 | Infection | Liver |
| Atp2a3 | 3.661625 | 5.912419 | 656.5512 | 6.70E-35 | 1.25E-33 | Infection | Liver |
| Ep300 | -0.05051 | 7.520522 | 1.013354 | 3.18E-01 | 3.44E-01 | Healthy control | Liver |
| Ep400 | -0.30977 | 7.505546 | 58.13294 | 1.57E-10 | 3.26E-10 | Healthy control | Liver |
| Epas1 | 0.763655 | 10.52625 | 246.4798 | 1.60E-23 | 7.89E-23 | Infection | Liver |
| Epb41 | -0.26647 | 8.848239 | 27.72524 | 1.78E-06 | 3.00E-06 | Healthy control | Liver |
| Epb41l1 | 1.424208 | 5.795791 | 127.8693 | 7.83E-17 | 2.31E-16 | Infection | Liver |
| Epb41l2 | 1.707867 | 5.601097 | 277.6853 | 8.26E-25 | 4.67E-24 | Infection | Liver |
| Epb41l3 | 0.078276 | 4.5996 | 0.356914 | 5.52E-01 | 5.79E-01 | Infection | Liver |
| Epb41l4a | 1.655529 | 3.471175 | 86.28349 | 1.40E-13 | 3.43E-13 | Infection | Liver |
| Scn4b | 3.655365 | 3.011966 | 255.0121 | 6.07E-30 | 5.90E-29 | Infection | Liver |
| Epc1 | 0.360882 | 6.447108 | 46.41803 | 4.13E-09 | 8.00E-09 | Infection | Liver |
| Epc2 | -0.15929 | 6.608935 | 5.86261 | 1.83E-02 | 2.34E-02 | Healthy control | Liver |
| Cldn8 | 3.654542 | 1.713319 | 36.21408 | 1.01E-07 | 1.83E-07 | Infection | Liver |
| Epha1 | -1.19427 | 7.149026 | 147.3286 | 3.49E-18 | 1.12E-17 | Healthy control | Liver |
| Epha10 | -0.04716 | 1.289811 | 0.006491 | 9.36E-01 | 9.43E-01 | Healthy control | Liver |
| Epha2 | -0.32802 | 6.326961 | 24.31869 | 6.21E-06 | 1.02E-05 | Healthy control | Liver |
| Gfi1 | 3.636319 | 3.547863 | 194.5714 | 4.16E-25 | 2.40E-24 | Infection | Liver |
| Epha4 | 1.696967 | 2.678117 | 68.60671 | 1.29E-12 | 2.98E-12 | Infection | Liver |
| Epha5 | 1.294475 | 1.340314 | 5.056102 | 2.81E-02 | 3.51E-02 | Infection | Liver |
| Epha7 | 0.344552 | 4.724001 | 8.822168 | 4.21E-03 | 5.71E-03 | Infection | Liver |
| Knl1 | 3.629064 | 2.999536 | 196.6838 | 6.73E-26 | 4.22E-25 | Infection | Liver |
| Socs1 | 3.625062 | 3.356945 | 255.8629 | 4.80E-30 | 4.73E-29 | Infection | Liver |
| Tubb2b | 3.622462 | 2.759486 | 133.9808 | 1.62E-20 | 6.25E-20 | Infection | Liver |
| Ephb4 | -1.07452 | 6.985799 | 307.5112 | 5.35E-26 | 3.39E-25 | Healthy control | Liver |
| Ephx2 | -2.3216 | 10.06424 | 392.1003 | 1.34E-28 | 1.13E-27 | Healthy control | Liver |
| Epm2a | -1.21959 | 5.100808 | 172.6254 | 1.05E-19 | 3.85E-19 | Healthy control | Liver |
| Knstrn | 3.621945 | 5.167555 | 476.2645 | 2.47E-31 | 2.83E-30 | Infection | Liver |
| Epo | 1.067865 | 1.336077 | 4.279235 | 4.27E-02 | 5.22E-02 | Infection | Liver |
| Epor | -1.0831 | 4.939609 | 39.00338 | 4.05E-08 | 7.43E-08 | Healthy control | Liver |
| Eps15 | 0.990091 | 8.975076 | 596.1331 | 6.04E-34 | 9.42E-33 | Infection | Liver |
| Eps15l1 | 0.861384 | 6.666202 | 277.9089 | 7.50E-25 | 4.26E-24 | Infection | Liver |
| Slamf6 | 3.614518 | 4.048287 | 402.5404 | 3.58E-33 | 5.08E-32 | Infection | Liver |
| Erap1 | 0.447758 | 7.536163 | 22.98153 | 1.03E-05 | 1.67E-05 | Infection | Liver |
| Erbb2 | 0.894952 | 3.594112 | 32.73834 | 3.10E-07 | 5.46E-07 | Infection | Liver |
| Erbb3 | -1.56364 | 8.339891 | 179.0435 | 5.55E-20 | 2.07E-19 | Healthy control | Liver |
| Erbb4 | 1.42938 | 4.845442 | 73.08389 | 4.00E-12 | 8.99E-12 | Infection | Liver |
| Erbin | -0.64742 | 8.25833 | 275.3267 | 9.46E-25 | 5.32E-24 | Healthy control | Liver |
| Erc2 | -0.66353 | 4.522839 | 28.8686 | 1.19E-06 | 2.03E-06 | Healthy control | Liver |
| Havcr2 | 3.613109 | 4.216473 | 311.0687 | 1.40E-28 | 1.17E-27 | Infection | Liver |
| Ercc2 | -0.51356 | 4.875891 | 23.78642 | 7.65E-06 | 1.25E-05 | Healthy control | Liver |
| Ercc6 | -0.21755 | 6.199715 | 13.8733 | 4.19E-04 | 6.16E-04 | Healthy control | Liver |
| Zfp979 | 3.608789 | 3.467147 | 180.7775 | 7.69E-24 | 3.93E-23 | Infection | Liver |
| Fgfr1 | 3.604078 | 6.436391 | 232.076 | 5.19E-22 | 2.25E-21 | Infection | Liver |

| | | | | | | | |
|---|---|---|---|---|---|---|---|
| Erg | 0.141288 | 4.150176 | 1.281265 | 2.62E-01 | 2.88E-01 | Infection | Liver |
| Erg28 | -0.42427 | 7.731958 | 26.81888 | 2.46E-06 | 4.13E-06 | Healthy control | Liver |
| Ern1 | -0.83557 | 8.589885 | 159.0808 | 6.14E-19 | 2.09E-18 | Healthy control | Liver |
| Ern2 | 0.553855 | 1.365251 | 0.973997 | 3.27E-01 | 3.53E-01 | Infection | Liver |
| Ero1l | 1.150509 | 7.439997 | 81.57677 | 6.52E-13 | 1.54E-12 | Infection | Liver |
| Ero1lb | -0.64578 | 7.303949 | 67.56802 | 1.41E-11 | 3.11E-11 | Healthy control | Liver |
| Erp44 | -0.40299 | 8.462 | 44.84246 | 6.57E-09 | 1.26E-08 | Healthy control | Liver |
| Esam | -0.94675 | 5.875361 | 91.09344 | 7.24E-14 | 1.80E-13 | Healthy control | Liver |
| Esd | -1.97876 | 8.382779 | 357.2773 | 1.03E-27 | 7.93E-27 | Healthy control | Liver |
| En2 | 3.586408 | 1.77197 | 59.29211 | 1.25E-10 | 2.61E-10 | Infection | Liver |
| Espn | 1.764852 | 1.587345 | 18.33675 | 4.49E-05 | 7.02E-05 | Infection | Liver |
| Esr1 | -1.0654 | 7.842118 | 150.3984 | 2.19E-18 | 7.18E-18 | Healthy control | Liver |
| Esr2 | 0.638595 | 1.184142 | 2.306477 | 1.51E-01 | 1.71E-01 | Infection | Liver |
| Esrra | 0.746933 | 6.135812 | 101.0743 | 9.59E-15 | 2.50E-14 | Infection | Liver |
| Esrrb | -1.45289 | 2.858422 | 28.27695 | 8.54E-07 | 1.47E-06 | Healthy control | Liver |
| Esrrg | -3.69277 | 3.287063 | 265.9782 | 3.32E-29 | 2.98E-28 | Healthy control | Liver |
| Ess2 | -0.5175 | 5.370288 | 51.5112 | 9.72E-10 | 1.95E-09 | Healthy control | Liver |
| Esyt2 | 0.547161 | 7.309004 | 51.98267 | 8.35E-10 | 1.68E-09 | Infection | Liver |
| Ccr2 | 3.578343 | 4.711435 | 473.7456 | 2.90E-32 | 3.66E-31 | Infection | Liver |
| Etfbkmt | 1.19926 | 6.024772 | 295.1582 | 1.60E-25 | 9.55E-25 | Infection | Liver |
| Etnk1 | 0.177605 | 7.246948 | 10.80848 | 1.65E-03 | 2.32E-03 | Infection | Liver |
| Etnppl | -3.52125 | 8.101151 | 169.6868 | 7.17E-19 | 2.44E-18 | Healthy control | Liver |
| Ets1 | -0.04887 | 7.179867 | 0.460779 | 5.00E-01 | 5.27E-01 | Healthy control | Liver |
| Ets2 | 0.037142 | 7.608395 | 0.315203 | 5.76E-01 | 6.02E-01 | Infection | Liver |
| Etv1 | 0.940777 | 3.484829 | 21.27876 | 1.96E-05 | 3.14E-05 | Infection | Liver |
| Etv2 | 0.913006 | 1.413317 | 2.50443 | 1.17E-01 | 1.35E-01 | Infection | Liver |
| Etv3 | -0.11152 | 6.590863 | 2.926144 | 9.20E-02 | 1.08E-01 | Healthy control | Liver |
| Lin28a | 3.572131 | 1.822117 | 52.18239 | 1.04E-09 | 2.08E-09 | Infection | Liver |
| Etv5 | 0.786478 | 6.458935 | 80.58642 | 6.77E-13 | 1.60E-12 | Infection | Liver |
| Etv6 | -0.07288 | 6.653643 | 1.295274 | 2.59E-01 | 2.85E-01 | Healthy control | Liver |
| Eva1a | -1.39793 | 8.191957 | 210.7886 | 7.90E-22 | 3.37E-21 | Healthy control | Liver |
| Evl | 1.733477 | 6.470868 | 238.6387 | 3.70E-23 | 1.76E-22 | Infection | Liver |
| Evpl | 1.144309 | 2.409304 | 19.62395 | 2.55E-05 | 4.04E-05 | Infection | Liver |
| Slc17a6 | 3.571538 | 1.899976 | 68.47835 | 1.13E-11 | 2.49E-11 | Infection | Liver |
| Evx2 | 1.269538 | 1.220138 | 5.945595 | 2.17E-02 | 2.75E-02 | Infection | Liver |
| Ewsr1 | -1.08999 | 6.823905 | 258.4059 | 4.91E-24 | 2.57E-23 | Healthy control | Liver |
| Exoc3l2 | -1.75521 | 4.74467 | 190.986 | 6.84E-21 | 2.69E-20 | Healthy control | Liver |
| Exoc4 | 0.804129 | 7.976554 | 254.9465 | 6.81E-24 | 3.51E-23 | Infection | Liver |
| Exosc9 | -0.15415 | 6.77915 | 7.621936 | 7.53E-03 | 1.00E-02 | Healthy control | Liver |
| Exph5 | -2.05221 | 6.702768 | 278.5118 | 7.60E-25 | 4.30E-24 | Healthy control | Liver |
| Ext1 | -0.29313 | 7.513394 | 24.87589 | 5.03E-06 | 8.32E-06 | Healthy control | Liver |
| Ext2 | 0.089147 | 8.001517 | 4.309272 | 4.20E-02 | 5.14E-02 | Infection | Liver |
| Extl1 | -2.34911 | 5.548964 | 513.1116 | 4.73E-32 | 5.80E-31 | Healthy control | Liver |
| Has2 | 3.571077 | 1.852993 | 67.15758 | 1.20E-11 | 2.64E-11 | Infection | Liver |
| Itln1 | 3.566639 | 1.709381 | 45.97724 | 5.68E-09 | 1.09E-08 | Infection | Liver |
| Eya3 | -0.17879 | 6.114127 | 8.018737 | 6.20E-03 | 8.31E-03 | Healthy control | Liver |
| Pde1b | 3.557293 | 3.940381 | 352.5961 | 1.94E-31 | 2.26E-30 | Infection | Liver |
| Ezh1 | 0.027716 | 6.60777 | 0.277012 | 6.01E-01 | 6.26E-01 | Infection | Liver |
| Ezh2 | 1.790305 | 6.153373 | 534.778 | 1.42E-32 | 1.86E-31 | Infection | Liver |
| Slc8a1 | 3.554612 | 7.41551 | 1478.28 | 1.34E-45 | 3.36E-43 | Infection | Liver |

| | | | | | | | |
|---|---|---|---|---|---|---|---|
| F11r | 0.23385 | 8.510106 | 7.197656 | 9.30E-03 | 1.23E-02 | Infection | Liver |
| Mrgprf | 3.550306 | 3.798257 | 278.856 | 1.01E-28 | 8.59E-28 | Infection | Liver |
| Sox4 | 3.549906 | 6.967889 | 337.322 | 3.67E-26 | 2.37E-25 | Infection | Liver |
| Cd200r1 | 3.547326 | 5.540978 | 644.2249 | 8.98E-35 | 1.65E-33 | Infection | Liver |
| Themis | 3.545083 | 4.619387 | 475.199 | 2.19E-32 | 2.79E-31 | Infection | Liver |
| Eya1 | 3.544672 | 1.752556 | 47.04255 | 3.57E-09 | 6.94E-09 | Infection | Liver |
| F2rl3 | 3.532096 | 2.449326 | 107.8919 | 3.03E-17 | 9.20E-17 | Infection | Liver |
| F8 | -0.68131 | 6.222989 | 27.62872 | 1.84E-06 | 3.11E-06 | Healthy control | Liver |
| F9 | -1.96834 | 8.875245 | 412.4222 | 1.93E-29 | 1.79E-28 | Healthy control | Liver |
| Faah | -2.24268 | 9.025502 | 791.7419 | 1.60E-37 | 4.75E-36 | Healthy control | Liver |
| Faap100 | -0.0013 | 5.400133 | 0.000198 | 9.89E-01 | 9.90E-01 | Healthy control | Liver |
| Fabp12 | 1.880084 | 1.379 | 8.723987 | 4.43E-03 | 6.00E-03 | Infection | Liver |
| Chil1 | 3.529162 | 3.105278 | 112.8567 | 1.52E-18 | 5.08E-18 | Infection | Liver |
| Fabp7 | -0.00599 | 5.228332 | 0.00165 | 9.68E-01 | 9.71E-01 | Healthy control | Liver |
| Fadd | 0.036733 | 8.206737 | 0.628817 | 4.31E-01 | 4.58E-01 | Infection | Liver |
| Fah | -2.00554 | 10.936 | 694.5361 | 7.31E-36 | 1.60E-34 | Healthy control | Liver |
| Fam104a | 0.962234 | 4.645851 | 106.3594 | 3.85E-15 | 1.02E-14 | Infection | Liver |
| Fam107b | -1.6144 | 8.415482 | 626.135 | 1.48E-34 | 2.59E-33 | Healthy control | Liver |
| Fam129a | 1.742832 | 6.479729 | 458.4714 | 1.06E-30 | 1.12E-29 | Infection | Liver |
| Fam136a | -1.51677 | 8.445128 | 490.1873 | 1.58E-31 | 1.86E-30 | Healthy control | Liver |
| Fam13b | -0.23772 | 7.575132 | 21.50829 | 1.81E-05 | 2.91E-05 | Healthy control | Liver |
| Fam192a | -1.09581 | 5.647395 | 201.997 | 2.44E-21 | 1.00E-20 | Healthy control | Liver |
| Fam20a | -0.6706 | 7.577384 | 142.5069 | 7.27E-18 | 2.29E-17 | Healthy control | Liver |
| Fam20c | -0.75558 | 8.235881 | 91.35404 | 6.68E-14 | 1.66E-13 | Healthy control | Liver |
| Fam210a | -1.60704 | 7.931201 | 357.2451 | 9.73E-28 | 7.57E-27 | Healthy control | Liver |
| Fam210b | -2.23895 | 7.795205 | 1282.218 | 9.32E-44 | 1.01E-41 | Healthy control | Liver |
| Fam214a | -1.68737 | 7.682462 | 810.3451 | 8.16E-38 | 2.54E-36 | Healthy control | Liver |
| Il33 | 3.516522 | 6.241216 | 353.3015 | 3.54E-27 | 2.52E-26 | Infection | Liver |
| Fam83d | 3.514334 | 3.580571 | 218.4378 | 4.06E-26 | 2.61E-25 | Infection | Liver |
| Fam92a | -0.16246 | 7.176831 | 7.48227 | 8.07E-03 | 1.07E-02 | Healthy control | Liver |
| Fancc | 0.773211 | 3.854929 | 40.11717 | 2.92E-08 | 5.40E-08 | Infection | Liver |
| Nr4a3 | 3.512202 | 2.970759 | 169.5129 | 9.04E-24 | 4.60E-23 | Infection | Liver |
| Fancl | 0.033152 | 6.575313 | 0.274976 | 6.02E-01 | 6.27E-01 | Infection | Liver |
| Mmp13 | 3.506247 | 5.711218 | 337.3975 | 8.39E-27 | 5.75E-26 | Infection | Liver |
| Slc2a3 | 3.505347 | 4.958675 | 458.6661 | 4.76E-31 | 5.24E-30 | Infection | Liver |
| Fas | -0.31127 | 6.414094 | 17.00767 | 1.10E-04 | 1.68E-04 | Healthy control | Liver |
| Muc1 | 3.504244 | 2.443753 | 89.86843 | 2.35E-15 | 6.32E-15 | Infection | Liver |
| Fasn | -1.59221 | 10.9742 | 261.2221 | 3.65E-24 | 1.95E-23 | Healthy control | Liver |
| Fat1 | -0.19593 | 7.596936 | 12.89455 | 6.43E-04 | 9.30E-04 | Healthy control | Liver |
| Fat4 | 0.817602 | 4.969221 | 52.43029 | 7.56E-10 | 1.52E-09 | Infection | Liver |
| Fbh1 | 0.311708 | 7.380162 | 41.86019 | 1.63E-08 | 3.05E-08 | Infection | Liver |
| Fbl | -0.43706 | 7.374333 | 47.70302 | 2.83E-09 | 5.54E-09 | Healthy control | Liver |
| Mybl2 | 3.501728 | 3.098151 | 187.1741 | 3.18E-25 | 1.85E-24 | Infection | Liver |
| Batf3 | 3.500225 | 6.773848 | 1404.756 | 7.52E-45 | 1.27E-42 | Infection | Liver |
| Clec10a | 3.499264 | 4.501004 | 282.6177 | 4.03E-26 | 2.59E-25 | Infection | Liver |
| Fbxl12 | -0.422 | 5.695705 | 28.1324 | 1.54E-06 | 2.61E-06 | Healthy control | Liver |
| Shcbp1l | 3.496014 | 2.047258 | 78.03326 | 5.39E-13 | 1.28E-12 | Infection | Liver |
| Sema3c | 3.493286 | 2.038124 | 61.14605 | 4.01E-11 | 8.61E-11 | Infection | Liver |
| Fbxl17 | -1.3314 | 6.401689 | 465.3415 | 7.27E-31 | 7.87E-30 | Healthy control | Liver |
| Fbxl19 | 1.127104 | 6.824617 | 347.7319 | 2.02E-27 | 1.51E-26 | Infection | Liver |

| | | | | | | | |
|---|---|---|---|---|---|---|---|
| Fbxl20 | -0.63178 | 7.537805 | 157.2567 | 8.01E-19 | 2.71E-18 | Healthy control | Liver |
| Fbxl3 | -0.4433 | 6.74368 | 72.14217 | 4.70E-12 | 1.06E-11 | Healthy control | Liver |
| Fbxl4 | -1.16467 | 6.477121 | 246.4262 | 1.66E-23 | 8.17E-23 | Healthy control | Liver |
| Fbxl7 | 0.086516 | 3.526906 | 0.337861 | 5.63E-01 | 5.89E-01 | Infection | Liver |
| Fbxo11 | -0.11331 | 6.227837 | 2.67297 | 1.07E-01 | 1.24E-01 | Healthy control | Liver |
| Fbxo17 | 0.737252 | 4.872187 | 63.07871 | 4.50E-11 | 9.65E-11 | Infection | Liver |
| Fbxo2 | 0.829613 | 1.783835 | 5.205649 | 2.41E-02 | 3.04E-02 | Infection | Liver |
| Fbxo21 | -2.65553 | 7.422745 | 924.4731 | 1.80E-39 | 7.45E-38 | Healthy control | Liver |
| Fbxo3 | -0.168 | 8.851128 | 6.419623 | 1.38E-02 | 1.78E-02 | Healthy control | Liver |
| Fbxo31 | -0.29634 | 7.268257 | 21.2584 | 2.00E-05 | 3.19E-05 | Healthy control | Liver |
| Fbxo32 | 1.201331 | 4.764158 | 50.09567 | 1.48E-09 | 2.93E-09 | Infection | Liver |
| Fbxo33 | -1.56878 | 5.844504 | 483.97 | 2.74E-31 | 3.11E-30 | Healthy control | Liver |
| Fbxo4 | -0.16009 | 6.049785 | 6.743772 | 1.17E-02 | 1.53E-02 | Healthy control | Liver |
| Smpd3 | 3.48962 | 3.763067 | 230.9258 | 4.78E-26 | 3.05E-25 | Infection | Liver |
| Fbxo5 | 1.527276 | 4.179418 | 118.6754 | 4.56E-16 | 1.27E-15 | Infection | Liver |
| Fbxw11 | -0.05034 | 7.300065 | 0.998909 | 3.21E-01 | 3.47E-01 | Healthy control | Liver |
| Fbxw7 | 0.453123 | 6.667549 | 52.33051 | 7.59E-10 | 1.53E-09 | Infection | Liver |
| Fcgr1 | 1.721925 | 6.107709 | 265.8443 | 2.42E-24 | 1.31E-23 | Infection | Liver |
| Fcgr2b | 0.669826 | 9.377414 | 60.03884 | 9.48E-11 | 2.00E-10 | Infection | Liver |
| Def6 | 3.487907 | 4.814259 | 466.1594 | 1.74E-31 | 2.05E-30 | Infection | Liver |
| Fcgr4 | 1.314199 | 6.390516 | 153.8415 | 1.33E-18 | 4.45E-18 | Infection | Liver |
| Fcgrt | 0.12382 | 10.29421 | 11.65565 | 1.12E-03 | 1.60E-03 | Infection | Liver |
| Megf11 | 3.485241 | 2.512823 | 104.5588 | 4.95E-17 | 1.48E-16 | Infection | Liver |
| Il13ra2 | 3.484579 | 4.849008 | 332.7583 | 2.29E-27 | 1.69E-26 | Infection | Liver |
| Fdx1 | -1.42496 | 10.21781 | 292.5262 | 1.95E-25 | 1.16E-24 | Healthy control | Liver |
| Fen1 | 0.057408 | 3.938318 | 0.130888 | 7.19E-01 | 7.39E-01 | Infection | Liver |
| Serpina3a | 3.483734 | 2.1064 | 86.25582 | 4.62E-14 | 1.16E-13 | Infection | Liver |
| Fermt3 | 1.901394 | 6.320146 | 419.2475 | 1.28E-29 | 1.20E-28 | Infection | Liver |
| Fezf1 | 0.890958 | 1.388009 | 2.418097 | 1.24E-01 | 1.43E-01 | Infection | Liver |
| Fezf2 | 0.299528 | 1.197895 | 0.30956 | 5.85E-01 | 6.10E-01 | Infection | Liver |
| Ffar1 | 1.913136 | 1.572761 | 16.58523 | 1.02E-04 | 1.57E-04 | Infection | Liver |
| Mageb4 | 3.482804 | 1.740028 | 23.82821 | 6.84E-06 | 1.12E-05 | Infection | Liver |
| Ffar3 | 0.47553 | 1.208506 | 0.71857 | 4.06E-01 | 4.34E-01 | Infection | Liver |
| Cd8a | 3.48175 | 4.650006 | 190.0054 | 2.18E-21 | 8.97E-21 | Infection | Liver |
| Fga | -0.18351 | 13.33505 | 1.810814 | 1.83E-01 | 2.05E-01 | Healthy control | Liver |
| Fgb | -0.37368 | 13.64716 | 16.3662 | 1.44E-04 | 2.18E-04 | Healthy control | Liver |
| Fgf1 | -1.51032 | 7.985489 | 162.2216 | 5.34E-19 | 1.83E-18 | Healthy control | Liver |
| Map6 | 3.481158 | 1.814719 | 62.52772 | 4.05E-11 | 8.70E-11 | Infection | Liver |
| Fgf12 | 0.979827 | 3.129058 | 35.09282 | 1.20E-07 | 2.16E-07 | Infection | Liver |
| Fgf15 | 0.664149 | 1.278718 | 1.213161 | 2.77E-01 | 3.03E-01 | Infection | Liver |
| Fgf16 | 1.564144 | 1.484097 | 10.6816 | 1.52E-03 | 2.14E-03 | Infection | Liver |
| Arhgef6 | 3.476868 | 6.853013 | 1412.816 | 6.10E-45 | 1.13E-42 | Infection | Liver |
| Sell | 3.476777 | 5.483621 | 354.7247 | 1.48E-27 | 1.13E-26 | Infection | Liver |
| Dkk2 | 3.475007 | 1.918459 | 65.22977 | 1.00E-11 | 2.22E-11 | Infection | Liver |
| Fgf21 | 0.955558 | 5.590786 | 46.47792 | 4.08E-09 | 7.92E-09 | Infection | Liver |
| Fgf22 | 0.386745 | 1.278915 | 0.473697 | 4.95E-01 | 5.22E-01 | Infection | Liver |
| Espl1 | 3.471906 | 4.282408 | 214.8729 | 2.40E-23 | 1.16E-22 | Infection | Liver |
| Fgf3 | 1.346202 | 1.310398 | 4.311389 | 4.29E-02 | 5.24E-02 | Infection | Liver |
| Fgf4 | 1.41444 | 1.302926 | 5.263479 | 2.62E-02 | 3.28E-02 | Infection | Liver |
| Fgf5 | 1.104683 | 1.29015 | 2.863992 | 9.74E-02 | 1.14E-01 | Infection | Liver |

| | | | | | | | |
|---|---|---|---|---|---|---|---|
| Fgf7 | 1.814957 | 3.056678 | 71.05917 | 1.74E-12 | 4.00E-12 | Infection | Liver |
| Fgf8 | -1.08869 | 1.28873 | 3.981962 | 5.10E-02 | 6.16E-02 | Healthy control | Liver |
| Fgf9 | 0.859589 | 1.387843 | 2.55927 | 1.14E-01 | 1.31E-01 | Infection | Liver |
| Fgfbp3 | 1.360694 | 4.103319 | 152.7791 | 2.01E-18 | 6.62E-18 | Infection | Liver |
| Lat | 3.467849 | 6.6536 | 596.3582 | 8.70E-34 | 1.32E-32 | Infection | Liver |
| Fgfr1op | -0.78413 | 5.766012 | 111.791 | 1.31E-15 | 3.57E-15 | Healthy control | Liver |
| Fgfr2 | 0.035149 | 8.355998 | 0.200482 | 6.56E-01 | 6.79E-01 | Infection | Liver |
| Fgfr3 | -0.58807 | 8.123846 | 31.20607 | 5.32E-07 | 9.25E-07 | Healthy control | Liver |
| Fgfr4 | -1.15081 | 7.997406 | 138.616 | 1.34E-17 | 4.16E-17 | Healthy control | Liver |
| Fgg | -0.49117 | 14.22486 | 18.77643 | 5.35E-05 | 8.32E-05 | Healthy control | Liver |
| Fgl1 | -0.12744 | 10.31266 | 1.056926 | 3.08E-01 | 3.34E-01 | Healthy control | Liver |
| Fgr | 1.831811 | 4.947023 | 257.5212 | 6.84E-24 | 3.52E-23 | Infection | Liver |
| Fh1 | -1.9974 | 8.719624 | 823.8799 | 4.95E-38 | 1.60E-36 | Healthy control | Liver |
| Uts2b | 3.46347 | 2.136837 | 66.3915 | 4.81E-12 | 1.08E-11 | Infection | Liver |
| Fhod1 | 0.887604 | 6.316832 | 187.659 | 1.32E-20 | 5.12E-20 | Infection | Liver |
| Fibcd1 | 1.262941 | 1.47879 | 5.832899 | 1.77E-02 | 2.27E-02 | Infection | Liver |
| Fign | -0.79349 | 5.358765 | 123.3713 | 1.74E-16 | 5.02E-16 | Healthy control | Liver |
| Fkbp5 | 0.049313 | 7.294044 | 0.525301 | 4.71E-01 | 4.99E-01 | Infection | Liver |
| Fli1 | 1.36583 | 6.561975 | 343.8955 | 2.74E-27 | 1.99E-26 | Infection | Liver |
| Flna | 1.863373 | 8.687778 | 298.1132 | 1.57E-25 | 9.38E-25 | Infection | Liver |
| Flnb | -0.00697 | 8.519966 | 0.023137 | 8.80E-01 | 8.91E-01 | Healthy control | Liver |
| Flot1 | -0.28722 | 6.731466 | 19.01904 | 4.85E-05 | 7.58E-05 | Healthy control | Liver |
| Flot2 | -0.58905 | 8.01939 | 211.5957 | 7.19E-22 | 3.09E-21 | Healthy control | Liver |
| Flt1 | 1.906614 | 6.579012 | 182.7249 | 3.50E-20 | 1.32E-19 | Infection | Liver |
| Gabrp | 3.451397 | 1.965135 | 54.95808 | 2.08E-10 | 4.32E-10 | Infection | Liver |
| Flt3l | 0.608633 | 4.128417 | 24.20955 | 6.60E-06 | 1.08E-05 | Infection | Liver |
| Flt4 | -0.79931 | 8.020556 | 84.05979 | 3.14E-13 | 7.56E-13 | Healthy control | Liver |
| Fmc1 | -1.74065 | 8.252228 | 945.178 | 8.53E-40 | 3.81E-38 | Healthy control | Liver |
| Fmo4 | -1.88121 | 6.280446 | 316.8161 | 2.69E-26 | 1.77E-25 | Healthy control | Liver |
| Cd6 | 3.45134 | 4.243929 | 402.3966 | 2.52E-31 | 2.88E-30 | Infection | Liver |
| Fmr1 | 1.636513 | 8.614581 | 743.5224 | 1.00E-36 | 2.69E-35 | Infection | Liver |
| Fn1 | 0.297923 | 12.66596 | 16.3926 | 1.43E-04 | 2.16E-04 | Infection | Liver |
| Wnt2b | 3.451144 | 1.825432 | 75.04638 | 1.81E-12 | 4.16E-12 | Infection | Liver |
| Ptpn22 | 3.447484 | 3.487386 | 211.7823 | 7.06E-26 | 4.42E-25 | Infection | Liver |
| Folr2 | 0.991185 | 5.984482 | 108.1008 | 2.53E-15 | 6.77E-15 | Infection | Liver |
| Fopnl | -1.22681 | 5.359886 | 182.1942 | 2.96E-20 | 1.12E-19 | Healthy control | Liver |
| Acss1 | 3.446097 | 4.307603 | 274.9974 | 2.85E-26 | 1.87E-25 | Infection | Liver |
| Plxdc2 | 3.443788 | 5.130273 | 277.1512 | 6.20E-25 | 3.54E-24 | Infection | Liver |
| Foxa1 | -1.32228 | 5.884048 | 146.911 | 3.94E-18 | 1.27E-17 | Healthy control | Liver |
| Foxa2 | -1.44993 | 7.17431 | 222.9428 | 2.01E-22 | 9.02E-22 | Healthy control | Liver |
| Angpt4 | 3.439967 | 2.014137 | 59.41088 | 4.50E-11 | 9.64E-11 | Infection | Liver |
| Foxc1 | 1.125559 | 1.783941 | 8.588928 | 4.02E-03 | 5.46E-03 | Infection | Liver |
| Birc5 | 3.4395 | 5.213097 | 481.8046 | 2.27E-31 | 2.61E-30 | Infection | Liver |
| Foxd1 | 1.507933 | 1.288544 | 6.145621 | 1.68E-02 | 2.16E-02 | Infection | Liver |
| Foxd2 | 1.674479 | 2.313284 | 46.9248 | 4.15E-10 | 8.48E-10 | Infection | Liver |
| Foxd3 | 0.835722 | 1.197232 | 2.707882 | 1.16E-01 | 1.34E-01 | Infection | Liver |
| Foxf1 | 0.976077 | 4.156132 | 45.67027 | 5.46E-09 | 1.05E-08 | Infection | Liver |
| Slc2a4 | 3.437119 | 3.680231 | 162.1697 | 9.28E-22 | 3.93E-21 | Infection | Liver |
| Foxh1 | -0.24565 | 1.444238 | 0.270648 | 6.04E-01 | 6.29E-01 | Healthy control | Liver |
| Foxi1 | 1.183245 | 1.289279 | 3.831211 | 5.59E-02 | 6.72E-02 | Infection | Liver |

| | | | | | | | |
|---|---|---|---|---|---|---|---|
| Foxi2 | 1.867627 | 1.323917 | 8.619657 | 5.23E-03 | 7.05E-03 | Infection | Liver |
| Spib | 3.435965 | 5.058199 | 251.0138 | 8.49E-24 | 4.32E-23 | Infection | Liver |
| Foxj2 | -0.8993 | 4.718819 | 74.64496 | 2.73E-12 | 6.19E-12 | Healthy control | Liver |
| Foxj3 | 0.965911 | 5.20382 | 151.5679 | 1.95E-18 | 6.44E-18 | Infection | Liver |
| Foxl2 | 1.565589 | 1.376404 | 7.009112 | 1.02E-02 | 1.33E-02 | Infection | Liver |
| Diaph3 | 3.433394 | 3.976673 | 265.1556 | 7.36E-27 | 5.08E-26 | Infection | Liver |
| Foxn1 | -0.49973 | 1.232516 | 0.732295 | 3.99E-01 | 4.27E-01 | Healthy control | Liver |
| Foxn4 | 1.589179 | 1.291181 | 5.853093 | 2.00E-02 | 2.55E-02 | Infection | Liver |
| Foxo1 | -0.64347 | 7.19081 | 62.17306 | 5.47E-11 | 1.17E-10 | Healthy control | Liver |
| Foxo3 | -1.30376 | 6.152867 | 200.337 | 2.86E-21 | 1.16E-20 | Healthy control | Liver |
| Foxo4 | -1.90237 | 4.242406 | 153.9323 | 5.75E-19 | 1.97E-18 | Healthy control | Liver |
| Foxp1 | 0.039345 | 7.511011 | 0.709623 | 4.03E-01 | 4.30E-01 | Infection | Liver |
| Foxp2 | -1.9747 | 3.931868 | 127.9118 | 1.57E-17 | 4.85E-17 | Healthy control | Liver |
| Itgb6 | 3.431746 | 2.061887 | 68.68225 | 3.87E-12 | 8.73E-12 | Infection | Liver |
| Foxp4 | 0.214022 | 7.864735 | 17.02495 | 1.10E-04 | 1.67E-04 | Infection | Liver |
| Fpr1 | -0.86926 | 4.7121 | 32.64791 | 3.22E-07 | 5.66E-07 | Healthy control | Liver |
| Fpr2 | 0.931005 | 5.043768 | 23.65097 | 8.16E-06 | 1.33E-05 | Infection | Liver |
| Fras1 | -0.15229 | 2.303952 | 0.383114 | 5.37E-01 | 5.64E-01 | Healthy control | Liver |
| Frk | 0.471298 | 4.877923 | 34.251 | 1.89E-07 | 3.37E-07 | Infection | Liver |
| Frmd7 | 0.911724 | 1.655496 | 3.973947 | 4.83E-02 | 5.86E-02 | Infection | Liver |
| Frmpd1 | 0.70698 | 1.330963 | 1.732777 | 1.93E-01 | 2.15E-01 | Infection | Liver |
| Frrs1 | 1.106534 | 5.855337 | 224.5757 | 1.71E-22 | 7.70E-22 | Infection | Liver |
| Plppr4 | 3.430966 | 1.844852 | 57.55337 | 1.52E-10 | 3.18E-10 | Infection | Liver |
| Frs2 | 0.345021 | 7.101852 | 76.57101 | 1.67E-12 | 3.85E-12 | Infection | Liver |
| Fryl | -0.461 | 6.018915 | 31.03648 | 5.55E-07 | 9.62E-07 | Healthy control | Liver |
| Ccl1 | 3.430196 | 2.432098 | 43.16925 | 2.65E-09 | 5.19E-09 | Infection | Liver |
| Itgb2 | 3.424152 | 7.761425 | 1383.895 | 9.47E-45 | 1.48E-42 | Infection | Liver |
| Fscn2 | 0.893557 | 1.296604 | 2.341281 | 1.32E-01 | 1.52E-01 | Infection | Liver |
| Fshb | 1.254657 | 1.221404 | 5.362497 | 2.86E-02 | 3.56E-02 | Infection | Liver |
| Fshr | 1.53033 | 1.299635 | 6.866532 | 1.22E-02 | 1.59E-02 | Infection | Liver |
| Gria4 | 3.423652 | 3.314527 | 141.7193 | 5.05E-21 | 2.01E-20 | Infection | Liver |
| Sema3a | 3.413246 | 5.123158 | 337.7253 | 3.75E-27 | 2.66E-26 | Infection | Liver |
| Fstl4 | 1.581324 | 1.748254 | 16.69331 | 7.92E-05 | 1.22E-04 | Infection | Liver |
| Cxcr2 | 3.411897 | 3.338037 | 148.4768 | 1.89E-21 | 7.81E-21 | Infection | Liver |
| Fthl17a | 0.638912 | 1.302817 | 1.137537 | 2.91E-01 | 3.17E-01 | Infection | Liver |
| Fto | -0.0303 | 7.3681 | 0.574824 | 4.51E-01 | 4.79E-01 | Healthy control | Liver |
| Ftsj1 | -0.8124 | 6.497532 | 157.7991 | 7.49E-19 | 2.54E-18 | Healthy control | Liver |
| Fubp1 | 0.678086 | 7.214283 | 249.7666 | 1.15E-23 | 5.75E-23 | Infection | Liver |
| Fundc1 | 0.726773 | 7.132086 | 180.5422 | 3.27E-20 | 1.23E-19 | Infection | Liver |
| Furin | -1.43354 | 9.954982 | 603.3524 | 4.27E-34 | 6.85E-33 | Healthy control | Liver |
| Fus | 0.975365 | 8.545947 | 485.6047 | 2.05E-31 | 2.37E-30 | Infection | Liver |
| Fut1 | 1.156544 | 2.489728 | 24.7752 | 3.00E-06 | 5.00E-06 | Infection | Liver |
| Nkd2 | 3.410193 | 3.597099 | 188.4423 | 1.06E-23 | 5.37E-23 | Infection | Liver |
| Fv1 | 0.408846 | 5.167365 | 23.29857 | 9.16E-06 | 1.49E-05 | Infection | Liver |
| Fxn | -1.40426 | 2.706365 | 35.71697 | 4.99E-08 | 9.12E-08 | Healthy control | Liver |
| Fxr1 | 0.508481 | 7.477841 | 74.71256 | 2.57E-12 | 5.82E-12 | Infection | Liver |
| Fyn | 0.88199 | 7.466591 | 167.5027 | 1.88E-19 | 6.71E-19 | Infection | Liver |
| Fzd1 | 0.679291 | 7.336919 | 29.09316 | 1.14E-06 | 1.95E-06 | Infection | Liver |
| Syt4 | 3.409412 | 1.734817 | 55.321 | 2.55E-10 | 5.26E-10 | Infection | Liver |
| Socs3 | 3.408172 | 7.091326 | 747.7174 | 9.66E-37 | 2.61E-35 | Infection | Liver |

| | | | | | | | |
|---|---|---|---|---|---|---|---|
| Plau | 3.401769 | 6.294179 | 275.4334 | 4.64E-24 | 2.44E-23 | Infection | Liver |
| Fzd4 | -0.8106 | 5.951624 | 143.7976 | 6.14E-18 | 1.94E-17 | Healthy control | Liver |
| Fzd5 | -0.09322 | 5.71222 | 0.949677 | 3.34E-01 | 3.60E-01 | Healthy control | Liver |
| Fzd6 | -0.21513 | 4.522678 | 3.034702 | 8.64E-02 | 1.01E-01 | Healthy control | Liver |
| Fzd7 | -0.73222 | 6.462556 | 109.1147 | 2.08E-15 | 5.61E-15 | Healthy control | Liver |
| Fzd8 | -1.71384 | 6.232824 | 517.4021 | 3.93E-32 | 4.89E-31 | Healthy control | Liver |
| G3bp1 | -0.64703 | 7.069153 | 142.6512 | 7.13E-18 | 2.25E-17 | Healthy control | Liver |
| G6pc | -1.12868 | 10.44194 | 22.10824 | 1.63E-05 | 2.61E-05 | Healthy control | Liver |
| Frzb | 3.399836 | 5.799733 | 315.9158 | 5.82E-26 | 3.67E-25 | Infection | Liver |
| G6pdx | 1.195719 | 3.158989 | 44.8743 | 5.14E-09 | 9.91E-09 | Infection | Liver |
| Gaa | -1.31442 | 7.165603 | 180.9162 | 3.13E-20 | 1.18E-19 | Healthy control | Liver |
| Gab1 | -0.28716 | 5.997425 | 12.7329 | 6.92E-04 | 9.98E-04 | Healthy control | Liver |
| Ccna2 | 3.399007 | 5.409968 | 402.0543 | 5.02E-29 | 4.40E-28 | Infection | Liver |
| Gabbr2 | -3.10259 | 4.953419 | 361.9851 | 3.19E-29 | 2.87E-28 | Healthy control | Liver |
| Gabpb1 | -0.85762 | 6.120447 | 209.49 | 9.54E-22 | 4.04E-21 | Healthy control | Liver |
| Gabra1 | 1.648192 | 1.394394 | 6.520538 | 1.30E-02 | 1.69E-02 | Infection | Liver |
| Gabra2 | 1.742053 | 1.265122 | 10.60689 | 2.19E-03 | 3.04E-03 | Infection | Liver |
| Gabra5 | 1.01149 | 1.274366 | 2.581646 | 1.15E-01 | 1.33E-01 | Infection | Liver |
| Gabra6 | 1.554378 | 1.301303 | 5.554409 | 2.24E-02 | 2.84E-02 | Infection | Liver |
| Gabrb2 | 1.946709 | 1.384635 | 11.81455 | 1.02E-03 | 1.46E-03 | Infection | Liver |
| Gabrb3 | 1.150747 | 1.35088 | 3.944951 | 5.11E-02 | 6.17E-02 | Infection | Liver |
| Gabrd | 0.056607 | 1.302653 | 0.008371 | 9.27E-01 | 9.35E-01 | Infection | Liver |
| Gabrg1 | 1.058986 | 1.355068 | 3.282381 | 7.44E-02 | 8.79E-02 | Infection | Liver |
| Gabrg2 | 1.303967 | 1.264508 | 5.287594 | 2.67E-02 | 3.34E-02 | Infection | Liver |
| Gabrg3 | 1.985758 | 1.34424 | 12.06759 | 9.95E-04 | 1.42E-03 | Infection | Liver |
| Cacna1b | 3.395707 | 1.990953 | 78.72986 | 3.97E-13 | 9.50E-13 | Infection | Liver |
| Trpc6 | 3.392602 | 1.876296 | 52.36761 | 4.96E-10 | 1.01E-09 | Infection | Liver |
| Gabrr2 | 0.154 | 2.131215 | 0.331304 | 5.66E-01 | 5.92E-01 | Infection | Liver |
| Amelx | 3.390941 | 2.129922 | 38.12555 | 2.70E-08 | 5.00E-08 | Infection | Liver |
| Gad2 | 1.656386 | 1.282348 | 4.665115 | 3.78E-02 | 4.65E-02 | Infection | Liver |
| Gadd45a | 0.218914 | 4.22964 | 2.143316 | 1.48E-01 | 1.68E-01 | Infection | Liver |
| Gak | -0.2598 | 7.83848 | 35.02597 | 1.44E-07 | 2.58E-07 | Healthy control | Liver |
| Epb42 | 3.389498 | 1.76811 | 40.46919 | 1.76E-08 | 3.29E-08 | Infection | Liver |
| Galc | 1.63433 | 5.696228 | 349.4308 | 1.90E-27 | 1.43E-26 | Infection | Liver |
| Galnt14 | -1.3546 | 2.509915 | 26.24259 | 1.71E-06 | 2.89E-06 | Healthy control | Liver |
| Galnt15 | 0.976721 | 3.109286 | 20.75702 | 2.25E-05 | 3.58E-05 | Infection | Liver |
| Fzd10 | 3.38568 | 1.632327 | 30.79328 | 5.44E-07 | 9.45E-07 | Infection | Liver |
| Galnt18 | 0.194282 | 4.675573 | 2.436757 | 1.24E-01 | 1.42E-01 | Infection | Liver |
| Galntl5 | -0.02556 | 1.278202 | 0.002022 | 9.64E-01 | 9.68E-01 | Healthy control | Liver |
| Galp | 1.33518 | 1.265666 | 5.824584 | 2.09E-02 | 2.66E-02 | Infection | Liver |
| Galr1 | 1.26189 | 1.254215 | 4.188074 | 4.81E-02 | 5.83E-02 | Infection | Liver |
| Galr2 | 0.72659 | 1.694096 | 3.097845 | 8.09E-02 | 9.51E-02 | Infection | Liver |
| Eomes | 3.381142 | 3.052745 | 158.9286 | 5.18E-23 | 2.43E-22 | Infection | Liver |
| Gar1 | 0.731064 | 5.831455 | 109.9142 | 1.81E-15 | 4.90E-15 | Infection | Liver |
| Garem1 | -0.68363 | 6.837243 | 49.99305 | 1.47E-09 | 2.90E-09 | Healthy control | Liver |
| Gars | -0.0729 | 8.264656 | 2.674661 | 1.07E-01 | 1.24E-01 | Healthy control | Liver |
| Gart | -0.64773 | 6.813555 | 113.399 | 9.48E-16 | 2.61E-15 | Healthy control | Liver |
| F2r | 3.375675 | 8.140183 | 487.4385 | 1.44E-30 | 1.51E-29 | Infection | Liver |
| Gas6 | 0.948696 | 8.100434 | 75.19846 | 2.46E-12 | 5.60E-12 | Infection | Liver |
| Gast | 0.340372 | 1.196518 | 0.387088 | 5.42E-01 | 5.68E-01 | Infection | Liver |

| | | | | | | | |
|---|---|---|---|---|---|---|---|
| Nlrp12 | 3.375369 | 2.293992 | 132.7998 | 1.89E-19 | 6.74E-19 | Infection | Liver |
| Gata2 | 1.609089 | 4.39224 | 75.67239 | 2.32E-12 | 5.27E-12 | Infection | Liver |
| Bmp8b | 3.367667 | 1.840214 | 75.48865 | 1.41E-12 | 3.25E-12 | Infection | Liver |
| Gata4 | -0.45662 | 6.780356 | 39.60295 | 3.29E-08 | 6.06E-08 | Healthy control | Liver |
| Gata5 | -0.43845 | 1.684526 | 1.237261 | 2.68E-01 | 2.94E-01 | Healthy control | Liver |
| Gata6 | 1.287942 | 5.779084 | 211.0799 | 7.93E-22 | 3.38E-21 | Infection | Liver |
| Gatad2b | -0.30588 | 7.195916 | 31.07431 | 5.46E-07 | 9.48E-07 | Healthy control | Liver |
| Gatd3a | -1.55148 | 9.69977 | 648.1359 | 5.43E-35 | 1.03E-33 | Healthy control | Liver |
| Gba | 0.218852 | 6.369623 | 9.720425 | 2.74E-03 | 3.78E-03 | Infection | Liver |
| Gba2 | 0.310644 | 6.309885 | 16.78451 | 1.21E-04 | 1.84E-04 | Infection | Liver |
| Gbf1 | 0.272244 | 7.825462 | 32.11327 | 3.82E-07 | 6.69E-07 | Infection | Liver |
| Hoxa9 | 3.362594 | 1.612554 | 46.58905 | 3.28E-09 | 6.39E-09 | Infection | Liver |
| Ccne1 | 3.362299 | 3.41151 | 187.7082 | 3.95E-24 | 2.09E-23 | Infection | Liver |
| Gbp3 | 0.542301 | 6.862963 | 14.10791 | 3.85E-04 | 5.67E-04 | Infection | Liver |
| Gbx1 | 1.441086 | 1.535057 | 8.811069 | 3.72E-03 | 5.07E-03 | Infection | Liver |
| Igfbp1 | 3.358798 | 10.07975 | 553.285 | 2.29E-32 | 2.91E-31 | Infection | Liver |
| Gcat | -2.91396 | 8.071002 | 1284.37 | 9.04E-44 | 1.01E-41 | Healthy control | Liver |
| Gcg | 0.688541 | 1.311883 | 1.553036 | 2.18E-01 | 2.42E-01 | Infection | Liver |
| Gcgr | -2.0128 | 9.875034 | 277.4958 | 1.40E-24 | 7.77E-24 | Healthy control | Liver |
| Gch1 | -2.28074 | 8.691528 | 706.0461 | 4.57E-36 | 1.04E-34 | Healthy control | Liver |
| Gck | -0.01398 | 8.566745 | 0.013763 | 9.07E-01 | 9.16E-01 | Healthy control | Liver |
| Gckr | -1.85403 | 9.853637 | 432.7179 | 5.10E-30 | 5.00E-29 | Healthy control | Liver |
| Gcm1 | 1.340404 | 1.311861 | 4.605871 | 3.65E-02 | 4.50E-02 | Infection | Liver |
| Gcm2 | 1.121475 | 1.245291 | 3.566178 | 6.74E-02 | 8.00E-02 | Infection | Liver |
| Sv2a | 3.35359 | 2.891695 | 172.1113 | 3.91E-24 | 2.08E-23 | Infection | Liver |
| Gcnt4 | -0.00674 | 4.045977 | 0.002365 | 9.61E-01 | 9.65E-01 | Healthy control | Liver |
| Irf8 | 3.348419 | 7.675569 | 1087.928 | 1.33E-41 | 9.53E-40 | Infection | Liver |
| Cdca5 | 3.344297 | 3.947467 | 293.4925 | 4.56E-28 | 3.67E-27 | Infection | Liver |
| Cfap69 | 3.336209 | 1.725829 | 41.46515 | 1.48E-08 | 2.79E-08 | Infection | Liver |
| Csf3r | 3.335377 | 5.251907 | 670.6002 | 3.00E-35 | 6.09E-34 | Infection | Liver |
| Gdf15 | 0.070111 | 4.672172 | 0.215031 | 6.44E-01 | 6.68E-01 | Infection | Liver |
| Gdf2 | -0.67059 | 7.4717 | 60.71771 | 7.95E-11 | 1.68E-10 | Healthy control | Liver |
| Gdf5 | 1.386046 | 1.279953 | 5.317127 | 2.61E-02 | 3.27E-02 | Infection | Liver |
| Eme1 | 3.333299 | 2.698392 | 105.8777 | 9.53E-18 | 2.99E-17 | Infection | Liver |
| Gdf9 | 1.169497 | 1.449781 | 5.730416 | 1.87E-02 | 2.39E-02 | Infection | Liver |
| Hoxd9 | 3.331937 | 1.717602 | 43.76954 | 8.08E-09 | 1.54E-08 | Infection | Liver |
| Fstl1 | 3.330038 | 8.270653 | 219.5253 | 2.11E-21 | 8.70E-21 | Infection | Liver |
| Gdpd2 | 0.698785 | 1.446769 | 1.824382 | 1.80E-01 | 2.02E-01 | Infection | Liver |
| Gem | 0.61875 | 5.825694 | 23.25445 | 9.28E-06 | 1.51E-05 | Infection | Liver |
| Gen1 | 0.819502 | 2.897159 | 19.00482 | 4.26E-05 | 6.68E-05 | Infection | Liver |
| Gfap | 0.906368 | 3.837403 | 18.74302 | 5.49E-05 | 8.53E-05 | Infection | Liver |
| Scn5a | 3.326323 | 1.927634 | 37.25503 | 5.41E-08 | 9.87E-08 | Infection | Liver |
| Srpx2 | 3.322157 | 6.624493 | 606.8577 | 4.41E-34 | 6.98E-33 | Infection | Liver |
| Gfpt1 | -0.5517 | 6.784028 | 43.8993 | 8.74E-09 | 1.66E-08 | Healthy control | Liver |
| Ncf1 | 3.32213 | 7.453481 | 997.8006 | 1.79E-40 | 9.54E-39 | Infection | Liver |
| Gfra1 | -0.99903 | 9.104653 | 136.8191 | 1.77E-17 | 5.47E-17 | Healthy control | Liver |
| Gfra2 | 1.222584 | 5.783421 | 125.1686 | 1.23E-16 | 3.58E-16 | Infection | Liver |
| Gfra3 | 0.98146 | 1.300114 | 2.964952 | 9.14E-02 | 1.07E-01 | Infection | Liver |
| Ggnbp1 | -0.97083 | 4.585975 | 59.84454 | 1.02E-10 | 2.15E-10 | Healthy control | Liver |
| Ggnbp2 | -0.72025 | 8.462056 | 261.922 | 3.42E-24 | 1.82E-23 | Healthy control | Liver |

| | | | | | | | |
|---|---|---|---|---|---|---|---|
| Ggps1 | 0.329396 | 5.127254 | 12.54121 | 7.54E-04 | 1.09E-03 | Infection | Liver |
| Gh | 0.556392 | 1.295695 | 0.968151 | 3.29E-01 | 3.56E-01 | Infection | Liver |
| Ghr | -1.96269 | 11.31984 | 656.779 | 3.70E-35 | 7.38E-34 | Healthy control | Liver |
| Ghrh | 0.411448 | 1.298697 | 0.503068 | 4.81E-01 | 5.08E-01 | Infection | Liver |
| Ghrhr | 0.556772 | 1.209714 | 0.944751 | 3.42E-01 | 3.69E-01 | Infection | Liver |
| Ghrl | 0.485118 | 1.267406 | 0.677723 | 4.15E-01 | 4.43E-01 | Infection | Liver |
| Ly6k | 3.312722 | 2.505271 | 117.2435 | 1.01E-18 | 3.40E-18 | Infection | Liver |
| Gigyf2 | -0.06641 | 7.078622 | 1.903536 | 1.73E-01 | 1.94E-01 | Healthy control | Liver |
| Gimap4 | 0.540134 | 6.597476 | 40.02583 | 2.88E-08 | 5.32E-08 | Infection | Liver |
| Gip | 0.95862 | 1.230148 | 2.425909 | 1.30E-01 | 1.49E-01 | Infection | Liver |
| Gipc1 | 0.265432 | 8.046987 | 45.63702 | 5.19E-09 | 1.00E-08 | Infection | Liver |
| Il5 | 3.31028 | 1.828935 | 32.80887 | 2.94E-07 | 5.19E-07 | Infection | Liver |
| Myc | 3.303729 | 6.482084 | 788.0049 | 2.40E-37 | 6.95E-36 | Infection | Liver |
| Gja3 | 0.958019 | 1.246839 | 1.903187 | 1.77E-01 | 1.98E-01 | Infection | Liver |
| Gja4 | 0.375197 | 4.98687 | 13.69918 | 4.53E-04 | 6.64E-04 | Infection | Liver |
| Blnk | 3.302955 | 6.176296 | 566.5106 | 3.61E-33 | 5.10E-32 | Infection | Liver |
| Gja8 | 0.227546 | 1.215836 | 0.151362 | 7.01E-01 | 7.22E-01 | Infection | Liver |
| Gjb1 | -1.46387 | 10.42175 | 269.3081 | 1.67E-24 | 9.16E-24 | Healthy control | Liver |
| Gjb2 | -2.69723 | 10.53391 | 1034.523 | 5.72E-41 | 3.54E-39 | Healthy control | Liver |
| Gjb6 | 1.383201 | 1.38029 | 5.175331 | 2.59E-02 | 3.24E-02 | Infection | Liver |
| Card9 | 3.301702 | 4.536077 | 428.9926 | 9.95E-31 | 1.06E-29 | Infection | Liver |
| Gjd2 | 0.431568 | 1.233137 | 0.512652 | 4.80E-01 | 5.07E-01 | Infection | Liver |
| Gk | -1.27228 | 8.322596 | 405.2446 | 3.13E-29 | 2.82E-28 | Healthy control | Liver |
| Gkn2 | 0.639125 | 1.266505 | 1.147514 | 2.90E-01 | 3.16E-01 | Infection | Liver |
| Gla | 0.991505 | 5.693033 | 154.9744 | 1.14E-18 | 3.84E-18 | Infection | Liver |
| Glb1 | 0.900186 | 6.957151 | 131.9608 | 3.90E-17 | 1.18E-16 | Infection | Liver |
| Gldc | -1.79706 | 8.274004 | 114.4297 | 1.99E-15 | 5.36E-15 | Healthy control | Liver |
| Gldn | 1.820207 | 1.395984 | 14.24804 | 3.35E-04 | 4.96E-04 | Infection | Liver |
| Glg1 | -0.04637 | 7.822085 | 0.346099 | 5.58E-01 | 5.85E-01 | Healthy control | Liver |
| Tnfsf11 | 3.297301 | 1.879088 | 56.86809 | 1.29E-10 | 2.69E-10 | Infection | Liver |
| Cfap45 | 3.29702 | 2.496687 | 105.0759 | 1.98E-17 | 6.07E-17 | Infection | Liver |
| Ighd | 3.294699 | 3.902834 | 131.5075 | 1.76E-18 | 5.83E-18 | Infection | Liver |
| Cxcl5 | 3.290172 | 1.93862 | 40.98848 | 1.31E-08 | 2.48E-08 | Infection | Liver |
| Glmp | -0.29408 | 6.95164 | 19.6449 | 3.78E-05 | 5.94E-05 | Healthy control | Liver |
| Glo1 | -1.6539 | 9.315199 | 661.3872 | 3.03E-35 | 6.11E-34 | Healthy control | Liver |
| Glp1r | -0.64568 | 1.36258 | 1.52447 | 2.20E-01 | 2.45E-01 | Healthy control | Liver |
| Cacna2d1 | 3.285795 | 5.30365 | 339.4845 | 4.55E-27 | 3.20E-26 | Infection | Liver |
| Glra2 | 1.158234 | 1.327084 | 3.204311 | 7.85E-02 | 9.24E-02 | Infection | Liver |
| Glra3 | 1.946032 | 1.317481 | 8.330729 | 6.09E-03 | 8.17E-03 | Infection | Liver |
| Glud1 | -0.97455 | 11.70184 | 156.6288 | 8.74E-19 | 2.96E-18 | Healthy control | Liver |
| Glyat | -1.87384 | 10.57475 | 317.0054 | 2.36E-26 | 1.56E-25 | Healthy control | Liver |
| Gm2a | -0.59686 | 9.612361 | 140.0253 | 1.07E-17 | 3.33E-17 | Healthy control | Liver |
| Cd300lf | 3.284684 | 5.238523 | 661.129 | 4.53E-35 | 8.93E-34 | Infection | Liver |
| Gmfb | -0.23386 | 6.928323 | 14.42476 | 3.29E-04 | 4.88E-04 | Healthy control | Liver |
| Gmnc | 1.658753 | 1.292905 | 5.725638 | 2.11E-02 | 2.68E-02 | Infection | Liver |
| Gmnn | -0.89971 | 5.980463 | 98.13658 | 1.73E-14 | 4.43E-14 | Healthy control | Liver |
| Gna11 | -0.63171 | 6.470767 | 95.53205 | 2.87E-14 | 7.28E-14 | Healthy control | Liver |
| Gna12 | -0.84627 | 7.9781 | 217.7501 | 3.55E-22 | 1.57E-21 | Healthy control | Liver |
| Gna13 | 0.782595 | 6.54516 | 180.5335 | 3.29E-20 | 1.24E-19 | Infection | Liver |
| Gna14 | -0.72423 | 4.57694 | 21.63499 | 1.73E-05 | 2.78E-05 | Healthy control | Liver |

| | | | | | | | |
|---|---|---|---|---|---|---|---|
| Fosl2 | 3.279158 | 6.466305 | 1170.917 | 1.93E-42 | 1.59E-40 | Infection | Liver |
| Gnai2 | 0.657323 | 9.94199 | 79.4028 | 8.78E-13 | 2.05E-12 | Infection | Liver |
| Gnai3 | -0.1124 | 8.674406 | 5.876222 | 1.82E-02 | 2.33E-02 | Healthy control | Liver |
| Gnal | -0.07829 | 1.699313 | 0.038646 | 8.44E-01 | 8.57E-01 | Healthy control | Liver |
| Thbs2 | 3.27774 | 6.489386 | 188.4344 | 6.34E-20 | 2.36E-19 | Infection | Liver |
| Gnaq | -0.26198 | 7.722654 | 25.21208 | 4.44E-06 | 7.36E-06 | Healthy control | Liver |
| Gnat1 | -1.01945 | 2.607541 | 24.64762 | 3.47E-06 | 5.77E-06 | Healthy control | Liver |
| Gnat2 | -0.1499 | 2.334223 | 0.3356 | 5.64E-01 | 5.90E-01 | Healthy control | Liver |
| Gnat3 | 1.603733 | 1.397407 | 9.729759 | 2.67E-03 | 3.69E-03 | Infection | Liver |
| Rcor2 | 3.268859 | 3.307894 | 214.0957 | 3.24E-26 | 2.11E-25 | Infection | Liver |
| Gnb1 | -1.13583 | 8.351871 | 171.6722 | 1.06E-19 | 3.89E-19 | Healthy control | Liver |
| Gnb3 | 1.898037 | 1.450952 | 15.41266 | 1.85E-04 | 2.79E-04 | Infection | Liver |
| Gnrh1 | 0.40606 | 2.737985 | 5.587699 | 2.07E-02 | 2.63E-02 | Infection | Liver |
| Ntrk3 | 3.268089 | 3.15016 | 225.9416 | 2.96E-27 | 2.14E-26 | Infection | Liver |
| Golga2 | 0.879148 | 7.181819 | 322.8856 | 1.46E-26 | 9.77E-26 | Infection | Liver |
| Golga3 | 0.68025 | 7.060334 | 240.4661 | 3.01E-23 | 1.44E-22 | Infection | Liver |
| Golga5 | 0.408902 | 6.939355 | 50.41418 | 1.30E-09 | 2.59E-09 | Infection | Liver |
| Gopc | -0.57603 | 7.604959 | 105.5606 | 4.02E-15 | 1.07E-14 | Healthy control | Liver |
| Gorasp1 | -0.6044 | 6.220786 | 75.85545 | 1.98E-12 | 4.53E-12 | Healthy control | Liver |
| Gosr1 | 0.221301 | 7.432251 | 12.27047 | 8.50E-04 | 1.22E-03 | Infection | Liver |
| Got2 | -1.59654 | 10.21768 | 404.1469 | 3.36E-29 | 3.01E-28 | Healthy control | Liver |
| Cdc6 | 3.264444 | 2.24858 | 101.0167 | 1.66E-16 | 4.79E-16 | Infection | Liver |
| Gpam | -0.21388 | 9.931591 | 4.946682 | 2.97E-02 | 3.69E-02 | Healthy control | Liver |
| Gpat2 | 0.632688 | 3.546062 | 13.84257 | 4.25E-04 | 6.24E-04 | Infection | Liver |
| Ly86 | 3.255543 | 6.819864 | 822.8662 | 5.89E-38 | 1.88E-36 | Infection | Liver |
| Meox1 | 3.255067 | 3.207536 | 137.9696 | 2.18E-20 | 8.34E-20 | Infection | Liver |
| Gpc4 | -1.37145 | 10.14136 | 604.7004 | 4.01E-34 | 6.47E-33 | Healthy control | Liver |
| Cenpf | 3.254905 | 4.127437 | 248.0479 | 5.54E-25 | 3.18E-24 | Infection | Liver |
| Gpd1 | -1.99573 | 8.618364 | 400.4823 | 4.33E-29 | 3.83E-28 | Healthy control | Liver |
| Gper1 | 0.064573 | 1.801933 | 0.030362 | 8.62E-01 | 8.74E-01 | Infection | Liver |
| Gphn | -1.70885 | 8.286507 | 560.6144 | 3.52E-33 | 5.01E-32 | Healthy control | Liver |
| Gpi1 | -0.44921 | 8.271665 | 64.38843 | 3.11E-11 | 6.71E-11 | Healthy control | Liver |
| Gpm6a | 1.959233 | 2.403333 | 37.11122 | 1.67E-08 | 3.13E-08 | Infection | Liver |
| Fgf23 | 3.245638 | 1.590405 | 34.29613 | 1.71E-07 | 3.06E-07 | Infection | Liver |
| Gpr101 | 1.190626 | 1.333113 | 4.425169 | 3.95E-02 | 4.85E-02 | Infection | Liver |
| Gpr107 | 0.493304 | 6.935624 | 108.1663 | 2.47E-15 | 6.62E-15 | Infection | Liver |
| Gpr139 | 0.422285 | 1.264931 | 0.61415 | 4.37E-01 | 4.65E-01 | Infection | Liver |
| Cd274 | 3.244569 | 7.201635 | 966.0037 | 4.82E-40 | 2.30E-38 | Infection | Liver |
| Gpr143 | 1.302092 | 1.3816 | 4.468144 | 3.80E-02 | 4.68E-02 | Infection | Liver |
| Gpr149 | 0.781141 | 1.268554 | 1.524973 | 2.24E-01 | 2.49E-01 | Infection | Liver |
| Gpr17 | 1.578281 | 1.242099 | 8.675781 | 5.80E-03 | 7.78E-03 | Infection | Liver |
| Gpr183 | 0.780866 | 4.052778 | 19.27932 | 4.44E-05 | 6.95E-05 | Infection | Liver |
| Gpr20 | 0.661989 | 1.373004 | 1.323814 | 2.53E-01 | 2.79E-01 | Infection | Liver |
| Camk2a | 3.239022 | 1.918232 | 49.66636 | 7.16E-10 | 1.45E-09 | Infection | Liver |
| Gpr26 | 0.743429 | 1.186681 | 2.221006 | 1.57E-01 | 1.78E-01 | Infection | Liver |
| Nfatc2 | 3.236768 | 4.113639 | 275.0957 | 3.08E-26 | 2.01E-25 | Infection | Liver |
| Gpr37l1 | 1.136822 | 1.285782 | 4.043522 | 4.97E-02 | 6.02E-02 | Infection | Liver |
| Slc1a3 | 3.230659 | 3.826578 | 267.1174 | 3.11E-27 | 2.24E-26 | Infection | Liver |
| Gpr4 | -0.34135 | 2.482773 | 2.481683 | 1.19E-01 | 1.37E-01 | Healthy control | Liver |
| Gpr50 | 0.248991 | 1.326997 | 0.16398 | 6.87E-01 | 7.09E-01 | Infection | Liver |

| | | | | | | | |
|---|---|---|---|---|---|---|---|
| Gpr6 | 0.008943 | 1.205327 | 0.000447 | 9.83E-01 | 9.85E-01 | Infection | Liver |
| Ccl19 | 3.223741 | 2.195062 | 79.94807 | 4.65E-14 | 1.17E-13 | Infection | Liver |
| Gpr83 | 1.568144 | 1.448306 | 7.711158 | 6.80E-03 | 9.08E-03 | Infection | Liver |
| Ifnlr1 | 3.222619 | 3.014047 | 130.5684 | 3.36E-20 | 1.26E-19 | Infection | Liver |
| Stat4 | 3.221334 | 3.958138 | 335.4447 | 2.22E-29 | 2.04E-28 | Infection | Liver |
| Il18rap | 3.220955 | 4.594665 | 297.8421 | 3.96E-26 | 2.55E-25 | Infection | Liver |
| Gprc6a | 1.412486 | 1.30053 | 5.4186 | 2.42E-02 | 3.06E-02 | Infection | Liver |
| Gps1 | -1.61894 | 8.369064 | 1327.599 | 3.17E-44 | 4.07E-42 | Healthy control | Liver |
| Gzmk | 3.216585 | 1.653425 | 40.0061 | 2.21E-08 | 4.11E-08 | Infection | Liver |
| Gpsm2 | -1.36358 | 6.417688 | 350.2575 | 1.74E-27 | 1.31E-26 | Healthy control | Liver |
| Gpt2 | -1.1302 | 9.75188 | 113.0418 | 1.12E-15 | 3.07E-15 | Healthy control | Liver |
| Gpx2 | 1.803337 | 1.30787 | 10.31497 | 2.30E-03 | 3.18E-03 | Infection | Liver |
| Gpx5 | 0.192962 | 1.358971 | 0.120735 | 7.29E-01 | 7.49E-01 | Infection | Liver |
| Gramd1a | 1.37623 | 7.173297 | 507.4461 | 5.98E-32 | 7.25E-31 | Infection | Liver |
| Nrip3 | 3.213822 | 1.750098 | 46.28088 | 3.01E-09 | 5.89E-09 | Infection | Liver |
| Grb10 | 1.601916 | 6.440798 | 111.1586 | 1.86E-15 | 5.03E-15 | Infection | Liver |
| Grb2 | 1.392903 | 7.493556 | 477.2971 | 3.34E-31 | 3.77E-30 | Infection | Liver |
| Grb7 | -1.65356 | 7.578274 | 165.3163 | 3.78E-19 | 1.31E-18 | Healthy control | Liver |
| Greb1 | 1.887698 | 1.326515 | 10.40424 | 2.26E-03 | 3.14E-03 | Infection | Liver |
| Grem1 | 1.663128 | 1.25483 | 7.875739 | 7.92E-03 | 1.05E-02 | Infection | Liver |
| Bpifb1 | 3.212857 | 1.569301 | 28.80985 | 1.26E-06 | 2.14E-06 | Infection | Liver |
| Grhl3 | 0.284845 | 1.692749 | 0.672321 | 4.14E-01 | 4.41E-01 | Infection | Liver |
| Gria1 | -0.06707 | 1.300641 | 0.013751 | 9.07E-01 | 9.16E-01 | Healthy control | Liver |
| Cdh3 | 3.210677 | 2.403079 | 97.13388 | 1.99E-16 | 5.71E-16 | Infection | Liver |
| Mybpc3 | 3.210416 | 1.761843 | 45.69179 | 3.97E-09 | 7.71E-09 | Infection | Liver |
| Igsf9b | 3.207395 | 1.659656 | 66.52472 | 1.16E-11 | 2.56E-11 | Infection | Liver |
| Grid1 | -1.90426 | 3.529511 | 102.6753 | 9.42E-16 | 2.60E-15 | Healthy control | Liver |
| Grid2 | 1.867289 | 1.380514 | 9.290308 | 3.45E-03 | 4.71E-03 | Infection | Liver |
| Grifin | 0.85688 | 1.39696 | 2.80201 | 9.80E-02 | 1.14E-01 | Infection | Liver |
| Grik1 | 1.713468 | 1.447516 | 7.477402 | 7.76E-03 | 1.03E-02 | Infection | Liver |
| Grik2 | 1.769949 | 1.652252 | 16.6114 | 8.72E-05 | 1.34E-04 | Infection | Liver |
| Grik3 | 0.146157 | 1.701332 | 0.132428 | 7.17E-01 | 7.37E-01 | Infection | Liver |
| Grik4 | -0.76714 | 2.805034 | 11.21083 | 1.25E-03 | 1.78E-03 | Healthy control | Liver |
| Grin1 | 1.799569 | 1.318953 | 10.36279 | 2.18E-03 | 3.03E-03 | Infection | Liver |
| Azin2 | 3.204499 | 2.546054 | 129.9029 | 2.73E-20 | 1.04E-19 | Infection | Liver |
| Grin2b | -0.37986 | 1.234435 | 0.4217 | 5.21E-01 | 5.49E-01 | Healthy control | Liver |
| Ccn2 | 3.20134 | 6.538001 | 228.6578 | 5.68E-22 | 2.46E-21 | Infection | Liver |
| Fabp3 | 3.201238 | 1.809831 | 86.51274 | 4.48E-14 | 1.13E-13 | Infection | Liver |
| Grk2 | 0.524817 | 8.599822 | 164.3629 | 2.91E-19 | 1.02E-18 | Infection | Liver |
| Eya4 | 3.200789 | 3.673433 | 206.2135 | 4.53E-24 | 2.38E-23 | Infection | Liver |
| Grk5 | 1.035355 | 5.055262 | 75.86092 | 2.04E-12 | 4.66E-12 | Infection | Liver |
| Grm1 | 1.122699 | 1.274462 | 4.066571 | 4.99E-02 | 6.04E-02 | Infection | Liver |
| Racgap1 | 3.199109 | 4.845468 | 304.7243 | 5.03E-26 | 3.20E-25 | Infection | Liver |
| Grm3 | 1.361976 | 1.232304 | 5.595126 | 2.45E-02 | 3.08E-02 | Infection | Liver |
| a | 3.19884 | 1.919863 | 74.65368 | 7.25E-13 | 1.71E-12 | Infection | Liver |
| Grm6 | 1.429396 | 1.909805 | 18.10639 | 3.95E-05 | 6.21E-05 | Infection | Liver |
| Grm8 | 1.019849 | 1.692754 | 7.162859 | 8.38E-03 | 1.11E-02 | Infection | Liver |
| Grn | 1.615818 | 10.19442 | 894.3506 | 4.35E-39 | 1.68E-37 | Infection | Liver |
| Grp | 1.935892 | 1.291795 | 9.601658 | 3.23E-03 | 4.43E-03 | Infection | Liver |
| Grpel1 | -1.50216 | 8.863786 | 428.0649 | 6.89E-30 | 6.65E-29 | Healthy control | Liver |

| | | | | | | | |
|---|---|---|---|---|---|---|---|
| Grpr | 0.529616 | 1.257268 | 0.69643 | 4.09E-01 | 4.37E-01 | Infection | Liver |
| Grxcr1 | 1.831803 | 1.266962 | 9.252364 | 4.16E-03 | 5.65E-03 | Infection | Liver |
| Grxcr2 | 0.810445 | 1.254799 | 1.707334 | 1.99E-01 | 2.22E-01 | Infection | Liver |
| Gsc | -0.25594 | 1.220568 | 0.201635 | 6.57E-01 | 6.80E-01 | Healthy control | Liver |
| Gsc2 | 1.832455 | 1.501861 | 10.14767 | 2.01E-03 | 2.80E-03 | Infection | Liver |
| Gsdma3 | 0.474023 | 1.269046 | 0.553075 | 4.61E-01 | 4.89E-01 | Infection | Liver |
| Gsdmd | 1.13258 | 9.099341 | 554.1627 | 4.87E-33 | 6.74E-32 | Infection | Liver |
| Gsdme | 0.300161 | 4.265297 | 7.88488 | 6.64E-03 | 8.88E-03 | Infection | Liver |
| Gsk3a | -0.73606 | 8.999619 | 419.7111 | 1.19E-29 | 1.12E-28 | Healthy control | Liver |
| Gsk3b | -0.28897 | 6.598646 | 32.43495 | 3.43E-07 | 6.01E-07 | Healthy control | Liver |
| Gsta1 | -2.32039 | 6.350058 | 287.5246 | 3.54E-25 | 2.06E-24 | Healthy control | Liver |
| Gstk1 | -1.82187 | 8.685707 | 459.294 | 9.76E-31 | 1.04E-29 | Healthy control | Liver |
| Gstp1 | -0.73941 | 9.371323 | 81.04972 | 6.08E-13 | 1.44E-12 | Healthy control | Liver |
| Gsx1 | -0.6585 | 1.256159 | 1.170551 | 2.86E-01 | 3.12E-01 | Healthy control | Liver |
| Gsx2 | 1.130925 | 1.291097 | 3.015138 | 8.86E-02 | 1.04E-01 | Infection | Liver |
| Gtf2a1 | -0.03363 | 7.521256 | 0.486313 | 4.88E-01 | 5.15E-01 | Healthy control | Liver |
| Gtf2a2 | 0.135199 | 7.925181 | 3.721012 | 5.82E-02 | 6.97E-02 | Infection | Liver |
| Gtf2b | -0.03271 | 5.935261 | 0.146226 | 7.03E-01 | 7.25E-01 | Healthy control | Liver |
| Gtf2e1 | -1.22237 | 5.454363 | 136.9104 | 1.91E-17 | 5.86E-17 | Healthy control | Liver |
| Gtf2e2 | 0.746723 | 4.970012 | 82.36132 | 4.76E-13 | 1.14E-12 | Infection | Liver |
| Gtf2h1 | 0.096237 | 6.401413 | 2.882102 | 9.45E-02 | 1.10E-01 | Infection | Liver |
| Gtf2i | -0.77285 | 9.159435 | 271.0408 | 1.42E-24 | 7.83E-24 | Healthy control | Liver |
| Gtpbp2 | 0.392945 | 7.662077 | 96.1415 | 2.52E-14 | 6.41E-14 | Infection | Liver |
| Nlgn2 | 3.196774 | 5.918508 | 477.1112 | 4.67E-31 | 5.16E-30 | Infection | Liver |
| Gucy1a1 | 0.524592 | 7.223646 | 12.58574 | 7.61E-04 | 1.10E-03 | Infection | Liver |
| Gucy2e | 1.574572 | 1.368406 | 10.50095 | 1.84E-03 | 2.58E-03 | Infection | Liver |
| Gucy2f | 1.740776 | 1.291031 | 6.811677 | 1.29E-02 | 1.68E-02 | Infection | Liver |
| Gusb | 1.165649 | 4.536576 | 89.71787 | 1.05E-13 | 2.58E-13 | Infection | Liver |
| Gyg | 1.005124 | 5.323389 | 69.38752 | 9.26E-12 | 2.05E-11 | Infection | Liver |
| Gys2 | -1.34244 | 7.869881 | 170.4469 | 1.26E-19 | 4.58E-19 | Healthy control | Liver |
| Pou3f1 | 3.196506 | 1.785664 | 44.98668 | 3.13E-09 | 6.10E-09 | Infection | Liver |
| Gzmb | 1.546288 | 2.946355 | 31.55103 | 3.08E-07 | 5.42E-07 | Infection | Liver |
| Traf3ip3 | 3.195021 | 5.124847 | 369.4544 | 4.97E-28 | 3.99E-27 | Infection | Liver |
| Alox12e | 3.193985 | 1.857563 | 53.88691 | 3.09E-10 | 6.35E-10 | Infection | Liver |
| H1foo | 1.452624 | 1.35303 | 6.211022 | 1.52E-02 | 1.97E-02 | Infection | Liver |
| Trem1 | 3.190096 | 1.63492 | 49.02356 | 1.40E-09 | 2.77E-09 | Infection | Liver |
| H2afv | -0.39699 | 5.448768 | 19.92362 | 3.39E-05 | 5.35E-05 | Healthy control | Liver |
| H2afx | 1.803774 | 6.605239 | 414.0125 | 1.77E-29 | 1.66E-28 | Infection | Liver |
| H2afy | -0.44495 | 7.890187 | 97.6783 | 1.85E-14 | 4.73E-14 | Healthy control | Liver |
| H2afz | 0.360089 | 7.69978 | 40.88 | 2.20E-08 | 4.10E-08 | Infection | Liver |
| Hace1 | 0.415355 | 5.107475 | 21.16726 | 2.08E-05 | 3.32E-05 | Infection | Liver |
| Hacl1 | -2.12964 | 10.87234 | 286.6864 | 6.10E-25 | 3.49E-24 | Healthy control | Liver |
| Hadh | -1.91796 | 10.80455 | 634.7568 | 9.91E-35 | 1.80E-33 | Healthy control | Liver |
| Hadhb | -2.1717 | 9.397328 | 2303.892 | 1.50E-51 | 4.85E-48 | Healthy control | Liver |
| Hal | -1.86384 | 11.04402 | 64.49979 | 5.32E-11 | 1.14E-10 | Healthy control | Liver |
| Hamp | -2.87201 | 12.93089 | 589.3115 | 8.39E-34 | 1.28E-32 | Healthy control | Liver |
| Hand1 | 1.095228 | 1.27904 | 3.266567 | 7.79E-02 | 9.17E-02 | Infection | Liver |
| Hand2 | 1.152713 | 5.239776 | 80.91956 | 6.46E-13 | 1.53E-12 | Infection | Liver |
| Hao1 | -2.15181 | 9.719571 | 385.9784 | 1.60E-28 | 1.34E-27 | Healthy control | Liver |
| Hap1 | 0.15744 | 3.902354 | 1.497036 | 2.26E-01 | 2.50E-01 | Infection | Liver |

| | | | | | | | |
|---|---|---|---|---|---|---|---|
| Hapln1 | 0.610789 | 1.955758 | 3.24945 | 7.39E-02 | 8.73E-02 | Infection | Liver |
| Edil3 | 3.187908 | 1.67756 | 35.77405 | 1.03E-07 | 1.85E-07 | Infection | Liver |
| Gja5 | 3.184355 | 3.430741 | 94.2039 | 1.28E-15 | 3.50E-15 | Infection | Liver |
| Hat1 | -0.45144 | 6.435366 | 54.48233 | 4.20E-10 | 8.56E-10 | Healthy control | Liver |
| Bub1 | 3.181702 | 2.909548 | 120.5084 | 4.16E-19 | 1.44E-18 | Infection | Liver |
| Izumo1r | 3.179302 | 2.779786 | 141.3873 | 1.31E-21 | 5.45E-21 | Infection | Liver |
| Hba-x | 0.492862 | 1.207241 | 0.8427 | 3.69E-01 | 3.96E-01 | Infection | Liver |
| Gas1 | 3.176012 | 8.937325 | 293.7351 | 1.65E-24 | 9.06E-24 | Infection | Liver |
| Hc | 0.362406 | 11.64381 | 11.6152 | 1.14E-03 | 1.63E-03 | Infection | Liver |
| Hcar2 | 1.384344 | 1.682697 | 14.37762 | 2.32E-04 | 3.46E-04 | Infection | Liver |
| Hcfc1 | -0.68917 | 7.925443 | 242.2208 | 2.50E-23 | 1.21E-22 | Healthy control | Liver |
| Hck | 1.740885 | 6.042789 | 235.9406 | 5.00E-23 | 2.35E-22 | Infection | Liver |
| Hcn1 | 1.609965 | 1.376531 | 6.663473 | 1.20E-02 | 1.56E-02 | Infection | Liver |
| Hcn2 | 0.062225 | 1.477142 | 0.016746 | 8.97E-01 | 9.07E-01 | Infection | Liver |
| Hcn3 | 1.140354 | 1.708336 | 10.00845 | 1.93E-03 | 2.70E-03 | Infection | Liver |
| Hcn4 | 0.040514 | 1.400521 | 0.006185 | 9.37E-01 | 9.44E-01 | Infection | Liver |
| Hcrt | 1.645623 | 1.413442 | 7.229948 | 8.92E-03 | 1.18E-02 | Infection | Liver |
| Hcrtr1 | 0.971068 | 1.267779 | 2.498119 | 1.22E-01 | 1.40E-01 | Infection | Liver |
| Hoxc6 | 3.171917 | 3.10352 | 135.9071 | 2.88E-20 | 1.09E-19 | Infection | Liver |
| Hdac1 | 0.864674 | 6.558011 | 257.2503 | 5.47E-24 | 2.85E-23 | Infection | Liver |
| Hdac2 | 0.512907 | 6.444455 | 78.11204 | 1.18E-12 | 2.74E-12 | Infection | Liver |
| Hdac3 | -1.03586 | 7.269821 | 447.532 | 2.03E-30 | 2.09E-29 | Healthy control | Liver |
| Hdac4 | 0.417553 | 6.745123 | 55.22759 | 3.42E-10 | 7.01E-10 | Infection | Liver |
| Hdac5 | -0.5325 | 6.792787 | 35.48025 | 1.24E-07 | 2.22E-07 | Healthy control | Liver |
| Hdac6 | -0.81537 | 6.726097 | 229.1708 | 1.01E-22 | 4.66E-22 | Healthy control | Liver |
| Hdac8 | 0.197723 | 4.883168 | 5.45381 | 2.27E-02 | 2.88E-02 | Infection | Liver |
| Hdac9 | 1.440089 | 3.446098 | 58.78012 | 1.14E-10 | 2.40E-10 | Infection | Liver |
| Hdc | 1.997357 | 3.15222 | 97.2098 | 3.65E-15 | 9.71E-15 | Infection | Liver |
| Hdgfl2 | 0.078136 | 7.119556 | 3.739056 | 5.76E-02 | 6.90E-02 | Infection | Liver |
| Hecw2 | 0.054139 | 5.621153 | 0.309702 | 5.80E-01 | 6.05E-01 | Infection | Liver |
| Hells | 1.839361 | 4.558938 | 298.2311 | 1.78E-25 | 1.06E-24 | Infection | Liver |
| Herc6 | -0.11176 | 6.60788 | 1.188185 | 2.80E-01 | 3.06E-01 | Healthy control | Liver |
| Hes1 | -0.42346 | 7.222956 | 26.32757 | 2.95E-06 | 4.91E-06 | Healthy control | Liver |
| Hes2 | 0.220362 | 1.21933 | 0.153259 | 6.98E-01 | 7.20E-01 | Infection | Liver |
| Hes5 | 1.374627 | 1.265776 | 4.147872 | 4.82E-02 | 5.85E-02 | Infection | Liver |
| Hes6 | -0.48307 | 7.474419 | 52.22844 | 7.80E-10 | 1.57E-09 | Healthy control | Liver |
| Hesx1 | 1.978396 | 1.541444 | 13.06377 | 5.05E-04 | 7.37E-04 | Infection | Liver |
| Hexa | 1.087773 | 8.541114 | 173.3258 | 8.49E-20 | 3.14E-19 | Infection | Liver |
| Src | 3.169507 | 3.855376 | 410.5486 | 1.75E-32 | 2.27E-31 | Infection | Liver |
| Lamb1 | 3.163648 | 7.484843 | 433.9632 | 1.83E-29 | 1.71E-28 | Infection | Liver |
| Heyl | 1.439016 | 4.375215 | 107.7148 | 3.16E-15 | 8.45E-15 | Infection | Liver |
| Hfe | -0.32093 | 4.287173 | 8.736929 | 4.39E-03 | 5.94E-03 | Healthy control | Liver |
| Dnmt3l | 3.163081 | 1.749338 | 52.97376 | 3.82E-10 | 7.82E-10 | Infection | Liver |
| Hgf | -0.33739 | 7.246002 | 13.07197 | 5.95E-04 | 8.62E-04 | Healthy control | Liver |
| Hgs | -0.54149 | 7.530323 | 140.7781 | 9.52E-18 | 2.98E-17 | Healthy control | Liver |
| Hhex | -0.97645 | 8.897458 | 145.2481 | 4.76E-18 | 1.52E-17 | Healthy control | Liver |
| Hibadh | -1.8327 | 10.19483 | 414.9022 | 1.63E-29 | 1.52E-28 | Healthy control | Liver |
| Hic1 | 1.644835 | 6.647589 | 82.47004 | 7.71E-13 | 1.81E-12 | Infection | Liver |
| Hif1a | -0.48882 | 7.513995 | 59.25211 | 1.17E-10 | 2.44E-10 | Healthy control | Liver |
| Hif1an | -0.1661 | 6.359357 | 9.722082 | 2.74E-03 | 3.78E-03 | Healthy control | Liver |

| | | | | | | | |
|---|---|---|---|---|---|---|---|
| Hif3a | 1.064979 | 1.402915 | 4.183961 | 4.39E-02 | 5.35E-02 | Infection | Liver |
| Hikeshi | -1.03205 | 2.62063 | 14.10426 | 3.11E-04 | 4.61E-04 | Healthy control | Liver |
| Hip1r | 0.098016 | 7.503896 | 0.931694 | 3.38E-01 | 3.64E-01 | Infection | Liver |
| Hipk2 | -1.04542 | 8.246766 | 535.1562 | 1.32E-32 | 1.73E-31 | Healthy control | Liver |
| Hira | 0.087069 | 5.735775 | 1.592192 | 2.12E-01 | 2.35E-01 | Infection | Liver |
| Hist1h2ba | 0.407243 | 1.171985 | 1.475336 | 2.50E-01 | 2.76E-01 | Infection | Liver |
| Hist1h2bc | -2.81969 | 9.549488 | 1338.391 | 2.48E-44 | 3.27E-42 | Healthy control | Liver |
| Hist1h4h | 1.126192 | 6.423822 | 118.2185 | 4.02E-16 | 1.13E-15 | Infection | Liver |
| Ikzf2 | 3.16248 | 4.703376 | 322.214 | 1.01E-26 | 6.82E-26 | Infection | Liver |
| Hist2h2be | -0.22961 | 4.165893 | 3.113345 | 8.25E-02 | 9.69E-02 | Healthy control | Liver |
| Hivep2 | 0.40213 | 6.070698 | 27.22142 | 2.13E-06 | 3.58E-06 | Infection | Liver |
| Hjv | -0.79945 | 7.939891 | 58.03298 | 1.65E-10 | 3.44E-10 | Healthy control | Liver |
| Itga4 | 3.161488 | 5.714625 | 759.1389 | 9.48E-37 | 2.58E-35 | Infection | Liver |
| Synpo | 3.159295 | 5.15327 | 316.7575 | 2.29E-26 | 1.52E-25 | Infection | Liver |
| Cygb | 3.157923 | 7.765714 | 244.1668 | 1.48E-22 | 6.70E-22 | Infection | Liver |
| Hlf | -1.59799 | 7.3609 | 346.7116 | 2.19E-27 | 1.63E-26 | Healthy control | Liver |
| Hmcn1 | 0.299987 | 4.583048 | 6.528728 | 1.30E-02 | 1.69E-02 | Infection | Liver |
| Abcb1b | 3.156936 | 3.05508 | 161.8874 | 7.16E-23 | 3.33E-22 | Infection | Liver |
| Hmga1 | -0.94948 | 4.150241 | 60.25582 | 8.35E-11 | 1.76E-10 | Healthy control | Liver |
| Hmga1b | -0.56244 | 3.622847 | 19.67302 | 3.63E-05 | 5.72E-05 | Healthy control | Liver |
| Hmga2 | 1.638334 | 3.752051 | 93.11991 | 4.38E-14 | 1.10E-13 | Infection | Liver |
| Hmgcr | 0.169366 | 8.692695 | 2.758122 | 1.02E-01 | 1.18E-01 | Infection | Liver |
| Hmox1 | 0.72725 | 7.446852 | 40.56386 | 2.51E-08 | 4.66E-08 | Infection | Liver |
| Hmx2 | 1.73318 | 1.402199 | 7.213361 | 9.04E-03 | 1.19E-02 | Infection | Liver |
| Hmx3 | 0.962803 | 1.300323 | 2.22721 | 1.42E-01 | 1.61E-01 | Infection | Liver |
| Hnf1a | -2.59707 | 7.571557 | 1267.71 | 1.38E-43 | 1.44E-41 | Healthy control | Liver |
| Hnf1b | -0.2758 | 5.425258 | 5.140935 | 2.68E-02 | 3.35E-02 | Healthy control | Liver |
| Hnf4a | -1.27647 | 10.86207 | 214.9591 | 4.87E-22 | 2.12E-21 | Healthy control | Liver |
| Hnrnpa1 | -0.31131 | 8.714469 | 16.39849 | 1.42E-04 | 2.16E-04 | Healthy control | Liver |
| Hnrnpdl | 0.403525 | 7.384063 | 65.94008 | 2.11E-11 | 4.59E-11 | Infection | Liver |
| Hnrnph1 | 0.506382 | 8.807152 | 111.8867 | 1.24E-15 | 3.40E-15 | Infection | Liver |
| Hnrnpk | 0.700985 | 8.825389 | 126.104 | 1.03E-16 | 3.01E-16 | Infection | Liver |
| Hnrnpm | 0.06396 | 9.35261 | 4.357013 | 4.09E-02 | 5.01E-02 | Infection | Liver |
| Homer1 | -0.31733 | 5.983111 | 15.482 | 2.10E-04 | 3.14E-04 | Healthy control | Liver |
| Homer3 | 1.642046 | 5.027545 | 129.7921 | 6.15E-17 | 1.83E-16 | Infection | Liver |
| Hopx | -0.95623 | 6.077869 | 50.33104 | 1.34E-09 | 2.66E-09 | Healthy control | Liver |
| Hoxa1 | 1.961687 | 3.294816 | 127.0745 | 1.50E-17 | 4.64E-17 | Infection | Liver |
| Hoxa10 | 1.580471 | 1.440667 | 6.735363 | 1.12E-02 | 1.46E-02 | Infection | Liver |
| Hoxa11 | 0.935919 | 1.300893 | 2.328555 | 1.34E-01 | 1.53E-01 | Infection | Liver |
| Hoxa13 | 1.613461 | 1.279196 | 6.022371 | 1.90E-02 | 2.43E-02 | Infection | Liver |
| Hoxa2 | 1.768886 | 3.145595 | 81.86481 | 1.51E-13 | 3.71E-13 | Infection | Liver |
| Hoxa3 | 1.319366 | 3.620421 | 59.68447 | 1.01E-10 | 2.13E-10 | Infection | Liver |
| Hoxa4 | 0.931377 | 2.502238 | 14.46963 | 2.65E-04 | 3.94E-04 | Infection | Liver |
| Hoxa5 | 1.196023 | 2.314962 | 21.18652 | 1.21E-05 | 1.96E-05 | Infection | Liver |
| Sema3b | 3.149752 | 3.175044 | 103.566 | 5.77E-17 | 1.72E-16 | Infection | Liver |
| Nphs2 | 3.145978 | 1.846846 | 51.23703 | 3.91E-10 | 8.00E-10 | Infection | Liver |
| Hoxb1 | 0.667124 | 1.265561 | 1.405468 | 2.43E-01 | 2.68E-01 | Infection | Liver |
| Hoxb13 | 1.181493 | 1.334189 | 3.641447 | 6.12E-02 | 7.31E-02 | Infection | Liver |
| Hoxb2 | 1.528297 | 2.789688 | 58.37582 | 4.11E-11 | 8.82E-11 | Infection | Liver |
| Ryr2 | 3.144005 | 1.572295 | 46.49227 | 3.67E-09 | 7.14E-09 | Infection | Liver |

| | | | | | | | |
|---|---|---|---|---|---|---|---|
| Hoxb4 | 0.972223 | 5.438774 | 62.78112 | 4.75E-11 | 1.02E-10 | Infection | Liver |
| Hoxb5 | -0.24153 | 4.349172 | 3.528707 | 6.49E-02 | 7.72E-02 | Healthy control | Liver |
| Hoxb6 | 1.530392 | 2.321137 | 39.24261 | 8.12E-09 | 1.55E-08 | Infection | Liver |
| Hoxb7 | 1.472384 | 1.953813 | 22.66888 | 5.25E-06 | 8.66E-06 | Infection | Liver |
| Plaat1 | 3.143787 | 1.448637 | 39.34803 | 3.54E-08 | 6.52E-08 | Infection | Liver |
| Gdpd1 | 3.143412 | 4.405035 | 649.356 | 3.41E-36 | 8.20E-35 | Infection | Liver |
| Hoxc10 | 1.427348 | 1.306526 | 3.874495 | 5.49E-02 | 6.60E-02 | Infection | Liver |
| Hoxc11 | 0.680056 | 1.216802 | 1.413236 | 2.45E-01 | 2.71E-01 | Infection | Liver |
| Hoxc12 | 0.765494 | 1.220508 | 1.727501 | 2.00E-01 | 2.23E-01 | Infection | Liver |
| Cd19 | 3.142318 | 3.053978 | 129.4923 | 5.79E-20 | 2.16E-19 | Infection | Liver |
| Crybb3 | 3.14093 | 5.977332 | 370.2553 | 5.51E-28 | 4.39E-27 | Infection | Liver |
| Hoxc8 | 1.595626 | 1.292603 | 5.652585 | 2.15E-02 | 2.73E-02 | Infection | Liver |
| L1cam | 3.138501 | 2.309788 | 89.01541 | 1.52E-15 | 4.12E-15 | Infection | Liver |
| Hoxd10 | 1.875641 | 1.323484 | 6.574436 | 1.35E-02 | 1.75E-02 | Infection | Liver |
| Hoxd11 | 1.209714 | 1.275688 | 3.611821 | 6.48E-02 | 7.71E-02 | Infection | Liver |
| Hoxd12 | 1.787633 | 1.357258 | 7.312799 | 8.80E-03 | 1.16E-02 | Infection | Liver |
| Hoxd13 | 1.9516 | 1.363205 | 5.812155 | 1.93E-02 | 2.46E-02 | Infection | Liver |
| Hoxd3 | 1.690884 | 1.33274 | 8.02788 | 6.52E-03 | 8.73E-03 | Infection | Liver |
| Hspb2 | 3.138481 | 3.946085 | 229.4701 | 2.46E-24 | 1.33E-23 | Infection | Liver |
| Rasgrf1 | 3.13642 | 1.804407 | 47.60087 | 1.69E-09 | 3.34E-09 | Infection | Liver |
| Hpcal4 | 0.319423 | 1.255144 | 0.266916 | 6.08E-01 | 6.33E-01 | Infection | Liver |
| Hpd | -2.48779 | 13.32943 | 621.5625 | 1.82E-34 | 3.09E-33 | Healthy control | Liver |
| Hpgds | 1.942607 | 4.04461 | 168.322 | 1.82E-19 | 6.51E-19 | Infection | Liver |
| Hprt | 0.025927 | 7.609015 | 0.239634 | 6.26E-01 | 6.50E-01 | Infection | Liver |
| Hps4 | -0.85667 | 6.297208 | 170.8701 | 1.21E-19 | 4.42E-19 | Healthy control | Liver |
| Hps5 | -0.05587 | 6.688882 | 1.010802 | 3.19E-01 | 3.45E-01 | Healthy control | Liver |
| Hps6 | 0.525646 | 5.256806 | 38.26963 | 5.06E-08 | 9.24E-08 | Infection | Liver |
| Hpse | 1.918741 | 5.195523 | 364.3076 | 7.19E-28 | 5.66E-27 | Infection | Liver |
| Aurkb | 3.124171 | 4.2317 | 252.9621 | 1.05E-24 | 5.89E-24 | Infection | Liver |
| Gzma | 3.123319 | 6.128631 | 118.1609 | 1.34E-15 | 3.65E-15 | Infection | Liver |
| Hras | -1.23386 | 3.833161 | 84.34176 | 1.90E-13 | 4.62E-13 | Healthy control | Liver |
| Hrg | -0.18479 | 11.44192 | 2.168042 | 1.46E-01 | 1.66E-01 | Healthy control | Liver |
| Hrh1 | 1.794185 | 2.026978 | 36.28521 | 1.73E-08 | 3.24E-08 | Infection | Liver |
| Hrh3 | 0.945009 | 1.195056 | 4.363015 | 5.10E-02 | 6.16E-02 | Infection | Liver |
| Hs3st2 | 0.588916 | 1.320233 | 1.138585 | 2.90E-01 | 3.16E-01 | Infection | Liver |
| Hsd17b3 | 0.115774 | 2.306567 | 0.199217 | 6.56E-01 | 6.80E-01 | Infection | Liver |
| Hsd17b4 | -2.45146 | 10.70057 | 1204.055 | 6.01E-43 | 5.28E-41 | Healthy control | Liver |
| Hsd17b7 | -0.6996 | 7.524842 | 49.14876 | 1.86E-09 | 3.67E-09 | Healthy control | Liver |
| Hsf1 | -0.24088 | 7.099523 | 24.09832 | 6.73E-06 | 1.11E-05 | Healthy control | Liver |
| Hsf2 | -1.53687 | 7.278286 | 834.0744 | 3.52E-38 | 1.14E-36 | Healthy control | Liver |
| Hspa1b | 0.836007 | 4.988494 | 32.16349 | 3.80E-07 | 6.66E-07 | Infection | Liver |
| Hspa5 | 0.783609 | 11.71248 | 65.73228 | 2.22E-11 | 4.82E-11 | Infection | Liver |
| Hspa9 | -1.25379 | 10.36079 | 282.9275 | 4.66E-25 | 2.68E-24 | Healthy control | Liver |
| Pycard | 3.115156 | 3.847257 | 283.3945 | 3.15E-27 | 2.26E-26 | Infection | Liver |
| Hspb3 | 1.432686 | 1.233057 | 5.447213 | 2.62E-02 | 3.28E-02 | Infection | Liver |
| Hspd1 | -1.40629 | 10.89693 | 461.5092 | 8.49E-31 | 9.08E-30 | Healthy control | Liver |
| Hspg2 | 1.098203 | 8.628048 | 67.21347 | 2.17E-11 | 4.71E-11 | Infection | Liver |
| Hsph1 | -0.87713 | 7.302427 | 109.4087 | 1.96E-15 | 5.30E-15 | Healthy control | Liver |
| Htr1a | 0.934113 | 1.301247 | 2.377856 | 1.29E-01 | 1.48E-01 | Infection | Liver |
| Saa2 | 3.115025 | 8.515066 | 92.62298 | 1.23E-13 | 3.02E-13 | Infection | Liver |

| | | | | | | | |
|---|---|---|---|---|---|---|---|
| Htr1d | 0.50448 | 1.326447 | 0.711233 | 4.02E-01 | 4.30E-01 | Infection | Liver |
| Htr1f | 1.324759 | 1.26865 | 4.639955 | 3.70E-02 | 4.55E-02 | Infection | Liver |
| Ndc80 | 3.114796 | 2.745049 | 175.5973 | 1.27E-24 | 7.04E-24 | Infection | Liver |
| Gdf7 | 3.113275 | 1.553976 | 37.27373 | 5.79E-08 | 1.06E-07 | Infection | Liver |
| Htr2c | 1.176431 | 1.338514 | 4.772224 | 3.28E-02 | 4.06E-02 | Infection | Liver |
| Htr3a | 0.038341 | 1.25485 | 0.004365 | 9.48E-01 | 9.53E-01 | Infection | Liver |
| Htr3b | 0.855033 | 1.232804 | 1.947856 | 1.73E-01 | 1.94E-01 | Infection | Liver |
| Htr4 | 1.678132 | 1.300264 | 7.91489 | 7.03E-03 | 9.37E-03 | Infection | Liver |
| Adamts20 | 3.111746 | 1.496258 | 40.67425 | 4.25E-08 | 7.80E-08 | Infection | Liver |
| Htra2 | -0.91928 | 5.240612 | 78.05473 | 1.25E-12 | 2.89E-12 | Healthy control | Liver |
| Htt | -0.24373 | 6.661835 | 14.21369 | 3.61E-04 | 5.33E-04 | Healthy control | Liver |
| Huwe1 | 0.112447 | 7.924003 | 5.188408 | 2.61E-02 | 3.27E-02 | Infection | Liver |
| Ibsp | 1.309135 | 1.312233 | 4.445512 | 3.97E-02 | 4.87E-02 | Infection | Liver |
| Icam1 | 1.89376 | 7.511423 | 513.615 | 4.25E-32 | 5.26E-31 | Infection | Liver |
| Ick | -1.1863 | 5.477558 | 207.5413 | 1.29E-21 | 5.40E-21 | Healthy control | Liver |
| Icos | 0.294208 | 3.69834 | 4.232974 | 4.38E-02 | 5.33E-02 | Infection | Liver |
| Trpv4 | 3.106734 | 3.516148 | 206.1998 | 2.27E-24 | 1.23E-23 | Infection | Liver |
| Id1 | -0.11885 | 6.043345 | 2.2357 | 1.40E-01 | 1.59E-01 | Healthy control | Liver |
| Id2 | -2.13983 | 9.443535 | 1431.577 | 3.21E-45 | 6.69E-43 | Healthy control | Liver |
| Id4 | -0.49527 | 4.693669 | 11.00586 | 1.51E-03 | 2.13E-03 | Healthy control | Liver |
| Idh1 | -2.47514 | 10.2885 | 1639.805 | 5.12E-47 | 1.60E-44 | Healthy control | Liver |
| Idh2 | -1.28442 | 9.297399 | 556.6046 | 4.30E-33 | 6.00E-32 | Healthy control | Liver |
| Idi1 | 0.281984 | 9.972511 | 6.884958 | 1.09E-02 | 1.43E-02 | Infection | Liver |
| Ido1 | 1.483677 | 1.376099 | 7.048395 | 9.91E-03 | 1.30E-02 | Infection | Liver |
| Ido2 | -2.68961 | 7.17357 | 1098.623 | 1.11E-41 | 8.19E-40 | Healthy control | Liver |
| Idua | 0.834537 | 4.891447 | 66.10853 | 2.11E-11 | 4.59E-11 | Infection | Liver |
| Ifi204 | 0.432982 | 5.579009 | 14.84189 | 2.76E-04 | 4.09E-04 | Infection | Liver |
| Ifi208 | 0.592823 | 2.187438 | 4.693104 | 3.25E-02 | 4.03E-02 | Infection | Liver |
| Selplg | 3.102071 | 6.566179 | 704.6771 | 5.65E-36 | 1.26E-34 | Infection | Liver |
| Ifi27 | 0.262457 | 8.348123 | 12.39537 | 8.03E-04 | 1.16E-03 | Infection | Liver |
| Ifi44 | 0.684785 | 6.456953 | 28.66313 | 1.29E-06 | 2.19E-06 | Infection | Liver |
| Ifih1 | -0.45813 | 7.203771 | 44.50467 | 7.28E-09 | 1.39E-08 | Healthy control | Liver |
| Ifit1 | -0.43518 | 4.770676 | 3.616576 | 6.19E-02 | 7.39E-02 | Healthy control | Liver |
| Ifit3 | 1.289013 | 6.037648 | 26.69813 | 3.01E-06 | 5.00E-06 | Infection | Liver |
| Ifnar1 | -1.02183 | 6.47982 | 301.645 | 9.04E-26 | 5.54E-25 | Healthy control | Liver |
| Ifnar2 | 0.622342 | 6.709894 | 82.4361 | 4.49E-13 | 1.07E-12 | Infection | Liver |
| Ifnb1 | 1.053395 | 1.245038 | 2.824049 | 1.02E-01 | 1.19E-01 | Infection | Liver |
| Rad51 | 3.101173 | 3.790191 | 287.4204 | 1.09E-27 | 8.41E-27 | Infection | Liver |
| Ifngr1 | 0.912191 | 7.771366 | 311.7448 | 3.68E-26 | 2.38E-25 | Infection | Liver |
| Ifngr2 | -0.23971 | 5.822003 | 6.984923 | 1.03E-02 | 1.36E-02 | Healthy control | Liver |
| Lum | 3.100109 | 9.832932 | 160.4965 | 2.27E-18 | 7.41E-18 | Infection | Liver |
| Ift122 | 0.789362 | 4.871814 | 57.88062 | 1.73E-10 | 3.60E-10 | Infection | Liver |
| Ift27 | -0.04956 | 4.66853 | 0.230371 | 6.33E-01 | 6.57E-01 | Healthy control | Liver |
| Ift88 | 0.759815 | 6.182584 | 133.6388 | 2.99E-17 | 9.10E-17 | Infection | Liver |
| Moxd1 | 3.098627 | 1.528023 | 26.30838 | 2.58E-06 | 4.31E-06 | Infection | Liver |
| Igf1 | -0.56956 | 11.03238 | 38.1878 | 5.14E-08 | 9.38E-08 | Healthy control | Liver |
| Igf1r | 1.745049 | 5.72675 | 268.1311 | 2.00E-24 | 1.09E-23 | Infection | Liver |
| Edn1 | 3.095027 | 3.13252 | 113.3834 | 5.20E-18 | 1.66E-17 | Infection | Liver |
| Igf2bp1 | 0.042053 | 1.270252 | 0.003713 | 9.52E-01 | 9.56E-01 | Infection | Liver |
| Igf2r | -0.06666 | 7.699929 | 1.07925 | 3.03E-01 | 3.29E-01 | Healthy control | Liver |

| | | | | | | | |
|---|---|---|---|---|---|---|---|
| Dusp5 | 3.090771 | 5.280079 | 808.5353 | 1.26E-37 | 3.83E-36 | Infection | Liver |
| Igfbp2 | -0.92368 | 9.116463 | 44.50032 | 9.05E-09 | 1.72E-08 | Healthy control | Liver |
| Igfbp3 | 0.527011 | 8.022281 | 7.867542 | 6.83E-03 | 9.12E-03 | Infection | Liver |
| Igfn1 | 0.202516 | 1.274922 | 0.130753 | 7.19E-01 | 7.39E-01 | Infection | Liver |
| Slco4c1 | 3.087449 | 1.860816 | 45.54917 | 2.70E-09 | 5.29E-09 | Infection | Liver |
| Ogn | 3.08615 | 3.811527 | 148.7321 | 1.45E-19 | 5.22E-19 | Infection | Liver |
| Tnfsf9 | 3.083023 | 1.635023 | 40.3302 | 1.94E-08 | 3.63E-08 | Infection | Liver |
| Rgs18 | 3.077281 | 3.404299 | 162.1035 | 1.02E-21 | 4.29E-21 | Infection | Liver |
| Igll1 | 0.408512 | 1.230402 | 0.537043 | 4.69E-01 | 4.97E-01 | Infection | Liver |
| Igsf11 | -1.62234 | 8.196518 | 339.4291 | 3.84E-27 | 2.71E-26 | Healthy control | Liver |
| Igsf21 | -0.51703 | 1.484914 | 1.346794 | 2.48E-01 | 2.74E-01 | Healthy control | Liver |
| Igsf5 | -0.72166 | 7.319307 | 44.04812 | 8.45E-09 | 1.61E-08 | Healthy control | Liver |
| Igsf8 | 1.638276 | 4.57437 | 213.294 | 7.69E-22 | 3.29E-21 | Infection | Liver |
| Aqp3 | 3.076468 | 2.103486 | 67.87379 | 1.20E-12 | 2.79E-12 | Infection | Liver |
| Igtp | 0.965771 | 7.17432 | 47.60186 | 3.27E-09 | 6.38E-09 | Infection | Liver |
| Ihh | 1.093627 | 1.976105 | 14.88191 | 1.84E-04 | 2.77E-04 | Infection | Liver |
| Ikbkb | 0.099679 | 8.441875 | 4.760329 | 3.28E-02 | 4.07E-02 | Infection | Liver |
| Ikbkg | 0.193117 | 8.488258 | 5.389231 | 2.35E-02 | 2.97E-02 | Infection | Liver |
| Cav1 | 3.072012 | 4.827066 | 199.7974 | 2.67E-21 | 1.09E-20 | Infection | Liver |
| Dpysl4 | 3.066063 | 2.051208 | 64.86891 | 3.40E-12 | 7.68E-12 | Infection | Liver |
| Mmp16 | 3.059988 | 1.848538 | 56.07718 | 8.19E-11 | 1.73E-10 | Infection | Liver |
| Il10ra | 1.677501 | 6.758768 | 454.4441 | 1.33E-30 | 1.40E-29 | Infection | Liver |
| Cd96 | 3.058512 | 2.899987 | 113.6722 | 1.58E-18 | 5.24E-18 | Infection | Liver |
| Il11ra1 | -0.24024 | 7.969676 | 33.22099 | 2.62E-07 | 4.64E-07 | Healthy control | Liver |
| Adm | 3.054023 | 5.383877 | 350.9197 | 1.83E-27 | 1.38E-26 | Infection | Liver |
| Runx3 | 3.04844 | 4.005736 | 278.4731 | 2.57E-26 | 1.69E-25 | Infection | Liver |
| Il12rb1 | 1.520907 | 2.379262 | 37.4457 | 1.71E-08 | 3.21E-08 | Infection | Liver |
| Pclaf | 3.046249 | 2.481386 | 102.9615 | 2.09E-17 | 6.39E-17 | Infection | Liver |
| Amy2a5 | 3.044547 | 5.866108 | 118.96 | 1.09E-15 | 2.99E-15 | Infection | Liver |
| Il13ra1 | 0.467628 | 8.690267 | 64.83547 | 2.78E-11 | 6.01E-11 | Infection | Liver |
| Col7a1 | 3.033018 | 4.412922 | 336.984 | 9.21E-28 | 7.18E-27 | Infection | Liver |
| Il15 | 0.643703 | 4.684158 | 41.90071 | 1.65E-08 | 3.10E-08 | Infection | Liver |
| Il15ra | -1.54676 | 6.887679 | 385.2717 | 1.29E-28 | 1.09E-27 | Healthy control | Liver |
| Il17a | 1.49894 | 1.517126 | 9.087157 | 3.28E-03 | 4.49E-03 | Infection | Liver |
| Il17f | 1.073861 | 1.280415 | 2.991194 | 9.05E-02 | 1.06E-01 | Infection | Liver |
| Il17ra | 0.980529 | 6.350025 | 237.4139 | 4.19E-23 | 1.98E-22 | Infection | Liver |
| Il17rb | 0.183816 | 5.312796 | 2.862414 | 9.56E-02 | 1.12E-01 | Infection | Liver |
| Il17rc | -0.5281 | 7.997918 | 77.75537 | 1.27E-12 | 2.95E-12 | Healthy control | Liver |
| Il18 | -0.86293 | 5.97182 | 52.425 | 7.45E-10 | 1.50E-09 | Healthy control | Liver |
| Parpbp | 3.03254 | 2.511821 | 108.6265 | 4.52E-18 | 1.44E-17 | Infection | Liver |
| Hr | 3.028663 | 2.920434 | 135.8609 | 2.08E-20 | 7.99E-20 | Infection | Liver |
| Il1a | 0.388972 | 5.774832 | 7.619885 | 7.54E-03 | 1.00E-02 | Infection | Liver |
| Il1f10 | 1.136131 | 1.221252 | 4.833896 | 3.62E-02 | 4.46E-02 | Infection | Liver |
| Il1f5 | 1.717485 | 1.351055 | 7.644599 | 7.55E-03 | 1.00E-02 | Infection | Liver |
| Il1r1 | 1.853536 | 7.983915 | 430.4898 | 5.91E-30 | 5.77E-29 | Infection | Liver |
| Il1rap | -1.84145 | 8.621115 | 423.2952 | 9.41E-30 | 8.92E-29 | Healthy control | Liver |
| Hmcn2 | 3.028368 | 3.780898 | 240.2855 | 1.99E-25 | 1.18E-24 | Infection | Liver |
| Ccr6 | 3.021436 | 2.467819 | 81.54342 | 7.41E-15 | 1.94E-14 | Infection | Liver |
| Fgf2 | 3.018785 | 3.797213 | 193.6812 | 1.62E-22 | 7.34E-22 | Infection | Liver |
| Pfkp | 3.017051 | 7.273811 | 769.9461 | 3.79E-37 | 1.08E-35 | Infection | Liver |

| | | | | | | | |
|---|---|---|---|---|---|---|---|
| Smarca1 | 3.014324 | 3.455554 | 207.8065 | 1.29E-24 | 7.19E-24 | Infection | Liver |
| Selp | 3.013744 | 4.364532 | 204.482 | 4.09E-22 | 1.79E-21 | Infection | Liver |
| Irf4 | 3.012564 | 4.918902 | 130.5975 | 7.62E-17 | 2.25E-16 | Infection | Liver |
| Il22ra1 | -1.34888 | 5.139541 | 170.9352 | 1.31E-19 | 4.75E-19 | Healthy control | Liver |
| Il22ra2 | 0.970577 | 1.382078 | 3.580672 | 6.24E-02 | 7.44E-02 | Infection | Liver |
| Il23a | 0.797224 | 1.401569 | 2.266236 | 1.36E-01 | 1.55E-01 | Infection | Liver |
| Il23r | 1.390647 | 1.357014 | 4.707304 | 3.37E-02 | 4.16E-02 | Infection | Liver |
| Il25 | 1.237402 | 1.222807 | 4.406189 | 4.53E-02 | 5.52E-02 | Infection | Liver |
| Il27 | 1.186138 | 2.098845 | 18.61324 | 3.29E-05 | 5.20E-05 | Infection | Liver |
| Fap | 3.012479 | 4.272389 | 229.3786 | 1.60E-23 | 7.89E-23 | Infection | Liver |
| Prl8a1 | 3.008739 | 1.588445 | 25.75266 | 3.04E-06 | 5.05E-06 | Infection | Liver |
| Il2rb | 1.74854 | 5.24618 | 91.73507 | 7.14E-14 | 1.77E-13 | Infection | Liver |
| Mmp3 | 3.007255 | 1.529945 | 34.98376 | 1.32E-07 | 2.38E-07 | Infection | Liver |
| Il3 | 0.99698 | 1.266183 | 2.284297 | 1.39E-01 | 1.58E-01 | Infection | Liver |
| Il31 | 1.828956 | 1.380296 | 6.668598 | 1.21E-02 | 1.58E-02 | Infection | Liver |
| Ccr5 | 3.005902 | 7.509663 | 1022.441 | 8.46E-41 | 5.04E-39 | Infection | Liver |
| Il34 | 0.754968 | 5.672896 | 30.81367 | 6.00E-07 | 1.04E-06 | Infection | Liver |
| Il3ra | 0.609487 | 1.519059 | 2.068236 | 1.53E-01 | 1.73E-01 | Infection | Liver |
| 6-Sep | 3.004501 | 5.868819 | 715.46 | 4.64E-36 | 1.05E-34 | Infection | Liver |
| Il4ra | 1.505213 | 7.100285 | 198.3412 | 3.48E-21 | 1.41E-20 | Infection | Liver |
| Tagln | 3.003152 | 6.774619 | 237.4881 | 2.11E-22 | 9.47E-22 | Infection | Liver |
| Grin2a | 2.996732 | 1.767868 | 38.78487 | 2.47E-08 | 4.58E-08 | Infection | Liver |
| Il6ra | -0.09587 | 7.191385 | 1.623591 | 2.07E-01 | 2.31E-01 | Healthy control | Liver |
| Il6st | -0.95268 | 8.687363 | 167.0163 | 2.01E-19 | 7.15E-19 | Healthy control | Liver |
| Il7 | 0.923624 | 3.233928 | 22.37155 | 1.22E-05 | 1.98E-05 | Infection | Liver |
| Prl7a1 | 2.996283 | 1.596043 | 17.33453 | 8.65E-05 | 1.33E-04 | Infection | Liver |
| Il9 | -0.25497 | 1.222202 | 0.177093 | 6.77E-01 | 7.00E-01 | Healthy control | Liver |
| Ilf2 | -0.33213 | 6.225408 | 17.057 | 1.08E-04 | 1.65E-04 | Healthy control | Liver |
| Ilk | 0.998923 | 6.921435 | 297.2384 | 1.30E-25 | 7.84E-25 | Infection | Liver |
| Ilkap | 0.217147 | 7.75216 | 23.53886 | 8.32E-06 | 1.36E-05 | Infection | Liver |
| Immt | -0.81356 | 9.325883 | 247.676 | 1.42E-23 | 7.02E-23 | Healthy control | Liver |
| Impact | 0.842317 | 7.63028 | 197.7698 | 3.72E-21 | 1.50E-20 | Infection | Liver |
| Impad1 | 0.319332 | 7.040386 | 33.20498 | 2.64E-07 | 4.66E-07 | Infection | Liver |
| Ina | 0.410472 | 1.205477 | 0.603632 | 4.45E-01 | 4.73E-01 | Infection | Liver |
| Ing3 | -0.14752 | 6.094647 | 6.063665 | 1.65E-02 | 2.13E-02 | Healthy control | Liver |
| Osr1 | 2.994485 | 1.667429 | 31.55834 | 3.44E-07 | 6.04E-07 | Infection | Liver |
| Inhba | 0.835963 | 6.120253 | 29.53259 | 9.81E-07 | 1.68E-06 | Infection | Liver |
| C1qtnf5 | 2.993988 | 6.765298 | 271.0901 | 6.51E-24 | 3.37E-23 | Infection | Liver |
| Adamts5 | 2.989918 | 6.411085 | 154.1886 | 4.96E-18 | 1.58E-17 | Infection | Liver |
| Ino80d | 0.487502 | 7.68273 | 145.3504 | 4.69E-18 | 1.50E-17 | Infection | Liver |
| Inpp1 | -0.80089 | 5.973637 | 113.4812 | 9.51E-16 | 2.62E-15 | Healthy control | Liver |
| Mki67 | 2.988126 | 6.20099 | 269.4751 | 3.85E-24 | 2.05E-23 | Infection | Liver |
| Inpp5b | -0.40005 | 7.636831 | 90.38378 | 8.18E-14 | 2.02E-13 | Healthy control | Liver |
| Psat1 | 2.987819 | 2.762015 | 142.5669 | 1.13E-21 | 4.75E-21 | Infection | Liver |
| Inpp5e | 0.389934 | 5.516658 | 28.97863 | 1.14E-06 | 1.95E-06 | Infection | Liver |
| Ezr | 2.984775 | 6.378037 | 1011.754 | 1.40E-40 | 7.88E-39 | Infection | Liver |
| Insm1 | -0.12653 | 1.208399 | 0.045241 | 8.34E-01 | 8.47E-01 | Healthy control | Liver |
| Insr | -1.5947 | 8.823966 | 765.0027 | 4.36E-37 | 1.23E-35 | Healthy control | Liver |
| Crisp1 | 2.984198 | 1.879309 | 23.28868 | 6.25E-06 | 1.03E-05 | Infection | Liver |
| Ints11 | -0.76081 | 6.68842 | 219.7837 | 2.86E-22 | 1.27E-21 | Healthy control | Liver |

| | | | | | | | |
|---|---|---|---|---|---|---|---|
| Ints4 | -0.81548 | 6.501808 | 173.2926 | 8.66E-20 | 3.20E-19 | Healthy control | Liver |
| Invs | -0.94509 | 5.659792 | 150.8029 | 2.19E-18 | 7.18E-18 | Healthy control | Liver |
| Ipmk | -0.07083 | 7.455595 | 0.715012 | 4.01E-01 | 4.29E-01 | Healthy control | Liver |
| Ipo9 | -0.01004 | 6.396282 | 0.021766 | 8.83E-01 | 8.93E-01 | Healthy control | Liver |
| Iqgap1 | 1.641075 | 7.034686 | 243.4416 | 2.21E-23 | 1.08E-22 | Infection | Liver |
| Iqgap2 | -1.33846 | 9.509632 | 194.2759 | 5.70E-21 | 2.26E-20 | Healthy control | Liver |
| Rad21l | 2.983989 | 1.658242 | 29.4807 | 6.74E-07 | 1.16E-06 | Infection | Liver |
| Irak1 | -0.2493 | 8.026573 | 40.02402 | 2.87E-08 | 5.32E-08 | Healthy control | Liver |
| Irak2 | -1.40596 | 6.222433 | 351.2571 | 1.65E-27 | 1.25E-26 | Healthy control | Liver |
| Pld5 | 2.974379 | 1.587748 | 36.95626 | 5.33E-08 | 9.72E-08 | Infection | Liver |
| Ireb2 | -0.84057 | 8.114831 | 249.7347 | 1.15E-23 | 5.75E-23 | Healthy control | Liver |
| Irf1 | 0.542317 | 8.48885 | 67.80912 | 1.33E-11 | 2.93E-11 | Infection | Liver |
| Irf3 | -0.67972 | 8.99656 | 323.7789 | 1.35E-26 | 9.05E-26 | Healthy control | Liver |
| Pmaip1 | 2.973618 | 3.977046 | 200.9579 | 2.33E-22 | 1.04E-21 | Infection | Liver |
| Irf5 | 1.709104 | 6.726086 | 398.9918 | 4.87E-29 | 4.29E-28 | Infection | Liver |
| Irf7 | 0.816781 | 3.827067 | 16.37581 | 1.45E-04 | 2.20E-04 | Infection | Liver |
| Ccl2 | 2.971513 | 5.56091 | 246.3245 | 2.70E-23 | 1.30E-22 | Infection | Liver |
| Irf9 | -0.73378 | 8.295785 | 163.4552 | 3.30E-19 | 1.15E-18 | Healthy control | Liver |
| Irs1 | -2.11994 | 6.921549 | 738.9375 | 1.33E-36 | 3.42E-35 | Healthy control | Liver |
| Irs2 | -1.01156 | 7.324267 | 120.7665 | 2.57E-16 | 7.29E-16 | Healthy control | Liver |
| Irx2 | 1.980001 | 1.339308 | 8.39303 | 5.33E-03 | 7.17E-03 | Infection | Liver |
| Irx3 | 0.785936 | 2.441147 | 9.184478 | 3.20E-03 | 4.39E-03 | Infection | Liver |
| Irx4 | 0.963332 | 1.233225 | 2.393146 | 1.33E-01 | 1.52E-01 | Infection | Liver |
| Rflna | 2.970755 | 2.314826 | 102.0343 | 3.16E-17 | 9.58E-17 | Infection | Liver |
| Irx6 | 1.743795 | 1.254426 | 9.811911 | 3.45E-03 | 4.72E-03 | Infection | Liver |
| Isca1 | -0.67564 | 7.62996 | 161.4182 | 4.41E-19 | 1.53E-18 | Healthy control | Liver |
| Isg15 | -0.29584 | 8.273536 | 3.308779 | 7.40E-02 | 8.74E-02 | Healthy control | Liver |
| Isg20 | 0.632275 | 4.876917 | 19.51237 | 4.01E-05 | 6.29E-05 | Infection | Liver |
| Isl1 | 1.791073 | 1.369939 | 7.290113 | 8.93E-03 | 1.18E-02 | Infection | Liver |
| Isl2 | 0.67569 | 1.337232 | 1.260942 | 2.66E-01 | 2.91E-01 | Infection | Liver |
| Itch | -0.37716 | 8.292303 | 90.95075 | 7.26E-14 | 1.80E-13 | Healthy control | Liver |
| Itga1 | 1.512411 | 8.968173 | 842.49 | 2.55E-38 | 8.40E-37 | Infection | Liver |
| Cacna1s | 2.966212 | 2.045012 | 57.55391 | 2.47E-11 | 5.35E-11 | Infection | Liver |
| Gdf10 | 2.965425 | 8.998426 | 695.8936 | 1.06E-35 | 2.31E-34 | Infection | Liver |
| Itga2b | 1.093729 | 5.102346 | 46.78061 | 3.79E-09 | 7.36E-09 | Infection | Liver |
| Itga3 | 0.288247 | 5.450255 | 8.80459 | 4.24E-03 | 5.74E-03 | Infection | Liver |
| Pdgfrb | 2.963649 | 9.024444 | 276.4443 | 7.17E-24 | 3.68E-23 | Infection | Liver |
| Itga5 | 0.399903 | 8.360627 | 22.61036 | 1.18E-05 | 1.92E-05 | Infection | Liver |
| Efnb3 | 2.959422 | 2.795147 | 164.1165 | 3.81E-23 | 1.81E-22 | Infection | Liver |
| Itga7 | 0.147007 | 5.303059 | 2.045227 | 1.58E-01 | 1.78E-01 | Infection | Liver |
| Itga8 | 1.765585 | 7.408485 | 128.205 | 1.66E-16 | 4.80E-16 | Infection | Liver |
| Itga9 | 1.031729 | 6.959548 | 259.1623 | 4.51E-24 | 2.38E-23 | Infection | Liver |
| Cd22 | 2.957837 | 4.539901 | 393.3219 | 4.41E-29 | 3.89E-28 | Infection | Liver |
| Itgal | 1.039758 | 6.913393 | 88.0639 | 1.33E-13 | 3.27E-13 | Infection | Liver |
| Cd8b1 | 2.956591 | 4.074504 | 185.502 | 2.70E-21 | 1.10E-20 | Infection | Liver |
| Itgav | 0.752409 | 7.443206 | 108.2076 | 2.45E-15 | 6.57E-15 | Infection | Liver |
| Rab38 | 2.952472 | 2.936554 | 135.7042 | 1.87E-20 | 7.20E-20 | Infection | Liver |
| Itgb1 | 1.488914 | 11.10176 | 1060.736 | 2.70E-41 | 1.80E-39 | Infection | Liver |
| Gdap1 | 2.950822 | 1.616053 | 22.18464 | 1.26E-05 | 2.04E-05 | Infection | Liver |
| Itgb3 | 1.341371 | 5.567253 | 102.5411 | 7.35E-15 | 1.92E-14 | Infection | Liver |

| | | | | | | | |
|---|---|---|---|---|---|---|---|
| Pvalb | 2.949784 | 1.424807 | 22.52229 | 1.22E-05 | 1.98E-05 | Infection | Liver |
| Itgb5 | 0.437514 | 8.174611 | 27.30267 | 2.07E-06 | 3.48E-06 | Infection | Liver |
| Fbxl13 | 2.947703 | 2.39687 | 101.4753 | 2.63E-17 | 8.02E-17 | Infection | Liver |
| Cabp1 | 2.946332 | 4.031138 | 253.2617 | 5.20E-25 | 2.98E-24 | Infection | Liver |
| Arg2 | 2.941999 | 3.882587 | 232.4169 | 2.62E-24 | 1.41E-23 | Infection | Liver |
| Blk | 2.940199 | 2.869392 | 119.5292 | 4.94E-19 | 1.70E-18 | Infection | Liver |
| Bdkrb2 | 2.937081 | 1.665931 | 34.27599 | 1.24E-07 | 2.23E-07 | Infection | Liver |
| Col24a1 | 2.93639 | 4.459095 | 248.782 | 6.86E-24 | 3.53E-23 | Infection | Liver |
| Itpkb | 0.69582 | 6.328656 | 55.76969 | 2.95E-10 | 6.08E-10 | Infection | Liver |
| Itpr1 | 0.450581 | 6.93719 | 55.78075 | 2.94E-10 | 6.06E-10 | Infection | Liver |
| Itpr2 | 0.003791 | 6.881312 | 0.003816 | 9.51E-01 | 9.56E-01 | Infection | Liver |
| Itpr3 | 1.681485 | 3.820363 | 115.2722 | 7.11E-16 | 1.97E-15 | Infection | Liver |
| Itsn1 | 0.885992 | 8.847214 | 601.208 | 4.74E-34 | 7.44E-33 | Infection | Liver |
| Itsn2 | 0.967027 | 8.319518 | 444.6846 | 2.40E-30 | 2.46E-29 | Infection | Liver |
| Ivd | -1.89616 | 9.363374 | 449.1418 | 1.81E-30 | 1.88E-29 | Healthy control | Liver |
| Pcdh11x | 2.93502 | 2.639151 | 82.22725 | 5.72E-15 | 1.51E-14 | Infection | Liver |
| Jade1 | -1.18024 | 6.541841 | 317.5622 | 2.33E-26 | 1.54E-25 | Healthy control | Liver |
| Jade2 | 0.666017 | 6.452481 | 94.02474 | 3.88E-14 | 9.79E-14 | Infection | Liver |
| Jade3 | -0.26992 | 4.615169 | 5.989729 | 1.72E-02 | 2.21E-02 | Healthy control | Liver |
| Jag1 | 1.939476 | 6.741996 | 362.6261 | 6.59E-28 | 5.22E-27 | Infection | Liver |
| Tnfrsf13c | 2.932873 | 2.846773 | 100.1889 | 3.87E-17 | 1.17E-16 | Infection | Liver |
| Jak1 | 0.674026 | 9.513759 | 290.6623 | 2.31E-25 | 1.36E-24 | Infection | Liver |
| Jak2 | 1.505646 | 7.786081 | 979.7368 | 2.92E-40 | 1.46E-38 | Infection | Liver |
| Jak3 | 1.325546 | 6.917852 | 392.4261 | 7.61E-29 | 6.57E-28 | Infection | Liver |
| Jam2 | 1.489181 | 5.451621 | 125.6547 | 1.16E-16 | 3.38E-16 | Infection | Liver |
| Jam3 | 1.345231 | 4.53635 | 92.70022 | 5.70E-14 | 1.43E-13 | Infection | Liver |
| Il12a | 2.927815 | 1.722171 | 35.00218 | 9.39E-08 | 1.70E-07 | Infection | Liver |
| Jarid2 | 0.475147 | 6.762782 | 61.47193 | 6.55E-11 | 1.39E-10 | Infection | Liver |
| Jazf1 | 1.503835 | 2.434707 | 31.77836 | 1.64E-07 | 2.94E-07 | Infection | Liver |
| Ltb4r1 | 2.926658 | 4.145557 | 206.142 | 3.17E-22 | 1.40E-21 | Infection | Liver |
| Jmjd6 | -0.23364 | 6.739123 | 20.62508 | 2.56E-05 | 4.07E-05 | Healthy control | Liver |
| Serpinb3b | 2.922432 | 2.084775 | 33.0335 | 1.07E-07 | 1.93E-07 | Infection | Liver |
| Jpt1 | 0.995927 | 7.494485 | 215.4474 | 4.62E-22 | 2.01E-21 | Infection | Liver |
| Jup | -0.74253 | 8.383234 | 64.15214 | 3.30E-11 | 7.11E-11 | Healthy control | Liver |
| Kansl2 | -0.98474 | 7.164913 | 323.1768 | 1.43E-26 | 9.60E-26 | Healthy control | Liver |
| Kat14 | -0.16214 | 7.704648 | 13.68537 | 4.54E-04 | 6.65E-04 | Healthy control | Liver |
| Kat2a | -0.43664 | 7.33278 | 81.12576 | 5.99E-13 | 1.42E-12 | Healthy control | Liver |
| Kat2b | -1.22207 | 7.507087 | 265.245 | 2.49E-24 | 1.34E-23 | Healthy control | Liver |
| Kat5 | -0.67969 | 6.310455 | 128.5033 | 6.97E-17 | 2.07E-16 | Healthy control | Liver |
| Kat6a | -0.08732 | 6.927497 | 2.412943 | 1.25E-01 | 1.44E-01 | Healthy control | Liver |
| Kat6b | 1.012363 | 5.35205 | 117.6876 | 4.55E-16 | 1.27E-15 | Infection | Liver |
| Kat8 | -1.22332 | 7.16989 | 312.1952 | 3.58E-26 | 2.32E-25 | Healthy control | Liver |
| Katna1 | -0.50107 | 6.172979 | 71.43314 | 5.59E-12 | 1.25E-11 | Healthy control | Liver |
| Ebf3 | 2.920212 | 1.582544 | 41.21491 | 1.33E-08 | 2.51E-08 | Infection | Liver |
| Kcna10 | 1.329965 | 1.309963 | 5.115352 | 2.82E-02 | 3.52E-02 | Infection | Liver |
| Cemip | 2.920088 | 1.635303 | 39.60955 | 1.88E-08 | 3.52E-08 | Infection | Liver |
| Ascl2 | 2.919304 | 1.714095 | 38.85879 | 2.12E-08 | 3.96E-08 | Infection | Liver |
| Kcnab3 | 0.517162 | 2.616373 | 5.266893 | 2.44E-02 | 3.08E-02 | Infection | Liver |
| Uhrf1 | 2.916199 | 3.988012 | 240.6319 | 2.21E-24 | 1.20E-23 | Infection | Liver |
| Kcnc2 | 1.361625 | 1.385592 | 6.152375 | 1.55E-02 | 1.99E-02 | Infection | Liver |

| | | | | | | | |
|---|---|---|---|---|---|---|---|
| Kcnc4 | 1.93292 | 1.372149 | 11.756 | 1.05E-03 | 1.50E-03 | Infection | Liver |
| Kcnf1 | 1.680899 | 1.296591 | 7.263689 | 9.78E-03 | 1.29E-02 | Infection | Liver |
| Inka2 | 2.907882 | 1.860475 | 52.57775 | 2.01E-10 | 4.17E-10 | Infection | Liver |
| Kcng2 | 0.48202 | 2.680749 | 4.070815 | 4.71E-02 | 5.72E-02 | Infection | Liver |
| Cd3d | 2.900042 | 4.322711 | 283.6121 | 1.78E-25 | 1.06E-24 | Infection | Liver |
| Nr0b1 | 2.899058 | 1.598512 | 22.59203 | 1.06E-05 | 1.72E-05 | Infection | Liver |
| Kcnh5 | 0.212742 | 1.39713 | 0.147577 | 7.02E-01 | 7.23E-01 | Infection | Liver |
| Kcnh8 | 0.549734 | 1.299177 | 0.858821 | 3.58E-01 | 3.85E-01 | Infection | Liver |
| Kcnj10 | 0.305588 | 4.83937 | 5.319166 | 2.44E-02 | 3.08E-02 | Infection | Liver |
| Kcnj11 | 1.079572 | 1.204857 | 4.826372 | 3.84E-02 | 4.72E-02 | Infection | Liver |
| Elf3 | 2.89304 | 4.622513 | 181.2692 | 2.50E-20 | 9.51E-20 | Infection | Liver |
| Kcnj3 | 1.088649 | 1.328097 | 3.752744 | 5.75E-02 | 6.89E-02 | Infection | Liver |
| Kcnj5 | -0.86792 | 1.776108 | 4.907205 | 2.87E-02 | 3.57E-02 | Healthy control | Liver |
| Kcnj6 | 0.907806 | 1.232351 | 2.336808 | 1.37E-01 | 1.56E-01 | Infection | Liver |
| Kcnj8 | -0.68245 | 5.465581 | 37.92965 | 5.65E-08 | 1.03E-07 | Healthy control | Liver |
| Kcnj9 | 0.899777 | 1.488512 | 3.285992 | 7.27E-02 | 8.61E-02 | Infection | Liver |
| Kcnk10 | -3.20789 | 3.075929 | 152.0182 | 2.84E-21 | 1.16E-20 | Healthy control | Liver |
| Kcnk18 | 1.402938 | 1.221671 | 6.133302 | 2.12E-02 | 2.69E-02 | Infection | Liver |
| Kcnk2 | 1.368378 | 2.420146 | 29.47961 | 4.10E-07 | 7.16E-07 | Infection | Liver |
| Kcnk5 | -2.02892 | 6.849911 | 394.3356 | 7.10E-29 | 6.15E-28 | Healthy control | Liver |
| Cngb3 | 2.89053 | 1.664401 | 18.96542 | 4.41E-05 | 6.90E-05 | Infection | Liver |
| Kcnmb2 | 0.894918 | 1.653676 | 4.851342 | 2.94E-02 | 3.66E-02 | Infection | Liver |
| Kcnn2 | -1.974 | 6.423658 | 241.3178 | 2.98E-23 | 1.43E-22 | Healthy control | Liver |
| Naaa | 2.88961 | 6.23124 | 764.5357 | 5.49E-37 | 1.52E-35 | Infection | Liver |
| Kcnq1 | -0.36125 | 5.170215 | 12.98417 | 6.20E-04 | 8.97E-04 | Healthy control | Liver |
| Gria2 | 2.889534 | 1.558187 | 34.47862 | 1.23E-07 | 2.22E-07 | Infection | Liver |
| Kcns2 | 1.686218 | 1.347566 | 7.352396 | 8.68E-03 | 1.15E-02 | Infection | Liver |
| Kcns3 | 1.495681 | 1.530425 | 9.211815 | 3.05E-03 | 4.19E-03 | Infection | Liver |
| Sh2d1a | 2.889502 | 3.084657 | 171.4689 | 4.59E-23 | 2.16E-22 | Infection | Liver |
| Kctd15 | -0.24311 | 4.417828 | 4.940019 | 2.98E-02 | 3.71E-02 | Healthy control | Liver |
| Kdf1 | -1.02127 | 2.112838 | 10.48951 | 1.62E-03 | 2.28E-03 | Healthy control | Liver |
| Kdm1a | -0.2258 | 5.341894 | 8.500837 | 4.91E-03 | 6.62E-03 | Healthy control | Liver |
| Kdm1b | -1.56018 | 7.10127 | 473.1692 | 4.34E-31 | 4.82E-30 | Healthy control | Liver |
| Kdm2a | -0.29057 | 7.768735 | 45.10379 | 6.08E-09 | 1.17E-08 | Healthy control | Liver |
| Kdm2b | -0.17658 | 5.567299 | 3.324376 | 7.30E-02 | 8.64E-02 | Healthy control | Liver |
| Kdm4a | -0.24609 | 7.422476 | 33.15065 | 2.69E-07 | 4.75E-07 | Healthy control | Liver |
| Kdm4b | -0.62743 | 5.81509 | 31.8486 | 4.20E-07 | 7.34E-07 | Healthy control | Liver |
| Kdm4c | 0.485744 | 6.133314 | 55.46526 | 3.21E-10 | 6.60E-10 | Infection | Liver |
| Kdm5a | 0.494301 | 7.886845 | 115.8094 | 6.12E-16 | 1.70E-15 | Infection | Liver |
| Kdm5b | -0.05133 | 7.952028 | 1.125251 | 2.93E-01 | 3.19E-01 | Healthy control | Liver |
| Kdm5c | 0.231778 | 8.516627 | 26.103 | 3.20E-06 | 5.32E-06 | Infection | Liver |
| Kdm6a | 0.484405 | 6.731964 | 66.11003 | 2.03E-11 | 4.42E-11 | Infection | Liver |
| Kdm6b | 0.862926 | 7.71119 | 256.0501 | 6.11E-24 | 3.17E-23 | Infection | Liver |
| Kdm7a | 0.117499 | 6.109605 | 3.608567 | 6.20E-02 | 7.40E-02 | Infection | Liver |
| Kdm8 | -0.05118 | 4.229024 | 0.201406 | 6.55E-01 | 6.79E-01 | Healthy control | Liver |
| Kdr | -1.74966 | 7.86637 | 344.2476 | 2.64E-27 | 1.93E-26 | Healthy control | Liver |
| Keap1 | -0.60648 | 6.74625 | 133.918 | 2.85E-17 | 8.67E-17 | Healthy control | Liver |
| Khdrbs1 | 0.72229 | 7.704724 | 208.2135 | 1.07E-21 | 4.48E-21 | Infection | Liver |
| Khdrbs2 | 1.797216 | 1.613712 | 12.09194 | 7.43E-04 | 1.07E-03 | Infection | Liver |
| Khsrp | -0.09927 | 7.521102 | 5.553085 | 2.16E-02 | 2.73E-02 | Healthy control | Liver |

| | | | | | | | |
|---|---|---|---|---|---|---|---|
| Kif19a | 0.420697 | 2.109476 | 1.818416 | 1.80E-01 | 2.02E-01 | Infection | Liver |
| Kif1b | -1.14206 | 9.147373 | 507.5476 | 5.89E-32 | 7.16E-31 | Healthy control | Liver |
| Kif1c | -1.32657 | 9.836075 | 696.1327 | 6.84E-36 | 1.51E-34 | Healthy control | Liver |
| Aire | 2.888323 | 1.514101 | 26.9239 | 2.47E-06 | 4.13E-06 | Infection | Liver |
| Ephb3 | 2.887847 | 3.897688 | 185.8517 | 1.74E-21 | 7.21E-21 | Infection | Liver |
| Cd180 | 2.887535 | 5.433064 | 545.457 | 1.24E-32 | 1.64E-31 | Infection | Liver |
| Pou3f3 | 2.884617 | 1.497017 | 24.62964 | 4.97E-06 | 8.22E-06 | Infection | Liver |
| Kif2b | 0.851991 | 1.256106 | 1.97705 | 1.68E-01 | 1.89E-01 | Infection | Liver |
| Ret | 2.884038 | 1.512819 | 22.77626 | 1.14E-05 | 1.85E-05 | Infection | Liver |
| Kif3a | 0.9639 | 3.858577 | 49.04297 | 2.02E-09 | 3.97E-09 | Infection | Liver |
| Kif3b | 0.495495 | 6.509538 | 45.80868 | 4.94E-09 | 9.55E-09 | Infection | Liver |
| Klc3 | 2.882473 | 2.699945 | 136.5472 | 1.23E-20 | 4.78E-20 | Infection | Liver |
| Kif5a | 0.524532 | 2.011643 | 3.198077 | 7.63E-02 | 9.01E-02 | Infection | Liver |
| Kif5b | 0.310911 | 8.753191 | 43.19357 | 1.08E-08 | 2.05E-08 | Infection | Liver |
| Mycn | 2.877715 | 4.723059 | 349.2455 | 1.28E-27 | 9.78E-27 | Infection | Liver |
| Kifap3 | 0.501362 | 5.32173 | 26.51795 | 2.76E-06 | 4.61E-06 | Infection | Liver |
| Rsph9 | 2.871175 | 2.844486 | 203.5973 | 3.49E-26 | 2.26E-25 | Infection | Liver |
| Kiss1 | -1.22519 | 1.622489 | 6.55176 | 1.20E-02 | 1.56E-02 | Healthy control | Liver |
| Kiss1r | -0.00853 | 2.622052 | 0.000935 | 9.76E-01 | 9.78E-01 | Healthy control | Liver |
| Kit | -0.26594 | 5.370437 | 3.477783 | 6.68E-02 | 7.94E-02 | Healthy control | Liver |
| Kitl | 1.285179 | 5.600377 | 103.3897 | 6.23E-15 | 1.64E-14 | Infection | Liver |
| Kl | -0.63675 | 1.424964 | 1.954941 | 1.65E-01 | 1.87E-01 | Healthy control | Liver |
| Klb | -1.41683 | 6.965722 | 141.8618 | 8.53E-18 | 2.68E-17 | Healthy control | Liver |
| Klc1 | 0.335232 | 7.189901 | 18.7785 | 5.35E-05 | 8.32E-05 | Infection | Liver |
| Klc2 | 0.210235 | 6.588152 | 23.50821 | 8.42E-06 | 1.37E-05 | Infection | Liver |
| Drd4 | 2.870559 | 1.493363 | 26.6293 | 2.38E-06 | 3.99E-06 | Infection | Liver |
| Klf1 | 1.472754 | 4.438187 | 22.10621 | 1.61E-05 | 2.58E-05 | Infection | Liver |
| Klf10 | -0.09875 | 5.527241 | 0.839326 | 3.63E-01 | 3.90E-01 | Healthy control | Liver |
| Klf11 | 0.295448 | 6.213062 | 8.844635 | 4.15E-03 | 5.64E-03 | Infection | Liver |
| Klf12 | -0.8943 | 8.163537 | 120.7502 | 2.57E-16 | 7.29E-16 | Healthy control | Liver |
| Klf15 | -1.32263 | 8.084486 | 169.3162 | 1.47E-19 | 5.29E-19 | Healthy control | Liver |
| Klf17 | 1.714839 | 1.462152 | 11.96765 | 8.43E-04 | 1.21E-03 | Infection | Liver |
| Klf4 | 1.758526 | 3.294024 | 79.85883 | 3.91E-13 | 9.37E-13 | Infection | Liver |
| Synpo2 | 2.870126 | 3.699556 | 204.9972 | 4.20E-23 | 1.98E-22 | Infection | Liver |
| Klhl18 | 1.096338 | 6.181994 | 248.0769 | 1.39E-23 | 6.89E-23 | Infection | Liver |
| Klhl21 | 0.018209 | 7.164243 | 0.11028 | 7.41E-01 | 7.60E-01 | Infection | Liver |
| Klhl22 | -0.02836 | 5.602023 | 0.106236 | 7.46E-01 | 7.64E-01 | Healthy control | Liver |
| Klhl3 | 0.192683 | 7.246264 | 4.372405 | 4.05E-02 | 4.97E-02 | Infection | Liver |
| Klhl36 | -0.51896 | 6.296076 | 63.18926 | 4.24E-11 | 9.09E-11 | Healthy control | Liver |
| Klhl41 | 1.383816 | 1.971018 | 19.47396 | 2.13E-05 | 3.39E-05 | Infection | Liver |
| Klk1b4 | -0.95412 | 1.72321 | 5.737878 | 1.82E-02 | 2.32E-02 | Healthy control | Liver |
| Klk5 | 0.039398 | 1.184772 | 0.006049 | 9.39E-01 | 9.46E-01 | Infection | Liver |
| Klk6 | 0.667832 | 1.18558 | 2.378342 | 1.45E-01 | 1.65E-01 | Infection | Liver |
| Klra8 | 1.789354 | 2.030349 | 25.36895 | 1.58E-06 | 2.67E-06 | Infection | Liver |
| Klrb1 | 1.89067 | 1.272649 | 7.963467 | 7.69E-03 | 1.02E-02 | Infection | Liver |
| Ptger3 | 2.865312 | 1.990641 | 53.89345 | 7.29E-11 | 1.55E-10 | Infection | Liver |
| Klrc1 | 1.685494 | 2.08088 | 31.85338 | 1.05E-07 | 1.90E-07 | Infection | Liver |
| Klrd1 | 1.871246 | 1.776614 | 18.53922 | 3.65E-05 | 5.75E-05 | Infection | Liver |
| Sele | 2.854356 | 2.086655 | 62.42027 | 3.92E-12 | 8.84E-12 | Infection | Liver |
| Klrk1 | 0.762469 | 4.447137 | 23.194 | 9.64E-06 | 1.57E-05 | Infection | Liver |

| | | | | | | | |
|---|---|---|---|---|---|---|---|
| Kmt2a | 1.072107 | 7.32793 | 362.1511 | 6.72E-28 | 5.31E-27 | Infection | Liver |
| Kmt2b | 1.009407 | 7.637779 | 435.513 | 4.29E-30 | 4.27E-29 | Infection | Liver |
| Kmt2c | -0.22918 | 6.887918 | 13.67095 | 4.57E-04 | 6.69E-04 | Healthy control | Liver |
| Kmt2d | 0.072995 | 8.201168 | 1.749312 | 1.91E-01 | 2.13E-01 | Infection | Liver |
| Kmt2e | 0.325513 | 8.311775 | 79.32864 | 8.93E-13 | 2.09E-12 | Infection | Liver |
| Kmt5b | 0.380329 | 7.417326 | 69.73865 | 8.33E-12 | 1.85E-11 | Infection | Liver |
| Kmt5c | -0.2628 | 6.311199 | 13.45089 | 5.03E-04 | 7.35E-04 | Healthy control | Liver |
| Kndc1 | 0.89874 | 1.268429 | 2.105495 | 1.54E-01 | 1.75E-01 | Infection | Liver |
| Tgfb2 | 2.8487 | 3.323687 | 113.5888 | 2.25E-17 | 6.88E-17 | Infection | Liver |
| Kcnma1 | 2.845494 | 2.039741 | 47.19162 | 6.81E-10 | 1.38E-09 | Infection | Liver |
| Kpna2 | 0.176688 | 6.473026 | 5.898436 | 1.80E-02 | 2.31E-02 | Infection | Liver |
| Kpnb1 | -0.72246 | 9.449902 | 441.2315 | 2.97E-30 | 3.03E-29 | Healthy control | Liver |
| Kras | 1.016596 | 8.21515 | 171.7928 | 1.05E-19 | 3.84E-19 | Infection | Liver |
| Kremen1 | 0.303083 | 4.536841 | 4.543704 | 3.70E-02 | 4.55E-02 | Infection | Liver |
| Kremen2 | 1.794619 | 1.496694 | 16.0614 | 1.33E-04 | 2.02E-04 | Infection | Liver |
| Krit1 | -0.06652 | 5.92957 | 1.153325 | 2.87E-01 | 3.13E-01 | Healthy control | Liver |
| Krt2 | 1.221535 | 1.258763 | 3.821553 | 5.79E-02 | 6.94E-02 | Infection | Liver |
| Krt20 | 1.129373 | 1.518928 | 5.140281 | 2.55E-02 | 3.20E-02 | Infection | Liver |
| Krt28 | 0.150642 | 1.362079 | 0.075645 | 7.84E-01 | 8.02E-01 | Infection | Liver |
| Krt40 | 0.76536 | 1.195457 | 2.778233 | 1.12E-01 | 1.30E-01 | Infection | Liver |
| Krt71 | 1.398544 | 1.309589 | 6.653781 | 1.26E-02 | 1.64E-02 | Infection | Liver |
| Krt76 | 0.161272 | 1.220305 | 0.073891 | 7.88E-01 | 8.05E-01 | Infection | Liver |
| Krt78 | 1.4506 | 1.446647 | 8.337883 | 4.99E-03 | 6.73E-03 | Infection | Liver |
| Krt79 | 0.497076 | 1.508979 | 0.970341 | 3.27E-01 | 3.53E-01 | Infection | Liver |
| Krt82 | 0.765731 | 1.2314 | 1.796834 | 1.90E-01 | 2.12E-01 | Infection | Liver |
| Cdkn2b | 2.845382 | 2.226021 | 71.86871 | 1.58E-13 | 3.86E-13 | Infection | Liver |
| Kyat1 | -3.55401 | 7.604822 | 1710.955 | 1.89E-47 | 6.73E-45 | Healthy control | Liver |
| Dpp10 | 2.8441 | 1.416846 | 15.3881 | 2.24E-04 | 3.35E-04 | Infection | Liver |
| L3hypdh | -1.58994 | 5.931164 | 308.9952 | 5.30E-26 | 3.37E-25 | Healthy control | Liver |
| L3mbtl2 | -0.56282 | 6.732819 | 93.04025 | 4.74E-14 | 1.19E-13 | Healthy control | Liver |
| Lacc1 | 0.833205 | 4.605549 | 60.90155 | 8.01E-11 | 1.70E-10 | Infection | Liver |
| Lag3 | 1.496757 | 4.781344 | 255.8634 | 8.03E-24 | 4.10E-23 | Infection | Liver |
| Lama1 | -1.53747 | 4.912668 | 144.4175 | 5.65E-18 | 1.79E-17 | Healthy control | Liver |
| Cd83 | 2.843561 | 5.312118 | 375.3995 | 3.63E-28 | 2.95E-27 | Infection | Liver |
| Lama3 | 1.000019 | 2.658647 | 17.10164 | 8.70E-05 | 1.34E-04 | Infection | Liver |
| Lama4 | 1.689491 | 6.08551 | 104.5429 | 6.34E-15 | 1.66E-14 | Infection | Liver |
| Lama5 | 1.20976 | 4.712699 | 104.6286 | 5.33E-15 | 1.41E-14 | Infection | Liver |
| Kif20a | 2.841524 | 5.415727 | 346.3778 | 3.17E-27 | 2.27E-26 | Infection | Liver |
| Lamb2 | 1.288831 | 6.45782 | 77.31474 | 1.69E-12 | 3.87E-12 | Infection | Liver |
| Lamb3 | 0.075713 | 3.963285 | 0.142439 | 7.07E-01 | 7.28E-01 | Infection | Liver |
| Lamc1 | 1.444129 | 7.785439 | 199.3277 | 3.08E-21 | 1.25E-20 | Infection | Liver |
| Lamp1 | -0.46408 | 10.76478 | 118.1213 | 4.06E-16 | 1.14E-15 | Healthy control | Liver |
| Lamp2 | 0.248422 | 11.21198 | 11.13968 | 1.42E-03 | 2.00E-03 | Infection | Liver |
| Cdh9 | 2.836181 | 1.884735 | 35.752 | 4.65E-08 | 8.51E-08 | Infection | Liver |
| Lamp5 | 0.951774 | 1.19806 | 3.406523 | 8.12E-02 | 9.55E-02 | Infection | Liver |
| Lamtor1 | -0.80096 | 8.233245 | 344.1498 | 2.64E-27 | 1.93E-26 | Healthy control | Liver |
| Lamtor2 | -1.18142 | 9.303066 | 616.1034 | 2.34E-34 | 3.90E-33 | Healthy control | Liver |
| Lamtor3 | 1.190317 | 7.380159 | 216.9968 | 3.88E-22 | 1.70E-21 | Infection | Liver |
| Lamtor4 | -1.16626 | 10.05324 | 666.0308 | 2.47E-35 | 5.10E-34 | Healthy control | Liver |
| Lamtor5 | -1.1571 | 6.223963 | 291.543 | 2.26E-25 | 1.33E-24 | Healthy control | Liver |

| | | | | | | | |
|---|---|---|---|---|---|---|---|
| Lancl2 | 0.033548 | 5.720831 | 0.226174 | 6.36E-01 | 6.60E-01 | Infection | Liver |
| Laptm4b | -0.82079 | 6.758282 | 97.94788 | 1.77E-14 | 4.51E-14 | Healthy control | Liver |
| Nmu | 2.835543 | 2.169795 | 34.19357 | 6.56E-08 | 1.19E-07 | Infection | Liver |
| Large2 | 1.423019 | 2.348595 | 26.82208 | 1.12E-06 | 1.92E-06 | Infection | Liver |
| Lasp1 | 0.000861 | 8.829023 | 0.000281 | 9.87E-01 | 9.88E-01 | Infection | Liver |
| Spn | 2.831658 | 4.011247 | 174.0954 | 1.87E-20 | 7.20E-20 | Infection | Liver |
| Lats1 | -0.67032 | 7.541788 | 173.4809 | 8.34E-20 | 3.08E-19 | Healthy control | Liver |
| Lats2 | -0.22523 | 5.960025 | 6.850902 | 1.11E-02 | 1.45E-02 | Healthy control | Liver |
| Lbr | 0.070812 | 6.241814 | 0.538866 | 4.66E-01 | 4.93E-01 | Infection | Liver |
| Lbx1 | 1.058904 | 1.24458 | 2.64948 | 1.13E-01 | 1.30E-01 | Infection | Liver |
| Gjc1 | 2.829551 | 4.139546 | 200.6618 | 6.63E-22 | 2.86E-21 | Infection | Liver |
| Lcp2 | 0.904667 | 6.888475 | 120.8436 | 2.53E-16 | 7.21E-16 | Infection | Liver |
| Ldah | -0.92066 | 7.656353 | 296.9148 | 1.33E-25 | 8.02E-25 | Healthy control | Liver |
| Ldb1 | 0.131836 | 7.849356 | 4.142945 | 4.60E-02 | 5.60E-02 | Infection | Liver |
| Ldb2 | -0.67457 | 5.056377 | 39.76838 | 3.18E-08 | 5.86E-08 | Healthy control | Liver |
| Ldha | -2.51599 | 10.37717 | 1107.172 | 7.50E-42 | 5.69E-40 | Healthy control | Liver |
| Ldhb | -0.07356 | 5.06132 | 0.272832 | 6.03E-01 | 6.28E-01 | Healthy control | Liver |
| Ldlr | -1.03468 | 7.735314 | 152.8206 | 1.53E-18 | 5.09E-18 | Healthy control | Liver |
| Ldlrad4 | 0.576139 | 6.369571 | 69.2788 | 9.34E-12 | 2.07E-11 | Infection | Liver |
| Lect2 | -0.87038 | 9.938772 | 106.546 | 3.33E-15 | 8.88E-15 | Healthy control | Liver |
| Lef1 | 1.086915 | 2.036738 | 10.95588 | 1.23E-03 | 1.74E-03 | Infection | Liver |
| Lefty1 | 1.624433 | 2.169265 | 31.89809 | 1.16E-07 | 2.10E-07 | Infection | Liver |
| Ano5 | 2.829069 | 1.695971 | 36.84819 | 4.32E-08 | 7.93E-08 | Infection | Liver |
| Leo1 | 0.013728 | 6.803876 | 0.05053 | 8.23E-01 | 8.38E-01 | Infection | Liver |
| Lep | -0.02876 | 1.232488 | 0.002758 | 9.58E-01 | 9.62E-01 | Healthy control | Liver |
| Lepr | 1.825771 | 5.621514 | 154.8872 | 1.19E-18 | 4.00E-18 | Infection | Liver |
| Leprot | 0.2705 | 6.470953 | 14.1655 | 3.69E-04 | 5.44E-04 | Infection | Liver |
| Letmd1 | -0.53508 | 6.856351 | 101.0174 | 9.65E-15 | 2.51E-14 | Healthy control | Liver |
| Lfng | 0.957747 | 5.973228 | 159.8345 | 5.61E-19 | 1.92E-18 | Infection | Liver |
| Lgals9 | -1.86474 | 11.89342 | 848.2806 | 2.08E-38 | 7.13E-37 | Healthy control | Liver |
| Pax5 | 2.82855 | 3.253227 | 123.4714 | 1.69E-18 | 5.61E-18 | Infection | Liver |
| Lgr4 | -0.41764 | 7.124973 | 10.81047 | 1.67E-03 | 2.34E-03 | Healthy control | Liver |
| Lgr5 | -1.16896 | 5.009471 | 56.57687 | 2.46E-10 | 5.08E-10 | Healthy control | Liver |
| Plk1 | 2.818248 | 3.560279 | 148.5665 | 1.06E-19 | 3.87E-19 | Infection | Liver |
| Lhb | 1.593821 | 1.384509 | 8.255 | 5.40E-03 | 7.27E-03 | Infection | Liver |
| Lhcgr | 0.530183 | 1.244294 | 0.811702 | 3.74E-01 | 4.01E-01 | Infection | Liver |
| Lhfpl5 | 1.469828 | 1.35679 | 5.514799 | 2.19E-02 | 2.78E-02 | Infection | Liver |
| Lhx1 | 1.140896 | 1.285596 | 3.545466 | 6.55E-02 | 7.78E-02 | Infection | Liver |
| Lhx2 | -1.59504 | 4.53308 | 117.7519 | 3.73E-16 | 1.05E-15 | Healthy control | Liver |
| Lhx3 | 1.023303 | 1.274342 | 3.932481 | 5.41E-02 | 6.51E-02 | Infection | Liver |
| Lhx4 | 1.216028 | 1.411153 | 4.208616 | 4.35E-02 | 5.31E-02 | Infection | Liver |
| Lhx5 | 1.170203 | 1.21774 | 5.43697 | 2.77E-02 | 3.45E-02 | Infection | Liver |
| Lhx6 | -0.62811 | 4.726825 | 13.84983 | 4.24E-04 | 6.23E-04 | Healthy control | Liver |
| Lhx8 | 1.903863 | 1.381537 | 7.095037 | 1.03E-02 | 1.35E-02 | Infection | Liver |
| Aim2 | 2.81757 | 5.525303 | 479.7456 | 4.24E-31 | 4.71E-30 | Infection | Liver |
| Far2 | 2.814771 | 1.793569 | 29.65012 | 6.30E-07 | 1.09E-06 | Infection | Liver |
| Lifr | -2.45848 | 10.52892 | 1027.668 | 6.98E-41 | 4.26E-39 | Healthy control | Liver |
| Lig4 | -0.65099 | 4.352683 | 23.77866 | 7.59E-06 | 1.24E-05 | Healthy control | Liver |
| Lilra5 | -0.12995 | 5.870246 | 0.914083 | 3.43E-01 | 3.69E-01 | Healthy control | Liver |
| Lima1 | -0.14059 | 8.564115 | 7.58588 | 7.67E-03 | 1.02E-02 | Healthy control | Liver |

| | | | | | | | |
|---|---|---|---|---|---|---|---|
| Limd1 | -0.19648 | 7.299559 | 7.93824 | 6.45E-03 | 8.63E-03 | Healthy control | Liver |
| Limk1 | 1.400362 | 5.79222 | 166.31 | 2.29E-19 | 8.12E-19 | Infection | Liver |
| Limk2 | 1.330038 | 6.444694 | 462.7673 | 8.08E-31 | 8.66E-30 | Infection | Liver |
| Nxph4 | 2.814389 | 1.534576 | 31.83495 | 3.11E-07 | 5.46E-07 | Infection | Liver |
| Lin28b | 1.743262 | 1.390782 | 7.837761 | 6.65E-03 | 8.89E-03 | Infection | Liver |
| Lin7c | -0.43545 | 7.382021 | 84.31825 | 2.97E-13 | 7.17E-13 | Healthy control | Liver |
| Lingo1 | 1.488929 | 1.987097 | 23.34869 | 3.86E-06 | 6.41E-06 | Infection | Liver |
| Lipe | -1.03551 | 5.82396 | 105.9083 | 3.90E-15 | 1.03E-14 | Healthy control | Liver |
| Lipg | -0.91573 | 6.656248 | 75.41496 | 2.19E-12 | 4.99E-12 | Healthy control | Liver |
| Ltbp1 | 2.814296 | 5.384196 | 378.9552 | 2.69E-28 | 2.20E-27 | Infection | Liver |
| Litaf | 0.426135 | 8.770255 | 108.8051 | 2.19E-15 | 5.88E-15 | Infection | Liver |
| Llgl2 | -1.96827 | 6.649077 | 477.709 | 3.56E-31 | 3.99E-30 | Healthy control | Liver |
| Lmna | 1.108711 | 8.819154 | 218.2231 | 3.36E-22 | 1.48E-21 | Infection | Liver |
| Lmnb1 | 1.274557 | 7.328538 | 303.6365 | 7.40E-26 | 4.61E-25 | Infection | Liver |
| Lmnb2 | -0.42332 | 4.939785 | 18.35111 | 6.40E-05 | 9.91E-05 | Healthy control | Liver |
| Lmo2 | 1.154257 | 5.350373 | 131.7022 | 4.24E-17 | 1.28E-16 | Infection | Liver |
| Lmo4 | 1.185976 | 7.350325 | 257.5031 | 5.30E-24 | 2.77E-23 | Infection | Liver |
| Lmx1a | 1.867799 | 1.290047 | 11.49433 | 1.48E-03 | 2.08E-03 | Infection | Liver |
| Lmx1b | -0.93731 | 1.265 | 2.674871 | 1.09E-01 | 1.26E-01 | Healthy control | Liver |
| Lnpk | -0.0867 | 5.313513 | 1.314661 | 2.56E-01 | 2.81E-01 | Healthy control | Liver |
| Slc6a8 | 2.811146 | 5.490583 | 213.1768 | 9.72E-22 | 4.11E-21 | Infection | Liver |
| Scn2a | 2.807974 | 2.015748 | 66.34178 | 1.33E-12 | 3.07E-12 | Infection | Liver |
| Prl3d1 | 2.807816 | 2.043674 | 30.65458 | 2.60E-07 | 4.59E-07 | Infection | Liver |
| Lpar2 | 0.960152 | 2.359009 | 14.98794 | 1.98E-04 | 2.98E-04 | Infection | Liver |
| Lpar5 | 1.290422 | 1.983382 | 22.27655 | 6.25E-06 | 1.03E-05 | Infection | Liver |
| Lpar6 | 0.070505 | 7.761261 | 1.085219 | 3.01E-01 | 3.28E-01 | Infection | Liver |
| Lpin1 | -0.33081 | 6.798081 | 11.50442 | 1.20E-03 | 1.70E-03 | Healthy control | Liver |
| Lpin2 | -0.18881 | 8.540681 | 4.651568 | 3.48E-02 | 4.30E-02 | Healthy control | Liver |
| Lpin3 | -0.00885 | 3.996051 | 0.004326 | 9.48E-01 | 9.53E-01 | Healthy control | Liver |
| Plcb2 | 2.806456 | 5.668791 | 625.8318 | 2.22E-34 | 3.70E-33 | Infection | Liver |
| Lpp | -0.43695 | 8.774053 | 79.09839 | 9.40E-13 | 2.19E-12 | Healthy control | Liver |
| Lratd2 | -0.57726 | 6.744488 | 34.0978 | 1.96E-07 | 3.49E-07 | Healthy control | Liver |
| Lrig1 | 0.196132 | 6.484202 | 7.035262 | 1.01E-02 | 1.33E-02 | Infection | Liver |
| Lrp1 | -0.53705 | 11.82172 | 125.6352 | 1.11E-16 | 3.25E-16 | Healthy control | Liver |
| Lrp12 | 0.704746 | 4.978086 | 48.3931 | 2.37E-09 | 4.65E-09 | Infection | Liver |
| Lrp2 | 1.213407 | 1.363945 | 5.571441 | 2.10E-02 | 2.67E-02 | Infection | Liver |
| Lrp5 | -0.15315 | 8.088476 | 10.42646 | 1.97E-03 | 2.75E-03 | Healthy control | Liver |
| Lrp6 | -0.4102 | 8.401768 | 69.59663 | 8.61E-12 | 1.91E-11 | Healthy control | Liver |
| Foxp3 | 2.803197 | 1.945127 | 61.19974 | 9.00E-12 | 2.00E-11 | Infection | Liver |
| Lrpprc | -1.47956 | 8.601373 | 711.727 | 3.60E-36 | 8.54E-35 | Healthy control | Liver |
| Lrrc15 | 0.299559 | 1.339439 | 0.305653 | 5.82E-01 | 6.07E-01 | Infection | Liver |
| Lrrc52 | 1.107662 | 1.256039 | 3.387628 | 7.33E-02 | 8.67E-02 | Infection | Liver |
| Tgfbr2 | 2.797435 | 8.883565 | 609.5025 | 4.95E-34 | 7.74E-33 | Infection | Liver |
| Lrrk1 | 0.394504 | 6.362788 | 19.62471 | 3.81E-05 | 5.99E-05 | Infection | Liver |
| Lrrk2 | 1.881368 | 6.23684 | 349.7578 | 1.78E-27 | 1.35E-26 | Infection | Liver |
| Lrrtm1 | 1.08697 | 1.243915 | 3.000712 | 9.24E-02 | 1.08E-01 | Infection | Liver |
| Casq2 | 2.796957 | 1.591814 | 36.07935 | 6.26E-08 | 1.14E-07 | Infection | Liver |
| Lsm14a | 1.158845 | 8.302664 | 677.7189 | 1.49E-35 | 3.17E-34 | Infection | Liver |
| Lsr | 0.498614 | 10.28998 | 26.56613 | 2.70E-06 | 4.51E-06 | Infection | Liver |
| Lss | 1.165142 | 8.587741 | 91.22109 | 8.72E-14 | 2.16E-13 | Infection | Liver |

| | | | | | | | |
|---|---|---|---|---|---|---|---|
| Wnt16 | 2.796493 | 1.799487 | 31.6938 | 2.19E-07 | 3.90E-07 | Infection | Liver |
| Lta4h | 0.62145 | 8.013007 | 120.7641 | 2.56E-16 | 7.28E-16 | Infection | Liver |
| Myf5 | 2.79207 | 1.801554 | 28.96051 | 6.32E-07 | 1.09E-06 | Infection | Liver |
| Rab7b | 2.790537 | 4.439146 | 378.7502 | 1.15E-28 | 9.78E-28 | Infection | Liver |
| Ltbr | -0.62311 | 7.871983 | 83.38707 | 3.64E-13 | 8.73E-13 | Healthy control | Liver |
| Luc7l3 | 0.707443 | 8.476119 | 239.0411 | 3.48E-23 | 1.66E-22 | Infection | Liver |
| Anln | 2.781426 | 5.340618 | 344.1319 | 3.80E-27 | 2.69E-26 | Infection | Liver |
| Ercc6l | 2.777244 | 2.931853 | 160.0862 | 2.39E-22 | 1.06E-21 | Infection | Liver |
| Lvrn | 1.628745 | 1.416477 | 10.12809 | 2.10E-03 | 2.92E-03 | Infection | Liver |
| Cr2 | 2.7762 | 2.468847 | 60.88193 | 2.82E-12 | 6.38E-12 | Infection | Liver |
| Ly6g5b | 0.30897 | 1.610789 | 0.553695 | 4.58E-01 | 4.86E-01 | Infection | Liver |
| Ly6g6d | 1.315689 | 1.582651 | 8.89278 | 3.50E-03 | 4.79E-03 | Infection | Liver |
| Esyt3 | 2.773399 | 1.753755 | 28.69166 | 7.44E-07 | 1.28E-06 | Infection | Liver |
| Ly6h | 1.471129 | 1.566892 | 11.24476 | 1.10E-03 | 1.57E-03 | Infection | Liver |
| Itga6 | 2.771341 | 3.39865 | 165.2497 | 3.10E-21 | 1.26E-20 | Infection | Liver |
| Pak1 | 2.771214 | 6.046591 | 934.8941 | 1.54E-39 | 6.53E-38 | Infection | Liver |
| Ly6m | 0.956268 | 1.339616 | 3.020776 | 8.70E-02 | 1.02E-01 | Infection | Liver |
| Ly75 | 1.014697 | 5.805639 | 115.0309 | 7.16E-16 | 1.98E-15 | Infection | Liver |
| Syk | 2.771055 | 8.094574 | 1038.149 | 5.23E-41 | 3.27E-39 | Infection | Liver |
| Ly96 | 0.649946 | 6.262189 | 128.8406 | 6.56E-17 | 1.95E-16 | Infection | Liver |
| Lyar | -0.72458 | 5.506532 | 67.26497 | 1.56E-11 | 3.42E-11 | Healthy control | Liver |
| Lyg1 | 0.340861 | 1.197312 | 0.390437 | 5.40E-01 | 5.67E-01 | Infection | Liver |
| Lyn | 0.731152 | 7.40023 | 190.9598 | 8.63E-21 | 3.39E-20 | Infection | Liver |
| Myf6 | 2.76848 | 1.672931 | 18.93274 | 4.01E-05 | 6.29E-05 | Infection | Liver |
| Lypd6 | 1.119828 | 1.730059 | 7.532645 | 6.93E-03 | 9.23E-03 | Infection | Liver |
| Lypd6b | 0.992748 | 1.318965 | 2.940727 | 9.23E-02 | 1.08E-01 | Infection | Liver |
| Lyplal1 | -1.34805 | 6.242516 | 226.5077 | 1.41E-22 | 6.39E-22 | Healthy control | Liver |
| Lyst | 0.898375 | 8.271497 | 383.69 | 1.39E-28 | 1.17E-27 | Infection | Liver |
| Lyve1 | 1.514706 | 5.398056 | 69.16271 | 1.10E-11 | 2.44E-11 | Infection | Liver |
| Lyzl6 | 1.360404 | 1.302406 | 4.12414 | 4.76E-02 | 5.78E-02 | Infection | Liver |
| Lztr1 | 0.167183 | 7.633292 | 23.40129 | 8.76E-06 | 1.43E-05 | Infection | Liver |
| Mad1l1 | -0.15277 | 6.637012 | 8.157198 | 5.79E-03 | 7.78E-03 | Healthy control | Liver |
| Mad2l1 | 1.726368 | 3.787715 | 132.7503 | 3.41E-17 | 1.03E-16 | Infection | Liver |
| Mad2l2 | -0.54932 | 7.87154 | 60.0014 | 9.58E-11 | 2.02E-10 | Healthy control | Liver |
| Meox2 | 2.765183 | 1.860161 | 35.97301 | 4.13E-08 | 7.59E-08 | Infection | Liver |
| Mael | 1.732807 | 1.30947 | 8.798765 | 4.52E-03 | 6.11E-03 | Infection | Liver |
| Maf | -0.26337 | 9.147548 | 17.18399 | 1.03E-04 | 1.57E-04 | Healthy control | Liver |
| Maf1 | -0.73881 | 6.389696 | 100.5941 | 1.05E-14 | 2.74E-14 | Healthy control | Liver |
| Mafa | 1.072006 | 1.77271 | 10.75418 | 1.33E-03 | 1.88E-03 | Infection | Liver |
| Mafb | 1.039284 | 7.720496 | 92.53057 | 5.75E-14 | 1.44E-13 | Infection | Liver |
| Maff | 1.866825 | 6.16433 | 221.047 | 2.55E-22 | 1.13E-21 | Infection | Liver |
| Mafk | -1.03161 | 3.915166 | 65.53692 | 2.02E-11 | 4.40E-11 | Healthy control | Liver |
| Mag | 1.983648 | 3.197491 | 108.6453 | 3.95E-16 | 1.11E-15 | Infection | Liver |
| Cacnb1 | 2.763973 | 4.617629 | 456.8562 | 1.10E-30 | 1.16E-29 | Infection | Liver |
| Magoh | -1.17539 | 6.858147 | 297.3179 | 1.30E-25 | 7.86E-25 | Healthy control | Liver |
| Maip1 | -0.22807 | 7.159627 | 13.48331 | 4.96E-04 | 7.25E-04 | Healthy control | Liver |
| Malt1 | 1.518198 | 5.684881 | 386.0127 | 1.27E-28 | 1.08E-27 | Infection | Liver |
| Maml1 | -0.70501 | 6.099839 | 85.20955 | 2.48E-13 | 6.00E-13 | Healthy control | Liver |
| Man2b2 | 0.385567 | 8.710568 | 44.47976 | 7.33E-09 | 1.40E-08 | Infection | Liver |
| Maoa | -0.3652 | 6.40459 | 21.14852 | 2.09E-05 | 3.33E-05 | Healthy control | Liver |

| | | | | | | | |
|---|---|---|---|---|---|---|---|
| Maob | -1.61828 | 9.160101 | 277.5774 | 7.66E-25 | 4.33E-24 | Healthy control | Liver |
| Cxcr3 | 2.758826 | 4.188972 | 406.8366 | 7.68E-30 | 7.36E-29 | Infection | Liver |
| Map1lc3a | -0.36109 | 8.164898 | 43.47942 | 9.91E-09 | 1.88E-08 | Healthy control | Liver |
| Map2 | 0.609966 | 4.532431 | 22.35579 | 1.32E-05 | 2.14E-05 | Infection | Liver |
| Map2k1 | -0.76544 | 7.872182 | 208.6506 | 1.01E-21 | 4.28E-21 | Healthy control | Liver |
| Map2k2 | -1.11389 | 8.657157 | 860.7725 | 1.36E-38 | 4.78E-37 | Healthy control | Liver |
| Map2k3 | -2.3493 | 6.395497 | 909.649 | 3.50E-39 | 1.39E-37 | Healthy control | Liver |
| Map2k4 | 0.396887 | 6.238725 | 26.32846 | 2.95E-06 | 4.91E-06 | Infection | Liver |
| Map2k5 | -0.76516 | 6.297161 | 149.2408 | 2.65E-18 | 8.62E-18 | Healthy control | Liver |
| Map3k1 | 1.233862 | 6.047406 | 211.7152 | 7.25E-22 | 3.11E-21 | Infection | Liver |
| Map3k11 | -0.63746 | 7.006269 | 136.4242 | 1.90E-17 | 5.85E-17 | Healthy control | Liver |
| Cdc20 | 2.756578 | 5.164755 | 190.6481 | 1.28E-20 | 4.98E-20 | Infection | Liver |
| Grm4 | 2.756383 | 1.767663 | 40.99861 | 1.14E-08 | 2.15E-08 | Infection | Liver |
| Map3k5 | -0.21736 | 7.323936 | 10.9452 | 1.55E-03 | 2.18E-03 | Healthy control | Liver |
| Map3k7 | -0.65442 | 7.675924 | 169.8618 | 1.36E-19 | 4.93E-19 | Healthy control | Liver |
| Map3k8 | 1.572682 | 4.876028 | 246.4721 | 1.99E-23 | 9.68E-23 | Infection | Liver |
| Map4 | -0.2747 | 7.864745 | 49.01279 | 1.94E-09 | 3.82E-09 | Healthy control | Liver |
| Cd3e | 2.755697 | 5.00575 | 318.3034 | 3.12E-26 | 2.04E-25 | Infection | Liver |
| Wnt7b | 2.752076 | 1.619106 | 34.45364 | 1.12E-07 | 2.02E-07 | Infection | Liver |
| Map7 | -0.74493 | 7.234005 | 52.29828 | 7.65E-10 | 1.54E-09 | Healthy control | Liver |
| Map9 | 0.278462 | 3.258219 | 3.446598 | 6.79E-02 | 8.06E-02 | Infection | Liver |
| Mapk1 | -0.46225 | 8.205546 | 125.5716 | 1.12E-16 | 3.29E-16 | Healthy control | Liver |
| Mapk10 | 1.805271 | 1.368388 | 9.631113 | 2.84E-03 | 3.92E-03 | Infection | Liver |
| Mapk11 | 0.564922 | 3.908921 | 15.55867 | 2.05E-04 | 3.07E-04 | Infection | Liver |
| Mapk14 | -1.68622 | 6.796452 | 892.9867 | 4.90E-39 | 1.84E-37 | Healthy control | Liver |
| Mapk3 | -0.83173 | 7.160955 | 149.5452 | 2.50E-18 | 8.13E-18 | Healthy control | Liver |
| Mapk7 | 1.308946 | 4.774388 | 122.085 | 2.29E-16 | 6.52E-16 | Infection | Liver |
| Mapk8 | 0.678529 | 6.90236 | 146.0727 | 4.21E-18 | 1.35E-17 | Infection | Liver |
| Mapk8ip1 | 1.718737 | 5.18553 | 166.0074 | 2.58E-19 | 9.10E-19 | Infection | Liver |
| Mapk8ip2 | 0.552721 | 1.35369 | 1.165761 | 2.84E-01 | 3.10E-01 | Infection | Liver |
| Mapk9 | -0.36173 | 7.694159 | 40.92598 | 2.17E-08 | 4.04E-08 | Healthy control | Liver |
| Mapkap1 | -1.44525 | 7.463433 | 961.5034 | 5.18E-40 | 2.45E-38 | Healthy control | Liver |
| Mapkapk2 | 0.381068 | 8.133936 | 61.71263 | 6.15E-11 | 1.31E-10 | Infection | Liver |
| Mapre1 | -0.39706 | 7.404772 | 47.18351 | 3.29E-09 | 6.42E-09 | Healthy control | Liver |
| Mapre2 | 1.134955 | 7.219284 | 526.4376 | 2.11E-32 | 2.70E-31 | Infection | Liver |
| Mapt | -2.02737 | 5.652259 | 228.3902 | 1.26E-22 | 5.76E-22 | Healthy control | Liver |
| 2-Mar | -0.90921 | 7.18261 | 363.8082 | 5.97E-28 | 4.75E-27 | Healthy control | Liver |
| 5-Mar | -1.83047 | 6.902148 | 841.3811 | 2.85E-38 | 9.34E-37 | Healthy control | Liver |
| Marcks | 1.3062 | 8.217337 | 302.062 | 8.45E-26 | 5.23E-25 | Infection | Liver |
| Marcksl1 | 1.991247 | 7.633131 | 781.5146 | 2.35E-37 | 6.85E-36 | Infection | Liver |
| Cyp27b1 | 2.751909 | 1.802335 | 27.83327 | 9.98E-07 | 1.71E-06 | Infection | Liver |
| Mark2 | 1.321243 | 7.360578 | 601.6116 | 4.69E-34 | 7.38E-33 | Infection | Liver |
| Mark3 | -0.32214 | 6.924802 | 29.9418 | 8.10E-07 | 1.39E-06 | Healthy control | Liver |
| Marveld2 | 0.189041 | 5.680219 | 2.628778 | 1.10E-01 | 1.27E-01 | Infection | Liver |
| Mas1 | 0.123783 | 2.470206 | 0.230804 | 6.32E-01 | 6.56E-01 | Infection | Liver |
| Mast2 | 0.726056 | 7.260002 | 155.9995 | 9.61E-19 | 3.24E-18 | Infection | Liver |
| Mast3 | -0.3377 | 7.141134 | 26.81674 | 2.47E-06 | 4.13E-06 | Healthy control | Liver |
| Matn1 | 1.725049 | 1.2965 | 9.543581 | 3.40E-03 | 4.65E-03 | Infection | Liver |
| Mavs | -0.92659 | 7.587728 | 130.1939 | 5.21E-17 | 1.55E-16 | Healthy control | Liver |
| Max | 1.257592 | 8.36988 | 363.5735 | 6.02E-28 | 4.78E-27 | Infection | Liver |

| | | | | | | | |
|---|---|---|---|---|---|---|---|
| Maz | 0.315376 | 6.95653 | 27.58212 | 1.87E-06 | 3.15E-06 | Infection | Liver |
| Mbd2 | 0.11085 | 8.37464 | 6.543905 | 1.29E-02 | 1.68E-02 | Infection | Liver |
| Mbd3 | -0.25241 | 8.734908 | 33.30439 | 2.55E-07 | 4.52E-07 | Healthy control | Liver |
| Mbd5 | -0.12718 | 5.263146 | 2.860314 | 9.57E-02 | 1.12E-01 | Healthy control | Liver |
| Mbl1 | -0.98674 | 9.779321 | 116.0751 | 5.83E-16 | 1.62E-15 | Healthy control | Liver |
| Mbl2 | -0.92996 | 10.83733 | 59.96076 | 1.10E-10 | 2.31E-10 | Healthy control | Liver |
| Mbnl1 | -0.30732 | 8.575107 | 46.94285 | 3.53E-09 | 6.88E-09 | Healthy control | Liver |
| Iqgap3 | 2.750141 | 2.677392 | 93.52478 | 2.11E-16 | 6.05E-16 | Infection | Liver |
| Mboat7 | -0.94113 | 8.238889 | 342.9549 | 2.90E-27 | 2.10E-26 | Healthy control | Liver |
| Mbtd1 | 0.544811 | 7.579757 | 138.1584 | 1.44E-17 | 4.46E-17 | Infection | Liver |
| Mc1r | 1.098658 | 1.329589 | 4.262465 | 4.35E-02 | 5.31E-02 | Infection | Liver |
| Mc3r | 0.617036 | 1.215572 | 1.3998 | 2.48E-01 | 2.73E-01 | Infection | Liver |
| Mc4r | 1.287139 | 1.332132 | 4.716931 | 3.41E-02 | 4.22E-02 | Infection | Liver |
| Mc5r | 1.697495 | 1.375231 | 8.011828 | 6.12E-03 | 8.21E-03 | Infection | Liver |
| Mcam | 1.684514 | 5.962317 | 132.289 | 4.11E-17 | 1.24E-16 | Infection | Liver |
| Mcl1 | 0.1831 | 9.898266 | 9.684862 | 2.79E-03 | 3.84E-03 | Infection | Liver |
| Mcm2 | 1.399698 | 5.203161 | 169.2869 | 1.61E-19 | 5.78E-19 | Infection | Liver |
| Mcph1 | 0.639194 | 5.568711 | 87.04371 | 1.68E-13 | 4.11E-13 | Infection | Liver |
| Mctp2 | -0.0557 | 5.954299 | 0.428922 | 5.15E-01 | 5.42E-01 | Healthy control | Liver |
| Mcu | -2.10896 | 8.228953 | 1235.5 | 2.81E-43 | 2.65E-41 | Healthy control | Liver |
| Trpc4 | 2.744341 | 1.483153 | 20.40883 | 2.77E-05 | 4.39E-05 | Infection | Liver |
| Mcur1 | -1.97501 | 5.650018 | 591.2462 | 9.90E-34 | 1.49E-32 | Healthy control | Liver |
| Nkain1 | 2.740747 | 2.627217 | 83.92751 | 5.42E-15 | 1.43E-14 | Infection | Liver |
| Mdm2 | 1.438825 | 7.641808 | 664.1932 | 2.69E-35 | 5.50E-34 | Infection | Liver |
| Mdm4 | 0.864907 | 7.992213 | 249.4178 | 1.19E-23 | 5.92E-23 | Infection | Liver |
| Mdp1 | -0.35635 | 6.644624 | 38.72186 | 4.34E-08 | 7.96E-08 | Healthy control | Liver |
| Me1 | -2.10088 | 7.66238 | 725.9228 | 2.07E-36 | 5.21E-35 | Healthy control | Liver |
| Meaf6 | 0.78881 | 5.16508 | 81.61995 | 5.53E-13 | 1.31E-12 | Infection | Liver |
| Mecom | 1.466603 | 3.892892 | 77.249 | 1.52E-12 | 3.51E-12 | Infection | Liver |
| Mecp2 | 0.011164 | 7.061429 | 0.045083 | 8.33E-01 | 8.47E-01 | Infection | Liver |
| Med1 | -0.45367 | 7.227011 | 138.4272 | 1.38E-17 | 4.29E-17 | Healthy control | Liver |
| Med12 | -0.14985 | 6.161 | 3.754592 | 5.71E-02 | 6.85E-02 | Healthy control | Liver |
| Med14 | -0.83864 | 5.537242 | 110.564 | 1.66E-15 | 4.50E-15 | Healthy control | Liver |
| Med15 | -0.32148 | 7.368669 | 41.01117 | 2.11E-08 | 3.94E-08 | Healthy control | Liver |
| Med21 | -0.70733 | 6.249665 | 106.5411 | 3.39E-15 | 9.01E-15 | Healthy control | Liver |
| Med24 | -0.55624 | 5.769867 | 83.34928 | 3.74E-13 | 8.96E-13 | Healthy control | Liver |
| Med6 | 0.299319 | 6.714608 | 28.67689 | 1.26E-06 | 2.15E-06 | Infection | Liver |
| Col4a1 | 2.740492 | 9.52125 | 599.7687 | 7.79E-34 | 1.20E-32 | Infection | Liver |
| Mef2a | -0.09025 | 7.422978 | 1.680756 | 2.00E-01 | 2.22E-01 | Healthy control | Liver |
| Mef2c | 1.787279 | 4.092555 | 156.2919 | 1.13E-18 | 3.80E-18 | Infection | Liver |
| Mef2d | -1.38634 | 6.991137 | 352.828 | 1.39E-27 | 1.06E-26 | Healthy control | Liver |
| Ins1 | 2.736565 | 1.606792 | 20.40759 | 2.31E-05 | 3.67E-05 | Infection | Liver |
| Mei4 | 1.191243 | 1.336593 | 3.365325 | 7.16E-02 | 8.48E-02 | Infection | Liver |
| Meiob | -1.39088 | 4.679487 | 91.1705 | 6.85E-14 | 1.70E-13 | Healthy control | Liver |
| Meis1 | 1.947782 | 6.664802 | 555.112 | 4.79E-33 | 6.64E-32 | Infection | Liver |
| Meis2 | -0.13678 | 6.456656 | 3.058626 | 8.52E-02 | 1.00E-01 | Healthy control | Liver |
| Chrm3 | 2.735537 | 1.800516 | 36.76926 | 3.31E-08 | 6.10E-08 | Infection | Liver |
| Memo1 | -0.35974 | 7.068475 | 38.47164 | 4.70E-08 | 8.60E-08 | Healthy control | Liver |
| Men1 | -0.1083 | 7.198163 | 4.073046 | 4.78E-02 | 5.80E-02 | Healthy control | Liver |
| Tenm4 | 2.735105 | 2.162409 | 69.3784 | 2.75E-13 | 6.64E-13 | Infection | Liver |

| | | | | | | | |
|---|---|---|---|---|---|---|---|
| Itgb7 | 2.726682 | 5.973526 | 439.3721 | 4.08E-30 | 4.11E-29 | Infection | Liver |
| Mep1a | 1.22613 | 1.294567 | 3.338274 | 7.39E-02 | 8.74E-02 | Infection | Liver |
| Mep1b | 0.468473 | 1.344984 | 0.668425 | 4.16E-01 | 4.44E-01 | Infection | Liver |
| Mepe | 1.244959 | 1.254876 | 4.643227 | 3.77E-02 | 4.64E-02 | Infection | Liver |
| Mertk | -0.27238 | 6.59022 | 14.39475 | 3.34E-04 | 4.94E-04 | Healthy control | Liver |
| Mesp1 | -0.92419 | 1.29156 | 2.671718 | 1.08E-01 | 1.25E-01 | Healthy control | Liver |
| Mesp2 | -0.06272 | 1.709181 | 0.023599 | 8.78E-01 | 8.90E-01 | Healthy control | Liver |
| Mest | 1.912245 | 5.889128 | 161.6766 | 4.70E-19 | 1.62E-18 | Infection | Liver |
| Met | -1.85237 | 8.34915 | 518.8983 | 3.17E-32 | 3.97E-31 | Healthy control | Liver |
| Metrnl | 1.698761 | 3.410998 | 76.60111 | 1.10E-12 | 2.55E-12 | Infection | Liver |
| Mettl14 | 0.11507 | 5.836961 | 3.268934 | 7.54E-02 | 8.89E-02 | Infection | Liver |
| Mettl21c | 1.739223 | 1.405776 | 6.750249 | 1.14E-02 | 1.49E-02 | Infection | Liver |
| Mettl3 | -0.27983 | 6.299853 | 30.32031 | 7.10E-07 | 1.22E-06 | Healthy control | Liver |
| Mettl7b | -1.84374 | 12.5036 | 441.991 | 2.83E-30 | 2.89E-29 | Healthy control | Liver |
| Ifng | 2.72589 | 2.279227 | 78.49061 | 1.30E-14 | 3.34E-14 | Infection | Liver |
| Mfn1 | -1.73081 | 7.19927 | 720.8803 | 2.56E-36 | 6.38E-35 | Healthy control | Liver |
| Mfn2 | -1.35122 | 8.180565 | 576.1043 | 1.62E-33 | 2.37E-32 | Healthy control | Liver |
| Mfrp | 1.194692 | 1.251989 | 5.199074 | 2.84E-02 | 3.53E-02 | Infection | Liver |
| Mfsd2a | -1.65309 | 6.344907 | 25.14779 | 5.33E-06 | 8.79E-06 | Healthy control | Liver |
| Foxm1 | 2.725785 | 4.143644 | 278.6891 | 2.48E-25 | 1.45E-24 | Infection | Liver |
| Mia | -0.52337 | 1.762825 | 2.092614 | 1.51E-01 | 1.71E-01 | Healthy control | Liver |
| Mia2 | -0.24963 | 10.5064 | 8.974096 | 3.90E-03 | 5.32E-03 | Healthy control | Liver |
| Mib1 | 0.288327 | 8.858259 | 58.6379 | 1.37E-10 | 2.86E-10 | Infection | Liver |
| Micu1 | -1.05604 | 6.62396 | 249.8588 | 1.16E-23 | 5.78E-23 | Healthy control | Liver |
| Micu2 | -0.13213 | 8.040219 | 7.656218 | 7.41E-03 | 9.85E-03 | Healthy control | Liver |
| Miga1 | 0.922753 | 3.545927 | 43.98511 | 8.53E-09 | 1.63E-08 | Infection | Liver |
| Miga2 | -1.49353 | 7.988279 | 609.8814 | 3.16E-34 | 5.16E-33 | Healthy control | Liver |
| Minar2 | 1.312937 | 1.345469 | 5.196521 | 2.60E-02 | 3.25E-02 | Infection | Liver |
| Mindy1 | -0.55691 | 7.884549 | 99.90838 | 1.19E-14 | 3.08E-14 | Healthy control | Liver |
| Mip | 0.895459 | 1.333634 | 2.263652 | 1.37E-01 | 1.57E-01 | Infection | Liver |
| Misp | 1.067637 | 2.576759 | 31.92553 | 2.12E-07 | 3.77E-07 | Infection | Liver |
| Pak3 | 2.72482 | 1.483203 | 13.19616 | 5.60E-04 | 8.13E-04 | Infection | Liver |
| S100b | 2.724201 | 1.471847 | 22.85968 | 1.14E-05 | 1.85E-05 | Infection | Liver |
| Gabbr1 | 2.720088 | 6.996388 | 785.8673 | 2.09E-37 | 6.12E-36 | Infection | Liver |
| Mkks | -0.50743 | 5.052841 | 27.11105 | 2.24E-06 | 3.76E-06 | Healthy control | Liver |
| Mkln1 | -0.70342 | 8.405676 | 241.9194 | 2.57E-23 | 1.24E-22 | Healthy control | Liver |
| Mknk1 | 0.464384 | 6.490961 | 71.69562 | 5.23E-12 | 1.17E-11 | Infection | Liver |
| Mknk2 | 0.175321 | 9.054965 | 11.48182 | 1.21E-03 | 1.72E-03 | Infection | Liver |
| Mkx | -0.62818 | 3.406796 | 16.35826 | 1.39E-04 | 2.10E-04 | Healthy control | Liver |
| Mlc1 | 1.303421 | 1.263421 | 5.007877 | 3.08E-02 | 3.82E-02 | Infection | Liver |
| Mlec | 0.431374 | 8.143677 | 83.7815 | 3.33E-13 | 8.02E-13 | Infection | Liver |
| Mlf1 | 1.272601 | 2.024017 | 9.313618 | 2.79E-03 | 3.84E-03 | Infection | Liver |
| Mlh1 | 0.186379 | 6.437627 | 8.543034 | 4.80E-03 | 6.49E-03 | Infection | Liver |
| Mlkl | 0.686804 | 6.292992 | 105.914 | 3.79E-15 | 1.00E-14 | Infection | Liver |
| Mllt10 | -0.59912 | 6.541367 | 97.02591 | 2.12E-14 | 5.42E-14 | Healthy control | Liver |
| Mllt6 | 0.268764 | 7.471215 | 10.89143 | 1.59E-03 | 2.23E-03 | Infection | Liver |
| Mlph | 1.371086 | 4.466399 | 119.3937 | 3.85E-16 | 1.08E-15 | Infection | Liver |
| Mlst8 | 0.590954 | 5.665117 | 79.64458 | 8.43E-13 | 1.98E-12 | Infection | Liver |
| Mlx | -0.73782 | 6.72587 | 169.8631 | 1.37E-19 | 4.97E-19 | Healthy control | Liver |
| Mlxipl | -1.77685 | 9.664477 | 257.6785 | 6.62E-24 | 3.42E-23 | Healthy control | Liver |

| | | | | | | | |
|---|---|---|---|---|---|---|---|
| Mlycd | -1.45947 | 7.743355 | 346.333 | 2.24E-27 | 1.66E-26 | Healthy control | Liver |
| Mmd2 | -0.98087 | 2.929584 | 7.52866 | 7.55E-03 | 1.00E-02 | Healthy control | Liver |
| Mme | -2.11198 | 7.635393 | 403.0668 | 3.69E-29 | 3.28E-28 | Healthy control | Liver |
| Il18r1 | 2.715148 | 2.708389 | 111.8646 | 2.51E-18 | 8.18E-18 | Infection | Liver |
| Disc1 | 2.714692 | 2.443801 | 89.52888 | 4.83E-16 | 1.35E-15 | Infection | Liver |
| Gja1 | 2.708699 | 7.774857 | 238.8545 | 2.12E-22 | 9.50E-22 | Infection | Liver |
| Mmp15 | -1.3898 | 6.463901 | 184.0172 | 2.14E-20 | 8.21E-20 | Healthy control | Liver |
| Cd48 | 2.707882 | 6.111346 | 639.7214 | 9.59E-35 | 1.75E-33 | Infection | Liver |
| Adcy1 | 2.707208 | 4.32717 | 325.0446 | 5.55E-27 | 3.86E-26 | Infection | Liver |
| Bcl11b | 2.706116 | 2.658586 | 116.2136 | 4.21E-19 | 1.46E-18 | Infection | Liver |
| Mmp28 | 1.67656 | 2.599885 | 68.91745 | 7.96E-13 | 1.87E-12 | Infection | Liver |
| Agrp | 2.704754 | 1.536989 | 23.41805 | 7.37E-06 | 1.21E-05 | Infection | Liver |
| Ly6i | 2.700247 | 1.552972 | 29.9602 | 1.31E-06 | 2.23E-06 | Infection | Liver |
| Il5ra | 2.698529 | 1.881068 | 44.86646 | 1.59E-09 | 3.13E-09 | Infection | Liver |
| Cacng3 | 2.698139 | 1.558272 | 24.6517 | 5.45E-06 | 8.99E-06 | Infection | Liver |
| Mms19 | -0.40054 | 6.827385 | 44.51302 | 7.27E-09 | 1.39E-08 | Healthy control | Liver |
| Mmut | -0.92625 | 9.357353 | 103.1809 | 6.32E-15 | 1.66E-14 | Healthy control | Liver |
| Fcgr3 | 2.69592 | 6.747192 | 548.9655 | 6.79E-33 | 9.23E-32 | Infection | Liver |
| Mob2 | -0.02777 | 7.388403 | 0.30192 | 5.85E-01 | 6.10E-01 | Healthy control | Liver |
| Mog | -0.19213 | 1.323995 | 0.111397 | 7.40E-01 | 7.59E-01 | Healthy control | Liver |
| Mos | 1.526676 | 1.315698 | 8.740466 | 4.80E-03 | 6.49E-03 | Infection | Liver |
| Mov10l1 | 1.419733 | 1.326415 | 7.168682 | 9.75E-03 | 1.28E-02 | Infection | Liver |
| Gpr22 | 2.69391 | 1.583838 | 25.31688 | 3.13E-06 | 5.21E-06 | Infection | Liver |
| Mpc1 | -1.98658 | 9.308667 | 512.001 | 4.61E-32 | 5.68E-31 | Healthy control | Liver |
| Mpc2 | -1.52722 | 9.500532 | 412.6736 | 1.89E-29 | 1.76E-28 | Healthy control | Liver |
| Mpi | -0.08493 | 5.484558 | 1.006032 | 3.20E-01 | 3.46E-01 | Healthy control | Liver |
| Tgm2 | 2.688503 | 11.34832 | 2173.574 | 8.99E-51 | 1.50E-47 | Infection | Liver |
| Mplkip | -0.02767 | 4.876125 | 0.109418 | 7.42E-01 | 7.61E-01 | Healthy control | Liver |
| Bcl11a | 2.688312 | 2.628774 | 79.63465 | 1.44E-14 | 3.70E-14 | Infection | Liver |
| Mpp3 | 0.834029 | 2.340131 | 12.85936 | 5.31E-04 | 7.75E-04 | Infection | Liver |
| Mprip | 0.045052 | 9.918868 | 1.205626 | 2.76E-01 | 3.03E-01 | Infection | Liver |
| Mrc1 | -0.00271 | 7.968934 | 0.000768 | 9.78E-01 | 9.80E-01 | Healthy control | Liver |
| Mre11a | 0.57872 | 6.306848 | 79.5019 | 8.63E-13 | 2.02E-12 | Infection | Liver |
| Vstm4 | 2.683979 | 7.51351 | 270.2531 | 7.67E-24 | 3.92E-23 | Infection | Liver |
| Mrm3 | -0.94848 | 5.474803 | 137.5054 | 1.71E-17 | 5.29E-17 | Healthy control | Liver |
| Mrnip | -2.24186 | 5.232508 | 408.2914 | 2.29E-29 | 2.10E-28 | Healthy control | Liver |
| Mrtfa | 0.395868 | 2.646645 | 4.512239 | 3.68E-02 | 4.54E-02 | Infection | Liver |
| Neurod1 | 2.68208 | 1.834872 | 21.71703 | 1.08E-05 | 1.75E-05 | Infection | Liver |
| Col4a2 | 2.677356 | 8.812779 | 609.7205 | 4.28E-34 | 6.85E-33 | Infection | Liver |
| Sgo2a | 2.675811 | 3.180736 | 167.1286 | 5.97E-22 | 2.58E-21 | Infection | Liver |
| Msh2 | 0.100769 | 5.770393 | 2.314207 | 1.33E-01 | 1.52E-01 | Infection | Liver |
| Epn3 | 2.673959 | 1.714849 | 38.08183 | 2.34E-08 | 4.36E-08 | Infection | Liver |
| Hoxa7 | 2.670564 | 1.489423 | 24.19356 | 5.37E-06 | 8.86E-06 | Infection | Liver |
| Msmo1 | 0.136147 | 9.072683 | 1.248802 | 2.68E-01 | 2.94E-01 | Infection | Liver |
| Msn | 1.880318 | 9.287276 | 733.4297 | 1.49E-36 | 3.82E-35 | Infection | Liver |
| Msr1 | 0.672884 | 6.218893 | 48.18857 | 2.46E-09 | 4.84E-09 | Infection | Liver |
| Msra | -2.0576 | 9.417499 | 818.0819 | 6.08E-38 | 1.91E-36 | Healthy control | Liver |
| Msrb1 | -1.88749 | 8.865955 | 643.497 | 6.71E-35 | 1.25E-33 | Healthy control | Liver |
| Mst1 | -0.64702 | 9.499729 | 49.48697 | 1.69E-09 | 3.33E-09 | Healthy control | Liver |
| Mst1r | 1.532974 | 2.021228 | 23.33662 | 3.97E-06 | 6.59E-06 | Infection | Liver |

| | | | | | | | |
|---|---|---|---|---|---|---|---|
| Hk2 | 2.668576 | 5.952175 | 346.0941 | 2.67E-27 | 1.94E-26 | Infection | Liver |
| Ctnnal1 | 2.664012 | 6.052992 | 421.4025 | 1.25E-29 | 1.17E-28 | Infection | Liver |
| Msx2 | 1.668818 | 1.482277 | 11.10768 | 1.25E-03 | 1.78E-03 | Infection | Liver |
| Mta1 | -0.71816 | 7.271541 | 133.0436 | 3.27E-17 | 9.92E-17 | Healthy control | Liver |
| Mta2 | 0.549295 | 8.643137 | 196.1795 | 4.51E-21 | 1.80E-20 | Infection | Liver |
| Mta3 | 0.198112 | 7.176812 | 17.28525 | 9.84E-05 | 1.51E-04 | Infection | Liver |
| Mtch1 | -0.24246 | 8.76294 | 27.25614 | 2.10E-06 | 3.53E-06 | Healthy control | Liver |
| Mtch2 | -0.59111 | 8.60683 | 72.0978 | 4.73E-12 | 1.06E-11 | Healthy control | Liver |
| F3 | 2.660728 | 5.137734 | 163.4531 | 4.71E-19 | 1.62E-18 | Infection | Liver |
| Mtf1 | -1.41775 | 6.001891 | 299.2162 | 1.20E-25 | 7.29E-25 | Healthy control | Liver |
| Mtf2 | 0.215053 | 5.691376 | 9.850881 | 2.58E-03 | 3.57E-03 | Infection | Liver |
| Mtfp1 | -2.44024 | 7.268104 | 960.7167 | 5.70E-40 | 2.64E-38 | Healthy control | Liver |
| Mthfr | 0.420875 | 5.194753 | 32.13448 | 3.83E-07 | 6.69E-07 | Infection | Liver |
| Mtif2 | -0.40189 | 7.385674 | 40.97688 | 2.14E-08 | 3.98E-08 | Healthy control | Liver |
| Mtmr12 | 0.378439 | 6.626467 | 29.69325 | 8.84E-07 | 1.52E-06 | Infection | Liver |
| Mtnr1a | 0.211845 | 1.729888 | 0.277843 | 5.99E-01 | 6.25E-01 | Infection | Liver |
| Mtnr1b | 0.635825 | 1.206914 | 1.381597 | 2.53E-01 | 2.78E-01 | Infection | Liver |
| Mtor | 0.009337 | 8.26095 | 0.022318 | 8.82E-01 | 8.92E-01 | Infection | Liver |
| Mtrex | -0.36384 | 6.061292 | 36.73176 | 8.25E-08 | 1.49E-07 | Healthy control | Liver |
| Mttp | -1.4053 | 8.637571 | 188.2309 | 1.30E-20 | 5.03E-20 | Healthy control | Liver |
| Mturn | 1.129896 | 5.03638 | 102.7184 | 7.33E-15 | 1.92E-14 | Infection | Liver |
| Mtus1 | -0.42552 | 9.020975 | 50.05959 | 1.44E-09 | 2.84E-09 | Healthy control | Liver |
| Mtx1 | -0.62122 | 6.63522 | 120.8071 | 2.56E-16 | 7.28E-16 | Healthy control | Liver |
| Pth | 2.658345 | 1.445943 | 20.58926 | 2.50E-05 | 3.98E-05 | Infection | Liver |
| Muc2 | 1.498423 | 2.14664 | 27.77252 | 6.42E-07 | 1.11E-06 | Infection | Liver |
| Muc20 | 1.791172 | 1.485477 | 11.27275 | 1.17E-03 | 1.66E-03 | Infection | Liver |
| Muc5ac | 0.339114 | 1.195386 | 0.393502 | 5.39E-01 | 5.65E-01 | Infection | Liver |
| Mus81 | -0.29712 | 5.83347 | 31.37491 | 4.94E-07 | 8.60E-07 | Healthy control | Liver |
| Musk | 0.797974 | 1.231523 | 1.651882 | 2.09E-01 | 2.33E-01 | Infection | Liver |
| Tent5c | 2.657203 | 5.261186 | 263.251 | 4.02E-24 | 2.13E-23 | Infection | Liver |
| Mxd4 | -0.57359 | 6.796382 | 61.27676 | 6.89E-11 | 1.47E-10 | Healthy control | Liver |
| Tmem173 | 2.656486 | 6.191715 | 624.9503 | 1.83E-34 | 3.10E-33 | Infection | Liver |
| Mybbp1a | -0.02438 | 7.651336 | 0.32638 | 5.70E-01 | 5.96E-01 | Healthy control | Liver |
| Gli1 | 2.65627 | 1.916615 | 48.34631 | 4.61E-10 | 9.39E-10 | Infection | Liver |
| Fcmr | 2.648746 | 4.114081 | 106.9163 | 1.57E-15 | 4.25E-15 | Infection | Liver |
| Mybph | 1.942785 | 1.53684 | 14.66059 | 2.50E-04 | 3.72E-04 | Infection | Liver |
| Atp1b2 | 2.644139 | 3.733496 | 118.3537 | 5.66E-17 | 1.69E-16 | Infection | Liver |
| Mycbp2 | 0.614923 | 6.048193 | 64.77895 | 2.84E-11 | 6.13E-11 | Infection | Liver |
| Mycl | -0.22413 | 4.834851 | 2.44583 | 1.23E-01 | 1.41E-01 | Healthy control | Liver |
| Rtn2 | 2.643979 | 2.884005 | 118.2424 | 2.03E-18 | 6.68E-18 | Infection | Liver |
| Myd88 | 0.929529 | 7.038891 | 250.3792 | 1.08E-23 | 5.45E-23 | Infection | Liver |
| Defa5 | 2.642068 | 1.750416 | 25.78998 | 2.17E-06 | 3.65E-06 | Infection | Liver |
| Cntn2 | 2.641057 | 1.435025 | 17.6329 | 8.12E-05 | 1.25E-04 | Infection | Liver |
| Myh1 | 0.947099 | 1.234646 | 2.234625 | 1.46E-01 | 1.66E-01 | Infection | Liver |
| Myh10 | 1.653503 | 5.722641 | 118.6054 | 4.01E-16 | 1.13E-15 | Infection | Liver |
| Trim63 | 2.636543 | 1.656453 | 30.64617 | 4.52E-07 | 7.87E-07 | Infection | Liver |
| Myh2 | 0.922971 | 1.23213 | 2.357297 | 1.35E-01 | 1.54E-01 | Infection | Liver |
| Myh3 | 0.781699 | 1.262124 | 1.773252 | 1.90E-01 | 2.13E-01 | Infection | Liver |
| Myh4 | 1.727404 | 1.38587 | 10.69568 | 1.72E-03 | 2.41E-03 | Infection | Liver |
| Myh7 | 1.543639 | 1.235372 | 6.072216 | 2.02E-02 | 2.58E-02 | Infection | Liver |

| | | | | | | | |
|---|---|---|---|---|---|---|---|
| Myh8 | 0.806301 | 1.429274 | 2.383992 | 1.26E-01 | 1.45E-01 | Infection | Liver |
| Myh9 | 0.245524 | 9.043574 | 22.73869 | 1.13E-05 | 1.83E-05 | Infection | Liver |
| Myl2 | 1.059324 | 1.354849 | 3.007862 | 8.74E-02 | 1.02E-01 | Infection | Liver |
| Sprr2f | 2.635301 | 1.439673 | 14.43971 | 3.14E-04 | 4.65E-04 | Infection | Liver |
| Myo10 | 1.619574 | 7.953495 | 551.2823 | 5.68E-33 | 7.79E-32 | Infection | Liver |
| Myo16 | 1.945706 | 1.490479 | 12.85737 | 5.90E-04 | 8.56E-04 | Infection | Liver |
| Myo18a | -0.64898 | 8.619588 | 170.2408 | 1.29E-19 | 4.69E-19 | Healthy control | Liver |
| Myo1c | -0.04588 | 8.241386 | 0.464035 | 4.98E-01 | 5.26E-01 | Healthy control | Liver |
| Myo1d | 0.399094 | 6.910835 | 25.4669 | 4.04E-06 | 6.71E-06 | Infection | Liver |
| Myo1e | -0.10991 | 8.83764 | 5.596583 | 2.11E-02 | 2.68E-02 | Healthy control | Liver |
| Speg | 2.634988 | 4.401997 | 372.2709 | 2.67E-28 | 2.18E-27 | Infection | Liver |
| Myo6 | -0.8826 | 7.721041 | 324.3874 | 1.29E-26 | 8.67E-26 | Healthy control | Liver |
| Myo7a | 0.383402 | 5.866798 | 9.334806 | 3.29E-03 | 4.50E-03 | Infection | Liver |
| Myo9a | 1.422456 | 6.30877 | 276.4628 | 8.68E-25 | 4.90E-24 | Infection | Liver |
| Myocd | 0.905329 | 1.286751 | 2.281031 | 1.38E-01 | 1.57E-01 | Infection | Liver |
| Myod1 | 0.590501 | 1.269197 | 0.829907 | 3.67E-01 | 3.94E-01 | Infection | Liver |
| Myog | 1.738849 | 1.315285 | 6.685414 | 1.25E-02 | 1.63E-02 | Infection | Liver |
| Myrf | -0.70838 | 6.270433 | 62.83543 | 4.64E-11 | 9.93E-11 | Healthy control | Liver |
| Myt1 | 1.945302 | 1.336556 | 9.699279 | 2.87E-03 | 3.96E-03 | Infection | Liver |
| Msi1 | 2.634087 | 3.510415 | 108.4053 | 3.05E-16 | 8.60E-16 | Infection | Liver |
| Naa15 | -0.20541 | 7.364932 | 13.25409 | 5.49E-04 | 7.97E-04 | Healthy control | Liver |
| Grk3 | 2.63322 | 5.918363 | 658.7826 | 4.29E-35 | 8.53E-34 | Infection | Liver |
| Nadk2 | -1.71293 | 8.476496 | 252.6539 | 1.07E-23 | 5.40E-23 | Healthy control | Liver |
| Nae1 | -0.22181 | 7.391003 | 20.46696 | 2.73E-05 | 4.32E-05 | Healthy control | Liver |
| Nagk | -0.4678 | 6.467293 | 44.09452 | 8.25E-09 | 1.57E-08 | Healthy control | Liver |
| Naip1 | 1.963454 | 1.980263 | 21.65427 | 8.66E-06 | 1.41E-05 | Infection | Liver |
| Naip5 | 0.494236 | 3.362043 | 9.876831 | 2.53E-03 | 3.50E-03 | Infection | Liver |
| Nampt | 1.878781 | 8.391876 | 712.2361 | 3.53E-36 | 8.41E-35 | Infection | Liver |
| Nppb | 2.631184 | 1.353445 | 20.07135 | 3.91E-05 | 6.15E-05 | Infection | Liver |
| Nat1 | -1.83074 | 2.849015 | 59.19645 | 1.78E-11 | 3.90E-11 | Healthy control | Liver |
| Nbn | -1.14123 | 6.313856 | 266.7156 | 2.24E-24 | 1.22E-23 | Healthy control | Liver |
| Nbr1 | -1.75055 | 9.490524 | 1448.309 | 2.25E-45 | 4.90E-43 | Healthy control | Liver |
| Saa1 | 2.629345 | 9.945464 | 79.46028 | 1.79E-12 | 4.11E-12 | Infection | Liver |
| Ncam2 | 1.72893 | 2.098002 | 29.25807 | 3.10E-07 | 5.45E-07 | Infection | Liver |
| Ncan | 0.461222 | 1.287858 | 0.600985 | 4.42E-01 | 4.70E-01 | Infection | Liver |
| Wnt6 | 2.629114 | 1.916001 | 41.72088 | 4.85E-09 | 9.39E-09 | Infection | Liver |
| Ncbp2 | -0.61175 | 6.315067 | 69.64593 | 8.57E-12 | 1.90E-11 | Healthy control | Liver |
| Ncr1 | 2.628616 | 3.522866 | 98.11314 | 2.34E-15 | 6.28E-15 | Infection | Liver |
| Ttll3 | 2.627395 | 4.033926 | 275.1088 | 2.44E-25 | 1.43E-24 | Infection | Liver |
| Nck1 | -0.60259 | 6.008341 | 51.98769 | 8.41E-10 | 1.69E-09 | Healthy control | Liver |
| Ncl | -1.10694 | 7.331234 | 220.5943 | 2.59E-22 | 1.15E-21 | Healthy control | Liver |
| Ncoa1 | -0.59785 | 6.301698 | 71.20467 | 5.89E-12 | 1.32E-11 | Healthy control | Liver |
| Ncoa2 | -1.21746 | 7.903452 | 875.0448 | 8.39E-39 | 3.04E-37 | Healthy control | Liver |
| Ncoa3 | 0.280945 | 7.36063 | 18.60695 | 5.73E-05 | 8.90E-05 | Infection | Liver |
| Ncor1 | -1.05979 | 8.752125 | 743.2063 | 1.02E-36 | 2.71E-35 | Healthy control | Liver |
| Ncor2 | -0.62461 | 7.583118 | 78.33169 | 1.12E-12 | 2.60E-12 | Healthy control | Liver |
| Prl3c1 | 2.625077 | 1.498814 | 20.1912 | 2.75E-05 | 4.35E-05 | Infection | Liver |
| Ncstn | 1.189899 | 7.801429 | 468.1706 | 5.72E-31 | 6.28E-30 | Infection | Liver |
| Ndc1 | 0.405854 | 6.051192 | 32.21637 | 3.70E-07 | 6.47E-07 | Infection | Liver |
| Gtse1 | 2.623943 | 3.697974 | 228.7931 | 5.73E-24 | 2.98E-23 | Infection | Liver |

| | | | | | | | |
|---|---|---|---|---|---|---|---|
| Nde1 | 0.057059 | 6.651337 | 1.010322 | 3.19E-01 | 3.45E-01 | Infection | Liver |
| Ndel1 | 1.040615 | 7.357106 | 293.653 | 1.78E-25 | 1.06E-24 | Infection | Liver |
| Ndn | -0.06443 | 4.82967 | 0.166102 | 6.85E-01 | 7.07E-01 | Healthy control | Liver |
| Ndor1 | 0.68838 | 6.23821 | 122.2004 | 2.01E-16 | 5.77E-16 | Infection | Liver |
| Ndrg2 | -2.21008 | 11.69396 | 487.0249 | 1.88E-31 | 2.20E-30 | Healthy control | Liver |
| Gfi1b | 2.622863 | 1.46265 | 17.30833 | 1.08E-04 | 1.65E-04 | Infection | Liver |
| Ndst1 | -1.82289 | 8.389932 | 676.1401 | 1.61E-35 | 3.38E-34 | Healthy control | Liver |
| Ndst2 | -0.30763 | 7.475938 | 42.08586 | 1.52E-08 | 2.85E-08 | Healthy control | Liver |
| Ndst3 | 0.839573 | 1.383897 | 2.670314 | 1.06E-01 | 1.23E-01 | Infection | Liver |
| Ndufa12 | -0.83586 | 3.773478 | 28.13439 | 1.46E-06 | 2.48E-06 | Healthy control | Liver |
| Ndufa9 | -1.50579 | 8.423258 | 578.5529 | 1.43E-33 | 2.11E-32 | Healthy control | Liver |
| Ndufaf6 | -1.09879 | 7.588919 | 184.2615 | 2.02E-20 | 7.77E-20 | Healthy control | Liver |
| Ndufs1 | -1.38763 | 9.245664 | 624.9644 | 1.55E-34 | 2.67E-33 | Healthy control | Liver |
| Ndufs4 | -0.55861 | 7.326924 | 98.87238 | 1.47E-14 | 3.76E-14 | Healthy control | Liver |
| Ndufv2 | -1.79529 | 7.21873 | 539.8763 | 1.06E-32 | 1.40E-31 | Healthy control | Liver |
| Neat1 | 1.547187 | 12.02647 | 197.4657 | 3.85E-21 | 1.55E-20 | Infection | Liver |
| Nectin2 | -1.02222 | 6.731963 | 169.7825 | 1.39E-19 | 5.02E-19 | Healthy control | Liver |
| Nectin3 | -0.74611 | 8.080947 | 106.0465 | 3.67E-15 | 9.75E-15 | Healthy control | Liver |
| Nedd4 | -0.64611 | 10.01498 | 76.47543 | 1.71E-12 | 3.92E-12 | Healthy control | Liver |
| Nedd4l | -0.5178 | 8.438104 | 30.40182 | 6.89E-07 | 1.19E-06 | Healthy control | Liver |
| Nefh | 0.333773 | 1.870464 | 1.102312 | 2.96E-01 | 3.22E-01 | Infection | Liver |
| Ffar4 | 2.621475 | 1.883385 | 47.48459 | 6.10E-10 | 1.23E-09 | Infection | Liver |
| Nefm | 1.241904 | 1.435132 | 5.678127 | 1.94E-02 | 2.47E-02 | Infection | Liver |
| Neil1 | -0.19378 | 6.42294 | 9.500278 | 3.04E-03 | 4.18E-03 | Healthy control | Liver |
| Nek2 | 0.659318 | 3.401596 | 11.3704 | 1.27E-03 | 1.80E-03 | Infection | Liver |
| Nek7 | -0.14143 | 7.603909 | 5.953991 | 1.75E-02 | 2.24E-02 | Healthy control | Liver |
| Nelfb | -0.58401 | 7.811874 | 176.2985 | 5.72E-20 | 2.13E-19 | Healthy control | Liver |
| Nelfe | -0.10427 | 7.274703 | 3.29594 | 7.42E-02 | 8.76E-02 | Healthy control | Liver |
| Nell1 | 0.795872 | 1.218434 | 2.023255 | 1.68E-01 | 1.89E-01 | Infection | Liver |
| Neo1 | 1.766161 | 6.753391 | 536.2036 | 1.28E-32 | 1.68E-31 | Infection | Liver |
| Dazl | 2.61819 | 1.885197 | 21.40295 | 1.18E-05 | 1.91E-05 | Infection | Liver |
| Neu1 | -0.6532 | 8.00606 | 135.8951 | 2.06E-17 | 6.31E-17 | Healthy control | Liver |
| Ccnb1 | 2.617984 | 3.154803 | 87.15434 | 9.77E-15 | 2.54E-14 | Infection | Liver |
| Neurod2 | 0.171048 | 1.280519 | 0.073545 | 7.87E-01 | 8.05E-01 | Infection | Liver |
| Neurod4 | 1.128055 | 1.348121 | 2.705068 | 1.05E-01 | 1.22E-01 | Infection | Liver |
| Neurod6 | 0.565422 | 1.245575 | 0.761585 | 3.89E-01 | 4.16E-01 | Infection | Liver |
| Neurog1 | 1.000466 | 1.206157 | 4.434044 | 4.64E-02 | 5.64E-02 | Infection | Liver |
| Neurog2 | 1.976241 | 1.285284 | 13.70457 | 5.49E-04 | 7.97E-04 | Infection | Liver |
| Neurog3 | 0.36377 | 1.256315 | 0.416763 | 5.22E-01 | 5.50E-01 | Infection | Liver |
| Nf1 | 0.538022 | 7.649713 | 169.0991 | 1.51E-19 | 5.44E-19 | Infection | Liver |
| Nf2 | -0.65931 | 6.476664 | 137.0411 | 1.73E-17 | 5.34E-17 | Healthy control | Liver |
| Nfat5 | 0.365865 | 7.30152 | 45.76474 | 5.00E-09 | 9.66E-09 | Infection | Liver |
| Nfatc1 | 0.803537 | 6.994684 | 150.0945 | 2.30E-18 | 7.50E-18 | Infection | Liver |
| Bard1 | 2.616191 | 3.3814 | 175.8482 | 1.04E-21 | 4.39E-21 | Infection | Liver |
| Nfatc3 | -0.72434 | 6.84342 | 155.6869 | 1.01E-18 | 3.40E-18 | Healthy control | Liver |
| Bex3 | 2.61569 | 5.689394 | 485.9412 | 2.63E-31 | 3.00E-30 | Infection | Liver |
| Nfe2l1 | -0.11681 | 9.27056 | 9.71755 | 2.74E-03 | 3.79E-03 | Healthy control | Liver |
| Nfe2l2 | -0.40875 | 8.605422 | 111.6737 | 1.29E-15 | 3.52E-15 | Healthy control | Liver |
| Nfia | -1.73093 | 7.583992 | 1248.594 | 2.07E-43 | 2.00E-41 | Healthy control | Liver |
| Nfib | -0.15957 | 8.571526 | 10.21095 | 2.18E-03 | 3.03E-03 | Healthy control | Liver |

| | | | | | | | |
|---|---|---|---|---|---|---|---|
| Nfil3 | -0.3093 | 7.803068 | 31.53124 | 4.66E-07 | 8.12E-07 | Healthy control | Liver |
| Nfix | -0.23983 | 8.738052 | 24.96739 | 4.86E-06 | 8.05E-06 | Healthy control | Liver |
| Nfkb1 | 0.228316 | 7.637654 | 16.2991 | 1.48E-04 | 2.24E-04 | Infection | Liver |
| Emilin1 | 2.612061 | 9.111963 | 376.5025 | 7.68E-28 | 6.03E-27 | Infection | Liver |
| Nfkbib | -0.67928 | 7.54181 | 157.0764 | 8.22E-19 | 2.78E-18 | Healthy control | Liver |
| Nfkbie | 1.753144 | 6.076477 | 464.664 | 7.44E-31 | 8.04E-30 | Infection | Liver |
| Nfu1 | -2.54887 | 4.354176 | 266.3931 | 1.16E-25 | 7.04E-25 | Healthy control | Liver |
| Nfya | 0.538942 | 5.253209 | 23.82415 | 7.50E-06 | 1.23E-05 | Infection | Liver |
| Nfyc | -0.66173 | 6.244434 | 109.3255 | 2.01E-15 | 5.43E-15 | Healthy control | Liver |
| Ngb | 1.135979 | 2.652943 | 22.02226 | 1.07E-05 | 1.74E-05 | Infection | Liver |
| Ngf | 0.326903 | 5.225909 | 5.770614 | 1.92E-02 | 2.46E-02 | Infection | Liver |
| Ngfr | -0.28464 | 5.395335 | 5.588624 | 2.12E-02 | 2.69E-02 | Healthy control | Liver |
| Nhlh1 | 1.185121 | 1.290794 | 3.627923 | 6.26E-02 | 7.47E-02 | Infection | Liver |
| Nhlh2 | 1.870138 | 1.566868 | 11.75627 | 9.18E-04 | 1.31E-03 | Infection | Liver |
| Nhsl1 | 0.820902 | 5.180488 | 64.31237 | 3.25E-11 | 6.99E-11 | Infection | Liver |
| Nid1 | 1.1954 | 7.583985 | 83.33177 | 4.64E-13 | 1.11E-12 | Infection | Liver |
| Nid2 | 1.101592 | 6.317137 | 197.26 | 4.00E-21 | 1.61E-20 | Infection | Liver |
| Ninl | 1.82881 | 4.032208 | 132.0524 | 4.39E-17 | 1.32E-16 | Infection | Liver |
| Nipbl | -0.28682 | 6.916678 | 31.79697 | 4.26E-07 | 7.43E-07 | Healthy control | Liver |
| Nipsnap2 | 0.036172 | 3.973602 | 0.072907 | 7.88E-01 | 8.05E-01 | Infection | Liver |
| Htr7 | 2.610718 | 2.697983 | 93.3278 | 2.72E-16 | 7.71E-16 | Infection | Liver |
| Grin2d | 2.607735 | 3.400522 | 148.2101 | 1.01E-19 | 3.71E-19 | Infection | Liver |
| Lipm | 2.607192 | 1.497802 | 16.87474 | 1.07E-04 | 1.64E-04 | Infection | Liver |
| Nkx2-1 | 0.713754 | 1.256225 | 1.196486 | 2.81E-01 | 3.07E-01 | Infection | Liver |
| Nkx2-2 | 1.64122 | 1.33756 | 6.206157 | 1.56E-02 | 2.01E-02 | Infection | Liver |
| Nkx2-5 | 1.23183 | 1.255992 | 4.118837 | 4.95E-02 | 6.00E-02 | Infection | Liver |
| Nkx3-1 | 1.710734 | 1.246254 | 7.423631 | 1.08E-02 | 1.41E-02 | Infection | Liver |
| Nkx3-2 | 0.157663 | 1.216798 | 0.082765 | 7.76E-01 | 7.94E-01 | Infection | Liver |
| Haspin | 2.605454 | 3.878481 | 374.4312 | 2.50E-29 | 2.28E-28 | Infection | Liver |
| Nkx6-2 | 0.236499 | 1.218666 | 0.170496 | 6.83E-01 | 7.06E-01 | Infection | Liver |
| Cxcl10 | 2.605418 | 5.592335 | 68.28678 | 2.28E-11 | 4.95E-11 | Infection | Liver |
| Nlgn3 | 1.832626 | 2.912727 | 103.6948 | 4.33E-16 | 1.21E-15 | Infection | Liver |
| Nlk | -0.01733 | 5.750459 | 0.076658 | 7.83E-01 | 8.01E-01 | Healthy control | Liver |
| Nlrc4 | 1.368585 | 3.419617 | 53.47759 | 4.92E-10 | 1.00E-09 | Infection | Liver |
| Kif5c | 2.604703 | 1.977424 | 61.20767 | 5.00E-12 | 1.12E-11 | Infection | Liver |
| Nlrp3 | 0.993798 | 4.175258 | 57.11424 | 2.23E-10 | 4.63E-10 | Infection | Liver |
| Nlrp4e | 1.356729 | 1.270533 | 4.03506 | 5.17E-02 | 6.23E-02 | Infection | Liver |
| Nlrp4f | 1.659514 | 1.433912 | 7.025988 | 9.75E-03 | 1.28E-02 | Infection | Liver |
| Nlrp6 | -1.69276 | 9.149945 | 433.2156 | 4.95E-30 | 4.86E-29 | Healthy control | Liver |
| Nlrp9b | 1.172771 | 1.334454 | 3.332728 | 7.32E-02 | 8.66E-02 | Infection | Liver |
| Nmb | 0.592436 | 3.422503 | 12.96139 | 6.22E-04 | 9.00E-04 | Infection | Liver |
| Nme5 | 0.475338 | 2.133907 | 2.397619 | 1.24E-01 | 1.43E-01 | Infection | Liver |
| Nmrk1 | -0.10315 | 5.231499 | 1.428675 | 2.36E-01 | 2.62E-01 | Healthy control | Liver |
| Nmt1 | -0.34357 | 7.436894 | 56.23896 | 2.60E-10 | 5.36E-10 | Healthy control | Liver |
| Dkk1 | 2.601065 | 1.514849 | 18.66555 | 5.02E-05 | 7.83E-05 | Infection | Liver |
| Nmur1 | 1.868331 | 1.466613 | 11.52992 | 1.08E-03 | 1.54E-03 | Infection | Liver |
| Nobox | 1.866404 | 1.281101 | 13.59292 | 6.00E-04 | 8.69E-04 | Infection | Liver |
| Noct | 0.537611 | 6.3364 | 5.244339 | 2.56E-02 | 3.22E-02 | Infection | Liver |
| Nod1 | -0.42905 | 6.196575 | 34.87257 | 1.52E-07 | 2.72E-07 | Healthy control | Liver |
| Nod2 | 0.770629 | 1.630298 | 4.046427 | 4.63E-02 | 5.63E-02 | Infection | Liver |

| | | | | | | | |
|---|---|---|---|---|---|---|---|
| Nodal | 0.198577 | 1.692038 | 0.206893 | 6.50E-01 | 6.73E-01 | Infection | Liver |
| Nog | 0.774733 | 1.292621 | 1.630484 | 2.07E-01 | 2.31E-01 | Infection | Liver |
| Nos1ap | 0.680537 | 4.72786 | 30.03519 | 7.97E-07 | 1.37E-06 | Infection | Liver |
| Piezo2 | 2.598093 | 5.397892 | 204.7682 | 2.12E-21 | 8.73E-21 | Infection | Liver |
| Nos3 | -0.45855 | 4.12485 | 11.61559 | 1.14E-03 | 1.63E-03 | Healthy control | Liver |
| Nosip | -0.1896 | 7.790641 | 23.41445 | 8.72E-06 | 1.42E-05 | Healthy control | Liver |
| Notch1 | -0.0062 | 7.503265 | 0.007346 | 9.32E-01 | 9.39E-01 | Healthy control | Liver |
| Notch2 | -0.73187 | 8.104269 | 237.3373 | 4.17E-23 | 1.97E-22 | Healthy control | Liver |
| Tnfsf4 | 2.597566 | 1.720317 | 37.44779 | 2.49E-08 | 4.63E-08 | Infection | Liver |
| Notch4 | 1.504234 | 5.062862 | 150.9904 | 2.22E-18 | 7.27E-18 | Infection | Liver |
| Noto | 0.904642 | 1.309616 | 2.08273 | 1.55E-01 | 1.75E-01 | Infection | Liver |
| Nova2 | -0.04381 | 3.077187 | 0.05354 | 8.18E-01 | 8.33E-01 | Healthy control | Liver |
| Nox1 | 0.678305 | 1.308996 | 1.421058 | 2.38E-01 | 2.63E-01 | Infection | Liver |
| Nox3 | 0.883771 | 1.26986 | 1.94081 | 1.71E-01 | 1.93E-01 | Infection | Liver |
| Nox4 | 0.339678 | 3.297308 | 3.406676 | 6.94E-02 | 8.22E-02 | Infection | Liver |
| Noxo1 | -0.23349 | 3.417182 | 2.034152 | 1.59E-01 | 1.79E-01 | Healthy control | Liver |
| Npas1 | 1.60481 | 1.300276 | 6.626684 | 1.30E-02 | 1.69E-02 | Infection | Liver |
| Npas2 | 0.114694 | 6.040726 | 2.255499 | 1.38E-01 | 1.58E-01 | Infection | Liver |
| Npas4 | 1.747346 | 1.489614 | 9.932184 | 2.25E-03 | 3.13E-03 | Infection | Liver |
| Npbwr1 | 1.528636 | 1.243874 | 8.032132 | 7.49E-03 | 9.96E-03 | Infection | Liver |
| Npc1 | -1.0789 | 8.750954 | 404.01 | 3.39E-29 | 3.03E-28 | Healthy control | Liver |
| Npc1l1 | 1.219245 | 1.266505 | 4.048354 | 5.06E-02 | 6.12E-02 | Infection | Liver |
| Npepps | -0.01283 | 7.432444 | 0.058891 | 8.09E-01 | 8.25E-01 | Healthy control | Liver |
| Npffr2 | 1.425092 | 1.288158 | 5.775598 | 2.03E-02 | 2.58E-02 | Infection | Liver |
| Nphs1 | 1.474765 | 1.323444 | 7.91386 | 6.85E-03 | 9.14E-03 | Infection | Liver |
| Onecut3 | 2.592665 | 1.838518 | 37.632 | 1.93E-08 | 3.61E-08 | Infection | Liver |
| Nploc4 | -0.58927 | 8.466933 | 161.0171 | 4.66E-19 | 1.61E-18 | Healthy control | Liver |
| Nppa | 1.300619 | 1.343593 | 5.541689 | 2.18E-02 | 2.76E-02 | Infection | Liver |
| Pianp | 2.590514 | 6.640369 | 700.471 | 6.17E-36 | 1.37E-34 | Infection | Liver |
| Npr1 | -0.84391 | 6.826254 | 115.3107 | 6.73E-16 | 1.87E-15 | Healthy control | Liver |
| Npsr1 | 1.416555 | 1.509781 | 6.108833 | 1.52E-02 | 1.96E-02 | Infection | Liver |
| Nptx2 | 1.56277 | 1.437885 | 8.773225 | 4.16E-03 | 5.64E-03 | Infection | Liver |
| Nptxr | 0.819738 | 2.186168 | 10.56039 | 1.55E-03 | 2.18E-03 | Infection | Liver |
| Kif2c | 2.588108 | 3.321234 | 147.2413 | 8.91E-20 | 3.28E-19 | Infection | Liver |
| Npy2r | 0.475068 | 1.209076 | 0.691429 | 4.15E-01 | 4.43E-01 | Infection | Liver |
| Npy4r | 1.098188 | 1.417558 | 4.804027 | 3.12E-02 | 3.87E-02 | Infection | Liver |
| Nqo1 | -0.53642 | 6.242945 | 20.58239 | 2.61E-05 | 4.14E-05 | Healthy control | Liver |
| Madcam1 | 2.585528 | 1.567586 | 28.99518 | 8.11E-07 | 1.39E-06 | Infection | Liver |
| Nr0b2 | -1.62395 | 8.943658 | 202.1682 | 3.07E-21 | 1.25E-20 | Healthy control | Liver |
| Nr1d1 | -3.26355 | 8.66445 | 975.33 | 3.45E-40 | 1.69E-38 | Healthy control | Liver |
| Nr1h2 | 0.042487 | 7.701075 | 0.924475 | 3.40E-01 | 3.66E-01 | Infection | Liver |
| Nr1h3 | -1.41656 | 9.538195 | 303.1525 | 7.67E-26 | 4.76E-25 | Healthy control | Liver |
| Nr1h4 | -1.40534 | 8.045295 | 376.6101 | 2.32E-28 | 1.91E-27 | Healthy control | Liver |
| Nr1i2 | -0.93425 | 7.489885 | 100.2686 | 1.11E-14 | 2.89E-14 | Healthy control | Liver |
| Nr1i3 | -3.20041 | 5.517996 | 474.0531 | 4.73E-32 | 5.80E-31 | Healthy control | Liver |
| Nr2c2 | -0.20584 | 6.383653 | 13.78103 | 4.36E-04 | 6.39E-04 | Healthy control | Liver |
| Six6 | 2.584408 | 1.423192 | 15.75748 | 1.79E-04 | 2.70E-04 | Infection | Liver |
| Nr2e3 | 0.795314 | 1.488517 | 2.906652 | 9.10E-02 | 1.07E-01 | Infection | Liver |
| Nr2f1 | 1.164357 | 5.064455 | 79.52727 | 8.91E-13 | 2.09E-12 | Infection | Liver |
| Nr2f2 | 0.135216 | 8.196295 | 6.394953 | 1.39E-02 | 1.81E-02 | Infection | Liver |

| | | | | | | | |
|---|---|---|---|---|---|---|---|
| Nr3c1 | 0.205523 | 7.608202 | 21.29517 | 1.97E-05 | 3.15E-05 | Infection | Liver |
| Nr3c2 | -1.07022 | 6.402249 | 175.4858 | 6.53E-20 | 2.43E-19 | Healthy control | Liver |
| Crxos | 2.580578 | 1.643051 | 21.49882 | 1.38E-05 | 2.23E-05 | Infection | Liver |
| Nr4a2 | 1.396467 | 2.310879 | 24.26497 | 3.14E-06 | 5.23E-06 | Infection | Liver |
| Pkm | 2.572462 | 7.950674 | 267.1983 | 1.07E-23 | 5.39E-23 | Infection | Liver |
| Nr5a1 | 0.576029 | 1.208872 | 1.11663 | 3.02E-01 | 3.28E-01 | Infection | Liver |
| Nr5a2 | -1.55993 | 7.007358 | 267.2321 | 2.08E-24 | 1.13E-23 | Healthy control | Liver |
| Nr6a1 | -1.69319 | 4.679909 | 182.0137 | 2.33E-20 | 8.87E-20 | Healthy control | Liver |
| Nrarp | 0.328715 | 2.769271 | 2.645516 | 1.08E-01 | 1.25E-01 | Infection | Liver |
| Nras | 1.309599 | 6.412344 | 448.9453 | 1.88E-30 | 1.94E-29 | Infection | Liver |
| Kif4 | 2.571306 | 3.381458 | 152.899 | 5.13E-20 | 1.92E-19 | Infection | Liver |
| Nrep | 1.60773 | 6.452815 | 179.9129 | 3.59E-20 | 1.35E-19 | Infection | Liver |
| Nrf1 | 0.28514 | 5.783207 | 18.38079 | 6.29E-05 | 9.74E-05 | Infection | Liver |
| Casq1 | 2.570955 | 1.778537 | 36.9927 | 3.50E-08 | 6.45E-08 | Infection | Liver |
| Nrip1 | -0.33958 | 5.968576 | 23.91162 | 7.24E-06 | 1.19E-05 | Healthy control | Liver |
| Col23a1 | 2.569477 | 2.953303 | 133.5495 | 1.10E-19 | 4.02E-19 | Infection | Liver |
| Nrl | 0.741038 | 1.185727 | 2.59106 | 1.29E-01 | 1.48E-01 | Infection | Liver |
| Nrn1 | -2.19378 | 10.86489 | 998.3218 | 1.66E-40 | 8.91E-39 | Healthy control | Liver |
| Nrp1 | 0.140755 | 8.824441 | 3.661292 | 6.02E-02 | 7.20E-02 | Infection | Liver |
| Nrp2 | -0.87202 | 6.194666 | 66.49946 | 1.86E-11 | 4.05E-11 | Healthy control | Liver |
| Nsf | -0.08377 | 6.678864 | 2.384864 | 1.27E-01 | 1.46E-01 | Healthy control | Liver |
| Nsmce3 | 0.62651 | 2.766373 | 9.134655 | 3.44E-03 | 4.71E-03 | Infection | Liver |
| Nsmf | -1.50365 | 9.529334 | 702.8065 | 5.18E-36 | 1.16E-34 | Healthy control | Liver |
| Nsun2 | -0.35995 | 7.37561 | 41.61738 | 1.75E-08 | 3.28E-08 | Healthy control | Liver |
| Nt5c1b | 1.328795 | 1.429773 | 5.13406 | 2.62E-02 | 3.28E-02 | Infection | Liver |
| Nt5e | -1.68217 | 8.713468 | 704.2648 | 4.90E-36 | 1.10E-34 | Healthy control | Liver |
| Ntf3 | 1.278612 | 3.313094 | 41.41383 | 1.65E-08 | 3.10E-08 | Infection | Liver |
| Ccnb3 | 2.567155 | 1.356655 | 16.06962 | 1.75E-04 | 2.64E-04 | Infection | Liver |
| Nthl1 | -1.16974 | 4.541308 | 112.4855 | 1.11E-15 | 3.04E-15 | Healthy control | Liver |
| Ntmt1 | 0.446962 | 5.112257 | 34.2632 | 1.87E-07 | 3.34E-07 | Infection | Liver |
| Ntn1 | -0.3981 | 7.016954 | 22.20127 | 1.39E-05 | 2.23E-05 | Healthy control | Liver |
| Ntn4 | -1.21789 | 4.656696 | 85.10408 | 2.51E-13 | 6.07E-13 | Healthy control | Liver |
| Cpne4 | 2.564083 | 1.736847 | 29.32806 | 5.45E-07 | 9.46E-07 | Infection | Liver |
| Ntng2 | 0.748588 | 6.153008 | 63.33838 | 4.08E-11 | 8.75E-11 | Infection | Liver |
| Rho | 2.560391 | 1.572069 | 24.05094 | 5.38E-06 | 8.87E-06 | Infection | Liver |
| Ntrk2 | -1.03703 | 6.156969 | 21.60738 | 1.94E-05 | 3.10E-05 | Healthy control | Liver |
| Dkk3 | 2.559685 | 6.226704 | 129.2871 | 1.79E-16 | 5.13E-16 | Infection | Liver |
| Mpl | 2.556973 | 1.961598 | 26.88354 | 1.12E-06 | 1.91E-06 | Infection | Liver |
| Ntsr1 | 1.231625 | 1.255383 | 4.435693 | 4.19E-02 | 5.13E-02 | Infection | Liver |
| Ntsr2 | 1.537886 | 1.331391 | 6.242941 | 1.54E-02 | 1.98E-02 | Infection | Liver |
| Nuak2 | -0.32562 | 6.28963 | 14.53028 | 3.15E-04 | 4.67E-04 | Healthy control | Liver |
| Nubpl | -0.97364 | 2.520035 | 18.41984 | 4.57E-05 | 7.14E-05 | Healthy control | Liver |
| Numb | -0.55464 | 7.479023 | 114.2438 | 8.11E-16 | 2.24E-15 | Healthy control | Liver |
| Nup155 | -0.41232 | 5.797252 | 32.87279 | 2.97E-07 | 5.23E-07 | Healthy control | Liver |
| Nup205 | 0.003579 | 5.60944 | 0.001927 | 9.65E-01 | 9.68E-01 | Infection | Liver |
| Nup62 | -0.98756 | 4.584134 | 45.03502 | 6.35E-09 | 1.22E-08 | Healthy control | Liver |
| Nup98 | 0.455657 | 7.154723 | 53.61945 | 5.30E-10 | 1.08E-09 | Infection | Liver |
| Map3k14 | 2.554767 | 6.15925 | 408.947 | 2.72E-29 | 2.47E-28 | Infection | Liver |
| Adgra2 | 2.551322 | 7.166449 | 819.8685 | 5.90E-38 | 1.88E-36 | Infection | Liver |
| Gdf1 | 2.549899 | 1.7875 | 39.67446 | 9.56E-09 | 1.82E-08 | Infection | Liver |

| | | | | | | | |
|---|---|---|---|---|---|---|---|
| Phex | 2.549255 | 1.711736 | 36.41183 | 3.79E-08 | 6.96E-08 | Infection | Liver |
| Nyap2 | 0.821987 | 1.287961 | 2.122699 | 1.51E-01 | 1.71E-01 | Infection | Liver |
| Oasl2 | 1.299643 | 7.652619 | 238.0108 | 3.89E-23 | 1.84E-22 | Infection | Liver |
| Oc90 | 0.574456 | 1.241072 | 0.909268 | 3.47E-01 | 3.73E-01 | Infection | Liver |
| Oca2 | -0.03278 | 1.230916 | 0.003371 | 9.54E-01 | 9.58E-01 | Healthy control | Liver |
| Ocln | -0.42256 | 6.530912 | 9.858284 | 2.58E-03 | 3.57E-03 | Healthy control | Liver |
| Ocstamp | -1.12402 | 1.491601 | 5.272355 | 2.35E-02 | 2.98E-02 | Healthy control | Liver |
| Gpr65 | 2.543555 | 5.12231 | 547.1509 | 1.11E-32 | 1.47E-31 | Infection | Liver |
| Odc1 | 0.036837 | 8.069305 | 0.477815 | 4.92E-01 | 5.19E-01 | Infection | Liver |
| Odf2 | 0.169336 | 5.876126 | 4.432381 | 3.92E-02 | 4.82E-02 | Infection | Liver |
| Odr4 | -0.64346 | 6.919949 | 168.5925 | 1.63E-19 | 5.85E-19 | Healthy control | Liver |
| Ofd1 | 1.337021 | 4.807136 | 228.4678 | 1.32E-22 | 6.04E-22 | Infection | Liver |
| Ogdh | -1.33549 | 8.079857 | 495.4488 | 1.17E-31 | 1.40E-30 | Healthy control | Liver |
| Ogg1 | 0.027 | 4.833426 | 0.105193 | 7.47E-01 | 7.65E-01 | Infection | Liver |
| Cd84 | 2.543162 | 7.473239 | 1023.086 | 8.18E-41 | 4.93E-39 | Infection | Liver |
| Ogt | 0.468121 | 7.412489 | 56.05049 | 2.73E-10 | 5.63E-10 | Infection | Liver |
| Olfm3 | -0.90292 | 2.816188 | 9.437523 | 2.91E-03 | 4.01E-03 | Healthy control | Liver |
| Olfr17 | 1.382425 | 1.233606 | 5.924029 | 2.10E-02 | 2.67E-02 | Infection | Liver |
| Olig1 | -1.94688 | 1.873617 | 27.4927 | 9.09E-07 | 1.56E-06 | Healthy control | Liver |
| Olig2 | -0.12476 | 1.289309 | 0.044731 | 8.33E-01 | 8.47E-01 | Healthy control | Liver |
| Olig3 | 0.27404 | 1.197406 | 0.262951 | 6.15E-01 | 6.39E-01 | Infection | Liver |
| Hbegf | 2.54122 | 4.511752 | 161.1829 | 4.90E-19 | 1.69E-18 | Infection | Liver |
| Omp | 1.864909 | 3.148158 | 105.572 | 7.23E-16 | 2.00E-15 | Infection | Liver |
| Onecut1 | -1.4552 | 6.561338 | 85.05587 | 3.23E-13 | 7.77E-13 | Healthy control | Liver |
| Onecut2 | -1.66478 | 7.955759 | 603.9137 | 4.21E-34 | 6.77E-33 | Healthy control | Liver |
| Ncf4 | 2.537714 | 5.067753 | 291.2996 | 3.11E-25 | 1.82E-24 | Infection | Liver |
| Opa1 | -0.48476 | 8.301201 | 65.82897 | 2.17E-11 | 4.71E-11 | Healthy control | Liver |
| Vdr | 2.533665 | 1.98956 | 45.44457 | 8.77E-10 | 1.76E-09 | Infection | Liver |
| Opn1mw | 1.463594 | 1.279711 | 5.546905 | 2.32E-02 | 2.93E-02 | Infection | Liver |
| Opn1sw | 0.35047 | 1.663988 | 0.783827 | 3.78E-01 | 4.05E-01 | Infection | Liver |
| Opn3 | 1.212706 | 2.369938 | 26.14794 | 1.55E-06 | 2.63E-06 | Infection | Liver |
| Opn4 | 1.535485 | 1.406489 | 7.004354 | 9.97E-03 | 1.31E-02 | Infection | Liver |
| Oprd1 | 1.380169 | 1.65828 | 11.00546 | 1.20E-03 | 1.70E-03 | Infection | Liver |
| Oprk1 | 1.239098 | 1.293492 | 3.941403 | 5.29E-02 | 6.36E-02 | Infection | Liver |
| Oprl1 | 1.493605 | 1.434172 | 6.457346 | 1.32E-02 | 1.72E-02 | Infection | Liver |
| Oprm1 | 0.184128 | 1.310497 | 0.113937 | 7.37E-01 | 0.756326 | Infection | Liver |
| Orai1 | -0.79206 | 4.085985 | 53.63925 | 5.21E-10 | 1.06E-09 | Healthy control | Liver |
| Adra2a | 2.533137 | 1.332136 | 14.69873 | 3.73E-04 | 0.00055 | Infection | Liver |
| Orm1 | 0.097802 | 11.71 | 0.41466 | 5.22E-01 | 0.549508 | Infection | Liver |
| Osgep | -0.30961 | 6.640918 | 26.3326 | 2.94E-06 | 4.91E-06 | Healthy control | Liver |
| Ager | 2.532142 | 2.682768 | 129.9215 | 7.43E-20 | 2.75E-19 | Infection | Liver |
| Dpysl3 | 2.527977 | 6.726404 | 132.4521 | 1.15E-16 | 3.35E-16 | Infection | Liver |
| Mitf | 2.526208 | 4.88477 | 377.1665 | 2.89E-28 | 2.36E-27 | Infection | Liver |
| Osr2 | 0.789164 | 1.354357 | 1.903142 | 1.73E-01 | 0.193917 | Infection | Liver |
| Otc | -2.48893 | 10.80153 | 407.7561 | 4.90E-29 | 4.31E-28 | Healthy control | Liver |
| Otof | 0.263767 | 1.544635 | 0.309393 | 5.79E-01 | 0.604862 | Infection | Liver |
| Otp | 0.205262 | 1.27574 | 0.127638 | 7.22E-01 | 0.742473 | Infection | Liver |
| Otud5 | -0.06335 | 7.940298 | 3.026144 | 8.68E-02 | 0.101794 | Healthy control | Liver |
| Otud7b | -0.54704 | 6.25727 | 53.55286 | 5.42E-10 | 1.10E-09 | Healthy control | Liver |
| Otx1 | -0.3674 | 1.173593 | 1.247903 | 2.88E-01 | 0.314495 | Healthy control | Liver |

| | | | | | | | |
|---|---|---|---|---|---|---|---|
| Otx2 | 1.996691 | 1.292316 | 10.0153 | 2.98E-03 | 0.004101 | Infection | Liver |
| Ovol1 | 0.826196 | 1.334351 | 2.00594 | 1.62E-01 | 0.182343 | Infection | Liver |
| Oxt | 0.724036 | 1.251988 | 1.470106 | 2.33E-01 | 0.257544 | Infection | Liver |
| Oxtr | 1.059858 | 1.317551 | 2.707475 | 1.05E-01 | 0.122137 | Infection | Liver |
| P2rx3 | -0.28602 | 2.805994 | 1.666535 | 2.01E-01 | 0.223622 | Healthy control | Liver |
| P2rx4 | -0.17615 | 6.574507 | 9.943142 | 2.47E-03 | 0.003417 | Healthy control | Liver |
| Kif21b | 2.525272 | 6.128205 | 634.1333 | 1.18E-34 | 2.09E-33 | Infection | Liver |
| P2ry1 | -0.28289 | 4.269561 | 4.253671 | 4.33E-02 | 0.052839 | Healthy control | Liver |
| P2ry13 | 0.273049 | 5.061591 | 3.495871 | 6.62E-02 | 0.078617 | Infection | Liver |
| P2ry2 | 1.262044 | 4.528573 | 99.89572 | 1.36E-14 | 3.49E-14 | Infection | Liver |
| P3h1 | 1.469009 | 6.812871 | 371.1007 | 3.50E-28 | 2.84E-27 | Infection | Liver |
| P3h3 | 1.992659 | 6.20901 | 103.6982 | 1.14E-14 | 2.96E-14 | Infection | Liver |
| P4hb | -0.53434 | 11.90333 | 100.1006 | 1.15E-14 | 2.97E-14 | Healthy control | Liver |
| Pa2g4 | -0.91914 | 7.39139 | 283.7381 | 4.36E-25 | 2.52E-24 | Healthy control | Liver |
| Pacsin3 | -0.7988 | 8.59727 | 97.51874 | 1.91E-14 | 4.88E-14 | Healthy control | Liver |
| Padi2 | 1.915068 | 2.128297 | 35.27723 | 2.82E-08 | 5.22E-08 | Infection | Liver |
| Cenpu | 2.522626 | 2.805113 | 110.8575 | 7.61E-18 | 2.40E-17 | Infection | Liver |
| Padi6 | 0.86036 | 1.197323 | 3.192919 | 9.11E-02 | 0.106523 | Infection | Liver |
| Pafah1b1 | -0.36874 | 7.896433 | 77.2543 | 1.43E-12 | 3.30E-12 | Healthy control | Liver |
| Pafah1b2 | 0.157212 | 6.922791 | 6.482908 | 0.013334 | 0.017289 | Infection | Liver |
| Pafah1b3 | 1.578789 | 4.816696 | 164.8862 | 3.19E-19 | 1.12E-18 | Infection | Liver |
| Pah | -1.9239 | 12.08394 | 448.9654 | 1.83E-30 | 1.89E-29 | Healthy control | Liver |
| Paip1 | 0.359988 | 9.367283 | 65.17449 | 2.55E-11 | 5.53E-11 | Infection | Liver |
| Uchl1 | 2.520726 | 1.554459 | 32.20273 | 2.55E-07 | 4.52E-07 | Infection | Liver |
| Pak2 | -0.0582 | 7.711144 | 1.482388 | 0.227915 | 0.252533 | Healthy control | Liver |
| Mpp2 | 2.520356 | 2.788392 | 114.8679 | 2.93E-18 | 9.50E-18 | Infection | Liver |
| Adamts6 | 2.519497 | 6.010975 | 551.57 | 6.62E-33 | 9.04E-32 | Infection | Liver |
| Pald1 | 0.062988 | 5.304948 | 0.560764 | 0.456731 | 0.484714 | Infection | Liver |
| Palmd | -1.80076 | 6.886317 | 663.6647 | 2.91E-35 | 5.93E-34 | Healthy control | Liver |
| Slc1a1 | 2.518417 | 1.649878 | 30.12737 | 6.10E-07 | 1.06E-06 | Infection | Liver |
| Panx2 | 1.597893 | 1.335189 | 6.985485 | 0.010673 | 0.013986 | Infection | Liver |
| Panx3 | 1.978604 | 1.324989 | 11.75117 | 0.001213 | 0.001721 | Infection | Liver |
| Pard3 | -1.57991 | 8.830924 | 876.4246 | 7.96E-39 | 2.91E-37 | Healthy control | Liver |
| Pard6a | -0.42177 | 4.156591 | 20.63076 | 2.57E-05 | 4.08E-05 | Healthy control | Liver |
| Pard6b | -1.28749 | 4.767464 | 128.4107 | 7.40E-17 | 2.19E-16 | Healthy control | Liver |
| Park7 | -1.67504 | 9.60418 | 1003.466 | 1.42E-40 | 7.91E-39 | Healthy control | Liver |
| Parp1 | -0.13453 | 7.573149 | 9.313294 | 0.003321 | 0.004545 | Healthy control | Liver |
| Parp2 | 0.260845 | 6.339758 | 18.68697 | 5.55E-05 | 8.62E-05 | Infection | Liver |
| Gli3 | 2.515741 | 4.481389 | 143.5242 | 5.89E-18 | 1.86E-17 | Infection | Liver |
| Parva | -0.29687 | 8.167353 | 49.70385 | 1.59E-09 | 3.14E-09 | Healthy control | Liver |
| Parvb | 1.019353 | 6.055557 | 120.3974 | 2.76E-16 | 7.82E-16 | Infection | Liver |
| Patj | -0.58531 | 4.836376 | 30.11702 | 7.74E-07 | 1.33E-06 | Healthy control | Liver |
| Patz1 | 0.941086 | 5.17687 | 99.03742 | 1.48E-14 | 3.78E-14 | Infection | Liver |
| Pawr | 0.767915 | 3.900382 | 37.22878 | 7.30E-08 | 1.32E-07 | Infection | Liver |
| Havcr1 | 2.514092 | 1.765671 | 37.62581 | 2.16E-08 | 4.03E-08 | Infection | Liver |
| Pax2 | 1.01309 | 1.26757 | 2.340409 | 0.134069 | 0.153405 | Infection | Liver |
| Pax3 | 0.725412 | 1.298686 | 1.655186 | 0.203887 | 0.227065 | Infection | Liver |
| Pax4 | 0.833405 | 1.195432 | 3.500105 | 0.076827 | 0.090579 | Infection | Liver |
| Ksr2 | 2.513995 | 3.327866 | 141.5756 | 2.92E-19 | 1.03E-18 | Infection | Liver |
| Pax6 | 1.3556 | 1.312848 | 4.709854 | 0.034347 | 0.042444 | Infection | Liver |

| | | | | | | | |
|---|---|---|---|---|---|---|---|
| Pax7 | 0.356912 | 1.23275 | 0.364815 | 0.550438 | 0.576824 | Infection | Liver |
| Pax8 | 1.26092 | 1.330662 | 3.991137 | 0.050406 | 0.06095 | Infection | Liver |
| Pax9 | 1.607692 | 1.413387 | 5.91587 | 0.017529 | 0.022471 | Infection | Liver |
| Paxbp1 | 0.283278 | 7.325861 | 34.18323 | 1.90E-07 | 3.39E-07 | Infection | Liver |
| Paxip1 | 0.003719 | 6.503015 | 0.004129 | 0.948968 | 0.954114 | Infection | Liver |
| Pbrm1 | 0.080635 | 8.396806 | 4.03741 | 0.048764 | 0.059108 | Infection | Liver |
| Pbx2 | -1.17595 | 6.443886 | 306.9557 | 5.75E-26 | 3.64E-25 | Healthy control | Liver |
| Pbx3 | 0.845229 | 3.97351 | 23.98751 | 7.20E-06 | 1.18E-05 | Infection | Liver |
| Pcbp3 | -1.04908 | 2.783177 | 32.10061 | 2.17E-07 | 3.85E-07 | Healthy control | Liver |
| Snai1 | 2.510973 | 5.608618 | 139.2166 | 2.30E-17 | 7.03E-17 | Infection | Liver |
| Inpp5d | 2.510578 | 7.078104 | 1069.127 | 2.23E-41 | 1.53E-39 | Infection | Liver |
| Pcdh20 | 1.300013 | 1.369362 | 5.165538 | 0.026139 | 0.032738 | Infection | Liver |
| Pcdh8 | 1.426877 | 1.382648 | 4.163016 | 0.045542 | 0.055443 | Infection | Liver |
| Pcdha2 | 0.382566 | 1.256693 | 0.387519 | 0.537268 | 0.564442 | Infection | Liver |
| Pcdha7 | 0.493952 | 1.286592 | 0.780585 | 0.381187 | 0.408263 | Infection | Liver |
| Pcgf1 | -0.55954 | 3.602652 | 18.20984 | 6.50E-05 | 0.000101 | Healthy control | Liver |
| Ikzf1 | 2.508198 | 6.428974 | 541.9289 | 9.95E-33 | 1.33E-31 | Infection | Liver |
| Pcif1 | -0.44263 | 5.637765 | 34.37791 | 1.79E-07 | 3.20E-07 | Healthy control | Liver |
| Lck | 2.505628 | 4.715154 | 271.95 | 1.81E-24 | 9.89E-24 | Infection | Liver |
| Pcm1 | 0.234315 | 6.158886 | 11.24899 | 0.001349 | 0.001906 | Infection | Liver |
| Pcmtd2 | -0.46667 | 7.432406 | 120.6915 | 2.60E-16 | 7.36E-16 | Healthy control | Liver |
| Pcna | 1.535758 | 7.194021 | 453.9537 | 1.36E-30 | 1.43E-29 | Infection | Liver |
| Pcnt | 0.559985 | 7.315955 | 75.60129 | 2.09E-12 | 4.77E-12 | Infection | Liver |
| Fscn1 | 2.501843 | 7.595123 | 510.0673 | 5.63E-32 | 6.86E-31 | Infection | Liver |
| Pcsk1 | 1.936624 | 1.598227 | 15.9405 | 0.000127 | 0.000193 | Infection | Liver |
| Pcsk4 | -2.45167 | 4.508643 | 417.3796 | 7.13E-31 | 7.75E-30 | Healthy control | Liver |
| Pcsk6 | -1.87996 | 8.868591 | 651.5565 | 4.68E-35 | 9.19E-34 | Healthy control | Liver |
| Pcsk9 | -0.25861 | 6.361243 | 3.195531 | 0.078701 | 0.092657 | Healthy control | Liver |
| Pcx | -1.76245 | 10.93108 | 380.4518 | 1.75E-28 | 1.46E-27 | Healthy control | Liver |
| Pdap1 | -0.20805 | 8.331582 | 19.95046 | 3.34E-05 | 5.28E-05 | Healthy control | Liver |
| Pdc | 0.69515 | 1.243949 | 1.249559 | 0.271778 | 0.297634 | Infection | Liver |
| Mnx1 | 2.499409 | 1.530359 | 18.08938 | 5.85E-05 | 9.07E-05 | Infection | Liver |
| Nes | 2.498886 | 5.377091 | 352.9618 | 1.79E-27 | 1.35E-26 | Infection | Liver |
| Pdcd4 | -1.19336 | 9.592241 | 224.0363 | 1.75E-22 | 7.90E-22 | Healthy control | Liver |
| Pde10a | 1.592025 | 4.298557 | 154.6378 | 1.51E-18 | 5.03E-18 | Infection | Liver |
| Pde12 | 0.171468 | 4.822024 | 3.271733 | 0.075258 | 0.088854 | Infection | Liver |
| Fzd3 | 2.498172 | 2.154146 | 51.02866 | 8.42E-11 | 1.78E-10 | Infection | Liver |
| Pde1c | -0.35046 | 2.365883 | 1.840466 | 0.178312 | 0.200007 | Healthy control | Liver |
| Pde3b | -0.77649 | 6.981415 | 92.18741 | 5.64E-14 | 1.41E-13 | Healthy control | Liver |
| Pde4c | -0.60241 | 6.854682 | 60.43024 | 8.58E-11 | 1.81E-10 | Healthy control | Liver |
| Pde6b | 0.76655 | 1.196274 | 2.711451 | 0.116067 | 0.134125 | Infection | Liver |
| Pde6c | -2.14485 | 2.260651 | 57.79988 | 1.78E-11 | 3.90E-11 | Healthy control | Liver |
| Pde6g | -1.88965 | 2.923066 | 91.87687 | 3.30E-15 | 8.80E-15 | Healthy control | Liver |
| Pde7b | -0.88266 | 5.664875 | 123.48 | 1.68E-16 | 4.84E-16 | Healthy control | Liver |
| Pde8a | -1.15549 | 7.680801 | 434.7076 | 4.53E-30 | 4.50E-29 | Healthy control | Liver |
| Pde8b | -1.63447 | 3.4278 | 93.09 | 8.73E-15 | 2.28E-14 | Healthy control | Liver |
| Pdgfa | 1.838459 | 5.759023 | 351.1739 | 1.68E-27 | 1.27E-26 | Infection | Liver |
| Spo11 | 2.496042 | 1.479916 | 30.71452 | 4.87E-07 | 8.48E-07 | Infection | Liver |
| Ccl20 | 2.496041 | 1.671196 | 39.3924 | 1.49E-08 | 2.81E-08 | Infection | Liver |
| Sptb | 2.494978 | 3.807141 | 56.5648 | 1.24E-10 | 2.60E-10 | Infection | Liver |

| | | | | | | | |
|---|---|---|---|---|---|---|---|
| Pdia3 | 0.656287 | 12.23442 | 160.8949 | 4.74E-19 | 1.63E-18 | Infection | Liver |
| Pdk1 | -1.57852 | 7.53996 | 354.7233 | 1.18E-27 | 9.08E-27 | Healthy control | Liver |
| Pdk4 | -0.69865 | 6.636082 | 16.17716 | 0.000164 | 0.000247 | Healthy control | Liver |
| Pdlim1 | -0.08272 | 1.786506 | 0.054034 | 0.816565 | 0.832361 | Healthy control | Liver |
| Col26a1 | 2.486868 | 1.401641 | 18.0573 | 7.08E-05 | 0.00011 | Infection | Liver |
| Pdpk1 | -0.16062 | 6.951988 | 10.3037 | 0.002085 | 0.002901 | Healthy control | Liver |
| Rpe65 | 2.486129 | 1.352092 | 11.4739 | 0.001395 | 0.001969 | Infection | Liver |
| Pdx1 | 0.471664 | 1.207784 | 0.765151 | 0.391576 | 0.418942 | Infection | Liver |
| Pdxk | -1.17892 | 6.285706 | 301.4352 | 9.35E-26 | 5.72E-25 | Healthy control | Liver |
| Pdyn | 1.334809 | 1.33735 | 5.260407 | 0.025567 | 0.032078 | Infection | Liver |
| Pdzd7 | 1.089852 | 2.625229 | 26.39823 | 1.82E-06 | 3.07E-06 | Infection | Liver |
| Pdzk1 | -1.30268 | 9.070781 | 237.7537 | 3.98E-23 | 1.89E-22 | Healthy control | Liver |
| Pecam1 | 1.083293 | 7.82801 | 173.0636 | 8.81E-20 | 3.25E-19 | Infection | Liver |
| Dtl | 2.48305 | 3.670231 | 149.7058 | 5.07E-19 | 1.74E-18 | Infection | Liver |
| Peg3 | -0.63211 | 4.886471 | 38.65468 | 4.52E-08 | 8.29E-08 | Healthy control | Liver |
| Per1 | 1.782482 | 5.840379 | 46.1159 | 6.34E-09 | 1.22E-08 | Infection | Liver |
| Per2 | -0.7598 | 5.001344 | 33.65019 | 2.31E-07 | 4.11E-07 | Healthy control | Liver |
| Per3 | 0.776325 | 3.362784 | 16.96529 | 0.000111 | 0.000169 | Infection | Liver |
| Pex3 | -0.61516 | 6.949519 | 43.81631 | 8.96E-09 | 1.70E-08 | Healthy control | Liver |
| Pfas | 0.104843 | 5.266918 | 1.248926 | 0.26799 | 0.294008 | Infection | Liver |
| Pfkfb1 | -0.37573 | 7.054921 | 11.58129 | 0.001159 | 0.001651 | Healthy control | Liver |
| Pfkfb2 | -0.49841 | 6.505583 | 33.48616 | 2.40E-07 | 4.26E-07 | Healthy control | Liver |
| Pfkl | 0.128755 | 8.579721 | 6.135717 | 0.015927 | 0.020533 | Infection | Liver |
| Gnaz | 2.482489 | 1.559057 | 23.18267 | 8.00E-06 | 1.31E-05 | Infection | Liver |
| Pfn4 | -0.40073 | 2.336481 | 1.539527 | 0.21783 | 0.241947 | Healthy control | Liver |
| Pgam5 | -1.55426 | 6.965614 | 649.9678 | 5.21E-35 | 1.01E-33 | Healthy control | Liver |
| Tfap2b | 2.482216 | 1.540202 | 19.42519 | 3.42E-05 | 5.39E-05 | Infection | Liver |
| Pgk1 | -1.63524 | 10.1411 | 797.579 | 1.28E-37 | 3.86E-36 | Healthy control | Liver |
| Pgk2 | -0.44319 | 1.25802 | 0.527558 | 0.471901 | 0.499437 | Healthy control | Liver |
| Pgpep1 | -1.12922 | 6.539099 | 185.0219 | 1.87E-20 | 7.19E-20 | Healthy control | Liver |
| Pgr | 1.698266 | 1.388992 | 10.32453 | 0.001978 | 0.002757 | Infection | Liver |
| Phb | -1.53857 | 7.300112 | 648.8301 | 5.39E-35 | 1.03E-33 | Healthy control | Liver |
| Phb2 | -0.71582 | 9.992612 | 177.9352 | 4.59E-20 | 1.72E-19 | Healthy control | Liver |
| Phc1 | 0.724343 | 4.37394 | 23.77454 | 7.75E-06 | 1.27E-05 | Infection | Liver |
| Phc2 | 0.39007 | 7.066971 | 82.29709 | 4.62E-13 | 1.10E-12 | Infection | Liver |
| Plp2 | 2.480397 | 4.488702 | 139.0806 | 1.29E-17 | 4.02E-17 | Infection | Liver |
| Phf10 | 0.78078 | 7.911837 | 325.0963 | 1.21E-26 | 8.18E-26 | Infection | Liver |
| Phf3 | -0.08301 | 6.341608 | 1.271781 | 0.263686 | 0.289604 | Healthy control | Liver |
| Phf6 | 0.493418 | 5.965271 | 54.68809 | 3.97E-10 | 8.12E-10 | Infection | Liver |
| Phlda2 | 1.586995 | 1.369151 | 9.273777 | 0.003394 | 0.004642 | Infection | Liver |
| Alx1 | 2.480162 | 1.364504 | 9.065455 | 0.00393 | 0.005344 | Infection | Liver |
| Phox2b | 0.607914 | 1.33978 | 1.011294 | 0.318591 | 0.344824 | Infection | Liver |
| Phtf1 | 1.059736 | 6.117101 | 244.6199 | 1.99E-23 | 9.68E-23 | Infection | Liver |
| Phykpl | -0.25367 | 5.830145 | 8.742621 | 0.004365 | 0.005914 | Healthy control | Liver |
| Pi4ka | 0.053814 | 7.24961 | 0.964199 | 0.329865 | 0.356038 | Infection | Liver |
| Trp73 | 2.478349 | 1.663878 | 40.41956 | 1.06E-08 | 2.01E-08 | Infection | Liver |
| Pias1 | 0.266836 | 6.602901 | 21.76938 | 1.64E-05 | 2.63E-05 | Infection | Liver |
| Pias2 | -0.19844 | 7.576772 | 20.94556 | 2.26E-05 | 3.60E-05 | Healthy control | Liver |
| Pias3 | 0.33425 | 5.211883 | 9.698039 | 0.002772 | 0.003824 | Infection | Liver |
| Picalm | 0.210001 | 9.096169 | 11.59473 | 0.001152 | 0.001641 | Infection | Liver |

| | | | | | | | |
|---|---|---|---|---|---|---|---|
| Pick1 | -0.48039 | 7.061496 | 65.96281 | 2.10E-11 | 4.58E-11 | Healthy control | Liver |
| Piezo1 | 1.318714 | 5.020806 | 125.5933 | 1.21E-16 | 3.54E-16 | Infection | Liver |
| Slit2 | 2.476897 | 1.657916 | 21.26068 | 1.38E-05 | 2.22E-05 | Infection | Liver |
| Pifo | 1.583634 | 1.304365 | 5.306696 | 0.025346 | 0.03184 | Infection | Liver |
| Pik3c3 | -0.96209 | 7.122536 | 365.3011 | 5.35E-28 | 4.28E-27 | Healthy control | Liver |
| Pik3ca | -0.18178 | 6.581302 | 4.132922 | 0.046252 | 0.056253 | Healthy control | Liver |
| Pik3cb | 0.2909 | 7.033924 | 11.7043 | 0.001096 | 0.001565 | Infection | Liver |
| Pik3cd | 1.91376 | 6.195751 | 292.589 | 2.02E-25 | 1.20E-24 | Infection | Liver |
| Pik3cg | 1.221381 | 4.260249 | 92.15474 | 6.61E-14 | 1.64E-13 | Infection | Liver |
| Pik3r1 | -0.57389 | 8.720015 | 34.10757 | 1.95E-07 | 3.47E-07 | Healthy control | Liver |
| Pik3r2 | -0.83292 | 7.375948 | 134.4157 | 2.62E-17 | 7.99E-17 | Healthy control | Liver |
| Pim1 | 1.140556 | 4.653071 | 95.62052 | 3.09E-14 | 7.84E-14 | Infection | Liver |
| Pim2 | -0.10512 | 5.06898 | 1.190659 | 0.279348 | 0.305531 | Healthy control | Liver |
| Pin1 | 1.386569 | 2.779529 | 52.62864 | 2.41E-10 | 4.99E-10 | Infection | Liver |
| Pink1 | -1.24149 | 10.0171 | 255.7078 | 6.30E-24 | 3.27E-23 | Healthy control | Liver |
| Pinlyp | 0.677462 | 1.467898 | 1.929385 | 0.167652 | 0.188769 | Infection | Liver |
| Pip | 1.881954 | 1.32115 | 11.40618 | 0.001397 | 0.001971 | Infection | Liver |
| Pip4k2c | 0.334243 | 7.070849 | 48.60908 | 2.18E-09 | 4.28E-09 | Infection | Liver |
| Pip4p1 | -0.07156 | 7.370968 | 1.321458 | 0.25465 | 0.280294 | Healthy control | Liver |
| Pip5k1c | -0.10739 | 7.619715 | 1.417819 | 0.238203 | 0.263349 | Healthy control | Liver |
| Pirb | 1.795088 | 6.498461 | 308.2587 | 5.07E-26 | 3.23E-25 | Infection | Liver |
| Pisd | -0.32148 | 7.114337 | 27.29036 | 2.08E-06 | 3.49E-06 | Healthy control | Liver |
| Pithd1 | -1.60172 | 5.244099 | 322.691 | 1.75E-26 | 1.16E-25 | Healthy control | Liver |
| Sidt1 | 2.472998 | 2.995367 | 131.9398 | 2.45E-19 | 8.68E-19 | Infection | Liver |
| Pitrm1 | -0.92934 | 7.46726 | 399.2 | 4.75E-29 | 4.18E-28 | Healthy control | Liver |
| Pitx1 | 1.234567 | 1.40623 | 5.099685 | 0.026696 | 0.03337 | Infection | Liver |
| Pitx2 | 1.90201 | 1.603198 | 17.04286 | 7.79E-05 | 0.00012 | Infection | Liver |
| Piwil1 | 1.701324 | 1.257493 | 6.472316 | 0.015076 | 0.019481 | Infection | Liver |
| Cd28 | 2.472866 | 2.806031 | 108.1075 | 1.66E-17 | 5.15E-17 | Infection | Liver |
| Piwil4 | 1.034857 | 1.49404 | 4.286698 | 0.040827 | 0.050007 | Infection | Liver |
| Pja1 | -0.80527 | 7.798376 | 233.4448 | 6.31E-23 | 2.95E-22 | Healthy control | Liver |
| Pja2 | -0.04799 | 7.797753 | 1.465297 | 0.230584 | 0.255321 | Healthy control | Liver |
| Pkd1 | 1.275577 | 7.309466 | 221.1391 | 2.43E-22 | 1.08E-21 | Infection | Liver |
| Pkd2 | 1.537619 | 7.742562 | 407.7716 | 2.64E-29 | 2.40E-28 | Infection | Liver |
| Pkd2l1 | 0.687297 | 1.221917 | 1.518501 | 0.228859 | 0.253467 | Infection | Liver |
| Hoxc5 | 2.472795 | 1.659539 | 34.30682 | 8.77E-08 | 1.59E-07 | Infection | Liver |
| Hoxd4 | 2.471603 | 1.728437 | 38.3935 | 1.80E-08 | 3.38E-08 | Infection | Liver |
| Pkig | 0.348841 | 6.00655 | 16.05683 | 0.000164 | 0.000248 | Infection | Liver |
| Pklr | -1.3664 | 9.667349 | 129.5043 | 7.91E-17 | 2.34E-16 | Healthy control | Liver |
| Prdm1 | 2.47138 | 3.967063 | 200.8659 | 1.21E-21 | 5.05E-21 | Infection | Liver |
| Pkn1 | 1.20378 | 7.350824 | 554.2647 | 4.89E-33 | 6.74E-32 | Infection | Liver |
| Pdgfra | 2.471326 | 7.387172 | 280.5841 | 1.63E-24 | 8.96E-24 | Infection | Liver |
| Pkp4 | -2.12303 | 7.84479 | 937.9486 | 1.09E-39 | 4.82E-38 | Healthy control | Liver |
| Pla2g10 | 1.67158 | 1.390423 | 5.241358 | 0.025225 | 0.031704 | Infection | Liver |
| Cd209d | 2.463749 | 1.630474 | 29.85321 | 4.42E-07 | 7.70E-07 | Infection | Liver |
| Pla2g2f | 0.411414 | 1.232981 | 0.4663 | 0.500001 | 0.52717 | Infection | Liver |
| Pla2g4a | 1.88551 | 5.436551 | 260.5995 | 4.39E-24 | 2.32E-23 | Infection | Liver |
| Pla2g6 | -1.90452 | 6.903081 | 820.0612 | 6.10E-38 | 1.91E-36 | Healthy control | Liver |
| Pla2r1 | 0.547677 | 6.281717 | 38.07809 | 5.33E-08 | 9.73E-08 | Infection | Liver |
| Cxcl1 | 2.462756 | 6.33129 | 137.9109 | 4.18E-17 | 1.26E-16 | Infection | Liver |

| | | | | | | | |
|---|---|---|---|---|---|---|---|
| Plac9a | -1.28807 | 1.507655 | 7.562392 | 0.006933 | 0.00924 | Healthy control | Liver |
| Shisa3 | 2.461361 | 1.575635 | 22.91847 | 9.31E-06 | 1.51E-05 | Infection | Liver |
| Plagl2 | -0.07925 | 5.317514 | 1.228867 | 0.271831 | 0.297634 | Healthy control | Liver |
| Gcnt3 | 2.46095 | 1.398402 | 16.87324 | 0.000116 | 0.000176 | Infection | Liver |
| Kirrel | 2.46021 | 5.722054 | 248.2983 | 1.53E-23 | 7.59E-23 | Infection | Liver |
| Gpr141b | 2.460054 | 2.077909 | 53.40943 | 4.05E-11 | 8.70E-11 | Infection | Liver |
| Plcb4 | 1.085196 | 4.588865 | 98.52019 | 1.75E-14 | 4.47E-14 | Infection | Liver |
| Plcg1 | 0.578344 | 7.660197 | 135.9667 | 2.04E-17 | 6.25E-17 | Infection | Liver |
| Stmn2 | 2.457473 | 1.882696 | 31.79696 | 1.57E-07 | 2.81E-07 | Infection | Liver |
| Pld1 | 0.097024 | 5.39556 | 1.878646 | 0.175336 | 0.196756 | Infection | Liver |
| Pld2 | -0.21531 | 4.840646 | 3.774017 | 0.056529 | 0.067846 | Healthy control | Liver |
| Gpa33 | 2.457438 | 1.577407 | 19.82437 | 2.71E-05 | 4.29E-05 | Infection | Liver |
| Plec | 0.760495 | 9.251824 | 161.5047 | 4.34E-19 | 1.51E-18 | Infection | Liver |
| Plekha1 | 1.716027 | 6.623261 | 707.4622 | 4.42E-36 | 1.02E-34 | Infection | Liver |
| Plekha2 | 1.807944 | 5.879812 | 304.5647 | 7.22E-26 | 4.51E-25 | Infection | Liver |
| Plekha5 | 0.578162 | 7.007884 | 68.26851 | 1.19E-11 | 2.63E-11 | Infection | Liver |
| Plekhg2 | 0.473136 | 6.098803 | 25.39862 | 4.15E-06 | 6.88E-06 | Infection | Liver |
| Plekhg4 | 1.351746 | 1.538041 | 8.1393 | 0.005299 | 0.007137 | Infection | Liver |
| Plg | -1.35826 | 13.45023 | 183.1343 | 2.33E-20 | 8.87E-20 | Healthy control | Liver |
| Plgrkt | 0.584979 | 5.632274 | 44.54541 | 7.23E-09 | 1.38E-08 | Infection | Liver |
| Plin1 | 1.863571 | 1.467171 | 9.564156 | 0.002742 | 0.003785 | Infection | Liver |
| Plin2 | -2.3826 | 10.33004 | 1391.164 | 7.63E-45 | 1.27E-42 | Healthy control | Liver |
| Plin5 | -1.65677 | 6.210222 | 148.0442 | 3.28E-18 | 1.06E-17 | Healthy control | Liver |
| Lpar1 | 2.452225 | 3.492986 | 157.8055 | 6.37E-20 | 2.37E-19 | Infection | Liver |
| Plk2 | -0.29404 | 7.032639 | 5.991675 | 0.017182 | 0.022066 | Healthy control | Liver |
| Plk3 | 1.042376 | 7.259612 | 158.4713 | 6.72E-19 | 2.29E-18 | Infection | Liver |
| Plk4 | 1.182368 | 2.642809 | 26.32164 | 1.91E-06 | 3.21E-06 | Infection | Liver |
| Pln | 1.749319 | 1.304929 | 6.38984 | 0.014815 | 0.019153 | Infection | Liver |
| Plod1 | 0.086491 | 8.143416 | 0.668504 | 0.416665 | 0.443982 | Infection | Liver |
| Plod3 | 0.730487 | 7.278107 | 85.24489 | 2.43E-13 | 5.90E-13 | Infection | Liver |
| Gnrhr | 2.448121 | 1.398727 | 9.82264 | 0.002643 | 0.003652 | Infection | Liver |
| Plpp3 | -2.10371 | 10.31689 | 899.6343 | 3.66E-39 | 1.43E-37 | Healthy control | Liver |
| Gabrq | 2.447023 | 1.895224 | 40.14742 | 6.17E-09 | 1.18E-08 | Infection | Liver |
| Npy1r | 2.444844 | 1.545381 | 20.14696 | 2.68E-05 | 4.25E-05 | Infection | Liver |
| Pls1 | -0.38392 | 5.047668 | 7.426614 | 0.008308 | 0.010992 | Healthy control | Liver |
| Pltp | 0.77917 | 9.786344 | 44.95315 | 6.85E-09 | 1.31E-08 | Infection | Liver |
| Plvap | -0.62618 | 7.795387 | 30.01199 | 8.18E-07 | 1.41E-06 | Healthy control | Liver |
| Ccl24 | 2.444463 | 6.627368 | 232.7235 | 1.32E-22 | 6.04E-22 | Infection | Liver |
| Plxna1 | 1.279899 | 5.693396 | 174.7822 | 7.24E-20 | 2.69E-19 | Infection | Liver |
| Plxna2 | -0.29172 | 4.988085 | 3.629066 | 0.061345 | 0.073309 | Healthy control | Liver |
| Plxna3 | 1.319962 | 3.458845 | 66.23005 | 1.78E-11 | 3.89E-11 | Infection | Liver |
| Plxna4 | 1.708358 | 4.215597 | 56.05026 | 2.93E-10 | 6.05E-10 | Infection | Liver |
| Plxnb2 | -0.06208 | 10.11997 | 3.246015 | 0.076353 | 0.090083 | Healthy control | Liver |
| Plxnc1 | -0.5905 | 7.030522 | 68.20716 | 1.21E-11 | 2.67E-11 | Healthy control | Liver |
| Plxnd1 | 0.921776 | 7.273583 | 127.7326 | 7.83E-17 | 2.31E-16 | Infection | Liver |
| Gpsm1 | 2.442854 | 5.364718 | 237.1367 | 5.29E-23 | 2.48E-22 | Infection | Liver |
| Pmch | 1.882534 | 1.367544 | 8.289458 | 0.005414 | 0.007285 | Infection | Liver |
| Pml | -0.55786 | 7.453579 | 142.3597 | 7.44E-18 | 2.35E-17 | Healthy control | Liver |
| Pmm2 | -0.91513 | 6.033064 | 187.3708 | 1.41E-20 | 5.45E-20 | Healthy control | Liver |
| Pms1 | -0.64875 | 5.07354 | 50.39956 | 1.34E-09 | 2.66E-09 | Healthy control | Liver |

| | | | | | | | |
|---|---|---|---|---|---|---|---|
| Pms2 | 0.7946 | 2.890454 | 17.402 | 8.38E-05 | 0.000129 | Infection | Liver |
| Pnkd | -1.68284 | 7.01345 | 325.4323 | 1.21E-26 | 8.18E-26 | Healthy control | Liver |
| Pnoc | 1.2188 | 1.28821 | 4.319475 | 0.043077 | 0.052583 | Infection | Liver |
| Pnpla1 | 1.177896 | 1.356207 | 4.142365 | 0.046085 | 0.056064 | Infection | Liver |
| Pnpla2 | -0.22226 | 8.634743 | 17.94019 | 7.52E-05 | 0.000116 | Healthy control | Liver |
| Pnpla3 | 0.819836 | 7.338653 | 19.0224 | 5.30E-05 | 8.25E-05 | Infection | Liver |
| Podxl | 0.737845 | 6.642916 | 87.03293 | 1.66E-13 | 4.06E-13 | Infection | Liver |
| Polb | -0.84849 | 5.92737 | 111.6646 | 1.33E-15 | 3.63E-15 | Healthy control | Liver |
| Polg | -0.27685 | 8.217054 | 56.31738 | 2.54E-10 | 5.25E-10 | Healthy control | Liver |
| Polk | 0.492636 | 5.114922 | 35.86754 | 1.10E-07 | 1.99E-07 | Infection | Liver |
| Polr1e | -0.40485 | 5.242301 | 21.17194 | 2.08E-05 | 3.32E-05 | Healthy control | Liver |
| Polr2a | -0.44139 | 8.11972 | 33.51979 | 2.37E-07 | 4.21E-07 | Healthy control | Liver |
| Polr2b | -0.54165 | 6.565112 | 108.3219 | 2.41E-15 | 6.47E-15 | Healthy control | Liver |
| Polr2e | -0.71923 | 7.093752 | 145.9871 | 4.27E-18 | 1.37E-17 | Healthy control | Liver |
| Polr2f | 0.682564 | 7.855272 | 163.721 | 3.18E-19 | 1.12E-18 | Infection | Liver |
| Polr2h | -0.19214 | 5.678383 | 5.71789 | 0.019779 | 0.025227 | Healthy control | Liver |
| Polr2j | -2.43185 | 5.926626 | 713.875 | 4.49E-36 | 1.03E-34 | Healthy control | Liver |
| Polr3e | 0.21099 | 6.092529 | 13.60824 | 0.00047 | 0.000688 | Infection | Liver |
| Pom121 | -0.08558 | 6.318682 | 2.230336 | 0.140289 | 0.159756 | Healthy control | Liver |
| Pomc | 1.373761 | 2.849334 | 52.15842 | 3.31E-10 | 6.81E-10 | Infection | Liver |
| Pon1 | -1.7053 | 11.06437 | 321.8433 | 1.58E-26 | 1.05E-25 | Healthy control | Liver |
| Pon3 | 0.461021 | 8.585483 | 35.98849 | 1.05E-07 | 1.89E-07 | Infection | Liver |
| Por | -1.30493 | 8.209955 | 591.2316 | 7.69E-34 | 1.19E-32 | Healthy control | Liver |
| Gbp2b | 2.440317 | 7.436822 | 156.8252 | 3.25E-18 | 1.05E-17 | Infection | Liver |
| Unc13b | 2.432074 | 3.989115 | 214.9588 | 2.20E-22 | 9.84E-22 | Infection | Liver |
| Ebf2 | 2.43117 | 1.416481 | 14.19518 | 0.000364 | 0.000537 | Infection | Liver |
| Pou2f1 | -0.01839 | 5.564326 | 0.071304 | 0.790316 | 0.807413 | Healthy control | Liver |
| Pou2f3 | 0.988536 | 1.519555 | 5.146562 | 0.02522 | 0.031704 | Infection | Liver |
| Cubn | 2.423699 | 1.580357 | 26.21684 | 1.99E-06 | 3.35E-06 | Infection | Liver |
| Pou3f2 | 0.610182 | 1.380962 | 1.027222 | 0.313713 | 0.340071 | Infection | Liver |
| Spdl1 | 2.423443 | 3.015936 | 120.0749 | 4.60E-18 | 1.47E-17 | Infection | Liver |
| Pou3f4 | 1.356128 | 1.308667 | 4.457156 | 0.039339 | 0.048291 | Infection | Liver |
| Pou4f1 | 0.961043 | 1.649691 | 7.196468 | 0.008285 | 0.010966 | Infection | Liver |
| Pou4f2 | 1.102044 | 1.278532 | 3.361061 | 0.073441 | 0.086831 | Infection | Liver |
| Sct | 2.421611 | 1.823812 | 37.93881 | 1.53E-08 | 2.88E-08 | Infection | Liver |
| Pou5f1 | 0.850467 | 1.217514 | 1.987591 | 0.171546 | 0.19285 | Infection | Liver |
| Pou6f1 | 0.82908 | 6.103132 | 81.54769 | 5.48E-13 | 1.30E-12 | Infection | Liver |
| Pou6f2 | 0.671759 | 1.302118 | 1.303334 | 0.258698 | 0.284312 | Infection | Liver |
| Ppara | -2.50538 | 8.82625 | 1074.083 | 1.88E-41 | 1.32E-39 | Healthy control | Liver |
| Pparg | -1.39985 | 5.644537 | 243.8987 | 2.38E-23 | 1.16E-22 | Healthy control | Liver |
| Ppargc1a | 0.012548 | 5.51006 | 0.007039 | 0.933402 | 0.940541 | Infection | Liver |
| Ppargc1b | 0.414765 | 6.361843 | 22.93235 | 1.05E-05 | 1.70E-05 | Infection | Liver |
| Ppfibp1 | 0.712382 | 8.763419 | 336.9537 | 4.65E-27 | 3.25E-26 | Infection | Liver |
| Ppif | -0.91901 | 7.566065 | 251.5853 | 9.58E-24 | 4.87E-23 | Healthy control | Liver |
| Ppl | 0.059587 | 5.11597 | 0.191294 | 0.663336 | 0.68637 | Infection | Liver |
| Ppm1d | 0.913712 | 4.156472 | 46.54106 | 4.23E-09 | 8.20E-09 | Infection | Liver |
| Epha3 | 2.416997 | 2.787426 | 118.844 | 1.52E-18 | 5.07E-18 | Infection | Liver |
| Ppp1cc | -1.65996 | 6.883347 | 463.7496 | 7.79E-31 | 8.39E-30 | Healthy control | Liver |
| Ppp1r10 | -0.58107 | 7.386342 | 25.15309 | 4.62E-06 | 7.64E-06 | Healthy control | Liver |
| Ppp1r16b | -0.20852 | 5.821012 | 8.430645 | 0.005074 | 0.006842 | Healthy control | Liver |

| | | | | | | | |
|---|---|---|---|---|---|---|---|
| Ppp1r1b | -1.78014 | 4.481685 | 210.9064 | 4.06E-22 | 1.78E-21 | Healthy control | Liver |
| Ppp1r1c | 1.459869 | 1.423923 | 6.368113 | 0.013538 | 0.017534 | Infection | Liver |
| Ppp1r2 | -0.3007 | 7.030869 | 26.67455 | 2.60E-06 | 4.35E-06 | Healthy control | Liver |
| Ppp1r3a | 0.36327 | 1.282403 | 0.424036 | 0.517899 | 0.545466 | Infection | Liver |
| Ppp1r7 | -0.91652 | 6.95198 | 214.2531 | 5.34E-22 | 2.31E-21 | Healthy control | Liver |
| Ppp2ca | -0.7855 | 8.856526 | 202.5623 | 2.08E-21 | 8.57E-21 | Healthy control | Liver |
| Ppp2r1a | -0.64984 | 7.749619 | 160.4372 | 5.07E-19 | 1.74E-18 | Healthy control | Liver |
| Ppp2r1b | -0.87828 | 5.752603 | 194.1614 | 6.15E-21 | 2.43E-20 | Healthy control | Liver |
| Ppp2r2a | -0.02308 | 4.812781 | 0.058012 | 0.810449 | 0.826621 | Healthy control | Liver |
| Tnf | 2.416978 | 2.982168 | 93.61362 | 1.28E-15 | 3.50E-15 | Infection | Liver |
| Ppp2r5c | -0.53994 | 8.56492 | 138.0532 | 1.46E-17 | 4.52E-17 | Healthy control | Liver |
| Ppp3cb | -1.13057 | 7.186602 | 365.759 | 5.18E-28 | 4.15E-27 | Healthy control | Liver |
| Ppp3r1 | -0.83352 | 6.835248 | 121.7583 | 2.17E-16 | 6.20E-16 | Healthy control | Liver |
| Ppp3r2 | 1.309494 | 1.255793 | 4.072072 | 0.051724 | 0.062348 | Infection | Liver |
| Ppp4r3a | 0.284864 | 8.696483 | 18.98732 | 4.91E-05 | 7.66E-05 | Infection | Liver |
| Ppp5c | -1.18241 | 7.468978 | 557.2717 | 4.21E-33 | 5.88E-32 | Healthy control | Liver |
| Ppp6c | -0.75017 | 7.760431 | 245.715 | 1.74E-23 | 8.52E-23 | Healthy control | Liver |
| Ppp6r3 | -0.70347 | 7.948663 | 316.7706 | 2.41E-26 | 1.59E-25 | Healthy control | Liver |
| Ppt1 | 0.022441 | 8.761707 | 0.16594 | 0.685118 | 0.7073 | Infection | Liver |
| Praf2 | 1.032722 | 5.815372 | 76.90532 | 1.56E-12 | 3.60E-12 | Infection | Liver |
| Prap1 | 0.762038 | 1.400559 | 1.602426 | 0.209096 | 0.232652 | Infection | Liver |
| Ebi3 | 2.416778 | 3.821441 | 220.5558 | 7.44E-23 | 3.45E-22 | Infection | Liver |
| Prdm13 | 0.964081 | 1.303465 | 2.102521 | 0.153118 | 0.17334 | Infection | Liver |
| Prdm14 | 0.275681 | 1.217649 | 0.262118 | 0.612913 | 0.637624 | Infection | Liver |
| Tespa1 | 2.416753 | 1.678843 | 23.95436 | 4.32E-06 | 7.16E-06 | Infection | Liver |
| Prdm9 | -0.17059 | 5.074209 | 2.793773 | 0.099587 | 0.115992 | Healthy control | Liver |
| Preb | -0.92979 | 8.63439 | 518.0815 | 3.30E-32 | 4.12E-31 | Healthy control | Liver |
| Prelp | 0.06102 | 8.246499 | 0.550239 | 0.46096 | 0.488891 | Infection | Liver |
| Prep | -0.39618 | 7.474566 | 66.42701 | 1.87E-11 | 4.09E-11 | Healthy control | Liver |
| Prf1 | 1.907182 | 2.687853 | 33.71064 | 9.26E-08 | 1.68E-07 | Infection | Liver |
| Prickle3 | 0.337664 | 4.821279 | 14.88984 | 0.000271 | 0.000403 | Infection | Liver |
| Primpol | 0.12506 | 5.085294 | 2.206373 | 0.14242 | 0.162073 | Infection | Liver |
| Prkaa1 | -0.71477 | 6.689301 | 89.96795 | 8.96E-14 | 2.21E-13 | Healthy control | Liver |
| Prkaa2 | -0.88807 | 8.478713 | 81.61508 | 5.36E-13 | 1.28E-12 | Healthy control | Liver |
| Prkab1 | -0.49135 | 8.21488 | 103.7031 | 5.73E-15 | 1.51E-14 | Healthy control | Liver |
| Prkaca | -0.28971 | 8.763329 | 34.64951 | 1.63E-07 | 2.91E-07 | Healthy control | Liver |
| Prkacb | -0.51091 | 7.718868 | 101.249 | 9.20E-15 | 2.40E-14 | Healthy control | Liver |
| Prkar1a | -0.37168 | 9.938294 | 91.32748 | 6.71E-14 | 1.67E-13 | Healthy control | Liver |
| Klrg1 | 2.414558 | 2.62853 | 65.90525 | 1.05E-12 | 2.44E-12 | Infection | Liver |
| Prkca | -0.23587 | 5.791119 | 7.541942 | 0.00784 | 0.010399 | Healthy control | Liver |
| Prkcd | 1.793126 | 7.673156 | 619.6446 | 2.00E-34 | 3.37E-33 | Infection | Liver |
| Prkce | -0.36605 | 6.7921 | 59.65742 | 1.05E-10 | 2.21E-10 | Healthy control | Liver |
| Mmp14 | 2.412464 | 9.508017 | 319.487 | 5.78E-26 | 3.65E-25 | Infection | Liver |
| Prkci | -0.92155 | 5.953546 | 200.7472 | 2.71E-21 | 1.11E-20 | Healthy control | Liver |
| Prkcq | 1.234175 | 5.281954 | 137.014 | 1.82E-17 | 5.61E-17 | Infection | Liver |
| Prkcz | -2.06459 | 5.161433 | 274.6885 | 9.64E-25 | 5.42E-24 | Healthy control | Liver |
| Prkd1 | 0.406282 | 2.079193 | 1.927604 | 0.167839 | 0.188937 | Infection | Liver |
| Prkd3 | -0.56075 | 9.632568 | 76.01589 | 1.90E-12 | 4.34E-12 | Healthy control | Liver |
| Prkdc | -0.51287 | 4.411122 | 17.4215 | 9.37E-05 | 0.000144 | Healthy control | Liver |
| Zc3h12a | 2.412149 | 5.574417 | 552.5249 | 7.01E-33 | 9.51E-32 | Infection | Liver |

| | | | | | | | |
|---|---|---|---|---|---|---|---|
| Prkn | -0.77808 | 4.267358 | 38.58739 | 4.47E-08 | 8.19E-08 | Healthy control | Liver |
| Prl | 1.784591 | 1.350042 | 6.578767 | 0.012968 | 0.016848 | Infection | Liver |
| Csf2rb2 | 2.412089 | 5.103732 | 261.1945 | 5.01E-24 | 2.62E-23 | Infection | Liver |
| Prl3a1 | 0.911561 | 1.324748 | 2.104887 | 0.152223 | 0.172443 | Infection | Liver |
| Prl3b1 | 1.000806 | 1.207605 | 4.574954 | 0.042871 | 0.052389 | Infection | Liver |
| F2rl2 | 2.411101 | 1.79582 | 29.40541 | 4.18E-07 | 7.29E-07 | Infection | Liver |
| Sim1 | 2.410101 | 1.569286 | 12.97992 | 0.000541 | 0.000788 | Infection | Liver |
| Itpa | 2.40951 | 4.380626 | 255.5431 | 6.63E-24 | 3.42E-23 | Infection | Liver |
| Katnal2 | 2.409263 | 1.544714 | 24.78186 | 4.27E-06 | 7.07E-06 | Infection | Liver |
| Aurka | 2.408902 | 4.180375 | 210.3946 | 7.23E-22 | 3.11E-21 | Infection | Liver |
| Prl8a6 | 1.904651 | 1.850032 | 15.88823 | 0.000115 | 0.000176 | Infection | Liver |
| Meis3 | 2.408111 | 5.107848 | 527.1764 | 3.02E-32 | 3.80E-31 | Infection | Liver |
| Prmt1 | -0.57475 | 2.083402 | 3.920627 | 0.050299 | 0.06085 | Healthy control | Liver |
| Prmt3 | 0.58206 | 6.585256 | 90.20424 | 8.52E-14 | 2.11E-13 | Infection | Liver |
| Prmt5 | -0.00337 | 6.228367 | 0.003289 | 0.954446 | 0.958659 | Healthy control | Liver |
| Prmt7 | -0.78239 | 5.501911 | 95.58641 | 2.94E-14 | 7.45E-14 | Healthy control | Liver |
| Prodh | -2.11804 | 8.755294 | 544.8025 | 7.95E-33 | 1.07E-31 | Healthy control | Liver |
| Prok1 | 0.668766 | 3.589916 | 22.76243 | 1.08E-05 | 1.75E-05 | Infection | Liver |
| Prokr2 | 1.462688 | 1.432109 | 7.840871 | 0.006347 | 0.008504 | Infection | Liver |
| Prom1 | -1.13896 | 6.436482 | 80.86918 | 6.42E-13 | 1.52E-12 | Healthy control | Liver |
| Prom2 | 0.563968 | 1.863431 | 2.336726 | 0.12886 | 0.147885 | Infection | Liver |
| Prop1 | 0.594803 | 1.246046 | 1.010231 | 0.321848 | 0.34791 | Infection | Liver |
| Prorp | -0.38347 | 5.136283 | 14.99115 | 0.000259 | 0.000386 | Healthy control | Liver |
| Prox1 | -1.37119 | 8.491969 | 145.9749 | 5.38E-18 | 1.71E-17 | Healthy control | Liver |
| Prpf19 | -1.03966 | 8.151265 | 361.3972 | 7.09E-28 | 5.59E-27 | Healthy control | Liver |
| Prpf3 | 0.059099 | 5.007975 | 0.396309 | 0.531276 | 0.558616 | Infection | Liver |
| Prpf31 | 0.126667 | 6.03579 | 3.644506 | 0.060788 | 0.072696 | Infection | Liver |
| Prpf4b | -0.15242 | 7.814864 | 5.80203 | 0.018926 | 0.024182 | Healthy control | Liver |
| Prpf8 | -1.81305 | 9.078988 | 1912.182 | 4.65E-49 | 3.33E-46 | Healthy control | Liver |
| Prph | 1.712218 | 1.316161 | 9.489006 | 0.003709 | 0.005055 | Infection | Liver |
| Prph2 | 0.990282 | 1.278775 | 2.922962 | 0.093913 | 0.109715 | Infection | Liver |
| Prps1 | -0.11733 | 6.080835 | 2.258716 | 0.137836 | 0.157248 | Healthy control | Liver |
| Prpsap2 | -0.89378 | 5.982506 | 187.7273 | 1.35E-20 | 5.23E-20 | Healthy control | Liver |
| Prr5 | 0.622522 | 5.12517 | 41.32406 | 1.94E-08 | 3.63E-08 | Infection | Liver |
| Prrc2b | -0.36448 | 7.637737 | 26.83802 | 2.45E-06 | 4.10E-06 | Healthy control | Liver |
| Pcp2 | 2.407942 | 1.481991 | 12.70634 | 0.000667 | 0.000964 | Infection | Liver |
| Adcyap1r1 | 2.406841 | 5.312675 | 174.3791 | 9.31E-20 | 3.42E-19 | Infection | Liver |
| Prss1 | 1.672849 | 1.404825 | 7.797301 | 0.006787 | 0.00907 | Infection | Liver |
| Prss21 | 0.664843 | 1.218652 | 1.445014 | 0.240262 | 0.265567 | Infection | Liver |
| Prss41 | -0.70601 | 2.025588 | 5.277071 | 0.023576 | 0.029788 | Healthy control | Liver |
| Prss42 | 1.365603 | 1.279184 | 4.761054 | 0.034464 | 0.042568 | Infection | Liver |
| Serpini1 | 2.405688 | 4.220416 | 298.1538 | 8.96E-26 | 5.50E-25 | Infection | Liver |
| Prune1 | 0.386693 | 6.055765 | 30.48887 | 6.70E-07 | 1.16E-06 | Infection | Liver |
| Actg2 | 2.405034 | 1.789404 | 30.50668 | 2.68E-07 | 4.73E-07 | Infection | Liver |
| Psd2 | 1.744268 | 1.453032 | 14.16546 | 0.00033 | 0.000488 | Infection | Liver |
| Psen1 | -0.14294 | 7.905766 | 9.103464 | 0.00367 | 0.005005 | Healthy control | Liver |
| Psen2 | -2.35673 | 10.43777 | 631.2112 | 1.17E-34 | 2.07E-33 | Healthy control | Liver |
| Psma4 | -1.23962 | 9.415387 | 510.9403 | 4.88E-32 | 5.96E-31 | Healthy control | Liver |
| Psmb11 | 1.142295 | 1.289118 | 4.021019 | 0.050318 | 0.060858 | Infection | Liver |
| Psmb6 | -0.7982 | 9.430508 | 246.1126 | 1.66E-23 | 8.17E-23 | Healthy control | Liver |

| | | | | | | | |
|---|---|---|---|---|---|---|---|
| Csta1 | 2.404786 | 1.330012 | 9.78972 | 0.003104 | 0.004263 | Infection | Liver |
| Psmc2 | -0.48189 | 8.751996 | 78.08091 | 1.18E-12 | 2.75E-12 | Healthy control | Liver |
| Psmd1 | -1.58564 | 8.569903 | 578.5763 | 1.43E-33 | 2.11E-32 | Healthy control | Liver |
| Psmd10 | -0.3195 | 5.371557 | 15.45166 | 0.000213 | 0.000319 | Healthy control | Liver |
| Psmd11 | -1.51586 | 6.777563 | 624.75 | 1.65E-34 | 2.83E-33 | Healthy control | Liver |
| Psmd12 | -0.68069 | 8.488794 | 100.1958 | 1.13E-14 | 2.92E-14 | Healthy control | Liver |
| Psmd13 | -0.23732 | 8.27058 | 27.76983 | 1.75E-06 | 2.95E-06 | Healthy control | Liver |
| Psmd14 | -1.2199 | 8.332819 | 335.4065 | 5.27E-27 | 3.66E-26 | Healthy control | Liver |
| Psmd3 | -0.93691 | 8.540164 | 266.1423 | 2.27E-24 | 1.23E-23 | Healthy control | Liver |
| Psmd5 | -0.99308 | 6.856104 | 225.9962 | 1.43E-22 | 6.50E-22 | Healthy control | Liver |
| Psmd6 | -0.84253 | 8.093005 | 240.9306 | 2.86E-23 | 1.37E-22 | Healthy control | Liver |
| Psmd9 | -1.40221 | 8.158876 | 818.9227 | 5.93E-38 | 1.88E-36 | Healthy control | Liver |
| Ptafr | 1.650718 | 3.907196 | 110.0382 | 1.88E-15 | 5.08E-15 | Infection | Liver |
| Ptar1 | -0.06744 | 6.49896 | 0.840674 | 0.362681 | 0.389443 | Healthy control | Liver |
| Ptbp1 | 0.446848 | 9.503755 | 169.5018 | 1.43E-19 | 5.15E-19 | Infection | Liver |
| Ptch1 | -1.06409 | 6.143475 | 146.5711 | 4.01E-18 | 1.28E-17 | Healthy control | Liver |
| Ptch2 | -1.64618 | 2.260254 | 42.44895 | 3.17E-09 | 6.20E-09 | Healthy control | Liver |
| Tlr11 | 2.402813 | 2.971866 | 106.9207 | 5.78E-17 | 1.72E-16 | Infection | Liver |
| Pten | -1.51169 | 8.928188 | 1175.457 | 1.24E-42 | 1.07E-40 | Healthy control | Liver |
| Nxn | 2.399708 | 5.459104 | 243.8837 | 2.52E-23 | 1.22E-22 | Infection | Liver |
| Ptgdr | 1.658203 | 1.293595 | 8.835343 | 0.004843 | 0.006541 | Infection | Liver |
| Tacstd2 | 2.399705 | 1.544513 | 22.96995 | 9.50E-06 | 1.54E-05 | Infection | Liver |
| Ptger2 | 1.529897 | 4.257153 | 122.0894 | 2.46E-16 | 7.00E-16 | Infection | Liver |
| Ubash3b | 2.398106 | 5.341312 | 351.3989 | 2.00E-27 | 1.49E-26 | Infection | Liver |
| Frrs1l | 2.396883 | 1.481742 | 12.59192 | 0.000714 | 0.00103 | Infection | Liver |
| Lgr6 | 2.395106 | 1.510806 | 17.92142 | 6.33E-05 | 9.81E-05 | Infection | Liver |
| Ptgfr | -0.11374 | 1.440089 | 0.040285 | 0.841372 | 0.854588 | Healthy control | Liver |
| Cd300lb | 2.394738 | 4.958023 | 276.9432 | 1.10E-24 | 6.13E-24 | Infection | Liver |
| Ptgr1 | 0.605086 | 5.943536 | 33.83743 | 2.14E-07 | 3.81E-07 | Infection | Liver |
| Ptgs1 | 1.555127 | 8.045209 | 460.8523 | 8.88E-31 | 9.48E-30 | Infection | Liver |
| Fut4 | 2.392701 | 3.399356 | 150.2517 | 1.70E-19 | 6.09E-19 | Infection | Liver |
| Nr2e1 | 2.391752 | 1.436931 | 16.43574 | 0.000139 | 0.000211 | Infection | Liver |
| Pth1r | -1.12438 | 7.098938 | 140.6238 | 9.80E-18 | 3.07E-17 | Healthy control | Liver |
| Pth2r | 1.826117 | 1.377603 | 9.844664 | 0.002579 | 0.003567 | Infection | Liver |
| Pthlh | 1.178066 | 1.393734 | 5.378036 | 0.022968 | 0.029049 | Infection | Liver |
| Ptk2 | 0.04905 | 7.262266 | 1.420119 | 0.237827 | 0.26305 | Infection | Liver |
| Ptk2b | 0.325981 | 7.472668 | 40.36059 | 2.59E-08 | 4.80E-08 | Infection | Liver |
| Ptk6 | -0.24514 | 1.792481 | 0.372658 | 0.542646 | 0.569017 | Healthy control | Liver |
| Ptp4a1 | -1.68831 | 8.512701 | 591.0951 | 7.74E-34 | 1.19E-32 | Healthy control | Liver |
| Ptpa | -0.5252 | 8.467329 | 86.76126 | 1.75E-13 | 4.28E-13 | Healthy control | Liver |
| Ptpn1 | 1.013238 | 7.53651 | 246.3334 | 1.63E-23 | 8.03E-23 | Infection | Liver |
| Ptpn11 | 0.002596 | 7.63924 | 0.001953 | 0.964893 | 0.968181 | Infection | Liver |
| Ptpn2 | 0.564793 | 7.255796 | 148.0157 | 3.13E-18 | 1.01E-17 | Infection | Liver |
| Glis3 | 2.389033 | 4.988448 | 159.0994 | 7.33E-19 | 2.49E-18 | Infection | Liver |
| Ptpn23 | 1.065376 | 7.372992 | 316.5133 | 2.47E-26 | 1.63E-25 | Infection | Liver |
| Ptpn5 | 1.723218 | 1.596201 | 16.50212 | 9.43E-05 | 0.000145 | Infection | Liver |
| Ptpn6 | 0.263447 | 7.034043 | 13.26364 | 0.000546 | 0.000795 | Infection | Liver |
| Ptpra | -1.00244 | 6.55913 | 199.5316 | 3.05E-21 | 1.24E-20 | Healthy control | Liver |
| Dppa3 | 2.387126 | 1.517546 | 15.28739 | 0.0002 | 0.0003 | Infection | Liver |
| Ptprf | -0.8661 | 7.035113 | 88.03224 | 1.34E-13 | 3.30E-13 | Healthy control | Liver |

| | | | | | | | |
|---|---|---|---|---|---|---|---|
| Ptprk | -1.09376 | 8.575342 | 365.4366 | 5.24E-28 | 4.20E-27 | Healthy control | Liver |
| Ptprn2 | -0.88394 | 3.333039 | 17.0045 | 0.000102 | 0.000156 | Healthy control | Liver |
| Ptprq | 0.848181 | 1.374891 | 2.157929 | 0.145978 | 0.165894 | Infection | Liver |
| Ptpru | 0.833454 | 3.134096 | 13.44758 | 0.000487 | 0.000712 | Infection | Liver |
| Ptprz1 | 0.337831 | 1.362649 | 0.378714 | 0.540189 | 0.566797 | Infection | Liver |
| Pttg1 | 1.271475 | 4.895594 | 133.9033 | 3.14E-17 | 9.55E-17 | Infection | Liver |
| Mstn | 2.38482 | 1.448308 | 13.63481 | 0.000433 | 0.000635 | Infection | Liver |
| Il12rb2 | 2.384742 | 2.270418 | 81.66255 | 3.18E-15 | 8.49E-15 | Infection | Liver |
| Pwp1 | 0.114747 | 5.923438 | 2.65646 | 0.108096 | 0.125174 | Infection | Liver |
| Pxn | -0.1787 | 5.592375 | 7.496528 | 0.008019 | 0.010623 | Healthy control | Liver |
| Ghsr | 2.381973 | 1.737021 | 16.29302 | 0.00011 | 0.000168 | Infection | Liver |
| Pycrl | -1.38207 | 6.347067 | 349.059 | 1.92E-27 | 1.44E-26 | Healthy control | Liver |
| Cxcl9 | 2.380817 | 7.79889 | 67.58551 | 2.56E-11 | 5.55E-11 | Infection | Liver |
| Pygl | -1.27505 | 10.83495 | 228.9212 | 1.03E-22 | 4.71E-22 | Healthy control | Liver |
| Pyroxd1 | -0.16956 | 6.13785 | 7.535922 | 0.007862 | 0.010425 | Healthy control | Liver |
| Pzp | -0.83819 | 13.74586 | 93.57755 | 4.23E-14 | 1.07E-13 | Healthy control | Liver |
| Cdca8 | 2.380747 | 4.511788 | 331.8542 | 9.10E-27 | 6.20E-26 | Infection | Liver |
| Qsox1 | -0.91872 | 9.74307 | 120.383 | 2.73E-16 | 7.74E-16 | Healthy control | Liver |
| Rab10 | -1.03957 | 8.712364 | 530.0885 | 1.72E-32 | 2.24E-31 | Healthy control | Liver |
| Rab11a | -0.86246 | 5.447599 | 99.5883 | 1.34E-14 | 3.44E-14 | Healthy control | Liver |
| Rab1a | -0.70435 | 8.456828 | 233.6356 | 6.17E-23 | 2.89E-22 | Healthy control | Liver |
| Rab20 | -0.87961 | 4.667147 | 50.84852 | 1.18E-09 | 2.35E-09 | Healthy control | Liver |
| Rab21 | 0.073286 | 8.032417 | 2.248127 | 0.138741 | 0.158109 | Infection | Liver |
| Rab23 | -0.32224 | 5.085222 | 16.64077 | 0.000129 | 0.000196 | Healthy control | Liver |
| Rab24 | -0.3005 | 6.698594 | 33.38536 | 2.48E-07 | 4.40E-07 | Healthy control | Liver |
| Rab27a | 1.027881 | 5.533348 | 79.5036 | 8.74E-13 | 2.05E-12 | Infection | Liver |
| Il2 | 2.378765 | 1.747268 | 19.36402 | 2.86E-05 | 4.52E-05 | Infection | Liver |
| Rab28 | -0.39567 | 7.133253 | 73.38198 | 3.50E-12 | 7.91E-12 | Healthy control | Liver |
| Rab29 | -0.08038 | 7.208307 | 3.038161 | 0.086178 | 0.101151 | Healthy control | Liver |
| Rab32 | 0.201747 | 8.341118 | 6.524621 | 0.013054 | 0.016942 | Infection | Liver |
| Nanog | 2.377928 | 1.521366 | 20.18936 | 2.49E-05 | 3.95E-05 | Infection | Liver |
| Rab39 | 0.07653 | 3.743049 | 0.275039 | 0.601795 | 0.626889 | Infection | Liver |
| Rab3a | 0.043743 | 5.531997 | 0.258248 | 0.613095 | 0.63768 | Infection | Liver |
| Rab3b | 0.070708 | 1.446904 | 0.023274 | 0.879035 | 0.890515 | Infection | Liver |
| Rab3d | 0.580265 | 4.87915 | 43.64515 | 9.64E-09 | 1.83E-08 | Infection | Liver |
| Rab3gap2 | 0.504255 | 6.916757 | 118.7725 | 3.63E-16 | 1.02E-15 | Infection | Liver |
| Rab43 | -1.31057 | 8.878002 | 664.7605 | 2.61E-35 | 5.36E-34 | Healthy control | Liver |
| Rab4b | -0.342 | 5.830415 | 19.90837 | 3.41E-05 | 5.37E-05 | Healthy control | Liver |
| Rab5a | -2.75177 | 6.518186 | 908.5968 | 3.66E-39 | 1.43E-37 | Healthy control | Liver |
| Rab5b | -1.45086 | 8.82047 | 763.7525 | 4.58E-37 | 1.29E-35 | Healthy control | Liver |
| Rab5c | -1.07116 | 9.679219 | 625.7693 | 1.50E-34 | 2.61E-33 | Healthy control | Liver |
| Rab6a | -1.16355 | 7.92808 | 400.523 | 4.32E-29 | 3.83E-28 | Healthy control | Liver |
| Rab7 | -0.40364 | 11.10403 | 89.92504 | 8.99E-14 | 2.22E-13 | Healthy control | Liver |
| Slc7a10 | 2.37702 | 1.325194 | 18.21356 | 7.92E-05 | 0.000122 | Infection | Liver |
| Rab8a | -0.62918 | 8.575342 | 163.7798 | 3.15E-19 | 1.11E-18 | Healthy control | Liver |
| Dlx5 | 2.376616 | 1.396116 | 13.96533 | 0.000412 | 0.000606 | Infection | Liver |
| Rabep1 | -0.37743 | 8.657658 | 79.05594 | 9.49E-13 | 2.21E-12 | Healthy control | Liver |
| Rabep2 | -1.05568 | 5.073781 | 111.0796 | 1.55E-15 | 4.22E-15 | Healthy control | Liver |
| Rac1 | -0.91598 | 9.648095 | 278.7934 | 6.83E-25 | 3.89E-24 | Healthy control | Liver |
| Hoxb9 | 2.376266 | 1.334114 | 15.96718 | 0.000197 | 0.000296 | Infection | Liver |

| | | | | | | | |
|---|---|---|---|---|---|---|---|
| Rack1 | -1.20366 | 5.683506 | 201.3195 | 2.66E-21 | 1.09E-20 | Healthy control | Liver |
| Daam2 | 2.375139 | 5.804315 | 157.2827 | 1.31E-18 | 4.40E-18 | Infection | Liver |
| Rad23b | -0.64804 | 9.99421 | 89.97038 | 8.90E-14 | 2.20E-13 | Healthy control | Liver |
| Rad50 | -0.15894 | 5.657484 | 4.202705 | 0.04451 | 0.054213 | Healthy control | Liver |
| Pkia | 2.374774 | 3.431092 | 175.3178 | 3.75E-21 | 1.51E-20 | Infection | Liver |
| Rad51b | 0.711439 | 7.337532 | 41.93814 | 1.62E-08 | 3.03E-08 | Infection | Liver |
| Rad51d | -0.41514 | 6.488245 | 71.06841 | 6.08E-12 | 1.36E-11 | Healthy control | Liver |
| Gnao1 | 2.37151 | 4.176847 | 197.95 | 2.81E-21 | 1.15E-20 | Infection | Liver |
| Rad9a | 1.032916 | 6.359779 | 226.2995 | 1.39E-22 | 6.31E-22 | Infection | Liver |
| Rae1 | -0.00373 | 5.981844 | 0.003125 | 0.955596 | 0.959622 | Healthy control | Liver |
| Raf1 | -0.93737 | 9.748368 | 635.5587 | 9.56E-35 | 1.75E-33 | Healthy control | Liver |
| Rag1 | 1.280844 | 1.411838 | 4.178954 | 0.044466 | 0.054173 | Infection | Liver |
| Rag2 | 1.288422 | 1.50705 | 7.697544 | 0.006571 | 0.008793 | Infection | Liver |
| Rai14 | 1.106907 | 8.045856 | 253.1526 | 8.15E-24 | 4.16E-23 | Infection | Liver |
| Rala | -0.91293 | 7.09707 | 359.3062 | 8.37E-28 | 6.54E-27 | Healthy control | Liver |
| Ramp3 | 1.289711 | 2.955154 | 34.9736 | 1.03E-07 | 1.85E-07 | Infection | Liver |
| Ranbp1 | -0.1794 | 6.014186 | 5.013371 | 0.028674 | 0.035716 | Healthy control | Liver |
| Ranbp2 | 0.726368 | 8.101833 | 197.0629 | 4.05E-21 | 1.63E-20 | Infection | Liver |
| Rangap1 | -0.39987 | 8.124047 | 59.4385 | 1.11E-10 | 2.33E-10 | Healthy control | Liver |
| Rap1a | -0.9182 | 8.160386 | 299.9706 | 1.02E-25 | 6.19E-25 | Healthy control | Liver |
| Rap1gap | -0.60715 | 5.638797 | 37.59842 | 6.27E-08 | 1.14E-07 | Healthy control | Liver |
| Rapgef1 | 0.233626 | 7.646 | 20.34433 | 2.86E-05 | 4.52E-05 | Infection | Liver |
| Rapgef3 | 0.034871 | 4.675903 | 0.112478 | 0.738457 | 0.757834 | Infection | Liver |
| Hoxb8 | 2.370976 | 2.442015 | 100.6735 | 3.31E-17 | 1.00E-16 | Infection | Liver |
| Rara | -1.28791 | 8.109886 | 264.4793 | 2.67E-24 | 1.43E-23 | Healthy control | Liver |
| Rarb | 0.462202 | 5.262189 | 8.512551 | 0.004879 | 0.006587 | Infection | Liver |
| Rarg | 1.506794 | 6.120147 | 215.3371 | 4.79E-22 | 2.08E-21 | Infection | Liver |
| Rasd1 | 0.690261 | 1.604413 | 2.415735 | 0.122553 | 0.141164 | Infection | Liver |
| Stra8 | 2.369325 | 1.447328 | 16.41543 | 0.00013 | 0.000198 | Infection | Liver |
| Rassf5 | -0.47408 | 6.220962 | 34.52503 | 1.70E-07 | 3.04E-07 | Healthy control | Liver |
| Rax | 0.715458 | 1.256188 | 1.417417 | 0.241254 | 0.266605 | Infection | Liver |
| Rb1 | 1.129085 | 6.672016 | 377.5643 | 2.19E-28 | 1.81E-27 | Infection | Liver |
| Rb1cc1 | -0.69026 | 4.714585 | 53.97596 | 5.01E-10 | 1.02E-09 | Healthy control | Liver |
| Rbbp4 | -1.06237 | 9.119658 | 844.348 | 2.39E-38 | 7.98E-37 | Healthy control | Liver |
| Rbbp7 | -0.1812 | 8.026227 | 11.91305 | 0.000998 | 0.001427 | Healthy control | Liver |
| Rbck1 | -0.91271 | 7.018682 | 194.3019 | 5.73E-21 | 2.27E-20 | Healthy control | Liver |
| Rbfox2 | 0.079723 | 6.244788 | 1.116629 | 0.294659 | 0.320736 | Infection | Liver |
| Rbfox3 | 0.030751 | 3.153276 | 0.023304 | 0.87913 | 0.890515 | Infection | Liver |
| Rbl1 | 0.339056 | 3.938633 | 6.94223 | 0.010593 | 0.013885 | Infection | Liver |
| Rbl2 | -0.22475 | 6.320123 | 13.83314 | 0.000426 | 0.000625 | Healthy control | Liver |
| Rbm8a | -1.94971 | 4.086638 | 223.8196 | 2.53E-23 | 1.22E-22 | Healthy control | Liver |
| Rbms1 | 0.853979 | 8.760562 | 300.015 | 1.01E-25 | 6.16E-25 | Infection | Liver |
| Whrn | 2.368977 | 3.874532 | 284.1195 | 1.45E-25 | 8.73E-25 | Infection | Liver |
| Rbp3 | 0.432936 | 1.296344 | 0.672403 | 0.416033 | 0.443592 | Infection | Liver |
| Rbpj | 1.067386 | 7.568384 | 284.237 | 4.15E-25 | 2.40E-24 | Infection | Liver |
| Rbpjl | 0.645577 | 1.252513 | 1.333728 | 0.255655 | 0.281215 | Infection | Liver |
| Rcan1 | -1.22136 | 5.966426 | 292.2498 | 2.19E-25 | 1.29E-24 | Healthy control | Liver |
| Rcc2 | 0.365538 | 6.439562 | 18.78763 | 5.33E-05 | 8.30E-05 | Infection | Liver |
| Rcn2 | 0.714225 | 5.804365 | 64.41886 | 3.11E-11 | 6.71E-11 | Infection | Liver |
| Rcor1 | -0.14802 | 7.306572 | 4.43664 | 0.039141 | 0.048072 | Healthy control | Liver |

| | | | | | | | |
|---|---|---|---|---|---|---|---|
| Cntnap3 | 2.368325 | 1.601035 | 15.88603 | 0.000151 | 0.000228 | Infection | Liver |
| Rdx | -1.27047 | 9.861575 | 706.8233 | 4.39E-36 | 1.02E-34 | Healthy control | Liver |
| Rec114 | 0.890156 | 2.282089 | 8.188957 | 0.005135 | 0.006921 | Infection | Liver |
| Rec8 | 1.509244 | 1.816228 | 16.90339 | 7.00E-05 | 0.000108 | Infection | Liver |
| Rel | 1.100118 | 5.76924 | 295.7831 | 1.53E-25 | 9.17E-25 | Infection | Liver |
| Rela | -0.52489 | 7.448565 | 194.5494 | 5.53E-21 | 2.19E-20 | Healthy control | Liver |
| Irx5 | 2.363775 | 1.999698 | 18.8102 | 3.20E-05 | 5.06E-05 | Infection | Liver |
| Reln | -0.07251 | 8.370576 | 0.515444 | 0.475443 | 0.502866 | Healthy control | Liver |
| Ren1 | -1.49255 | 1.534689 | 11.68463 | 0.0009 | 0.001291 | Healthy control | Liver |
| Rest | 1.078141 | 7.306686 | 264.5491 | 2.66E-24 | 1.42E-23 | Infection | Liver |
| Spdye4a | 2.363219 | 1.390376 | 12.74028 | 0.000702 | 0.001013 | Infection | Liver |
| Retn | 1.21759 | 1.389862 | 5.924084 | 0.017282 | 0.022172 | Infection | Liver |
| Retreg3 | -1.41108 | 8.072814 | 956.4133 | 6.00E-40 | 2.75E-38 | Healthy control | Liver |
| Rev1 | -0.59395 | 5.933346 | 77.34512 | 1.42E-12 | 3.28E-12 | Healthy control | Liver |
| Tubg2 | 2.361343 | 2.449312 | 93.95579 | 1.87E-16 | 5.36E-16 | Infection | Liver |
| Rfx2 | -0.69989 | 2.343987 | 8.31005 | 0.004895 | 0.006607 | Healthy control | Liver |
| Rfx3 | 0.310265 | 4.684556 | 10.69248 | 0.001747 | 0.002446 | Infection | Liver |
| Rfx4 | -3.71615 | 4.918429 | 668.3175 | 1.10E-39 | 4.84E-38 | Healthy control | Liver |
| Rfx5 | -0.00105 | 5.416295 | 0.000203 | 0.988686 | 0.989621 | Healthy control | Liver |
| Lefty2 | 2.355104 | 1.552367 | 16.65491 | 0.000103 | 0.000157 | Infection | Liver |
| Myh11 | 2.354506 | 4.077846 | 86.49092 | 1.57E-13 | 3.85E-13 | Infection | Liver |
| Rgmb | 1.716098 | 5.823565 | 495.2985 | 1.29E-31 | 1.54E-30 | Infection | Liver |
| Rgr | 1.088004 | 1.243018 | 2.735285 | 0.107456 | 0.124578 | Infection | Liver |
| Efcab3 | 2.352155 | 1.325303 | 12.71087 | 0.000802 | 0.001154 | Infection | Liver |
| Rgs3 | -0.14564 | 6.658841 | 2.583456 | 0.112958 | 0.130623 | Healthy control | Liver |
| Rgs6 | 0.876156 | 1.568098 | 3.56326 | 0.061393 | 0.073349 | Infection | Liver |
| Rgs9 | 0.653854 | 3.099479 | 12.45238 | 0.000754 | 0.001087 | Infection | Liver |
| Rhbdf2 | 1.333301 | 6.267972 | 183.7247 | 2.19E-20 | 8.39E-20 | Infection | Liver |
| Rbp1 | 2.351963 | 9.621262 | 649.7549 | 5.05E-35 | 9.85E-34 | Infection | Liver |
| Rheb | -0.2865 | 7.677063 | 26.60934 | 2.66E-06 | 4.44E-06 | Healthy control | Liver |
| Nfe2 | 2.349588 | 4.180096 | 127.8183 | 7.06E-17 | 2.10E-16 | Infection | Liver |
| Rhoa | -0.73547 | 10.08307 | 262.4554 | 3.24E-24 | 1.73E-23 | Healthy control | Liver |
| Rhoc | 1.667315 | 7.728019 | 315.8168 | 2.62E-26 | 1.72E-25 | Infection | Liver |
| Rhod | 0.010203 | 7.825162 | 0.01375 | 0.907024 | 0.915828 | Infection | Liver |
| Rhog | -1.2754 | 6.179698 | 180.9505 | 3.22E-20 | 1.22E-19 | Healthy control | Liver |
| Rhoj | 1.37621 | 5.81156 | 126.8293 | 9.30E-17 | 2.73E-16 | Infection | Liver |
| Rhot2 | -0.77717 | 7.451086 | 251.42 | 9.75E-24 | 4.94E-23 | Healthy control | Liver |
| Rhox13 | 1.260784 | 1.300905 | 4.107117 | 0.047963 | 0.058192 | Infection | Liver |
| Rhox5 | -0.14344 | 1.921907 | 0.168698 | 0.681995 | 0.704513 | Healthy control | Liver |
| Rhpn1 | 1.150754 | 1.575484 | 7.508429 | 0.007106 | 0.009459 | Infection | Liver |
| Rhpn2 | -0.93378 | 4.996422 | 44.52512 | 7.45E-09 | 1.42E-08 | Healthy control | Liver |
| Ric8a | 0.208576 | 6.708003 | 17.0154 | 0.00011 | 0.000168 | Infection | Liver |
| Ric8b | -0.89027 | 6.006248 | 128.1597 | 7.51E-17 | 2.22E-16 | Healthy control | Liver |
| Rictor | 0.433451 | 6.760173 | 59.39582 | 1.12E-10 | 2.36E-10 | Infection | Liver |
| Rif1 | 0.262132 | 6.338907 | 21.2694 | 1.99E-05 | 3.18E-05 | Infection | Liver |
| Kif20b | 2.346473 | 3.571898 | 246.4853 | 1.31E-24 | 7.24E-24 | Infection | Liver |
| Ring1 | -0.34355 | 6.952877 | 38.75337 | 4.30E-08 | 7.88E-08 | Healthy control | Liver |
| Riok3 | -0.61281 | 8.29226 | 249.7767 | 1.15E-23 | 5.73E-23 | Healthy control | Liver |
| Riox1 | -0.55201 | 4.804726 | 31.02754 | 5.64E-07 | 9.78E-07 | Healthy control | Liver |
| Riox2 | -0.55957 | 5.227092 | 55.16383 | 3.56E-10 | 7.30E-10 | Healthy control | Liver |

| | | | | | | | |
|---|---|---|---|---|---|---|---|
| Ripk1 | 0.133225 | 6.228755 | 4.369973 | 0.040596 | 0.049737 | Infection | Liver |
| Ripk2 | 0.428289 | 5.588217 | 21.33979 | 1.94E-05 | 3.11E-05 | Infection | Liver |
| Ripk3 | 1.14676 | 4.221168 | 62.3782 | 5.70E-11 | 1.22E-10 | Infection | Liver |
| Rita1 | 0.288359 | 4.036039 | 5.451024 | 0.022778 | 0.028816 | Infection | Liver |
| Rlim | 0.034558 | 7.497335 | 0.54095 | 0.464753 | 0.4926 | Infection | Liver |
| Rln1 | 0.99848 | 1.379372 | 3.00016 | 0.087298 | 0.102297 | Infection | Liver |
| Rln3 | 0.306761 | 1.451252 | 0.437804 | 0.509512 | 0.536858 | Infection | Liver |
| Rmdn3 | 0.46286 | 8.146355 | 23.67899 | 7.89E-06 | 1.29E-05 | Infection | Liver |
| Ifi209 | 2.346051 | 4.175333 | 180.7913 | 2.93E-20 | 1.11E-19 | Infection | Liver |
| Rnaseh2b | -0.35463 | 5.176598 | 16.87976 | 0.000117 | 0.000178 | Healthy control | Liver |
| Rnaset2a | -0.66579 | 10.92277 | 238.8376 | 3.55E-23 | 1.69E-22 | Healthy control | Liver |
| Rnd2 | -1.0827 | 5.785148 | 108.1017 | 2.60E-15 | 6.96E-15 | Healthy control | Liver |
| Rnf10 | -0.48753 | 9.473756 | 148.7895 | 2.78E-18 | 9.02E-18 | Healthy control | Liver |
| Rnf13 | 0.064931 | 9.891777 | 0.958925 | 0.331181 | 0.357304 | Infection | Liver |
| Rnf130 | -0.94994 | 8.352353 | 451.2113 | 1.60E-30 | 1.66E-29 | Healthy control | Liver |
| Rnf14 | -0.33367 | 8.940338 | 46.18759 | 4.41E-09 | 8.54E-09 | Healthy control | Liver |
| Rnf144a | 0.53486 | 4.863314 | 21.91524 | 1.56E-05 | 2.51E-05 | Infection | Liver |
| Rnf19a | 0.140916 | 6.965035 | 9.414201 | 0.003166 | 0.004345 | Infection | Liver |
| Rnf2 | -0.16563 | 5.98967 | 5.953863 | 0.017497 | 0.022435 | Healthy control | Liver |
| Rnf20 | 0.007336 | 6.167794 | 0.012544 | 0.911176 | 0.919441 | Infection | Liver |
| Rnf219 | 1.231768 | 4.568257 | 100.2881 | 1.25E-14 | 3.22E-14 | Infection | Liver |
| Rnf220 | 0.665861 | 7.073631 | 146.51 | 3.94E-18 | 1.27E-17 | Infection | Liver |
| Rnf31 | -0.24289 | 5.477861 | 10.93277 | 0.001561 | 0.002194 | Healthy control | Liver |
| Rnh1 | -1.11793 | 9.003999 | 409.6552 | 2.32E-29 | 2.12E-28 | Healthy control | Liver |
| Robo1 | 1.522522 | 5.324911 | 191.1947 | 9.09E-21 | 3.56E-20 | Infection | Liver |
| Icosl | 2.34374 | 3.723312 | 169.4634 | 4.58E-20 | 1.72E-19 | Infection | Liver |
| Robo3 | -0.14223 | 1.394527 | 0.097306 | 0.755736 | 0.774456 | Healthy control | Liver |
| Rock1 | -0.37581 | 7.333815 | 67.51756 | 1.43E-11 | 3.14E-11 | Healthy control | Liver |
| Rock2 | 0.919057 | 8.171695 | 430.2629 | 5.98E-30 | 5.83E-29 | Infection | Liver |
| Ror1 | 1.183008 | 4.916123 | 66.37721 | 1.98E-11 | 4.33E-11 | Infection | Liver |
| Phox2a | 2.339461 | 1.451692 | 21.31475 | 1.75E-05 | 2.81E-05 | Infection | Liver |
| Rora | -1.2715 | 6.950831 | 427.3382 | 7.38E-30 | 7.11E-29 | Healthy control | Liver |
| Rorb | 1.80745 | 1.356929 | 8.505784 | 0.004918 | 0.006634 | Infection | Liver |
| Rorc | -1.73638 | 7.097492 | 313.2053 | 3.34E-26 | 2.17E-25 | Healthy control | Liver |
| Ros1 | 0.882163 | 1.365079 | 2.288836 | 0.134798 | 0.154098 | Infection | Liver |
| Rp1l1 | 0.938908 | 1.265149 | 2.542742 | 0.118312 | 0.136531 | Infection | Liver |
| Rp2 | 0.801704 | 6.541647 | 168.0625 | 1.75E-19 | 6.28E-19 | Infection | Liver |
| Rpa1 | 0.293255 | 7.326291 | 19.63683 | 3.79E-05 | 5.96E-05 | Infection | Liver |
| Ms4a1 | 2.338953 | 3.895065 | 106.0413 | 2.17E-15 | 5.84E-15 | Infection | Liver |
| Rph3a | 0.873852 | 1.287331 | 2.014706 | 0.162035 | 0.182815 | Infection | Liver |
| Rph3al | -0.09573 | 4.161231 | 0.744537 | 0.391494 | 0.418942 | Healthy control | Liver |
| Rpn1 | -0.45775 | 8.944077 | 89.0716 | 1.07E-13 | 2.65E-13 | Healthy control | Liver |
| Rprml | 0.529986 | 1.207637 | 0.949505 | 0.340365 | 0.366739 | Infection | Liver |
| Rptor | -1.96337 | 7.017908 | 613.6454 | 2.78E-34 | 4.57E-33 | Healthy control | Liver |
| Rragd | 1.037337 | 4.886728 | 97.32995 | 2.13E-14 | 5.43E-14 | Infection | Liver |
| Rras2 | -0.43977 | 6.646906 | 63.61936 | 3.79E-11 | 8.15E-11 | Healthy control | Liver |
| Rsad1 | -0.71837 | 5.023716 | 34.56426 | 1.70E-07 | 3.04E-07 | Healthy control | Liver |
| Rsbn1 | -0.01499 | 6.12463 | 0.054665 | 0.815891 | 0.831843 | Healthy control | Liver |
| Rsf1 | 0.201493 | 7.371439 | 17.3213 | 9.70E-05 | 0.000149 | Infection | Liver |
| Chek1 | 2.338422 | 3.588714 | 202.529 | 2.36E-22 | 1.05E-21 | Infection | Liver |

| | | | | | | | |
|---|---|---|---|---|---|---|---|
| Rsph10b | 1.852431 | 1.774809 | 25.76958 | 1.46E-06 | 2.47E-06 | Infection | Liver |
| Eya2 | 2.334915 | 2.133337 | 55.37836 | 1.63E-11 | 3.56E-11 | Infection | Liver |
| Rspo1 | 1.662229 | 1.489755 | 13.56308 | 0.000423 | 0.000622 | Infection | Liver |
| Rspo2 | -0.77729 | 1.430887 | 2.091895 | 0.151095 | 0.17136 | Healthy control | Liver |
| Rspo3 | -1.012 | 6.175822 | 73.93437 | 3.12E-12 | 7.05E-12 | Healthy control | Liver |
| Rsrp1 | 1.474418 | 11.61875 | 624.9156 | 1.55E-34 | 2.67E-33 | Infection | Liver |
| Rtbdn | 1.900481 | 1.269528 | 8.464175 | 0.00607 | 0.008147 | Infection | Liver |
| Rtcb | -0.63482 | 7.465967 | 165.4665 | 2.50E-19 | 8.83E-19 | Healthy control | Liver |
| Rtf1 | 0.039831 | 6.828637 | 0.677445 | 0.413555 | 0.441325 | Infection | Liver |
| Rtkn2 | 0.756098 | 1.758358 | 3.813465 | 0.052945 | 0.063728 | Infection | Liver |
| Tet1 | 2.334641 | 3.133268 | 81.06836 | 7.29E-14 | 1.81E-13 | Infection | Liver |
| Rtn3 | 0.049518 | 9.482157 | 1.055129 | 0.308233 | 0.334492 | Infection | Liver |
| Rtn4 | 0.222201 | 9.55688 | 20.98928 | 2.22E-05 | 3.54E-05 | Infection | Liver |
| Rtn4r | 0.576609 | 1.338103 | 1.232873 | 0.270573 | 0.296387 | Infection | Liver |
| Rubcn | -0.21372 | 6.862099 | 22.40847 | 1.28E-05 | 2.07E-05 | Healthy control | Liver |
| Rundc1 | -0.40149 | 5.308761 | 30.18733 | 7.50E-07 | 1.29E-06 | Healthy control | Liver |
| Pcdh10 | 2.331723 | 1.515607 | 23.07566 | 8.07E-06 | 1.32E-05 | Infection | Liver |
| Sp9 | 2.328827 | 1.714872 | 15.86411 | 0.000132 | 0.000201 | Infection | Liver |
| Sntn | 2.326979 | 1.511561 | 14.23285 | 0.000318 | 0.000472 | Infection | Liver |
| Ruvbl1 | -0.719 | 7.485277 | 145.7232 | 4.44E-18 | 1.42E-17 | Healthy control | Liver |
| Ruvbl2 | -0.50938 | 6.681418 | 49.75721 | 1.57E-09 | 3.10E-09 | Healthy control | Liver |
| Rxfp1 | 1.707214 | 1.303238 | 7.555742 | 0.008288 | 0.010968 | Infection | Liver |
| Rxfp2 | 0.42816 | 1.232673 | 0.517907 | 0.477411 | 0.504735 | Infection | Liver |
| Cd248 | 2.326619 | 5.501648 | 138.4153 | 2.12E-17 | 6.49E-17 | Infection | Liver |
| Rxra | -2.22598 | 10.12201 | 1130.929 | 3.96E-42 | 3.10E-40 | Healthy control | Liver |
| Rxrb | -0.51886 | 6.960687 | 92.03422 | 5.82E-14 | 1.45E-13 | Healthy control | Liver |
| Rxrg | -1.70639 | 5.568513 | 272.3288 | 1.46E-24 | 8.05E-24 | Healthy control | Liver |
| Rybp | 0.715423 | 6.774626 | 142.0173 | 7.87E-18 | 2.47E-17 | Infection | Liver |
| Ryk | -0.79159 | 7.034061 | 131.0586 | 4.53E-17 | 1.36E-16 | Healthy control | Liver |
| Ryr1 | 1.220685 | 1.300039 | 4.7502 | 0.033811 | 0.041802 | Infection | Liver |
| Bfsp1 | 2.325301 | 2.298441 | 57.15492 | 1.24E-11 | 2.73E-11 | Infection | Liver |
| Ryr3 | 1.313988 | 1.344558 | 5.29628 | 0.024615 | 0.031015 | Infection | Liver |
| S100a1 | -0.42596 | 9.155369 | 72.18284 | 4.64E-12 | 1.04E-11 | Healthy control | Liver |
| Prrxl1 | 2.322636 | 1.475339 | 14.90242 | 0.00024 | 0.000358 | Infection | Liver |
| S1pr1 | -1.7573 | 8.635799 | 615.237 | 2.45E-34 | 4.05E-33 | Healthy control | Liver |
| S1pr2 | 1.301007 | 6.608609 | 205.6317 | 1.46E-21 | 6.06E-21 | Infection | Liver |
| S1pr3 | 0.516151 | 4.961436 | 15.95163 | 0.000172 | 0.000259 | Infection | Liver |
| S1pr5 | -2.23132 | 5.003757 | 246.7658 | 1.09E-23 | 5.49E-23 | Healthy control | Liver |
| Sox14 | 2.321459 | 1.382177 | 11.29142 | 0.001349 | 0.001906 | Infection | Liver |
| Odam | 2.32117 | 1.407075 | 17.98858 | 7.79E-05 | 0.00012 | Infection | Liver |
| Sacm1l | -0.82585 | 7.712693 | 238.1724 | 3.82E-23 | 1.81E-22 | Healthy control | Liver |
| Sag | 1.082149 | 2.022063 | 15.58066 | 0.000132 | 0.000201 | Infection | Liver |
| Sall1 | -2.4346 | 7.365109 | 343.8372 | 4.70E-27 | 3.29E-26 | Healthy control | Liver |
| Sall3 | 0.236307 | 1.219879 | 0.164044 | 0.688919 | 0.710785 | Infection | Liver |
| Sall4 | 0.54115 | 1.186068 | 1.473645 | 0.242798 | 0.268248 | Infection | Liver |
| Was | 2.32104 | 3.841179 | 202.5736 | 8.79E-22 | 3.73E-21 | Infection | Liver |
| Plagl1 | 2.318301 | 4.184843 | 75.27925 | 2.26E-12 | 5.14E-12 | Infection | Liver |
| Samd8 | -0.74339 | 7.020654 | 170.1438 | 1.32E-19 | 4.77E-19 | Healthy control | Liver |
| Samhd1 | 0.578775 | 8.006054 | 75.38599 | 2.19E-12 | 5.00E-12 | Infection | Liver |
| Sap130 | -0.66431 | 7.688062 | 315.0547 | 2.79E-26 | 1.83E-25 | Healthy control | Liver |

| | | | | | | | |
|---|---|---|---|---|---|---|---|
| Sap18 | -0.59375 | 6.942399 | 115.3845 | 6.63E-16 | 1.84E-15 | Healthy control | Liver |
| Saraf | -0.69437 | 8.960403 | 206.2535 | 1.34E-21 | 5.59E-21 | Healthy control | Liver |
| Sardh | -2.4734 | 10.93366 | 714.872 | 3.15E-36 | 7.66E-35 | Healthy control | Liver |
| Sarm1 | 1.205402 | 2.448614 | 26.80566 | 1.31E-06 | 2.22E-06 | Infection | Liver |
| Sart3 | 0.103688 | 6.447706 | 3.37519 | 0.070876 | 0.083998 | Infection | Liver |
| Sass6 | 0.307354 | 4.254546 | 6.245932 | 0.015079 | 0.019481 | Infection | Liver |
| Satb2 | 1.769218 | 2.76961 | 66.36213 | 2.86E-12 | 6.48E-12 | Infection | Liver |
| Saysd1 | 0.0395 | 4.892148 | 0.153709 | 0.696339 | 0.718292 | Infection | Liver |
| Scamp4 | -0.92567 | 6.822373 | 148.9605 | 2.73E-18 | 8.88E-18 | Healthy control | Liver |
| Scarb1 | -1.97639 | 7.70145 | 632.4867 | 1.12E-34 | 2.00E-33 | Healthy control | Liver |
| Scarb2 | 0.687374 | 10.16327 | 208.1503 | 1.07E-21 | 4.50E-21 | Infection | Liver |
| Scgb1b27 | 1.374126 | 1.451988 | 5.437687 | 0.022074 | 0.027968 | Infection | Liver |
| Scimp | 1.07492 | 3.091685 | 28.22145 | 1.27E-06 | 2.15E-06 | Infection | Liver |
| Robo2 | 2.318014 | 6.335399 | 336.2334 | 5.20E-27 | 3.62E-26 | Infection | Liver |
| Scmh1 | 0.656186 | 5.346315 | 81.59471 | 5.51E-13 | 1.31E-12 | Infection | Liver |
| Scn10a | 1.111159 | 1.3927 | 3.601164 | 0.061471 | 0.073425 | Infection | Liver |
| Scn1a | 1.422109 | 1.278617 | 5.355461 | 0.025246 | 0.031722 | Infection | Liver |
| Prss50 | 2.314004 | 1.431336 | 15.09751 | 0.000233 | 0.000348 | Infection | Liver |
| Scn3b | 1.500155 | 1.328332 | 7.258117 | 0.009268 | 0.012216 | Infection | Liver |
| Scn4a | 1.113806 | 1.32351 | 3.583689 | 0.063162 | 0.075266 | Infection | Liver |
| Pkib | 2.313008 | 5.546384 | 334.7111 | 6.72E-27 | 4.64E-26 | Infection | Liver |
| Sv2c | 2.312401 | 1.633772 | 26.11403 | 1.78E-06 | 3.00E-06 | Infection | Liver |
| Scn8a | 1.749954 | 1.458716 | 9.196428 | 0.003347 | 0.004579 | Infection | Liver |
| Scn9a | 1.519945 | 1.334597 | 6.806971 | 0.011488 | 0.015019 | Infection | Liver |
| Scnn1a | -2.13792 | 6.547465 | 405.8235 | 3.39E-29 | 3.03E-28 | Healthy control | Liver |
| Scnn1b | 0.881909 | 1.226514 | 2.317275 | 0.138376 | 0.157793 | Infection | Liver |
| Scnn1g | 1.456117 | 1.288221 | 5.331269 | 0.025547 | 0.03206 | Infection | Liver |
| Scrib | 0.366718 | 7.453798 | 52.66082 | 6.91E-10 | 1.40E-09 | Infection | Liver |
| Scrt1 | 1.059934 | 1.285489 | 3.520627 | 0.066872 | 0.079365 | Infection | Liver |
| Scrt2 | 0.206303 | 1.265033 | 0.151448 | 0.698967 | 0.720558 | Infection | Liver |
| Hey1 | 2.311163 | 4.03355 | 149.9845 | 1.61E-18 | 5.35E-18 | Infection | Liver |
| Scx | 0.932989 | 1.504596 | 4.757244 | 0.031296 | 0.03883 | Infection | Liver |
| Sdc1 | -0.54554 | 10.34019 | 38.52363 | 4.62E-08 | 8.46E-08 | Healthy control | Liver |
| Sdc2 | -1.44598 | 8.630361 | 442.1218 | 2.82E-30 | 2.88E-29 | Healthy control | Liver |
| Sdcbp | -0.68241 | 8.168548 | 70.18632 | 7.47E-12 | 1.67E-11 | Healthy control | Liver |
| Sdccag8 | 1.090757 | 7.029425 | 332.8162 | 6.52E-27 | 4.51E-26 | Infection | Liver |
| Sdha | -1.59269 | 10.31399 | 557.5795 | 4.09E-33 | 5.75E-32 | Healthy control | Liver |
| Sdhb | -1.61133 | 10.80773 | 650.8036 | 4.82E-35 | 9.42E-34 | Healthy control | Liver |
| Sdhc | -1.13314 | 10.13678 | 210.1672 | 8.47E-22 | 3.60E-21 | Healthy control | Liver |
| Cfap61 | 2.310897 | 1.680617 | 20.94405 | 1.54E-05 | 2.48E-05 | Infection | Liver |
| Celsr3 | 2.309011 | 1.653428 | 29.43276 | 4.77E-07 | 8.30E-07 | Infection | Liver |
| Sdr16c5 | 0.012516 | 1.272492 | 0.000458 | 0.983006 | 0.984777 | Infection | Liver |
| Sec1 | 1.508623 | 1.351104 | 6.752105 | 0.011619 | 0.015175 | Infection | Liver |
| Sec22b | -0.67775 | 8.496779 | 170.7628 | 1.20E-19 | 4.39E-19 | Healthy control | Liver |
| Sec23a | -0.79875 | 7.824127 | 173.8767 | 7.90E-20 | 2.92E-19 | Healthy control | Liver |
| Sec24c | -0.81104 | 8.573757 | 529.466 | 1.78E-32 | 2.30E-31 | Healthy control | Liver |
| Sec61a1 | -1.22652 | 8.796444 | 348.992 | 1.81E-27 | 1.37E-26 | Healthy control | Liver |
| Sel1l | -0.51317 | 9.404111 | 103.2474 | 6.24E-15 | 1.64E-14 | Healthy control | Liver |
| Gbp2 | 2.307026 | 7.186522 | 274.8336 | 1.82E-24 | 9.96E-24 | Infection | Liver |
| Selenof | 1.022738 | 8.910119 | 467.2678 | 6.01E-31 | 6.57E-30 | Infection | Liver |

| | | | | | | | |
|---|---|---|---|---|---|---|---|
| Selenoh | 0.392317 | 4.297841 | 8.042343 | 0.006146 | 0.008241 | Infection | Liver |
| Selenoi | -1.23715 | 7.026714 | 250.9765 | 1.03E-23 | 5.20E-23 | Healthy control | Liver |
| Selenok | 0.402041 | 2.120554 | 1.94859 | 0.1656 | 0.186626 | Infection | Liver |
| Selenon | 1.340994 | 5.28827 | 95.5125 | 2.98E-14 | 7.55E-14 | Infection | Liver |
| Selenoo | -1.62289 | 8.735616 | 482.236 | 2.49E-31 | 2.85E-30 | Healthy control | Liver |
| Selenos | -1.05756 | 7.309317 | 350.813 | 1.59E-27 | 1.21E-26 | Healthy control | Liver |
| Selenov | -0.89548 | 1.196663 | 3.882118 | 0.063417 | 0.075534 | Healthy control | Liver |
| Prkg2 | 2.306285 | 1.877154 | 37.71793 | 1.46E-08 | 2.74E-08 | Infection | Liver |
| Kcnj2 | 2.305365 | 4.236041 | 160.1707 | 4.46E-19 | 1.54E-18 | Infection | Liver |
| Dbn1 | 2.304989 | 6.543027 | 288.3898 | 3.84E-25 | 2.23E-24 | Infection | Liver |
| Sdk2 | 2.304637 | 4.561997 | 103.2361 | 7.16E-15 | 1.87E-14 | Infection | Liver |
| Casp4 | 2.304536 | 4.737282 | 370.59 | 5.42E-28 | 4.32E-27 | Infection | Liver |
| Mixl1 | 2.304159 | 1.374697 | 14.22159 | 0.000374 | 0.000552 | Infection | Liver |
| Sema3d | 0.703712 | 6.69711 | 60.53744 | 8.34E-11 | 1.76E-10 | Infection | Liver |
| Sema3e | 1.806486 | 1.413616 | 13.89328 | 0.000389 | 0.000573 | Infection | Liver |
| Sema3f | 0.21181 | 5.924998 | 8.626573 | 0.004615 | 0.006244 | Infection | Liver |
| Sema4a | -1.51625 | 7.652364 | 338.1997 | 4.25E-27 | 3.00E-26 | Healthy control | Liver |
| Camk2d | 2.303736 | 6.808744 | 847.7353 | 2.22E-38 | 7.52E-37 | Infection | Liver |
| Sema4g | -1.37924 | 10.36098 | 271.9356 | 1.30E-24 | 7.22E-24 | Healthy control | Liver |
| Apba1 | 2.299069 | 3.466232 | 129.3776 | 1.06E-17 | 3.32E-17 | Infection | Liver |
| Sema5b | -1.15133 | 2.97095 | 33.27424 | 1.52E-07 | 2.73E-07 | Healthy control | Liver |
| Sema6a | -1.35724 | 6.070569 | 194.0406 | 6.28E-21 | 2.48E-20 | Healthy control | Liver |
| Sema6c | 0.743442 | 4.930715 | 24.56924 | 5.70E-06 | 9.39E-06 | Infection | Liver |
| Sema6d | -0.26062 | 7.003067 | 12.80914 | 0.000668 | 0.000965 | Healthy control | Liver |
| Senp3 | 0.133296 | 5.6639 | 3.53351 | 0.064746 | 0.077025 | Infection | Liver |
| 10-Sep | -1.32698 | 5.839056 | 292.977 | 2.11E-25 | 1.25E-24 | Healthy control | Liver |
| 11-Sep | 1.451527 | 9.108461 | 324.7087 | 1.25E-26 | 8.42E-26 | Infection | Liver |
| 2-Sep | 0.794638 | 9.308972 | 554.6105 | 4.76E-33 | 6.62E-32 | Infection | Liver |
| Cidea | 2.29797 | 1.869322 | 36.08615 | 2.63E-08 | 4.88E-08 | Infection | Liver |
| 4-Sep | 0.890716 | 8.244188 | 122.0045 | 2.07E-16 | 5.91E-16 | Infection | Liver |
| Gramd1b | 2.297401 | 6.115989 | 410.0869 | 2.46E-29 | 2.25E-28 | Infection | Liver |
| Kcnab2 | 2.29735 | 4.327872 | 178.1013 | 5.07E-20 | 1.90E-19 | Infection | Liver |
| 7-Sep | 0.221272 | 8.183898 | 26.631 | 2.64E-06 | 4.41E-06 | Infection | Liver |
| Hfm1 | 2.294389 | 2.134418 | 25.29662 | 1.82E-06 | 3.07E-06 | Infection | Liver |
| 9-Sep | -0.11531 | 8.862231 | 3.748027 | 0.057332 | 0.068744 | Healthy control | Liver |
| Pcgf2 | 2.289804 | 5.308928 | 252.2685 | 1.09E-23 | 5.49E-23 | Infection | Liver |
| Serpina3n | -0.3594 | 11.50017 | 34.11522 | 1.94E-07 | 3.47E-07 | Healthy control | Liver |
| Serpinb13 | 1.271535 | 1.221294 | 5.664551 | 0.024739 | 0.031148 | Infection | Liver |
| Tbx5 | 2.288908 | 1.326547 | 12.28746 | 0.000931 | 0.001334 | Infection | Liver |
| Serpinb5 | 1.974799 | 1.618071 | 15.42817 | 0.000161 | 0.000243 | Infection | Liver |
| Serpinc1 | -1.62176 | 13.34444 | 275.639 | 9.17E-25 | 5.16E-24 | Healthy control | Liver |
| Csf2rb | 2.288849 | 6.305973 | 673.3171 | 1.97E-35 | 4.12E-34 | Infection | Liver |
| Hpse2 | 2.283963 | 1.336521 | 11.42155 | 0.001344 | 0.001899 | Infection | Liver |
| Serpinf2 | -0.71819 | 11.8764 | 54.55269 | 4.10E-10 | 8.37E-10 | Healthy control | Liver |
| Cd33 | 2.28206 | 3.555455 | 181.7383 | 4.08E-21 | 1.64E-20 | Infection | Liver |
| Rfx6 | 2.28164 | 1.302086 | 10.86762 | 0.002023 | 0.002818 | Infection | Liver |
| Sesn1 | 0.233696 | 6.585268 | 8.80371 | 0.004237 | 0.005744 | Infection | Liver |
| Sesn2 | 0.78135 | 6.959597 | 123.6106 | 1.57E-16 | 4.55E-16 | Infection | Liver |
| Set | -0.10879 | 7.161631 | 2.717847 | 0.104181 | 0.121061 | Healthy control | Liver |
| Setbp1 | 1.396037 | 5.001049 | 116.5627 | 5.81E-16 | 1.62E-15 | Infection | Liver |

| | | | | | | | |
|---|---|---|---|---|---|---|---|
| Setd1a | -0.14159 | 7.979243 | 12.16517 | 0.000891 | 0.001278 | Healthy control | Liver |
| Setd1b | -0.77022 | 7.796459 | 240.5405 | 2.98E-23 | 1.43E-22 | Healthy control | Liver |
| Setd5 | 0.732444 | 7.740578 | 272.7907 | 1.20E-24 | 6.71E-24 | Infection | Liver |
| Setd6 | -0.17795 | 7.278507 | 3.936495 | 0.051584 | 0.062209 | Healthy control | Liver |
| Setdb1 | 0.195758 | 6.37634 | 12.81136 | 0.000668 | 0.000965 | Infection | Liver |
| Setx | 0.210162 | 6.582937 | 11.53088 | 0.001186 | 0.001686 | Infection | Liver |
| Sez6 | 0.765127 | 1.222284 | 1.65032 | 0.209815 | 0.233355 | Infection | Liver |
| Sez6l2 | 0.70051 | 2.864946 | 12.03524 | 0.000886 | 0.001272 | Infection | Liver |
| Sf1 | -0.34496 | 9.433814 | 81.79569 | 5.15E-13 | 1.23E-12 | Healthy control | Liver |
| Sf3a1 | -0.34856 | 7.321171 | 41.72142 | 1.70E-08 | 3.18E-08 | Healthy control | Liver |
| Sf3b6 | -1.04281 | 7.208584 | 255.6296 | 6.41E-24 | 3.32E-23 | Healthy control | Liver |
| Sfpq | 0.883277 | 9.306411 | 246.409 | 1.61E-23 | 7.95E-23 | Infection | Liver |
| Nyap1 | 2.281206 | 3.032913 | 131.2292 | 7.31E-19 | 2.48E-18 | Infection | Liver |
| Sfrp2 | 1.514102 | 1.243555 | 7.027168 | 0.011955 | 0.015589 | Infection | Liver |
| Sfrp4 | 1.998895 | 1.487627 | 14.96792 | 0.000221 | 0.00033 | Infection | Liver |
| Sfxn1 | -1.1171 | 8.24976 | 51.44071 | 1.34E-09 | 2.66E-09 | Healthy control | Liver |
| Sfxn3 | 1.298774 | 4.687482 | 73.41005 | 3.74E-12 | 8.44E-12 | Infection | Liver |
| Sgca | 1.882157 | 1.343878 | 11.54284 | 0.001327 | 0.001878 | Infection | Liver |
| Cacna2d2 | 2.278414 | 1.735008 | 27.92224 | 7.53E-07 | 1.30E-06 | Infection | Liver |
| Sgms2 | -0.91645 | 7.712531 | 100.3041 | 1.11E-14 | 2.87E-14 | Healthy control | Liver |
| App | 2.276587 | 10.47114 | 1247.148 | 2.08E-43 | 2.00E-41 | Infection | Liver |
| Map3k15 | 2.273238 | 3.488273 | 170.5048 | 1.66E-20 | 6.42E-20 | Infection | Liver |
| Sgpl1 | -0.2333 | 6.81776 | 16.26158 | 0.000151 | 0.000228 | Healthy control | Liver |
| Sgsh | 0.272457 | 5.661469 | 14.04952 | 0.000388 | 0.000572 | Infection | Liver |
| Sgta | -0.50171 | 8.183653 | 118.4654 | 3.82E-16 | 1.08E-15 | Healthy control | Liver |
| Zfpm2 | 2.271356 | 4.616261 | 228.5052 | 1.33E-22 | 6.09E-22 | Infection | Liver |
| Sh3bp1 | 1.579947 | 6.564992 | 210.6604 | 8.11E-22 | 3.45E-21 | Infection | Liver |
| Sh3bp2 | -0.29081 | 7.172549 | 30.67936 | 6.26E-07 | 1.08E-06 | Healthy control | Liver |
| Sh3bp5 | -1.49556 | 8.535095 | 912.0757 | 2.45E-39 | 9.88E-38 | Healthy control | Liver |
| Sh3gl1 | 0.847033 | 7.142852 | 224.0708 | 1.76E-22 | 7.91E-22 | Infection | Liver |
| Ercc1 | 2.268449 | 4.654493 | 149.6797 | 2.71E-18 | 8.79E-18 | Infection | Liver |
| Batf | 2.265757 | 4.500271 | 301.3014 | 1.25E-25 | 7.54E-25 | Infection | Liver |
| Sh3rf2 | 0.616492 | 1.276406 | 0.97962 | 0.327731 | 0.353964 | Infection | Liver |
| Sh3tc2 | -0.34534 | 2.902113 | 3.692197 | 0.058606 | 0.070136 | Healthy control | Liver |
| Shank3 | 0.298809 | 5.263666 | 10.57364 | 0.001842 | 0.002577 | Infection | Liver |
| Sharpin | -1.14435 | 7.315613 | 433.8124 | 4.82E-30 | 4.74E-29 | Healthy control | Liver |
| Shc1 | 0.0794 | 8.156653 | 3.405556 | 0.069649 | 0.082563 | Infection | Liver |
| Vstm2a | 2.264413 | 1.383494 | 9.353353 | 0.003269 | 0.004478 | Infection | Liver |
| She | 0.108441 | 6.205976 | 2.446519 | 0.122769 | 0.14138 | Infection | Liver |
| Shh | -0.37475 | 1.420334 | 0.504635 | 0.479015 | 0.506323 | Healthy control | Liver |
| Cd72 | 2.26134 | 7.3698 | 761.8914 | 5.00E-37 | 1.39E-35 | Infection | Liver |
| Shld1 | -0.76249 | 3.492266 | 25.6039 | 3.51E-06 | 5.84E-06 | Healthy control | Liver |
| Shmt1 | -1.54527 | 10.59309 | 196.3321 | 5.37E-21 | 2.14E-20 | Healthy control | Liver |
| Shoc2 | -1.16395 | 7.270825 | 682.8665 | 1.22E-35 | 2.63E-34 | Healthy control | Liver |
| Shox2 | 0.766005 | 1.260444 | 1.280553 | 0.265014 | 0.29087 | Infection | Liver |
| Shroom3 | -0.10985 | 6.211901 | 1.698109 | 0.197254 | 0.220021 | Healthy control | Liver |
| Shtn1 | -0.28296 | 6.743929 | 18.77205 | 5.36E-05 | 8.34E-05 | Healthy control | Liver |
| Siah2 | -1.65492 | 7.226363 | 671.1537 | 2.04E-35 | 4.25E-34 | Healthy control | Liver |
| Tubb1 | 2.259181 | 2.549675 | 40.17351 | 8.56E-09 | 1.63E-08 | Infection | Liver |
| Sidt2 | 0.360505 | 7.348361 | 28.49671 | 1.35E-06 | 2.29E-06 | Infection | Liver |

| | | | | | | | |
|---|---|---|---|---|---|---|---|
| Siglec1 | -0.30066 | 6.551076 | 4.011589 | 0.049578 | 0.060035 | Healthy control | Liver |
| Sigmar1 | -0.43587 | 9.311457 | 21.63753 | 1.72E-05 | 2.77E-05 | Healthy control | Liver |
| Foxg1 | 2.257551 | 1.422443 | 16.16397 | 0.000147 | 0.000222 | Infection | Liver |
| Sin3a | -0.20832 | 7.2917 | 20.60129 | 2.58E-05 | 4.10E-05 | Healthy control | Liver |
| Sin3b | 0.653792 | 6.210588 | 105.5165 | 4.08E-15 | 1.08E-14 | Infection | Liver |
| Sinhcaf | 1.709167 | 4.068735 | 152.8005 | 1.92E-18 | 6.34E-18 | Infection | Liver |
| Sirpa | 1.452926 | 8.474961 | 511.7384 | 4.68E-32 | 5.75E-31 | Infection | Liver |
| Sirt1 | 0.211121 | 5.6288 | 7.170292 | 0.009431 | 0.012424 | Infection | Liver |
| Sirt2 | -0.56764 | 7.897725 | 138.952 | 1.27E-17 | 3.95E-17 | Healthy control | Liver |
| Sirt3 | -1.56715 | 8.062403 | 438.5665 | 3.54E-30 | 3.59E-29 | Healthy control | Liver |
| Sirt4 | -0.98508 | 5.620425 | 147.493 | 3.61E-18 | 1.16E-17 | Healthy control | Liver |
| Sirt5 | -0.7143 | 6.05699 | 130.6883 | 4.89E-17 | 1.46E-16 | Healthy control | Liver |
| Sirt6 | -0.43185 | 5.08587 | 17.96517 | 7.48E-05 | 0.000116 | Healthy control | Liver |
| Sirt7 | -0.91487 | 7.893506 | 424.6193 | 8.65E-30 | 8.23E-29 | Healthy control | Liver |
| Six1 | 1.907096 | 1.683867 | 11.7896 | 0.000847 | 0.001218 | Infection | Liver |
| Six2 | 0.087445 | 1.277351 | 0.022932 | 0.880257 | 0.890935 | Infection | Liver |
| Six3 | 0.756464 | 1.278047 | 1.729526 | 0.195056 | 0.217666 | Infection | Liver |
| Foxb1 | 2.254121 | 1.324848 | 13.98278 | 0.000446 | 0.000653 | Infection | Liver |
| Fasl | 2.251513 | 3.035841 | 97.74955 | 1.07E-15 | 2.94E-15 | Infection | Liver |
| Ska2 | 1.081373 | 5.297289 | 156.9697 | 8.80E-19 | 2.97E-18 | Infection | Liver |
| Skp1a | -0.23761 | 7.766547 | 9.169858 | 0.003556 | 0.004854 | Healthy control | Liver |
| Skp2 | 0.711684 | 3.850789 | 36.30667 | 9.82E-08 | 1.78E-07 | Infection | Liver |
| Spsb1 | 2.250481 | 5.421203 | 149.2626 | 3.38E-18 | 1.09E-17 | Infection | Liver |
| Pou4f3 | 2.247992 | 1.54081 | 16.89854 | 9.29E-05 | 0.000143 | Infection | Liver |
| Slc11a1 | 0.210164 | 6.351208 | 3.798862 | 0.055719 | 0.066905 | Infection | Liver |
| Slc11a2 | -0.3693 | 6.74945 | 46.84362 | 3.64E-09 | 7.08E-09 | Healthy control | Liver |
| Slc12a2 | 0.479192 | 5.409916 | 30.75785 | 6.12E-07 | 1.06E-06 | Infection | Liver |
| Slc12a5 | -0.89032 | 2.884508 | 26.18492 | 2.20E-06 | 3.70E-06 | Healthy control | Liver |
| Slc12a7 | -0.58265 | 7.650028 | 75.37191 | 2.20E-12 | 5.01E-12 | Healthy control | Liver |
| Slc13a4 | -0.44953 | 1.296781 | 0.595572 | 0.443512 | 0.471384 | Healthy control | Liver |
| Fgf10 | 2.245657 | 1.674095 | 29.06608 | 5.94E-07 | 1.03E-06 | Infection | Liver |
| Slc15a1 | 0.642777 | 1.409731 | 1.328379 | 0.252024 | 0.277648 | Infection | Liver |
| Slc16a1 | -0.27228 | 8.41811 | 16.52124 | 0.000135 | 0.000205 | Healthy control | Liver |
| Slc16a2 | -1.35474 | 8.463817 | 381.3826 | 1.64E-28 | 1.37E-27 | Healthy control | Liver |
| Fbxo41 | 2.240458 | 1.377818 | 11.69855 | 0.001137 | 0.001622 | Infection | Liver |
| Slc16a8 | 0.67695 | 1.253551 | 1.514738 | 0.225756 | 0.250306 | Infection | Liver |
| Slc17a2 | -1.44728 | 7.983842 | 188.7065 | 1.23E-20 | 4.77E-20 | Healthy control | Liver |
| Il22 | 2.240375 | 1.704728 | 15.6916 | 0.000143 | 0.000217 | Infection | Liver |
| Slc17a7 | 1.913475 | 1.468479 | 15.75207 | 0.000157 | 0.000237 | Infection | Liver |
| Slc17a8 | -1.87371 | 5.369839 | 212.5078 | 7.10E-22 | 3.06E-21 | Healthy control | Liver |
| Kcna4 | 2.239868 | 1.356797 | 16.38748 | 0.000151 | 0.000228 | Infection | Liver |
| Slc18a3 | 0.641292 | 1.185308 | 2.22408 | 0.157548 | 0.178154 | Infection | Liver |
| Slc19a1 | 0.229235 | 5.319144 | 7.269946 | 0.008976 | 0.01184 | Infection | Liver |
| Slc19a2 | -1.24057 | 8.953974 | 280.8547 | 5.65E-25 | 3.23E-24 | Healthy control | Liver |
| Slc19a3 | -0.00267 | 1.355439 | 8.14E-05 | 0.992825 | 0.993222 | Healthy control | Liver |
| Sh3gl3 | 2.238289 | 1.664706 | 30.83766 | 2.43E-07 | 4.31E-07 | Infection | Liver |
| Plcg2 | 2.237874 | 5.05015 | 304.7239 | 9.28E-26 | 5.68E-25 | Infection | Liver |
| Slc20a2 | -1.88125 | 7.513836 | 744.148 | 1.00E-36 | 2.69E-35 | Healthy control | Liver |
| Slc22a18 | -0.90549 | 7.54949 | 91.55823 | 6.41E-14 | 1.60E-13 | Healthy control | Liver |
| Slc22a2 | 1.167718 | 1.289474 | 3.403208 | 0.071155 | 0.084288 | Infection | Liver |

| | | | | | | | |
|---|---|---|---|---|---|---|---|
| Slc22a5 | -2.23359 | 5.526759 | 584.6712 | 1.26E-33 | 1.88E-32 | Healthy control | Liver |
| Slc22a6 | 0.351951 | 1.266275 | 0.312171 | 0.579267 | 0.60501 | Infection | Liver |
| Slc22a8 | 0.587209 | 1.354228 | 1.123136 | 0.292605 | 0.318707 | Infection | Liver |
| Slc24a3 | 1.668987 | 3.545559 | 99.72676 | 9.68E-15 | 2.52E-14 | Infection | Liver |
| Slc24a4 | 0.797357 | 1.311195 | 1.969182 | 0.165837 | 0.186852 | Infection | Liver |
| Slc25a17 | -0.31842 | 7.625547 | 17.69323 | 8.32E-05 | 0.000128 | Healthy control | Liver |
| Slc25a20 | -1.20531 | 9.683446 | 277.7464 | 7.53E-25 | 4.27E-24 | Healthy control | Liver |
| Slc25a22 | -1.36399 | 9.856272 | 125.9964 | 1.41E-16 | 4.09E-16 | Healthy control | Liver |
| Slc25a28 | -0.10679 | 7.170962 | 5.373788 | 0.023682 | 0.029914 | Healthy control | Liver |
| Slc25a31 | 0.26029 | 1.527078 | 0.328416 | 0.567628 | 0.593844 | Infection | Liver |
| Slc25a51 | -1.02742 | 6.256265 | 92.13102 | 5.79E-14 | 1.45E-13 | Healthy control | Liver |
| Slc26a3 | 1.034691 | 1.282445 | 3.273763 | 0.076568 | 0.090295 | Infection | Liver |
| Slc26a4 | -3.22351 | 3.163178 | 152.244 | 2.71E-21 | 1.11E-20 | Healthy control | Liver |
| Slc26a5 | 0.768719 | 1.829301 | 4.042272 | 0.046378 | 0.056379 | Infection | Liver |
| Slc2a1 | 1.319567 | 6.167281 | 68.07086 | 1.46E-11 | 3.20E-11 | Infection | Liver |
| Slc2a2 | -1.18091 | 9.812636 | 87.49721 | 1.91E-13 | 4.66E-13 | Healthy control | Liver |
| Bcl2 | 2.237615 | 5.280587 | 525.9341 | 3.02E-32 | 3.80E-31 | Infection | Liver |
| Prl8a2 | 2.236513 | 1.558398 | 10.42534 | 0.00181 | 0.002533 | Infection | Liver |
| Slc2a5 | -2.13751 | 4.615841 | 121.8875 | 1.22E-16 | 3.55E-16 | Healthy control | Liver |
| Slc30a10 | -1.18403 | 8.404898 | 71.39451 | 8.17E-12 | 1.82E-11 | Healthy control | Liver |
| Slc30a3 | 0.578948 | 2.529609 | 5.229661 | 0.024705 | 0.031112 | Infection | Liver |
| Slc30a8 | -0.33215 | 1.255038 | 0.331176 | 0.568198 | 0.594316 | Healthy control | Liver |
| Slc31a1 | -1.55559 | 7.257593 | 466.4097 | 6.46E-31 | 7.04E-30 | Healthy control | Liver |
| Slc32a1 | 0.97848 | 1.312828 | 2.47607 | 0.121065 | 0.139579 | Infection | Liver |
| Slc39a5 | -0.97612 | 2.609038 | 18.62284 | 4.26E-05 | 6.67E-05 | Healthy control | Liver |
| Slc39a8 | -1.91629 | 5.447161 | 585.9947 | 1.28E-33 | 1.89E-32 | Healthy control | Liver |
| Slc3a1 | 0.528583 | 7.321898 | 5.642105 | 0.020851 | 0.026539 | Infection | Liver |
| Slc40a1 | -0.95549 | 9.703232 | 86.97923 | 1.73E-13 | 4.22E-13 | Healthy control | Liver |
| Slc44a5 | 1.658121 | 1.34565 | 7.813358 | 0.007073 | 0.009417 | Infection | Liver |
| Slc45a1 | 0.791782 | 1.197302 | 2.874377 | 0.106162 | 0.123192 | Infection | Liver |
| Slc45a2 | -0.83621 | 1.430967 | 2.898929 | 0.091459 | 0.106948 | Healthy control | Liver |
| Slc47a1 | -1.70091 | 9.678555 | 303.1503 | 7.68E-26 | 4.76E-25 | Healthy control | Liver |
| Inha | 2.233987 | 2.544987 | 87.16574 | 1.73E-15 | 4.70E-15 | Infection | Liver |
| Slc4a10 | 1.284737 | 1.533688 | 7.246242 | 0.008358 | 0.011054 | Infection | Liver |
| Slc4a7 | 0.49414 | 6.077242 | 43.45896 | 1.00E-08 | 1.90E-08 | Infection | Liver |
| Slc4a9 | 0.467485 | 2.654582 | 3.619609 | 0.060751 | 0.072669 | Infection | Liver |
| Slc52a3 | 1.154619 | 2.651919 | 25.42679 | 2.73E-06 | 4.57E-06 | Infection | Liver |
| Slc5a12 | 1.623714 | 1.334062 | 6.717113 | 0.012153 | 0.015831 | Infection | Liver |
| Slc5a2 | 1.378912 | 1.279074 | 4.528719 | 0.038694 | 0.047546 | Infection | Liver |
| Slc5a7 | 1.788723 | 1.412971 | 9.907668 | 0.002418 | 0.003351 | Infection | Liver |
| Slc6a11 | -0.26069 | 1.196132 | 0.257954 | 0.617899 | 0.642143 | Healthy control | Liver |
| Slc6a14 | 1.006903 | 1.301895 | 2.784788 | 0.10119 | 0.117777 | Infection | Liver |
| Slc6a19 | 1.350917 | 1.486521 | 6.713168 | 0.011049 | 0.014464 | Infection | Liver |
| Slc6a3 | 1.697283 | 1.311909 | 7.071787 | 0.010296 | 0.013518 | Infection | Liver |
| Slc6a4 | 1.117773 | 1.97255 | 10.3368 | 0.001665 | 0.002337 | Infection | Liver |
| Slc6a5 | 0.259308 | 1.221506 | 0.193495 | 0.663824 | 0.686734 | Infection | Liver |
| Grin2c | 2.23238 | 2.385258 | 59.64765 | 6.40E-12 | 1.43E-11 | Infection | Liver |
| Slc6a9 | -1.00811 | 5.915283 | 87.86902 | 1.42E-13 | 3.48E-13 | Healthy control | Liver |
| Hoxb3 | 2.23103 | 5.288648 | 239.1897 | 4.20E-23 | 1.98E-22 | Infection | Liver |
| Dlg4 | 2.226431 | 5.730821 | 211.7658 | 7.74E-22 | 3.31E-21 | Infection | Liver |

| | | | | | | | |
|---|---|---|---|---|---|---|---|
| Slc8a3 | -0.55276 | 1.665825 | 1.965691 | 0.163456 | 0.184376 | Healthy control | Liver |
| Slc8b1 | -0.64223 | 8.298035 | 78.8591 | 9.92E-13 | 2.31E-12 | Healthy control | Liver |
| Slc9a1 | 1.016566 | 5.837226 | 151.457 | 1.91E-18 | 6.30E-18 | Infection | Liver |
| Slc9a3 | 1.899178 | 1.325669 | 8.971972 | 0.004119 | 0.005595 | Infection | Liver |
| Slc9a3r1 | -2.15984 | 8.442336 | 1012.396 | 1.10E-40 | 6.34E-39 | Healthy control | Liver |
| Slc9a4 | 0.582338 | 1.21152 | 0.992614 | 0.329861 | 0.356038 | Infection | Liver |
| Slc9a8 | -0.90901 | 6.623796 | 229.3461 | 9.97E-23 | 4.60E-22 | Healthy control | Liver |
| Slc9a9 | 1.055576 | 6.432838 | 164.9819 | 2.69E-19 | 9.50E-19 | Infection | Liver |
| Slco1a1 | -3.40602 | 6.891489 | 452.7813 | 4.26E-30 | 4.27E-29 | Healthy control | Liver |
| Slco3a1 | 1.121839 | 5.490019 | 192.2592 | 7.68E-21 | 3.02E-20 | Infection | Liver |
| Cavin1 | 2.22578 | 6.986152 | 250.8235 | 1.82E-23 | 8.89E-23 | Infection | Liver |
| Slit1 | 0.332057 | 1.275961 | 0.276185 | 0.601654 | 0.626889 | Infection | Liver |
| Wls | 2.223496 | 6.611144 | 359.9297 | 8.16E-28 | 6.39E-27 | Infection | Liver |
| Pou1f1 | 2.221458 | 1.3574 | 10.62857 | 0.001958 | 0.002732 | Infection | Liver |
| Slitrk1 | -0.13411 | 1.268165 | 0.04859 | 0.82654 | 0.841501 | Healthy control | Liver |
| Slitrk3 | 0.566738 | 1.18524 | 1.900332 | 0.189261 | 0.211672 | Infection | Liver |
| Slk | -0.05243 | 9.18022 | 2.863164 | 0.09554 | 0.111485 | Healthy control | Liver |
| Slurp2 | 0.852347 | 1.298258 | 2.289263 | 0.136151 | 0.155539 | Infection | Liver |
| Smad1 | -0.19948 | 7.260683 | 11.58725 | 0.001156 | 0.001647 | Healthy control | Liver |
| Smad2 | -0.2949 | 7.562848 | 40.51422 | 2.47E-08 | 4.58E-08 | Healthy control | Liver |
| Smad3 | 1.059597 | 6.947917 | 265.9762 | 2.32E-24 | 1.25E-23 | Infection | Liver |
| Smad4 | 0.115605 | 7.64413 | 9.434199 | 0.003136 | 0.004305 | Infection | Liver |
| Smad5 | -0.27267 | 6.306701 | 10.59631 | 0.001821 | 0.002548 | Healthy control | Liver |
| Smad6 | -0.81193 | 5.613459 | 68.34068 | 1.20E-11 | 2.64E-11 | Healthy control | Liver |
| Smad7 | 0.104633 | 7.593059 | 3.090901 | 0.083559 | 0.098146 | Infection | Liver |
| Smad9 | -2.24571 | 3.326188 | 133.312 | 6.01E-19 | 2.05E-18 | Healthy control | Liver |
| Smap1 | -0.52285 | 6.830807 | 80.1061 | 7.52E-13 | 1.77E-12 | Healthy control | Liver |
| Smap2 | 0.228723 | 7.55868 | 29.43771 | 9.66E-07 | 1.65E-06 | Infection | Liver |
| Hk1 | 2.220125 | 5.711002 | 268.7821 | 1.97E-24 | 1.08E-23 | Infection | Liver |
| Smarca2 | -0.45342 | 8.784953 | 147.4149 | 3.42E-18 | 1.10E-17 | Healthy control | Liver |
| Smarca4 | -0.37555 | 8.530532 | 67.34441 | 1.49E-11 | 3.27E-11 | Healthy control | Liver |
| Smarca5 | -0.49222 | 6.619833 | 66.66634 | 1.77E-11 | 3.87E-11 | Healthy control | Liver |
| Smarcb1 | -0.00695 | 7.062036 | 0.012976 | 0.909669 | 0.918122 | Healthy control | Liver |
| Smarcc1 | -0.37157 | 6.902571 | 39.3512 | 3.56E-08 | 6.54E-08 | Healthy control | Liver |
| Smarcc2 | 0.042531 | 8.452535 | 1.32735 | 0.253604 | 0.279204 | Infection | Liver |
| Smarcd1 | 0.812928 | 5.931051 | 125.6146 | 1.13E-16 | 3.31E-16 | Infection | Liver |
| Smarcd2 | -0.77257 | 7.614275 | 253.5381 | 7.86E-24 | 4.02E-23 | Healthy control | Liver |
| Smarcd3 | 1.680741 | 4.861708 | 150.8894 | 2.37E-18 | 7.74E-18 | Infection | Liver |
| Smarce1 | -0.42024 | 6.663419 | 43.10722 | 1.11E-08 | 2.11E-08 | Healthy control | Liver |
| Smc1a | 0.550247 | 8.45986 | 122.6628 | 1.84E-16 | 5.30E-16 | Infection | Liver |
| Smc3 | -0.13163 | 6.801297 | 7.085811 | 0.009835 | 0.012929 | Healthy control | Liver |
| Smc5 | -0.61502 | 6.932728 | 135.5612 | 2.18E-17 | 6.67E-17 | Healthy control | Liver |
| Smcp | 0.739286 | 1.286116 | 1.94189 | 0.169947 | 0.191181 | Infection | Liver |
| Smdt1 | -1.61046 | 9.224119 | 1001.744 | 1.50E-40 | 8.16E-39 | Healthy control | Liver |
| Smg9 | -0.27537 | 6.102064 | 13.16882 | 0.00057 | 0.000827 | Healthy control | Liver |
| Lurap1 | 2.219985 | 2.454908 | 99.21315 | 5.40E-17 | 1.61E-16 | Infection | Liver |
| Smim7 | 0.451469 | 7.677434 | 55.55486 | 3.12E-10 | 6.42E-10 | Infection | Liver |
| Smn1 | -0.41132 | 5.787613 | 33.08054 | 2.77E-07 | 4.88E-07 | Healthy control | Liver |
| Smo | 0.975259 | 5.921833 | 54.18779 | 4.56E-10 | 9.29E-10 | Infection | Liver |
| Smoc1 | -1.37163 | 8.922402 | 119.7382 | 4.32E-16 | 1.21E-15 | Healthy control | Liver |

| | | | | | | | |
|---|---|---|---|---|---|---|---|
| Ephb2 | 2.219282 | 3.018653 | 73.15707 | 4.20E-13 | 1.00E-12 | Infection | Liver |
| Smpd1 | -1.71265 | 7.268628 | 679.0102 | 1.45E-35 | 3.11E-34 | Healthy control | Liver |
| Pitpnm1 | 2.219048 | 6.094571 | 691.4487 | 9.29E-36 | 2.03E-34 | Infection | Liver |
| Smpd4 | -0.83112 | 4.963334 | 86.7189 | 1.88E-13 | 4.57E-13 | Healthy control | Liver |
| Smtn | 1.167805 | 7.739464 | 109.3734 | 2.20E-15 | 5.93E-15 | Infection | Liver |
| Smurf1 | 1.0401 | 7.317857 | 425.5134 | 8.17E-30 | 7.81E-29 | Infection | Liver |
| Smurf2 | -0.76287 | 7.725918 | 163.4401 | 3.32E-19 | 1.16E-18 | Healthy control | Liver |
| Smyd2 | -0.86337 | 7.57016 | 225.0197 | 1.58E-22 | 7.15E-22 | Healthy control | Liver |
| Smyd3 | -0.08281 | 4.973196 | 1.266977 | 0.264602 | 0.290482 | Healthy control | Liver |
| Gnai1 | 2.214127 | 4.309565 | 196.466 | 4.94E-21 | 1.97E-20 | Infection | Liver |
| Snai2 | -1.31857 | 5.496872 | 163.9854 | 3.35E-19 | 1.17E-18 | Healthy control | Liver |
| Snai3 | 0.645732 | 1.849362 | 3.565477 | 0.061252 | 0.073216 | Infection | Liver |
| Snap25 | 1.302691 | 1.228552 | 6.159519 | 0.019164 | 0.024473 | Infection | Liver |
| Snap29 | -1.22477 | 6.51146 | 341.1516 | 3.47E-27 | 2.47E-26 | Healthy control | Liver |
| Snap91 | 0.295983 | 1.171812 | 0.925265 | 0.357044 | 0.383801 | Infection | Liver |
| Snapin | -1.11646 | 6.167009 | 236.4937 | 4.79E-23 | 2.26E-22 | Healthy control | Liver |
| Sncb | 1.310368 | 1.295414 | 6.289811 | 0.015333 | 0.019788 | Infection | Liver |
| Grm2 | 2.210997 | 1.30094 | 12.13112 | 0.001184 | 0.001683 | Infection | Liver |
| Snf8 | -0.54994 | 8.650152 | 126.427 | 9.73E-17 | 2.86E-16 | Healthy control | Liver |
| Snph | 1.196748 | 1.25393 | 4.791002 | 0.034875 | 0.043043 | Infection | Liver |
| Snrk | -2.14347 | 7.341055 | 876.2662 | 8.43E-39 | 3.04E-37 | Healthy control | Liver |
| Snrpa | 0.221288 | 7.339064 | 21.82607 | 1.60E-05 | 2.57E-05 | Infection | Liver |
| Snrpd1 | -0.50229 | 2.967653 | 6.91777 | 0.010406 | 0.013647 | Healthy control | Liver |
| Snrpn | -0.20383 | 5.417197 | 4.507813 | 0.037663 | 0.046348 | Healthy control | Liver |
| Snta1 | -1.02899 | 6.288339 | 164.6179 | 2.88E-19 | 1.01E-18 | Healthy control | Liver |
| Sntb1 | -1.32756 | 8.870945 | 308.1588 | 4.99E-26 | 3.19E-25 | Healthy control | Liver |
| Cbl | 2.21037 | 7.248349 | 885.9189 | 5.91E-39 | 2.21E-37 | Infection | Liver |
| Snu13 | 0.543586 | 5.036589 | 50.72324 | 1.22E-09 | 2.42E-09 | Infection | Liver |
| Snx27 | 0.118732 | 6.512652 | 4.245739 | 0.04346 | 0.053025 | Infection | Liver |
| Snx30 | 0.391612 | 6.388087 | 45.73477 | 5.05E-09 | 9.75E-09 | Infection | Liver |
| Snx4 | 0.433194 | 6.879036 | 68.32223 | 1.18E-11 | 2.60E-11 | Infection | Liver |
| Snx5 | 1.294499 | 9.415721 | 892.7632 | 4.59E-39 | 1.76E-37 | Infection | Liver |
| Snx6 | -0.07201 | 6.857865 | 1.128195 | 0.292192 | 0.318327 | Healthy control | Liver |
| Chrnb4 | 2.205584 | 1.316763 | 18.12422 | 8.34E-05 | 0.000128 | Infection | Liver |
| Socs2 | 0.616898 | 7.495288 | 147.5282 | 3.37E-18 | 1.09E-17 | Infection | Liver |
| Tdh | 2.203283 | 1.547926 | 19.34083 | 3.61E-05 | 5.68E-05 | Infection | Liver |
| Socs6 | 0.486522 | 6.526072 | 99.6351 | 1.26E-14 | 3.26E-14 | Infection | Liver |
| Sod1 | -1.71201 | 11.92164 | 626.8981 | 1.42E-34 | 2.49E-33 | Healthy control | Liver |
| Sod2 | -1.6019 | 9.562847 | 244.1985 | 2.20E-23 | 1.07E-22 | Healthy control | Liver |
| Sohlh2 | 0.840244 | 1.291424 | 1.618232 | 0.209136 | 0.232652 | Infection | Liver |
| Sorbs1 | -0.93598 | 8.914987 | 235.3924 | 5.11E-23 | 2.40E-22 | Healthy control | Liver |
| Sorcs3 | 0.107111 | 1.674217 | 0.067734 | 0.795066 | 0.811769 | Infection | Liver |
| Relb | 2.202709 | 5.667812 | 624.3018 | 1.97E-34 | 3.33E-33 | Infection | Liver |
| Pakap | 2.201839 | 6.396566 | 490.1686 | 1.67E-31 | 1.97E-30 | Infection | Liver |
| Sos1 | -0.98048 | 7.272345 | 435.6984 | 4.27E-30 | 4.27E-29 | Healthy control | Liver |
| Sost | 0.585677 | 1.434433 | 1.336526 | 0.250363 | 0.275939 | Infection | Liver |
| Sostdc1 | 0.507477 | 1.285589 | 0.801264 | 0.374943 | 0.401748 | Infection | Liver |
| Sox1 | 0.456672 | 1.29052 | 0.531315 | 0.469258 | 0.49706 | Infection | Liver |
| Sox10 | 0.817678 | 1.253566 | 1.958724 | 0.170033 | 0.191235 | Infection | Liver |
| Sox11 | 1.63492 | 2.209308 | 17.86045 | 4.79E-05 | 7.48E-05 | Infection | Liver |

| | | | | | | | |
|---|---|---|---|---|---|---|---|
| Sox13 | 0.734536 | 5.313123 | 77.02959 | 1.54E-12 | 3.55E-12 | Infection | Liver |
| Rsph1 | 2.201613 | 2.01904 | 39.89812 | 5.09E-09 | 9.81E-09 | Infection | Liver |
| Sox17 | 0.325178 | 4.267674 | 7.275319 | 0.008968 | 0.011833 | Infection | Liver |
| Sox18 | -1.08426 | 6.050819 | 102.3049 | 7.71E-15 | 2.01E-14 | Healthy control | Liver |
| Sox2 | 0.910157 | 1.292792 | 1.898668 | 0.17467 | 0.196053 | Infection | Liver |
| Sox21 | 1.233078 | 1.327685 | 3.943075 | 0.05147 | 0.062117 | Infection | Liver |
| Sox3 | 1.313181 | 1.22226 | 4.981959 | 0.034576 | 0.042696 | Infection | Liver |
| Sox30 | 0.935845 | 1.266809 | 2.125319 | 0.1527 | 0.172945 | Infection | Liver |
| Samd11 | 2.19745 | 1.708581 | 28.82151 | 4.97E-07 | 8.64E-07 | Infection | Liver |
| Sox5 | -1.55292 | 6.559873 | 477.1748 | 3.56E-31 | 3.99E-30 | Healthy control | Liver |
| Sox6 | -0.85137 | 7.011403 | 75.58097 | 2.10E-12 | 4.80E-12 | Healthy control | Liver |
| Sox8 | 0.984293 | 1.471 | 4.894575 | 0.029278 | 0.036422 | Infection | Liver |
| Tnfsf14 | 2.194902 | 2.936459 | 82.44728 | 3.42E-14 | 8.65E-14 | Infection | Liver |
| Sp1 | 0.098313 | 7.210115 | 5.747457 | 0.019471 | 0.02484 | Infection | Liver |
| Sp3 | 0.648084 | 8.115619 | 264.9606 | 2.54E-24 | 1.37E-23 | Infection | Liver |
| Sp5 | -1.18965 | 2.592551 | 18.2948 | 4.77E-05 | 7.45E-05 | Healthy control | Liver |
| Sp7 | 1.184327 | 1.253558 | 4.522296 | 0.040376 | 0.049504 | Infection | Liver |
| Sp8 | 0.921319 | 1.289974 | 2.293551 | 0.136479 | 0.155843 | Infection | Liver |
| Rab8b | 2.194243 | 5.84615 | 411.6155 | 2.29E-29 | 2.10E-28 | Infection | Liver |
| Spaca6 | 1.480552 | 3.573543 | 95.61166 | 2.54E-14 | 6.45E-14 | Infection | Liver |
| Spaca9 | 1.264575 | 1.906633 | 16.48646 | 8.35E-05 | 0.000129 | Infection | Liver |
| Spag9 | -0.58055 | 7.627907 | 161.0432 | 4.65E-19 | 1.61E-18 | Healthy control | Liver |
| Spast | -0.3293 | 8.7827 | 70.80668 | 6.44E-12 | 1.44E-11 | Healthy control | Liver |
| Spata25 | 0.639115 | 1.524979 | 2.086295 | 0.151397 | 0.171547 | Infection | Liver |
| Dock2 | 2.191929 | 6.474894 | 389.5973 | 9.56E-29 | 8.20E-28 | Infection | Liver |
| Spatc1l | -0.4681 | 1.363146 | 0.804453 | 0.372204 | 0.399069 | Healthy control | Liver |
| Spdef | 0.849474 | 1.267057 | 2.181087 | 0.147084 | 0.167009 | Infection | Liver |
| Mfsd2b | 2.190636 | 4.189794 | 76.49266 | 1.77E-12 | 4.06E-12 | Infection | Liver |
| Itk | 2.190263 | 3.674622 | 101.5273 | 4.90E-15 | 1.29E-14 | Infection | Liver |
| Trpm2 | 2.190147 | 5.185347 | 176.4549 | 6.68E-20 | 2.48E-19 | Infection | Liver |
| Mxd1 | 2.188513 | 6.689792 | 647.0672 | 5.94E-35 | 1.12E-33 | Infection | Liver |
| Sphk2 | -1.49064 | 8.207709 | 203.7949 | 1.90E-21 | 7.83E-21 | Healthy control | Liver |
| Spi1 | 0.762597 | 6.379129 | 37.18422 | 7.18E-08 | 1.30E-07 | Infection | Liver |
| Cdh7 | 2.186831 | 1.732555 | 22.47096 | 7.05E-06 | 1.16E-05 | Infection | Liver |
| Adcyap1 | 2.185005 | 1.471974 | 13.08747 | 0.000533 | 0.000778 | Infection | Liver |
| Spink12 | 0.490617 | 1.26773 | 0.528509 | 0.471106 | 0.498806 | Infection | Liver |
| Spink8 | 0.873474 | 1.301187 | 1.975902 | 0.165535 | 0.186595 | Infection | Liver |
| Chia1 | 2.183604 | 1.305079 | 9.898259 | 0.002927 | 0.004027 | Infection | Liver |
| Marco | 2.183435 | 5.32215 | 33.86669 | 2.76E-07 | 4.88E-07 | Infection | Liver |
| Spock3 | 1.55523 | 2.393612 | 32.72362 | 1.07E-07 | 1.92E-07 | Infection | Liver |
| Nfkb2 | 2.182362 | 6.91227 | 1298.889 | 6.38E-44 | 7.60E-42 | Infection | Liver |
| Spop | -0.35392 | 8.916538 | 75.13192 | 2.33E-12 | 5.28E-12 | Healthy control | Liver |
| Spopl | 0.288911 | 6.176676 | 28.32396 | 1.44E-06 | 2.43E-06 | Infection | Liver |
| Spred1 | 0.319477 | 7.855702 | 13.2758 | 0.000543 | 0.000791 | Infection | Liver |
| Sprr2d | 0.620319 | 1.217725 | 1.058643 | 0.313136 | 0.339519 | Infection | Liver |
| Prkcg | 2.182276 | 2.330975 | 73.46996 | 4.53E-14 | 1.14E-13 | Infection | Liver |
| Spry1 | 0.880915 | 5.932623 | 65.38935 | 2.44E-11 | 5.29E-11 | Infection | Liver |
| Spry2 | 0.877779 | 4.591197 | 40.08284 | 2.92E-08 | 5.39E-08 | Infection | Liver |
| Spry4 | -0.20723 | 6.348842 | 4.794751 | 0.032237 | 0.039955 | Healthy control | Liver |
| Klrb1c | 2.179302 | 1.778322 | 22.83653 | 5.86E-06 | 9.65E-06 | Infection | Liver |

| | | | | | | | |
|---|---|---|---|---|---|---|---|
| Spsb2 | -0.98893 | 7.279028 | 343.0837 | 2.90E-27 | 2.10E-26 | Healthy control | Liver |
| Spata4 | 2.176032 | 1.481371 | 18.44295 | 5.16E-05 | 8.04E-05 | Infection | Liver |
| Sptan1 | 0.370831 | 9.129712 | 110.7537 | 1.53E-15 | 4.15E-15 | Infection | Liver |
| Nkx6-1 | 2.175215 | 1.419355 | 14.22377 | 0.000338 | 0.000499 | Infection | Liver |
| Sptbn2 | -1.10227 | 7.07376 | 124.2251 | 1.42E-16 | 4.12E-16 | Healthy control | Liver |
| Sptlc2 | 0.544791 | 7.839084 | 142.1442 | 7.68E-18 | 2.42E-17 | Infection | Liver |
| Sqstm1 | -1.00013 | 10.48086 | 355.2763 | 1.12E-27 | 8.61E-27 | Healthy control | Liver |
| Sra1 | -0.7366 | 7.87601 | 183.6562 | 2.18E-20 | 8.35E-20 | Healthy control | Liver |
| Ptf1a | 2.174874 | 1.390988 | 11.08735 | 0.001506 | 0.002121 | Infection | Liver |
| Srd5a3 | 0.459399 | 6.564853 | 59.68524 | 1.04E-10 | 2.19E-10 | Infection | Liver |
| Srebf1 | -1.21547 | 9.501624 | 218.9398 | 3.10E-22 | 1.37E-21 | Healthy control | Liver |
| Srebf2 | -0.05538 | 8.843656 | 0.827516 | 0.366438 | 0.393224 | Healthy control | Liver |
| Srf | -0.62544 | 5.402088 | 54.54044 | 4.20E-10 | 8.58E-10 | Healthy control | Liver |
| Srgap1 | 0.198943 | 3.661735 | 1.77051 | 0.188093 | 0.210412 | Infection | Liver |
| Srp72 | 0.100658 | 8.004888 | 5.2801 | 0.024882 | 0.03132 | Infection | Liver |
| Pdgfb | 2.172872 | 5.134745 | 259.3579 | 5.59E-24 | 2.91E-23 | Infection | Liver |
| Srr | -2.35817 | 6.96954 | 546.3233 | 8.15E-33 | 1.09E-31 | Healthy control | Liver |
| Srrm1 | -0.29974 | 8.286686 | 60.9836 | 7.42E-11 | 1.57E-10 | Healthy control | Liver |
| Srrm4 | 0.379778 | 6.33355 | 6.365042 | 0.014218 | 0.018397 | Infection | Liver |
| Srsf1 | -1.3467 | 7.761118 | 612.5841 | 2.79E-34 | 4.57E-33 | Healthy control | Liver |
| Srsf2 | -0.16227 | 9.681949 | 15.18638 | 0.000237 | 0.000354 | Healthy control | Liver |
| Srsf4 | 0.223782 | 8.657526 | 32.57128 | 3.27E-07 | 5.73E-07 | Infection | Liver |
| Sry | 0.963617 | 1.234269 | 2.464593 | 0.127259 | 0.146282 | Infection | Liver |
| Ss18l1 | -1.01596 | 4.90915 | 121.7697 | 2.31E-16 | 6.59E-16 | Healthy control | Liver |
| Ssbp2 | 0.970197 | 4.287128 | 43.49274 | 1.04E-08 | 1.98E-08 | Infection | Liver |
| Ssr2 | -0.83962 | 7.617092 | 123.2315 | 1.68E-16 | 4.83E-16 | Healthy control | Liver |
| Ssrp1 | 0.26183 | 7.61235 | 32.7203 | 3.11E-07 | 5.46E-07 | Infection | Liver |
| Sst | 1.683276 | 1.361461 | 5.516404 | 0.022027 | 0.027915 | Infection | Liver |
| Sstr1 | -0.28305 | 1.393649 | 0.37431 | 0.542059 | 0.56852 | Healthy control | Liver |
| Sstr2 | 1.985154 | 1.405147 | 13.11643 | 0.00058 | 0.000841 | Infection | Liver |
| Sstr3 | 1.178794 | 2.537786 | 29.69374 | 4.40E-07 | 7.68E-07 | Infection | Liver |
| Sstr4 | 1.442061 | 1.44161 | 9.192309 | 0.003269 | 0.004478 | Infection | Liver |
| Piwil2 | 2.172227 | 1.784457 | 30.68942 | 2.34E-07 | 4.15E-07 | Infection | Liver |
| Stac | 1.688524 | 1.528442 | 11.77962 | 0.000909 | 0.001304 | Infection | Liver |
| Stag2 | -0.75043 | 7.417551 | 274.7653 | 1.00E-24 | 5.62E-24 | Healthy control | Liver |
| Stam2 | 0.100976 | 6.200998 | 2.879404 | 0.094629 | 0.110448 | Infection | Liver |
| Star | -0.21273 | 2.907652 | 1.115307 | 0.294441 | 0.320568 | Healthy control | Liver |
| Stat1 | 0.47274 | 8.45796 | 16.50564 | 0.000139 | 0.00021 | Infection | Liver |
| Stat2 | -0.21196 | 8.409903 | 14.88858 | 0.00027 | 0.000401 | Healthy control | Liver |
| Stat3 | -0.56233 | 8.427804 | 147.4864 | 3.39E-18 | 1.09E-17 | Healthy control | Liver |
| Drc7 | 2.170635 | 1.381416 | 15.0278 | 0.00027 | 0.000402 | Infection | Liver |
| Stat5a | -0.90797 | 6.055844 | 190.2261 | 9.82E-21 | 3.84E-20 | Healthy control | Liver |
| Stat5b | -1.7001 | 8.098605 | 845.0995 | 2.35E-38 | 7.90E-37 | Healthy control | Liver |
| Stat6 | -0.05909 | 7.69748 | 0.739483 | 0.393068 | 0.420448 | Healthy control | Liver |
| Stau2 | 0.482275 | 5.652471 | 29.94318 | 8.12E-07 | 1.40E-06 | Infection | Liver |
| Kcnh1 | 2.169035 | 1.444563 | 16.23449 | 0.000152 | 0.000229 | Infection | Liver |
| Stim1 | -0.24545 | 7.057821 | 14.7953 | 0.000281 | 0.000417 | Healthy control | Liver |
| Stim2 | -0.22476 | 6.701727 | 16.68566 | 0.000126 | 0.000192 | Healthy control | Liver |
| Stimate | -0.55809 | 5.233948 | 51.95035 | 8.63E-10 | 1.73E-09 | Healthy control | Liver |
| Stip1 | -1.57248 | 7.467728 | 545.0249 | 7.95E-33 | 1.07E-31 | Healthy control | Liver |

| | | | | | | | |
|---|---|---|---|---|---|---|---|
| Stk11 | -0.30762 | 8.606244 | 48.5597 | 2.21E-09 | 4.34E-09 | Healthy control | Liver |
| Stk3 | 0.472229 | 6.267597 | 51.10708 | 1.07E-09 | 2.14E-09 | Infection | Liver |
| Stk32a | 1.728483 | 1.31648 | 8.39559 | 0.005799 | 0.007786 | Infection | Liver |
| Stk38 | -0.35973 | 7.347011 | 78.1031 | 1.18E-12 | 2.74E-12 | Healthy control | Liver |
| Stk38l | 0.454302 | 6.100369 | 47.65859 | 2.88E-09 | 5.63E-09 | Infection | Liver |
| Stk39 | 1.2647 | 5.601745 | 194.4371 | 5.84E-21 | 2.31E-20 | Infection | Liver |
| Stk4 | 0.557903 | 7.142636 | 91.03884 | 7.14E-14 | 1.77E-13 | Infection | Liver |
| Dbx2 | 2.16618 | 1.313732 | 13.67784 | 0.000548 | 0.000796 | Infection | Liver |
| Stom | 0.070697 | 8.318727 | 1.534323 | 0.220033 | 0.244285 | Infection | Liver |
| Stoml3 | 1.466455 | 1.26823 | 5.487637 | 0.024981 | 0.031437 | Infection | Liver |
| Stra6l | -0.93763 | 8.67963 | 86.93986 | 1.74E-13 | 4.25E-13 | Healthy control | Liver |
| Pygb | 2.163253 | 6.059268 | 636.6118 | 1.02E-34 | 1.83E-33 | Infection | Liver |
| Strn3 | -0.71254 | 7.719599 | 112.3138 | 1.15E-15 | 3.15E-15 | Healthy control | Liver |
| Stub1 | -0.00294 | 7.932079 | 0.005246 | 0.942487 | 0.94855 | Healthy control | Liver |
| Stx1a | 1.715037 | 4.447514 | 132.1953 | 4.59E-17 | 1.38E-16 | Infection | Liver |
| Stx8 | 0.607279 | 6.875267 | 125.6893 | 1.11E-16 | 3.24E-16 | Infection | Liver |
| Stxbp1 | 0.870371 | 5.848091 | 92.70254 | 5.12E-14 | 1.28E-13 | Infection | Liver |
| Stxbp4 | 0.549697 | 5.470574 | 37.75895 | 5.94E-08 | 1.08E-07 | Infection | Liver |
| Stxbp5l | 1.24046 | 1.638697 | 6.456829 | 0.012352 | 0.016078 | Infection | Liver |
| Sucnr1 | -2.56234 | 4.156237 | 188.2565 | 4.03E-22 | 1.77E-21 | Healthy control | Liver |
| Sufu | -0.10264 | 5.864876 | 2.159173 | 0.146674 | 0.16661 | Healthy control | Liver |
| Tgfbr1 | 2.1623 | 7.613893 | 1215.243 | 4.63E-43 | 4.14E-41 | Infection | Liver |
| Sulf2 | -0.43404 | 6.2486 | 7.823833 | 0.006865 | 0.009161 | Healthy control | Liver |
| Sult2a8 | -3.30125 | 10.54736 | 225.3021 | 1.15E-21 | 4.81E-21 | Healthy control | Liver |
| Sumf1 | -0.70731 | 3.882877 | 18.43894 | 6.07E-05 | 9.41E-05 | Healthy control | Liver |
| Sumo1 | -0.60094 | 7.146037 | 93.15581 | 4.62E-14 | 1.16E-13 | Healthy control | Liver |
| Sumo3 | -1.40352 | 7.967323 | 699.5933 | 5.97E-36 | 1.33E-34 | Healthy control | Liver |
| Sun1 | -0.16982 | 6.606048 | 5.084887 | 0.027599 | 0.034463 | Healthy control | Liver |
| Supt16 | 0.023832 | 6.906501 | 0.176099 | 0.676167 | 0.698924 | Infection | Liver |
| Supt20 | -0.1221 | 6.684989 | 4.805509 | 0.03205 | 0.039734 | Healthy control | Liver |
| Supt3 | -0.35541 | 5.289352 | 15.15538 | 0.000241 | 0.00036 | Healthy control | Liver |
| Supt5 | -0.14575 | 8.04409 | 11.79909 | 0.00105 | 0.001501 | Healthy control | Liver |
| Supt6 | -0.67835 | 8.188235 | 312.0993 | 3.57E-26 | 2.31E-25 | Healthy control | Liver |
| Suv39h1 | 0.115284 | 4.449349 | 1.186434 | 0.280206 | 0.306202 | Infection | Liver |
| Suz12 | -0.19562 | 6.775654 | 15.27618 | 0.000229 | 0.000342 | Healthy control | Liver |
| Dst | 2.16154 | 6.416569 | 375.932 | 2.53E-28 | 2.07E-27 | Infection | Liver |
| Panx1 | 2.159017 | 4.21434 | 164.6161 | 2.97E-19 | 1.05E-18 | Infection | Liver |
| Svil | 0.535648 | 8.417088 | 100.7518 | 1.01E-14 | 2.63E-14 | Infection | Liver |
| Syce2 | 0.371556 | 3.294808 | 7.790846 | 0.006894 | 0.009195 | Infection | Liver |
| Cryge | 2.152747 | 1.326482 | 9.307807 | 0.003677 | 0.005013 | Infection | Liver |
| Msln | 2.152715 | 1.381114 | 11.03408 | 0.001745 | 0.002444 | Infection | Liver |
| Cd160 | 2.149958 | 2.555855 | 52.92433 | 7.11E-11 | 1.51E-10 | Infection | Liver |
| Syn1 | 1.17358 | 3.098515 | 45.11199 | 4.49E-09 | 8.69E-09 | Infection | Liver |
| Syn2 | 1.774075 | 1.343483 | 9.47353 | 0.003175 | 0.004355 | Infection | Liver |
| Syn3 | -0.55118 | 4.3016 | 18.76261 | 5.40E-05 | 8.40E-05 | Healthy control | Liver |
| Syncrip | 0.401356 | 7.881587 | 37.51502 | 6.38E-08 | 1.16E-07 | Infection | Liver |
| Syne1 | -0.37311 | 6.875971 | 29.59608 | 9.14E-07 | 1.57E-06 | Healthy control | Liver |
| Syne4 | 1.588129 | 3.049825 | 74.01444 | 9.76E-13 | 2.27E-12 | Infection | Liver |
| Syngr3 | 0.040066 | 1.744188 | 0.010463 | 0.918683 | 0.926643 | Infection | Liver |
| Synj2bp | -0.05141 | 7.309288 | 0.647725 | 0.423936 | 0.451249 | Healthy control | Liver |

| | | | | | | | |
|---|---|---|---|---|---|---|---|
| Cxcr5 | 2.149383 | 1.961646 | 39.76069 | 5.73E-09 | 1.10E-08 | Infection | Liver |
| Clspn | 2.148666 | 3.789344 | 219.8571 | 1.34E-22 | 6.12E-22 | Infection | Liver |
| Syp | 0.163407 | 2.871123 | 0.93985 | 0.335601 | 0.361839 | Infection | Liver |
| Sypl2 | 0.476388 | 1.278983 | 0.593824 | 0.444829 | 0.472684 | Infection | Liver |
| Syt2 | 1.029414 | 1.277035 | 2.793121 | 0.101312 | 0.117891 | Infection | Liver |
| Trdn | 2.148306 | 1.750365 | 22.06616 | 8.06E-06 | 1.32E-05 | Infection | Liver |
| Syt6 | 1.764612 | 1.463529 | 9.741799 | 0.002598 | 0.003592 | Infection | Liver |
| Syt9 | 0.654884 | 3.252276 | 9.969962 | 0.002414 | 0.003346 | Infection | Liver |
| Sytl5 | 1.452986 | 5.952036 | 89.15486 | 1.17E-13 | 2.89E-13 | Infection | Liver |
| Szt2 | -0.40256 | 5.701073 | 26.59052 | 2.69E-06 | 4.49E-06 | Healthy control | Liver |
| T | 1.946401 | 1.27773 | 13.53518 | 0.000636 | 0.00092 | Infection | Liver |
| Taar5 | 1.511571 | 1.277977 | 6.953452 | 0.011697 | 0.015265 | Infection | Liver |
| Tab1 | 0.270203 | 4.826068 | 6.758145 | 0.011612 | 0.015169 | Infection | Liver |
| Tac1 | 1.967016 | 1.3504 | 9.249501 | 0.004035 | 0.005483 | Infection | Liver |
| Map4k4 | 2.146837 | 8.528098 | 652.9799 | 4.39E-35 | 8.69E-34 | Infection | Liver |
| Tacr1 | 1.652666 | 1.247468 | 5.951236 | 0.020614 | 0.026251 | Infection | Liver |
| Tacr2 | -0.21563 | 1.219891 | 0.138555 | 0.712889 | 0.7334 | Healthy control | Liver |
| Tacr3 | 1.17409 | 1.348715 | 3.367879 | 0.071086 | 0.084227 | Infection | Liver |
| Cltrn | 2.146121 | 1.62989 | 11.67639 | 0.000944 | 0.001351 | Infection | Liver |
| Taf11 | -0.38018 | 5.455797 | 24.97551 | 4.88E-06 | 8.07E-06 | Healthy control | Liver |
| Taf12 | 0.024829 | 7.432189 | 0.212474 | 0.646413 | 0.669968 | Infection | Liver |
| Taf15 | 0.033831 | 8.34914 | 0.413914 | 0.52231 | 0.549538 | Infection | Liver |
| Taf4 | -0.46386 | 6.78247 | 83.39266 | 3.64E-13 | 8.74E-13 | Healthy control | Liver |
| Taf5l | -0.90112 | 5.833473 | 116.8655 | 5.25E-16 | 1.46E-15 | Healthy control | Liver |
| Taf6l | 0.329947 | 2.949393 | 3.85148 | 0.053731 | 0.064642 | Infection | Liver |
| Taf9 | -0.97351 | 5.79805 | 150.0455 | 2.43E-18 | 7.94E-18 | Healthy control | Liver |
| Prm1 | 2.144417 | 1.309599 | 13.89191 | 0.000476 | 0.000697 | Infection | Liver |
| Tal1 | 1.51304 | 3.576689 | 58.36038 | 1.30E-10 | 2.72E-10 | Infection | Liver |
| Tap1 | 1.226845 | 7.617086 | 194.5962 | 5.49E-21 | 2.18E-20 | Infection | Liver |
| Tardbp | -0.2199 | 9.423596 | 50.87606 | 1.14E-09 | 2.27E-09 | Healthy control | Liver |
| Tas1r1 | 0.882176 | 1.442677 | 3.435934 | 0.066805 | 0.079323 | Infection | Liver |
| Tas1r2 | -0.22443 | 1.195882 | 0.18378 | 0.67336 | 0.696167 | Healthy control | Liver |
| Tas1r3 | -0.91524 | 2.9965 | 22.69698 | 9.09E-06 | 1.48E-05 | Healthy control | Liver |
| Tbc1d1 | 0.739761 | 5.683588 | 74.63202 | 2.64E-12 | 5.99E-12 | Infection | Liver |
| Tbc1d22a | 0.554937 | 7.236194 | 133.2979 | 3.13E-17 | 9.52E-17 | Infection | Liver |
| Tbck | -0.39057 | 6.980837 | 77.70409 | 1.29E-12 | 2.99E-12 | Healthy control | Liver |
| Tbk1 | 0.197712 | 7.317232 | 15.59301 | 0.0002 | 0.0003 | Infection | Liver |
| Tbl1xr1 | -0.12862 | 7.125373 | 5.670562 | 0.020268 | 0.025817 | Healthy control | Liver |
| Tbp | -0.36286 | 6.19944 | 22.48267 | 1.25E-05 | 2.01E-05 | Healthy control | Liver |
| Tbpl1 | 0.427776 | 6.013698 | 43.22188 | 1.08E-08 | 2.04E-08 | Infection | Liver |
| Tbpl2 | 0.791841 | 1.228088 | 2.321872 | 0.138492 | 0.157889 | Infection | Liver |
| Tbr1 | 1.182274 | 1.278437 | 3.384716 | 0.072983 | 0.086356 | Infection | Liver |
| Lrrtm3 | 2.143332 | 1.417968 | 14.0921 | 0.000364 | 0.000537 | Infection | Liver |
| Tbx18 | 1.007535 | 1.266568 | 2.709238 | 0.107754 | 0.124894 | Infection | Liver |
| Tbx2 | 1.033896 | 6.483452 | 46.8363 | 4.08E-09 | 7.92E-09 | Infection | Liver |
| Tbx20 | 1.145594 | 7.128918 | 58.14195 | 2.09E-10 | 4.34E-10 | Infection | Liver |
| Tbx21 | 1.54182 | 3.420575 | 47.47186 | 2.51E-09 | 4.93E-09 | Infection | Liver |
| Tbx3 | -2.30536 | 7.203742 | 439.2187 | 3.62E-30 | 3.66E-29 | Healthy control | Liver |
| Tbx4 | 1.422026 | 1.578764 | 9.2706 | 0.002922 | 0.004022 | Infection | Liver |
| Hey2 | 2.140052 | 2.620107 | 70.50884 | 3.46E-13 | 8.33E-13 | Infection | Liver |

| | | | | | | | |
|---|---|---|---|---|---|---|---|
| Tbx6 | 0.571848 | 2.482621 | 5.26243 | 0.024254 | 0.030591 | Infection | Liver |
| Tbxa2r | -1.54137 | 2.986488 | 54.80191 | 1.07E-10 | 2.25E-10 | Healthy control | Liver |
| Tcaf1 | 1.022855 | 4.96636 | 52.39352 | 7.65E-10 | 1.54E-09 | Infection | Liver |
| Tcam1 | 1.638905 | 1.510459 | 14.40941 | 0.000268 | 0.000398 | Infection | Liver |
| Tceal5 | -0.39155 | 2.251328 | 1.55703 | 0.21515 | 0.23913 | Healthy control | Liver |
| Tceal9 | 1.52338 | 6.395951 | 193.8009 | 6.14E-21 | 2.43E-20 | Infection | Liver |
| Tcf12 | -0.31727 | 6.807457 | 28.56265 | 1.32E-06 | 2.24E-06 | Healthy control | Liver |
| Tcf21 | 0.419118 | 4.357709 | 4.979971 | 0.029227 | 0.036368 | Infection | Liver |
| Tcf3 | -0.07123 | 6.679021 | 1.036266 | 0.312563 | 0.339045 | Healthy control | Liver |
| Tcf7 | 0.571323 | 6.300699 | 36.58756 | 8.62E-08 | 1.56E-07 | Infection | Liver |
| Tcf7l1 | -0.17206 | 5.905209 | 4.080265 | 0.047624 | 0.057824 | Healthy control | Liver |
| Tcf7l2 | -0.89232 | 6.700688 | 254.2577 | 7.41E-24 | 3.80E-23 | Healthy control | Liver |
| Tcof1 | 0.009923 | 6.086644 | 0.020637 | 0.886228 | 0.896254 | Infection | Liver |
| Tcp1 | -0.83551 | 8.403476 | 393.0907 | 7.19E-29 | 6.22E-28 | Healthy control | Liver |
| Tctn1 | 1.334248 | 4.704148 | 155.7445 | 1.17E-18 | 3.93E-18 | Infection | Liver |
| Tdg | -0.58528 | 6.214395 | 107.4045 | 2.88E-15 | 7.69E-15 | Healthy control | Liver |
| Tdgf1 | 1.184344 | 1.670764 | 7.907648 | 0.005709 | 0.007678 | Infection | Liver |
| Zfp42 | 2.139389 | 1.295959 | 13.6633 | 0.00055 | 0.000799 | Infection | Liver |
| Sertad1 | 2.138909 | 4.163711 | 220.338 | 2.95E-22 | 1.31E-21 | Infection | Liver |
| Tdrkh | -0.83835 | 4.474195 | 64.55028 | 3.08E-11 | 6.64E-11 | Healthy control | Liver |
| Tead1 | 1.831629 | 7.298799 | 710.7427 | 3.79E-36 | 8.87E-35 | Infection | Liver |
| Tead2 | 1.381866 | 4.208288 | 50.81816 | 1.25E-09 | 2.49E-09 | Infection | Liver |
| Tead3 | -0.41199 | 6.015581 | 43.42963 | 1.01E-08 | 1.92E-08 | Healthy control | Liver |
| Tec | -0.19502 | 8.266885 | 11.41789 | 0.001249 | 0.00177 | Healthy control | Liver |
| Tecpr2 | -0.31237 | 5.928573 | 31.072 | 5.48E-07 | 9.51E-07 | Healthy control | Liver |
| Tecr | -1.11851 | 7.338897 | 140.3657 | 1.02E-17 | 3.18E-17 | Healthy control | Liver |
| Tectb | 0.312621 | 1.637775 | 0.718265 | 0.398239 | 0.425889 | Infection | Liver |
| Tek | -0.81528 | 6.789854 | 52.34055 | 7.57E-10 | 1.53E-09 | Healthy control | Liver |
| Tenm1 | 0.209961 | 1.37614 | 0.115364 | 0.735052 | 0.754649 | Infection | Liver |
| Ptx3 | 2.137037 | 1.411192 | 11.70265 | 0.001088 | 0.001554 | Infection | Liver |
| Ncaph | 2.134298 | 4.219103 | 163.3222 | 3.63E-19 | 1.26E-18 | Infection | Liver |
| Tepsin | -0.99269 | 6.442887 | 286.6197 | 3.44E-25 | 2.00E-24 | Healthy control | Liver |
| Terf1 | -0.50941 | 6.019893 | 60.72302 | 8.00E-11 | 1.69E-10 | Healthy control | Liver |
| Terf2 | 0.213302 | 7.097842 | 21.45347 | 1.85E-05 | 2.97E-05 | Infection | Liver |
| Tert | -1.665 | 5.970542 | 440.0663 | 3.78E-30 | 3.82E-29 | Healthy control | Liver |
| Tesmin | -0.35743 | 2.334341 | 1.309192 | 0.255487 | 0.281092 | Healthy control | Liver |
| Qrfpr | 2.130462 | 1.817835 | 13.45375 | 0.000379 | 0.000559 | Infection | Liver |
| Sycp1 | 2.129261 | 2.172908 | 26.57131 | 1.10E-06 | 1.87E-06 | Infection | Liver |
| Tet2 | -0.33797 | 5.6699 | 16.89819 | 0.000116 | 0.000176 | Healthy control | Liver |
| Tet3 | 0.102003 | 6.624652 | 3.051527 | 0.085507 | 0.100387 | Infection | Liver |
| Tex101 | 1.090467 | 1.356837 | 3.526146 | 0.064704 | 0.076992 | Infection | Liver |
| Tex19.1 | 0.125956 | 1.320516 | 0.040361 | 0.841443 | 0.854588 | Infection | Liver |
| Tfam | -0.2632 | 7.45828 | 17.11374 | 0.000106 | 0.000162 | Healthy control | Liver |
| Tfap2a | 0.981015 | 1.440952 | 3.802593 | 0.05431 | 0.065292 | Infection | Liver |
| Kcnh2 | 2.128964 | 1.463094 | 15.08466 | 0.000228 | 0.000341 | Infection | Liver |
| Tfap2c | 1.294862 | 1.25723 | 3.707345 | 0.061958 | 0.073954 | Infection | Liver |
| Tfap2d | 0.288557 | 1.218432 | 0.267615 | 0.609257 | 0.633951 | Infection | Liver |
| Tfeb | 0.337761 | 6.651174 | 33.65049 | 2.27E-07 | 4.04E-07 | Infection | Liver |
| Tfpi | 1.482056 | 7.016435 | 320.191 | 1.83E-26 | 1.22E-25 | Infection | Liver |
| Tfrc | 1.564218 | 7.045606 | 356.9729 | 9.96E-28 | 7.74E-27 | Infection | Liver |

| | | | | | | | |
|---|---|---|---|---|---|---|---|
| Tgfa | -0.37989 | 6.474399 | 17.49887 | 9.02E-05 | 0.000138 | Healthy control | Liver |
| Tgfb1 | 1.243258 | 6.920422 | 113.7336 | 9.56E-16 | 2.63E-15 | Infection | Liver |
| Tgfb1i1 | 1.877639 | 5.534175 | 191.5154 | 8.73E-21 | 3.42E-20 | Infection | Liver |
| Prdm16 | 2.126709 | 1.579403 | 14.57533 | 0.000259 | 0.000386 | Infection | Liver |
| Fbxl16 | 2.126214 | 1.890695 | 39.99302 | 5.04E-09 | 9.73E-09 | Infection | Liver |
| Adam28 | 2.126047 | 1.641912 | 17.26217 | 7.09E-05 | 0.00011 | Infection | Liver |
| Scin | 2.123425 | 1.464076 | 12.65785 | 0.000652 | 0.000942 | Infection | Liver |
| Tgfbr3 | 0.571735 | 8.250784 | 50.23811 | 1.36E-09 | 2.71E-09 | Infection | Liver |
| Tgif2 | 1.781961 | 3.367026 | 116.9775 | 2.13E-16 | 6.09E-16 | Infection | Liver |
| Hoxd1 | 2.123035 | 1.463245 | 10.14125 | 0.002106 | 0.002928 | Infection | Liver |
| Tgs1 | -1.1603 | 3.492673 | 63.00376 | 2.52E-11 | 5.45E-11 | Healthy control | Liver |
| Th | 1.81E-16 | 1.160624 | 0 | 1 | 1 | Infection | Liver |
| Thap12 | 0.70033 | 8.245845 | 188.0594 | 1.24E-20 | 4.82E-20 | Infection | Liver |
| Thbd | 0.308779 | 4.025501 | 4.713362 | 0.033724 | 0.041716 | Infection | Liver |
| Ppm1h | 2.122707 | 5.782979 | 452.4205 | 1.69E-30 | 1.76E-29 | Infection | Liver |
| Inpp4b | 2.122654 | 5.508006 | 348.2639 | 2.28E-27 | 1.68E-26 | Infection | Liver |
| Adgrg2 | 2.12228 | 2.35179 | 45.95475 | 5.99E-10 | 1.21E-09 | Infection | Liver |
| Akt3 | 2.118731 | 3.809826 | 161.2896 | 2.49E-19 | 8.80E-19 | Infection | Liver |
| Thpo | -1.03395 | 4.768802 | 104.2882 | 5.42E-15 | 1.43E-14 | Healthy control | Liver |
| Thra | 0.429404 | 7.03814 | 29.78236 | 8.56E-07 | 1.47E-06 | Infection | Liver |
| Thrap3 | 0.030773 | 8.193557 | 0.442932 | 0.508125 | 0.535511 | Infection | Liver |
| Thrb | -0.7615 | 7.813343 | 75.89865 | 1.95E-12 | 4.46E-12 | Healthy control | Liver |
| Thumpd1 | -0.6244 | 6.378046 | 56.67376 | 2.32E-10 | 4.80E-10 | Healthy control | Liver |
| Tiam1 | 1.745209 | 5.250361 | 217.2264 | 4.29E-22 | 1.88E-21 | Infection | Liver |
| Ticam1 | -0.74438 | 4.036742 | 44.21954 | 7.78E-09 | 1.49E-08 | Healthy control | Liver |
| Kif24 | 2.115708 | 3.431659 | 142.8187 | 1.54E-18 | 5.12E-18 | Infection | Liver |
| Tie1 | 0.59167 | 6.355788 | 41.99212 | 1.56E-08 | 2.94E-08 | Infection | Liver |
| Tigar | -0.82422 | 4.609942 | 35.98247 | 1.06E-07 | 1.91E-07 | Healthy control | Liver |
| Slc14a1 | 2.114637 | 2.736288 | 101.4168 | 1.49E-16 | 4.32E-16 | Infection | Liver |
| Timeless | 1.771436 | 4.092778 | 137.2867 | 1.93E-17 | 5.92E-17 | Infection | Liver |
| Timm50 | 0.221249 | 7.695213 | 13.24121 | 0.000552 | 0.000801 | Infection | Liver |
| Timmdc1 | -0.41346 | 7.415311 | 28.78771 | 1.22E-06 | 2.07E-06 | Healthy control | Liver |
| Kcng1 | 2.110854 | 1.464021 | 12.18637 | 0.000843 | 0.001212 | Infection | Liver |
| Timp4 | 0.630274 | 1.244635 | 1.028593 | 0.317695 | 0.34409 | Infection | Liver |
| Tiparp | 1.75322 | 6.314798 | 427.2734 | 7.56E-30 | 7.26E-29 | Infection | Liver |
| Tirap | -1.29815 | 7.119205 | 585.0772 | 1.06E-33 | 1.58E-32 | Healthy control | Liver |
| Tjp1 | 0.639875 | 8.01742 | 132.4818 | 3.57E-17 | 1.08E-16 | Infection | Liver |
| Tjp2 | 0.009559 | 5.608434 | 0.014458 | 0.904673 | 0.913984 | Infection | Liver |
| Tjp3 | -0.84673 | 7.314727 | 94.59475 | 3.45E-14 | 8.71E-14 | Healthy control | Liver |
| Tlcd1 | -1.14415 | 7.693928 | 161.5323 | 4.34E-19 | 1.51E-18 | Healthy control | Liver |
| Tle1 | -1.74652 | 8.62102 | 477.391 | 3.31E-31 | 3.75E-30 | Healthy control | Liver |
| Tle4 | -0.56317 | 6.357433 | 53.49898 | 5.51E-10 | 1.12E-09 | Healthy control | Liver |
| Tle6 | 0.133999 | 3.161519 | 0.888979 | 0.349172 | 0.375581 | Infection | Liver |
| Tll1 | 1.66771 | 1.634567 | 13.72486 | 0.000339 | 0.000501 | Infection | Liver |
| Tln1 | 0.927783 | 8.020076 | 221.7896 | 2.26E-22 | 1.01E-21 | Infection | Liver |
| Tlr1 | 1.817667 | 2.954063 | 103.164 | 6.10E-16 | 1.70E-15 | Infection | Liver |
| Rab27b | 2.109129 | 3.808571 | 226.6079 | 6.82E-23 | 3.18E-22 | Infection | Liver |
| Tlr12 | -0.00202 | 2.598053 | 0.000161 | 0.989909 | 0.990502 | Healthy control | Liver |
| Tlr13 | 1.678779 | 5.018166 | 227.323 | 1.46E-22 | 6.63E-22 | Infection | Liver |
| Tlr2 | 1.725036 | 5.670074 | 171.3711 | 1.16E-19 | 4.25E-19 | Infection | Liver |

| | | | | | | | |
|---|---|---|---|---|---|---|---|
| Tlr3 | 0.196497 | 4.610973 | 3.035357 | 0.086358 | 0.101338 | Infection | Liver |
| Tlr4 | 1.754785 | 4.559608 | 209.2351 | 1.27E-21 | 5.29E-21 | Infection | Liver |
| Tlr5 | -1.84406 | 5.386857 | 310.3885 | 4.71E-26 | 3.01E-25 | Healthy control | Liver |
| Tlr6 | 1.686412 | 5.052655 | 373.3361 | 3.78E-28 | 3.06E-27 | Infection | Liver |
| Tlr7 | 1.503615 | 4.385978 | 99.66439 | 1.44E-14 | 3.70E-14 | Infection | Liver |
| Tlr8 | 0.900406 | 4.074941 | 35.2474 | 1.39E-07 | 2.50E-07 | Infection | Liver |
| Tlr9 | 1.997317 | 3.959562 | 180.7464 | 2.99E-20 | 1.13E-19 | Infection | Liver |
| Tlx1 | 1.112261 | 1.209425 | 4.201262 | 0.052646 | 0.063413 | Infection | Liver |
| Tlx3 | -0.2537 | 1.220565 | 0.201714 | 0.656971 | 0.679989 | Healthy control | Liver |
| Tmco5 | 1.701878 | 1.300111 | 7.891734 | 0.007061 | 0.009404 | Infection | Liver |
| Tmem106b | -0.88422 | 8.37835 | 250.1512 | 1.10E-23 | 5.53E-23 | Healthy control | Liver |
| Cpa4 | 2.108894 | 1.492881 | 17.15068 | 8.38E-05 | 0.000129 | Infection | Liver |
| Tmem109 | 0.532961 | 7.867561 | 175.9242 | 6.00E-20 | 2.24E-19 | Infection | Liver |
| Tdrd1 | 2.108105 | 1.50261 | 14.13117 | 0.000317 | 0.00047 | Infection | Liver |
| Tmem120a | -2.14787 | 7.868733 | 988.006 | 2.31E-40 | 1.18E-38 | Healthy control | Liver |
| Tmem163 | -1.32688 | 1.98563 | 21.98804 | 8.56E-06 | 1.40E-05 | Healthy control | Liver |
| G6pc2 | 2.104343 | 1.424688 | 10.44386 | 0.001918 | 0.00268 | Infection | Liver |
| Tmem259 | -0.68261 | 8.33994 | 341.0803 | 3.36E-27 | 2.40E-26 | Healthy control | Liver |
| Tmem35a | -1.21396 | 3.022086 | 32.55809 | 2.04E-07 | 3.63E-07 | Healthy control | Liver |
| Tmem41b | -0.84475 | 5.254091 | 76.01866 | 1.98E-12 | 4.53E-12 | Healthy control | Liver |
| Tmem43 | 0.588883 | 4.896127 | 35.04011 | 1.45E-07 | 2.61E-07 | Infection | Liver |
| Tmem63b | -1.40087 | 8.209629 | 988.045 | 2.27E-40 | 1.17E-38 | Healthy control | Liver |
| Tmem94 | -1.48702 | 7.816054 | 1290.307 | 7.57E-44 | 8.81E-42 | Healthy control | Liver |
| Tmem98 | -1.37315 | 6.601605 | 222.7531 | 2.08E-22 | 9.35E-22 | Healthy control | Liver |
| Tmigd1 | 1.222266 | 1.601043 | 4.415329 | 0.037761 | 0.046445 | Infection | Liver |
| Zfp57 | 2.101071 | 2.893775 | 70.01162 | 9.22E-13 | 2.15E-12 | Infection | Liver |
| Tmprss2 | 0.798037 | 6.347414 | 42.63405 | 1.28E-08 | 2.43E-08 | Infection | Liver |
| Tmprss4 | 0.787638 | 2.038385 | 5.886221 | 0.016772 | 0.021578 | Infection | Liver |
| Gpr39 | 2.100061 | 6.851709 | 342.2696 | 3.12E-27 | 2.25E-26 | Infection | Liver |
| 8-Sep | 2.099833 | 6.68667 | 259.5596 | 5.62E-24 | 2.92E-23 | Infection | Liver |
| Tnfaip3 | 1.454014 | 5.141333 | 168.3325 | 1.85E-19 | 6.59E-19 | Infection | Liver |
| Tnfrsf11a | 1.596406 | 5.081114 | 206.1025 | 1.57E-21 | 6.53E-21 | Infection | Liver |
| Tnfrsf11b | -1.32781 | 5.536642 | 127.3755 | 8.96E-17 | 2.64E-16 | Healthy control | Liver |
| Rims2 | 2.098899 | 1.649925 | 22.11107 | 8.44E-06 | 1.38E-05 | Infection | Liver |
| Cd247 | 2.098106 | 3.050929 | 87.28013 | 1.87E-14 | 4.78E-14 | Infection | Liver |
| Tnfrsf14 | 0.14231 | 7.713002 | 7.468864 | 0.008125 | 0.010758 | Infection | Liver |
| Pax1 | 2.096665 | 1.304193 | 9.58374 | 0.003264 | 0.004473 | Infection | Liver |
| Tnfrsf1a | 0.564719 | 9.230717 | 163.9077 | 3.10E-19 | 1.09E-18 | Infection | Liver |
| Tnfrsf1b | -0.07978 | 7.729678 | 1.343316 | 0.250798 | 0.276358 | Healthy control | Liver |
| Tnfrsf4 | 0.881554 | 3.062568 | 20.41592 | 2.52E-05 | 4.00E-05 | Infection | Liver |
| Gpr84 | 2.095837 | 1.591497 | 20.08337 | 2.19E-05 | 3.50E-05 | Infection | Liver |
| Tnfsf10 | 0.362968 | 5.564144 | 11.27421 | 0.001335 | 0.001887 | Infection | Liver |
| Ckb | 2.09519 | 7.077598 | 277.2974 | 1.03E-24 | 5.78E-24 | Infection | Liver |
| Tnfsf12 | 0.250349 | 6.01294 | 8.524604 | 0.004848 | 0.006546 | Infection | Liver |
| Tnfsf13 | 1.392396 | 6.06477 | 245.9757 | 1.74E-23 | 8.51E-23 | Infection | Liver |
| Cldn10 | 2.094005 | 1.476267 | 13.72893 | 0.000417 | 0.000613 | Infection | Liver |
| Hist2h2bb | 2.091935 | 8.370988 | 300.2088 | 1.87E-25 | 1.11E-24 | Infection | Liver |
| Smgc | 2.089469 | 1.419234 | 12.01672 | 0.000928 | 0.001329 | Infection | Liver |
| Cdh26 | 2.089421 | 1.494976 | 17.07717 | 8.58E-05 | 0.000132 | Infection | Liver |
| Tnks2 | 0.624308 | 8.401897 | 314.0027 | 3.04E-26 | 1.99E-25 | Infection | Liver |

| | | | | | | | |
|---|---|---|---|---|---|---|---|
| Tnnc1 | 0.950738 | 2.034332 | 8.757192 | 0.003719 | 0.005066 | Infection | Liver |
| Tnni3k | 0.704181 | 1.304172 | 1.242285 | 0.269924 | 0.295757 | Infection | Liver |
| Il1rapl2 | 2.085553 | 1.464593 | 11.05571 | 0.001379 | 0.001947 | Infection | Liver |
| Tnp1 | 0.429946 | 1.311239 | 0.58398 | 0.447656 | 0.475486 | Infection | Liver |
| Tnpo1 | 0.258385 | 8.807951 | 51.37079 | 9.91E-10 | 1.98E-09 | Infection | Liver |
| Tns2 | -1.20518 | 6.556578 | 164.2718 | 3.00E-19 | 1.05E-18 | Healthy control | Liver |
| Tnxb | 1.31588 | 5.906887 | 33.33677 | 3.08E-07 | 5.41E-07 | Infection | Liver |
| Togaram1 | -0.02085 | 5.8084 | 0.118614 | 0.731685 | 0.751501 | Healthy control | Liver |
| Arrb2 | 2.083771 | 8.514153 | 1164.237 | 1.66E-42 | 1.41E-40 | Infection | Liver |
| Top2b | 0.324119 | 8.112619 | 56.06515 | 2.72E-10 | 5.61E-10 | Infection | Liver |
| Topbp1 | 0.38625 | 5.768565 | 14.78337 | 0.000282 | 0.000419 | Infection | Liver |
| Tor1a | -0.3685 | 8.598028 | 61.01722 | 7.35E-11 | 1.56E-10 | Healthy control | Liver |
| Tor2a | -0.01453 | 7.263117 | 0.059423 | 0.808197 | 0.824671 | Healthy control | Liver |
| Tox | -1.47024 | 5.771972 | 322.703 | 1.71E-26 | 1.14E-25 | Healthy control | Liver |
| Tox3 | 1.577241 | 1.453558 | 11.81667 | 0.000916 | 0.001313 | Infection | Liver |
| Tpbpa | 1.809321 | 1.534959 | 12.10456 | 0.000788 | 0.001134 | Infection | Liver |
| Tpcn1 | -0.88325 | 8.212512 | 467.6844 | 5.88E-31 | 6.44E-30 | Healthy control | Liver |
| Casr | 2.083715 | 1.299283 | 12.69768 | 0.000857 | 0.001232 | Infection | Liver |
| Tph2 | -0.19005 | 1.267205 | 0.1098 | 0.741938 | 0.760783 | Healthy control | Liver |
| Tpi1 | -1.51848 | 9.676525 | 795.4089 | 1.39E-37 | 4.16E-36 | Healthy control | Liver |
| Tppp | -1.8784 | 4.348379 | 146.9029 | 2.01E-18 | 6.63E-18 | Healthy control | Liver |
| Tpr | 0.975323 | 8.405041 | 356.6119 | 1.01E-27 | 7.86E-27 | Infection | Liver |
| Gpc6 | 2.082702 | 5.667671 | 243.2536 | 2.47E-23 | 1.20E-22 | Infection | Liver |
| Cyct | 2.079005 | 1.499386 | 18.30315 | 5.16E-05 | 8.04E-05 | Infection | Liver |
| Traf2 | 0.382984 | 7.19415 | 49.664 | 1.61E-09 | 3.17E-09 | Infection | Liver |
| Traf3 | 0.368686 | 6.048252 | 29.47479 | 9.55E-07 | 1.63E-06 | Infection | Liver |
| Traf3ip2 | 0.446403 | 5.19922 | 27.60315 | 1.87E-06 | 3.15E-06 | Infection | Liver |
| Pdlim4 | 2.076081 | 4.114611 | 239.3688 | 3.77E-23 | 1.79E-22 | Infection | Liver |
| Traf6 | -0.79576 | 6.099895 | 174.9943 | 7.01E-20 | 2.60E-19 | Healthy control | Liver |
| Trappc10 | 0.243093 | 7.851762 | 37.79677 | 5.83E-08 | 1.06E-07 | Infection | Liver |
| Gad1 | 2.070197 | 1.470408 | 10.85972 | 0.001529 | 0.00215 | Infection | Liver |
| Treh | -0.483 | 1.837022 | 1.79862 | 0.182451 | 0.20442 | Healthy control | Liver |
| Il27ra | 2.069565 | 5.45412 | 406.8334 | 3.41E-29 | 3.04E-28 | Infection | Liver |
| Fgf17 | 2.068302 | 1.407575 | 16.13652 | 0.000152 | 0.00023 | Infection | Liver |
| Trex1 | -1.18368 | 5.343525 | 145.0096 | 5.43E-18 | 1.72E-17 | Healthy control | Liver |
| Trh | 1.696864 | 1.324422 | 5.707843 | 0.021425 | 0.027201 | Infection | Liver |
| Mymk | 2.068157 | 1.284854 | 14.27472 | 0.000458 | 0.000671 | Infection | Liver |
| Trhr2 | 0.706365 | 1.288942 | 1.565058 | 0.216938 | 0.241064 | Infection | Liver |
| Trib2 | 1.652286 | 3.570993 | 110.4532 | 1.15E-15 | 3.15E-15 | Infection | Liver |
| Trib3 | -0.48796 | 5.65602 | 13.35058 | 0.000527 | 0.000769 | Healthy control | Liver |
| Trim17 | 0.843031 | 1.329307 | 2.302186 | 0.134166 | 0.153482 | Infection | Liver |
| Trim2 | -0.79453 | 7.095119 | 170.4707 | 1.26E-19 | 4.58E-19 | Healthy control | Liver |
| Trim21 | 0.241484 | 5.955029 | 14.0145 | 0.000394 | 0.00058 | Infection | Liver |
| Trim24 | -0.84716 | 7.852893 | 191.9325 | 7.64E-21 | 3.00E-20 | Healthy control | Liver |
| Trim27 | 0.342381 | 7.930605 | 52.22869 | 7.79E-10 | 1.57E-09 | Infection | Liver |
| Trim28 | -0.51555 | 8.470146 | 80.92767 | 6.24E-13 | 1.48E-12 | Healthy control | Liver |
| Cd5 | 2.067139 | 2.446009 | 45.21846 | 8.80E-10 | 1.76E-09 | Infection | Liver |
| Trim32 | -0.32477 | 5.068275 | 11.54773 | 0.00118 | 0.001678 | Healthy control | Liver |
| Trim33 | -0.09213 | 6.274104 | 2.508977 | 0.11818 | 0.13641 | Healthy control | Liver |
| Trim36 | 1.445625 | 1.501271 | 10.13619 | 0.001958 | 0.002732 | Infection | Liver |

| | | | | | | | |
|---|---|---|---|---|---|---|---|
| Sh3gl2 | 2.066504 | 1.500915 | 15.10162 | 0.000207 | 0.00031 | Infection | Liver |
| Trim65 | -1.07889 | 5.976009 | 208.4279 | 1.10E-21 | 4.60E-21 | Healthy control | Liver |
| Trim7 | -0.64437 | 6.849911 | 51.28485 | 1.02E-09 | 2.03E-09 | Healthy control | Liver |
| Trip13 | 0.93627 | 4.308122 | 48.35401 | 2.49E-09 | 4.88E-09 | Infection | Liver |
| Trmt13 | 0.084898 | 5.393007 | 1.102332 | 0.297754 | 0.323893 | Infection | Liver |
| Trmt2a | -0.39289 | 5.31572 | 28.83079 | 1.21E-06 | 2.06E-06 | Healthy control | Liver |
| Trp53 | 0.65866 | 8.498848 | 331.0093 | 7.48E-27 | 5.15E-26 | Infection | Liver |
| Trp53bp1 | 0.140642 | 5.668741 | 3.829073 | 0.054787 | 0.065829 | Infection | Liver |
| Trp53cor1 | 0.34712 | 2.394067 | 2.326271 | 0.130761 | 0.149894 | Infection | Liver |
| Trp63 | 1.945264 | 1.583527 | 16.78299 | 8.67E-05 | 0.000133 | Infection | Liver |
| Psmb9 | 2.06407 | 3.882084 | 161.8019 | 3.24E-19 | 1.13E-18 | Infection | Liver |
| Trpa1 | 1.394464 | 1.413001 | 5.967393 | 0.016905 | 0.021727 | Infection | Liver |
| Trpc1 | 1.759546 | 3.388471 | 122.4873 | 7.06E-17 | 2.10E-16 | Infection | Liver |
| Trpc3 | -1.19027 | 2.650746 | 22.26198 | 9.03E-06 | 1.47E-05 | Healthy control | Liver |
| Cfc1 | 2.064036 | 1.293608 | 8.643379 | 0.005228 | 0.007045 | Infection | Liver |
| Trpc5 | 1.788517 | 1.346301 | 8.221555 | 0.005792 | 0.007783 | Infection | Liver |
| Cdh23 | 2.062312 | 1.401432 | 13.23692 | 0.000535 | 0.000779 | Infection | Liver |
| Trpc7 | 1.915238 | 1.273333 | 13.16683 | 0.000767 | 0.001105 | Infection | Liver |
| Trpm1 | 0.32057 | 1.406344 | 0.355555 | 0.552383 | 0.578621 | Infection | Liver |
| Abcc1 | 2.059734 | 5.146672 | 350.4632 | 2.15E-27 | 1.59E-26 | Infection | Liver |
| Trpm5 | 0.57939 | 1.376211 | 1.059302 | 0.306439 | 0.332762 | Infection | Liver |
| Trpm6 | -1.94101 | 2.758234 | 67.37009 | 1.38E-12 | 3.18E-12 | Healthy control | Liver |
| Trpm7 | -0.56806 | 7.174732 | 160.6148 | 4.95E-19 | 1.70E-18 | Healthy control | Liver |
| Trpm8 | -0.80794 | 1.323697 | 2.32318 | 0.131922 | 0.151191 | Healthy control | Liver |
| Trpv1 | -0.98801 | 1.445429 | 4.320277 | 0.040102 | 0.049179 | Healthy control | Liver |
| Trpv2 | 1.063233 | 4.631428 | 72.27427 | 4.87E-12 | 1.09E-11 | Infection | Liver |
| Trpv3 | 0.627193 | 1.326157 | 1.02901 | 0.314289 | 0.340621 | Infection | Liver |
| Kcnc1 | 2.058829 | 1.697412 | 27.58951 | 9.11E-07 | 1.56E-06 | Infection | Liver |
| Trpv5 | 1.066494 | 1.266489 | 3.503761 | 0.069167 | 0.08203 | Infection | Liver |
| Trpv6 | 0.731092 | 1.245629 | 1.335321 | 0.255879 | 0.281338 | Infection | Liver |
| Trrap | -0.16369 | 7.789978 | 13.17673 | 0.000568 | 0.000824 | Healthy control | Liver |
| Tsc1 | -1.01838 | 6.700221 | 229.151 | 1.02E-22 | 4.68E-22 | Healthy control | Liver |
| Tsc2 | -0.7969 | 7.338689 | 208.3707 | 1.05E-21 | 4.41E-21 | Healthy control | Liver |
| Tsc22d3 | -0.40214 | 8.396657 | 9.25165 | 0.003462 | 0.004729 | Healthy control | Liver |
| Tsg101 | 0.159774 | 2.810878 | 0.473886 | 0.493356 | 0.520603 | Infection | Liver |
| Tshb | 1.447814 | 1.276005 | 5.72053 | 0.021559 | 0.02735 | Infection | Liver |
| Tshr | 1.795151 | 1.466805 | 9.36664 | 0.003087 | 0.004241 | Infection | Liver |
| Tshz1 | -1.51353 | 5.877907 | 305.3349 | 7.28E-26 | 4.54E-25 | Healthy control | Liver |
| Tslp | 1.912347 | 2.2666 | 41.1176 | 3.33E-09 | 6.48E-09 | Infection | Liver |
| Tspyl2 | 1.18811 | 6.165712 | 266.9803 | 2.14E-24 | 1.16E-23 | Infection | Liver |
| Ttbk1 | -0.71602 | 2.437548 | 4.887892 | 0.029623 | 0.036833 | Healthy control | Liver |
| Ttc8 | 0.171791 | 5.030447 | 5.70535 | 0.019918 | 0.025397 | Infection | Liver |
| Ly6g6e | 2.05757 | 2.411789 | 57.8133 | 1.05E-11 | 2.32E-11 | Infection | Liver |
| Sort1 | 2.054586 | 4.699539 | 214.67 | 6.91E-22 | 2.98E-21 | Infection | Liver |
| Ttll6 | 0.77165 | 1.253987 | 1.804818 | 0.187084 | 0.209377 | Infection | Liver |
| Ttn | -0.33419 | 1.655212 | 0.661363 | 0.417556 | 0.444836 | Healthy control | Liver |
| Tuba3a | 1.645331 | 1.253208 | 10.83902 | 0.002099 | 0.002919 | Infection | Liver |
| Nr4a1 | 2.052029 | 3.385948 | 83.14591 | 5.83E-14 | 1.46E-13 | Infection | Liver |
| P2rx7 | 2.051283 | 5.694105 | 357.1679 | 1.09E-27 | 8.42E-27 | Infection | Liver |
| St6galnac5 | 2.0493 | 1.470939 | 12.08733 | 0.000865 | 0.001242 | Infection | Liver |

| | | | | | | | |
|---|---|---|---|---|---|---|---|
| Tubg1 | -0.69973 | 6.441552 | 117.5161 | 4.56E-16 | 1.27E-15 | Healthy control | Liver |
| Nxph2 | 2.046575 | 1.502362 | 10.54791 | 0.001697 | 0.00238 | Infection | Liver |
| Tubgcp2 | -0.07334 | 6.223454 | 1.809901 | 0.183317 | 0.205253 | Healthy control | Liver |
| Tufm | -1.25194 | 7.685545 | 360.4726 | 7.63E-28 | 5.99E-27 | Healthy control | Liver |
| Twf1 | -0.93844 | 7.104396 | 258.9089 | 4.64E-24 | 2.44E-23 | Healthy control | Liver |
| Twf2 | 0.725884 | 6.060217 | 57.80245 | 1.72E-10 | 3.57E-10 | Infection | Liver |
| Twist1 | 0.999155 | 2.865995 | 13.12055 | 0.000539 | 0.000785 | Infection | Liver |
| Twist2 | 1.872118 | 1.645117 | 22.61984 | 6.43E-06 | 1.06E-05 | Infection | Liver |
| Twnk | -0.77325 | 5.62915 | 79.07639 | 9.71E-13 | 2.26E-12 | Healthy control | Liver |
| Txnip | -0.10353 | 9.332176 | 1.375345 | 0.245284 | 0.270775 | Healthy control | Liver |
| Tyk2 | -0.0673 | 6.712946 | 1.850626 | 0.178531 | 0.200207 | Healthy control | Liver |
| Tyr | 1.355721 | 1.312148 | 5.042488 | 0.028793 | 0.035838 | Infection | Liver |
| Tyro3 | 1.41724 | 3.661894 | 70.4926 | 6.91E-12 | 1.54E-11 | Infection | Liver |
| Tyrp1 | 1.839477 | 1.41425 | 9.278209 | 0.003227 | 0.004425 | Infection | Liver |
| Tyw5 | 0.413023 | 6.145939 | 30.90824 | 5.79E-07 | 1.00E-06 | Infection | Liver |
| Uba1y | 0.03714 | 1.184539 | 0.005766 | 0.940608 | 0.94685 | Infection | Liver |
| Uba6 | -0.07529 | 6.194283 | 1.243892 | 0.268936 | 0.294852 | Healthy control | Liver |
| Ubac1 | -0.29222 | 6.975059 | 15.38427 | 0.000218 | 0.000327 | Healthy control | Liver |
| Ubap1 | -0.56454 | 6.873982 | 73.11285 | 3.74E-12 | 8.43E-12 | Healthy control | Liver |
| Notch3 | 2.045052 | 4.524542 | 96.70118 | 2.57E-14 | 6.52E-14 | Infection | Liver |
| Ube2b | 0.487606 | 8.856597 | 89.28116 | 1.03E-13 | 2.54E-13 | Infection | Liver |
| Ube2d1 | 0.287409 | 5.803154 | 20.00021 | 3.28E-05 | 5.18E-05 | Infection | Liver |
| Ube2g1 | -1.40415 | 7.424359 | 724.8195 | 2.15E-36 | 5.38E-35 | Healthy control | Liver |
| Ube2g2 | -0.51307 | 7.045276 | 75.53875 | 2.12E-12 | 4.84E-12 | Healthy control | Liver |
| Ube2i | -0.06474 | 7.93842 | 1.006785 | 0.31949 | 0.345659 | Healthy control | Liver |
| Ube2m | -0.71599 | 7.848278 | 144.4799 | 5.36E-18 | 1.70E-17 | Healthy control | Liver |
| Ube2n | -1.62759 | 4.214975 | 176.1767 | 3.04E-20 | 1.15E-19 | Healthy control | Liver |
| Ube2r2 | -0.53232 | 8.716897 | 59.58124 | 1.07E-10 | 2.24E-10 | Healthy control | Liver |
| Ube2u | 1.367457 | 1.361831 | 6.877796 | 0.010902 | 0.014275 | Infection | Liver |
| Ube3a | -0.16515 | 8.649835 | 7.809664 | 0.006867 | 0.009162 | Healthy control | Liver |
| Ublcp1 | -1.719 | 4.712506 | 267.1629 | 1.70E-24 | 9.33E-24 | Healthy control | Liver |
| Ubr5 | -0.02492 | 8.077225 | 0.396969 | 0.530924 | 0.558362 | Healthy control | Liver |
| Ubr7 | 0.429221 | 7.051989 | 79.83711 | 7.98E-13 | 1.87E-12 | Infection | Liver |
| Ubtf | 0.57545 | 8.902569 | 124.1644 | 1.43E-16 | 4.13E-16 | Infection | Liver |
| Ubxn7 | -0.06943 | 7.416887 | 2.46547 | 0.121353 | 0.139878 | Healthy control | Liver |
| Fstl5 | 2.044224 | 1.358209 | 8.888347 | 0.004172 | 0.005661 | Infection | Liver |
| Ucn3 | 0.964029 | 1.281505 | 2.040849 | 0.160008 | 0.180772 | Infection | Liver |
| Ucp1 | 1.125949 | 1.357872 | 3.606119 | 0.061769 | 0.073764 | Infection | Liver |
| Ucp3 | 1.959917 | 1.389014 | 13.31642 | 0.000593 | 0.00086 | Infection | Liver |
| Ufd1 | -0.24665 | 7.526526 | 21.44924 | 1.85E-05 | 2.97E-05 | Healthy control | Liver |
| Ufl1 | -0.60435 | 7.2948 | 153.4113 | 1.40E-18 | 4.69E-18 | Healthy control | Liver |
| Ptprc | 2.041576 | 6.684559 | 330.5931 | 7.92E-27 | 5.43E-26 | Infection | Liver |
| Chrna4 | 2.039369 | 5.469658 | 128.5467 | 8.43E-17 | 2.48E-16 | Infection | Liver |
| Uhrf2 | 0.121264 | 6.497329 | 6.034028 | 0.016785 | 0.021589 | Infection | Liver |
| Ulk1 | -0.89636 | 8.143783 | 132.2944 | 3.68E-17 | 1.11E-16 | Healthy control | Liver |
| Evx1 | 2.038446 | 1.289058 | 11.5586 | 0.001395 | 0.001969 | Infection | Liver |
| Rnase1 | 2.036682 | 1.792168 | 26.51764 | 1.17E-06 | 2.00E-06 | Infection | Liver |
| Unc93b1 | -0.46291 | 7.530499 | 62.27289 | 5.33E-11 | 1.14E-10 | Healthy control | Liver |
| Uncx | 1.84193 | 1.379319 | 8.673908 | 0.004489 | 0.006077 | Infection | Liver |
| Ung | 0.049277 | 5.055719 | 0.24856 | 0.619825 | 0.643877 | Infection | Liver |

| | | | | | | | |
|---|---|---|---|---|---|---|---|
| Upk1b | 1.947043 | 1.424646 | 9.540261 | 0.002833 | 0.003903 | Infection | Liver |
| Upk3b | 1.873223 | 1.709163 | 14.83993 | 0.000199 | 0.000299 | Infection | Liver |
| Uri1 | -0.62763 | 7.219589 | 172.9774 | 8.94E-20 | 3.29E-19 | Healthy control | Liver |
| Urm1 | -0.89847 | 4.182852 | 55.05843 | 3.43E-10 | 7.03E-10 | Healthy control | Liver |
| Usp14 | -0.68772 | 7.051723 | 188.7009 | 1.15E-20 | 4.49E-20 | Healthy control | Liver |
| Usp18 | 0.154582 | 5.906031 | 1.307569 | 0.257141 | 0.282663 | Infection | Liver |
| Usp2 | 1.127513 | 3.773289 | 54.92048 | 3.96E-10 | 8.09E-10 | Infection | Liver |
| Usp22 | -1.38914 | 6.053444 | 260.593 | 4.21E-24 | 2.23E-23 | Healthy control | Liver |
| Usp25 | 0.292319 | 8.019133 | 51.67237 | 9.10E-10 | 1.82E-09 | Infection | Liver |
| Usp3 | 0.44437 | 7.076099 | 114.0688 | 8.38E-16 | 2.31E-15 | Infection | Liver |
| Usp32 | 0.239675 | 7.334007 | 39.63252 | 3.25E-08 | 6.00E-08 | Infection | Liver |
| Usp47 | -0.30179 | 7.87032 | 49.5903 | 1.64E-09 | 3.24E-09 | Healthy control | Liver |
| Usp50 | 1.293557 | 1.301887 | 5.674759 | 0.020963 | 0.026669 | Infection | Liver |
| Usp54 | 0.09473 | 4.219561 | 0.617974 | 0.434762 | 0.462379 | Infection | Liver |
| Usp7 | 0.245666 | 7.739943 | 34.10271 | 1.95E-07 | 3.48E-07 | Infection | Liver |
| Utp25 | -0.90622 | 5.913177 | 124.0262 | 1.51E-16 | 4.38E-16 | Healthy control | Liver |
| Utrn | -0.67638 | 7.097666 | 159.9761 | 5.43E-19 | 1.86E-18 | Healthy control | Liver |
| Uts2 | 1.548423 | 1.280342 | 6.186647 | 0.016956 | 0.021787 | Infection | Liver |
| Dsg1a | 2.035261 | 1.831058 | 17.67144 | 5.36E-05 | 8.33E-05 | Infection | Liver |
| Vamp2 | 0.655103 | 6.975138 | 122.2992 | 1.97E-16 | 5.65E-16 | Infection | Liver |
| Vamp3 | 1.158931 | 6.813663 | 297.7065 | 1.25E-25 | 7.54E-25 | Infection | Liver |
| Vangl1 | 0.466342 | 5.3778 | 32.40961 | 3.48E-07 | 6.09E-07 | Infection | Liver |
| Vangl2 | 1.089554 | 3.259712 | 32.72433 | 2.84E-07 | 5.01E-07 | Infection | Liver |
| Vasp | 1.682593 | 4.71264 | 229.0538 | 1.33E-22 | 6.05E-22 | Infection | Liver |
| Vat1 | 0.67611 | 8.101135 | 39.04966 | 4.03E-08 | 7.41E-08 | Infection | Liver |
| Ptger1 | 2.034445 | 2.007117 | 35.52413 | 2.43E-08 | 4.51E-08 | Infection | Liver |
| Vav2 | -0.59201 | 5.609941 | 50.21399 | 1.39E-09 | 2.76E-09 | Healthy control | Liver |
| Vax1 | 1.756879 | 1.262744 | 11.00504 | 0.002054 | 0.002859 | Infection | Liver |
| Irak3 | 2.034051 | 3.724749 | 211.9947 | 3.69E-22 | 1.62E-21 | Infection | Liver |
| Vcam1 | 1.374437 | 5.98084 | 183.3887 | 2.31E-20 | 8.81E-20 | Infection | Liver |
| Large1 | 2.033718 | 5.965019 | 384.9888 | 1.38E-28 | 1.16E-27 | Infection | Liver |
| Vcl | 0.135788 | 7.806844 | 3.771753 | 0.056572 | 0.067881 | Infection | Liver |
| Vcp | -1.20577 | 8.418784 | 353.2616 | 1.31E-27 | 1.00E-26 | Healthy control | Liver |
| Vdac1 | -1.25824 | 9.538203 | 588.3773 | 8.79E-34 | 1.33E-32 | Healthy control | Liver |
| Vdac2 | -0.25425 | 8.051397 | 24.94789 | 4.90E-06 | 8.10E-06 | Healthy control | Liver |
| Zic3 | 2.033704 | 1.507623 | 13.89982 | 0.000362 | 0.000535 | Infection | Liver |
| Vegfa | -0.08609 | 8.976786 | 1.320135 | 0.254885 | 0.280491 | Healthy control | Liver |
| Vegfb | -0.63747 | 6.840402 | 91.67235 | 6.28E-14 | 1.56E-13 | Healthy control | Liver |
| Vegfc | 0.239312 | 4.468366 | 3.022581 | 0.087012 | 0.102034 | Infection | Liver |
| Vegfd | 1.435482 | 4.312415 | 85.16216 | 2.84E-13 | 6.85E-13 | Infection | Liver |
| Vgll2 | 0.013919 | 1.207889 | 0.000582 | 0.980968 | 0.983129 | Infection | Liver |
| Vhl | 0.306112 | 4.99797 | 10.13428 | 0.002261 | 0.003138 | Infection | Liver |
| Vil1 | 0.744237 | 2.403231 | 7.413617 | 0.007746 | 0.01028 | Infection | Liver |
| Vip | 0.800123 | 1.398811 | 2.368698 | 0.127484 | 0.146487 | Infection | Liver |
| Vipas39 | -0.21734 | 5.00668 | 8.146678 | 0.005832 | 0.007829 | Healthy control | Liver |
| Vipr1 | -1.83307 | 6.78297 | 429.202 | 6.81E-30 | 6.58E-29 | Healthy control | Liver |
| Vipr2 | 1.700919 | 1.399709 | 11.20355 | 0.001343 | 0.001898 | Infection | Liver |
| Virma | 0.068877 | 7.05601 | 1.678121 | 0.199873 | 0.222744 | Infection | Liver |
| Vit | 1.23612 | 1.385887 | 3.922936 | 0.051584 | 0.062209 | Infection | Liver |
| Vldlr | -0.91247 | 6.37577 | 19.48302 | 4.40E-05 | 6.89E-05 | Healthy control | Liver |

| | | | | | | | |
|---|---|---|---|---|---|---|---|
| Vps13a | 0.404641 | 6.069368 | 33.7621 | 2.19E-07 | 3.90E-07 | Infection | Liver |
| Vps25 | -0.08801 | 6.996891 | 1.726049 | 0.193653 | 0.216245 | Healthy control | Liver |
| Vps26b | -0.56011 | 6.277063 | 92.03978 | 5.85E-14 | 1.46E-13 | Healthy control | Liver |
| Vps28 | -1.33998 | 8.015501 | 560.8584 | 3.48E-33 | 4.97E-32 | Healthy control | Liver |
| Vps51 | -0.05904 | 6.706172 | 1.383574 | 0.243894 | 0.269343 | Healthy control | Liver |
| Vps54 | -0.17719 | 6.715728 | 11.55563 | 0.001173 | 0.001669 | Healthy control | Liver |
| Vps72 | -1.06063 | 6.565039 | 337.4801 | 4.60E-27 | 3.23E-26 | Healthy control | Liver |
| Vps9d1 | -0.86595 | 6.339346 | 176.8644 | 5.42E-20 | 2.03E-19 | Healthy control | Liver |
| Vsig1 | 1.834117 | 1.277163 | 8.82953 | 0.004703 | 0.006362 | Infection | Liver |
| Vsir | 1.410731 | 5.655335 | 125.4295 | 1.19E-16 | 3.46E-16 | Infection | Liver |
| Hcrtr2 | 2.033399 | 1.440513 | 10.99321 | 0.00141 | 0.001989 | Infection | Liver |
| Egr1 | 2.031837 | 6.561581 | 81.22013 | 1.16E-12 | 2.69E-12 | Infection | Liver |
| Vstm5 | 0.590681 | 2.39829 | 5.783044 | 0.018245 | 0.02333 | Infection | Liver |
| Vsx1 | 1.437841 | 1.311732 | 5.954218 | 0.018089 | 0.023154 | Infection | Liver |
| Vsx2 | 1.171014 | 1.256578 | 4.032275 | 0.051562 | 0.062209 | Infection | Liver |
| Vtcn1 | -0.38428 | 5.306016 | 11.3446 | 0.001294 | 0.001831 | Healthy control | Liver |
| Vtn | -0.97074 | 12.54315 | 79.64712 | 8.31E-13 | 1.95E-12 | Healthy control | Liver |
| Vwa8 | -1.36466 | 9.437978 | 566.9607 | 2.54E-33 | 3.65E-32 | Healthy control | Liver |
| Vwc2l | 1.545876 | 1.297649 | 6.891054 | 0.011721 | 0.015292 | Infection | Liver |
| Vwf | 1.886152 | 5.460918 | 51.5592 | 1.39E-09 | 2.75E-09 | Infection | Liver |
| Vxn | 1.43912 | 1.28841 | 7.136288 | 0.010342 | 0.013571 | Infection | Liver |
| Wap | 1.598214 | 1.335158 | 6.799867 | 0.01156 | 0.015105 | Infection | Liver |
| Wapl | -0.56635 | 8.560746 | 114.5033 | 7.73E-16 | 2.14E-15 | Healthy control | Liver |
| Opalin | 2.03172 | 1.335054 | 11.19376 | 0.001507 | 0.002123 | Infection | Liver |
| Washc1 | -0.4429 | 6.891892 | 52.52124 | 7.19E-10 | 1.45E-09 | Healthy control | Liver |
| Washc2 | 0.36353 | 8.166108 | 60.87651 | 7.63E-11 | 1.62E-10 | Infection | Liver |
| Washc3 | -1.03904 | 6.169736 | 292.8107 | 2.01E-25 | 1.19E-24 | Healthy control | Liver |
| Washc4 | -0.4925 | 6.489507 | 95.36978 | 2.96E-14 | 7.51E-14 | Healthy control | Liver |
| Washc5 | 0.268628 | 6.805233 | 31.33711 | 4.99E-07 | 8.67E-07 | Infection | Liver |
| Wasl | -1.05431 | 7.849672 | 330.5419 | 7.80E-27 | 5.36E-26 | Healthy control | Liver |
| Wdr34 | -1.58479 | 4.420529 | 151.8271 | 1.40E-18 | 4.69E-18 | Healthy control | Liver |
| Wdr36 | -0.34302 | 6.860874 | 62.87215 | 4.58E-11 | 9.80E-11 | Healthy control | Liver |
| Wdr37 | 0.112906 | 5.16617 | 1.539609 | 0.219269 | 0.243492 | Infection | Liver |
| Wdr4 | -0.15961 | 5.248459 | 2.9849 | 0.088935 | 0.104118 | Healthy control | Liver |
| Wdr45 | -0.72169 | 8.089406 | 215.5631 | 4.56E-22 | 1.99E-21 | Healthy control | Liver |
| Wdr5 | -0.5152 | 7.091664 | 103.009 | 6.56E-15 | 1.72E-14 | Healthy control | Liver |
| Wdr59 | -0.71792 | 6.610221 | 158.0173 | 7.24E-19 | 2.46E-18 | Healthy control | Liver |
| Wdr61 | -2.1036 | 7.470512 | 877.7133 | 7.90E-39 | 2.91E-37 | Healthy control | Liver |
| Wdr62 | 0.259153 | 4.509987 | 7.902092 | 0.006577 | 0.008799 | Infection | Liver |
| Wdr82 | 0.105415 | 7.127276 | 2.791618 | 0.09969 | 0.116085 | Infection | Liver |
| Wdr83 | -0.44039 | 5.873054 | 41.42649 | 1.87E-08 | 3.49E-08 | Healthy control | Liver |
| Wdtc1 | -0.57592 | 8.608898 | 71.34808 | 5.66E-12 | 1.27E-11 | Healthy control | Liver |
| Wee1 | 1.707389 | 4.964129 | 196.723 | 5.06E-21 | 2.02E-20 | Infection | Liver |
| Wfdc18 | 0.849996 | 2.901915 | 19.74262 | 3.14E-05 | 4.96E-05 | Infection | Liver |
| Lta | 2.031003 | 1.808507 | 27.99092 | 6.17E-07 | 1.07E-06 | Infection | Liver |
| Wif1 | -0.68254 | 1.879802 | 3.299004 | 0.07193 | 0.085166 | Healthy control | Liver |
| Wipf1 | 1.564427 | 4.923318 | 216.8338 | 4.72E-22 | 2.06E-21 | Infection | Liver |
| Wiz | -0.4739 | 7.071761 | 54.39459 | 4.29E-10 | 8.74E-10 | Healthy control | Liver |
| Gmds | 2.030515 | 3.928994 | 230.5684 | 7.08E-23 | 3.29E-22 | Infection | Liver |
| Wnt1 | 0.664234 | 1.183786 | 2.492236 | 0.136358 | 0.15574 | Infection | Liver |

| | | | | | | | |
|---|---|---|---|---|---|---|---|
| Wnt10a | 1.542698 | 1.63465 | 11.14607 | 0.001192 | 0.001693 | Infection | Liver |
| Wnt10b | 0.733312 | 1.336167 | 2.036412 | 0.158454 | 0.179097 | Infection | Liver |
| Cdhr1 | 2.027726 | 1.496092 | 18.01213 | 5.90E-05 | 9.15E-05 | Infection | Liver |
| Lamp3 | 2.026017 | 1.551065 | 14.56439 | 0.000256 | 0.000381 | Infection | Liver |
| Wnt2 | -1.97324 | 3.923058 | 105.672 | 1.07E-15 | 2.94E-15 | Healthy control | Liver |
| Tmprss15 | 2.025411 | 1.343294 | 10.98188 | 0.001678 | 0.002354 | Infection | Liver |
| Wnt3 | 1.033205 | 1.336301 | 2.664463 | 0.107865 | 0.124965 | Infection | Liver |
| Wnt3a | 0.643959 | 1.244338 | 1.110551 | 0.299446 | 0.325592 | Infection | Liver |
| Wnt4 | 1.975838 | 5.006546 | 41.17133 | 2.72E-08 | 5.03E-08 | Infection | Liver |
| Wnt5a | 0.304722 | 4.990724 | 3.927106 | 0.051876 | 0.062515 | Infection | Liver |
| Wnt5b | -1.60164 | 3.835902 | 130.5403 | 1.62E-17 | 5.00E-17 | Healthy control | Liver |
| Fam49b | 2.021427 | 7.351205 | 1055.659 | 3.18E-41 | 2.04E-39 | Infection | Liver |
| Wnt7a | -0.03039 | 1.29652 | 0.003456 | 0.95332 | 0.957912 | Healthy control | Liver |
| Folr1 | 2.02098 | 1.956517 | 41.03127 | 3.37E-09 | 6.56E-09 | Infection | Liver |
| Wnt8a | -0.03746 | 1.376626 | 0.006155 | 0.937645 | 0.944247 | Healthy control | Liver |
| Wnt8b | 0.98685 | 1.210074 | 3.731186 | 0.065362 | 0.077701 | Infection | Liver |
| Wnt9a | 0.756922 | 3.000232 | 15.30254 | 0.000213 | 0.000318 | Infection | Liver |
| Wnt9b | -1.42438 | 2.583789 | 19.15085 | 3.02E-05 | 4.78E-05 | Healthy control | Liver |
| Wrap53 | 0.268199 | 5.780419 | 17.86026 | 7.78E-05 | 0.00012 | Infection | Liver |
| Wrap73 | -0.56588 | 6.325695 | 73.05593 | 3.80E-12 | 8.57E-12 | Healthy control | Liver |
| Wrnip1 | -1.05685 | 8.575387 | 455.7749 | 1.21E-30 | 1.27E-29 | Healthy control | Liver |
| Wsb1 | 1.071672 | 6.749313 | 150.3124 | 2.23E-18 | 7.29E-18 | Infection | Liver |
| Vav1 | 2.019652 | 6.375225 | 424.7952 | 8.88E-30 | 8.44E-29 | Infection | Liver |
| Wtip | 0.913246 | 4.848002 | 46.70515 | 3.89E-09 | 7.56E-09 | Infection | Liver |
| Wwox | -0.65572 | 7.461438 | 85.98146 | 2.08E-13 | 5.05E-13 | Healthy control | Liver |
| Wwtr1 | 1.827035 | 5.894337 | 346.2167 | 2.42E-27 | 1.77E-26 | Infection | Liver |
| Xcl1 | 1.464916 | 3.273198 | 33.72362 | 1.83E-07 | 3.27E-07 | Infection | Liver |
| Ntng1 | 2.018428 | 1.383868 | 10.21246 | 0.002269 | 0.003149 | Infection | Liver |
| Xiap | -0.71571 | 8.299758 | 100.0784 | 1.15E-14 | 2.99E-14 | Healthy control | Liver |
| Xist | 0.766741 | 9.658764 | 6.263939 | 0.015153 | 0.01957 | Infection | Liver |
| Xkr6 | 0.52486 | 1.981335 | 2.256066 | 0.135716 | 0.155113 | Infection | Liver |
| Xpa | -0.19208 | 6.678403 | 8.917651 | 0.004012 | 0.005454 | Healthy control | Liver |
| Xpo1 | 0.662299 | 7.276207 | 215.7973 | 4.45E-22 | 1.94E-21 | Infection | Liver |
| Xrcc1 | -0.28253 | 6.198941 | 19.23332 | 4.46E-05 | 6.97E-05 | Healthy control | Liver |
| Xrcc2 | -0.28084 | 3.387274 | 3.969969 | 0.050478 | 0.061023 | Healthy control | Liver |
| Xrcc4 | -0.01525 | 6.066061 | 0.040037 | 0.842049 | 0.85503 | Healthy control | Liver |
| Xrcc5 | -1.7801 | 5.576277 | 330.766 | 9.35E-27 | 6.35E-26 | Healthy control | Liver |
| Xrcc6 | -0.95649 | 6.682413 | 221.6876 | 2.32E-22 | 1.03E-21 | Healthy control | Liver |
| Yap1 | -1.25341 | 5.815178 | 271.8728 | 1.45E-24 | 8.02E-24 | Healthy control | Liver |
| Yars | 0.060074 | 7.760731 | 1.371385 | 0.245958 | 0.271443 | Infection | Liver |
| Ybx3 | -0.34144 | 7.271085 | 9.137276 | 0.003623 | 0.004943 | Healthy control | Liver |
| Yeats2 | 0.02966 | 5.244802 | 0.144182 | 0.705433 | 0.726475 | Infection | Liver |
| Yeats4 | 0.1956 | 7.646151 | 16.29291 | 0.000149 | 0.000225 | Infection | Liver |
| Yes1 | -0.7102 | 6.783896 | 169.1466 | 1.51E-19 | 5.44E-19 | Healthy control | Liver |
| Yif1a | -0.48655 | 5.560106 | 38.07852 | 5.37E-08 | 9.80E-08 | Healthy control | Liver |
| Yipf1 | -0.43955 | 5.697768 | 33.14994 | 2.70E-07 | 4.77E-07 | Healthy control | Liver |
| Ywhae | -0.53497 | 9.527651 | 75.55105 | 2.11E-12 | 4.81E-12 | Healthy control | Liver |
| Ywhag | -0.19261 | 8.862354 | 26.37251 | 2.90E-06 | 4.84E-06 | Healthy control | Liver |
| Ywhaz | 0.847145 | 9.865325 | 185.6557 | 1.68E-20 | 6.49E-20 | Infection | Liver |
| Yy1 | 0.170189 | 8.632621 | 22.11044 | 1.44E-05 | 2.31E-05 | Infection | Liver |

| | | | | | | | |
|---|---|---|---|---|---|---|---|
| Zap70 | 0.013008 | 5.692888 | 0.012971 | 0.909685 | 0.918122 | Infection | Liver |
| Zbp1 | 1.163876 | 6.208423 | 83.66993 | 3.44E-13 | 8.27E-13 | Infection | Liver |
| Zbtb16 | 0.815899 | 8.44476 | 16.6735 | 0.000137 | 0.000208 | Infection | Liver |
| Zbtb4 | 0.772938 | 5.090424 | 103.1337 | 6.70E-15 | 1.75E-14 | Infection | Liver |
| Zbtb46 | -0.04265 | 5.452918 | 0.201483 | 0.655062 | 0.678581 | Healthy control | Liver |
| Zbtb7a | -0.76551 | 7.106498 | 210.9797 | 7.76E-22 | 3.32E-21 | Healthy control | Liver |
| Zbtb7b | -1.12893 | 6.361638 | 231.3611 | 8.14E-23 | 3.77E-22 | Healthy control | Liver |
| Prl2c2 | 2.017786 | 1.58574 | 13.46048 | 0.000406 | 0.000597 | Infection | Liver |
| Zc3h11a | -0.17023 | 8.174384 | 8.81074 | 0.004222 | 0.005727 | Healthy control | Liver |
| Slamf9 | 2.014059 | 3.435492 | 104.7302 | 1.85E-15 | 5.00E-15 | Infection | Liver |
| Zc3h14 | -0.47675 | 6.345851 | 86.10802 | 2.03E-13 | 4.94E-13 | Healthy control | Liver |
| Zc3hc1 | -0.94374 | 6.546741 | 220.8539 | 2.55E-22 | 1.14E-21 | Healthy control | Liver |
| Zcchc12 | 1.685823 | 1.310317 | 6.833648 | 0.011672 | 0.01524 | Infection | Liver |
| Zdhhc18 | -0.60111 | 6.177389 | 80.39137 | 7.10E-13 | 1.68E-12 | Healthy control | Liver |
| Zdhhc2 | 0.63744 | 3.844728 | 18.5582 | 5.93E-05 | 9.20E-05 | Infection | Liver |
| Zdhhc6 | -1.47701 | 8.318281 | 625.1948 | 1.54E-34 | 2.67E-33 | Healthy control | Liver |
| Zdhhc7 | -0.9643 | 6.628108 | 362.2859 | 6.78E-28 | 5.35E-27 | Healthy control | Liver |
| Zdhhc8 | 0.442318 | 6.675302 | 39.85474 | 3.03E-08 | 5.60E-08 | Infection | Liver |
| Zeb1 | -0.39388 | 6.766437 | 47.44785 | 3.05E-09 | 5.97E-09 | Healthy control | Liver |
| Zeb2 | 0.822484 | 7.25277 | 69.77526 | 8.26E-12 | 1.84E-11 | Infection | Liver |
| Zfand2a | 0.61743 | 5.478687 | 69.81623 | 8.29E-12 | 1.85E-11 | Infection | Liver |
| Zfand2b | -0.84576 | 7.140003 | 231.0305 | 8.23E-23 | 3.81E-22 | Healthy control | Liver |
| Zfhx3 | -0.65535 | 7.09017 | 127.349 | 8.36E-17 | 2.47E-16 | Healthy control | Liver |
| Zfp106 | -0.92781 | 7.332144 | 283.8277 | 4.33E-25 | 2.50E-24 | Healthy control | Liver |
| Zfp111 | -0.71453 | 5.401392 | 75.50035 | 2.21E-12 | 5.03E-12 | Healthy control | Liver |
| Zfp148 | -0.36248 | 7.277836 | 50.33774 | 1.33E-09 | 2.64E-09 | Healthy control | Liver |
| Zfp219 | -0.35497 | 6.112122 | 34.90122 | 1.50E-07 | 2.69E-07 | Healthy control | Liver |
| Zfp24 | 0.484623 | 6.798333 | 38.38865 | 4.82E-08 | 8.82E-08 | Infection | Liver |
| Zfp281 | -0.48344 | 6.399916 | 42.07384 | 1.53E-08 | 2.87E-08 | Healthy control | Liver |
| Agtr1b | 2.009456 | 1.338723 | 8.650474 | 0.004747 | 0.00642 | Infection | Liver |
| Zfp503 | 0.865637 | 5.792104 | 43.03009 | 1.14E-08 | 2.16E-08 | Infection | Liver |
| Zfp536 | 1.199016 | 1.223804 | 4.707755 | 0.038843 | 0.047718 | Infection | Liver |
| Diras1 | 2.007348 | 4.144161 | 49.60566 | 1.84E-09 | 3.63E-09 | Infection | Liver |
| Zfp750 | -0.53086 | 4.751213 | 17.46118 | 9.21E-05 | 0.000141 | Healthy control | Liver |
| Zfp819 | 0.55875 | 1.210988 | 0.875474 | 0.360125 | 0.386864 | Infection | Liver |
| Sncg | 2.006201 | 1.681073 | 19.88033 | 2.05E-05 | 3.27E-05 | Infection | Liver |
| Zfpl1 | -0.56581 | 4.053598 | 19.47215 | 4.04E-05 | 6.34E-05 | Healthy control | Liver |
| Lrrc6 | 2.006198 | 1.284488 | 11.80826 | 0.00125 | 0.001772 | Infection | Liver |
| Zfyve9 | -1.02767 | 5.755205 | 197.287 | 4.23E-21 | 1.70E-20 | Healthy control | Liver |
| Zic1 | 1.251407 | 1.220759 | 5.171477 | 0.031298 | 0.03883 | Infection | Liver |
| Zic2 | 1.425157 | 1.228486 | 7.693793 | 0.009539 | 0.01256 | Infection | Liver |
| Lgi3 | 2.005433 | 1.611929 | 22.79301 | 6.29E-06 | 1.04E-05 | Infection | Liver |
| Zic4 | 1.292986 | 1.4058 | 5.919425 | 0.017199 | 0.022077 | Infection | Liver |
| Zkscan1 | -1.16904 | 7.466304 | 423.1248 | 9.58E-30 | 9.07E-29 | Healthy control | Liver |
| Zmpste24 | -0.41059 | 7.930787 | 64.70825 | 2.87E-11 | 6.20E-11 | Healthy control | Liver |
| Zmynd10 | 1.148428 | 1.697904 | 9.073655 | 0.003122 | 0.004286 | Infection | Liver |
| Zmynd19 | 0.602014 | 4.835296 | 42.03564 | 1.58E-08 | 2.96E-08 | Infection | Liver |
| Zp1 | 0.948458 | 1.196757 | 3.887011 | 0.064025 | 0.076221 | Infection | Liver |
| Zp3 | 1.088846 | 1.321936 | 3.617984 | 0.062331 | 0.074364 | Infection | Liver |
| Zscan2 | -0.27449 | 4.903494 | 7.786034 | 0.006959 | 0.009272 | Healthy control | Liver |

| | | | | | | | |
|---|---|---|---|---|---|---|---|
| Zscan21 | -0.12348 | 6.529611 | 4.639868 | 0.035048 | 0.043246 | Healthy control | Liver |
| Zswim5 | -1.5798 | 4.089557 | 110.5963 | 8.01E-16 | 2.21E-15 | Healthy control | Liver |
| Zswim9 | -0.19079 | 4.663733 | 4.267471 | 0.042974 | 0.052483 | Healthy control | Liver |
| Zup1 | 0.590423 | 5.559924 | 62.42492 | 5.18E-11 | 1.11E-10 | Infection | Liver |
| Zyx | -0.01554 | 6.466056 | 5.38E-02 | 0.817282 | 0.832922 | Healthy control | Liver |
| Zzef1 | 1.31E-01 | 7.718387 | 8.559119 | 0.004766 | 0.006443 | Infection | Liver |
| Aqp5 | 2.01E+00 | 1.598114 | 14.84769 | 0.000221 | 0.00033 | Infection | Liver |
| A1cf | -2.56E+00 | 6.977706 | 8.827726 | 0.006625 | 0.023289 | Healthy Control | Intestine |
| A2m | -2.28E+00 | 1.361967 | 0.081651 | 0.777677 | 0.93628 | Healthy Control | Intestine |
| Aatf | 1.52E-01 | 7.785804 | 0.314206 | 0.580827 | 0.807672 | Infection | Intestine |
| Abca1 | 2.82E+00 | 6.954426 | 111.0169 | 5.11E-10 | 1.90E-08 | Infection | Intestine |
| Abca13 | 2.83E+00 | 2.055983 | 1.240267 | 0.276997 | 0.478979 | Infection | Intestine |
| Abca3 | -2.12E+00 | 9.356633 | 83.22749 | 6.25E-09 | 1.50E-07 | Healthy Control | Intestine |
| Abca4 | -3.45E+00 | 1.766961 | 0.211793 | 0.648643 | 0.86016 | Healthy Control | Intestine |
| Abca7 | 5.46E-01 | 8.064698 | 6.117801 | 0.021675 | 0.063268 | Infection | Intestine |
| Abca8a | 6.51E-01 | 6.445064 | 0.460231 | 0.507042 | 0.741745 | Infection | Intestine |
| Abca8b | 2.14E+00 | 4.610628 | 6.305353 | 0.019839 | 0.058802 | Infection | Intestine |
| Abcb1a | -4.70E+00 | 10.41723 | 124.1962 | 7.84E-11 | 4.27E-09 | Healthy Control | Intestine |
| Abcb1b | 1.52E+00 | 3.966711 | 2.608009 | 0.11927 | 0.24857 | Infection | Intestine |
| Abcb4 | -3.27E+00 | 2.049047 | 0.222557 | 0.64131 | 0.855379 | Healthy Control | Intestine |
| Abcb8 | -8.21E-01 | 6.630968 | 2.395604 | 0.134971 | 0.274326 | Healthy Control | Intestine |
| Abcc1 | 8.28E-02 | 7.341113 | 0.065851 | 0.799862 | 0.945927 | Infection | Intestine |
| Abcc6 | -2.08E+00 | 5.737649 | 2.721261 | 0.112628 | 0.237998 | Healthy Control | Intestine |
| Abcc8 | -6.86E+00 | 4.387959 | 1.762452 | 0.199527 | 0.3709 | Healthy Control | Intestine |
| Abcc9 | 1.85E+00 | 5.548512 | 6.581083 | 0.017687 | 0.053499 | Infection | Intestine |
| Abcd2 | 4.90E+00 | 3.690301 | 120.9675 | 2.58E-10 | 1.07E-08 | Infection | Intestine |
| Abcd3 | -1.69E+00 | 10.14242 | 91.85702 | 2.87E-09 | 7.92E-08 | Healthy Control | Intestine |
| Abcf1 | -5.01E-01 | 8.460078 | 4.618735 | 0.043 | 0.110787 | Healthy Control | Intestine |
| Abcg2 | -1.42E+00 | 8.044235 | 10.71919 | 0.003413 | 0.013143 | Healthy Control | Intestine |
| Abcg8 | -3.19E-01 | 4.414022 | 0.019723 | 0.889606 | 0.969599 | Healthy Control | Intestine |
| Abhd17c | -3.88E+00 | 9.712112 | 145.4516 | 1.67E-11 | 1.09E-09 | Healthy Control | Intestine |
| Abhd4 | -1.26E+00 | 8.039783 | 13.41151 | 0.001348 | 0.005928 | Healthy Control | Intestine |
| Abhd5 | -1.44E+00 | 7.518743 | 9.621536 | 0.004954 | 0.018154 | Healthy Control | Intestine |
| Abi1 | 8.92E-01 | 9.41476 | 41.60447 | 1.80E-06 | 1.81E-05 | Infection | Intestine |
| Abi3bp | -1.82E+00 | 7.658607 | 7.630658 | 0.010931 | 0.035442 | Healthy Control | Intestine |
| Abitram | -1.89E+00 | 6.280218 | 4.656274 | 0.04108 | 0.106375 | Healthy Control | Intestine |
| Abl1 | -7.66E-01 | 8.75586 | 11.50177 | 0.00265 | 0.010527 | Healthy Control | Intestine |
| Abracl | -2.24E+00 | 6.502434 | 8.624937 | 0.007271 | 0.025172 | Healthy Control | Intestine |
| Abraxas1 | 6.20E-01 | 5.122854 | 0.641537 | 0.431074 | 0.663374 | Infection | Intestine |
| Abraxas2 | -5.36E-01 | 7.742983 | 4.037832 | 0.056912 | 0.138908 | Healthy Control | Intestine |
| Acaa2 | -1.95E+00 | 9.367769 | 46.34086 | 8.16E-07 | 9.43E-06 | Healthy Control | Intestine |
| Acaca | -1.09E+00 | 8.207303 | 13.59095 | 0.001284 | 0.0057 | Healthy Control | Intestine |
| Acacb | -3.35E+00 | 6.5245 | 11.1754 | 0.002907 | 0.011406 | Healthy Control | Intestine |
| Acan | -3.08E+00 | 1.523244 | 0.155584 | 0.696201 | 0.892492 | Healthy Control | Intestine |
| Ace | -9.95E-01 | 7.2693 | 3.223259 | 0.085818 | 0.192302 | Healthy Control | Intestine |
| Ace2 | -8.44E+00 | 5.965838 | 6.251748 | 0.021334 | 0.062308 | Healthy Control | Intestine |
| Acer1 | -3.53E+00 | 8.440732 | 42.55278 | 9.59E-07 | 1.08E-05 | Healthy Control | Intestine |
| Ache | -7.24E+00 | 4.59852 | 2.544544 | 0.12654 | 0.259933 | Healthy Control | Intestine |
| Ackr1 | -1.23E+00 | 5.108595 | 0.264944 | 0.613871 | 0.834611 | Healthy Control | Intestine |
| Ackr2 | -2.80E+00 | 1.632414 | 0.148759 | 0.702232 | 0.894649 | Healthy Control | Intestine |

| | | | | | | | |
|---|---|---|---|---|---|---|---|
| Ackr3 | -8.41E-03 | 6.885272 | 0.000127 | 0.991117 | 1 | Healthy Control | Intestine |
| Ackr4 | -3.65E-01 | 3.778195 | 0.035534 | 0.852129 | 0.956553 | Healthy Control | Intestine |
| Acly | -1.04E+00 | 11.33349 | 43.18553 | 1.37E-06 | 1.46E-05 | Healthy Control | Intestine |
| Acod1 | 4.77E+00 | 1.363021 | 3.395064 | 0.077758 | 0.178557 | Infection | Intestine |
| Acot6 | -9.57E-01 | 5.474141 | 1.188573 | 0.286501 | 0.49109 | Healthy Control | Intestine |
| Acox1 | -6.33E-01 | 10.06146 | 7.625906 | 0.011445 | 0.036893 | Healthy Control | Intestine |
| Acp2 | -1.92E+00 | 7.597637 | 11.50501 | 0.002427 | 0.009766 | Healthy Control | Intestine |
| Acp5 | -3.71E+00 | 9.447604 | 92.92059 | 1.13E-09 | 3.67E-08 | Healthy Control | Intestine |
| Acr | -2.89E+00 | 1.522093 | 0.107072 | 0.745556 | 0.92349 | Healthy Control | Intestine |
| Acsl1 | -6.22E-02 | 6.512641 | 0.017444 | 0.896076 | 0.969735 | Healthy Control | Intestine |
| Acsl4 | 8.78E-01 | 7.614155 | 9.974557 | 0.004596 | 0.016978 | Infection | Intestine |
| Acsl6 | -4.26E+00 | 2.202995 | 0.356646 | 0.555965 | 0.784337 | Healthy Control | Intestine |
| Acss1 | -1.09E+00 | 9.099195 | 21.22109 | 0.00014 | 0.000827 | Healthy Control | Intestine |
| Acss2 | -3.24E+00 | 10.23405 | 137.9236 | 3.26E-11 | 2.02E-09 | Healthy Control | Intestine |
| Actc1 | 1.39E+00 | 7.636046 | 7.679467 | 0.013121 | 0.041466 | Infection | Intestine |
| Actg2 | 1.70E+00 | 8.740445 | 10.04083 | 0.007712 | 0.026441 | Infection | Intestine |
| Actl6a | -1.45E+00 | 7.076095 | 5.031415 | 0.034448 | 0.091776 | Healthy Control | Intestine |
| Actl6b | 2.95E+00 | 1.828531 | 2.670923 | 0.111887 | 0.236631 | Infection | Intestine |
| Actn1 | 2.60E-01 | 9.479493 | 2.484917 | 0.129344 | 0.264607 | Infection | Intestine |
| Actn2 | 4.97E+00 | 1.320952 | 3.531499 | 0.075688 | 0.174687 | Infection | Intestine |
| Actr1a | -5.67E-01 | 7.472368 | 2.665489 | 0.116541 | 0.24468 | Healthy Control | Intestine |
| Actr1b | -3.55E-01 | 8.560887 | 2.203187 | 0.152064 | 0.301359 | Healthy Control | Intestine |
| Actr2 | -1.08E+00 | 10.79656 | 76.2137 | 1.45E-08 | 3.00E-07 | Healthy Control | Intestine |
| Acvr1 | -1.74E+00 | 6.96729 | 9.107154 | 0.00592 | 0.021131 | Healthy Control | Intestine |
| Acvr1b | -3.11E-01 | 8.599334 | 1.274337 | 0.271213 | 0.471094 | Healthy Control | Intestine |
| Acvr1c | -8.04E+00 | 5.310087 | 6.007808 | 0.023664 | 0.068121 | Healthy Control | Intestine |
| Acvr2a | -4.79E-01 | 7.743303 | 2.770133 | 0.110215 | 0.233687 | Healthy Control | Intestine |
| Acvr2b | -1.06E+00 | 5.431033 | 1.445457 | 0.24074 | 0.429794 | Healthy Control | Intestine |
| Acvrl1 | -6.35E-01 | 8.210615 | 1.273325 | 0.274431 | 0.476092 | Healthy Control | Intestine |
| Ada | -8.82E+00 | 5.99696 | 14.37977 | 0.00116 | 0.005216 | Healthy Control | Intestine |
| Adad1 | 3.88E+00 | 1.363798 | 2.793882 | 0.110505 | 0.234152 | Infection | Intestine |
| Adad2 | -1.52E+00 | 1.275042 | 0.038572 | 0.846592 | 0.954066 | Healthy Control | Intestine |
| Adam10 | -1.75E+00 | 9.243646 | 56.93779 | 1.55E-07 | 2.27E-06 | Healthy Control | Intestine |
| Adam17 | -1.97E+00 | 6.39313 | 7.761388 | 0.010205 | 0.033494 | Healthy Control | Intestine |
| Adam18 | 4.35E+00 | 1.599831 | 8.296886 | 0.006517 | 0.023022 | Infection | Intestine |
| Adam28 | -2.41E+00 | 1.523632 | 0.09114 | 0.765812 | 0.931224 | Healthy Control | Intestine |
| Adamdec1 | -1.66E-01 | 11.19241 | 0.161195 | 0.694825 | 0.891642 | Healthy Control | Intestine |
| Adamts1 | -8.34E-02 | 5.760443 | 0.007794 | 0.930397 | 0.969735 | Healthy Control | Intestine |
| Adamts13 | -5.79E+00 | 3.255286 | 1.297381 | 0.268435 | 0.467555 | Healthy Control | Intestine |
| Adamts14 | 7.08E-01 | 4.107854 | 0.256872 | 0.616789 | 0.837115 | Infection | Intestine |
| Adamts18 | -3.94E+00 | 1.922302 | 0.422779 | 0.520881 | 0.755917 | Healthy Control | Intestine |
| Adamts20 | 2.13E+00 | 2.086366 | 1.155484 | 0.291923 | 0.49859 | Infection | Intestine |
| Adamts3 | 3.81E-01 | 3.849418 | 0.113237 | 0.739494 | 0.920185 | Infection | Intestine |
| Adamts5 | 1.28E-01 | 4.919373 | 0.013942 | 0.906961 | 0.969735 | Infection | Intestine |
| Adamts6 | 4.72E+00 | 4.380876 | 131.8416 | 1.08E-10 | 5.66E-09 | Infection | Intestine |
| Adar | -7.29E-01 | 7.91533 | 4.46603 | 0.046135 | 0.117176 | Healthy Control | Intestine |
| Adarb1 | 2.22E+00 | 3.587794 | 6.26065 | 0.019643 | 0.058393 | Infection | Intestine |
| Adcy1 | 1.43E+00 | 2.465305 | 0.602807 | 0.445371 | 0.679113 | Infection | Intestine |
| Adcy2 | 2.54E+00 | 3.735722 | 9.201105 | 0.005922 | 0.021131 | Infection | Intestine |
| Adcy3 | 1.78E+00 | 4.586789 | 5.463786 | 0.028563 | 0.078999 | Infection | Intestine |

| | | | | | | | |
|---|---|---|---|---|---|---|---|
| Adcy8 | -2.16E+00 | 1.320562 | 0.057586 | 0.812884 | 0.950244 | Healthy Control | Intestine |
| Adcyap1 | -2.36E+00 | 1.403737 | 0.082899 | 0.775565 | 0.935312 | Healthy Control | Intestine |
| Adcyap1r1 | 2.55E+00 | 3.846787 | 6.444967 | 0.018338 | 0.055101 | Infection | Intestine |
| Add2 | -5.06E+00 | 2.577109 | 0.623445 | 0.43778 | 0.670807 | Healthy Control | Intestine |
| Adgra2 | 3.44E+00 | 6.305653 | 125.9179 | 1.60E-10 | 7.48E-09 | Infection | Intestine |
| Adgra3 | -1.15E+00 | 7.791207 | 9.941995 | 0.004409 | 0.016375 | Healthy Control | Intestine |
| Adgrb1 | -4.85E+00 | 2.48286 | 0.766151 | 0.391516 | 0.621456 | Healthy Control | Intestine |
| Adgre1 | 1.95E-01 | 7.436733 | 0.095942 | 0.760226 | 0.928591 | Infection | Intestine |
| Adgre5 | 7.62E-02 | 8.333766 | 0.06597 | 0.79971 | 0.945927 | Infection | Intestine |
| Adgrf1 | 5.69E-01 | 3.487202 | 0.087621 | 0.76979 | 0.932779 | Infection | Intestine |
| Adgrf5 | -1.13E+00 | 6.557041 | 0.503741 | 0.487788 | 0.721592 | Healthy Control | Intestine |
| Adgrg1 | -2.82E+00 | 7.920342 | 28.47386 | 1.63E-05 | 0.000128 | Healthy Control | Intestine |
| Adgrg2 | 2.83E+00 | 2.006459 | 1.710146 | 0.201478 | 0.373694 | Infection | Intestine |
| Adgrg5 | 1.25E+00 | 4.990897 | 1.688849 | 0.206741 | 0.381617 | Infection | Intestine |
| Adgrg6 | -7.65E-01 | 4.087272 | 0.142966 | 0.708927 | 0.897243 | Healthy Control | Intestine |
| Adgrg7 | -5.30E+00 | 8.474717 | 37.26025 | 3.57E-06 | 3.31E-05 | Healthy Control | Intestine |
| Adgrl1 | 1.71E+00 | 6.476638 | 20.82525 | 0.000156 | 0.000907 | Infection | Intestine |
| Adgrl2 | -1.29E+00 | 6.872982 | 6.130709 | 0.020821 | 0.061205 | Healthy Control | Intestine |
| Adgrl3 | 2.30E-01 | 5.372128 | 0.064828 | 0.801192 | 0.946496 | Infection | Intestine |
| Adgrv1 | 1.24E+00 | 2.804495 | 0.388811 | 0.538406 | 0.770735 | Infection | Intestine |
| Adh5 | -2.04E+00 | 9.229659 | 58.85229 | 1.25E-07 | 1.90E-06 | Healthy Control | Intestine |
| Adipoq | -3.59E+00 | 2.028973 | 0.34053 | 0.564445 | 0.792599 | Healthy Control | Intestine |
| Adipor1 | 7.56E-01 | 9.945221 | 44.13649 | 1.16E-06 | 1.28E-05 | Infection | Intestine |
| Adm | 1.95E+00 | 4.122938 | 5.166979 | 0.032502 | 0.087383 | Infection | Intestine |
| Adora1 | -5.02E-01 | 3.839646 | 0.059632 | 0.809338 | 0.94887 | Healthy Control | Intestine |
| Adora2a | 2.92E+00 | 3.683336 | 13.69084 | 0.001211 | 0.005406 | Infection | Intestine |
| Adora2b | 2.31E+00 | 2.657051 | 2.508563 | 0.125552 | 0.258329 | Infection | Intestine |
| Adprhl1 | -4.95E-01 | 1.228692 | 0.008683 | 0.927069 | 0.969735 | Healthy Control | Intestine |
| Adra1a | 2.60E+00 | 2.391301 | 2.272927 | 0.143818 | 0.288442 | Infection | Intestine |
| Adra1b | 3.47E+00 | 1.792797 | 1.963901 | 0.172003 | 0.330917 | Infection | Intestine |
| Adra2a | -2.52E+00 | 5.405699 | 3.770085 | 0.065425 | 0.156034 | Healthy Control | Intestine |
| Adrb1 | 1.07E+00 | 4.197177 | 0.604053 | 0.444531 | 0.678358 | Infection | Intestine |
| Adrb2 | 1.15E+00 | 4.650933 | 1.659076 | 0.210116 | 0.385997 | Infection | Intestine |
| Adrb3 | 2.89E+00 | 1.830297 | 1.70515 | 0.200712 | 0.372687 | Infection | Intestine |
| Adtrp | -3.89E+00 | 7.223209 | 17.9853 | 0.000314 | 0.001696 | Healthy Control | Intestine |
| Aebp2 | 1.18E+00 | 8.177917 | 28.62295 | 2.34E-05 | 0.000174 | Infection | Intestine |
| Afap1l2 | 7.41E-01 | 6.226742 | 1.514187 | 0.231449 | 0.417689 | Infection | Intestine |
| Afdn | -2.38E+00 | 9.138482 | 64.25626 | 5.02E-08 | 9.00E-07 | Healthy Control | Intestine |
| Afg3l2 | -2.08E+00 | 9.23743 | 34.35241 | 6.55E-06 | 5.71E-05 | Healthy Control | Intestine |
| Afp | -9.77E-01 | 1.317821 | 0.017928 | 0.894683 | 0.969735 | Healthy Control | Intestine |
| Ager | 1.95E+00 | 3.024486 | 1.914401 | 0.178892 | 0.341285 | Infection | Intestine |
| Agl | 1.08E+00 | 7.324686 | 17.89856 | 0.00035 | 0.001863 | Infection | Intestine |
| Ago1 | -4.20E-01 | 7.712453 | 1.531796 | 0.228829 | 0.414292 | Healthy Control | Intestine |
| Ago2 | 8.70E-02 | 7.755697 | 0.119846 | 0.732511 | 0.914908 | Infection | Intestine |
| Ago3 | 8.86E-02 | 6.395626 | 0.027865 | 0.868901 | 0.960878 | Infection | Intestine |
| Agpat3 | -4.88E-01 | 9.102204 | 7.025436 | 0.014684 | 0.045461 | Healthy Control | Intestine |
| Agpat4 | -2.89E+00 | 9.879589 | 110.1971 | 3.80E-10 | 1.48E-08 | Healthy Control | Intestine |
| Agps | -1.03E+00 | 8.864923 | 18.83165 | 0.000266 | 0.001462 | Healthy Control | Intestine |
| Agr3 | -6.21E+00 | 3.539653 | 1.529789 | 0.230756 | 0.417192 | Healthy Control | Intestine |
| Agrn | 2.17E-01 | 7.080352 | 0.326277 | 0.573671 | 0.80173 | Infection | Intestine |

| | | | | | | | |
|---|---|---|---|---|---|---|---|
| Agrp | -3.22E+00 | 1.599838 | 0.202182 | 0.655833 | 0.865113 | Healthy Control | Intestine |
| Agt | -2.02E+00 | 6.262145 | 4.520928 | 0.043904 | 0.11271 | Healthy Control | Intestine |
| Agtr1a | -7.11E+00 | 4.635498 | 2.262982 | 0.148313 | 0.295681 | Healthy Control | Intestine |
| Agtr1b | -4.64E+00 | 2.541401 | 0.55578 | 0.464355 | 0.699958 | Healthy Control | Intestine |
| Agtr2 | 4.86E+00 | 1.444146 | 4.170429 | 0.049722 | 0.124393 | Infection | Intestine |
| Agtrap | 9.00E-01 | 6.437837 | 3.584817 | 0.07143 | 0.166859 | Infection | Intestine |
| Agxt2 | 4.79E+00 | 1.444883 | 5.250384 | 0.02932 | 0.080615 | Infection | Intestine |
| Ahi1 | 1.40E+00 | 4.829465 | 5.086193 | 0.03379 | 0.090311 | Infection | Intestine |
| Ahr | -2.19E+00 | 7.055559 | 9.53658 | 0.004989 | 0.01827 | Healthy Control | Intestine |
| Aicda | 4.36E+00 | 1.484611 | 3.985818 | 0.055972 | 0.137284 | Infection | Intestine |
| Aida | -1.32E+00 | 6.923178 | 5.36086 | 0.029491 | 0.081026 | Healthy Control | Intestine |
| Aif1 | 9.23E-01 | 7.597153 | 5.078966 | 0.034974 | 0.092879 | Infection | Intestine |
| Aif1l | 5.07E+00 | 2.957881 | 59.53802 | 7.55E-08 | 1.26E-06 | Infection | Intestine |
| Aifm1 | -1.38E+00 | 7.616401 | 7.098055 | 0.013873 | 0.043294 | Healthy Control | Intestine |
| Aim2 | -2.02E+00 | 8.429055 | 30.51395 | 1.24E-05 | 1.00E-04 | Healthy Control | Intestine |
| Aip | -1.02E+00 | 7.454145 | 6.134121 | 0.021184 | 0.061943 | Healthy Control | Intestine |
| Aire | -3.50E+00 | 1.80243 | 0.183806 | 0.671214 | 0.877539 | Healthy Control | Intestine |
| Ajuba | 3.11E+00 | 4.654072 | 30.49464 | 1.55E-05 | 0.000122 | Infection | Intestine |
| Ak4 | 1.13E+00 | 3.914673 | 0.772105 | 0.388432 | 0.617887 | Infection | Intestine |
| Akap5 | -4.28E+00 | 2.204152 | 0.671814 | 0.420931 | 0.652985 | Healthy Control | Intestine |
| Akap9 | -3.56E-01 | 8.812611 | 2.668034 | 0.116758 | 0.244864 | Healthy Control | Intestine |
| Akip1 | 2.36E-01 | 6.361247 | 0.254817 | 0.61863 | 0.837906 | Infection | Intestine |
| Akr1b3 | 1.41E-01 | 8.862314 | 0.596222 | 0.44832 | 0.682363 | Infection | Intestine |
| Akt1 | -1.72E+00 | 9.206091 | 34.48753 | 6.63E-06 | 5.75E-05 | Healthy Control | Intestine |
| Akt1s1 | -1.28E+00 | 8.158278 | 17.95618 | 0.00033 | 0.001773 | Healthy Control | Intestine |
| Akt2 | -7.35E-01 | 6.632434 | 1.680769 | 0.207453 | 0.382086 | Healthy Control | Intestine |
| Akt3 | 1.53E+00 | 3.812465 | 2.597328 | 0.119861 | 0.24949 | Infection | Intestine |
| Aldh1a2 | 5.09E+00 | 5.971041 | 469.0122 | 3.15E-16 | 1.13E-13 | Infection | Intestine |
| Aldh1a3 | 3.28E+00 | 5.519404 | 41.94505 | 1.71E-06 | 1.75E-05 | Infection | Intestine |
| Aldh1l1 | -1.03E+01 | 7.783753 | 35.68131 | 8.00E-06 | 6.83E-05 | Healthy Control | Intestine |
| Aldh1l2 | 5.43E+00 | 3.098994 | 80.80115 | 7.65E-09 | 1.81E-07 | Infection | Intestine |
| Aldh2 | -2.70E+00 | 8.727488 | 47.30222 | 4.50E-07 | 5.65E-06 | Healthy Control | Intestine |
| Aldh3a1 | -3.39E+00 | 2.128183 | 0.172305 | 0.681754 | 0.883047 | Healthy Control | Intestine |
| Aldh4a1 | -1.63E+00 | 6.728917 | 6.96617 | 0.014438 | 0.044837 | Healthy Control | Intestine |
| Aldh9a1 | -3.13E+00 | 7.451743 | 23.91806 | 5.24E-05 | 0.00035 | Healthy Control | Intestine |
| Aldoa | -1.55E+00 | 11.85887 | 178.4484 | 5.63E-12 | 4.55E-10 | Healthy Control | Intestine |
| Aldoc | -2.77E+00 | 6.14331 | 8.029699 | 0.009526 | 0.031581 | Healthy Control | Intestine |
| Alg13 | 9.52E-01 | 5.49374 | 1.877602 | 0.183858 | 0.348484 | Infection | Intestine |
| Alk | 5.09E+00 | 1.45016 | 4.742032 | 0.038015 | 0.099687 | Infection | Intestine |
| Alkbh5 | -6.28E-01 | 8.126117 | 5.523851 | 0.028121 | 0.078126 | Healthy Control | Intestine |
| Alox12 | -6.98E+00 | 3.982977 | 2.014536 | 0.171443 | 0.330155 | Healthy Control | Intestine |
| Alox12b | -3.57E-01 | 1.228299 | 0.00618 | 0.938438 | 0.969735 | Healthy Control | Intestine |
| Alox12e | 3.91E+00 | 1.923715 | 7.410737 | 0.010265 | 0.033631 | Infection | Intestine |
| Alox15 | 6.57E+00 | 5.503793 | 173.4855 | 4.56E-10 | 1.74E-08 | Infection | Intestine |
| Alox5 | 3.78E+00 | 5.501238 | 65.42361 | 5.31E-08 | 9.43E-07 | Infection | Intestine |
| Alpl | 1.65E+00 | 5.782953 | 8.324579 | 0.008619 | 0.028996 | Infection | Intestine |
| Als2 | 1.29E+00 | 5.732892 | 9.034103 | 0.006487 | 0.022951 | Infection | Intestine |
| Alx1 | -3.57E-01 | 1.227067 | 0.006351 | 0.937596 | 0.969735 | Healthy Control | Intestine |
| Alx4 | -2.62E+00 | 1.632757 | 0.102544 | 0.751004 | 0.92508 | Healthy Control | Intestine |
| Ambra1 | 3.64E-01 | 8.512031 | 4.099419 | 0.055333 | 0.135982 | Infection | Intestine |

| | | | | | | | |
|---|---|---|---|---|---|---|---|
| Amd1 | -1.43E+00 | 8.457987 | 21.64025 | 0.00012 | 0.000728 | Healthy Control | Intestine |
| Amelx | 2.99E+00 | 2.061485 | 0.574863 | 0.457751 | 0.692088 | Infection | Intestine |
| Amer1 | -2.72E-01 | 5.812188 | 0.168116 | 0.685442 | 0.885046 | Healthy Control | Intestine |
| Amfr | -1.80E-01 | 10.39153 | 2.355749 | 0.139205 | 0.281105 | Healthy Control | Intestine |
| Amh | -3.42E+00 | 2.078643 | 0.22553 | 0.639451 | 0.854004 | Healthy Control | Intestine |
| Amhr2 | -3.83E+00 | 1.802993 | 0.289594 | 0.59464 | 0.820046 | Healthy Control | Intestine |
| Amigo3 | -7.36E+00 | 4.405244 | 3.146712 | 0.091478 | 0.20234 | Healthy Control | Intestine |
| Amy1 | 2.10E+00 | 2.138624 | 1.003152 | 0.325715 | 0.542424 | Infection | Intestine |
| Amy2a5 | 3.18E+00 | 5.188638 | 63.36201 | 6.89E-08 | 1.18E-06 | Infection | Intestine |
| Anapc11 | -1.81E+00 | 6.177699 | 4.60072 | 0.042144 | 0.108918 | Healthy Control | Intestine |
| Anapc15 | -5.06E+00 | 2.636435 | 0.909243 | 0.351667 | 0.575496 | Healthy Control | Intestine |
| Anapc16 | 6.29E-02 | 7.981516 | 0.029211 | 0.865866 | 0.960453 | Infection | Intestine |
| Anapc2 | -1.42E+00 | 9.098619 | 24.52053 | 6.00E-05 | 0.000395 | Healthy Control | Intestine |
| Anapc4 | -4.66E-01 | 8.174859 | 3.161434 | 0.08935 | 0.198353 | Healthy Control | Intestine |
| Anapc5 | 1.07E+00 | 9.347568 | 25.90871 | 4.35E-05 | 0.000297 | Infection | Intestine |
| Ang | 5.88E-01 | 3.379324 | 0.103772 | 0.750285 | 0.92508 | Infection | Intestine |
| Angpt1 | 2.38E+00 | 3.193056 | 5.686982 | 0.025267 | 0.071584 | Infection | Intestine |
| Angpt2 | -1.39E-01 | 3.934556 | 0.006634 | 0.935799 | 0.969735 | Healthy Control | Intestine |
| Angpt4 | 2.06E+00 | 2.232883 | 1.234276 | 0.276639 | 0.478691 | Infection | Intestine |
| Angptl3 | -1.83E+00 | 1.361829 | 0.057243 | 0.81282 | 0.950244 | Healthy Control | Intestine |
| Angptl4 | 4.22E+00 | 4.287635 | 61.09991 | 9.59E-08 | 1.54E-06 | Infection | Intestine |
| Angptl8 | -3.08E+00 | 1.486221 | 0.158673 | 0.69392 | 0.890937 | Healthy Control | Intestine |
| Ank | -1.96E-01 | 9.222761 | 0.757763 | 0.393516 | 0.623079 | Healthy Control | Intestine |
| Ank1 | -3.48E+00 | 1.889812 | 0.278698 | 0.601736 | 0.825737 | Healthy Control | Intestine |
| Ank2 | 4.34E+00 | 3.893084 | 53.80089 | 2.69E-07 | 3.59E-06 | Infection | Intestine |
| Ankle2 | 1.35E-02 | 8.404093 | 0.00356 | 0.952968 | 0.980184 | Infection | Intestine |
| Ankrd11 | -6.70E-01 | 7.985392 | 5.114961 | 0.033966 | 0.090734 | Healthy Control | Intestine |
| Ankrd13a | -4.47E-01 | 9.70711 | 7.488281 | 0.012112 | 0.038841 | Healthy Control | Intestine |
| Ankrd13b | -7.16E-01 | 6.763248 | 2.080603 | 0.162327 | 0.316855 | Healthy Control | Intestine |
| Ankrd13d | 2.76E+00 | 3.108994 | 5.323418 | 0.030002 | 0.082056 | Infection | Intestine |
| Anks6 | 1.51E-01 | 4.761937 | 0.030352 | 0.863132 | 0.958908 | Infection | Intestine |
| Anln | 2.06E+00 | 6.642323 | 23.48234 | 7.86E-05 | 0.000504 | Infection | Intestine |
| Ano1 | -3.71E+00 | 10.09426 | 74.81823 | 1.44E-08 | 3.00E-07 | Healthy Control | Intestine |
| Ano5 | 2.25E-01 | 3.821632 | 0.01794 | 0.894625 | 0.969735 | Infection | Intestine |
| Anp32e | -8.38E-01 | 6.195655 | 1.827054 | 0.188987 | 0.355398 | Healthy Control | Intestine |
| Anxa2 | -2.54E+00 | 11.73926 | 166.8561 | 1.09E-11 | 7.83E-10 | Healthy Control | Intestine |
| Anxa6 | 1.25E+00 | 8.656542 | 29.40151 | 1.96E-05 | 0.00015 | Infection | Intestine |
| Aox3 | -1.71E+00 | 1.318024 | 0.051098 | 0.823394 | 0.953685 | Healthy Control | Intestine |
| Ap1b1 | 5.52E-01 | 9.409303 | 13.01081 | 0.001582 | 0.006767 | Infection | Intestine |
| Ap1g1 | -1.95E-01 | 8.88109 | 0.964419 | 0.336863 | 0.556274 | Healthy Control | Intestine |
| Ap1s1 | -1.28E+00 | 8.0389 | 10.22967 | 0.004103 | 0.015422 | Healthy Control | Intestine |
| Ap2a1 | 9.34E-01 | 7.060316 | 6.492275 | 0.018413 | 0.055262 | Infection | Intestine |
| Ap2a2 | 8.38E-01 | 8.553022 | 17.28513 | 0.000417 | 0.002179 | Infection | Intestine |
| Ap2b1 | -2.79E-01 | 9.569257 | 3.413208 | 0.078297 | 0.179548 | Healthy Control | Intestine |
| Ap3b1 | -1.03E+00 | 8.654514 | 19.10789 | 0.000248 | 0.001378 | Healthy Control | Intestine |
| Ap3b2 | -4.64E+00 | 2.229897 | 0.826696 | 0.372628 | 0.598504 | Healthy Control | Intestine |
| Ap3m2 | 1.71E+00 | 5.251967 | 8.447917 | 0.008177 | 0.027752 | Infection | Intestine |
| Ap3s2 | -1.37E+00 | 6.758544 | 6.218235 | 0.019896 | 0.058936 | Healthy Control | Intestine |
| Ap4s1 | -6.59E-02 | 6.353421 | 0.012161 | 0.913144 | 0.969735 | Healthy Control | Intestine |
| Ap5z1 | -7.89E-01 | 6.178058 | 1.436019 | 0.242418 | 0.431889 | Healthy Control | Intestine |

| | | | | | | | |
|---|---|---|---|---|---|---|---|
| Apaf1 | -3.40E-01 | 7.758145 | 1.483079 | 0.236102 | 0.423479 | Healthy Control | Intestine |
| Apba1 | 4.10E-01 | 4.917651 | 0.195212 | 0.662502 | 0.869783 | Infection | Intestine |
| Apba3 | -9.19E-01 | 7.573972 | 4.050103 | 0.056312 | 0.137915 | Healthy Control | Intestine |
| Apc | 2.33E-01 | 8.330578 | 1.332194 | 0.26093 | 0.457358 | Infection | Intestine |
| Apc2 | 1.09E+00 | 2.930274 | 0.317386 | 0.57851 | 0.806375 | Infection | Intestine |
| Apcs | -9.71E-16 | 1.180616 | 0 | 1 | 1 | Healthy Control | Intestine |
| Apex1 | 1.93E+00 | 3.555315 | 4.556058 | 0.043565 | 0.111954 | Infection | Intestine |
| Aph1c | -5.04E+00 | 2.794002 | 0.873049 | 0.361257 | 0.586019 | Healthy Control | Intestine |
| Apip | -2.73E+00 | 7.085225 | 15.11853 | 0.000686 | 0.003334 | Healthy Control | Intestine |
| Apln | 1.24E+00 | 3.107747 | 0.480817 | 0.49502 | 0.727774 | Infection | Intestine |
| Aplnr | -2.36E+00 | 5.364862 | 1.691782 | 0.207377 | 0.382086 | Healthy Control | Intestine |
| Aplp2 | -9.11E-01 | 9.309572 | 14.73969 | 0.000905 | 0.004255 | Healthy Control | Intestine |
| Apmap | 5.55E-01 | 7.770746 | 3.71794 | 0.066976 | 0.159354 | Infection | Intestine |
| Apoa1 | -1.34E+00 | 1.402674 | 0.033627 | 0.855851 | 0.956553 | Healthy Control | Intestine |
| Apoa4 | 7.31E+00 | 1.527698 | 29.10766 | 1.51E-05 | 0.000119 | Infection | Intestine |
| Apob | -1.08E+00 | 4.208279 | 0.216636 | 0.646631 | 0.858916 | Healthy Control | Intestine |
| Apobec1 | -2.23E+00 | 8.096288 | 7.698027 | 0.011902 | 0.038266 | Healthy Control | Intestine |
| Apobec3 | 6.05E-02 | 7.646726 | 0.039063 | 0.845132 | 0.954066 | Infection | Intestine |
| App | 1.29E+00 | 10.8179 | 116.0079 | 3.40E-10 | 1.35E-08 | Infection | Intestine |
| Appbp2 | -2.42E+00 | 7.926027 | 21.91749 | 9.32E-05 | 0.000581 | Healthy Control | Intestine |
| Appl1 | 2.33E-02 | 8.401501 | 0.010734 | 0.918431 | 0.969735 | Infection | Intestine |
| Appl2 | -3.05E+00 | 9.956786 | 90.63442 | 2.32E-09 | 6.74E-08 | Healthy Control | Intestine |
| Aqp2 | -3.57E-01 | 1.228299 | 0.00618 | 0.938438 | 0.969735 | Healthy Control | Intestine |
| Aqp3 | -1.41E+00 | 1.402006 | 0.03234 | 0.85857 | 0.956553 | Healthy Control | Intestine |
| Aqp4 | -1.26E+01 | 9.421341 | 33.46261 | 5.85E-05 | 0.000386 | Healthy Control | Intestine |
| Aqp5 | -6.42E-01 | 1.273555 | 0.012735 | 0.91133 | 0.969735 | Healthy Control | Intestine |
| Aqp6 | -1.65E+00 | 1.317354 | 0.040195 | 0.842926 | 0.954066 | Healthy Control | Intestine |
| Ar | 1.71E+00 | 4.66511 | 4.565818 | 0.043722 | 0.1123 | Infection | Intestine |
| Araf | -6.45E-01 | 8.665221 | 5.340767 | 0.030696 | 0.083695 | Healthy Control | Intestine |
| Arap1 | 1.58E+00 | 9.055562 | 139.0588 | 6.22E-11 | 3.46E-09 | Infection | Intestine |
| Arc | -3.76E+00 | 2.261112 | 0.293305 | 0.592903 | 0.818553 | Healthy Control | Intestine |
| Arf2 | 1.71E+00 | 6.402688 | 22.91371 | 9.06E-05 | 0.000568 | Infection | Intestine |
| Arf6 | -4.08E-01 | 8.130652 | 1.963876 | 0.175115 | 0.336001 | Healthy Control | Intestine |
| Arfip2 | -2.83E+00 | 8.255556 | 35.95906 | 3.30E-06 | 3.10E-05 | Healthy Control | Intestine |
| Arg1 | 7.86E+00 | 7.269301 | 488.984 | 8.29E-13 | 9.88E-11 | Infection | Intestine |
| Arg2 | 2.35E+00 | 5.532331 | 28.84079 | 2.24E-05 | 0.000168 | Infection | Intestine |
| Arhgap17 | -2.54E+00 | 7.473211 | 18.88402 | 0.000227 | 0.00128 | Healthy Control | Intestine |
| Arhgap31 | 4.88E-01 | 6.647451 | 0.90628 | 0.351362 | 0.575186 | Infection | Intestine |
| Arhgap35 | -1.17E+00 | 7.828431 | 10.15172 | 0.004123 | 0.015472 | Healthy Control | Intestine |
| Arhgap44 | 3.89E-01 | 4.696992 | 0.151992 | 0.70001 | 0.89357 | Infection | Intestine |
| Arhgef1 | 2.60E+00 | 9.314773 | 612.0675 | 1.90E-17 | 1.36E-14 | Infection | Intestine |
| Arhgef11 | -1.92E+00 | 8.592896 | 40.99379 | 1.68E-06 | 1.74E-05 | Healthy Control | Intestine |
| Arhgef12 | -3.90E-01 | 8.832952 | 2.600853 | 0.12121 | 0.251567 | Healthy Control | Intestine |
| Arhgef15 | -8.66E-01 | 5.496322 | 0.332013 | 0.570557 | 0.799164 | Healthy Control | Intestine |
| Arhgef28 | -2.97E+00 | 9.38469 | 50.48921 | 3.62E-07 | 4.71E-06 | Healthy Control | Intestine |
| Arhgef3 | 5.45E-01 | 7.310608 | 3.721232 | 0.066843 | 0.159114 | Infection | Intestine |
| Arhgef6 | 3.73E+00 | 5.854589 | 128.921 | 1.28E-10 | 6.46E-09 | Infection | Intestine |
| Arhgef7 | -2.14E-01 | 7.666359 | 0.494828 | 0.48919 | 0.722598 | Healthy Control | Intestine |
| Arid1a | -2.10E+00 | 8.675587 | 52.82754 | 2.36E-07 | 3.22E-06 | Healthy Control | Intestine |
| Arid1b | -3.17E+00 | 8.077325 | 50.49475 | 2.27E-07 | 3.13E-06 | Healthy Control | Intestine |

| | | | | | | | |
|---|---|---|---|---|---|---|---|
| Arid2 | -8.91E-01 | 8.079778 | 8.164765 | 0.00911 | 0.030341 | Healthy Control | Intestine |
| Arid3a | 2.05E+00 | 5.058091 | 10.6735 | 0.003512 | 0.013442 | Infection | Intestine |
| Arid3b | -3.38E-02 | 5.776543 | 0.003336 | 0.954424 | 0.981314 | Healthy Control | Intestine |
| Arid5b | -6.26E-01 | 7.66003 | 1.851414 | 0.187282 | 0.35312 | Healthy Control | Intestine |
| Arl13b | 2.24E-02 | 6.746838 | 0.002723 | 0.958841 | 0.984203 | Infection | Intestine |
| Armc12 | -9.77E-01 | 1.318419 | 0.021469 | 0.884821 | 0.96867 | Healthy Control | Intestine |
| Arnt | 8.36E-01 | 7.928037 | 11.81548 | 0.002378 | 0.009598 | Infection | Intestine |
| Arnt2 | 8.50E-01 | 3.040477 | 0.213644 | 0.647909 | 0.859523 | Infection | Intestine |
| Arntl | -2.22E-01 | 8.049923 | 0.427721 | 0.519878 | 0.755446 | Healthy Control | Intestine |
| Arr3 | -6.41E-01 | 1.272019 | 0.010776 | 0.918405 | 0.969735 | Healthy Control | Intestine |
| Arrb1 | 7.78E-01 | 8.687448 | 19.17186 | 0.000244 | 0.001357 | Infection | Intestine |
| Arrb2 | 3.98E+00 | 7.078182 | 350.9183 | 6.29E-15 | 1.57E-12 | Infection | Intestine |
| Arrdc3 | 4.69E-01 | 8.561012 | 6.517856 | 0.018216 | 0.0548 | Infection | Intestine |
| Arx | -3.60E+00 | 2.05107 | 0.405753 | 0.529542 | 0.763504 | Healthy Control | Intestine |
| Asap1 | 5.80E-01 | 7.796192 | 5.670996 | 0.026426 | 0.074363 | Infection | Intestine |
| Asap2 | -7.98E-03 | 7.450435 | 0.000681 | 0.979415 | 0.998971 | Healthy Control | Intestine |
| Asb9 | -1.17E+00 | 1.360601 | 0.031962 | 0.859492 | 0.956553 | Healthy Control | Intestine |
| Ascc2 | 9.24E-01 | 7.64272 | 13.31226 | 0.001434 | 0.006231 | Infection | Intestine |
| Ascl1 | -4.90E+00 | 2.336856 | 0.550879 | 0.466002 | 0.701807 | Healthy Control | Intestine |
| Ascl2 | -3.36E+00 | 6.914049 | 7.191039 | 0.013444 | 0.04222 | Healthy Control | Intestine |
| Asf1a | -3.40E+00 | 6.762674 | 15.35773 | 0.000703 | 0.003399 | Healthy Control | Intestine |
| Ash1l | -8.10E-01 | 8.758526 | 15.42116 | 0.00073 | 0.003504 | Healthy Control | Intestine |
| Ash2l | -1.44E+00 | 8.854721 | 28.27887 | 2.45E-05 | 0.000181 | Healthy Control | Intestine |
| Asic1 | -1.37E+00 | 1.228088 | 0.037791 | 0.848672 | 0.954066 | Healthy Control | Intestine |
| Asic3 | -2.84E+00 | 1.766214 | 0.137008 | 0.714192 | 0.901474 | Healthy Control | Intestine |
| Asna1 | -1.98E+00 | 8.61689 | 32.95414 | 8.44E-06 | 7.17E-05 | Healthy Control | Intestine |
| Aspa | 2.43E+00 | 4.693116 | 12.89551 | 0.001621 | 0.006908 | Infection | Intestine |
| Asxl3 | -3.96E+00 | 1.89855 | 0.378333 | 0.544202 | 0.775484 | Healthy Control | Intestine |
| Atad3a | -6.60E-02 | 6.944405 | 0.017159 | 0.896952 | 0.969735 | Healthy Control | Intestine |
| Atf1 | -7.09E-01 | 7.578802 | 3.82114 | 0.063165 | 0.151586 | Healthy Control | Intestine |
| Atf2 | -6.02E-01 | 7.191873 | 1.91964 | 0.179293 | 0.341841 | Healthy Control | Intestine |
| Atf4 | 1.55E+00 | 10.24704 | 107.3171 | 6.95E-10 | 2.42E-08 | Infection | Intestine |
| Atf5 | 2.34E+00 | 8.704662 | 83.96384 | 2.89E-08 | 5.55E-07 | Infection | Intestine |
| Atf6 | -8.01E-01 | 6.853461 | 3.17882 | 0.087532 | 0.19527 | Healthy Control | Intestine |
| Atf6b | -2.00E+00 | 8.66766 | 44.18143 | 1.02E-06 | 1.14E-05 | Healthy Control | Intestine |
| Atg101 | -3.49E+00 | 8.899152 | 62.691 | 4.33E-08 | 7.90E-07 | Healthy Control | Intestine |
| Atg13 | -8.58E-01 | 8.618068 | 12.44742 | 0.001912 | 0.007954 | Healthy Control | Intestine |
| Atg14 | 1.43E-01 | 6.30037 | 0.069437 | 0.794536 | 0.943769 | Infection | Intestine |
| Atg16l1 | -4.36E-01 | 6.863102 | 1.034136 | 0.319883 | 0.534669 | Healthy Control | Intestine |
| Atg3 | -1.11E-01 | 7.47137 | 0.097338 | 0.757983 | 0.928591 | Healthy Control | Intestine |
| Atg5 | -2.73E+00 | 7.93558 | 27.4087 | 2.23E-05 | 0.000167 | Healthy Control | Intestine |
| Atg7 | -6.99E-01 | 7.401763 | 3.298037 | 0.082708 | 0.187348 | Healthy Control | Intestine |
| Atg9a | -3.15E+00 | 7.439405 | 37.18205 | 2.58E-06 | 2.48E-05 | Healthy Control | Intestine |
| Atl1 | -5.58E-01 | 4.660071 | 0.246174 | 0.624428 | 0.84099 | Healthy Control | Intestine |
| Atm | -1.94E+00 | 6.861011 | 10.98267 | 0.002926 | 0.011444 | Healthy Control | Intestine |
| Atn1 | -2.58E+00 | 8.673371 | 67.62357 | 2.44E-08 | 4.78E-07 | Healthy Control | Intestine |
| Atoh1 | -3.22E+00 | 7.044231 | 12.55431 | 0.001809 | 0.007596 | Healthy Control | Intestine |
| Atoh7 | 3.71E+00 | 1.559578 | 2.914735 | 0.09619 | 0.210185 | Infection | Intestine |
| Atp1b2 | 3.66E-01 | 5.677565 | 0.293628 | 0.593138 | 0.818651 | Infection | Intestine |
| Atp2a2 | -1.11E+00 | 10.52875 | 66.82344 | 4.41E-08 | 8.02E-07 | Healthy Control | Intestine |

| | | | | | | | |
|---|---|---|---|---|---|---|---|
| Atp2a3 | -2.87E+00 | 11.5587 | 204.7614 | 1.49E-12 | 1.62E-10 | Healthy Control | Intestine |
| Atp2b2 | -4.57E+00 | 2.395353 | 0.398818 | 0.533765 | 0.765939 | Healthy Control | Intestine |
| Atp2b3 | -1.88E+00 | 1.363249 | 0.04632 | 0.831517 | 0.954066 | Healthy Control | Intestine |
| Atp2b4 | -1.42E-01 | 6.643881 | 0.063472 | 0.803384 | 0.947264 | Healthy Control | Intestine |
| Atp4a | -2.16E+00 | 1.319625 | 0.0783 | 0.782593 | 0.940206 | Healthy Control | Intestine |
| Atp4b | 5.22E+00 | 1.276006 | 4.563616 | 0.049701 | 0.124393 | Infection | Intestine |
| Atp5f1 | -2.13E+00 | 11.12587 | 231.5684 | 4.36E-13 | 5.74E-11 | Healthy Control | Intestine |
| Atp5o | -2.55E+00 | 11.25386 | 316.6824 | 1.85E-14 | 3.85E-12 | Healthy Control | Intestine |
| Atp6ap2 | -1.46E+00 | 9.861064 | 54.77911 | 2.24E-07 | 3.10E-06 | Healthy Control | Intestine |
| Atp6v0a1 | 1.44E-02 | 8.049887 | 0.003854 | 0.951066 | 0.978831 | Infection | Intestine |
| Atp6v0d2 | 4.87E+00 | 2.184584 | 21.27029 | 7.03E-05 | 0.000455 | Infection | Intestine |
| Atp6v1a | -8.17E-01 | 8.98206 | 14.26604 | 0.001049 | 0.004826 | Healthy Control | Intestine |
| Atp6v1b2 | 2.80E-01 | 8.809631 | 2.406389 | 0.135245 | 0.274771 | Infection | Intestine |
| Atp7a | -1.13E+00 | 8.111411 | 12.13653 | 0.001988 | 0.008217 | Healthy Control | Intestine |
| Atp7b | -4.31E+00 | 2.686732 | 0.495525 | 0.488186 | 0.721792 | Healthy Control | Intestine |
| Atr | 2.28E-01 | 5.935228 | 0.161602 | 0.6914 | 0.889526 | Infection | Intestine |
| Atraid | -2.54E+00 | 8.249262 | 34.51664 | 4.86E-06 | 4.36E-05 | Healthy Control | Intestine |
| Atrx | 5.90E-01 | 8.010119 | 6.755142 | 0.01646 | 0.050151 | Infection | Intestine |
| Atxn1 | -1.65E+00 | 7.009133 | 10.0218 | 0.004211 | 0.015765 | Healthy Control | Intestine |
| Atxn10 | -6.64E-04 | 8.091304 | 4.55E-06 | 0.998318 | 1 | Healthy Control | Intestine |
| Atxn3 | -1.42E+00 | 7.347573 | 7.196832 | 0.013135 | 0.041487 | Healthy Control | Intestine |
| Aurka | -7.80E-01 | 5.765246 | 0.581529 | 0.452908 | 0.686424 | Healthy Control | Intestine |
| Aurkb | -1.18E+00 | 6.163604 | 1.377273 | 0.251711 | 0.445166 | Healthy Control | Intestine |
| Avil | 7.21E-01 | 3.902905 | 0.361404 | 0.553496 | 0.782305 | Infection | Intestine |
| Avpr1b | -3.68E+00 | 1.884977 | 0.360547 | 0.554563 | 0.783336 | Healthy Control | Intestine |
| Avpr2 | -2.22E+00 | 1.52073 | 0.137162 | 0.713771 | 0.901397 | Healthy Control | Intestine |
| Axin1 | 3.75E-01 | 6.569767 | 0.879107 | 0.358527 | 0.582912 | Infection | Intestine |
| Axin2 | -1.09E+00 | 8.235892 | 7.435765 | 0.012214 | 0.039118 | Healthy Control | Intestine |
| Axl | 1.45E+00 | 8.451195 | 26.08615 | 5.06E-05 | 0.000339 | Infection | Intestine |
| Azi2 | 2.21E-01 | 8.951665 | 1.56659 | 0.223973 | 0.406678 | Infection | Intestine |
| Azin2 | 5.31E+00 | 1.63647 | 13.48759 | 0.000741 | 0.003539 | Infection | Intestine |
| B3glct | -8.46E+00 | 5.716616 | 9.820247 | 0.005273 | 0.019143 | Healthy Control | Intestine |
| B3gnt2 | -1.32E+00 | 8.297574 | 17.33868 | 0.000395 | 0.002071 | Healthy Control | Intestine |
| B3gnt6 | -1.03E+01 | 6.292002 | 23.22767 | 9.17E-05 | 0.000575 | Healthy Control | Intestine |
| B4galnt1 | -1.88E+00 | 9.22642 | 16.33205 | 0.00064 | 0.003146 | Healthy Control | Intestine |
| B4galnt2 | -2.82E+00 | 10.823 | 14.67575 | 0.002117 | 0.008649 | Healthy Control | Intestine |
| B9d2 | 1.40E+00 | 4.229271 | 1.343622 | 0.257809 | 0.453216 | Infection | Intestine |
| Babam2 | -1.50E+00 | 7.456915 | 10.16489 | 0.004022 | 0.015162 | Healthy Control | Intestine |
| Bace1 | -5.54E-01 | 6.482759 | 0.818939 | 0.37478 | 0.601329 | Healthy Control | Intestine |
| Bace2 | -2.33E+00 | 8.352681 | 24.38401 | 4.88E-05 | 0.000328 | Healthy Control | Intestine |
| Bach1 | -6.18E-02 | 8.287542 | 0.046396 | 0.831457 | 0.954066 | Healthy Control | Intestine |
| Bach2 | -3.20E-01 | 6.157047 | 0.252651 | 0.61979 | 0.838379 | Healthy Control | Intestine |
| Bad | 1.11E+00 | 8.257301 | 31.86344 | 1.16E-05 | 9.42E-05 | Infection | Intestine |
| Bag2 | -6.13E+00 | 3.508965 | 1.768498 | 0.198811 | 0.369842 | Healthy Control | Intestine |
| Baiap2 | 2.50E-01 | 7.795759 | 0.889826 | 0.355858 | 0.580457 | Infection | Intestine |
| Bak1 | 8.30E-01 | 7.834142 | 13.89758 | 0.001185 | 0.005313 | Infection | Intestine |
| Banp | -3.94E-01 | 6.68094 | 0.761467 | 0.391794 | 0.621459 | Healthy Control | Intestine |
| Bap1 | -1.49E+00 | 7.383834 | 11.00047 | 0.002961 | 0.011561 | Healthy Control | Intestine |
| Bard1 | -5.21E-02 | 4.622918 | 0.002045 | 0.964294 | 0.987775 | Healthy Control | Intestine |
| Barhl1 | 4.53E+00 | 1.40514 | 3.463491 | 0.07341 | 0.170845 | Infection | Intestine |

| | | | | | | | |
|---|---|---|---|---|---|---|---|
| Barhl2 | -2.24E+00 | 1.361538 | 0.095114 | 0.760412 | 0.928591 | Healthy Control | Intestine |
| Barx1 | -1.92E+00 | 1.590262 | 0.060139 | 0.808842 | 0.94887 | Healthy Control | Intestine |
| Batf | 1.44E+00 | 4.363775 | 3.141862 | 0.088979 | 0.197879 | Infection | Intestine |
| Batf3 | 3.63E+00 | 5.408247 | 76.75302 | 1.38E-08 | 2.91E-07 | Infection | Intestine |
| Bax | 2.79E-01 | 8.257148 | 1.502941 | 0.233316 | 0.419702 | Infection | Intestine |
| Baz1a | 7.50E-01 | 7.361322 | 5.628128 | 0.026937 | 0.07563 | Infection | Intestine |
| Baz1b | 3.33E-01 | 8.686546 | 3.914988 | 0.06064 | 0.146649 | Infection | Intestine |
| Bbc3 | -3.13E-01 | 5.419164 | 0.073257 | 0.789008 | 0.941823 | Healthy Control | Intestine |
| Bbox1 | -3.27E+00 | 1.768533 | 0.202844 | 0.655661 | 0.865113 | Healthy Control | Intestine |
| Bbs1 | -1.52E+00 | 5.105642 | 1.300696 | 0.265836 | 0.464007 | Healthy Control | Intestine |
| Bbs2 | 1.80E-02 | 5.992166 | 0.000863 | 0.976811 | 0.997128 | Infection | Intestine |
| Bbs4 | -1.54E-02 | 5.513835 | 0.000233 | 0.987948 | 1 | Healthy Control | Intestine |
| Bbs5 | -1.41E-01 | 4.626177 | 0.017243 | 0.896594 | 0.969735 | Healthy Control | Intestine |
| Bbs7 | 8.39E-01 | 5.747754 | 1.885521 | 0.182916 | 0.347242 | Infection | Intestine |
| Bbs9 | -5.86E-01 | 6.250115 | 0.623441 | 0.437579 | 0.670704 | Healthy Control | Intestine |
| Bcam | -1.39E+00 | 6.897485 | 1.118662 | 0.304595 | 0.515702 | Healthy Control | Intestine |
| Bcar1 | 2.44E-01 | 7.549654 | 0.922615 | 0.347302 | 0.570217 | Infection | Intestine |
| Bcl10 | -1.22E+00 | 8.315181 | 18.61379 | 0.000273 | 0.001494 | Healthy Control | Intestine |
| Bcl11a | -7.44E+00 | 4.817227 | 3.244227 | 0.086935 | 0.19411 | Healthy Control | Intestine |
| Bcl11b | -8.04E+00 | 5.53826 | 7.517975 | 0.012644 | 0.040238 | Healthy Control | Intestine |
| Bcl2 | 1.23E+00 | 6.515359 | 11.34059 | 0.002802 | 0.011046 | Infection | Intestine |
| Bcl2a1a | 2.07E+00 | 5.596844 | 15.7548 | 0.000654 | 0.003202 | Infection | Intestine |
| Bcl2l1 | -7.74E-02 | 7.826953 | 0.08911 | 0.768131 | 0.932538 | Healthy Control | Intestine |
| Bcl2l11 | -1.45E-01 | 6.578587 | 0.069505 | 0.794439 | 0.943769 | Healthy Control | Intestine |
| Bcl2l12 | 5.22E-02 | 6.10465 | 0.007326 | 0.932529 | 0.969735 | Infection | Intestine |
| Bcl2l2 | -4.71E-01 | 8.521409 | 2.846033 | 0.105869 | 0.226681 | Healthy Control | Intestine |
| Bcl3 | -3.75E-01 | 6.375605 | 0.381274 | 0.542845 | 0.775095 | Healthy Control | Intestine |
| Bcl6 | 3.07E-01 | 7.081779 | 0.726878 | 0.403149 | 0.633866 | Infection | Intestine |
| Bcl7a | 6.29E-02 | 6.128708 | 0.009191 | 0.924455 | 0.969735 | Infection | Intestine |
| Bcl7b | 8.55E-01 | 8.206795 | 11.96085 | 0.002259 | 0.00918 | Infection | Intestine |
| Bcl7c | -1.24E+00 | 7.567979 | 12.76424 | 0.001636 | 0.006963 | Healthy Control | Intestine |
| Bclaf1 | 9.67E-01 | 8.6011 | 26.74531 | 3.58E-05 | 0.000252 | Infection | Intestine |
| Bco1 | -4.30E+00 | 2.585586 | 0.405416 | 0.530503 | 0.76467 | Healthy Control | Intestine |
| Bcor | -1.11E-01 | 7.579433 | 0.121617 | 0.730575 | 0.91327 | Healthy Control | Intestine |
| Bcr | 1.94E-01 | 7.860445 | 0.506416 | 0.484243 | 0.71874 | Infection | Intestine |
| Bdh1 | -3.16E+00 | 9.334457 | 59.60002 | 8.59E-08 | 1.40E-06 | Healthy Control | Intestine |
| Bdkrb1 | 5.06E+00 | 1.600642 | 7.754486 | 0.008612 | 0.028996 | Infection | Intestine |
| Bdkrb2 | 2.04E+00 | 6.782269 | 26.77304 | 3.56E-05 | 0.000251 | Infection | Intestine |
| Bdnf | 2.63E+00 | 1.836455 | 1.25193 | 0.274566 | 0.476092 | Infection | Intestine |
| Becn1 | -8.74E-01 | 8.693629 | 14.63161 | 0.000936 | 0.004379 | Healthy Control | Intestine |
| Bex3 | 1.89E-01 | 7.531505 | 0.251423 | 0.621091 | 0.839021 | Infection | Intestine |
| Bfsp1 | 2.38E-01 | 5.173878 | 0.063998 | 0.802558 | 0.947102 | Infection | Intestine |
| Bglap | 4.50E-02 | 3.056684 | 0.000565 | 0.981266 | 1 | Infection | Intestine |
| Bglap2 | -5.23E+00 | 2.58876 | 0.781227 | 0.387051 | 0.616278 | Healthy Control | Intestine |
| Bhlha15 | -7.59E+00 | 4.202173 | 2.737669 | 0.113864 | 0.240305 | Healthy Control | Intestine |
| Bhlhe22 | 5.19E+00 | 2.101631 | 26.84833 | 1.02E-05 | 8.45E-05 | Infection | Intestine |
| Bhlhe40 | 4.70E-01 | 7.773032 | 2.222349 | 0.150336 | 0.298999 | Infection | Intestine |
| Bhmt | 3.05E+00 | 1.801322 | 2.017876 | 0.164566 | 0.319432 | Infection | Intestine |
| Bicd2 | 1.34E+00 | 7.462122 | 28.66431 | 2.32E-05 | 0.000173 | Infection | Intestine |
| Bid | 1.51E+00 | 6.078923 | 14.16857 | 0.001054 | 0.00484 | Infection | Intestine |

| | | | | | | | |
|---|---|---|---|---|---|---|---|
| Bin1 | 1.84E+00 | 7.793116 | 55.59549 | 1.98E-07 | 2.78E-06 | Infection | Intestine |
| Birc2 | -2.93E+00 | 6.226537 | 7.710144 | 0.010871 | 0.035268 | Healthy Control | Intestine |
| Birc3 | -3.42E-01 | 7.728223 | 0.836725 | 0.370168 | 0.595648 | Healthy Control | Intestine |
| Birc5 | -5.14E-01 | 6.809886 | 0.685205 | 0.416431 | 0.649201 | Healthy Control | Intestine |
| Blk | -4.53E+00 | 2.380602 | 0.252885 | 0.620163 | 0.838493 | Healthy Control | Intestine |
| Blmh | 8.42E-01 | 7.438159 | 9.808177 | 0.004891 | 0.017965 | Infection | Intestine |
| Blnk | -2.04E+00 | 9.62379 | 57.57157 | 1.44E-07 | 2.15E-06 | Healthy Control | Intestine |
| Bloc1s5 | -2.98E-01 | 5.244738 | 0.104303 | 0.749528 | 0.924856 | Healthy Control | Intestine |
| Bloc1s6 | -3.41E-01 | 6.475218 | 0.39352 | 0.536596 | 0.768583 | Healthy Control | Intestine |
| Bmf | -2.16E+00 | 5.616925 | 3.318369 | 0.081666 | 0.185742 | Healthy Control | Intestine |
| Bmi1 | 1.94E+00 | 7.371704 | 112.0605 | 4.71E-10 | 1.79E-08 | Infection | Intestine |
| Bmp1 | 2.90E+00 | 8.46353 | 278.6419 | 6.63E-14 | 1.11E-11 | Infection | Intestine |
| Bmp10 | -1.78E+00 | 1.362331 | 0.054079 | 0.81794 | 0.952901 | Healthy Control | Intestine |
| Bmp15 | -9.77E-01 | 1.317821 | 0.017928 | 0.894683 | 0.969735 | Healthy Control | Intestine |
| Bmp2 | -7.59E-01 | 7.325678 | 1.90935 | 0.18021 | 0.343015 | Healthy Control | Intestine |
| Bmp3 | -3.30E+00 | 6.735178 | 9.965934 | 0.00446 | 0.016552 | Healthy Control | Intestine |
| Bmp4 | 3.69E-01 | 8.118048 | 1.19527 | 0.286201 | 0.490826 | Infection | Intestine |
| Bmp5 | -1.42E+00 | 7.065014 | 6.206099 | 0.020125 | 0.059544 | Healthy Control | Intestine |
| Bmp6 | 5.14E-01 | 7.120313 | 1.197343 | 0.285684 | 0.490276 | Infection | Intestine |
| Bmp7 | -8.35E+00 | 5.449318 | 6.810924 | 0.016854 | 0.051195 | Healthy Control | Intestine |
| Bmp8b | -2.78E+00 | 1.482965 | 0.175714 | 0.678159 | 0.881326 | Healthy Control | Intestine |
| Bmpr1a | -5.31E-01 | 6.823594 | 1.48505 | 0.235426 | 0.42272 | Healthy Control | Intestine |
| Bmpr1b | 1.89E+00 | 2.271842 | 1.350544 | 0.255518 | 0.450396 | Infection | Intestine |
| Bmpr2 | -3.90E-02 | 9.280751 | 0.062181 | 0.805417 | 0.947891 | Healthy Control | Intestine |
| Bnc1 | 4.97E+00 | 1.321203 | 4.118623 | 0.056704 | 0.138603 | Infection | Intestine |
| Bnc2 | 6.05E-01 | 5.557306 | 0.767664 | 0.390023 | 0.620012 | Infection | Intestine |
| Bnip3 | 2.28E+00 | 5.626337 | 32.34428 | 1.04E-05 | 8.56E-05 | Infection | Intestine |
| Boc | -2.73E+00 | 6.712743 | 16.29593 | 0.000513 | 0.002587 | Healthy Control | Intestine |
| Borcs8 | -1.44E+00 | 7.163676 | 9.466249 | 0.00522 | 0.018978 | Healthy Control | Intestine |
| Bpifb1 | -7.56E-01 | 1.274871 | 0.014406 | 0.905832 | 0.969735 | Healthy Control | Intestine |
| Braf | -1.56E+00 | 6.460151 | 5.487284 | 0.027631 | 0.077275 | Healthy Control | Intestine |
| Brca1 | 1.53E+00 | 4.647562 | 5.055891 | 0.034532 | 0.091951 | Infection | Intestine |
| Brca2 | 4.34E-01 | 4.14418 | 0.116602 | 0.735647 | 0.916538 | Infection | Intestine |
| Brcc3 | 5.18E-01 | 6.056766 | 0.95415 | 0.33888 | 0.558773 | Infection | Intestine |
| Brd2 | -5.50E-01 | 9.563588 | 11.41692 | 0.002731 | 0.010806 | Healthy Control | Intestine |
| Brd4 | -4.14E-01 | 8.733983 | 4.16023 | 0.053698 | 0.13268 | Healthy Control | Intestine |
| Brd7 | -1.13E+00 | 8.307993 | 13.04815 | 0.001539 | 0.006632 | Healthy Control | Intestine |
| Brd8 | 7.04E-01 | 8.30344 | 9.884058 | 0.004753 | 0.01749 | Infection | Intestine |
| Brip1 | -2.54E-01 | 5.387048 | 0.074214 | 0.787579 | 0.941409 | Healthy Control | Intestine |
| Brpf1 | -5.08E-01 | 7.73509 | 2.902776 | 0.102419 | 0.220995 | Healthy Control | Intestine |
| Brpf3 | -3.41E+00 | 8.30213 | 44.51967 | 6.21E-07 | 7.50E-06 | Healthy Control | Intestine |
| Brs3 | -3.57E-01 | 1.228299 | 0.00618 | 0.938438 | 0.969735 | Healthy Control | Intestine |
| Brsk2 | -3.53E+00 | 1.947291 | 0.198116 | 0.659885 | 0.867941 | Healthy Control | Intestine |
| Brwd3 | -2.67E+00 | 6.949959 | 15.61703 | 0.000609 | 0.003009 | Healthy Control | Intestine |
| Bsn | -6.84E+00 | 4.175982 | 2.397478 | 0.137427 | 0.278102 | Healthy Control | Intestine |
| Bsnd | -9.71E-16 | 1.180616 | 0 | 1 | 1 | Healthy Control | Intestine |
| Bsx | -9.71E-16 | 1.180616 | 0 | 1 | 1 | Healthy Control | Intestine |
| Btbd2 | -8.91E+00 | 6.318489 | 13.89511 | 0.001347 | 0.005928 | Healthy Control | Intestine |
| Btbd9 | 7.52E-01 | 6.965256 | 5.838435 | 0.024481 | 0.069908 | Infection | Intestine |
| Btc | -1.76E+00 | 5.002502 | 1.802508 | 0.193007 | 0.3616 | Healthy Control | Intestine |

| | | | | | | | |
|---|---|---|---|---|---|---|---|
| Btk | 1.95E+00 | 4.859874 | 10.37121 | 0.003827 | 0.014513 | Infection | Intestine |
| Btla | -6.18E+00 | 3.539201 | 0.855696 | 0.366174 | 0.591312 | Healthy Control | Intestine |
| Btrc | 6.43E-01 | 6.49062 | 3.009606 | 0.096702 | 0.210748 | Infection | Intestine |
| Bub1 | -1.74E+00 | 5.089715 | 1.896301 | 0.181982 | 0.345599 | Healthy Control | Intestine |
| Bub1b | 1.64E+00 | 6.732888 | 17.78133 | 0.000362 | 0.001926 | Infection | Intestine |
| C1ql1 | -4.74E+00 | 2.547559 | 0.620496 | 0.439803 | 0.672877 | Healthy Control | Intestine |
| C1qtnf12 | 1.53E+00 | 4.55827 | 5.77004 | 0.024562 | 0.070028 | Infection | Intestine |
| C1qtnf5 | 2.71E+00 | 5.909103 | 42.5922 | 1.53E-06 | 1.59E-05 | Infection | Intestine |
| C2 | 1.34E+00 | 5.205025 | 1.543323 | 0.226501 | 0.410821 | Infection | Intestine |
| C2cd6 | -1.52E+00 | 1.274134 | 0.042767 | 0.838583 | 0.954066 | Healthy Control | Intestine |
| C3 | 3.07E+00 | 8.360407 | 10.32098 | 0.00766 | 0.026304 | Infection | Intestine |
| C3ar1 | -1.96E+00 | 6.457971 | 2.597429 | 0.120081 | 0.249741 | Healthy Control | Intestine |
| C4b | 1.72E+00 | 8.901917 | 14.70606 | 0.001955 | 0.00811 | Infection | Intestine |
| C4bp | -8.55E+00 | 6.02581 | 8.973177 | 0.007202 | 0.024967 | Healthy Control | Intestine |
| C5ar1 | -7.55E+00 | 4.936514 | 2.960256 | 0.100932 | 0.218541 | Healthy Control | Intestine |
| C5ar2 | 2.46E+00 | 2.077624 | 1.32188 | 0.259579 | 0.455629 | Infection | Intestine |
| C6 | 4.02E+00 | 3.311817 | 8.70938 | 0.00815 | 0.027679 | Infection | Intestine |
| C7 | 5.77E+00 | 1.642164 | 13.86546 | 0.001117 | 0.005067 | Infection | Intestine |
| Cab39l | -9.30E-01 | 7.693379 | 7.349464 | 0.012676 | 0.040315 | Healthy Control | Intestine |
| Cabp1 | -6.74E+00 | 4.021546 | 2.392603 | 0.137815 | 0.278748 | Healthy Control | Intestine |
| Cacna1a | -1.25E+00 | 5.067726 | 0.778696 | 0.386875 | 0.616217 | Healthy Control | Intestine |
| Cacna1b | -1.37E+00 | 4.322412 | 0.67715 | 0.419738 | 0.651756 | Healthy Control | Intestine |
| Cacna1c | 2.68E+00 | 5.39104 | 36.61555 | 4.54E-06 | 4.13E-05 | Infection | Intestine |
| Cacna1d | 6.43E-01 | 5.546573 | 0.937092 | 0.342929 | 0.564581 | Infection | Intestine |
| Cacna1g | -3.38E-01 | 4.111002 | 0.041471 | 0.840432 | 0.954066 | Healthy Control | Intestine |
| Cacna1h | -2.12E-01 | 5.972528 | 0.067408 | 0.797394 | 0.944347 | Healthy Control | Intestine |
| Cacna1s | -2.80E-01 | 3.407474 | 0.026771 | 0.871588 | 0.962378 | Healthy Control | Intestine |
| Cacna2d1 | 3.54E+00 | 5.452176 | 94.67309 | 2.19E-09 | 6.43E-08 | Infection | Intestine |
| Cacna2d2 | -2.29E+00 | 5.474572 | 3.248603 | 0.085115 | 0.190985 | Healthy Control | Intestine |
| Cacnb1 | 1.40E+00 | 6.08631 | 10.12533 | 0.004295 | 0.016022 | Infection | Intestine |
| Cacng1 | -8.61E-01 | 1.274702 | 0.021049 | 0.886432 | 0.96867 | Healthy Control | Intestine |
| Cacng2 | -2.28E+00 | 1.362874 | 0.095976 | 0.759553 | 0.928591 | Healthy Control | Intestine |
| Cacng3 | -1.37E+00 | 1.228088 | 0.037791 | 0.848672 | 0.954066 | Healthy Control | Intestine |
| Cadm1 | -6.18E-01 | 5.854894 | 0.530886 | 0.473289 | 0.708095 | Healthy Control | Intestine |
| Cadm3 | -5.37E-01 | 5.603836 | 0.22489 | 0.639565 | 0.854004 | Healthy Control | Intestine |
| Cadps2 | -4.65E+00 | 8.90187 | 85.74015 | 2.40E-09 | 6.88E-08 | Healthy Control | Intestine |
| Calb1 | 5.33E+00 | 1.276779 | 3.834058 | 0.069519 | 0.163539 | Infection | Intestine |
| Calb2 | -7.62E+00 | 3.894313 | 0.415334 | 0.532103 | 0.765638 | Healthy Control | Intestine |
| Calcr | -3.57E-01 | 1.227387 | 0.006293 | 0.937883 | 0.969735 | Healthy Control | Intestine |
| Calcrl | 7.78E-01 | 7.239353 | 1.102615 | 0.310406 | 0.523372 | Infection | Intestine |
| Calu | 6.55E-01 | 9.325956 | 21.46224 | 0.000131 | 0.000781 | Infection | Intestine |
| Camk1d | -2.74E+00 | 8.752422 | 45.28406 | 6.71E-07 | 7.94E-06 | Healthy Control | Intestine |
| Camk2a | 8.01E-01 | 2.574078 | 0.13286 | 0.718907 | 0.904688 | Infection | Intestine |
| Camk2b | -1.05E+00 | 4.287031 | 0.30678 | 0.585277 | 0.810829 | Healthy Control | Intestine |
| Camk2d | 9.75E-01 | 8.583229 | 17.47857 | 0.000394 | 0.002067 | Infection | Intestine |
| Camk2g | -1.55E+00 | 6.50076 | 5.211803 | 0.031405 | 0.085441 | Healthy Control | Intestine |
| Camk4 | 6.28E-01 | 2.818118 | 0.092281 | 0.764182 | 0.930099 | Infection | Intestine |
| Camkv | -2.54E+00 | 1.484626 | 0.136578 | 0.714045 | 0.901474 | Healthy Control | Intestine |
| Camsap1 | 8.94E-01 | 6.501584 | 4.040849 | 0.056844 | 0.138811 | Infection | Intestine |
| Camsap2 | 5.13E-01 | 7.004975 | 1.689097 | 0.207168 | 0.382086 | Infection | Intestine |

| | | | | | | | |
|---|---|---|---|---|---|---|---|
| Camsap3 | -5.87E+00 | 8.994503 | 106.6717 | 5.84E-10 | 2.09E-08 | Healthy Control | Intestine |
| Camta2 | -3.97E-02 | 8.713034 | 0.041189 | 0.841058 | 0.954066 | Healthy Control | Intestine |
| Canx | -5.90E-01 | 10.25708 | 8.854758 | 0.007021 | 0.024458 | Healthy Control | Intestine |
| Cap2 | 1.23E+00 | 5.214456 | 2.457132 | 0.130984 | 0.267151 | Infection | Intestine |
| Capn1 | -1.68E+00 | 9.106259 | 43.66081 | 1.26E-06 | 1.36E-05 | Healthy Control | Intestine |
| Capn10 | -4.73E-01 | 7.489678 | 1.788759 | 0.194673 | 0.364038 | Healthy Control | Intestine |
| Capn2 | -6.70E-01 | 9.691704 | 8.440644 | 0.008261 | 0.027999 | Healthy Control | Intestine |
| Caprin1 | -8.36E-02 | 10.06838 | 0.323355 | 0.57541 | 0.803039 | Healthy Control | Intestine |
| Car4 | -6.66E-01 | 9.83247 | 1.175186 | 0.298847 | 0.507645 | Healthy Control | Intestine |
| Car7 | 8.74E-01 | 3.016759 | 0.209967 | 0.651107 | 0.862286 | Infection | Intestine |
| Card9 | 1.40E+00 | 4.151331 | 2.290128 | 0.143213 | 0.28746 | Infection | Intestine |
| Carm1 | -1.16E+00 | 7.841373 | 7.513627 | 0.011777 | 0.037914 | Healthy Control | Intestine |
| Carmil1 | -2.95E+00 | 7.844582 | 23.50642 | 6.03E-05 | 0.000395 | Healthy Control | Intestine |
| Carns1 | 2.94E+00 | 3.67102 | 11.34197 | 0.002728 | 0.010805 | Infection | Intestine |
| Cartpt | -2.41E+00 | 1.443828 | 0.10408 | 0.749283 | 0.924781 | Healthy Control | Intestine |
| Casc4 | -1.69E+00 | 6.698745 | 4.945238 | 0.036125 | 0.095428 | Healthy Control | Intestine |
| Caskin1 | -2.51E+00 | 1.48449 | 0.10754 | 0.745063 | 0.923383 | Healthy Control | Intestine |
| Casp1 | -2.43E+00 | 9.655821 | 62.48291 | 7.32E-08 | 1.23E-06 | Healthy Control | Intestine |
| Casp12 | -1.34E-01 | 5.90206 | 0.034495 | 0.854257 | 0.956553 | Healthy Control | Intestine |
| Casp2 | -1.87E+00 | 6.843296 | 7.023537 | 0.013881 | 0.043294 | Healthy Control | Intestine |
| Casp3 | 1.85E-01 | 7.655748 | 0.420958 | 0.523245 | 0.7574 | Infection | Intestine |
| Casp4 | -2.74E-01 | 7.331122 | 0.33935 | 0.566132 | 0.794298 | Healthy Control | Intestine |
| Casp7 | -2.09E+00 | 8.407905 | 22.09589 | 9.53E-05 | 0.000594 | Healthy Control | Intestine |
| Casp8 | -3.17E-01 | 8.757097 | 2.628709 | 0.119343 | 0.248619 | Healthy Control | Intestine |
| Casp9 | -1.16E+00 | 6.448264 | 4.478382 | 0.044917 | 0.114898 | Healthy Control | Intestine |
| Casq1 | 4.47E+00 | 1.636451 | 7.208431 | 0.010598 | 0.034496 | Infection | Intestine |
| Casq2 | 8.99E-01 | 2.928831 | 0.2546 | 0.618638 | 0.837906 | Infection | Intestine |
| Casr | -1.61E+00 | 1.444939 | 0.049758 | 0.82512 | 0.953685 | Healthy Control | Intestine |
| Casz1 | -2.19E+00 | 7.817595 | 12.9142 | 0.001473 | 0.006386 | Healthy Control | Intestine |
| Cat | -2.18E+00 | 8.747693 | 38.31965 | 2.95E-06 | 2.80E-05 | Healthy Control | Intestine |
| Cav1 | 1.01E+00 | 8.277037 | 5.801862 | 0.028201 | 0.078126 | Infection | Intestine |
| Cav2 | -2.86E-02 | 7.450378 | 0.005399 | 0.942086 | 0.972388 | Healthy Control | Intestine |
| Cav3 | 1.12E+00 | 2.886323 | 0.313347 | 0.580998 | 0.807686 | Infection | Intestine |
| Cavin1 | 3.73E-01 | 8.619436 | 0.614841 | 0.444617 | 0.678358 | Infection | Intestine |
| Cbfb | -7.54E-01 | 7.091403 | 2.805217 | 0.10733 | 0.229221 | Healthy Control | Intestine |
| Cbl | 1.20E+00 | 8.748497 | 63.08007 | 7.10E-08 | 1.20E-06 | Infection | Intestine |
| Cblb | 5.59E-01 | 6.28372 | 1.068321 | 0.312403 | 0.525677 | Infection | Intestine |
| Cblc | -5.19E+00 | 8.509463 | 72.66127 | 1.57E-08 | 3.21E-07 | Healthy Control | Intestine |
| Cblif | -9.71E-16 | 1.180616 | 0 | 1 | 1 | Healthy Control | Intestine |
| Cbln1 | 3.22E+00 | 1.737884 | 1.937231 | 0.172816 | 0.332226 | Infection | Intestine |
| Cbln2 | -1.62E+00 | 1.482211 | 0.062206 | 0.804978 | 0.947891 | Healthy Control | Intestine |
| Cbln3 | -6.82E+00 | 4.182941 | 1.975341 | 0.175452 | 0.336389 | Healthy Control | Intestine |
| Cbln4 | -3.96E+00 | 1.708058 | 0.323947 | 0.57402 | 0.801993 | Healthy Control | Intestine |
| Cbs | -4.99E+00 | 9.216749 | 48.41207 | 4.44E-07 | 5.60E-06 | Healthy Control | Intestine |
| Cbx1 | 3.53E-01 | 5.80436 | 0.430693 | 0.518135 | 0.753568 | Infection | Intestine |
| Cbx2 | -6.46E+00 | 3.947889 | 2.085918 | 0.164399 | 0.319238 | Healthy Control | Intestine |
| Cbx3 | 8.98E-01 | 10.28197 | 71.40663 | 2.52E-08 | 4.89E-07 | Infection | Intestine |
| Cbx5 | 5.58E-01 | 7.562568 | 3.671956 | 0.068549 | 0.162144 | Infection | Intestine |
| Cbx7 | -2.79E-01 | 6.475613 | 0.262791 | 0.613138 | 0.833923 | Healthy Control | Intestine |
| Ccar2 | -5.92E-01 | 8.377962 | 5.798776 | 0.024929 | 0.070864 | Healthy Control | Intestine |

| | | | | | | | |
|---|---|---|---|---|---|---|---|
| Ccdc130 | -8.49E-01 | 6.226246 | 2.063092 | 0.163865 | 0.31869 | Healthy Control | Intestine |
| Ccdc184 | -2.18E+00 | 1.519372 | 0.069274 | 0.794544 | 0.943769 | Healthy Control | Intestine |
| Cckbr | -7.56E-01 | 1.27352 | 0.015516 | 0.902291 | 0.969735 | Healthy Control | Intestine |
| Ccl1 | 5.70E+00 | 1.229995 | 6.086704 | 0.031631 | 0.085684 | Infection | Intestine |
| Ccl11 | 4.71E+00 | 7.490916 | 84.64034 | 6.30E-07 | 7.58E-06 | Infection | Intestine |
| Ccl12 | -4.43E-02 | 5.652484 | 0.00103 | 0.974721 | 0.995807 | Healthy Control | Intestine |
| Ccl17 | 6.64E-01 | 3.486546 | 0.105512 | 0.748601 | 0.924698 | Infection | Intestine |
| Ccl19 | -1.89E+00 | 1.40182 | 0.048999 | 0.826467 | 0.953685 | Healthy Control | Intestine |
| Ccl2 | 2.89E+00 | 5.282583 | 14.06078 | 0.001305 | 0.005771 | Infection | Intestine |
| Ccl20 | -3.62E+00 | 2.301506 | 0.172428 | 0.682668 | 0.883743 | Healthy Control | Intestine |
| Ccl21a | 6.15E-01 | 7.984526 | 0.153404 | 0.702352 | 0.894649 | Infection | Intestine |
| Ccl21b | 2.31E-01 | 6.967577 | 0.010894 | 0.918602 | 0.969735 | Infection | Intestine |
| Ccl24 | 6.50E+00 | 4.961742 | 455.452 | 4.67E-16 | 1.56E-13 | Infection | Intestine |
| Ccl25 | -8.00E-01 | 4.370027 | 0.28404 | 0.599199 | 0.823611 | Healthy Control | Intestine |
| Ccl3 | 3.95E-01 | 3.443454 | 0.039188 | 0.844809 | 0.954066 | Infection | Intestine |
| Ccl7 | 5.94E+00 | 5.065396 | 110.1879 | 8.19E-09 | 1.90E-07 | Infection | Intestine |
| Ccl8 | 3.40E+00 | 7.287753 | 16.45369 | 0.001536 | 0.006622 | Infection | Intestine |
| Ccn1 | 1.81E+00 | 6.67624 | 19.19754 | 0.000243 | 0.001355 | Infection | Intestine |
| Ccn2 | 1.79E+00 | 7.680943 | 14.56028 | 0.001484 | 0.006421 | Infection | Intestine |
| Ccna1 | -3.29E-01 | 3.713227 | 0.03123 | 0.861326 | 0.957326 | Healthy Control | Intestine |
| Ccna2 | -7.65E-01 | 7.920583 | 1.967466 | 0.175746 | 0.336565 | Healthy Control | Intestine |
| Ccnb1 | -2.09E+00 | 5.507028 | 1.953923 | 0.175639 | 0.33649 | Healthy Control | Intestine |
| Ccnb3 | -9.71E-16 | 1.180616 | 0 | 1 | 1 | Healthy Control | Intestine |
| Ccnd1 | 2.91E-01 | 8.947537 | 1.596425 | 0.219763 | 0.400194 | Infection | Intestine |
| Ccnd2 | -9.58E-01 | 10.08488 | 14.00522 | 0.001143 | 0.005154 | Healthy Control | Intestine |
| Ccnd3 | -3.92E+00 | 9.070926 | 91.30685 | 1.06E-09 | 3.49E-08 | Healthy Control | Intestine |
| Ccne1 | 2.50E+00 | 3.840809 | 10.93945 | 0.003064 | 0.011917 | Infection | Intestine |
| Ccnt1 | -2.03E+00 | 7.273178 | 14.70052 | 0.000793 | 0.003769 | Healthy Control | Intestine |
| Ccp110 | -5.58E-01 | 4.158391 | 0.129872 | 0.721877 | 0.90683 | Healthy Control | Intestine |
| Ccr1 | 3.57E-01 | 6.141976 | 0.29605 | 0.591532 | 0.817111 | Infection | Intestine |
| Ccr10 | -1.01E+00 | 4.073096 | 0.218552 | 0.644859 | 0.857415 | Healthy Control | Intestine |
| Ccr2 | 8.38E-01 | 4.648045 | 0.523084 | 0.476802 | 0.711649 | Infection | Intestine |
| Ccr3 | 7.47E+00 | 2.570045 | 128.8453 | 4.73E-13 | 6.07E-11 | Infection | Intestine |
| Ccr4 | -5.03E+00 | 2.451249 | 0.881794 | 0.358339 | 0.582912 | Healthy Control | Intestine |
| Ccr5 | 4.02E+00 | 5.079015 | 76.72722 | 1.39E-08 | 2.91E-07 | Infection | Intestine |
| Ccr6 | -5.99E+00 | 3.540167 | 1.192724 | 0.287913 | 0.493087 | Healthy Control | Intestine |
| Ccr7 | -5.79E+00 | 3.137886 | 0.21542 | 0.649623 | 0.861198 | Healthy Control | Intestine |
| Ccr9 | -4.08E+00 | 2.163022 | 0.466822 | 0.501166 | 0.734437 | Healthy Control | Intestine |
| Ccser2 | -2.69E-01 | 6.456328 | 0.267207 | 0.610088 | 0.83218 | Healthy Control | Intestine |
| Cct3 | -7.24E-01 | 9.091406 | 8.931747 | 0.006822 | 0.023881 | Healthy Control | Intestine |
| Cct5 | -8.29E-01 | 10.4864 | 36.2123 | 4.86E-06 | 4.36E-05 | Healthy Control | Intestine |
| Cct7 | -9.17E-01 | 9.259917 | 22.63472 | 9.74E-05 | 0.000605 | Healthy Control | Intestine |
| Cct8 | -5.36E-01 | 9.793063 | 13.45909 | 0.001365 | 0.005989 | Healthy Control | Intestine |
| Cd109 | 1.04E+00 | 4.605915 | 0.668965 | 0.421646 | 0.653734 | Infection | Intestine |
| Cd14 | -3.06E+00 | 8.561155 | 32.05811 | 8.87E-06 | 7.43E-05 | Healthy Control | Intestine |
| Cd151 | 2.79E-01 | 9.080277 | 3.097493 | 0.092443 | 0.203774 | Infection | Intestine |
| Cd160 | -4.56E+00 | 2.639901 | 0.372007 | 0.548133 | 0.778138 | Healthy Control | Intestine |
| Cd163 | 3.02E+00 | 6.42514 | 19.38455 | 0.000495 | 0.002517 | Infection | Intestine |
| Cd177 | -4.22E+00 | 9.548938 | 30.12376 | 3.11E-05 | 0.000222 | Healthy Control | Intestine |
| Cd180 | -8.38E-01 | 4.329286 | 0.24246 | 0.627196 | 0.84311 | Healthy Control | Intestine |

| | | | | | | | |
|---|---|---|---|---|---|---|---|
| Cd19 | -4.50E+00 | 2.39435 | 0.174231 | 0.681152 | 0.882694 | Healthy Control | Intestine |
| Cd1d1 | -1.74E+00 | 6.143388 | 2.993751 | 0.096972 | 0.211153 | Healthy Control | Intestine |
| Cd2 | -7.38E+00 | 4.384492 | 1.866002 | 0.187268 | 0.35312 | Healthy Control | Intestine |
| Cd200 | 1.19E+00 | 6.49348 | 3.808149 | 0.064891 | 0.154909 | Infection | Intestine |
| Cd200r1 | 4.44E+00 | 4.02869 | 71.74038 | 2.53E-08 | 4.90E-07 | Infection | Intestine |
| Cd207 | -7.36E+00 | 4.156078 | 1.075837 | 0.314623 | 0.52817 | Healthy Control | Intestine |
| Cd209a | -3.02E-01 | 4.334593 | 0.029088 | 0.866141 | 0.96049 | Healthy Control | Intestine |
| Cd209b | -5.54E+00 | 3.220003 | 1.155728 | 0.295299 | 0.503325 | Healthy Control | Intestine |
| Cd209d | -5.57E+00 | 3.091052 | 0.718197 | 0.406945 | 0.63781 | Healthy Control | Intestine |
| Cd22 | -5.74E+00 | 3.283397 | 0.846592 | 0.368545 | 0.594374 | Healthy Control | Intestine |
| Cd226 | 1.73E-01 | 3.410975 | 0.00714 | 0.93341 | 0.969735 | Infection | Intestine |
| Cd244a | 4.54E+00 | 4.552369 | 63.59768 | 6.78E-08 | 1.17E-06 | Infection | Intestine |
| Cd247 | -5.72E+00 | 3.0311 | 1.068123 | 0.313991 | 0.527558 | Healthy Control | Intestine |
| Cd248 | 3.72E-01 | 7.668924 | 0.149679 | 0.704829 | 0.896371 | Infection | Intestine |
| Cd24a | 6.92E-01 | 11.14168 | 12.29416 | 0.002639 | 0.010493 | Infection | Intestine |
| Cd27 | -3.10E+00 | 1.520847 | 0.142426 | 0.708505 | 0.897011 | Healthy Control | Intestine |
| Cd274 | 3.47E+00 | 5.300789 | 69.23392 | 3.32E-08 | 6.22E-07 | Infection | Intestine |
| Cd276 | 1.33E-01 | 4.436601 | 0.017096 | 0.897053 | 0.969735 | Infection | Intestine |
| Cd28 | -6.51E-01 | 5.017191 | 0.180721 | 0.674552 | 0.879138 | Healthy Control | Intestine |
| Cd2ap | -2.94E+00 | 9.430755 | 109.644 | 3.40E-10 | 1.35E-08 | Healthy Control | Intestine |
| Cd2bp2 | -5.73E-02 | 6.681819 | 0.019257 | 0.890874 | 0.969735 | Healthy Control | Intestine |
| Cd300a | 1.56E-01 | 4.279985 | 0.025749 | 0.87389 | 0.963801 | Infection | Intestine |
| Cd300lb | 1.15E+00 | 4.016073 | 1.250987 | 0.275141 | 0.47669 | Infection | Intestine |
| Cd300ld | -9.76E-02 | 3.779545 | 0.003179 | 0.955529 | 0.981696 | Healthy Control | Intestine |
| Cd300lf | 5.36E+00 | 3.593651 | 150.6313 | 3.54E-11 | 2.13E-09 | Infection | Intestine |
| Cd320 | -4.28E+00 | 2.131035 | 0.483423 | 0.494037 | 0.726969 | Healthy Control | Intestine |
| Cd33 | -1.98E+00 | 5.472186 | 1.760945 | 0.197712 | 0.368209 | Healthy Control | Intestine |
| Cd34 | 1.61E+00 | 7.563754 | 17.26451 | 0.000496 | 0.002519 | Infection | Intestine |
| Cd36 | -8.77E-01 | 7.8829 | 2.503369 | 0.128859 | 0.26394 | Healthy Control | Intestine |
| Cd37 | 4.30E+00 | 5.250218 | 90.30945 | 3.33E-09 | 8.88E-08 | Infection | Intestine |
| Cd38 | -1.84E+00 | 10.02098 | 69.64378 | 3.02E-08 | 5.77E-07 | Healthy Control | Intestine |
| Cd3d | -8.23E-02 | 3.556241 | 0.001501 | 0.969438 | 0.991826 | Healthy Control | Intestine |
| Cd3e | 5.45E-01 | 4.010205 | 0.13867 | 0.712838 | 0.900446 | Infection | Intestine |
| Cd3eap | -7.80E-01 | 4.01927 | 0.299126 | 0.589984 | 0.815873 | Healthy Control | Intestine |
| Cd3g | -6.98E+00 | 4.128311 | 1.446071 | 0.243371 | 0.432948 | Healthy Control | Intestine |
| Cd4 | -7.18E-01 | 5.786715 | 0.406811 | 0.529506 | 0.763504 | Healthy Control | Intestine |
| Cd40 | 1.79E+00 | 3.967625 | 3.553339 | 0.072032 | 0.168187 | Infection | Intestine |
| Cd40lg | -4.06E+00 | 2.080243 | 0.383988 | 0.541347 | 0.774023 | Healthy Control | Intestine |
| Cd44 | 1.49E+00 | 7.874562 | 17.1641 | 0.000517 | 0.002604 | Infection | Intestine |
| Cd47 | 1.49E+00 | 8.977851 | 124.5956 | 1.75E-10 | 7.89E-09 | Infection | Intestine |
| Cd48 | 1.06E+00 | 5.763382 | 2.86113 | 0.10459 | 0.224231 | Infection | Intestine |
| Cd5 | -4.40E+00 | 2.157961 | 0.398974 | 0.533475 | 0.765939 | Healthy Control | Intestine |
| Cd53 | -1.90E+00 | 6.232959 | 3.751444 | 0.064563 | 0.154347 | Healthy Control | Intestine |
| Cd55 | -4.98E-02 | 8.152268 | 0.008423 | 0.927941 | 0.969735 | Healthy Control | Intestine |
| Cd59a | 2.08E+00 | 4.269134 | 5.469472 | 0.028555 | 0.078999 | Infection | Intestine |
| Cd59b | 1.79E+00 | 3.921589 | 3.27334 | 0.083012 | 0.187782 | Infection | Intestine |
| Cd5l | -6.42E-01 | 1.2729 | 0.010432 | 0.919715 | 0.969735 | Healthy Control | Intestine |
| Cd6 | 7.21E-01 | 2.898097 | 0.10563 | 0.748249 | 0.924698 | Infection | Intestine |
| Cd68 | -4.66E-01 | 7.534132 | 0.525993 | 0.476506 | 0.711632 | Healthy Control | Intestine |
| Cd69 | 5.43E-02 | 3.171863 | 0.000706 | 0.979057 | 0.998809 | Infection | Intestine |

| | | | | | | | |
|---|---|---|---|---|---|---|---|
| Cd70 | -4.95E-01 | 1.228692 | 0.008683 | 0.927069 | 0.969735 | Healthy Control | Intestine |
| Cd72 | 9.29E-02 | 6.470858 | 0.013664 | 0.908019 | 0.969735 | Infection | Intestine |
| Cd79a | -8.15E+00 | 5.149171 | 1.508876 | 0.238891 | 0.426951 | Healthy Control | Intestine |
| Cd79b | -8.20E+00 | 5.429848 | 1.523808 | 0.237006 | 0.424388 | Healthy Control | Intestine |
| Cd80 | 1.15E+00 | 3.868128 | 0.839214 | 0.368801 | 0.594404 | Infection | Intestine |
| Cd81 | -2.34E+00 | 9.61722 | 84.58571 | 5.16E-09 | 1.28E-07 | Healthy Control | Intestine |
| Cd82 | -2.20E+00 | 8.175298 | 27.53369 | 2.41E-05 | 0.000179 | Healthy Control | Intestine |
| Cd83 | -1.83E+00 | 6.491783 | 3.423609 | 0.0764 | 0.175801 | Healthy Control | Intestine |
| Cd84 | 3.43E+00 | 6.464081 | 171.4953 | 8.36E-12 | 6.44E-10 | Infection | Intestine |
| Cd86 | -7.50E+00 | 4.657476 | 2.383294 | 0.138479 | 0.279978 | Healthy Control | Intestine |
| Cd8a | -5.09E+00 | 2.957801 | 0.659205 | 0.42618 | 0.658881 | Healthy Control | Intestine |
| Cd8b1 | -4.53E+00 | 2.733172 | 0.288034 | 0.596551 | 0.822229 | Healthy Control | Intestine |
| Cd9 | -2.67E+00 | 10.25382 | 99.64586 | 1.20E-09 | 3.83E-08 | Healthy Control | Intestine |
| Cd93 | -2.17E-01 | 6.453353 | 0.13207 | 0.719631 | 0.905372 | Healthy Control | Intestine |
| Cd96 | -5.19E+00 | 2.80488 | 1.039053 | 0.320347 | 0.535266 | Healthy Control | Intestine |
| Cdc16 | -4.90E-01 | 8.075917 | 2.781289 | 0.1096 | 0.232829 | Healthy Control | Intestine |
| Cdc20 | -1.48E+00 | 6.838294 | 4.057904 | 0.055213 | 0.135753 | Healthy Control | Intestine |
| Cdc23 | 7.46E-01 | 8.277779 | 12.62605 | 0.001801 | 0.00757 | Infection | Intestine |
| Cdc25a | -4.20E-01 | 7.150463 | 1.463707 | 0.238883 | 0.426951 | Healthy Control | Intestine |
| Cdc27 | 5.21E-01 | 7.31787 | 2.549611 | 0.124692 | 0.257214 | Infection | Intestine |
| Cdc37 | -2.30E+00 | 9.60955 | 124.51 | 1.58E-10 | 7.48E-09 | Healthy Control | Intestine |
| Cdc42 | -1.55E-01 | 6.629107 | 0.109396 | 0.74385 | 0.923082 | Healthy Control | Intestine |
| Cdc42bpb | 8.02E-01 | 8.4364 | 12.93858 | 0.001621 | 0.006908 | Infection | Intestine |
| Cdc42ep4 | -1.06E+00 | 8.141321 | 12.16669 | 0.002073 | 0.008513 | Healthy Control | Intestine |
| Cdc5l | 1.66E+00 | 6.968463 | 36.32037 | 4.78E-06 | 4.32E-05 | Infection | Intestine |
| Cdc6 | 1.88E+00 | 3.517069 | 3.146465 | 0.089042 | 0.197879 | Infection | Intestine |
| Cdc73 | -9.26E-01 | 7.999729 | 7.833346 | 0.010394 | 0.033942 | Healthy Control | Intestine |
| Cdca5 | 6.43E-01 | 5.68655 | 0.887056 | 0.35625 | 0.580719 | Infection | Intestine |
| Cdca8 | 6.52E-01 | 6.419866 | 1.374082 | 0.253674 | 0.447776 | Infection | Intestine |
| Cdcp1 | -4.06E+00 | 8.343182 | 77.35487 | 5.26E-09 | 1.29E-07 | Healthy Control | Intestine |
| Cdh1 | -3.22E+00 | 12.04438 | 502.4612 | 1.57E-16 | 6.53E-14 | Healthy Control | Intestine |
| Cdh10 | 4.26E+00 | 1.4854 | 4.240949 | 0.047156 | 0.118984 | Infection | Intestine |
| Cdh11 | 1.40E+00 | 5.703813 | 7.500691 | 0.011998 | 0.038525 | Infection | Intestine |
| Cdh12 | 4.39E+00 | 1.445814 | 4.154426 | 0.049871 | 0.124702 | Infection | Intestine |
| Cdh13 | -1.20E+00 | 5.353123 | 1.198353 | 0.284679 | 0.489055 | Healthy Control | Intestine |
| Cdh2 | 1.08E+00 | 3.300821 | 0.641332 | 0.431309 | 0.663533 | Infection | Intestine |
| Cdh20 | -1.17E+00 | 1.361862 | 0.028669 | 0.866851 | 0.96049 | Healthy Control | Intestine |
| Cdh23 | -3.02E+00 | 1.563433 | 0.134867 | 0.716855 | 0.90301 | Healthy Control | Intestine |
| Cdh26 | 2.13E-01 | 3.3455 | 0.015437 | 0.902245 | 0.969735 | Infection | Intestine |
| Cdh3 | 6.59E-01 | 2.884493 | 0.111089 | 0.742079 | 0.921798 | Infection | Intestine |
| Cdh4 | 1.41E+00 | 2.697492 | 0.536744 | 0.470941 | 0.706162 | Infection | Intestine |
| Cdh5 | -1.15E+00 | 6.998569 | 0.633796 | 0.438192 | 0.671028 | Healthy Control | Intestine |
| Cdh6 | 4.55E+00 | 3.890263 | 86.12171 | 5.36E-09 | 1.30E-07 | Infection | Intestine |
| Cdh7 | 3.74E+00 | 1.446553 | 2.613226 | 0.116887 | 0.24493 | Infection | Intestine |
| Cdh8 | -3.57E-01 | 1.227387 | 0.006293 | 0.937883 | 0.969735 | Healthy Control | Intestine |
| Cdh9 | 4.34E+00 | 1.363553 | 3.243732 | 0.084702 | 0.190398 | Infection | Intestine |
| Cdhr1 | -5.03E-01 | 4.995675 | 0.227711 | 0.63737 | 0.852893 | Healthy Control | Intestine |
| Cdipt | -2.80E-01 | 8.388341 | 1.20455 | 0.284401 | 0.488912 | Healthy Control | Intestine |
| Cdk1 | 2.59E+00 | 5.220403 | 25.13934 | 5.24E-05 | 0.00035 | Infection | Intestine |
| Cdk2 | 3.07E-01 | 6.792687 | 0.783538 | 0.385551 | 0.615063 | Infection | Intestine |

| | | | | | | | |
|---|---|---|---|---|---|---|---|
| Cdk4 | 1.82E+00 | 3.999167 | 4.073344 | 0.055025 | 0.135358 | Infection | Intestine |
| Cdk5 | 1.52E+00 | 6.664737 | 22.27366 | 0.000106 | 0.000654 | Infection | Intestine |
| Cdk5r1 | 4.47E+00 | 2.799097 | 25.67774 | 3.82E-05 | 0.000265 | Infection | Intestine |
| Cdk5r2 | -6.05E+00 | 3.207369 | 1.997348 | 0.173194 | 0.332825 | Healthy Control | Intestine |
| Cdk5rap2 | 1.06E+00 | 6.891367 | 9.01429 | 0.00661 | 0.023268 | Infection | Intestine |
| Cdk6 | -7.50E-01 | 7.723733 | 3.161323 | 0.089166 | 0.198033 | Healthy Control | Intestine |
| Cdk7 | -8.74E+00 | 5.922859 | 12.20201 | 0.002317 | 0.009378 | Healthy Control | Intestine |
| Cdk8 | 5.94E-02 | 8.137924 | 0.05423 | 0.818029 | 0.952901 | Infection | Intestine |
| Cdk9 | 1.12E+00 | 8.466979 | 29.82947 | 1.79E-05 | 0.000138 | Infection | Intestine |
| Cdkn1a | 8.47E-01 | 7.744398 | 4.439894 | 0.048065 | 0.121034 | Infection | Intestine |
| Cdkn1b | -1.32E-01 | 9.62062 | 0.594168 | 0.449089 | 0.682909 | Healthy Control | Intestine |
| Cdkn1c | 5.18E-01 | 5.109282 | 0.516663 | 0.479257 | 0.713187 | Infection | Intestine |
| Cdkn2a | 5.79E+00 | 2.431373 | 41.32898 | 7.15E-07 | 8.41E-06 | Infection | Intestine |
| Cdkn2b | 5.70E-01 | 5.805358 | 0.535294 | 0.471827 | 0.706752 | Infection | Intestine |
| Cdkn2c | 6.80E-01 | 6.491254 | 2.09656 | 0.161778 | 0.316105 | Infection | Intestine |
| Cdkn2d | 1.09E-01 | 6.172664 | 0.037878 | 0.847402 | 0.954066 | Infection | Intestine |
| Cdx1 | -4.28E+00 | 11.89244 | 592.5646 | 2.28E-17 | 1.43E-14 | Healthy Control | Intestine |
| Cdx2 | -3.57E+00 | 10.32421 | 206.571 | 1.00E-12 | 1.14E-10 | Healthy Control | Intestine |
| Ceacam1 | -3.56E+00 | 12.48966 | 326.7772 | 1.33E-14 | 2.90E-12 | Healthy Control | Intestine |
| Cebpa | 3.72E-02 | 6.973485 | 0.00629 | 0.937501 | 0.969735 | Infection | Intestine |
| Cebpg | -7.49E-01 | 8.220642 | 5.660903 | 0.026394 | 0.074313 | Healthy Control | Intestine |
| Celf6 | -5.28E+00 | 2.703 | 0.826281 | 0.3741 | 0.600431 | Healthy Control | Intestine |
| Celsr1 | -3.69E+00 | 8.45006 | 39.85501 | 1.82E-06 | 1.83E-05 | Healthy Control | Intestine |
| Celsr2 | -1.29E+00 | 4.863217 | 0.600878 | 0.4462 | 0.68017 | Healthy Control | Intestine |
| Celsr3 | -5.44E+00 | 2.607038 | 0.530415 | 0.47449 | 0.709467 | Healthy Control | Intestine |
| Cemip | -3.90E+00 | 1.946624 | 0.254469 | 0.61811 | 0.837736 | Healthy Control | Intestine |
| Cenpa | 3.42E-01 | 7.126365 | 0.392824 | 0.537323 | 0.769405 | Infection | Intestine |
| Cenpe | 3.44E-01 | 5.854935 | 0.269906 | 0.608422 | 0.831483 | Infection | Intestine |
| Cenpf | -3.56E+00 | 7.017945 | 10.49531 | 0.003624 | 0.013817 | Healthy Control | Intestine |
| Cenps | -6.11E+00 | 3.594937 | 2.230012 | 0.151224 | 0.300096 | Healthy Control | Intestine |
| Cenpu | 1.55E+00 | 3.752087 | 1.887617 | 0.181877 | 0.345532 | Infection | Intestine |
| Cep104 | -1.02E+00 | 6.345161 | 3.632696 | 0.06857 | 0.162144 | Healthy Control | Intestine |
| Cep152 | 3.91E-01 | 6.112474 | 0.576245 | 0.455664 | 0.689541 | Infection | Intestine |
| Cep164 | -5.40E-01 | 6.344671 | 0.816956 | 0.37519 | 0.601601 | Healthy Control | Intestine |
| Cep250 | 1.28E+00 | 6.738454 | 10.48397 | 0.003809 | 0.014456 | Infection | Intestine |
| Cep290 | 5.62E-01 | 5.91042 | 0.994308 | 0.32906 | 0.54636 | Infection | Intestine |
| Cer1 | -3.57E-01 | 1.227417 | 0.006288 | 0.937908 | 0.969735 | Healthy Control | Intestine |
| Cercam | 3.00E+00 | 6.442821 | 81.24067 | 8.38E-09 | 1.93E-07 | Infection | Intestine |
| Cers1 | 2.10E+00 | 3.01452 | 2.484417 | 0.127205 | 0.260979 | Infection | Intestine |
| Cers2 | 3.04E-01 | 8.206328 | 1.677521 | 0.208821 | 0.38404 | Infection | Intestine |
| Cers4 | -5.79E-01 | 8.324142 | 5.521938 | 0.028178 | 0.078126 | Healthy Control | Intestine |
| Cers5 | -1.25E+00 | 8.317162 | 16.57345 | 0.000481 | 0.00246 | Healthy Control | Intestine |
| Cers6 | -3.70E+00 | 8.041378 | 59.49804 | 6.07E-08 | 1.06E-06 | Healthy Control | Intestine |
| Cetn2 | -1.56E+00 | 6.473743 | 5.688195 | 0.025264 | 0.071584 | Healthy Control | Intestine |
| Cetn3 | -7.92E-01 | 7.838614 | 6.350407 | 0.019473 | 0.057991 | Healthy Control | Intestine |
| Cfap299 | -4.85E+00 | 2.793678 | 0.473072 | 0.499177 | 0.732165 | Healthy Control | Intestine |
| Cfap36 | 6.33E-02 | 7.168692 | 0.041183 | 0.841057 | 0.954066 | Infection | Intestine |
| Cfap44 | -1.81E+00 | 1.554598 | 0.073495 | 0.789067 | 0.941823 | Healthy Control | Intestine |
| Cfap45 | -4.76E+00 | 2.677371 | 0.612916 | 0.441614 | 0.674718 | Healthy Control | Intestine |
| Cfap52 | -6.59E+00 | 3.759566 | 1.892501 | 0.184359 | 0.349188 | Healthy Control | Intestine |

| | | | | | | | |
|---|---|---|---|---|---|---|---|
| Cfap53 | -3.12E+00 | 1.667139 | 0.150162 | 0.70087 | 0.89357 | Healthy Control | Intestine |
| Cfap57 | -3.57E-01 | 1.228299 | 0.00618 | 0.938438 | 0.969735 | Healthy Control | Intestine |
| Cfap61 | 2.34E+00 | 2.006612 | 1.760569 | 0.194957 | 0.364435 | Infection | Intestine |
| Cfap65 | -2.83E+00 | 1.703706 | 0.168534 | 0.684522 | 0.884771 | Healthy Control | Intestine |
| Cfap69 | -4.66E+00 | 2.160248 | 0.547553 | 0.466461 | 0.702287 | Healthy Control | Intestine |
| Cfb | -1.84E+00 | 8.124282 | 12.49674 | 0.001729 | 0.007302 | Healthy Control | Intestine |
| Cfc1 | 5.04E+00 | 1.40516 | 3.793304 | 0.061373 | 0.148135 | Infection | Intestine |
| Cflar | -8.35E-01 | 7.779018 | 4.976441 | 0.036143 | 0.095428 | Healthy Control | Intestine |
| Cftr | -4.47E+00 | 8.777696 | 87.11281 | 1.95E-09 | 5.93E-08 | Healthy Control | Intestine |
| Cga | -1.37E+00 | 1.228863 | 0.033199 | 0.858049 | 0.956553 | Healthy Control | Intestine |
| Cgas | 3.64E-01 | 3.210577 | 0.046024 | 0.832097 | 0.954066 | Infection | Intestine |
| Cgn | -3.08E+00 | 9.760028 | 153.3948 | 1.62E-11 | 1.07E-09 | Healthy Control | Intestine |
| Ch25h | 4.32E+00 | 3.607087 | 53.26571 | 2.91E-07 | 3.87E-06 | Infection | Intestine |
| Champ1 | -8.82E-01 | 6.745101 | 3.002468 | 0.096192 | 0.210185 | Healthy Control | Intestine |
| Chat | -4.44E+00 | 2.082329 | 0.329249 | 0.571197 | 0.799611 | Healthy Control | Intestine |
| Chchd3 | -1.50E+00 | 9.279806 | 42.06602 | 1.65E-06 | 1.71E-05 | Healthy Control | Intestine |
| Chd1 | 7.08E-03 | 8.422009 | 0.000914 | 0.976152 | 0.996659 | Infection | Intestine |
| Chd4 | -2.25E-01 | 10.27529 | 2.30381 | 0.143425 | 0.287769 | Healthy Control | Intestine |
| Chd6 | 2.75E-01 | 6.963678 | 0.623978 | 0.437977 | 0.670904 | Infection | Intestine |
| Chd7 | -2.15E+00 | 7.742888 | 25.37859 | 3.84E-05 | 0.000265 | Healthy Control | Intestine |
| Chek1 | 1.52E+00 | 4.533495 | 4.443399 | 0.046108 | 0.117176 | Infection | Intestine |
| Chek2 | -9.94E-01 | 6.024292 | 1.388562 | 0.250056 | 0.443266 | Healthy Control | Intestine |
| Chia1 | -1.17E+00 | 1.359858 | 0.026626 | 0.871638 | 0.962378 | Healthy Control | Intestine |
| Chic2 | 1.84E+00 | 6.745056 | 42.94448 | 1.43E-06 | 1.51E-05 | Infection | Intestine |
| Chil1 | -1.78E+00 | 1.359469 | 0.07024 | 0.793087 | 0.943769 | Healthy Control | Intestine |
| Chil3 | 1.05E+01 | 4.130829 | 719.0253 | 4.83E-24 | 2.42E-20 | Infection | Intestine |
| Chil4 | -1.37E+00 | 1.228863 | 0.033199 | 0.858049 | 0.956553 | Healthy Control | Intestine |
| Chka | -1.01E+00 | 6.788799 | 4.948253 | 0.035876 | 0.095002 | Healthy Control | Intestine |
| Chmp3 | -8.28E-01 | 8.273319 | 6.659026 | 0.017097 | 0.051808 | Healthy Control | Intestine |
| Chmp4c | -5.72E+00 | 8.913208 | 74.32128 | 1.40E-08 | 2.93E-07 | Healthy Control | Intestine |
| Chrd | 1.93E+00 | 3.536567 | 2.289702 | 0.143003 | 0.287268 | Infection | Intestine |
| Chrm1 | -9.72E+00 | 6.532197 | 17.82359 | 0.000427 | 0.00222 | Healthy Control | Intestine |
| Chrm2 | -5.93E+00 | 2.845591 | 1.628361 | 0.216409 | 0.395237 | Healthy Control | Intestine |
| Chrm3 | -7.06E+00 | 3.50571 | 3.151868 | 0.08884 | 0.197746 | Healthy Control | Intestine |
| Chrm4 | 2.04E+00 | 2.009123 | 1.140619 | 0.294764 | 0.502584 | Infection | Intestine |
| Chrm5 | -4.95E-01 | 1.228692 | 0.008683 | 0.927069 | 0.969735 | Healthy Control | Intestine |
| Chrna1 | -8.79E-01 | 1.316323 | 0.024248 | 0.877627 | 0.966433 | Healthy Control | Intestine |
| Chrna2 | -1.25E+00 | 1.360808 | 0.036437 | 0.850189 | 0.955127 | Healthy Control | Intestine |
| Chrna4 | -8.80E-01 | 1.317885 | 0.020487 | 0.887443 | 0.969534 | Healthy Control | Intestine |
| Chrna6 | 4.81E+00 | 1.409028 | 8.149989 | 0.009923 | 0.03263 | Infection | Intestine |
| Chrna7 | -2.88E+00 | 1.666663 | 0.22592 | 0.637786 | 0.853063 | Healthy Control | Intestine |
| Chrnb1 | 8.56E-01 | 4.763973 | 0.651294 | 0.427553 | 0.660391 | Infection | Intestine |
| Chrnb2 | 5.61E-01 | 4.018844 | 0.17857 | 0.676268 | 0.880238 | Infection | Intestine |
| Chrnb3 | -2.32E+00 | 1.401894 | 0.101833 | 0.752018 | 0.925646 | Healthy Control | Intestine |
| Chrnb4 | -2.22E+00 | 5.241152 | 2.584942 | 0.122285 | 0.253179 | Healthy Control | Intestine |
| Chrnd | -8.62E-01 | 1.275087 | 0.020018 | 0.889226 | 0.969599 | Healthy Control | Intestine |
| Chrng | -9.71E-16 | 1.180616 | 0 | 1 | 1 | Healthy Control | Intestine |
| Chst10 | -5.54E+00 | 2.922215 | 1.05314 | 0.317294 | 0.531584 | Healthy Control | Intestine |
| Chst8 | -5.01E+00 | 2.816167 | 0.554561 | 0.465087 | 0.70085 | Healthy Control | Intestine |
| Chtop | 5.04E-02 | 7.964194 | 0.023935 | 0.878472 | 0.96715 | Infection | Intestine |

| | | | | | | | |
|---|---|---|---|---|---|---|---|
| Chuk | 1.25E-01 | 7.647498 | 0.167734 | 0.686119 | 0.885076 | Infection | Intestine |
| Ciart | -3.77E+00 | 2.031918 | 0.262703 | 0.612774 | 0.833799 | Healthy Control | Intestine |
| Cib2 | -8.69E-02 | 4.592848 | 0.006421 | 0.936774 | 0.969735 | Healthy Control | Intestine |
| Cic | -1.85E-01 | 8.383015 | 0.57926 | 0.454762 | 0.688816 | Healthy Control | Intestine |
| Cidea | -4.31E+00 | 1.920714 | 0.533246 | 0.471151 | 0.706162 | Healthy Control | Intestine |
| Cideb | -3.68E+00 | 7.898774 | 26.9166 | 2.74E-05 | 0.0002 | Healthy Control | Intestine |
| Ciita | 3.87E-01 | 6.652226 | 0.156652 | 0.697451 | 0.892753 | Infection | Intestine |
| Cisd2 | -1.29E+00 | 8.83222 | 25.374 | 4.86E-05 | 0.000327 | Healthy Control | Intestine |
| Cish | 1.52E+00 | 6.385733 | 15.01379 | 0.000822 | 0.003887 | Infection | Intestine |
| Cit | -5.96E-01 | 6.735679 | 1.226197 | 0.279689 | 0.483186 | Healthy Control | Intestine |
| Cited1 | -7.56E-01 | 1.274309 | 0.015326 | 0.902889 | 0.969735 | Healthy Control | Intestine |
| Cited2 | -1.37E+00 | 8.655633 | 23.37715 | 7.99E-05 | 0.000511 | Healthy Control | Intestine |
| Cited4 | -8.67E+00 | 5.897163 | 9.227983 | 0.006549 | 0.023104 | Healthy Control | Intestine |
| Ckb | 1.37E+00 | 7.786019 | 16.53427 | 0.000521 | 0.002624 | Infection | Intestine |
| Cklf | -5.54E+00 | 3.208534 | 1.123287 | 0.302025 | 0.512347 | Healthy Control | Intestine |
| Cks1b | 1.50E+00 | 5.793362 | 9.51057 | 0.005439 | 0.019659 | Infection | Intestine |
| Clca3a2 | -3.15E+00 | 6.491998 | 4.731796 | 0.040377 | 0.104838 | Healthy Control | Intestine |
| Clcn4 | -2.16E+00 | 6.43343 | 5.797159 | 0.024043 | 0.069013 | Healthy Control | Intestine |
| Clcn7 | -6.80E-01 | 6.966098 | 2.974474 | 0.098138 | 0.213229 | Healthy Control | Intestine |
| Clcnka | -6.42E-01 | 1.273114 | 0.013189 | 0.909769 | 0.969735 | Healthy Control | Intestine |
| Clcnkb | -4.70E+00 | 2.038688 | 0.581385 | 0.455422 | 0.689399 | Healthy Control | Intestine |
| Cldn1 | -1.06E-01 | 4.639273 | 0.006838 | 0.934796 | 0.969735 | Healthy Control | Intestine |
| Cldn10 | 3.60E+00 | 1.896307 | 4.072319 | 0.051724 | 0.128884 | Infection | Intestine |
| Cldn12 | -1.67E+00 | 7.957445 | 13.30885 | 0.00131 | 0.005787 | Healthy Control | Intestine |
| Cldn14 | -6.34E+00 | 3.964399 | 1.44059 | 0.242431 | 0.431889 | Healthy Control | Intestine |
| Cldn16 | -2.15E+00 | 1.317599 | 0.073256 | 0.78953 | 0.941823 | Healthy Control | Intestine |
| Cldn17 | -3.05E+00 | 1.633014 | 0.281979 | 0.598913 | 0.823556 | Healthy Control | Intestine |
| Cldn19 | -9.71E-16 | 1.180616 | 0 | 1 | 1 | Healthy Control | Intestine |
| Cldn2 | -5.21E+00 | 9.752222 | 58.34701 | 9.94E-08 | 1.58E-06 | Healthy Control | Intestine |
| Cldn23 | -1.86E+00 | 5.820663 | 4.313464 | 0.049214 | 0.123616 | Healthy Control | Intestine |
| Cldn6 | 5.70E+00 | 1.229995 | 6.086704 | 0.031631 | 0.085684 | Infection | Intestine |
| Cldn7 | -4.83E+00 | 11.5827 | 707.3861 | 2.33E-18 | 2.34E-15 | Healthy Control | Intestine |
| Cldn8 | -4.04E+00 | 8.559572 | 40.68594 | 1.51E-06 | 1.57E-05 | Healthy Control | Intestine |
| Cldn9 | -3.92E+00 | 2.15692 | 0.504201 | 0.48458 | 0.718888 | Healthy Control | Intestine |
| Clec10a | 1.90E+00 | 4.078282 | 2.195018 | 0.151929 | 0.301211 | Infection | Intestine |
| Clec14a | -2.50E+00 | 6.67506 | 5.606124 | 0.0263 | 0.074131 | Healthy Control | Intestine |
| Clec1b | -1.26E+00 | 1.400425 | 0.029211 | 0.865731 | 0.960453 | Healthy Control | Intestine |
| Clec4e | 6.69E+00 | 1.230035 | 8.855416 | 0.012839 | 0.040654 | Infection | Intestine |
| Clec4n | 2.85E-01 | 5.52246 | 0.084858 | 0.773347 | 0.934664 | Infection | Intestine |
| Clec9a | -7.14E+00 | 4.399212 | 0.653102 | 0.432553 | 0.665037 | Healthy Control | Intestine |
| Clgn | -3.21E+00 | 1.446713 | 0.145226 | 0.707649 | 0.897011 | Healthy Control | Intestine |
| Clic3 | 4.14E+00 | 1.631032 | 4.699677 | 0.038503 | 0.100756 | Infection | Intestine |
| Clic4 | 1.89E-02 | 8.753227 | 0.004498 | 0.947139 | 0.975994 | Infection | Intestine |
| Clic5 | -4.03E+00 | 8.680636 | 29.82118 | 2.00E-05 | 0.000153 | Healthy Control | Intestine |
| Clic6 | -1.12E+01 | 8.636239 | 67.76925 | 8.12E-08 | 1.34E-06 | Healthy Control | Intestine |
| Clk1 | 1.35E-02 | 9.009434 | 0.005801 | 0.939985 | 0.97062 | Infection | Intestine |
| Clmp | 6.05E-01 | 6.952314 | 1.351211 | 0.257642 | 0.453216 | Infection | Intestine |
| Cln3 | 4.35E-01 | 6.346453 | 0.889046 | 0.355844 | 0.580457 | Infection | Intestine |
| Cln6 | -1.70E+00 | 8.308005 | 21.72795 | 0.000113 | 0.00069 | Healthy Control | Intestine |
| Cln8 | 4.86E-01 | 5.645552 | 0.595533 | 0.448019 | 0.682112 | Infection | Intestine |

| | | | | | | | |
|---|---|---|---|---|---|---|---|
| Clock | -7.24E-01 | 8.366693 | 6.841842 | 0.015714 | 0.0482 | Healthy Control | Intestine |
| Clspn | -1.85E+00 | 5.366756 | 2.182905 | 0.153159 | 0.302929 | Healthy Control | Intestine |
| Cltb | -6.64E-01 | 8.137147 | 4.338029 | 0.049137 | 0.123484 | Healthy Control | Intestine |
| Cltc | -8.94E-01 | 10.90034 | 46.83109 | 7.47E-07 | 8.73E-06 | Healthy Control | Intestine |
| Cltrn | -2.97E+00 | 1.595756 | 0.204413 | 0.654296 | 0.864378 | Healthy Control | Intestine |
| Cmah | -1.65E+00 | 6.805827 | 1.601446 | 0.223535 | 0.406029 | Healthy Control | Intestine |
| Cmip | -2.36E+00 | 9.406703 | 90.09056 | 2.88E-09 | 7.92E-08 | Healthy Control | Intestine |
| Cmklr1 | -1.77E+00 | 6.280889 | 3.519523 | 0.072582 | 0.169234 | Healthy Control | Intestine |
| Cmtm6 | 6.27E-01 | 9.358429 | 20.70652 | 0.00016 | 0.00093 | Infection | Intestine |
| Cndp1 | -7.56E-01 | 1.274175 | 0.01535 | 0.902812 | 0.969735 | Healthy Control | Intestine |
| Cnga2 | -7.56E-01 | 1.274175 | 0.01535 | 0.902812 | 0.969735 | Healthy Control | Intestine |
| Cngb3 | -9.77E-01 | 1.318065 | 0.022209 | 0.882868 | 0.96867 | Healthy Control | Intestine |
| Cnmd | -9.71E-16 | 1.180616 | 0 | 1 | 1 | Healthy Control | Intestine |
| Cnn1 | 7.28E-01 | 7.292228 | 0.709346 | 0.414238 | 0.64681 | Infection | Intestine |
| Cnot11 | -1.23E-03 | 5.688369 | 2.88E-06 | 0.998661 | 1 | Healthy Control | Intestine |
| Cnot2 | 7.71E-01 | 7.216118 | 7.794078 | 0.010685 | 0.03471 | Infection | Intestine |
| Cnot4 | -3.54E-01 | 6.08988 | 0.268893 | 0.608865 | 0.831569 | Healthy Control | Intestine |
| Cnot8 | -1.93E+00 | 6.868967 | 6.114331 | 0.020936 | 0.061506 | Healthy Control | Intestine |
| Cnot9 | -5.62E-01 | 7.362623 | 2.626553 | 0.119142 | 0.24857 | Healthy Control | Intestine |
| Cnp | -1.90E+00 | 8.105528 | 29.12343 | 1.71E-05 | 0.000133 | Healthy Control | Intestine |
| Cnppd1 | 6.67E-01 | 8.627752 | 6.979212 | 0.014968 | 0.046253 | Infection | Intestine |
| Cnpy1 | -9.77E-01 | 1.318419 | 0.021469 | 0.884821 | 0.96867 | Healthy Control | Intestine |
| Cnr2 | -6.07E+00 | 3.396236 | 1.099223 | 0.307018 | 0.518708 | Healthy Control | Intestine |
| Cntf | 1.89E-01 | 3.554336 | 0.013681 | 0.907916 | 0.969735 | Infection | Intestine |
| Cntn1 | -5.07E+00 | 2.796264 | 0.68996 | 0.415598 | 0.648465 | Healthy Control | Intestine |
| Cntn2 | -2.16E+00 | 1.318852 | 0.08494 | 0.773823 | 0.934669 | Healthy Control | Intestine |
| Cntn3 | -4.63E+00 | 2.550001 | 0.436228 | 0.514799 | 0.750025 | Healthy Control | Intestine |
| Cntn4 | -3.86E+00 | 1.863564 | 0.335974 | 0.5667 | 0.794649 | Healthy Control | Intestine |
| Cntn6 | -6.13E+00 | 3.879122 | 1.047001 | 0.318579 | 0.533202 | Healthy Control | Intestine |
| Cntnap1 | 4.47E-01 | 5.048945 | 0.178196 | 0.676672 | 0.8804 | Infection | Intestine |
| Cntnap3 | -5.84E+00 | 2.362837 | 1.22198 | 0.281444 | 0.485072 | Healthy Control | Intestine |
| Cntnap4 | 1.71E+00 | 2.752753 | 0.525682 | 0.474836 | 0.709562 | Infection | Intestine |
| Cntnap5b | -3.57E-01 | 1.227509 | 0.006273 | 0.937979 | 0.969735 | Healthy Control | Intestine |
| Cntrob | 3.18E-01 | 5.260992 | 0.118225 | 0.733957 | 0.915572 | Infection | Intestine |
| Coa7 | -2.23E+00 | 5.598158 | 2.980829 | 0.097858 | 0.212766 | Healthy Control | Intestine |
| Col10a1 | -9.71E-16 | 1.180616 | 0 | 1 | 1 | Healthy Control | Intestine |
| Col12a1 | 6.18E-02 | 6.439534 | 0.007423 | 0.93208 | 0.969735 | Infection | Intestine |
| Col13a1 | -5.10E-01 | 3.774466 | 0.06799 | 0.796718 | 0.943769 | Healthy Control | Intestine |
| Col14a1 | 4.78E-01 | 8.649344 | 0.553054 | 0.470182 | 0.705343 | Infection | Intestine |
| Col15a1 | 2.75E+00 | 8.441204 | 71.52397 | 3.01E-07 | 4.00E-06 | Infection | Intestine |
| Col18a1 | 6.88E-01 | 8.534964 | 3.06375 | 0.097883 | 0.212766 | Infection | Intestine |
| Col1a1 | 1.74E+00 | 9.841738 | 45.64499 | 5.01E-06 | 4.47E-05 | Infection | Intestine |
| Col1a2 | 3.28E+00 | 12.65092 | 433.5202 | 6.98E-13 | 8.74E-11 | Infection | Intestine |
| Col23a1 | 1.09E+00 | 4.916016 | 2.1618 | 0.154815 | 0.30512 | Infection | Intestine |
| Col24a1 | 4.84E+00 | 3.691554 | 78.80788 | 1.20E-08 | 2.60E-07 | Infection | Intestine |
| Col26a1 | -2.44E+00 | 1.526273 | 0.096185 | 0.760321 | 0.928591 | Healthy Control | Intestine |
| Col4a1 | 1.96E+00 | 9.476031 | 74.01699 | 9.06E-08 | 1.47E-06 | Infection | Intestine |
| Col4a2 | 1.87E+00 | 8.893711 | 32.90993 | 3.26E-05 | 0.000231 | Infection | Intestine |
| Col4a3 | 2.55E+00 | 2.518202 | 1.937805 | 0.175291 | 0.336209 | Infection | Intestine |
| Col4a3bp | -4.26E-01 | 8.222042 | 2.46947 | 0.130442 | 0.266419 | Healthy Control | Intestine |

| | | | | | | | |
|---|---|---|---|---|---|---|---|
| Col4a5 | 3.94E-01 | 6.473648 | 0.546031 | 0.467752 | 0.703385 | Infection | Intestine |
| Col5a1 | 1.67E+00 | 8.380876 | 25.74166 | 8.09E-05 | 0.000517 | Infection | Intestine |
| Col6a1 | 2.94E+00 | 8.907707 | 131.4783 | 3.26E-09 | 8.78E-08 | Infection | Intestine |
| Col6a2 | 2.13E+00 | 10.81197 | 70.47094 | 8.92E-07 | 1.02E-05 | Infection | Intestine |
| Col6a3 | 2.65E+00 | 9.830965 | 103.9983 | 4.75E-08 | 8.59E-07 | Infection | Intestine |
| Col7a1 | 3.80E+00 | 3.665101 | 50.60366 | 4.12E-07 | 5.26E-06 | Infection | Intestine |
| Col9a2 | -3.00E+00 | 1.627886 | 0.16655 | 0.686531 | 0.885082 | Healthy Control | Intestine |
| Col9a3 | -4.13E+00 | 2.2281 | 0.416307 | 0.525292 | 0.759344 | Healthy Control | Intestine |
| Colgalt2 | 1.25E+00 | 3.079724 | 1.037515 | 0.318841 | 0.533462 | Infection | Intestine |
| Colq | -7.39E-01 | 3.980037 | 0.230952 | 0.63558 | 0.851056 | Healthy Control | Intestine |
| Commd1 | -2.34E+00 | 8.851571 | 72.81173 | 1.65E-08 | 3.35E-07 | Healthy Control | Intestine |
| Commd10 | -1.22E+00 | 6.549402 | 4.875897 | 0.037052 | 0.097572 | Healthy Control | Intestine |
| Comt | -9.84E-01 | 8.64304 | 12.21729 | 0.002053 | 0.008444 | Healthy Control | Intestine |
| Cop1 | 1.82E-01 | 6.81727 | 0.199985 | 0.658986 | 0.867142 | Infection | Intestine |
| Copg1 | -1.52E-02 | 8.613176 | 0.004314 | 0.948231 | 0.976918 | Healthy Control | Intestine |
| Coprs | -5.79E-01 | 6.479569 | 1.058714 | 0.314048 | 0.527558 | Healthy Control | Intestine |
| Cops2 | -2.21E-01 | 8.897909 | 0.635678 | 0.433887 | 0.666066 | Healthy Control | Intestine |
| Cops5 | -9.33E-01 | 6.555888 | 1.936334 | 0.176892 | 0.338243 | Healthy Control | Intestine |
| Cops8 | -6.03E-01 | 7.908727 | 3.246287 | 0.085364 | 0.191457 | Healthy Control | Intestine |
| Cops9 | -2.73E-01 | 6.966425 | 0.345832 | 0.562293 | 0.790462 | Healthy Control | Intestine |
| Coq8a | -2.86E+00 | 8.465469 | 24.53273 | 4.72E-05 | 0.000319 | Healthy Control | Intestine |
| Coro1c | 2.49E-01 | 9.636788 | 2.839649 | 0.106225 | 0.22725 | Infection | Intestine |
| Coro2b | 1.23E+00 | 5.553491 | 6.529903 | 0.017885 | 0.054032 | Infection | Intestine |
| Coro6 | 1.73E+00 | 2.164335 | 1.053809 | 0.313464 | 0.527108 | Infection | Intestine |
| Cort | 6.20E-01 | 3.2444 | 0.130033 | 0.721687 | 0.906818 | Infection | Intestine |
| Cox7a1 | 2.21E+00 | 1.949192 | 1.201605 | 0.282051 | 0.485873 | Infection | Intestine |
| Cox8b | -1.89E+00 | 1.401878 | 0.051671 | 0.82188 | 0.953086 | Healthy Control | Intestine |
| Cpa4 | -1.52E+00 | 1.273479 | 0.043253 | 0.837684 | 0.954066 | Healthy Control | Intestine |
| Cpeb2 | -1.05E+00 | 6.9919 | 3.266772 | 0.083527 | 0.188519 | Healthy Control | Intestine |
| Cpeb4 | -3.45E-01 | 8.946868 | 3.235659 | 0.085931 | 0.192383 | Healthy Control | Intestine |
| Cplx3 | -1.89E+00 | 1.402407 | 0.060055 | 0.808247 | 0.94887 | Healthy Control | Intestine |
| Cpn1 | -1.56E+00 | 4.59193 | 1.183102 | 0.28869 | 0.494079 | Healthy Control | Intestine |
| Cpne4 | -3.94E+00 | 1.952416 | 0.218363 | 0.644176 | 0.857095 | Healthy Control | Intestine |
| Cpox | -2.14E+00 | 7.932672 | 16.56354 | 0.000452 | 0.002334 | Healthy Control | Intestine |
| Cps1 | -8.70E-01 | 5.097434 | 0.157094 | 0.695891 | 0.892492 | Healthy Control | Intestine |
| Cpsf6 | -3.87E-02 | 8.693933 | 0.045998 | 0.832176 | 0.954066 | Healthy Control | Intestine |
| Cpt1a | -2.85E+00 | 9.36178 | 60.09495 | 9.07E-08 | 1.47E-06 | Healthy Control | Intestine |
| Cpt1c | 2.75E+00 | 4.79299 | 26.70501 | 3.59E-05 | 0.000252 | Infection | Intestine |
| Cpt2 | -1.89E+00 | 8.075937 | 20.81471 | 0.00014 | 0.000824 | Healthy Control | Intestine |
| Cptp | -3.16E+00 | 6.485652 | 12.54362 | 0.001778 | 0.007493 | Healthy Control | Intestine |
| Cr1l | -2.88E-01 | 7.933345 | 0.788388 | 0.384236 | 0.613356 | Healthy Control | Intestine |
| Cr2 | -4.19E+00 | 2.131008 | 0.36101 | 0.553411 | 0.782305 | Healthy Control | Intestine |
| Crabp1 | 1.66E+00 | 2.836107 | 0.669744 | 0.420817 | 0.652985 | Infection | Intestine |
| Crabp2 | 4.22E+00 | 1.980861 | 11.48178 | 0.001888 | 0.007869 | Infection | Intestine |
| Crb1 | 3.92E+00 | 1.364607 | 3.329367 | 0.083426 | 0.188376 | Infection | Intestine |
| Crb2 | -2.01E+00 | 1.557632 | 0.06327 | 0.803239 | 0.947264 | Healthy Control | Intestine |
| Crbn | 1.26E+00 | 7.506173 | 22.32374 | 0.000105 | 0.000649 | Infection | Intestine |
| Creb1 | -3.87E-01 | 7.838438 | 1.052599 | 0.316064 | 0.530056 | Healthy Control | Intestine |
| Crebbp | -7.99E-01 | 8.459742 | 11.72755 | 0.002443 | 0.009806 | Healthy Control | Intestine |
| Crebrf | 1.69E+00 | 8.050978 | 95.76132 | 1.95E-09 | 5.93E-08 | Infection | Intestine |

| | | | | | | | |
|---|---|---|---|---|---|---|---|
| Creg1 | 8.44E-01 | 5.428483 | 2.045072 | 0.166289 | 0.321779 | Infection | Intestine |
| Creld1 | 1.83E+00 | 5.643474 | 10.41897 | 0.003885 | 0.014699 | Infection | Intestine |
| Crh | -1.71E+00 | 1.318797 | 0.046958 | 0.830578 | 0.954066 | Healthy Control | Intestine |
| Crhbp | -9.71E-16 | 1.180616 | 0 | 1 | 1 | Healthy Control | Intestine |
| Crhr1 | -2.33E+00 | 1.403932 | 0.104375 | 0.74905 | 0.924753 | Healthy Control | Intestine |
| Crhr2 | 3.57E+00 | 1.486379 | 3.196245 | 0.083594 | 0.188585 | Infection | Intestine |
| Crisp1 | -2.70E+00 | 1.560445 | 0.135377 | 0.715189 | 0.901639 | Healthy Control | Intestine |
| Crk | -8.42E-01 | 9.613431 | 26.56361 | 3.74E-05 | 0.000261 | Healthy Control | Intestine |
| Crlf2 | -4.62E-01 | 5.972758 | 0.27689 | 0.603568 | 0.826665 | Healthy Control | Intestine |
| Crmp1 | -3.76E+00 | 2.150021 | 0.378171 | 0.544092 | 0.775484 | Healthy Control | Intestine |
| Crocc | 1.08E+00 | 5.330586 | 2.308412 | 0.142477 | 0.286441 | Infection | Intestine |
| Crtc1 | -3.13E+00 | 1.524611 | 0.18286 | 0.672107 | 0.877942 | Healthy Control | Intestine |
| Crtc2 | 2.37E-01 | 6.298348 | 0.267868 | 0.609782 | 0.83218 | Infection | Intestine |
| Crx | -1.52E+00 | 1.275042 | 0.038572 | 0.846592 | 0.954066 | Healthy Control | Intestine |
| Crxos | -4.61E+00 | 2.305972 | 0.656684 | 0.426133 | 0.658881 | Healthy Control | Intestine |
| Cry1 | -1.34E+00 | 5.104396 | 2.063653 | 0.165078 | 0.320054 | Healthy Control | Intestine |
| Cry2 | 9.72E-01 | 6.350115 | 3.359826 | 0.080434 | 0.18344 | Infection | Intestine |
| Cryaa | 4.28E+00 | 1.363484 | 3.532155 | 0.073546 | 0.171082 | Infection | Intestine |
| Cryab | 9.08E-01 | 6.545511 | 3.027624 | 0.095939 | 0.209818 | Infection | Intestine |
| Cryba2 | -5.07E+00 | 2.619283 | 0.757269 | 0.394503 | 0.623582 | Healthy Control | Intestine |
| Cryba4 | -4.06E+00 | 2.113425 | 0.400587 | 0.532501 | 0.765785 | Healthy Control | Intestine |
| Crybb3 | -5.29E-01 | 3.92564 | 0.167413 | 0.686344 | 0.885076 | Healthy Control | Intestine |
| Crygb | -9.71E-16 | 1.180616 | 0 | 1 | 1 | Healthy Control | Intestine |
| Crygc | -1.08E+00 | 1.35936 | 0.026842 | 0.871226 | 0.962378 | Healthy Control | Intestine |
| Crygd | -7.56E-01 | 1.273813 | 0.014863 | 0.90436 | 0.969735 | Healthy Control | Intestine |
| Cryge | -9.71E-16 | 1.180616 | 0 | 1 | 1 | Healthy Control | Intestine |
| Crygf | -3.57E-01 | 1.227417 | 0.006288 | 0.937908 | 0.969735 | Healthy Control | Intestine |
| Crygs | 6.14E+00 | 1.275304 | 6.507456 | 0.022915 | 0.066385 | Infection | Intestine |
| Crym | -2.14E+00 | 5.593184 | 4.409928 | 0.047081 | 0.118855 | Healthy Control | Intestine |
| Cs | -7.93E-01 | 10.83304 | 23.06987 | 8.69E-05 | 0.000549 | Healthy Control | Intestine |
| Csde1 | -1.73E+00 | 10.39593 | 91.98539 | 2.84E-09 | 7.88E-08 | Healthy Control | Intestine |
| Cse1l | -1.85E+00 | 6.726155 | 8.797511 | 0.006644 | 0.023339 | Healthy Control | Intestine |
| Csf1 | 1.71E+00 | 7.755346 | 38.3411 | 3.25E-06 | 3.06E-05 | Infection | Intestine |
| Csf1r | -1.00E+00 | 8.360751 | 6.834798 | 0.015847 | 0.048519 | Healthy Control | Intestine |
| Csf2 | -2.76E+00 | 1.596848 | 0.135895 | 0.714685 | 0.901639 | Healthy Control | Intestine |
| Csf2rb | 3.19E+00 | 5.648946 | 46.26197 | 8.26E-07 | 9.52E-06 | Infection | Intestine |
| Csf2rb2 | 4.24E-01 | 4.9259 | 0.208741 | 0.651841 | 0.862801 | Infection | Intestine |
| Csf3 | -1.78E+00 | 1.360333 | 0.064533 | 0.801477 | 0.946496 | Healthy Control | Intestine |
| Csf3r | 1.67E+00 | 3.711483 | 1.764357 | 0.196419 | 0.366648 | Infection | Intestine |
| Csk | -6.67E-01 | 7.468654 | 3.460051 | 0.075975 | 0.175107 | Healthy Control | Intestine |
| Csnk1a1 | -7.25E-01 | 9.379804 | 17.18854 | 0.00043 | 0.002229 | Healthy Control | Intestine |
| Csnk1d | 1.55E-01 | 9.738978 | 1.227024 | 0.280058 | 0.483309 | Infection | Intestine |
| Csnk1e | 2.12E+00 | 6.914324 | 44.85396 | 1.04E-06 | 1.15E-05 | Infection | Intestine |
| Csnk2a1 | -2.22E-01 | 8.354488 | 0.617862 | 0.440305 | 0.673346 | Healthy Control | Intestine |
| Csnk2a2 | -1.27E+00 | 7.208145 | 6.782897 | 0.01571 | 0.0482 | Healthy Control | Intestine |
| Cspg4 | 3.26E+00 | 5.66078 | 55.70708 | 1.95E-07 | 2.75E-06 | Infection | Intestine |
| Cspg5 | 1.98E+00 | 2.675655 | 2.962167 | 0.098236 | 0.213348 | Infection | Intestine |
| Cst7 | 1.67E+00 | 2.444295 | 0.750111 | 0.394972 | 0.624125 | Infection | Intestine |
| Csta1 | -2.68E+00 | 1.595796 | 0.098891 | 0.755659 | 0.927619 | Healthy Control | Intestine |
| Ctbp1 | -2.56E+00 | 9.349914 | 104.1132 | 7.45E-10 | 2.54E-08 | Healthy Control | Intestine |

| | | | | | | | |
|---|---|---|---|---|---|---|---|
| Ctbp2 | -4.02E-01 | 8.146973 | 2.167111 | 0.155142 | 0.305644 | Healthy Control | Intestine |
| Ctcf | 9.17E-02 | 8.393096 | 0.189653 | 0.667493 | 0.874273 | Infection | Intestine |
| Ctdsp1 | -2.81E-01 | 9.927518 | 3.651206 | 0.069269 | 0.163489 | Healthy Control | Intestine |
| Ctdspl2 | 8.30E-01 | 6.708453 | 3.841691 | 0.062856 | 0.151145 | Infection | Intestine |
| Cth | -1.25E+00 | 8.882424 | 26.51462 | 3.77E-05 | 0.000262 | Healthy Control | Intestine |
| Ctla4 | 1.91E+00 | 2.113654 | 0.876576 | 0.357762 | 0.582236 | Infection | Intestine |
| Ctnna1 | -3.28E-01 | 10.21755 | 7.196289 | 0.013663 | 0.042791 | Healthy Control | Intestine |
| Ctnna3 | 2.07E+00 | 2.213071 | 0.788287 | 0.38253 | 0.611556 | Infection | Intestine |
| Ctnnal1 | 1.29E+00 | 6.407571 | 10.96024 | 0.003198 | 0.012372 | Infection | Intestine |
| Ctnnb1 | -1.88E+00 | 11.34247 | 145.727 | 4.00E-11 | 2.33E-09 | Healthy Control | Intestine |
| Ctnnbip1 | -2.06E-01 | 6.65806 | 0.163137 | 0.690018 | 0.888369 | Healthy Control | Intestine |
| Ctnnd1 | -7.83E-01 | 11.00374 | 37.2071 | 4.01E-06 | 3.67E-05 | Healthy Control | Intestine |
| Ctr9 | -3.32E-01 | 8.147962 | 1.55365 | 0.225803 | 0.409703 | Healthy Control | Intestine |
| Ctsk | 2.49E+00 | 5.881579 | 23.56024 | 7.60E-05 | 0.000488 | Infection | Intestine |
| Ctsl | -5.67E-01 | 7.729003 | 1.825068 | 0.190446 | 0.357604 | Healthy Control | Intestine |
| Ctsq | 5.42E+00 | 1.321064 | 6.075822 | 0.025212 | 0.071509 | Infection | Intestine |
| Cttn | -1.01E+00 | 9.659683 | 22.82069 | 9.28E-05 | 0.00058 | Healthy Control | Intestine |
| Cttnbp2 | -2.60E+00 | 6.061793 | 5.933856 | 0.0232 | 0.067015 | Healthy Control | Intestine |
| Cubn | -5.48E+00 | 3.254152 | 0.628134 | 0.435802 | 0.66839 | Healthy Control | Intestine |
| Cul1 | -1.56E+00 | 8.467479 | 27.07447 | 3.07E-05 | 0.000219 | Healthy Control | Intestine |
| Cul3 | -7.87E-01 | 9.586319 | 23.62988 | 7.58E-05 | 0.000487 | Healthy Control | Intestine |
| Cul4a | -1.26E+00 | 8.432637 | 25.01399 | 5.21E-05 | 0.000348 | Healthy Control | Intestine |
| Cux1 | -9.53E-01 | 8.591297 | 10.72938 | 0.003458 | 0.013277 | Healthy Control | Intestine |
| Cux2 | 2.38E+00 | 1.893632 | 1.263158 | 0.269963 | 0.469369 | Infection | Intestine |
| Cwc27 | -8.26E-02 | 6.271133 | 0.021331 | 0.885156 | 0.96867 | Healthy Control | Intestine |
| Cx3cl1 | 1.17E+00 | 5.48924 | 4.116579 | 0.054241 | 0.13389 | Infection | Intestine |
| Cx3cr1 | -2.48E+00 | 7.572582 | 9.899258 | 0.004351 | 0.016216 | Healthy Control | Intestine |
| Cxadr | -4.59E+00 | 10.0561 | 285.2568 | 8.50E-15 | 1.93E-12 | Healthy Control | Intestine |
| Cxcl1 | 3.85E+00 | 2.109688 | 7.03627 | 0.013541 | 0.042501 | Infection | Intestine |
| Cxcl10 | -6.47E+00 | 3.508471 | 0.56779 | 0.462235 | 0.697813 | Healthy Control | Intestine |
| Cxcl12 | 1.09E+00 | 9.386477 | 10.39369 | 0.005574 | 0.020062 | Infection | Intestine |
| Cxcl13 | -8.99E-01 | 4.939522 | 0.403939 | 0.53139 | 0.765288 | Healthy Control | Intestine |
| Cxcl14 | 8.49E-01 | 6.900015 | 2.780067 | 0.109776 | 0.232856 | Infection | Intestine |
| Cxcl16 | -1.05E-01 | 7.944698 | 0.043755 | 0.836587 | 0.954066 | Healthy Control | Intestine |
| Cxcl2 | -3.57E-01 | 1.228299 | 0.00618 | 0.938438 | 0.969735 | Healthy Control | Intestine |
| Cxcl3 | 6.16E+00 | 1.365587 | 10.83642 | 0.00371 | 0.01409 | Infection | Intestine |
| Cxcl5 | 4.45E+00 | 1.931005 | 8.718719 | 0.005612 | 0.020181 | Infection | Intestine |
| Cxcl9 | -7.19E+00 | 4.226896 | 0.157538 | 0.699628 | 0.89357 | Healthy Control | Intestine |
| Cxcr1 | -6.42E-01 | 1.273555 | 0.012735 | 0.91133 | 0.969735 | Healthy Control | Intestine |
| Cxcr2 | 7.01E+00 | 1.409473 | 16.40024 | 0.000645 | 0.003166 | Infection | Intestine |
| Cxcr3 | 1.32E-01 | 3.418293 | 0.00882 | 0.92601 | 0.969735 | Infection | Intestine |
| Cxcr4 | 1.84E-01 | 5.923124 | 0.105747 | 0.74784 | 0.924698 | Infection | Intestine |
| Cxcr5 | -3.16E+00 | 1.730115 | 0.190194 | 0.666127 | 0.872997 | Healthy Control | Intestine |
| Cxcr6 | -7.31E-01 | 4.050122 | 0.118743 | 0.73364 | 0.915572 | Healthy Control | Intestine |
| Cxxc1 | 8.55E-01 | 7.626649 | 10.38587 | 0.003951 | 0.014929 | Infection | Intestine |
| Cxxc5 | -2.70E-01 | 7.246578 | 0.298613 | 0.590211 | 0.815962 | Healthy Control | Intestine |
| Cyb561 | -3.08E+00 | 8.263168 | 34.23144 | 4.99E-06 | 4.46E-05 | Healthy Control | Intestine |
| Cyb5a | 4.73E-01 | 9.320705 | 10.31866 | 0.004046 | 0.015241 | Infection | Intestine |
| Cyb5r3 | 2.72E-01 | 9.158815 | 1.798929 | 0.193651 | 0.362398 | Infection | Intestine |
| Cybb | 1.63E+00 | 6.496138 | 6.72666 | 0.017994 | 0.054296 | Infection | Intestine |

| | | | | | | | |
|---|---|---|---|---|---|---|---|
| Cybc1 | -4.34E-01 | 7.257147 | 1.770482 | 0.196718 | 0.367041 | Healthy Control | Intestine |
| Cyct | -3.77E+00 | 2.156959 | 0.294908 | 0.591969 | 0.817489 | Healthy Control | Intestine |
| Cyfip1 | -5.36E-01 | 9.004994 | 9.033224 | 0.006558 | 0.02312 | Healthy Control | Intestine |
| Cyfip2 | 1.51E+00 | 6.671134 | 7.998129 | 0.010313 | 0.033721 | Infection | Intestine |
| Cygb | 2.04E+00 | 7.296631 | 31.17185 | 1.63E-05 | 0.000128 | Infection | Intestine |
| Cyld | 7.83E-01 | 8.450237 | 22.07559 | 0.000112 | 0.000687 | Infection | Intestine |
| Cyp11a1 | -4.64E+00 | 2.912244 | 0.382764 | 0.541476 | 0.774023 | Healthy Control | Intestine |
| Cyp11b2 | -9.71E-16 | 1.180616 | 0 | 1 | 1 | Healthy Control | Intestine |
| Cyp17a1 | -1.37E+00 | 1.228088 | 0.037791 | 0.848672 | 0.954066 | Healthy Control | Intestine |
| Cyp19a1 | -1.37E+00 | 1.228088 | 0.037791 | 0.848672 | 0.954066 | Healthy Control | Intestine |
| Cyp1a1 | -1.94E+00 | 1.403386 | 0.055803 | 0.814907 | 0.951358 | Healthy Control | Intestine |
| Cyp1a2 | -8.80E-01 | 1.318018 | 0.016319 | 0.89947 | 0.969735 | Healthy Control | Intestine |
| Cyp1b1 | 2.33E+00 | 2.010345 | 1.078569 | 0.31185 | 0.525277 | Infection | Intestine |
| Cyp21a1 | -1.83E+00 | 1.361299 | 0.05979 | 0.808786 | 0.94887 | Healthy Control | Intestine |
| Cyp24a1 | -4.95E-01 | 1.228515 | 0.008877 | 0.926263 | 0.969735 | Healthy Control | Intestine |
| Cyp26a1 | 7.32E+00 | 1.722135 | 42.64617 | 4.14E-07 | 5.27E-06 | Infection | Intestine |
| Cyp26b1 | 2.68E+00 | 2.539229 | 3.609591 | 0.067074 | 0.159513 | Infection | Intestine |
| Cyp27a1 | -2.23E-02 | 7.228619 | 0.003164 | 0.955652 | 0.981696 | Healthy Control | Intestine |
| Cyp27b1 | 2.97E+00 | 2.03926 | 4.410959 | 0.044325 | 0.113615 | Infection | Intestine |
| Cyp2a5 | -4.95E-01 | 1.228515 | 0.008877 | 0.926263 | 0.969735 | Healthy Control | Intestine |
| Cyp2c29 | 5.33E+00 | 1.275264 | 4.750983 | 0.046013 | 0.117162 | Infection | Intestine |
| Cyp2e1 | -2.54E+00 | 1.4848 | 0.111874 | 0.740116 | 0.920731 | Healthy Control | Intestine |
| Cyp2g1 | 5.45E+00 | 1.361791 | 3.595064 | 0.070708 | 0.165481 | Infection | Intestine |
| Cyp2r1 | -4.36E+00 | 2.460065 | 0.422597 | 0.522424 | 0.757167 | Healthy Control | Intestine |
| Cyp3a13 | -1.04E+01 | 7.408711 | 21.58525 | 0.000159 | 0.000925 | Healthy Control | Intestine |
| Cyp7a1 | -9.71E-16 | 1.180616 | 0 | 1 | 1 | Healthy Control | Intestine |
| Cyp7b1 | 4.06E+00 | 3.435831 | 26.22174 | 4.05E-05 | 0.000279 | Infection | Intestine |
| Cyp8b1 | -1.71E+00 | 1.320053 | 0.045602 | 0.833003 | 0.954066 | Healthy Control | Intestine |
| Cyren | 2.78E+00 | 3.135467 | 6.246287 | 0.019696 | 0.058418 | Infection | Intestine |
| Cysltr1 | 3.04E+00 | 4.979816 | 32.95079 | 9.35E-06 | 7.79E-05 | Infection | Intestine |
| D1Ertd622e | -9.43E+00 | 6.576217 | 18.77095 | 0.00033 | 0.001773 | Healthy Control | Intestine |
| D1Pas1 | 4.49E+00 | 1.319505 | 3.88395 | 0.064399 | 0.154102 | Infection | Intestine |
| Daam2 | 1.90E-01 | 5.386098 | 0.068439 | 0.795855 | 0.943769 | Infection | Intestine |
| Dab2 | -2.50E-01 | 7.142529 | 0.155811 | 0.696792 | 0.892753 | Healthy Control | Intestine |
| Dach1 | 5.91E-01 | 6.477482 | 1.352171 | 0.257316 | 0.452926 | Infection | Intestine |
| Dact2 | -2.04E+00 | 7.399088 | 10.9037 | 0.003006 | 0.0117 | Healthy Control | Intestine |
| Dag1 | -2.14E+00 | 8.967587 | 60.96859 | 8.45E-08 | 1.39E-06 | Healthy Control | Intestine |
| Dagla | -8.71E+00 | 5.864461 | 8.491077 | 0.008642 | 0.029053 | Healthy Control | Intestine |
| Daglb | -4.67E-01 | 6.678696 | 0.758997 | 0.392705 | 0.622311 | Healthy Control | Intestine |
| Dao | -2.65E+00 | 1.665541 | 0.127603 | 0.723468 | 0.908144 | Healthy Control | Intestine |
| Dapk1 | 1.06E+00 | 7.25333 | 10.30645 | 0.004071 | 0.015321 | Infection | Intestine |
| Daxx | 1.92E+00 | 6.584081 | 47.48559 | 6.75E-07 | 7.97E-06 | Infection | Intestine |
| Dazl | -2.58E+00 | 1.825035 | 0.103831 | 0.75023 | 0.92508 | Healthy Control | Intestine |
| Dbh | -3.76E+00 | 1.670209 | 0.217792 | 0.644509 | 0.857177 | Healthy Control | Intestine |
| Dbn1 | 2.10E+00 | 6.967804 | 25.69329 | 4.59E-05 | 0.000312 | Infection | Intestine |
| Dbnl | -2.48E-01 | 8.865338 | 2.058498 | 0.165569 | 0.320633 | Healthy Control | Intestine |
| Dbp | 1.35E+00 | 6.787134 | 1.639141 | 0.221832 | 0.403521 | Infection | Intestine |
| Dbx1 | -1.07E+00 | 1.318449 | 0.020183 | 0.888478 | 0.969599 | Healthy Control | Intestine |
| Dbx2 | 6.35E+00 | 1.36411 | 12.58686 | 0.002089 | 0.008563 | Infection | Intestine |
| Dcaf1 | -2.60E+00 | 7.480589 | 18.29404 | 0.000263 | 0.001453 | Healthy Control | Intestine |

| | | | | | | | |
|---|---|---|---|---|---|---|---|
| Dcaf11 | -2.95E+00 | 7.276394 | 19.2276 | 0.000195 | 0.001112 | Healthy Control | Intestine |
| Dcaf7 | -2.21E-01 | 8.681493 | 1.153991 | 0.294478 | 0.502268 | Healthy Control | Intestine |
| Dcc | -9.71E-16 | 1.180616 | 0 | 1 | 1 | Healthy Control | Intestine |
| Dct | -3.62E+00 | 1.917986 | 0.277162 | 0.602912 | 0.826445 | Healthy Control | Intestine |
| Dctn1 | 3.91E-01 | 9.40176 | 7.480333 | 0.012148 | 0.038934 | Infection | Intestine |
| Dctn2 | -5.61E-01 | 9.479527 | 14.07677 | 0.001117 | 0.005067 | Healthy Control | Intestine |
| Dctn3 | -2.09E-01 | 6.329077 | 0.145272 | 0.706551 | 0.896577 | Healthy Control | Intestine |
| Dctn4 | 8.98E-01 | 6.74228 | 7.997861 | 0.009833 | 0.032448 | Infection | Intestine |
| Dcx | -3.70E+00 | 1.998678 | 0.224867 | 0.639277 | 0.854004 | Healthy Control | Intestine |
| Ddah1 | -2.92E+00 | 7.002337 | 16.4846 | 0.000474 | 0.002434 | Healthy Control | Intestine |
| Ddb1 | 1.14E-01 | 9.130859 | 0.425022 | 0.52126 | 0.755917 | Infection | Intestine |
| Ddc | -9.38E+00 | 6.829884 | 20.8364 | 0.000192 | 0.0011 | Healthy Control | Intestine |
| Ddit3 | 4.67E-01 | 6.436822 | 1.217872 | 0.281489 | 0.485072 | Infection | Intestine |
| Ddit4 | -4.06E-01 | 8.807245 | 1.335689 | 0.26071 | 0.457134 | Healthy Control | Intestine |
| Ddr1 | -6.82E-01 | 8.127405 | 5.746607 | 0.02546 | 0.072047 | Healthy Control | Intestine |
| Ddx19b | 7.72E-01 | 6.558016 | 2.584897 | 0.122177 | 0.253154 | Infection | Intestine |
| Ddx21 | -1.18E+00 | 8.266155 | 20.9331 | 0.000145 | 0.00085 | Healthy Control | Intestine |
| Ddx4 | 4.66E+00 | 1.275965 | 4.05477 | 0.063807 | 0.152904 | Infection | Intestine |
| Ddx41 | -9.01E-01 | 6.822987 | 4.299015 | 0.049349 | 0.123744 | Healthy Control | Intestine |
| Ddx42 | -2.14E+00 | 8.235083 | 42.47919 | 1.11E-06 | 1.23E-05 | Healthy Control | Intestine |
| Ddx56 | -1.69E+00 | 7.601831 | 15.42408 | 0.000658 | 0.003222 | Healthy Control | Intestine |
| Ddx58 | 7.36E-01 | 6.773121 | 2.891553 | 0.103214 | 0.222231 | Infection | Intestine |
| Ddx6 | 1.02E+00 | 9.577221 | 63.45104 | 6.75E-08 | 1.17E-06 | Infection | Intestine |
| Decr1 | -2.04E+00 | 8.223999 | 26.50955 | 3.23E-05 | 0.000229 | Healthy Control | Intestine |
| Def6 | 1.70E+00 | 4.892205 | 9.936423 | 0.004518 | 0.016741 | Infection | Intestine |
| Defa5 | 4.96E+00 | 1.488956 | 5.062834 | 0.031766 | 0.085956 | Infection | Intestine |
| Dek | 8.78E-01 | 8.414455 | 21.34192 | 0.000136 | 0.000803 | Infection | Intestine |
| Dennd1a | 9.67E-02 | 6.557618 | 0.039133 | 0.844957 | 0.954066 | Infection | Intestine |
| Derl2 | 9.75E-01 | 6.891259 | 9.3022 | 0.005906 | 0.021115 | Infection | Intestine |
| Des | 1.56E+00 | 8.190806 | 12.50592 | 0.002924 | 0.011443 | Infection | Intestine |
| Dffb | -5.80E-01 | 5.966191 | 0.424091 | 0.521047 | 0.755917 | Healthy Control | Intestine |
| Dgat1 | -2.29E-02 | 7.141044 | 0.003289 | 0.954782 | 0.981445 | Healthy Control | Intestine |
| Dgcr2 | -1.34E+00 | 9.4663 | 50.79856 | 4.02E-07 | 5.16E-06 | Healthy Control | Intestine |
| Dgcr8 | 1.00E+00 | 6.697796 | 5.607774 | 0.027165 | 0.076141 | Infection | Intestine |
| Dgka | -4.61E-02 | 7.255696 | 0.015789 | 0.901112 | 0.969735 | Healthy Control | Intestine |
| Dgkg | 5.27E-01 | 6.407113 | 0.989704 | 0.33062 | 0.548586 | Infection | Intestine |
| Dgkq | -6.76E-02 | 7.317046 | 0.038693 | 0.845859 | 0.954066 | Healthy Control | Intestine |
| Dglucy | -1.21E+00 | 7.469627 | 8.095989 | 0.009171 | 0.030524 | Healthy Control | Intestine |
| Dhfr | -2.24E+00 | 6.602182 | 5.541954 | 0.026865 | 0.075469 | Healthy Control | Intestine |
| Dhh | 2.52E+00 | 2.530814 | 3.868635 | 0.060027 | 0.145308 | Infection | Intestine |
| Dhodh | -7.82E+00 | 5.178934 | 4.912821 | 0.038539 | 0.100757 | Healthy Control | Intestine |
| Dhx33 | 1.22E+00 | 6.611352 | 9.510014 | 0.005448 | 0.019663 | Infection | Intestine |
| Dhx58 | 3.30E-01 | 5.97243 | 0.25387 | 0.619057 | 0.838248 | Infection | Intestine |
| Diablo | -4.69E-01 | 6.817313 | 1.111453 | 0.302791 | 0.513242 | Healthy Control | Intestine |
| Diaph1 | 7.57E-01 | 9.420086 | 23.30158 | 8.20E-05 | 0.000522 | Infection | Intestine |
| Diaph3 | 8.05E-01 | 5.217361 | 1.357896 | 0.255705 | 0.450566 | Infection | Intestine |
| Dicer1 | -4.76E-01 | 7.920529 | 2.06942 | 0.164402 | 0.319238 | Healthy Control | Intestine |
| Dio1 | -2.75E+00 | 6.049093 | 5.532788 | 0.027815 | 0.077486 | Healthy Control | Intestine |
| Dio2 | 3.24E+00 | 1.668525 | 1.798235 | 0.188423 | 0.35447 | Infection | Intestine |
| Dio3 | -1.78E+00 | 1.359439 | 0.053902 | 0.818234 | 0.952901 | Healthy Control | Intestine |

| | | | | | | | |
|---|---|---|---|---|---|---|---|
| Dip2a | -4.16E-01 | 8.594084 | 3.213045 | 0.086922 | 0.19411 | Healthy Control | Intestine |
| Diras1 | 3.04E+00 | 3.449662 | 15.0997 | 0.000732 | 0.003508 | Infection | Intestine |
| Disc1 | 1.74E+00 | 2.281218 | 1.000309 | 0.327049 | 0.544104 | Infection | Intestine |
| Disp2 | -6.41E+00 | 3.706807 | 1.529373 | 0.230764 | 0.417192 | Healthy Control | Intestine |
| Dixdc1 | 1.61E-01 | 4.212295 | 0.015089 | 0.903246 | 0.969735 | Infection | Intestine |
| Dkk1 | -1.94E+00 | 1.405142 | 0.051586 | 0.821908 | 0.953086 | Healthy Control | Intestine |
| Dkk2 | 3.17E+00 | 2.625349 | 5.542825 | 0.026286 | 0.074131 | Infection | Intestine |
| Dkk3 | -8.15E-02 | 7.869802 | 0.033652 | 0.856143 | 0.956553 | Healthy Control | Intestine |
| Dkk4 | -7.55E-01 | 1.273079 | 0.015675 | 0.901795 | 0.969735 | Healthy Control | Intestine |
| Dlg1 | 3.33E-01 | 9.027161 | 4.431557 | 0.047041 | 0.118812 | Infection | Intestine |
| Dlg3 | -2.20E+00 | 7.833849 | 23.96208 | 5.58E-05 | 0.00037 | Healthy Control | Intestine |
| Dlg4 | 5.42E-01 | 6.054883 | 1.287615 | 0.268401 | 0.467555 | Infection | Intestine |
| Dlg5 | 1.55E-01 | 7.47741 | 0.363007 | 0.553047 | 0.782305 | Infection | Intestine |
| Dlgap4 | -2.55E-01 | 8.226461 | 1.386084 | 0.251744 | 0.445166 | Healthy Control | Intestine |
| Dlk1 | -9.71E-16 | 1.180616 | 0 | 1 | 1 | Healthy Control | Intestine |
| Dll1 | -4.69E-01 | 6.794082 | 0.755389 | 0.393563 | 0.623079 | Healthy Control | Intestine |
| Dll3 | 2.45E+00 | 1.832356 | 1.044934 | 0.314412 | 0.527993 | Infection | Intestine |
| Dll4 | -4.64E-01 | 5.897388 | 0.31614 | 0.579068 | 0.806375 | Healthy Control | Intestine |
| Dlx1 | -1.15E+00 | 1.320248 | 0.026284 | 0.87327 | 0.96333 | Healthy Control | Intestine |
| Dlx2 | -1.99E+00 | 1.403455 | 0.051859 | 0.821537 | 0.953086 | Healthy Control | Intestine |
| Dlx3 | -9.71E-16 | 1.180616 | 0 | 1 | 1 | Healthy Control | Intestine |
| Dlx4 | -4.94E-01 | 1.22813 | 0.009389 | 0.924171 | 0.969735 | Healthy Control | Intestine |
| Dlx5 | -4.95E-01 | 1.228692 | 0.008683 | 0.927069 | 0.969735 | Healthy Control | Intestine |
| Dlx6 | -8.80E-01 | 1.318204 | 0.019988 | 0.888814 | 0.969599 | Healthy Control | Intestine |
| Dmap1 | 5.02E-01 | 6.523023 | 0.793636 | 0.382619 | 0.611556 | Infection | Intestine |
| Dmbx1 | -1.25E+00 | 1.36134 | 0.028397 | 0.867561 | 0.96063 | Healthy Control | Intestine |
| Dmc1 | -4.41E+00 | 2.185345 | 0.482932 | 0.493896 | 0.726969 | Healthy Control | Intestine |
| Dmd | -1.94E+00 | 6.436809 | 5.581797 | 0.026527 | 0.074561 | Healthy Control | Intestine |
| Dmp1 | 6.27E+00 | 1.908291 | 42.85523 | 1.21E-07 | 1.86E-06 | Infection | Intestine |
| Dmrt1 | -1.37E+00 | 1.229023 | 0.032546 | 0.859435 | 0.956553 | Healthy Control | Intestine |
| Dmrt2 | -1.83E+00 | 1.360792 | 0.048566 | 0.827333 | 0.953685 | Healthy Control | Intestine |
| Dmrta1 | -4.67E+00 | 1.926355 | 0.714208 | 0.406007 | 0.637538 | Healthy Control | Intestine |
| Dmrta2 | -4.49E+00 | 2.386569 | 0.652627 | 0.428159 | 0.661048 | Healthy Control | Intestine |
| Dnah11 | -2.75E+00 | 1.444809 | 0.135723 | 0.715403 | 0.901639 | Healthy Control | Intestine |
| Dnah12 | -3.57E-01 | 1.227509 | 0.006273 | 0.937979 | 0.969735 | Healthy Control | Intestine |
| Dnah5 | -6.19E+00 | 2.997318 | 1.089103 | 0.3077 | 0.519547 | Healthy Control | Intestine |
| Dnajb6 | 1.58E-01 | 8.291933 | 0.599744 | 0.447005 | 0.68119 | Infection | Intestine |
| Dnajc5 | 4.35E-01 | 8.357017 | 4.278311 | 0.050683 | 0.126418 | Infection | Intestine |
| Dnajc7 | -9.75E-01 | 8.833949 | 14.59768 | 0.000943 | 0.004402 | Healthy Control | Intestine |
| Dner | -5.74E+00 | 3.405086 | 1.02613 | 0.323334 | 0.539358 | Healthy Control | Intestine |
| Dnm1 | 2.57E+00 | 7.249293 | 42.43614 | 3.43E-06 | 3.21E-05 | Infection | Intestine |
| Dnm1l | -1.44E+00 | 8.702783 | 32.74973 | 9.37E-06 | 7.79E-05 | Healthy Control | Intestine |
| Dnm2 | -4.59E-01 | 10.13616 | 9.667599 | 0.00515 | 0.018735 | Healthy Control | Intestine |
| Dnmt1 | 5.21E-01 | 7.593166 | 3.224329 | 0.086451 | 0.193203 | Infection | Intestine |
| Dnmt3a | -2.25E-01 | 7.293506 | 0.320398 | 0.576973 | 0.804772 | Healthy Control | Intestine |
| Dnmt3b | 2.43E+00 | 3.769839 | 6.393463 | 0.018691 | 0.055927 | Infection | Intestine |
| Dnmt3l | -6.32E+00 | 3.688292 | 1.513972 | 0.233039 | 0.419487 | Healthy Control | Intestine |
| Dnttip1 | -1.06E+00 | 8.050427 | 12.88573 | 0.001616 | 0.006894 | Healthy Control | Intestine |
| Dnttip2 | -1.79E+00 | 7.033286 | 12.87473 | 0.001464 | 0.006358 | Healthy Control | Intestine |
| Doc2a | -3.57E-01 | 1.228299 | 0.00618 | 0.938438 | 0.969735 | Healthy Control | Intestine |

| | | | | | | | |
|---|---|---|---|---|---|---|---|
| Dock2 | 3.09E-01 | 6.591518 | 0.246237 | 0.62464 | 0.84103 | Infection | Intestine |
| Dock3 | -2.44E+00 | 1.444839 | 0.06434 | 0.801428 | 0.946496 | Healthy Control | Intestine |
| Dock5 | 5.75E-01 | 7.50372 | 3.077246 | 0.093446 | 0.205532 | Infection | Intestine |
| Dock7 | 1.23E+00 | 7.771809 | 25.6624 | 4.63E-05 | 0.000313 | Infection | Intestine |
| Dock8 | 2.90E-01 | 6.937825 | 0.598694 | 0.447341 | 0.681494 | Infection | Intestine |
| Dot1l | 9.23E-01 | 6.635228 | 7.032582 | 0.014585 | 0.045237 | Infection | Intestine |
| Dpf2 | -2.40E-01 | 8.467025 | 1.228862 | 0.279719 | 0.483186 | Healthy Control | Intestine |
| Dpf3 | -9.00E-01 | 5.150894 | 0.634269 | 0.433541 | 0.666066 | Healthy Control | Intestine |
| Dpm1 | -1.66E-01 | 7.973024 | 0.241231 | 0.628226 | 0.843445 | Healthy Control | Intestine |
| Dpp10 | 4.28E+00 | 1.527911 | 5.455827 | 0.027056 | 0.075878 | Infection | Intestine |
| Dpp4 | -2.18E+00 | 7.233545 | 15.21254 | 0.000695 | 0.003364 | Healthy Control | Intestine |
| Dpp9 | 1.44E-01 | 7.777388 | 0.24479 | 0.625725 | 0.841585 | Infection | Intestine |
| Dppa3 | -2.28E+00 | 1.362379 | 0.093449 | 0.762637 | 0.929438 | Healthy Control | Intestine |
| Dpy30 | -2.11E+00 | 9.349558 | 82.06926 | 7.06E-09 | 1.68E-07 | Healthy Control | Intestine |
| Dpyd | -2.27E+00 | 7.276349 | 12.44935 | 0.001664 | 0.007066 | Healthy Control | Intestine |
| Dpysl2 | -1.94E+00 | 8.917089 | 41.55349 | 1.70E-06 | 1.74E-05 | Healthy Control | Intestine |
| Dpysl3 | 4.31E-01 | 7.863705 | 1.51427 | 0.231623 | 0.417689 | Infection | Intestine |
| Dpysl4 | 2.86E+00 | 2.190174 | 3.368194 | 0.076911 | 0.176694 | Infection | Intestine |
| Dpysl5 | -4.44E+00 | 2.034896 | 0.397248 | 0.533818 | 0.765939 | Healthy Control | Intestine |
| Drc7 | -2.06E+00 | 1.274832 | 0.084916 | 0.774502 | 0.934669 | Healthy Control | Intestine |
| Drd1 | -3.06E+00 | 1.765614 | 0.123291 | 0.727997 | 0.912201 | Healthy Control | Intestine |
| Drd2 | -1.17E+00 | 1.359666 | 0.026742 | 0.871363 | 0.962378 | Healthy Control | Intestine |
| Drd3 | 3.85E+00 | 1.48583 | 2.530359 | 0.12194 | 0.252871 | Infection | Intestine |
| Drd4 | -1.74E+00 | 1.484032 | 0.041035 | 0.840883 | 0.954066 | Healthy Control | Intestine |
| Drd5 | -1.17E+00 | 1.361479 | 0.029597 | 0.864735 | 0.96005 | Healthy Control | Intestine |
| Drosha | -1.27E+00 | 8.04324 | 14.31449 | 0.001002 | 0.004632 | Healthy Control | Intestine |
| Dsc1 | -1.94E+00 | 1.40322 | 0.056246 | 0.814187 | 0.950961 | Healthy Control | Intestine |
| Dsc3 | -1.34E+00 | 1.401481 | 0.046968 | 0.830031 | 0.954066 | Healthy Control | Intestine |
| Dscam | -2.97E+00 | 1.596084 | 0.153536 | 0.697831 | 0.892753 | Healthy Control | Intestine |
| Dscaml1 | -1.83E+00 | 1.361084 | 0.060735 | 0.807312 | 0.94887 | Healthy Control | Intestine |
| Dsg1a | 2.76E+00 | 1.600493 | 1.789081 | 0.191053 | 0.358475 | Infection | Intestine |
| Dsg1b | 3.34E+00 | 1.485717 | 2.153461 | 0.153533 | 0.303368 | Infection | Intestine |
| Dsg2 | -3.49E+00 | 9.726109 | 115.2666 | 1.98E-10 | 8.71E-09 | Healthy Control | Intestine |
| Dsp | -4.60E+00 | 10.44149 | 207.4408 | 4.16E-13 | 5.63E-11 | Healthy Control | Intestine |
| Dspp | -1.52E+00 | 1.273388 | 0.043339 | 0.837524 | 0.954066 | Healthy Control | Intestine |
| Dst | 3.39E-01 | 9.209884 | 4.917328 | 0.037331 | 0.098212 | Infection | Intestine |
| Dtl | -4.35E-01 | 5.944648 | 0.424195 | 0.521025 | 0.755917 | Healthy Control | Intestine |
| Dtnbp1 | -1.99E+00 | 7.63774 | 18.41317 | 0.000256 | 0.001418 | Healthy Control | Intestine |
| Dusp1 | 7.11E-01 | 6.131294 | 1.551468 | 0.225726 | 0.409703 | Infection | Intestine |
| Dusp11 | 1.07E+00 | 7.450019 | 15.87109 | 0.000637 | 0.003139 | Infection | Intestine |
| Dusp15 | -3.57E-01 | 1.227387 | 0.006293 | 0.937883 | 0.969735 | Healthy Control | Intestine |
| Dusp3 | 1.00E+00 | 6.761684 | 7.284969 | 0.013172 | 0.041497 | Infection | Intestine |
| Dusp4 | 3.72E+00 | 4.570142 | 58.75968 | 1.19E-07 | 1.84E-06 | Infection | Intestine |
| Dusp5 | 2.91E+00 | 5.856187 | 56.92188 | 1.60E-07 | 2.32E-06 | Infection | Intestine |
| Dusp6 | 2.00E-01 | 9.134867 | 0.859341 | 0.364081 | 0.588691 | Infection | Intestine |
| Dusp7 | 1.20E+00 | 5.809827 | 5.265942 | 0.031425 | 0.085449 | Infection | Intestine |
| Dut | -7.79E+00 | 5.026294 | 5.647441 | 0.027692 | 0.077277 | Healthy Control | Intestine |
| Duxbl1 | -3.69E+00 | 2.078721 | 0.414623 | 0.525546 | 0.759493 | Healthy Control | Intestine |
| Dvl1 | -9.78E-02 | 8.58285 | 0.223019 | 0.641448 | 0.855379 | Healthy Control | Intestine |
| Dvl2 | -8.25E-01 | 5.854223 | 1.338092 | 0.25866 | 0.454346 | Healthy Control | Intestine |

| | | | | | | | |
|---|---|---|---|---|---|---|---|
| Dvl3 | -1.55E+00 | 7.795269 | 19.08111 | 0.000223 | 0.001256 | Healthy Control | Intestine |
| Dxo | 1.13E+00 | 7.176875 | 8.582332 | 0.007815 | 0.026722 | Infection | Intestine |
| Dydc2 | -4.95E-01 | 1.228692 | 0.008683 | 0.927069 | 0.969735 | Healthy Control | Intestine |
| Dync1h1 | -1.25E+00 | 9.875202 | 49.01322 | 5.31E-07 | 6.55E-06 | Healthy Control | Intestine |
| Dync1i1 | -6.55E+00 | 3.753102 | 2.177213 | 0.155875 | 0.306822 | Healthy Control | Intestine |
| Dync1i2 | -3.65E+00 | 7.954752 | 40.46183 | 1.40E-06 | 1.48E-05 | Healthy Control | Intestine |
| Dync1li1 | -1.25E+00 | 8.32483 | 16.4409 | 0.000522 | 0.002624 | Healthy Control | Intestine |
| Dync2h1 | 1.21E+00 | 5.537024 | 5.177791 | 0.032735 | 0.087868 | Infection | Intestine |
| Dyrk1a | -1.36E+00 | 8.257759 | 19.45308 | 0.000215 | 0.001214 | Healthy Control | Intestine |
| Dysf | -5.99E-01 | 5.701468 | 0.370586 | 0.54824 | 0.778138 | Healthy Control | Intestine |
| E2f1 | 1.44E-01 | 5.372971 | 0.037412 | 0.848253 | 0.954066 | Infection | Intestine |
| E2f2 | 1.18E+00 | 6.286004 | 4.859856 | 0.038101 | 0.099859 | Infection | Intestine |
| E2f3 | -4.25E-01 | 5.756219 | 0.292956 | 0.593405 | 0.818794 | Healthy Control | Intestine |
| E2f4 | -9.71E-01 | 8.208409 | 10.52017 | 0.0037 | 0.014063 | Healthy Control | Intestine |
| E2f5 | -1.55E+00 | 7.871115 | 22.97628 | 7.79E-05 | 0.0005 | Healthy Control | Intestine |
| Ebf2 | -2.63E+00 | 1.668226 | 0.120639 | 0.730835 | 0.91327 | Healthy Control | Intestine |
| Ebf3 | 1.65E+00 | 2.894688 | 1.072679 | 0.31075 | 0.523777 | Infection | Intestine |
| Ebi3 | 2.18E+00 | 4.094701 | 9.765076 | 0.004755 | 0.01749 | Infection | Intestine |
| Ece1 | -1.10E+00 | 9.70417 | 28.89772 | 2.20E-05 | 0.000166 | Healthy Control | Intestine |
| Ecel1 | -3.57E-01 | 1.228299 | 0.00618 | 0.938438 | 0.969735 | Healthy Control | Intestine |
| Echdc1 | -1.65E+00 | 7.736962 | 12.38187 | 0.001834 | 0.007676 | Healthy Control | Intestine |
| Echs1 | -1.91E+00 | 10.22345 | 127.7771 | 1.41E-10 | 6.97E-09 | Healthy Control | Intestine |
| Eci3 | -6.73E+00 | 4.373779 | 1.902173 | 0.183292 | 0.347561 | Healthy Control | Intestine |
| Ecpas | -2.05E-01 | 8.614379 | 0.745673 | 0.397253 | 0.626197 | Healthy Control | Intestine |
| Ecscr | -7.87E+00 | 4.860358 | 4.504283 | 0.046624 | 0.118156 | Healthy Control | Intestine |
| Ect2 | -2.02E+00 | 6.972957 | 4.893782 | 0.036405 | 0.095968 | Healthy Control | Intestine |
| Eda | 7.11E-01 | 3.243469 | 0.206847 | 0.653477 | 0.863896 | Infection | Intestine |
| Edar | 9.17E-01 | 2.944215 | 0.199912 | 0.658985 | 0.867142 | Infection | Intestine |
| Edc4 | -3.18E-01 | 8.352482 | 1.929625 | 0.178836 | 0.341285 | Healthy Control | Intestine |
| Edil3 | -6.92E-01 | 6.109599 | 0.311561 | 0.582085 | 0.808493 | Healthy Control | Intestine |
| Edn1 | -3.96E+00 | 7.434509 | 8.194907 | 0.009685 | 0.032067 | Healthy Control | Intestine |
| Edn3 | -6.55E+00 | 3.45452 | 2.075642 | 0.165441 | 0.320511 | Healthy Control | Intestine |
| Ednra | 1.22E+00 | 5.813892 | 3.491468 | 0.074986 | 0.173547 | Infection | Intestine |
| Ednrb | -7.18E-02 | 6.480361 | 0.00981 | 0.921981 | 0.969735 | Healthy Control | Intestine |
| Eea1 | 3.18E-01 | 8.405611 | 2.254686 | 0.147575 | 0.294554 | Infection | Intestine |
| Eed | -6.61E-01 | 8.420558 | 7.09744 | 0.014188 | 0.044116 | Healthy Control | Intestine |
| Eepd1 | 9.17E-01 | 6.64326 | 5.678265 | 0.026327 | 0.074167 | Infection | Intestine |
| Efcab14 | -2.73E+00 | 8.418815 | 69.35469 | 1.64E-08 | 3.33E-07 | Healthy Control | Intestine |
| Efcab3 | -1.52E+00 | 1.273358 | 0.043368 | 0.83747 | 0.954066 | Healthy Control | Intestine |
| Efemp1 | -7.06E-02 | 7.402675 | 0.018918 | 0.891864 | 0.969735 | Healthy Control | Intestine |
| Efna1 | -2.19E+00 | 8.468605 | 14.31801 | 0.001058 | 0.004852 | Healthy Control | Intestine |
| Efna2 | -5.33E+00 | 2.933454 | 1.149284 | 0.296679 | 0.505264 | Healthy Control | Intestine |
| Efna3 | -6.30E+00 | 3.949745 | 1.034567 | 0.321383 | 0.536717 | Healthy Control | Intestine |
| Efna5 | 2.15E+00 | 3.825752 | 5.123455 | 0.033165 | 0.088784 | Infection | Intestine |
| Efnb1 | -4.17E+00 | 9.033656 | 91.41299 | 1.18E-09 | 3.81E-08 | Healthy Control | Intestine |
| Efnb2 | -2.99E+00 | 8.060246 | 28.87926 | 1.55E-05 | 0.000122 | Healthy Control | Intestine |
| Efnb3 | 1.92E+00 | 3.536875 | 3.610025 | 0.069427 | 0.163539 | Infection | Intestine |
| Egfl7 | -1.00E+00 | 7.2153 | 0.433433 | 0.521125 | 0.755917 | Healthy Control | Intestine |
| Egfr | 3.57E-01 | 7.553554 | 1.723107 | 0.20288 | 0.375875 | Infection | Intestine |
| Egln1 | -3.82E-01 | 7.872855 | 1.846933 | 0.187916 | 0.354038 | Healthy Control | Intestine |

| | | | | | | | |
|---|---|---|---|---|---|---|---|
| Egln2 | -1.96E+00 | 8.799635 | 40.84357 | 1.82E-06 | 1.82E-05 | Healthy Control | Intestine |
| Egln3 | 7.35E-01 | 6.670039 | 1.847873 | 0.187524 | 0.353444 | Infection | Intestine |
| Egr1 | 4.54E-02 | 6.216824 | 0.000783 | 0.978049 | 0.997985 | Infection | Intestine |
| Egr2 | 1.80E+00 | 2.488638 | 0.857095 | 0.363056 | 0.587794 | Infection | Intestine |
| Egr3 | 4.09E+00 | 3.485994 | 16.72726 | 0.00047 | 0.00242 | Infection | Intestine |
| Egr4 | 6.70E+00 | 1.322684 | 12.4169 | 0.003235 | 0.012486 | Infection | Intestine |
| Ehd4 | -1.36E+00 | 7.852232 | 16.54396 | 0.000495 | 0.002517 | Healthy Control | Intestine |
| Ehhadh | -4.16E+00 | 7.564782 | 30.01555 | 1.45E-05 | 0.000115 | Healthy Control | Intestine |
| Ehmt1 | -7.27E-01 | 6.637551 | 1.825013 | 0.189514 | 0.355987 | Healthy Control | Intestine |
| Ehmt2 | -3.47E-01 | 9.672526 | 5.066925 | 0.034818 | 0.092564 | Healthy Control | Intestine |
| Eif2a | 1.28E-01 | 8.611596 | 0.308222 | 0.584432 | 0.810434 | Infection | Intestine |
| Eif2ak2 | -1.76E+00 | 7.80567 | 14.84296 | 0.000789 | 0.003753 | Healthy Control | Intestine |
| Eif2ak3 | 3.18E-01 | 8.294494 | 1.87716 | 0.184626 | 0.34956 | Infection | Intestine |
| Eif2ak4 | 1.61E-01 | 6.247143 | 0.10875 | 0.744613 | 0.923113 | Infection | Intestine |
| Eif2b1 | -1.91E+00 | 9.091785 | 52.90698 | 2.56E-07 | 3.45E-06 | Healthy Control | Intestine |
| Eif2s1 | 2.25E+00 | 4.373642 | 14.21651 | 0.000997 | 0.00461 | Infection | Intestine |
| Eif2s2 | 6.24E-02 | 7.937119 | 0.027792 | 0.869129 | 0.960878 | Infection | Intestine |
| Eif3i | -7.78E-02 | 8.328261 | 0.077043 | 0.78396 | 0.940206 | Healthy Control | Intestine |
| Eif4a3 | -4.35E-01 | 9.357655 | 5.333367 | 0.030795 | 0.083918 | Healthy Control | Intestine |
| Eif4e | -1.17E+00 | 7.207374 | 9.638787 | 0.004923 | 0.018054 | Healthy Control | Intestine |
| Eif4ebp1 | -6.74E+00 | 4.04542 | 2.487225 | 0.13068 | 0.266797 | Healthy Control | Intestine |
| Eif4ebp2 | -2.74E-01 | 9.82423 | 2.644368 | 0.11829 | 0.247041 | Healthy Control | Intestine |
| Eif4enif1 | -1.13E+00 | 8.164983 | 18.23207 | 0.000308 | 0.00167 | Healthy Control | Intestine |
| Eif4g1 | -1.62E+00 | 9.938292 | 92.64118 | 2.67E-09 | 7.55E-08 | Healthy Control | Intestine |
| Eif4g3 | -6.41E-01 | 8.525625 | 6.582504 | 0.017718 | 0.053561 | Healthy Control | Intestine |
| Eipr1 | -4.56E-01 | 5.820721 | 0.424891 | 0.520639 | 0.755917 | Healthy Control | Intestine |
| Elac2 | -1.00E+00 | 6.674161 | 2.379933 | 0.136123 | 0.276107 | Healthy Control | Intestine |
| Elavl1 | -1.51E+00 | 9.517472 | 60.23923 | 1.04E-07 | 1.64E-06 | Healthy Control | Intestine |
| Elavl2 | -3.51E+00 | 2.124147 | 0.245199 | 0.625616 | 0.841585 | Healthy Control | Intestine |
| Elavl3 | -4.59E+00 | 2.18221 | 0.468932 | 0.499924 | 0.733046 | Healthy Control | Intestine |
| Elavl4 | -5.01E+00 | 2.285186 | 0.517321 | 0.479134 | 0.713187 | Healthy Control | Intestine |
| Elf2 | -5.42E-01 | 8.490972 | 6.35302 | 0.019551 | 0.058188 | Healthy Control | Intestine |
| Elf3 | -2.79E+00 | 10.19631 | 92.74254 | 2.08E-09 | 6.16E-08 | Healthy Control | Intestine |
| Elk1 | 1.88E+00 | 5.734136 | 24.69712 | 5.63E-05 | 0.000373 | Infection | Intestine |
| Elmo1 | -7.13E-01 | 8.270557 | 7.193569 | 0.013668 | 0.042791 | Healthy Control | Intestine |
| Elmo3 | -3.62E+00 | 7.929079 | 27.61879 | 2.16E-05 | 0.000163 | Healthy Control | Intestine |
| Eln | 1.94E+00 | 8.607856 | 20.88127 | 0.00038 | 0.002003 | Infection | Intestine |
| Elob | -8.07E-01 | 7.463698 | 3.94242 | 0.059464 | 0.144083 | Healthy Control | Intestine |
| Eloc | -1.80E+00 | 6.054266 | 4.741644 | 0.039644 | 0.103262 | Healthy Control | Intestine |
| Elovl1 | -2.77E+00 | 8.103262 | 40.00056 | 1.53E-06 | 1.59E-05 | Healthy Control | Intestine |
| Elovl4 | -4.09E+00 | 2.002565 | 0.495955 | 0.487616 | 0.721551 | Healthy Control | Intestine |
| Elp2 | 1.16E-01 | 6.83076 | 0.068073 | 0.796579 | 0.943769 | Infection | Intestine |
| Emc1 | -2.94E-01 | 8.615839 | 1.576708 | 0.222549 | 0.404384 | Healthy Control | Intestine |
| Emc10 | 3.52E-03 | 8.201322 | 0.000156 | 0.990147 | 1 | Infection | Intestine |
| Emc2 | -1.20E+00 | 7.280751 | 6.781196 | 0.015767 | 0.048306 | Healthy Control | Intestine |
| Emc3 | -2.02E+00 | 8.314862 | 48.41924 | 4.52E-07 | 5.65E-06 | Healthy Control | Intestine |
| Emc4 | -1.53E+00 | 7.586477 | 13.09791 | 0.001426 | 0.006201 | Healthy Control | Intestine |
| Emc6 | -1.71E+00 | 8.894434 | 38.82841 | 2.80E-06 | 2.67E-05 | Healthy Control | Intestine |
| Emd | -6.62E-01 | 6.982948 | 2.203999 | 0.151244 | 0.300096 | Healthy Control | Intestine |
| Eme1 | -6.00E+00 | 3.678524 | 1.338031 | 0.261142 | 0.45757 | Healthy Control | Intestine |

| | | | | | | | |
|---|---|---|---|---|---|---|---|
| Emilin1 | 1.15E+00 | 7.537838 | 9.367677 | 0.005773 | 0.0207 | Infection | Intestine |
| Emsy | -3.19E-02 | 7.236005 | 0.009539 | 0.92308 | 0.969735 | Healthy Control | Intestine |
| Emx1 | -6.41E-01 | 1.272368 | 0.013926 | 0.907296 | 0.969735 | Healthy Control | Intestine |
| Emx2 | 7.06E+00 | 1.277946 | 14.29634 | 0.003136 | 0.012161 | Infection | Intestine |
| En1 | -6.42E-01 | 1.273598 | 0.012729 | 0.91135 | 0.969735 | Healthy Control | Intestine |
| En2 | -2.30E+00 | 1.522366 | 0.087581 | 0.769178 | 0.932758 | Healthy Control | Intestine |
| Enah | 4.37E+00 | 6.919649 | 470.4387 | 3.01E-16 | 1.13E-13 | Infection | Intestine |
| Eng | -1.63E+00 | 7.622962 | 2.34567 | 0.144988 | 0.29044 | Healthy Control | Intestine |
| Eno1 | -1.58E+00 | 7.491629 | 7.915429 | 0.009721 | 0.032122 | Healthy Control | Intestine |
| Eno2 | 3.15E+00 | 5.665668 | 63.12281 | 7.12E-08 | 1.20E-06 | Infection | Intestine |
| Enpp1 | 4.59E-01 | 7.02505 | 2.138792 | 0.157701 | 0.309479 | Infection | Intestine |
| Enpp6 | -6.96E+00 | 3.80573 | 3.492648 | 0.076623 | 0.176113 | Healthy Control | Intestine |
| Entpd1 | -1.65E-01 | 7.685812 | 0.129297 | 0.722615 | 0.907301 | Healthy Control | Intestine |
| Eomes | -4.05E+00 | 2.034562 | 0.284477 | 0.598478 | 0.823556 | Healthy Control | Intestine |
| Ep300 | -2.38E-01 | 8.338027 | 1.044478 | 0.317998 | 0.532407 | Healthy Control | Intestine |
| Ep400 | -3.57E-01 | 8.43109 | 2.895696 | 0.103053 | 0.222044 | Healthy Control | Intestine |
| Epas1 | 6.40E-01 | 10.24112 | 16.65476 | 0.000502 | 0.002543 | Infection | Intestine |
| Epb41 | -1.07E+00 | 8.606328 | 9.356431 | 0.005782 | 0.020719 | Healthy Control | Intestine |
| Epb41l1 | 9.70E-02 | 7.819447 | 0.077355 | 0.783536 | 0.940206 | Infection | Intestine |
| Epb41l2 | 1.38E+00 | 6.39296 | 9.910645 | 0.004701 | 0.01734 | Infection | Intestine |
| Epb41l3 | -3.11E+00 | 7.658715 | 16.76258 | 0.000396 | 0.002072 | Healthy Control | Intestine |
| Epb41l4a | -8.45E-01 | 6.33758 | 1.167283 | 0.290686 | 0.497156 | Healthy Control | Intestine |
| Epb42 | -2.97E+00 | 2.023531 | 0.183341 | 0.673246 | 0.878962 | Healthy Control | Intestine |
| Epc1 | 4.03E-01 | 6.908474 | 0.751407 | 0.395441 | 0.624669 | Infection | Intestine |
| Epc2 | -3.12E-01 | 7.359329 | 0.441866 | 0.513116 | 0.748008 | Healthy Control | Intestine |
| Epcam | -5.60E+00 | 13.65796 | 1769.728 | 2.35E-22 | 5.89E-19 | Healthy Control | Intestine |
| Epha1 | -1.53E+00 | 6.484391 | 5.524175 | 0.027231 | 0.076284 | Healthy Control | Intestine |
| Epha10 | -2.55E+00 | 1.519569 | 0.141896 | 0.709227 | 0.897243 | Healthy Control | Intestine |
| Epha2 | -1.49E-01 | 5.031844 | 0.022646 | 0.881612 | 0.96867 | Healthy Control | Intestine |
| Epha3 | 3.61E+00 | 2.004936 | 4.45066 | 0.04306 | 0.110827 | Infection | Intestine |
| Epha4 | -9.02E+00 | 6.364284 | 12.92209 | 0.001833 | 0.007676 | Healthy Control | Intestine |
| Epha5 | -4.48E+00 | 2.413451 | 0.552232 | 0.465229 | 0.700854 | Healthy Control | Intestine |
| Epha7 | 2.44E+00 | 3.521108 | 7.107179 | 0.013563 | 0.042542 | Infection | Intestine |
| Ephb1 | 3.02E+00 | 1.737887 | 2.416735 | 0.129048 | 0.264218 | Infection | Intestine |
| Ephb2 | -2.04E+00 | 7.955031 | 12.2071 | 0.001924 | 0.008 | Healthy Control | Intestine |
| Ephb3 | -7.78E-01 | 7.719441 | 4.049165 | 0.056239 | 0.137804 | Healthy Control | Intestine |
| Ephb4 | -4.94E+00 | 9.127062 | 108.1377 | 2.39E-10 | 1.01E-08 | Healthy Control | Intestine |
| Ephx2 | -2.61E+00 | 6.042899 | 4.862529 | 0.03774 | 0.099069 | Healthy Control | Intestine |
| Epm2a | -5.98E-01 | 4.150172 | 0.147564 | 0.704429 | 0.896371 | Healthy Control | Intestine |
| Epn3 | -1.10E+01 | 8.021324 | 46.39558 | 1.34E-06 | 1.43E-05 | Healthy Control | Intestine |
| Epo | -6.42E-01 | 1.272459 | 0.013821 | 0.907644 | 0.969735 | Healthy Control | Intestine |
| Epor | -1.85E-01 | 3.935367 | 0.017345 | 0.896382 | 0.969735 | Healthy Control | Intestine |
| Eps15 | 1.47E+00 | 9.166858 | 95.07813 | 2.08E-09 | 6.16E-08 | Infection | Intestine |
| Eps15l1 | 8.02E-01 | 7.437609 | 8.214408 | 0.00903 | 0.030137 | Infection | Intestine |
| Epx | -9.71E-16 | 1.180616 | 0 | 1 | 1 | Healthy Control | Intestine |
| Erap1 | -7.98E-01 | 7.046005 | 3.383333 | 0.078798 | 0.180285 | Healthy Control | Intestine |
| Erbb2 | -3.64E+00 | 8.6003 | 46.18163 | 4.55E-07 | 5.68E-06 | Healthy Control | Intestine |
| Erbb3 | -3.78E+00 | 9.083405 | 85.6958 | 2.05E-09 | 6.16E-08 | Healthy Control | Intestine |
| Erbb4 | -1.37E+00 | 1.228088 | 0.037791 | 0.848672 | 0.954066 | Healthy Control | Intestine |
| Erbin | -1.60E+00 | 10.33641 | 54.38854 | 2.36E-07 | 3.22E-06 | Healthy Control | Intestine |

| | | | | | | | |
|---|---|---|---|---|---|---|---|
| Erc2 | -3.00E+00 | 1.598068 | 0.171844 | 0.681057 | 0.882694 | Healthy Control | Intestine |
| Ercc1 | 1.68E+00 | 5.624612 | 5.803839 | 0.024708 | 0.070356 | Infection | Intestine |
| Ercc2 | -1.03E+00 | 5.933916 | 1.493138 | 0.233565 | 0.41983 | Healthy Control | Intestine |
| Ercc6 | -2.49E-02 | 6.706285 | 0.001874 | 0.965861 | 0.988784 | Healthy Control | Intestine |
| Ercc6l | 1.14E+00 | 4.38331 | 1.707283 | 0.203735 | 0.377152 | Infection | Intestine |
| Ereg | 1.56E+00 | 6.155794 | 8.402864 | 0.008391 | 0.028382 | Infection | Intestine |
| Erg | 2.34E+00 | 4.189185 | 7.262949 | 0.012979 | 0.041045 | Infection | Intestine |
| Erg28 | -9.02E-01 | 7.838249 | 7.228773 | 0.013283 | 0.041821 | Healthy Control | Intestine |
| Ern1 | -7.30E-01 | 8.195491 | 5.966631 | 0.023063 | 0.066698 | Healthy Control | Intestine |
| Ern2 | -5.06E+00 | 9.027344 | 75.86518 | 9.15E-09 | 2.06E-07 | Healthy Control | Intestine |
| Ero1l | 4.70E-01 | 8.02008 | 3.393018 | 0.079133 | 0.180803 | Infection | Intestine |
| Ero1lb | -5.10E-02 | 6.098417 | 0.004977 | 0.944359 | 0.974332 | Healthy Control | Intestine |
| Erp44 | -1.87E+00 | 8.988786 | 47.21571 | 6.58E-07 | 7.81E-06 | Healthy Control | Intestine |
| Esam | -7.04E-01 | 6.183352 | 0.315185 | 0.5808 | 0.807672 | Healthy Control | Intestine |
| Esd | -1.56E+00 | 8.103699 | 14.75558 | 0.000845 | 0.003988 | Healthy Control | Intestine |
| Espl1 | 5.94E-02 | 5.884329 | 0.005318 | 0.942492 | 0.972606 | Infection | Intestine |
| Espn | -7.70E+00 | 4.928093 | 4.083915 | 0.057029 | 0.139126 | Healthy Control | Intestine |
| Esr1 | 4.68E-01 | 5.555257 | 0.442724 | 0.512335 | 0.747523 | Infection | Intestine |
| Esr2 | -6.83E+00 | 4.640691 | 1.9309 | 0.180082 | 0.342902 | Healthy Control | Intestine |
| Esrra | -4.82E-01 | 7.691595 | 2.255961 | 0.147207 | 0.293944 | Healthy Control | Intestine |
| Esrrb | -3.12E+00 | 1.52374 | 0.191978 | 0.664544 | 0.871778 | Healthy Control | Intestine |
| Esrrg | -6.57E+00 | 4.022341 | 2.454633 | 0.131014 | 0.267151 | Healthy Control | Intestine |
| Ess2 | -7.17E-01 | 6.604309 | 2.269852 | 0.145292 | 0.290816 | Healthy Control | Intestine |
| Esyt2 | -2.58E-01 | 8.777393 | 1.050811 | 0.316577 | 0.530739 | Healthy Control | Intestine |
| Esyt3 | -4.40E+00 | 2.183934 | 0.381686 | 0.542821 | 0.775095 | Healthy Control | Intestine |
| Etfbkmt | -3.73E-01 | 7.18795 | 0.732525 | 0.401165 | 0.631322 | Healthy Control | Intestine |
| Etnk1 | -2.21E-01 | 8.004249 | 0.670449 | 0.421675 | 0.653734 | Healthy Control | Intestine |
| Etnppl | -6.48E+00 | 3.436503 | 1.712395 | 0.203795 | 0.377152 | Healthy Control | Intestine |
| Ets1 | -1.17E+00 | 6.927942 | 3.376506 | 0.078732 | 0.180265 | Healthy Control | Intestine |
| Ets2 | -2.05E+00 | 10.2688 | 95.69837 | 1.98E-09 | 5.98E-08 | Healthy Control | Intestine |
| Etv1 | 8.44E-01 | 4.470976 | 0.673569 | 0.419797 | 0.651756 | Infection | Intestine |
| Etv2 | -1.83E+00 | 1.363495 | 0.051349 | 0.822539 | 0.953596 | Healthy Control | Intestine |
| Etv3 | -1.20E+00 | 8.169649 | 13.20801 | 0.001423 | 0.006196 | Healthy Control | Intestine |
| Etv4 | -2.25E-01 | 4.604441 | 0.031381 | 0.860958 | 0.95713 | Healthy Control | Intestine |
| Etv5 | -1.87E+00 | 8.973492 | 4.477692 | 0.053254 | 0.131812 | Healthy Control | Intestine |
| Etv6 | -9.24E-01 | 8.104168 | 9.367899 | 0.005706 | 0.020476 | Healthy Control | Intestine |
| Eva1a | -6.23E-02 | 4.786212 | 0.002561 | 0.960059 | 0.985251 | Healthy Control | Intestine |
| Evl | -1.34E+00 | 6.632227 | 2.62512 | 0.118066 | 0.246677 | Healthy Control | Intestine |
| Evpl | -2.94E+00 | 6.457165 | 9.435268 | 0.005446 | 0.019663 | Healthy Control | Intestine |
| Evx1 | -5.27E+00 | 2.878764 | 0.86811 | 0.362056 | 0.587125 | Healthy Control | Intestine |
| Evx2 | -6.74E+00 | 4.830493 | 0.642261 | 0.432177 | 0.664663 | Healthy Control | Intestine |
| Ewsr1 | -1.69E+00 | 8.301423 | 31.69638 | 1.04E-05 | 8.53E-05 | Healthy Control | Intestine |
| Exoc3l2 | -4.41E-01 | 3.821967 | 0.028674 | 0.867242 | 0.96049 | Healthy Control | Intestine |
| Exoc4 | 8.18E-01 | 8.210013 | 14.12427 | 0.0011 | 0.005011 | Infection | Intestine |
| Exosc9 | -2.25E-01 | 7.071909 | 0.362699 | 0.55311 | 0.782305 | Healthy Control | Intestine |
| Exph5 | -3.50E+00 | 9.046529 | 40.10103 | 1.56E-06 | 1.62E-05 | Healthy Control | Intestine |
| Ext1 | 6.53E-01 | 7.81577 | 7.114268 | 0.014149 | 0.04402 | Infection | Intestine |
| Ext2 | 5.12E-01 | 8.484811 | 5.567399 | 0.027685 | 0.077277 | Infection | Intestine |
| Extl1 | -3.10E+00 | 1.96771 | 0.179918 | 0.675427 | 0.879372 | Healthy Control | Intestine |
| Eya1 | 3.83E+00 | 1.768964 | 4.670735 | 0.037475 | 0.098477 | Infection | Intestine |

| | | | | | | | |
|---|---|---|---|---|---|---|---|
| Eya2 | -5.43E+00 | 2.815181 | 0.694472 | 0.414139 | 0.64681 | Healthy Control | Intestine |
| Eya3 | -3.00E+00 | 7.261508 | 20.12022 | 0.000155 | 0.000901 | Healthy Control | Intestine |
| Eya4 | 1.74E+00 | 2.440493 | 0.912688 | 0.34871 | 0.572154 | Infection | Intestine |
| Ezh1 | 3.83E-01 | 8.054623 | 3.793148 | 0.064473 | 0.154205 | Infection | Intestine |
| Ezh2 | 5.17E-01 | 7.444982 | 4.232093 | 0.051844 | 0.129055 | Infection | Intestine |
| Ezr | -1.88E+00 | 10.13889 | 90.06466 | 3.39E-09 | 8.98E-08 | Healthy Control | Intestine |
| F11r | -2.15E+00 | 9.837879 | 77.41726 | 1.24E-08 | 2.65E-07 | Healthy Control | Intestine |
| F13a1 | 5.56E-01 | 7.064307 | 0.622586 | 0.440415 | 0.673346 | Infection | Intestine |
| F2r | 1.97E+00 | 7.361508 | 62.04536 | 8.14E-08 | 1.34E-06 | Infection | Intestine |
| F2rl1 | -6.41E-01 | 8.051013 | 4.439533 | 0.046756 | 0.118272 | Healthy Control | Intestine |
| F2rl2 | -3.20E+00 | 1.702166 | 0.150627 | 0.700481 | 0.89357 | Healthy Control | Intestine |
| F2rl3 | 6.63E-01 | 3.637868 | 0.249758 | 0.621927 | 0.839021 | Infection | Intestine |
| F3 | 5.28E-02 | 6.486635 | 0.008672 | 0.926595 | 0.969735 | Infection | Intestine |
| F8 | 3.01E+00 | 1.975209 | 1.824792 | 0.188371 | 0.35447 | Infection | Intestine |
| F9 | -3.64E+00 | 1.526477 | 0.285227 | 0.600212 | 0.824505 | Healthy Control | Intestine |
| Faah | -9.41E-01 | 6.847908 | 3.813654 | 0.062913 | 0.151145 | Healthy Control | Intestine |
| Faap100 | -8.48E-01 | 6.464739 | 1.786576 | 0.193932 | 0.362789 | Healthy Control | Intestine |
| Fabp12 | -1.83E+00 | 1.361645 | 0.058008 | 0.811599 | 0.949933 | Healthy Control | Intestine |
| Fabp3 | -3.83E+00 | 2.176871 | 0.36093 | 0.553619 | 0.782305 | Healthy Control | Intestine |
| Fabp7 | -4.35E+00 | 2.530134 | 0.436386 | 0.515646 | 0.750821 | Healthy Control | Intestine |
| Fadd | -1.23E-01 | 8.509541 | 0.304083 | 0.586949 | 0.812799 | Healthy Control | Intestine |
| Fah | 2.35E-01 | 6.49791 | 0.262864 | 0.613198 | 0.833923 | Infection | Intestine |
| Fam104a | 9.00E-01 | 4.831453 | 1.631052 | 0.213877 | 0.39147 | Infection | Intestine |
| Fam107b | -2.53E+00 | 9.937149 | 79.29441 | 1.02E-08 | 2.26E-07 | Healthy Control | Intestine |
| Fam129a | -1.05E+00 | 8.772863 | 22.06013 | 0.000108 | 0.000662 | Healthy Control | Intestine |
| Fam136a | -2.13E+00 | 8.252125 | 28.46399 | 1.97E-05 | 0.000151 | Healthy Control | Intestine |
| Fam13b | -6.44E-03 | 8.371565 | 0.000638 | 0.980071 | 0.999437 | Healthy Control | Intestine |
| Fam192a | -9.85E-01 | 6.684482 | 2.16458 | 0.154325 | 0.304514 | Healthy Control | Intestine |
| Fam20a | 1.74E+00 | 4.999749 | 10.03982 | 0.004399 | 0.016347 | Infection | Intestine |
| Fam20c | 7.73E-01 | 5.131686 | 0.968426 | 0.335108 | 0.554562 | Infection | Intestine |
| Fam210a | -9.77E-01 | 7.627991 | 7.027253 | 0.014357 | 0.044612 | Healthy Control | Intestine |
| Fam210b | -1.80E+00 | 7.387314 | 13.28702 | 0.001292 | 0.00573 | Healthy Control | Intestine |
| Fam214a | -9.55E-01 | 6.527943 | 1.792755 | 0.193139 | 0.361691 | Healthy Control | Intestine |
| Fam49b | 1.56E+00 | 8.273437 | 86.01967 | 5.08E-09 | 1.27E-07 | Infection | Intestine |
| Fam83d | 4.31E-01 | 6.037081 | 0.282184 | 0.600508 | 0.824505 | Infection | Intestine |
| Fam92a | 3.98E-01 | 6.683438 | 1.028803 | 0.32143 | 0.536717 | Infection | Intestine |
| Fancc | 7.13E-01 | 5.542921 | 0.975247 | 0.333816 | 0.553156 | Infection | Intestine |
| Fanci | -5.89E+00 | 3.375657 | 1.397875 | 0.251212 | 0.444841 | Healthy Control | Intestine |
| Fancl | 8.05E-01 | 6.416023 | 3.62205 | 0.070177 | 0.164855 | Infection | Intestine |
| Fap | 2.61E+00 | 5.527416 | 24.32623 | 6.40E-05 | 0.000417 | Infection | Intestine |
| Far2 | -2.21E-01 | 4.809253 | 0.031722 | 0.860068 | 0.956566 | Healthy Control | Intestine |
| Fas | 6.66E-01 | 5.657079 | 1.331641 | 0.260416 | 0.456777 | Infection | Intestine |
| Fasl | -4.35E+00 | 2.44741 | 0.347405 | 0.56141 | 0.789783 | Healthy Control | Intestine |
| Fasn | -3.36E+00 | 9.495091 | 103.4673 | 4.75E-10 | 1.79E-08 | Healthy Control | Intestine |
| Fat1 | -2.40E+00 | 10.54541 | 165.4674 | 1.09E-11 | 7.83E-10 | Healthy Control | Intestine |
| Fat4 | -4.53E-01 | 6.910231 | 0.780061 | 0.386462 | 0.615928 | Healthy Control | Intestine |
| Fbh1 | -3.62E-01 | 8.58421 | 2.694791 | 0.115014 | 0.24222 | Healthy Control | Intestine |
| Fbl | -1.48E+00 | 7.757507 | 11.13207 | 0.002874 | 0.011309 | Healthy Control | Intestine |
| Fbln1 | 2.10E-01 | 9.00995 | 0.174499 | 0.682554 | 0.883743 | Infection | Intestine |
| Fbln2 | 1.84E+00 | 7.484726 | 15.59501 | 0.001066 | 0.004879 | Infection | Intestine |

| | | | | | | | |
|---|---|---|---|---|---|---|---|
| Fbn1 | 2.46E+00 | 9.699344 | 44.69242 | 1.67E-05 | 0.00013 | Infection | Intestine |
| Fbxl12 | -8.95E-01 | 6.801542 | 3.275153 | 0.083135 | 0.187837 | Healthy Control | Intestine |
| Fbxl13 | 1.56E+00 | 2.638916 | 0.648096 | 0.428384 | 0.661048 | Infection | Intestine |
| Fbxl16 | -4.40E+00 | 2.296371 | 0.512843 | 0.481578 | 0.715666 | Healthy Control | Intestine |
| Fbxl17 | -2.10E+00 | 7.337742 | 21.65142 | 9.85E-05 | 0.000611 | Healthy Control | Intestine |
| Fbxl19 | -6.35E-01 | 8.420451 | 4.60036 | 0.04333 | 0.111407 | Healthy Control | Intestine |
| Fbxl20 | 2.94E-01 | 7.974958 | 1.753263 | 0.199198 | 0.370426 | Infection | Intestine |
| Fbxl3 | -1.38E+00 | 8.514678 | 20.50089 | 0.000165 | 0.000954 | Healthy Control | Intestine |
| Fbxl4 | -8.37E-01 | 6.486146 | 1.878775 | 0.183228 | 0.347561 | Healthy Control | Intestine |
| Fbxl7 | 1.44E+00 | 4.392038 | 2.516587 | 0.125888 | 0.258806 | Infection | Intestine |
| Fbxo11 | -1.11E+00 | 7.368647 | 5.766445 | 0.024748 | 0.070431 | Healthy Control | Intestine |
| Fbxo17 | 2.24E+00 | 4.321191 | 9.803 | 0.004731 | 0.017442 | Infection | Intestine |
| Fbxo2 | 1.65E+00 | 2.234106 | 0.555956 | 0.462959 | 0.698695 | Infection | Intestine |
| Fbxo21 | -2.05E+00 | 6.752657 | 8.699759 | 0.006915 | 0.024157 | Healthy Control | Intestine |
| Fbxo3 | 1.22E+00 | 8.530777 | 50.6659 | 4.10E-07 | 5.24E-06 | Infection | Intestine |
| Fbxo31 | 5.75E-01 | 7.238009 | 2.998419 | 0.097406 | 0.212006 | Infection | Intestine |
| Fbxo32 | -1.39E+00 | 8.0927 | 15.50598 | 0.000638 | 0.003142 | Healthy Control | Intestine |
| Fbxo33 | -1.35E+00 | 7.028241 | 4.806882 | 0.03828 | 0.100278 | Healthy Control | Intestine |
| Fbxo4 | 2.73E-01 | 6.286344 | 0.24738 | 0.623674 | 0.840408 | Infection | Intestine |
| Fbxo41 | 3.43E+00 | 1.562926 | 2.588027 | 0.116198 | 0.244304 | Infection | Intestine |
| Fbxo5 | 1.44E-01 | 5.894127 | 0.048871 | 0.826984 | 0.953685 | Infection | Intestine |
| Fbxw11 | -7.14E-01 | 7.870968 | 4.459086 | 0.046164 | 0.117188 | Healthy Control | Intestine |
| Fbxw7 | 1.18E+00 | 7.170801 | 14.30089 | 0.001041 | 0.004793 | Infection | Intestine |
| Fcgr1 | 8.51E-01 | 4.943976 | 0.679436 | 0.417975 | 0.650212 | Infection | Intestine |
| Fcgr2b | 2.69E+00 | 7.803058 | 125.2055 | 1.67E-10 | 7.68E-09 | Infection | Intestine |
| Fcgr3 | 1.74E+00 | 6.216071 | 10.36623 | 0.003971 | 0.014992 | Infection | Intestine |
| Fcgr4 | -5.16E-01 | 5.680608 | 0.168572 | 0.684933 | 0.885006 | Healthy Control | Intestine |
| Fcgrt | 9.86E-01 | 8.497903 | 10.25603 | 0.004391 | 0.016332 | Infection | Intestine |
| Fcmr | 3.38E-01 | 3.211808 | 0.009195 | 0.924573 | 0.969735 | Infection | Intestine |
| Fcrls | -1.62E+00 | 6.103408 | 2.539827 | 0.123854 | 0.255781 | Healthy Control | Intestine |
| Fdx1 | -2.70E+00 | 8.590764 | 43.3589 | 9.31E-07 | 1.06E-05 | Healthy Control | Intestine |
| Fen1 | -1.29E+00 | 5.676412 | 1.601099 | 0.2177 | 0.39687 | Healthy Control | Intestine |
| Fermt1 | -5.10E+00 | 9.294136 | 91.24996 | 1.29E-09 | 4.07E-08 | Healthy Control | Intestine |
| Fermt3 | 1.95E+00 | 5.784062 | 18.88051 | 0.000263 | 0.001453 | Infection | Intestine |
| Fezf1 | -2.34E+00 | 1.522871 | 0.096842 | 0.757808 | 0.928591 | Healthy Control | Intestine |
| Fezf2 | -9.71E-16 | 1.180616 | 0 | 1 | 1 | Healthy Control | Intestine |
| Ffar1 | -4.65E+00 | 2.503386 | 0.376771 | 0.546026 | 0.776921 | Healthy Control | Intestine |
| Ffar2 | -9.65E-01 | 8.365964 | 2.09073 | 0.168047 | 0.324428 | Healthy Control | Intestine |
| Ffar3 | 3.97E+00 | 1.486811 | 2.642532 | 0.114316 | 0.241056 | Infection | Intestine |
| Ffar4 | -3.91E+00 | 6.734483 | 11.58289 | 0.002637 | 0.010493 | Healthy Control | Intestine |
| Fga | -4.94E-01 | 1.22813 | 0.009389 | 0.924171 | 0.969735 | Healthy Control | Intestine |
| Fgb | -7.56E-01 | 1.273429 | 0.015546 | 0.902199 | 0.969735 | Healthy Control | Intestine |
| Fgf1 | 1.31E+00 | 7.861686 | 20.95963 | 0.00015 | 0.000878 | Infection | Intestine |
| Fgf10 | 1.36E+00 | 4.558134 | 3.588656 | 0.070272 | 0.164908 | Infection | Intestine |
| Fgf12 | -2.06E-01 | 4.716202 | 0.026378 | 0.872289 | 0.962672 | Healthy Control | Intestine |
| Fgf15 | -4.94E-01 | 1.22813 | 0.009389 | 0.924171 | 0.969735 | Healthy Control | Intestine |
| Fgf16 | 3.35E+00 | 1.733375 | 1.694757 | 0.202032 | 0.374444 | Infection | Intestine |
| Fgf17 | -2.90E+00 | 1.66856 | 0.1606 | 0.691249 | 0.889526 | Healthy Control | Intestine |
| Fgf18 | 1.77E+00 | 2.649477 | 0.874263 | 0.358446 | 0.582912 | Infection | Intestine |
| Fgf2 | 1.50E+00 | 4.427645 | 3.597965 | 0.070284 | 0.164908 | Infection | Intestine |

| | | | | | | | |
|---|---|---|---|---|---|---|---|
| Fgf21 | -1.78E+00 | 1.360119 | 0.039512 | 0.843991 | 0.954066 | Healthy Control | Intestine |
| Fgf22 | 5.33E+00 | 1.275385 | 4.744831 | 0.046136 | 0.117176 | Infection | Intestine |
| Fgf23 | 6.70E+00 | 1.321453 | 12.67839 | 0.003004 | 0.0117 | Infection | Intestine |
| Fgf3 | -2.72E+00 | 1.405616 | 0.109024 | 0.744491 | 0.923113 | Healthy Control | Intestine |
| Fgf4 | -1.37E+00 | 1.228088 | 0.037791 | 0.848672 | 0.954066 | Healthy Control | Intestine |
| Fgf5 | 5.06E+00 | 1.318835 | 4.52696 | 0.046192 | 0.1172 | Infection | Intestine |
| Fgf7 | -6.32E-02 | 6.311969 | 0.009356 | 0.923782 | 0.969735 | Healthy Control | Intestine |
| Fgf8 | -1.52E+00 | 1.274131 | 0.039118 | 0.845524 | 0.954066 | Healthy Control | Intestine |
| Fgf9 | -7.11E+00 | 4.383308 | 2.202368 | 0.153563 | 0.303368 | Healthy Control | Intestine |
| Fgfbp3 | 7.11E-01 | 4.236814 | 0.494527 | 0.488779 | 0.722204 | Infection | Intestine |
| Fgfr1 | 1.96E+00 | 7.221869 | 33.13546 | 8.94E-06 | 7.47E-05 | Infection | Intestine |
| Fgfr1op | -2.09E-01 | 6.350357 | 0.148846 | 0.703158 | 0.895221 | Healthy Control | Intestine |
| Fgfr2 | -7.21E-01 | 8.819995 | 3.554 | 0.073762 | 0.171426 | Healthy Control | Intestine |
| Fgfr3 | -3.85E+00 | 8.554676 | 40.89883 | 1.44E-06 | 1.51E-05 | Healthy Control | Intestine |
| Fgfr4 | -1.56E+00 | 7.311307 | 5.351531 | 0.029698 | 0.081462 | Healthy Control | Intestine |
| Fgg | -2.08E+00 | 1.444722 | 0.083389 | 0.774658 | 0.934669 | Healthy Control | Intestine |
| Fgl1 | -1.32E+00 | 1.362994 | 0.044329 | 0.835473 | 0.954066 | Healthy Control | Intestine |
| Fgr | 3.01E+00 | 4.564762 | 21.88821 | 0.000113 | 0.00069 | Infection | Intestine |
| Fh1 | -3.77E+00 | 9.099135 | 155.5618 | 5.27E-12 | 4.32E-10 | Healthy Control | Intestine |
| Fhl2 | 1.14E+00 | 6.482855 | 8.408794 | 0.00834 | 0.028249 | Infection | Intestine |
| Fhod1 | 4.21E-01 | 7.557417 | 2.127359 | 0.158919 | 0.311369 | Infection | Intestine |
| Fibcd1 | -3.66E+00 | 1.705875 | 0.205633 | 0.65353 | 0.863896 | Healthy Control | Intestine |
| Fign | 3.11E+00 | 3.387102 | 8.849347 | 0.006677 | 0.023433 | Infection | Intestine |
| Fkbp5 | -1.46E-01 | 8.235866 | 0.181662 | 0.674134 | 0.879138 | Healthy Control | Intestine |
| Fli1 | 1.69E+00 | 6.509897 | 14.09926 | 0.00111 | 0.005047 | Infection | Intestine |
| Flna | 9.18E-01 | 10.05411 | 10.01092 | 0.006307 | 0.022361 | Infection | Intestine |
| Flnb | -1.12E+00 | 10.3605 | 40.53669 | 2.18E-06 | 2.14E-05 | Healthy Control | Intestine |
| Flot1 | -1.94E-01 | 7.822231 | 0.40373 | 0.531744 | 0.765576 | Healthy Control | Intestine |
| Flot2 | 5.95E-01 | 7.839953 | 5.667719 | 0.026452 | 0.074394 | Infection | Intestine |
| Flt1 | 3.36E+00 | 6.256988 | 17.33991 | 0.001061 | 0.004859 | Infection | Intestine |
| Flt3 | -6.83E+00 | 4.053958 | 1.143579 | 0.297804 | 0.506561 | Healthy Control | Intestine |
| Flt3l | -3.36E-01 | 4.641349 | 0.100559 | 0.753879 | 0.926115 | Healthy Control | Intestine |
| Flt4 | 8.69E-01 | 6.56884 | 0.276587 | 0.608309 | 0.831483 | Infection | Intestine |
| Fmc1 | -4.02E+00 | 8.942064 | 71.76583 | 1.05E-08 | 2.31E-07 | Healthy Control | Intestine |
| Fmo4 | -4.66E+00 | 2.94063 | 0.414656 | 0.526672 | 0.760461 | Healthy Control | Intestine |
| Fmod | -4.35E+00 | 2.48121 | 0.52196 | 0.477556 | 0.71235 | Healthy Control | Intestine |
| Fmr1 | 5.94E-01 | 8.630665 | 12.38835 | 0.001952 | 0.008103 | Infection | Intestine |
| Fn1 | 2.92E+00 | 10.49051 | 53.06285 | 8.71E-06 | 7.34E-05 | Infection | Intestine |
| Fndc5 | -6.98E+00 | 4.384968 | 2.63731 | 0.120221 | 0.249928 | Healthy Control | Intestine |
| Folr1 | -4.27E+00 | 1.896905 | 0.487125 | 0.493719 | 0.726929 | Healthy Control | Intestine |
| Folr2 | 8.93E-01 | 5.115433 | 0.971601 | 0.334237 | 0.553487 | Infection | Intestine |
| Fopnl | -1.83E+00 | 7.103403 | 8.437331 | 0.007827 | 0.026729 | Healthy Control | Intestine |
| Fosl1 | 3.18E+00 | 1.701957 | 1.897585 | 0.17717 | 0.338575 | Infection | Intestine |
| Fosl2 | 8.43E-01 | 8.492092 | 6.909462 | 0.015942 | 0.048781 | Infection | Intestine |
| Foxa1 | -3.54E+00 | 8.204597 | 38.88599 | 1.77E-06 | 1.79E-05 | Healthy Control | Intestine |
| Foxa2 | -1.10E+01 | 8.218378 | 53.33851 | 4.90E-07 | 6.10E-06 | Healthy Control | Intestine |
| Foxb1 | -1.83E+00 | 1.361176 | 0.059181 | 0.809743 | 0.94887 | Healthy Control | Intestine |
| Foxc1 | -5.10E+00 | 2.544325 | 0.438292 | 0.51528 | 0.750506 | Healthy Control | Intestine |
| Foxc2 | 1.01E+00 | 2.520886 | 0.22062 | 0.642745 | 0.856423 | Infection | Intestine |
| Foxd1 | -7.56E-01 | 1.27456 | 0.014684 | 0.904934 | 0.969735 | Healthy Control | Intestine |

| | | | | | | | |
|---|---|---|---|---|---|---|---|
| Foxd2 | -3.50E+00 | 8.97256 | 81.83284 | 3.32E-09 | 8.88E-08 | Healthy Control | Intestine |
| Foxd3 | -3.86E+00 | 2.250629 | 0.356255 | 0.556238 | 0.784375 | Healthy Control | Intestine |
| Foxf1 | 6.45E-01 | 5.957589 | 0.986295 | 0.33126 | 0.549466 | Infection | Intestine |
| Foxg1 | -1.68E+00 | 1.4827 | 0.049243 | 0.825924 | 0.953685 | Healthy Control | Intestine |
| Foxh1 | -2.03E+00 | 1.404274 | 0.068065 | 0.796579 | 0.943769 | Healthy Control | Intestine |
| Foxi1 | -8.79E-01 | 1.316791 | 0.017191 | 0.896835 | 0.969735 | Healthy Control | Intestine |
| Foxi2 | -6.42E-01 | 1.272688 | 0.013552 | 0.908542 | 0.969735 | Healthy Control | Intestine |
| Foxj1 | -2.35E+00 | 1.591174 | 0.112207 | 0.740643 | 0.921009 | Healthy Control | Intestine |
| Foxj2 | -2.83E+00 | 7.154227 | 20.17118 | 0.000156 | 0.000906 | Healthy Control | Intestine |
| Foxj3 | 5.77E-01 | 6.335081 | 2.217301 | 0.150514 | 0.299117 | Infection | Intestine |
| Foxl2 | 4.33E+00 | 1.445307 | 2.786569 | 0.106166 | 0.227219 | Infection | Intestine |
| Foxm1 | 3.03E-01 | 5.86764 | 0.157301 | 0.695339 | 0.892073 | Infection | Intestine |
| Foxn1 | -4.91E+00 | 2.84555 | 0.856807 | 0.363954 | 0.588676 | Healthy Control | Intestine |
| Foxn4 | -1.83E+00 | 1.361963 | 0.05693 | 0.813323 | 0.950395 | Healthy Control | Intestine |
| Foxo1 | -1.44E+00 | 7.69248 | 13.79246 | 0.001139 | 0.005141 | Healthy Control | Intestine |
| Foxo3 | -2.94E+00 | 7.850303 | 31.80298 | 8.64E-06 | 7.31E-05 | Healthy Control | Intestine |
| Foxo4 | -2.20E+00 | 5.611208 | 3.388009 | 0.07874 | 0.180265 | Healthy Control | Intestine |
| Foxp1 | -1.61E+00 | 10.11082 | 108.2386 | 6.41E-10 | 2.24E-08 | Healthy Control | Intestine |
| Foxp2 | -5.41E-01 | 5.092529 | 0.19025 | 0.666548 | 0.873264 | Healthy Control | Intestine |
| Foxp3 | 3.31E+00 | 1.928033 | 3.260172 | 0.080393 | 0.183432 | Infection | Intestine |
| Foxp4 | -8.90E-01 | 9.273819 | 21.78757 | 0.000121 | 0.000731 | Healthy Control | Intestine |
| Fpr1 | -2.94E+00 | 1.525747 | 0.134294 | 0.717335 | 0.903164 | Healthy Control | Intestine |
| Fpr2 | -6.28E+00 | 3.572556 | 1.055589 | 0.316708 | 0.530779 | Healthy Control | Intestine |
| Fras1 | -4.97E+00 | 2.466698 | 0.615428 | 0.44168 | 0.674718 | Healthy Control | Intestine |
| Frk | -2.29E+00 | 8.213868 | 28.49088 | 2.03E-05 | 0.000154 | Healthy Control | Intestine |
| Frmd7 | -2.97E+00 | 1.917938 | 0.120065 | 0.732309 | 0.914884 | Healthy Control | Intestine |
| Frmpd1 | -3.39E+00 | 1.564349 | 0.168432 | 0.68535 | 0.885046 | Healthy Control | Intestine |
| Frrs1 | 9.87E-01 | 6.26722 | 3.139618 | 0.089934 | 0.199383 | Infection | Intestine |
| Frrs1l | -4.15E+00 | 2.199232 | 0.417455 | 0.524178 | 0.758326 | Healthy Control | Intestine |
| Frs2 | 3.62E-01 | 7.841591 | 1.51153 | 0.231993 | 0.418068 | Infection | Intestine |
| Fryl | -1.53E+00 | 8.790063 | 16.44319 | 0.000527 | 0.002646 | Healthy Control | Intestine |
| Frzb | 1.55E+00 | 4.499221 | 3.513176 | 0.073239 | 0.170529 | Infection | Intestine |
| Fscn1 | 2.50E+00 | 7.279162 | 29.32242 | 5.41E-05 | 0.00036 | Infection | Intestine |
| Fscn2 | -2.97E+00 | 1.559698 | 0.158042 | 0.693546 | 0.890686 | Healthy Control | Intestine |
| Fshb | -9.71E-16 | 1.180616 | 0 | 1 | 1 | Healthy Control | Intestine |
| Fshr | -6.42E-01 | 1.273689 | 0.012641 | 0.911654 | 0.969735 | Healthy Control | Intestine |
| Fst | 1.18E+00 | 3.491281 | 0.923893 | 0.346265 | 0.569262 | Infection | Intestine |
| Fstl1 | 6.34E-01 | 9.411215 | 2.918832 | 0.107806 | 0.229943 | Infection | Intestine |
| Fstl4 | -1.89E+00 | 4.856179 | 1.608903 | 0.218149 | 0.397545 | Healthy Control | Intestine |
| Fstl5 | -1.78E+00 | 1.361453 | 0.05816 | 0.811328 | 0.949838 | Healthy Control | Intestine |
| Fthl17a | -1.37E+00 | 1.229023 | 0.032546 | 0.859435 | 0.956553 | Healthy Control | Intestine |
| Fto | 2.18E-01 | 7.559739 | 0.661655 | 0.424766 | 0.657507 | Infection | Intestine |
| Ftsj1 | -7.78E-01 | 6.652343 | 2.163521 | 0.154528 | 0.304794 | Healthy Control | Intestine |
| Fubp1 | -2.92E-03 | 7.938543 | 0.000122 | 0.991281 | 1 | Healthy Control | Intestine |
| Fundc1 | 8.68E-02 | 7.900273 | 0.129509 | 0.722404 | 0.907264 | Infection | Intestine |
| Furin | 7.24E-03 | 9.339406 | 0.00218 | 0.963186 | 0.987044 | Infection | Intestine |
| Fus | 1.70E+00 | 8.789449 | 93.70064 | 2.37E-09 | 6.82E-08 | Infection | Intestine |
| Fut1 | -3.75E-01 | 3.769822 | 0.052089 | 0.82156 | 0.953086 | Healthy Control | Intestine |
| Fut4 | -3.78E+00 | 9.342892 | 64.82077 | 4.10E-08 | 7.52E-07 | Healthy Control | Intestine |
| Fv1 | -1.54E+00 | 5.848286 | 2.76787 | 0.109123 | 0.232059 | Healthy Control | Intestine |

| | | | | | | | |
|---|---|---|---|---|---|---|---|
| Fxn | -6.30E+00 | 3.865214 | 2.092019 | 0.163789 | 0.318667 | Healthy Control | Intestine |
| Fxr1 | 9.01E-01 | 7.290439 | 9.213901 | 0.006116 | 0.02176 | Infection | Intestine |
| Fyn | 1.98E+00 | 7.094244 | 28.25545 | 2.54E-05 | 0.000187 | Infection | Intestine |
| Fzd1 | 2.34E-01 | 8.789737 | 0.549014 | 0.46694 | 0.702797 | Infection | Intestine |
| Fzd10 | -6.14E+00 | 3.338645 | 1.516016 | 0.232 | 0.418068 | Healthy Control | Intestine |
| Fzd2 | 3.48E+00 | 5.08447 | 47.70919 | 6.54E-07 | 7.80E-06 | Infection | Intestine |
| Fzd3 | -2.07E-01 | 3.701294 | 0.016689 | 0.898359 | 0.969735 | Healthy Control | Intestine |
| Fzd4 | 1.38E-01 | 5.532133 | 0.021723 | 0.884073 | 0.96867 | Infection | Intestine |
| Fzd5 | -2.82E+00 | 8.666037 | 52.23752 | 2.24E-07 | 3.10E-06 | Healthy Control | Intestine |
| Fzd6 | -2.50E+00 | 5.972906 | 4.593759 | 0.043022 | 0.110787 | Healthy Control | Intestine |
| Fzd7 | -1.52E+00 | 6.01456 | 2.566169 | 0.122004 | 0.252899 | Healthy Control | Intestine |
| Fzd8 | -1.82E+00 | 6.185736 | 4.785748 | 0.038497 | 0.100756 | Healthy Control | Intestine |
| G3bp1 | -2.06E+00 | 8.707494 | 50.07568 | 3.68E-07 | 4.78E-06 | Healthy Control | Intestine |
| G6pc | -1.71E+00 | 1.317989 | 0.052791 | 0.820546 | 0.953035 | Healthy Control | Intestine |
| G6pc2 | 4.28E+00 | 1.363129 | 3.642215 | 0.069511 | 0.163539 | Infection | Intestine |
| G6pdx | -2.17E+00 | 5.768422 | 3.824046 | 0.062922 | 0.151145 | Healthy Control | Intestine |
| Gaa | -2.82E+00 | 7.14529 | 17.91644 | 0.000289 | 0.001574 | Healthy Control | Intestine |
| Gab1 | -7.93E-01 | 7.466271 | 3.007321 | 0.096521 | 0.210445 | Healthy Control | Intestine |
| Gabbr1 | 1.97E+00 | 7.591029 | 81.9524 | 7.75E-09 | 1.82E-07 | Infection | Intestine |
| Gabbr2 | -1.40E+00 | 1.363136 | 0.035701 | 0.852687 | 0.956553 | Healthy Control | Intestine |
| Gabpb1 | -2.00E+00 | 7.289135 | 15.73568 | 0.000564 | 0.002813 | Healthy Control | Intestine |
| Gabra1 | 4.76E+00 | 1.526594 | 8.319149 | 0.006623 | 0.023289 | Infection | Intestine |
| Gabra2 | -8.61E-01 | 1.274141 | 0.016052 | 0.900735 | 0.969735 | Healthy Control | Intestine |
| Gabra5 | -2.20E+00 | 1.319993 | 0.087356 | 0.770732 | 0.93308 | Healthy Control | Intestine |
| Gabra6 | -9.71E-16 | 1.180616 | 0 | 1 | 1 | Healthy Control | Intestine |
| Gabrb2 | 2.80E+00 | 2.104046 | 1.922728 | 0.177973 | 0.339791 | Infection | Intestine |
| Gabrb3 | 2.11E+00 | 1.949789 | 0.968758 | 0.333211 | 0.552336 | Infection | Intestine |
| Gabrd | -2.99E+00 | 1.446909 | 0.172397 | 0.681777 | 0.883047 | Healthy Control | Intestine |
| Gabrg1 | -3.57E-01 | 1.228299 | 0.00618 | 0.938438 | 0.969735 | Healthy Control | Intestine |
| Gabrg2 | -1.37E+00 | 1.228863 | 0.033199 | 0.858049 | 0.956553 | Healthy Control | Intestine |
| Gabrg3 | -3.03E+00 | 1.483947 | 0.180443 | 0.674349 | 0.879138 | Healthy Control | Intestine |
| Gabrp | -6.20E+00 | 3.860414 | 0.918608 | 0.349427 | 0.573142 | Healthy Control | Intestine |
| Gabrq | 4.08E+00 | 1.444335 | 4.366624 | 0.045214 | 0.115481 | Infection | Intestine |
| Gabrr2 | -4.25E+00 | 2.158449 | 0.474313 | 0.497951 | 0.730795 | Healthy Control | Intestine |
| Gad1 | -2.99E+00 | 1.446003 | 0.151164 | 0.700944 | 0.89357 | Healthy Control | Intestine |
| Gad2 | -6.33E+00 | 3.809653 | 1.674147 | 0.210564 | 0.386677 | Healthy Control | Intestine |
| Gadd45a | -6.29E-01 | 5.737114 | 0.46827 | 0.500419 | 0.733557 | Healthy Control | Intestine |
| Gak | -6.19E-01 | 9.065669 | 11.75414 | 0.002429 | 0.009767 | Healthy Control | Intestine |
| Gal | 4.73E+00 | 1.702318 | 6.87501 | 0.013155 | 0.041497 | Infection | Intestine |
| Galc | 1.55E+00 | 5.394841 | 8.126458 | 0.009253 | 0.030737 | Infection | Intestine |
| Galnt14 | -2.35E+00 | 1.594645 | 0.08233 | 0.776701 | 0.935926 | Healthy Control | Intestine |
| Galnt15 | -1.31E+00 | 5.582216 | 0.882012 | 0.356941 | 0.581657 | Healthy Control | Intestine |
| Galnt17 | -3.83E-01 | 5.738745 | 0.082268 | 0.776641 | 0.935926 | Healthy Control | Intestine |
| Galnt18 | 1.18E+00 | 5.276165 | 2.394909 | 0.135633 | 0.275225 | Infection | Intestine |
| Galntl5 | 4.15E+00 | 1.404921 | 3.503447 | 0.072074 | 0.168205 | Infection | Intestine |
| Galp | -9.71E-16 | 1.180616 | 0 | 1 | 1 | Healthy Control | Intestine |
| Galr1 | -4.30E+00 | 2.267739 | 0.420075 | 0.52479 | 0.758839 | Healthy Control | Intestine |
| Galr2 | 3.24E+00 | 1.767233 | 2.343404 | 0.135411 | 0.274984 | Infection | Intestine |
| Gap43 | -1.12E-01 | 3.942461 | 0.002521 | 0.960542 | 0.985267 | Healthy Control | Intestine |
| Gar1 | -3.61E-01 | 6.893794 | 0.608939 | 0.443176 | 0.676796 | Healthy Control | Intestine |

| | | | | | | | |
|---|---|---|---|---|---|---|---|
| Garem1 | -7.78E-01 | 6.727156 | 1.912398 | 0.179636 | 0.342314 | Healthy Control | Intestine |
| Gars | -7.20E-01 | 9.300331 | 16.14853 | 0.000586 | 0.002911 | Healthy Control | Intestine |
| Gart | -6.16E-01 | 7.314413 | 2.476123 | 0.129449 | 0.264714 | Healthy Control | Intestine |
| Gas1 | 3.81E+00 | 5.617663 | 85.86245 | 5.17E-09 | 1.28E-07 | Infection | Intestine |
| Gas6 | -2.02E+00 | 10.07003 | 55.49607 | 1.95E-07 | 2.75E-06 | Healthy Control | Intestine |
| Gast | -3.57E-01 | 1.227417 | 0.006288 | 0.937908 | 0.969735 | Healthy Control | Intestine |
| Gata1 | 7.68E+00 | 1.684236 | 37.14414 | 1.66E-06 | 1.71E-05 | Infection | Intestine |
| Gata2 | 2.22E+00 | 4.259936 | 5.193136 | 0.032414 | 0.087192 | Infection | Intestine |
| Gata3 | 5.34E-01 | 4.701325 | 0.210902 | 0.650102 | 0.861262 | Infection | Intestine |
| Gata4 | -1.17E+00 | 1.36141 | 0.024352 | 0.877194 | 0.966168 | Healthy Control | Intestine |
| Gata5 | -2.93E+00 | 1.670685 | 0.169115 | 0.683604 | 0.88417 | Healthy Control | Intestine |
| Gata6 | -1.89E+00 | 7.485751 | 16.70369 | 0.000429 | 0.002225 | Healthy Control | Intestine |
| Gatad2b | -4.15E-01 | 8.213295 | 3.505081 | 0.074633 | 0.172924 | Healthy Control | Intestine |
| Gatd3a | -2.07E+00 | 9.00178 | 48.15507 | 5.59E-07 | 6.88E-06 | Healthy Control | Intestine |
| Gba | -6.00E-01 | 7.085043 | 2.383155 | 0.136415 | 0.276581 | Healthy Control | Intestine |
| Gba2 | 1.56E+00 | 6.957578 | 39.00457 | 2.90E-06 | 2.75E-05 | Infection | Intestine |
| Gbf1 | 6.18E-01 | 8.39871 | 8.252985 | 0.008895 | 0.029764 | Infection | Intestine |
| Gbp2 | 1.25E+00 | 6.244726 | 4.525682 | 0.044854 | 0.114796 | Infection | Intestine |
| Gbp2b | 7.34E-01 | 6.821174 | 0.710359 | 0.411724 | 0.643971 | Infection | Intestine |
| Gbp3 | -1.05E+00 | 5.870002 | 0.906971 | 0.35024 | 0.5741 | Healthy Control | Intestine |
| Gbx1 | 3.62E+00 | 1.485569 | 2.562057 | 0.1192 | 0.24857 | Infection | Intestine |
| Gbx2 | -2.88E+00 | 1.362623 | 0.145164 | 0.708506 | 0.897011 | Healthy Control | Intestine |
| Gcat | -8.59E+00 | 6.050537 | 9.698229 | 0.005512 | 0.01985 | Healthy Control | Intestine |
| Gcg | -3.02E+00 | 7.640324 | 9.58105 | 0.005836 | 0.020884 | Healthy Control | Intestine |
| Gcgr | -2.63E+00 | 1.363598 | 0.155625 | 0.697584 | 0.892753 | Healthy Control | Intestine |
| Gch1 | -1.45E+00 | 5.768045 | 1.895231 | 0.181522 | 0.344988 | Healthy Control | Intestine |
| Gck | -4.67E+00 | 2.47962 | 0.523105 | 0.476788 | 0.711649 | Healthy Control | Intestine |
| Gckr | -2.32E+00 | 1.402636 | 0.095816 | 0.75921 | 0.928591 | Healthy Control | Intestine |
| Gcm1 | -2.04E+00 | 1.4432 | 0.088959 | 0.767526 | 0.932403 | Healthy Control | Intestine |
| Gcm2 | -3.57E-01 | 1.227067 | 0.006351 | 0.937596 | 0.969735 | Healthy Control | Intestine |
| Gcnt3 | -3.15E+00 | 10.07804 | 57.09385 | 1.92E-07 | 2.73E-06 | Healthy Control | Intestine |
| Gcnt4 | 1.31E+00 | 4.127392 | 1.60056 | 0.217574 | 0.396784 | Infection | Intestine |
| Gdap1 | -3.09E+00 | 1.666599 | 0.213309 | 0.647268 | 0.859335 | Healthy Control | Intestine |
| Gdf1 | -4.29E+00 | 2.227732 | 0.518793 | 0.478339 | 0.713187 | Healthy Control | Intestine |
| Gdf10 | 8.44E-01 | 2.879578 | 0.072219 | 0.791049 | 0.942631 | Infection | Intestine |
| Gdf11 | 1.64E+00 | 3.694903 | 1.860591 | 0.184978 | 0.349961 | Infection | Intestine |
| Gdf15 | 6.82E-01 | 3.004586 | 0.155276 | 0.696829 | 0.892753 | Infection | Intestine |
| Gdf2 | -7.56E-01 | 1.273429 | 0.015546 | 0.902199 | 0.969735 | Healthy Control | Intestine |
| Gdf5 | -4.94E-01 | 1.22813 | 0.009389 | 0.924171 | 0.969735 | Healthy Control | Intestine |
| Gdf7 | -4.01E+00 | 1.94662 | 0.334692 | 0.569243 | 0.797768 | Healthy Control | Intestine |
| Gdf9 | -1.52E+00 | 1.274322 | 0.038472 | 0.846787 | 0.954066 | Healthy Control | Intestine |
| Gdnf | 1.29E+00 | 4.959316 | 3.293975 | 0.082775 | 0.187413 | Infection | Intestine |
| Gdpd1 | -2.95E+00 | 8.711082 | 56.01312 | 1.11E-07 | 1.74E-06 | Healthy Control | Intestine |
| Gdpd2 | 1.88E-01 | 3.482484 | 0.007463 | 0.931916 | 0.969735 | Infection | Intestine |
| Gem | 1.17E+00 | 6.034282 | 2.213935 | 0.152725 | 0.302311 | Infection | Intestine |
| Gen1 | 2.05E-01 | 4.189674 | 0.022962 | 0.880821 | 0.968458 | Infection | Intestine |
| Gfap | 2.27E+00 | 2.859045 | 2.642732 | 0.115875 | 0.243727 | Infection | Intestine |
| Gfi1 | -2.92E+00 | 6.628932 | 8.311234 | 0.008717 | 0.029266 | Healthy Control | Intestine |
| Gfi1b | 6.00E+00 | 1.276247 | 5.575755 | 0.032212 | 0.086743 | Infection | Intestine |
| Gfpt1 | -4.52E+00 | 12.02438 | 298.1757 | 3.33E-14 | 5.96E-12 | Healthy Control | Intestine |

| | | | | | | | |
|---|---|---|---|---|---|---|---|
| Gfpt2 | 2.65E+00 | 4.827767 | 9.333508 | 0.005835 | 0.020884 | Infection | Intestine |
| Gfra1 | 1.80E+00 | 3.65718 | 3.054991 | 0.093098 | 0.204946 | Infection | Intestine |
| Gfra2 | 3.21E+00 | 5.933045 | 29.42444 | 3.68E-05 | 0.000257 | Infection | Intestine |
| Gfra3 | 9.32E-01 | 3.837292 | 0.27802 | 0.602718 | 0.826406 | Infection | Intestine |
| Ggnbp1 | -5.67E+00 | 3.336941 | 0.886841 | 0.357734 | 0.582236 | Healthy Control | Intestine |
| Ggnbp2 | 1.97E-02 | 8.964227 | 0.014436 | 0.905464 | 0.969735 | Infection | Intestine |
| Ggps1 | -3.65E+00 | 6.872752 | 14.41395 | 0.000965 | 0.004482 | Healthy Control | Intestine |
| Gh | -1.37E+00 | 1.228088 | 0.037791 | 0.848672 | 0.954066 | Healthy Control | Intestine |
| Ghr | -9.16E-01 | 7.999463 | 6.150464 | 0.021112 | 0.061842 | Healthy Control | Intestine |
| Ghrh | -2.15E+00 | 1.317599 | 0.073256 | 0.78953 | 0.941823 | Healthy Control | Intestine |
| Ghrhr | -3.58E+00 | 1.887577 | 0.272062 | 0.606116 | 0.829653 | Healthy Control | Intestine |
| Ghrl | -1.71E+00 | 1.320364 | 0.044908 | 0.834259 | 0.954066 | Healthy Control | Intestine |
| Ghsr | 4.84E+00 | 1.362251 | 4.458155 | 0.04515 | 0.115376 | Infection | Intestine |
| Gigyf2 | -9.20E-01 | 7.924542 | 8.170511 | 0.00907 | 0.030229 | Healthy Control | Intestine |
| Gimap4 | -2.12E+00 | 6.25477 | 1.819909 | 0.192562 | 0.360901 | Healthy Control | Intestine |
| Gip | -9.71E-16 | 1.180616 | 0 | 1 | 1 | Healthy Control | Intestine |
| Gipc1 | -2.57E-01 | 9.498731 | 2.16576 | 0.155413 | 0.306057 | Healthy Control | Intestine |
| Gipr | 2.33E+00 | 2.548931 | 2.460054 | 0.129662 | 0.265042 | Infection | Intestine |
| Gja1 | 1.17E+00 | 8.155126 | 11.49742 | 0.002998 | 0.011688 | Infection | Intestine |
| Gja3 | -9.71E-16 | 1.180616 | 0 | 1 | 1 | Healthy Control | Intestine |
| Gja4 | -1.33E+00 | 4.7975 | 0.55688 | 0.463127 | 0.698738 | Healthy Control | Intestine |
| Gja5 | -6.67E+00 | 4.155191 | 0.493858 | 0.493671 | 0.726929 | Healthy Control | Intestine |
| Gja8 | -3.57E-01 | 1.227417 | 0.006288 | 0.937908 | 0.969735 | Healthy Control | Intestine |
| Gjb1 | -9.78E+00 | 7.108984 | 22.64425 | 0.000123 | 0.000738 | Healthy Control | Intestine |
| Gjb2 | -9.32E+00 | 6.432455 | 11.9565 | 0.002514 | 0.010066 | Healthy Control | Intestine |
| Gjb6 | -9.71E-16 | 1.180616 | 0 | 1 | 1 | Healthy Control | Intestine |
| Gjc1 | 1.49E+00 | 5.22384 | 5.476248 | 0.028625 | 0.079081 | Infection | Intestine |
| Gjd2 | -2.44E+00 | 1.44451 | 0.116918 | 0.734694 | 0.915807 | Healthy Control | Intestine |
| Gk | -6.72E-01 | 8.454457 | 2.218026 | 0.149714 | 0.297998 | Healthy Control | Intestine |
| Gkn2 | -1.37E+00 | 1.229023 | 0.032546 | 0.859435 | 0.956553 | Healthy Control | Intestine |
| Gla | 1.89E-01 | 6.333685 | 0.13968 | 0.712113 | 0.900212 | Infection | Intestine |
| Glb1 | -1.04E+00 | 8.624644 | 15.49733 | 0.000711 | 0.003428 | Healthy Control | Intestine |
| Gldc | -3.50E+00 | 1.699707 | 0.268041 | 0.60858 | 0.831483 | Healthy Control | Intestine |
| Gldn | 4.97E+00 | 1.320548 | 4.142798 | 0.05605 | 0.137407 | Infection | Intestine |
| Glg1 | -2.13E+00 | 9.468497 | 101.5757 | 1.08E-09 | 3.53E-08 | Healthy Control | Intestine |
| Gli1 | 4.64E-01 | 5.560468 | 0.252893 | 0.619824 | 0.838379 | Infection | Intestine |
| Gli2 | 4.42E-01 | 4.650859 | 0.207299 | 0.65288 | 0.863896 | Infection | Intestine |
| Gli3 | 1.60E+00 | 5.152739 | 5.48171 | 0.028595 | 0.079043 | Infection | Intestine |
| Glis3 | -5.55E-01 | 7.430122 | 2.061173 | 0.164874 | 0.319782 | Healthy Control | Intestine |
| Glmp | -1.26E+00 | 7.695477 | 10.5363 | 0.003583 | 0.013692 | Healthy Control | Intestine |
| Glo1 | -1.27E+00 | 8.241915 | 11.72 | 0.002398 | 0.009666 | Healthy Control | Intestine |
| Glp1r | 5.44E-01 | 2.443738 | 0.111479 | 0.74088 | 0.921009 | Infection | Intestine |
| Glra1 | 3.47E+00 | 1.562322 | 2.078523 | 0.158413 | 0.310621 | Infection | Intestine |
| Glra2 | -1.37E+00 | 1.228088 | 0.037791 | 0.848672 | 0.954066 | Healthy Control | Intestine |
| Glra3 | -4.95E-01 | 1.228515 | 0.008877 | 0.926263 | 0.969735 | Healthy Control | Intestine |
| Glud1 | -1.07E+00 | 9.543855 | 34.65223 | 6.62E-06 | 5.75E-05 | Healthy Control | Intestine |
| Glyat | -2.38E+00 | 1.55883 | 0.111097 | 0.741076 | 0.921009 | Healthy Control | Intestine |
| Gm2a | -1.48E-01 | 8.968148 | 0.212256 | 0.649769 | 0.861198 | Healthy Control | Intestine |
| Gmds | -4.16E+00 | 10.276 | 171.4537 | 4.38E-12 | 3.72E-10 | Healthy Control | Intestine |
| Gmfb | -7.04E-01 | 8.465568 | 5.070941 | 0.034651 | 0.092169 | Healthy Control | Intestine |

| | | | | | | | |
|---|---|---|---|---|---|---|---|
| Gmnc | -4.95E-01 | 1.228515 | 0.008877 | 0.926263 | 0.969735 | Healthy Control | Intestine |
| Gmnn | -1.94E+00 | 7.278896 | 11.33312 | 0.002552 | 0.010181 | Healthy Control | Intestine |
| Gna11 | -1.20E+00 | 8.486377 | 12.17337 | 0.002088 | 0.008563 | Healthy Control | Intestine |
| Gna12 | 1.18E+00 | 6.768755 | 12.48422 | 0.001892 | 0.00788 | Infection | Intestine |
| Gna13 | -7.85E-01 | 7.924803 | 5.76044 | 0.025184 | 0.071469 | Healthy Control | Intestine |
| Gna14 | -3.55E+00 | 8.600186 | 34.55269 | 5.54E-06 | 4.88E-05 | Healthy Control | Intestine |
| Gnai1 | -4.39E-02 | 7.354768 | 0.006098 | 0.938547 | 0.969735 | Healthy Control | Intestine |
| Gnai2 | 2.52E-01 | 10.22475 | 4.334255 | 0.049302 | 0.123744 | Infection | Intestine |
| Gnai3 | 5.43E-02 | 9.298777 | 0.134815 | 0.717032 | 0.90301 | Infection | Intestine |
| Gnal | 7.98E-01 | 2.804458 | 0.200839 | 0.658352 | 0.867142 | Infection | Intestine |
| Gnao1 | 1.19E+00 | 6.705696 | 7.666744 | 0.011268 | 0.036416 | Infection | Intestine |
| Gnaq | -4.91E-01 | 8.79586 | 4.298198 | 0.050187 | 0.125304 | Healthy Control | Intestine |
| Gnat1 | -9.71E-16 | 1.180616 | 0 | 1 | 1 | Healthy Control | Intestine |
| Gnat2 | -5.06E+00 | 2.433977 | 0.716605 | 0.406716 | 0.637769 | Healthy Control | Intestine |
| Gnat3 | -5.21E+00 | 2.188961 | 0.717675 | 0.403755 | 0.634599 | Healthy Control | Intestine |
| Gnaz | -3.71E+00 | 2.107537 | 0.296604 | 0.590865 | 0.816577 | Healthy Control | Intestine |
| Gnb1 | -2.42E+00 | 10.10418 | 117.7096 | 2.90E-10 | 1.17E-08 | Healthy Control | Intestine |
| Gnb3 | -3.57E-01 | 1.228299 | 0.00618 | 0.938438 | 0.969735 | Healthy Control | Intestine |
| Gnrh1 | -5.30E+00 | 3.031143 | 0.881807 | 0.359011 | 0.58313 | Healthy Control | Intestine |
| Gnrhr | -2.31E+00 | 1.558314 | 0.108294 | 0.744509 | 0.923113 | Healthy Control | Intestine |
| Golga2 | 4.41E-01 | 7.714614 | 3.018088 | 0.096475 | 0.210437 | Infection | Intestine |
| Golga3 | -8.44E-01 | 8.418984 | 12.63186 | 0.001787 | 0.007517 | Healthy Control | Intestine |
| Golga5 | -8.40E-02 | 7.800047 | 0.058677 | 0.810862 | 0.949736 | Healthy Control | Intestine |
| Gopc | -9.94E-01 | 8.429065 | 10.71322 | 0.003489 | 0.013363 | Healthy Control | Intestine |
| Gorasp1 | -2.64E+00 | 7.41388 | 14.99293 | 0.00073 | 0.003504 | Healthy Control | Intestine |
| Gosr1 | 2.57E-01 | 7.57371 | 0.584244 | 0.45285 | 0.686424 | Infection | Intestine |
| Got2 | -1.91E+00 | 8.4251 | 31.62037 | 1.07E-05 | 8.72E-05 | Healthy Control | Intestine |
| Gpa33 | -5.03E+00 | 12.55029 | 1326.861 | 4.00E-21 | 6.68E-18 | Healthy Control | Intestine |
| Gpam | -7.17E-01 | 7.836071 | 3.409162 | 0.07828 | 0.179548 | Healthy Control | Intestine |
| Gpat2 | 3.69E+00 | 1.738015 | 2.011906 | 0.167085 | 0.322821 | Infection | Intestine |
| Gpbar1 | 3.45E+00 | 1.739209 | 4.99524 | 0.032612 | 0.08763 | Infection | Intestine |
| Gpc3 | 1.23E+00 | 4.794268 | 1.302117 | 0.265537 | 0.463971 | Infection | Intestine |
| Gpc4 | -2.97E+00 | 9.482128 | 54.60856 | 1.56E-07 | 2.27E-06 | Healthy Control | Intestine |
| Gpc6 | 2.23E+00 | 5.700127 | 18.47901 | 0.000297 | 0.001616 | Infection | Intestine |
| Gpd1 | -9.01E+00 | 6.280747 | 15.9844 | 0.000718 | 0.00346 | Healthy Control | Intestine |
| Gper1 | 6.86E-01 | 3.275627 | 0.146444 | 0.705494 | 0.896371 | Infection | Intestine |
| Gphn | -1.30E+00 | 7.211939 | 6.782305 | 0.015692 | 0.048192 | Healthy Control | Intestine |
| Gpi1 | -1.22E+00 | 9.710912 | 44.35213 | 1.13E-06 | 1.25E-05 | Healthy Control | Intestine |
| Gpm6a | -5.70E+00 | 2.953341 | 0.84878 | 0.368113 | 0.593868 | Healthy Control | Intestine |
| Gpnmb | -7.99E+00 | 5.543533 | 4.13041 | 0.055747 | 0.136933 | Healthy Control | Intestine |
| Gpr101 | -6.42E-01 | 1.272809 | 0.01342 | 0.908986 | 0.969735 | Healthy Control | Intestine |
| Gpr107 | -4.41E-01 | 7.732667 | 2.821475 | 0.10718 | 0.228998 | Healthy Control | Intestine |
| Gpr139 | -1.41E+00 | 1.40466 | 0.03335 | 0.856402 | 0.956553 | Healthy Control | Intestine |
| Gpr141b | -5.86E+00 | 3.321767 | 0.78473 | 0.386438 | 0.615928 | Healthy Control | Intestine |
| Gpr143 | -9.71E-16 | 1.180616 | 0 | 1 | 1 | Healthy Control | Intestine |
| Gpr149 | -2.89E+00 | 1.523927 | 0.117144 | 0.734309 | 0.915783 | Healthy Control | Intestine |
| Gpr17 | -5.19E+00 | 2.836572 | 0.703179 | 0.411206 | 0.643482 | Healthy Control | Intestine |
| Gpr183 | -1.68E+00 | 4.984558 | 1.382034 | 0.252125 | 0.445671 | Healthy Control | Intestine |
| Gpr20 | -3.40E+00 | 6.155798 | 5.869359 | 0.02452 | 0.069982 | Healthy Control | Intestine |
| Gpr22 | 3.76E+00 | 1.526185 | 2.205812 | 0.146502 | 0.292886 | Infection | Intestine |

| | | | | | | | |
|---|---|---|---|---|---|---|---|
| Gpr26 | -3.57E-01 | 1.227387 | 0.006293 | 0.937883 | 0.969735 | Healthy Control | Intestine |
| Gpr35 | -2.65E-01 | 6.358797 | 0.19366 | 0.663932 | 0.871205 | Healthy Control | Intestine |
| Gpr37l1 | 9.99E-01 | 3.137616 | 0.251588 | 0.620411 | 0.838493 | Infection | Intestine |
| Gpr39 | 1.06E+00 | 6.108581 | 4.110697 | 0.054933 | 0.135199 | Infection | Intestine |
| Gpr4 | 4.77E-01 | 2.776737 | 0.054006 | 0.818436 | 0.952901 | Infection | Intestine |
| Gpr50 | -6.42E-01 | 1.272718 | 0.010492 | 0.919483 | 0.969735 | Healthy Control | Intestine |
| Gpr6 | 5.22E+00 | 1.276391 | 4.451111 | 0.052242 | 0.129789 | Infection | Intestine |
| Gpr65 | 2.70E+00 | 4.641914 | 23.02297 | 8.61E-05 | 0.000544 | Infection | Intestine |
| Gpr83 | 1.87E+00 | 2.700242 | 1.115442 | 0.300211 | 0.509645 | Infection | Intestine |
| Gpr84 | -2.25E+00 | 1.362964 | 0.085762 | 0.772131 | 0.934096 | Healthy Control | Intestine |
| Gpr88 | 3.72E+00 | 2.250606 | 6.660106 | 0.015445 | 0.047493 | Infection | Intestine |
| Gprc5b | 1.80E+00 | 5.904356 | 9.699285 | 0.005069 | 0.018467 | Infection | Intestine |
| Gprc6a | 4.10E+00 | 1.406014 | 2.842028 | 0.104131 | 0.223629 | Infection | Intestine |
| Gps1 | -1.67E+00 | 8.9399 | 39.72304 | 2.43E-06 | 2.35E-05 | Healthy Control | Intestine |
| Gpsm1 | 1.03E+00 | 5.745812 | 2.720845 | 0.113049 | 0.238787 | Infection | Intestine |
| Gpsm2 | -2.32E+00 | 8.5816 | 31.1366 | 1.14E-05 | 9.29E-05 | Healthy Control | Intestine |
| Gpt2 | -8.05E-01 | 8.381462 | 5.336408 | 0.03068 | 0.083695 | Healthy Control | Intestine |
| Gpx2 | -2.59E+00 | 7.043489 | 10.55353 | 0.003448 | 0.013246 | Healthy Control | Intestine |
| Gpx5 | -3.57E-01 | 1.227387 | 0.006293 | 0.937883 | 0.969735 | Healthy Control | Intestine |
| Gramd1a | 2.17E+00 | 7.08818 | 63.04267 | 7.17E-08 | 1.21E-06 | Infection | Intestine |
| Gramd1b | 8.34E-01 | 6.851187 | 4.991878 | 0.035959 | 0.095142 | Infection | Intestine |
| Grb10 | 1.97E+00 | 6.475399 | 20.15377 | 0.000187 | 0.00107 | Infection | Intestine |
| Grb2 | 1.38E+00 | 8.05283 | 45.04886 | 1.00E-06 | 1.13E-05 | Infection | Intestine |
| Grb7 | -4.86E+00 | 8.248505 | 35.3046 | 4.85E-06 | 4.36E-05 | Healthy Control | Intestine |
| Greb1 | -8.90E+00 | 5.964146 | 6.278385 | 0.021071 | 0.061759 | Healthy Control | Intestine |
| Grem1 | 2.77E+00 | 6.427409 | 8.876018 | 0.010979 | 0.035574 | Infection | Intestine |
| Grhl2 | -3.16E+00 | 7.468432 | 20.74135 | 0.000132 | 0.000784 | Healthy Control | Intestine |
| Grhl3 | -5.37E+00 | 3.350882 | 0.789565 | 0.384436 | 0.61348 | Healthy Control | Intestine |
| Gria1 | -3.12E+00 | 1.524095 | 0.151482 | 0.699985 | 0.89357 | Healthy Control | Intestine |
| Gria2 | 3.05E+00 | 2.174362 | 3.802853 | 0.061058 | 0.147453 | Infection | Intestine |
| Gria3 | 3.53E+00 | 4.423002 | 51.96607 | 3.42E-07 | 4.49E-06 | Infection | Intestine |
| Gria4 | -8.29E-01 | 4.446478 | 0.312607 | 0.581533 | 0.808205 | Healthy Control | Intestine |
| Grid1 | -2.28E+00 | 1.363258 | 0.092338 | 0.764007 | 0.930099 | Healthy Control | Intestine |
| Grid2 | -1.37E+00 | 1.229023 | 0.032546 | 0.859435 | 0.956553 | Healthy Control | Intestine |
| Grifin | -1.07E+00 | 1.31889 | 0.020035 | 0.888883 | 0.969599 | Healthy Control | Intestine |
| Grik1 | -2.95E+00 | 1.559526 | 0.172756 | 0.680244 | 0.882694 | Healthy Control | Intestine |
| Grik2 | -4.37E+00 | 2.252655 | 0.329209 | 0.571934 | 0.80042 | Healthy Control | Intestine |
| Grik3 | -1.40E+00 | 4.643887 | 0.716039 | 0.406466 | 0.637769 | Healthy Control | Intestine |
| Grik4 | 3.26E+00 | 1.94862 | 1.908762 | 0.180454 | 0.343219 | Infection | Intestine |
| Grin1 | -3.64E+00 | 1.916916 | 0.263523 | 0.611953 | 0.833585 | Healthy Control | Intestine |
| Grin2a | -1.52E+00 | 1.273388 | 0.043339 | 0.837524 | 0.954066 | Healthy Control | Intestine |
| Grin2b | -1.08E+00 | 1.360994 | 0.02378 | 0.87873 | 0.967178 | Healthy Control | Intestine |
| Grin2c | -9.71E-16 | 1.180616 | 0 | 1 | 1 | Healthy Control | Intestine |
| Grin2d | -2.01E-02 | 5.77281 | 0.001436 | 0.970091 | 0.992279 | Healthy Control | Intestine |
| Grk2 | 9.49E-02 | 9.625237 | 0.49387 | 0.489643 | 0.722876 | Infection | Intestine |
| Grk3 | 1.27E-01 | 5.988805 | 0.061273 | 0.806671 | 0.948508 | Infection | Intestine |
| Grk5 | 8.09E-01 | 6.011743 | 1.645148 | 0.212818 | 0.390244 | Infection | Intestine |
| Grm1 | -2.36E+00 | 1.405533 | 0.074577 | 0.786825 | 0.941182 | Healthy Control | Intestine |
| Grm2 | -9.77E-01 | 1.318411 | 0.021477 | 0.884798 | 0.96867 | Healthy Control | Intestine |
| Grm3 | 6.14E+00 | 1.275333 | 6.502241 | 0.022961 | 0.066478 | Infection | Intestine |

| | | | | | | | |
|---|---|---|---|---|---|---|---|
| Grm4 | -5.07E+00 | 2.586364 | 0.654665 | 0.426951 | 0.659869 | Healthy Control | Intestine |
| Grm6 | 4.80E+00 | 1.739008 | 12.64046 | 0.001017 | 0.004698 | Infection | Intestine |
| Grm8 | -3.57E-01 | 1.228299 | 0.00618 | 0.938438 | 0.969735 | Healthy Control | Intestine |
| Grn | 1.92E+00 | 9.407942 | 174.5625 | 6.98E-12 | 5.54E-10 | Infection | Intestine |
| Grp | -8.79E-01 | 1.316262 | 0.018104 | 0.894148 | 0.969735 | Healthy Control | Intestine |
| Grpel1 | -1.61E+00 | 8.237151 | 30.30146 | 1.47E-05 | 0.000116 | Healthy Control | Intestine |
| Grpr | -9.64E-01 | 4.578366 | 0.397539 | 0.535175 | 0.767123 | Healthy Control | Intestine |
| Grxcr1 | -3.57E-01 | 1.227387 | 0.006293 | 0.937883 | 0.969735 | Healthy Control | Intestine |
| Grxcr2 | 4.84E+00 | 1.363429 | 3.877543 | 0.060378 | 0.146086 | Infection | Intestine |
| Gsc | -1.07E+00 | 1.31916 | 0.027694 | 0.869536 | 0.961117 | Healthy Control | Intestine |
| Gsc2 | 3.69E+00 | 1.447967 | 2.254703 | 0.144356 | 0.28929 | Infection | Intestine |
| Gsdma3 | -3.57E-01 | 1.227417 | 0.006288 | 0.937908 | 0.969735 | Healthy Control | Intestine |
| Gsdmd | 1.55E+00 | 9.369741 | 73.56807 | 1.96E-08 | 3.93E-07 | Infection | Intestine |
| Gsdme | 4.77E-02 | 5.762808 | 0.003794 | 0.951409 | 0.978983 | Infection | Intestine |
| Gsk3a | -6.82E-01 | 9.264704 | 12.59177 | 0.001823 | 0.007645 | Healthy Control | Intestine |
| Gsk3b | 4.98E-02 | 7.827073 | 0.041852 | 0.839798 | 0.954066 | Infection | Intestine |
| Gsta1 | -1.76E+00 | 6.328104 | 2.216749 | 0.148872 | 0.296678 | Healthy Control | Intestine |
| Gstk1 | -1.04E+01 | 7.57676 | 34.23417 | 1.05E-05 | 8.56E-05 | Healthy Control | Intestine |
| Gstp1 | -2.34E+00 | 9.825161 | 54.62187 | 2.29E-07 | 3.15E-06 | Healthy Control | Intestine |
| Gsx1 | 6.14E+00 | 1.274983 | 6.569448 | 0.022381 | 0.064988 | Infection | Intestine |
| Gsx2 | -9.71E-16 | 1.180616 | 0 | 1 | 1 | Healthy Control | Intestine |
| Gtf2a1 | 5.74E-01 | 8.059657 | 6.680439 | 0.016993 | 0.051556 | Infection | Intestine |
| Gtf2a2 | 2.85E-01 | 8.254824 | 1.837347 | 0.189162 | 0.355594 | Infection | Intestine |
| Gtf2b | -2.83E-01 | 6.420616 | 0.306106 | 0.585364 | 0.810829 | Healthy Control | Intestine |
| Gtf2e1 | 9.24E-02 | 5.472717 | 0.017616 | 0.895515 | 0.969735 | Infection | Intestine |
| Gtf2e2 | -7.63E-01 | 5.719619 | 0.699856 | 0.411146 | 0.643482 | Healthy Control | Intestine |
| Gtf2h1 | 2.18E-01 | 7.509467 | 0.50255 | 0.485833 | 0.720189 | Infection | Intestine |
| Gtf2i | -4.29E-01 | 9.116029 | 4.568215 | 0.044057 | 0.112986 | Healthy Control | Intestine |
| Gtpbp2 | 1.68E-01 | 8.94937 | 0.748714 | 0.396312 | 0.625454 | Infection | Intestine |
| Gtse1 | 9.76E-01 | 4.700344 | 2.268273 | 0.14518 | 0.290709 | Infection | Intestine |
| Gucy1a1 | -8.92E-01 | 7.606652 | 0.709003 | 0.413758 | 0.646667 | Healthy Control | Intestine |
| Gucy2e | -9.71E-16 | 1.180616 | 0 | 1 | 1 | Healthy Control | Intestine |
| Gucy2f | 4.97E+00 | 1.31993 | 4.206254 | 0.054373 | 0.134084 | Infection | Intestine |
| Gusb | 4.91E-01 | 5.23099 | 0.431415 | 0.51759 | 0.752996 | Infection | Intestine |
| Gyg | -2.03E+00 | 6.985794 | 14.71376 | 0.000798 | 0.003785 | Healthy Control | Intestine |
| Gys2 | 4.97E+00 | 1.320427 | 4.148628 | 0.055893 | 0.137181 | Infection | Intestine |
| Gzma | 9.61E-01 | 3.086772 | 0.094622 | 0.761389 | 0.928591 | Infection | Intestine |
| Gzmb | -2.97E+00 | 1.891447 | 0.164546 | 0.68859 | 0.887279 | Healthy Control | Intestine |
| Gzmk | -6.42E-01 | 1.274344 | 0.012111 | 0.91352 | 0.969735 | Healthy Control | Intestine |
| H19 | 3.76E+00 | 2.284368 | 5.329898 | 0.02878 | 0.079424 | Infection | Intestine |
| H1foo | -2.43E+00 | 1.406608 | 0.131103 | 0.720931 | 0.906205 | Healthy Control | Intestine |
| H2-DMb2 | -1.38E+00 | 4.644645 | 0.738802 | 0.39922 | 0.629052 | Healthy Control | Intestine |
| H2afv | 4.18E-01 | 5.873333 | 0.527346 | 0.4751 | 0.709743 | Infection | Intestine |
| H2afx | 8.59E-01 | 7.858252 | 10.94388 | 0.003231 | 0.012486 | Infection | Intestine |
| H2afy | -1.90E+00 | 8.942437 | 54.80561 | 2.05E-07 | 2.88E-06 | Healthy Control | Intestine |
| H2afz | -1.03E+00 | 9.051029 | 20.9033 | 0.000151 | 0.000883 | Healthy Control | Intestine |
| Hace1 | 2.40E-01 | 6.172608 | 0.196203 | 0.661937 | 0.869498 | Infection | Intestine |
| Hacl1 | 3.36E-01 | 5.708706 | 0.279982 | 0.601713 | 0.825737 | Infection | Intestine |
| Hadh | -3.23E+00 | 10.92418 | 219.0057 | 7.47E-13 | 9.13E-11 | Healthy Control | Intestine |
| Hadhb | -2.86E+00 | 9.835076 | 107.9758 | 5.97E-10 | 2.12E-08 | Healthy Control | Intestine |

| | | | | | | | |
|---|---|---|---|---|---|---|---|
| Hal | -1.17E-01 | 5.610008 | 0.004718 | 0.945973 | 0.975194 | Healthy Control | Intestine |
| Hamp | -1.34E+00 | 1.401998 | 0.028579 | 0.866997 | 0.96049 | Healthy Control | Intestine |
| Hand1 | -4.44E+00 | 2.08839 | 0.3759 | 0.545283 | 0.776804 | Healthy Control | Intestine |
| Hand2 | 8.58E-01 | 2.769745 | 0.200538 | 0.658667 | 0.867142 | Infection | Intestine |
| Hao1 | -3.53E+00 | 1.670608 | 0.257109 | 0.616364 | 0.836864 | Healthy Control | Intestine |
| Hap1 | -7.15E+00 | 4.250259 | 3.497403 | 0.076357 | 0.175801 | Healthy Control | Intestine |
| Hapln1 | -7.21E+00 | 4.910959 | 2.922655 | 0.103015 | 0.222044 | Healthy Control | Intestine |
| Has2 | 2.76E+00 | 3.756982 | 5.996778 | 0.022264 | 0.064724 | Infection | Intestine |
| Haspin | 1.02E+00 | 4.831508 | 1.443838 | 0.241567 | 0.431117 | Infection | Intestine |
| Hat1 | -9.43E-01 | 7.514453 | 5.378438 | 0.029822 | 0.081668 | Healthy Control | Intestine |
| Havcr1 | -3.25E+00 | 1.486495 | 0.220041 | 0.643479 | 0.857095 | Healthy Control | Intestine |
| Havcr2 | 1.84E-01 | 5.307986 | 0.04457 | 0.834559 | 0.954066 | Infection | Intestine |
| Hba-x | -3.57E-01 | 1.227387 | 0.006293 | 0.937883 | 0.969735 | Healthy Control | Intestine |
| Hbegf | 8.50E-01 | 4.930571 | 0.754535 | 0.393763 | 0.623198 | Infection | Intestine |
| Hc | 2.54E+00 | 2.357781 | 3.551394 | 0.068741 | 0.162473 | Infection | Intestine |
| Hcar2 | -4.09E+00 | 2.27672 | 0.33017 | 0.571183 | 0.799611 | Healthy Control | Intestine |
| Hcfc1 | -5.88E-01 | 8.754028 | 7.852672 | 0.010446 | 0.03409 | Healthy Control | Intestine |
| Hck | 3.68E+00 | 5.808698 | 92.83348 | 2.61E-09 | 7.43E-08 | Infection | Intestine |
| Hcn1 | -1.83E+00 | 1.36231 | 0.049723 | 0.825323 | 0.953685 | Healthy Control | Intestine |
| Hcn2 | -6.72E+00 | 3.880251 | 2.68608 | 0.117093 | 0.245258 | Healthy Control | Intestine |
| Hcn3 | -3.52E+00 | 1.732173 | 0.257796 | 0.616304 | 0.836864 | Healthy Control | Intestine |
| Hcn4 | -3.42E+00 | 1.59937 | 0.217996 | 0.644226 | 0.857095 | Healthy Control | Intestine |
| Hcrt | -9.76E-01 | 1.317502 | 0.018574 | 0.892814 | 0.969735 | Healthy Control | Intestine |
| Hcrtr1 | -2.09E+00 | 1.444487 | 0.06839 | 0.795408 | 0.943769 | Healthy Control | Intestine |
| Hcrtr2 | -1.89E+00 | 1.401998 | 0.062915 | 0.803831 | 0.947487 | Healthy Control | Intestine |
| Hdac1 | -1.10E+00 | 8.189276 | 18.81604 | 0.000265 | 0.00146 | Healthy Control | Intestine |
| Hdac2 | 2.51E-01 | 7.66884 | 0.605719 | 0.44474 | 0.678358 | Infection | Intestine |
| Hdac3 | -1.84E+00 | 8.321819 | 24.34439 | 5.76E-05 | 0.00038 | Healthy Control | Intestine |
| Hdac4 | -3.41E-01 | 8.040403 | 1.643801 | 0.2133 | 0.390699 | Healthy Control | Intestine |
| Hdac5 | -1.53E+00 | 7.275505 | 10.24496 | 0.003865 | 0.014646 | Healthy Control | Intestine |
| Hdac6 | 7.23E-01 | 6.61574 | 3.394758 | 0.078967 | 0.180589 | Infection | Intestine |
| Hdac8 | -1.11E+00 | 5.554195 | 1.298478 | 0.265686 | 0.464007 | Healthy Control | Intestine |
| Hdac9 | 1.46E+00 | 3.719712 | 2.056216 | 0.164177 | 0.319173 | Infection | Intestine |
| Hdc | 1.76E+00 | 1.892627 | 0.431642 | 0.516237 | 0.751463 | Infection | Intestine |
| Hdgfl2 | 1.13E+00 | 7.084025 | 14.02263 | 0.001138 | 0.005141 | Infection | Intestine |
| Hecw2 | -7.31E+00 | 4.132644 | 1.715257 | 0.205361 | 0.37949 | Healthy Control | Intestine |
| Hells | 4.29E-01 | 6.338544 | 0.537725 | 0.471038 | 0.706162 | Infection | Intestine |
| Herc6 | 3.79E-01 | 6.244134 | 0.420178 | 0.523341 | 0.7574 | Infection | Intestine |
| Hes1 | -1.81E+00 | 9.398898 | 27.08443 | 3.04E-05 | 0.000218 | Healthy Control | Intestine |
| Hes2 | -5.99E+00 | 3.362631 | 1.377405 | 0.254236 | 0.44861 | Healthy Control | Intestine |
| Hes5 | -1.34E+00 | 1.400951 | 0.037867 | 0.847145 | 0.954066 | Healthy Control | Intestine |
| Hes6 | -7.87E-01 | 8.349864 | 5.955295 | 0.023063 | 0.066698 | Healthy Control | Intestine |
| Hesx1 | -1.68E+00 | 1.480699 | 0.053178 | 0.819223 | 0.953035 | Healthy Control | Intestine |
| Hexa | -5.48E-01 | 9.475009 | 9.142168 | 0.006289 | 0.022329 | Healthy Control | Intestine |
| Hey1 | -1.25E+00 | 4.521877 | 0.461212 | 0.504044 | 0.738006 | Healthy Control | Intestine |
| Hey2 | 4.92E-01 | 3.276799 | 0.069288 | 0.794725 | 0.943769 | Infection | Intestine |
| Heyl | 1.91E+00 | 3.866646 | 3.496979 | 0.073808 | 0.171455 | Infection | Intestine |
| Hfe | 1.76E-01 | 3.445125 | 0.012204 | 0.913019 | 0.969735 | Infection | Intestine |
| Hfm1 | -1.79E+00 | 1.483656 | 0.059808 | 0.808659 | 0.94887 | Healthy Control | Intestine |
| Hgf | 1.23E+00 | 6.093312 | 4.203995 | 0.052365 | 0.130029 | Infection | Intestine |

| | | | | | | | |
|---|---|---|---|---|---|---|---|
| Hgs | -1.38E-02 | 7.979185 | 0.003141 | 0.955814 | 0.981696 | Healthy Control | Intestine |
| Hhex | 3.35E-01 | 5.356437 | 0.111256 | 0.741661 | 0.921507 | Infection | Intestine |
| Hibadh | -1.75E+00 | 8.936539 | 33.34735 | 7.75E-06 | 6.64E-05 | Healthy Control | Intestine |
| Hic1 | 8.56E-01 | 7.364114 | 4.823644 | 0.03903 | 0.101921 | Infection | Intestine |
| Hif1a | -1.11E+00 | 8.595153 | 15.16577 | 0.000753 | 0.003594 | Healthy Control | Intestine |
| Hif1an | -7.86E-01 | 7.371729 | 3.270619 | 0.083909 | 0.189126 | Healthy Control | Intestine |
| Hif3a | -6.58E+00 | 3.830806 | 2.014147 | 0.171475 | 0.330155 | Healthy Control | Intestine |
| Hikeshi | -6.08E+00 | 3.295302 | 1.873797 | 0.186546 | 0.352131 | Healthy Control | Intestine |
| Hip1r | 2.51E-01 | 8.42492 | 1.248541 | 0.276011 | 0.477769 | Infection | Intestine |
| Hipk2 | -3.71E-01 | 8.325041 | 1.695625 | 0.206482 | 0.38128 | Healthy Control | Intestine |
| Hira | 1.32E-01 | 7.047102 | 0.102173 | 0.752254 | 0.925708 | Infection | Intestine |
| Hist1h2ba | -3.57E-01 | 1.228165 | 0.006192 | 0.93838 | 0.969735 | Healthy Control | Intestine |
| Hist1h2bc | -4.00E+00 | 9.825222 | 71.6884 | 1.55E-08 | 3.18E-07 | Healthy Control | Intestine |
| Hist1h4h | -3.49E-01 | 7.26894 | 0.370695 | 0.54886 | 0.778399 | Healthy Control | Intestine |
| Hist2h2bb | 4.09E-01 | 10.42639 | 1.373566 | 0.261279 | 0.45765 | Infection | Intestine |
| Hist2h2be | -1.35E+00 | 5.621494 | 2.194441 | 0.151639 | 0.300754 | Healthy Control | Intestine |
| Hivep2 | -1.39E+00 | 7.656987 | 11.44929 | 0.00255 | 0.01018 | Healthy Control | Intestine |
| Hjv | -6.42E-01 | 1.272658 | 0.010513 | 0.919403 | 0.969735 | Healthy Control | Intestine |
| Hk1 | -2.10E+00 | 8.78806 | 39.12801 | 2.41E-06 | 2.34E-05 | Healthy Control | Intestine |
| Hk2 | -1.43E+00 | 8.739709 | 12.04364 | 0.002274 | 0.009225 | Healthy Control | Intestine |
| Hkdc1 | -8.77E+00 | 6.113448 | 5.189119 | 0.03406 | 0.090887 | Healthy Control | Intestine |
| Hlf | 9.63E-02 | 4.906654 | 0.007517 | 0.931624 | 0.969735 | Infection | Intestine |
| Hmcn1 | -7.14E-02 | 5.975906 | 0.01007 | 0.920938 | 0.969735 | Healthy Control | Intestine |
| Hmcn2 | 5.72E-01 | 8.803654 | 0.677845 | 0.425796 | 0.658695 | Infection | Intestine |
| Hmga1 | -3.96E+00 | 7.3147 | 25.71353 | 4.11E-05 | 0.000283 | Healthy Control | Intestine |
| Hmga1b | -1.23E+00 | 5.732927 | 1.374421 | 0.252682 | 0.446496 | Healthy Control | Intestine |
| Hmga2 | 8.24E-01 | 6.51185 | 2.310025 | 0.142734 | 0.286843 | Infection | Intestine |
| Hmgcr | -1.75E+00 | 9.212102 | 30.01038 | 1.67E-05 | 0.000131 | Healthy Control | Intestine |
| Hmox1 | 3.90E+00 | 5.336785 | 121.8452 | 2.19E-10 | 9.31E-09 | Infection | Intestine |
| Hmx2 | 1.53E+00 | 2.110774 | 0.753305 | 0.3939 | 0.623218 | Infection | Intestine |
| Hmx3 | 3.99E+00 | 1.446703 | 2.904323 | 0.098568 | 0.213978 | Infection | Intestine |
| Hnf1a | -4.29E+00 | 7.570526 | 36.82378 | 3.68E-06 | 3.40E-05 | Healthy Control | Intestine |
| Hnf1b | -9.76E+00 | 7.035334 | 16.13918 | 0.000687 | 0.003334 | Healthy Control | Intestine |
| Hnf4a | -3.67E+00 | 11.22925 | 285.9842 | 4.35E-14 | 7.51E-12 | Healthy Control | Intestine |
| Hnrnpa1 | -2.26E+00 | 11.15412 | 204.6403 | 1.50E-12 | 1.62E-10 | Healthy Control | Intestine |
| Hnrnpdl | -3.82E-01 | 8.605544 | 2.750427 | 0.111549 | 0.236115 | Healthy Control | Intestine |
| Hnrnph1 | 7.30E-01 | 9.754005 | 37.72861 | 3.64E-06 | 3.37E-05 | Infection | Intestine |
| Hnrnpk | 1.45E+00 | 9.155633 | 126.9147 | 1.47E-10 | 7.16E-09 | Infection | Intestine |
| Hnrnpm | -3.35E-01 | 9.896834 | 5.173241 | 0.033138 | 0.088759 | Healthy Control | Intestine |
| Homer1 | -8.32E-01 | 6.412947 | 1.414238 | 0.246003 | 0.437319 | Healthy Control | Intestine |
| Homer3 | 1.05E+00 | 5.294633 | 2.05972 | 0.164686 | 0.319543 | Infection | Intestine |
| Hopx | -3.97E+00 | 8.334113 | 41.05028 | 1.29E-06 | 1.39E-05 | Healthy Control | Intestine |
| Hoxa1 | 4.27E-01 | 3.339609 | 0.072839 | 0.789619 | 0.941823 | Infection | Intestine |
| Hoxa10 | -2.39E+00 | 8.764771 | 57.87799 | 1.30E-07 | 1.97E-06 | Healthy Control | Intestine |
| Hoxa11 | 1.42E+00 | 4.597313 | 2.87587 | 0.103083 | 0.222044 | Infection | Intestine |
| Hoxa13 | -1.04E+01 | 7.611401 | 49.27435 | 3.53E-07 | 4.61E-06 | Healthy Control | Intestine |
| Hoxa2 | -7.37E+00 | 4.478719 | 4.092741 | 0.056809 | 0.138794 | Healthy Control | Intestine |
| Hoxa3 | -2.52E+00 | 7.264473 | 14.39005 | 0.00092 | 0.004316 | Healthy Control | Intestine |
| Hoxa4 | -1.09E-01 | 5.06301 | 0.016714 | 0.898261 | 0.969735 | Healthy Control | Intestine |
| Hoxa5 | -2.01E+00 | 5.386179 | 2.445129 | 0.132007 | 0.268956 | Healthy Control | Intestine |

| | | | | | | | |
|---|---|---|---|---|---|---|---|
| Hoxa7 | -1.09E+00 | 7.103213 | 4.157095 | 0.053246 | 0.131812 | Healthy Control | Intestine |
| Hoxa9 | -1.28E+00 | 7.953426 | 13.32364 | 0.001475 | 0.006387 | Healthy Control | Intestine |
| Hoxb1 | -9.71E-16 | 1.180616 | 0 | 1 | 1 | Healthy Control | Intestine |
| Hoxb13 | -1.21E+01 | 9.503797 | 175.676 | 3.50E-12 | 3.18E-10 | Healthy Control | Intestine |
| Hoxb2 | 2.59E+00 | 2.574374 | 2.633963 | 0.115587 | 0.243224 | Infection | Intestine |
| Hoxb3 | 1.05E+00 | 6.821973 | 1.151506 | 0.29755 | 0.506473 | Infection | Intestine |
| Hoxb4 | 6.30E-01 | 6.187039 | 0.517501 | 0.478943 | 0.713187 | Infection | Intestine |
| Hoxb5 | 3.07E-01 | 7.452153 | 0.093724 | 0.763418 | 0.929847 | Infection | Intestine |
| Hoxb6 | 4.91E-02 | 6.891414 | 0.001952 | 0.96526 | 0.988562 | Infection | Intestine |
| Hoxb7 | -7.35E+00 | 4.976962 | 3.150574 | 0.091288 | 0.202068 | Healthy Control | Intestine |
| Hoxb8 | -3.11E+00 | 7.017512 | 6.513482 | 0.018993 | 0.056729 | Healthy Control | Intestine |
| Hoxb9 | -4.74E+00 | 8.996751 | 90.73932 | 9.61E-10 | 3.21E-08 | Healthy Control | Intestine |
| Hoxc10 | 3.22E+00 | 1.868988 | 1.873158 | 0.185477 | 0.350641 | Infection | Intestine |
| Hoxc11 | 3.22E+00 | 1.774399 | 1.897769 | 0.186783 | 0.352445 | Infection | Intestine |
| Hoxc12 | -9.71E-16 | 1.180616 | 0 | 1 | 1 | Healthy Control | Intestine |
| Hoxc5 | -1.15E+00 | 1.319302 | 0.029155 | 0.866595 | 0.96049 | Healthy Control | Intestine |
| Hoxc6 | 4.10E+00 | 2.610315 | 9.340313 | 0.004282 | 0.015996 | Infection | Intestine |
| Hoxc8 | 6.19E+00 | 1.711427 | 22.96582 | 2.79E-05 | 0.000202 | Infection | Intestine |
| Hoxd1 | -2.35E+00 | 1.633263 | 0.076794 | 0.783948 | 0.940206 | Healthy Control | Intestine |
| Hoxd10 | 2.78E+00 | 3.98363 | 2.880779 | 0.103365 | 0.222462 | Infection | Intestine |
| Hoxd11 | 1.59E+00 | 5.606864 | 0.5089 | 0.48709 | 0.721412 | Infection | Intestine |
| Hoxd12 | 4.52E+00 | 1.557788 | 5.026203 | 0.035871 | 0.095002 | Infection | Intestine |
| Hoxd13 | -3.84E+00 | 8.790809 | 2.058152 | 0.179783 | 0.342463 | Healthy Control | Intestine |
| Hoxd3 | -3.80E+00 | 2.214571 | 0.402746 | 0.531005 | 0.764952 | Healthy Control | Intestine |
| Hoxd4 | 2.11E+00 | 3.604978 | 2.875442 | 0.103027 | 0.222044 | Infection | Intestine |
| Hoxd9 | 3.49E+00 | 5.42732 | 13.75962 | 0.002055 | 0.008447 | Infection | Intestine |
| Hpcal4 | -4.59E+00 | 2.13974 | 0.517244 | 0.478959 | 0.713187 | Healthy Control | Intestine |
| Hpd | -5.68E+00 | 8.499938 | 30.07518 | 3.60E-05 | 0.000253 | Healthy Control | Intestine |
| Hpgds | -8.81E-01 | 5.819475 | 1.187275 | 0.286551 | 0.49109 | Healthy Control | Intestine |
| Hprt | -3.67E-01 | 8.542338 | 1.811373 | 0.192186 | 0.360466 | Healthy Control | Intestine |
| Hps4 | -9.13E-01 | 6.858489 | 2.944907 | 0.099296 | 0.215463 | Healthy Control | Intestine |
| Hps5 | -1.74E-01 | 8.377704 | 0.649052 | 0.429144 | 0.661421 | Healthy Control | Intestine |
| Hps6 | -1.90E+00 | 5.320043 | 2.225856 | 0.149502 | 0.297697 | Healthy Control | Intestine |
| Hpse | -5.76E-01 | 7.013058 | 1.307809 | 0.264676 | 0.462951 | Healthy Control | Intestine |
| Hpse2 | 2.23E+00 | 3.605704 | 3.527174 | 0.072525 | 0.169181 | Infection | Intestine |
| Hr | -5.50E-01 | 6.52096 | 0.831915 | 0.370989 | 0.596778 | Healthy Control | Intestine |
| Hras | -9.82E-01 | 4.378096 | 0.463944 | 0.502759 | 0.73634 | Healthy Control | Intestine |
| Hrg | 3.07E+00 | 1.795489 | 2.247262 | 0.145985 | 0.291971 | Infection | Intestine |
| Hrh1 | -1.96E+00 | 5.580041 | 2.798658 | 0.107984 | 0.230224 | Healthy Control | Intestine |
| Hrh3 | -2.16E+00 | 1.318372 | 0.088982 | 0.768665 | 0.932538 | Healthy Control | Intestine |
| Hs3st2 | -2.28E+00 | 1.362804 | 0.068836 | 0.795417 | 0.943769 | Healthy Control | Intestine |
| Hsd17b3 | -1.37E+00 | 1.229023 | 0.032546 | 0.859435 | 0.956553 | Healthy Control | Intestine |
| Hsd17b4 | -2.06E+00 | 9.077583 | 43.00862 | 1.36E-06 | 1.45E-05 | Healthy Control | Intestine |
| Hsd17b7 | -1.40E+00 | 6.424434 | 3.2736 | 0.082672 | 0.187348 | Healthy Control | Intestine |
| Hsf1 | -2.35E-01 | 7.686755 | 0.481011 | 0.495269 | 0.727926 | Healthy Control | Intestine |
| Hsf2 | 4.87E-02 | 5.958564 | 0.007296 | 0.932654 | 0.969735 | Infection | Intestine |
| Hspa1b | 4.85E-01 | 6.710864 | 0.049931 | 0.826828 | 0.953685 | Infection | Intestine |
| Hspa5 | 1.29E+00 | 10.63686 | 47.39204 | 1.76E-06 | 1.79E-05 | Infection | Intestine |
| Hspa9 | -1.33E+00 | 9.919335 | 49.52609 | 4.91E-07 | 6.10E-06 | Healthy Control | Intestine |
| Hspb2 | -6.05E+00 | 3.750021 | 1.287837 | 0.270033 | 0.469369 | Healthy Control | Intestine |

| | | | | | | | |
|---|---|---|---|---|---|---|---|
| Hspb3 | -1.37E+00 | 1.228863 | 0.033199 | 0.858049 | 0.956553 | Healthy Control | Intestine |
| Hspd1 | -2.06E+00 | 10.21947 | 132.1387 | 1.03E-10 | 5.49E-09 | Healthy Control | Intestine |
| Hspg2 | -2.40E-02 | 9.20487 | 0.010979 | 0.917507 | 0.969735 | Healthy Control | Intestine |
| Hsph1 | 1.47E-01 | 7.254709 | 0.155051 | 0.697551 | 0.892753 | Infection | Intestine |
| Htr1a | -6.42E-01 | 1.272779 | 0.013453 | 0.908876 | 0.969735 | Healthy Control | Intestine |
| Htr1b | 2.94E+00 | 2.779817 | 8.316383 | 0.008066 | 0.027448 | Infection | Intestine |
| Htr1d | 4.77E+00 | 1.36373 | 3.643591 | 0.06829 | 0.161712 | Infection | Intestine |
| Htr1f | -3.25E+00 | 1.764807 | 0.225854 | 0.638041 | 0.853108 | Healthy Control | Intestine |
| Htr2a | 5.54E+00 | 1.707364 | 14.13263 | 0.00068 | 0.003319 | Infection | Intestine |
| Htr2b | -2.74E+00 | 1.594041 | 0.124957 | 0.725928 | 0.910547 | Healthy Control | Intestine |
| Htr2c | -9.77E-01 | 1.31829 | 0.021938 | 0.88358 | 0.96867 | Healthy Control | Intestine |
| Htr3a | -5.00E+00 | 2.045585 | 0.369229 | 0.548047 | 0.778138 | Healthy Control | Intestine |
| Htr3b | -1.37E+00 | 1.228863 | 0.033199 | 0.858049 | 0.956553 | Healthy Control | Intestine |
| Htr4 | -9.86E+00 | 7.02622 | 27.90059 | 3.72E-05 | 0.000259 | Healthy Control | Intestine |
| Htr7 | 1.46E+00 | 2.697391 | 0.641975 | 0.430533 | 0.662949 | Infection | Intestine |
| Htra2 | -2.69E+00 | 6.06665 | 6.622862 | 0.01707 | 0.051757 | Healthy Control | Intestine |
| Htt | -1.06E+00 | 7.805294 | 11.16537 | 0.002916 | 0.01143 | Healthy Control | Intestine |
| Huwe1 | -2.66E-01 | 8.897029 | 1.97958 | 0.173546 | 0.333374 | Healthy Control | Intestine |
| Ibsp | 4.74E+00 | 1.563221 | 7.83279 | 0.008744 | 0.029337 | Infection | Intestine |
| Icam1 | 6.32E-01 | 6.393433 | 0.718576 | 0.406568 | 0.637769 | Infection | Intestine |
| Ick | -2.36E+00 | 7.871297 | 39.38729 | 1.72E-06 | 1.75E-05 | Healthy Control | Intestine |
| Icos | -6.55E+00 | 3.655001 | 1.076365 | 0.31211 | 0.525537 | Healthy Control | Intestine |
| Icosl | -1.34E+00 | 6.828713 | 2.525509 | 0.125074 | 0.257733 | Healthy Control | Intestine |
| Id1 | -3.06E+00 | 9.455467 | 63.84672 | 4.07E-08 | 7.50E-07 | Healthy Control | Intestine |
| Id2 | -8.10E-01 | 7.449389 | 4.643855 | 0.04218 | 0.108954 | Healthy Control | Intestine |
| Id4 | 8.67E-01 | 6.631468 | 0.589461 | 0.455791 | 0.689541 | Infection | Intestine |
| Idh1 | -3.55E+00 | 9.836614 | 151.8556 | 1.62E-11 | 1.07E-09 | Healthy Control | Intestine |
| Idh2 | -1.30E+00 | 9.325941 | 31.40593 | 1.28E-05 | 0.000102 | Healthy Control | Intestine |
| Idi1 | -2.68E+00 | 9.109609 | 44.98945 | 6.95E-07 | 8.19E-06 | Healthy Control | Intestine |
| Ido1 | -5.20E+00 | 2.540926 | 0.79606 | 0.379919 | 0.607921 | Healthy Control | Intestine |
| Ido2 | -2.73E+00 | 1.445019 | 0.141993 | 0.70917 | 0.897243 | Healthy Control | Intestine |
| Idua | 4.33E-01 | 5.228504 | 0.287105 | 0.597058 | 0.822701 | Infection | Intestine |
| Ifi204 | 7.25E-01 | 5.231215 | 0.805349 | 0.378711 | 0.606665 | Infection | Intestine |
| Ifi208 | -3.20E+00 | 1.702805 | 0.17937 | 0.67472 | 0.879138 | Healthy Control | Intestine |
| Ifi209 | 1.32E+00 | 3.22685 | 0.800385 | 0.379981 | 0.607921 | Infection | Intestine |
| Ifi27 | 7.88E-01 | 7.474225 | 8.333072 | 0.008614 | 0.028996 | Infection | Intestine |
| Ifi44 | -3.40E-01 | 5.027221 | 0.055118 | 0.81647 | 0.952516 | Healthy Control | Intestine |
| Ifih1 | -1.38E+00 | 7.914992 | 9.75729 | 0.004792 | 0.017611 | Healthy Control | Intestine |
| Ifit1 | -7.24E+00 | 4.102147 | 1.489377 | 0.236784 | 0.424245 | Healthy Control | Intestine |
| Ifit3 | 4.21E+00 | 3.902015 | 50.73097 | 4.23E-07 | 5.36E-06 | Infection | Intestine |
| Ifnar1 | -2.49E+00 | 7.388786 | 18.69626 | 0.000226 | 0.001271 | Healthy Control | Intestine |
| Ifnar2 | 1.61E+00 | 5.448574 | 7.110182 | 0.014051 | 0.043769 | Infection | Intestine |
| Ifnb1 | -9.71E-16 | 1.180616 | 0 | 1 | 1 | Healthy Control | Intestine |
| Ifng | -3.07E+00 | 1.668581 | 0.113012 | 0.739133 | 0.919965 | Healthy Control | Intestine |
| Ifngr1 | -1.12E+00 | 10.13329 | 48.02917 | 6.20E-07 | 7.50E-06 | Healthy Control | Intestine |
| Ifngr2 | -2.70E+00 | 8.684652 | 50.15725 | 2.83E-07 | 3.78E-06 | Healthy Control | Intestine |
| Ifnlr1 | -9.30E-01 | 5.292671 | 0.936285 | 0.343091 | 0.564599 | Healthy Control | Intestine |
| Ift122 | 1.21E+00 | 5.782578 | 5.506397 | 0.028308 | 0.078379 | Infection | Intestine |
| Ift27 | 4.42E-01 | 5.609114 | 0.498939 | 0.486969 | 0.721412 | Infection | Intestine |
| Ift88 | 8.15E-01 | 6.751748 | 3.964843 | 0.059082 | 0.143367 | Infection | Intestine |

| | | | | | | | |
|---|---|---|---|---|---|---|---|
| Igdcc4 | -1.65E+00 | 8.193023 | 15.11911 | 0.000773 | 0.00368 | Healthy Control | Intestine |
| Igf1 | 3.31E+00 | 6.20885 | 95.88056 | 1.94E-09 | 5.93E-08 | Infection | Intestine |
| Igf1r | 4.65E-01 | 7.693794 | 3.242386 | 0.085631 | 0.191971 | Infection | Intestine |
| Igf2 | -4.60E-01 | 5.822156 | 0.240279 | 0.628428 | 0.843445 | Healthy Control | Intestine |
| Igf2bp1 | -3.40E+00 | 1.772792 | 0.149633 | 0.70273 | 0.894903 | Healthy Control | Intestine |
| Igf2r | -1.98E+00 | 9.077284 | 42.36991 | 1.44E-06 | 1.51E-05 | Healthy Control | Intestine |
| Igfbp1 | -8.61E-01 | 1.274525 | 0.021868 | 0.88426 | 0.96867 | Healthy Control | Intestine |
| Igfbp2 | -5.98E+00 | 3.1644 | 1.161508 | 0.293338 | 0.500493 | Healthy Control | Intestine |
| Igfbp3 | -8.81E-01 | 10.35636 | 6.888847 | 0.018394 | 0.055236 | Healthy Control | Intestine |
| Igfn1 | -3.47E+00 | 1.976076 | 0.155002 | 0.697684 | 0.892753 | Healthy Control | Intestine |
| Igha | 3.31E-01 | 11.04002 | 0.605463 | 0.450984 | 0.684875 | Infection | Intestine |
| Ighd | -5.57E+00 | 2.97372 | 0.497438 | 0.488649 | 0.722204 | Healthy Control | Intestine |
| Ighm | -6.58E-01 | 7.905896 | 0.454867 | 0.5114 | 0.746375 | Healthy Control | Intestine |
| Igkc | -9.71E-01 | 11.28171 | 3.625788 | 0.080694 | 0.183951 | Healthy Control | Intestine |
| Igll1 | -9.71E-16 | 1.180616 | 0 | 1 | 1 | Healthy Control | Intestine |
| Igsf11 | 9.97E-02 | 3.344836 | 0.004885 | 0.944914 | 0.974503 | Infection | Intestine |
| Igsf21 | -3.41E+00 | 1.915529 | 0.19736 | 0.660634 | 0.868632 | Healthy Control | Intestine |
| Igsf5 | -2.95E+00 | 7.734057 | 30.66848 | 1.03E-05 | 8.52E-05 | Healthy Control | Intestine |
| Igsf8 | -8.13E-01 | 5.340887 | 0.67765 | 0.418585 | 0.650873 | Healthy Control | Intestine |
| Igsf9b | -2.31E+00 | 6.705093 | 8.409703 | 0.007828 | 0.026729 | Healthy Control | Intestine |
| Igtp | 5.57E-01 | 4.812283 | 0.325911 | 0.57341 | 0.80173 | Infection | Intestine |
| Ihh | -9.99E+00 | 7.066697 | 35.25711 | 8.64E-06 | 7.31E-05 | Healthy Control | Intestine |
| Ikbkb | 9.29E-01 | 8.221434 | 16.71834 | 0.000494 | 0.002514 | Infection | Intestine |
| Ikbkg | 4.45E-01 | 7.654267 | 3.324644 | 0.081986 | 0.186131 | Infection | Intestine |
| Ikzf1 | 6.64E-01 | 5.657599 | 0.646642 | 0.429581 | 0.661686 | Infection | Intestine |
| Ikzf2 | 3.14E-01 | 6.893806 | 0.286735 | 0.597564 | 0.822945 | Infection | Intestine |
| Il10 | -4.27E+00 | 2.226619 | 0.263855 | 0.612117 | 0.833585 | Healthy Control | Intestine |
| Il10ra | 2.22E-01 | 6.453361 | 0.107792 | 0.745654 | 0.92349 | Infection | Intestine |
| Il11 | 4.30E+00 | 2.595778 | 31.5747 | 7.09E-06 | 6.10E-05 | Infection | Intestine |
| Il11ra1 | 7.31E-01 | 6.837836 | 3.338417 | 0.081349 | 0.185191 | Infection | Intestine |
| Il12a | -3.02E+00 | 1.631098 | 0.248242 | 0.62191 | 0.839021 | Healthy Control | Intestine |
| Il12b | 1.82E+00 | 2.169267 | 0.490399 | 0.489853 | 0.722937 | Infection | Intestine |
| Il12rb1 | -4.67E+00 | 2.467765 | 0.487505 | 0.49274 | 0.725914 | Healthy Control | Intestine |
| Il12rb2 | -3.93E+00 | 2.134015 | 0.356702 | 0.555973 | 0.784337 | Healthy Control | Intestine |
| Il13 | 2.01E+00 | 2.008269 | 0.913951 | 0.347279 | 0.570217 | Infection | Intestine |
| Il13ra1 | -5.88E-01 | 9.614315 | 5.941739 | 0.023377 | 0.067411 | Healthy Control | Intestine |
| Il13ra2 | 6.86E+00 | 2.744095 | 86.0094 | 5.01E-10 | 1.87E-08 | Infection | Intestine |
| Il15 | 3.82E-01 | 5.749285 | 0.399904 | 0.533402 | 0.765939 | Infection | Intestine |
| Il15ra | -2.24E+00 | 5.611954 | 3.219217 | 0.086108 | 0.192608 | Healthy Control | Intestine |
| Il17a | 4.85E+00 | 1.361464 | 4.751657 | 0.039134 | 0.102141 | Infection | Intestine |
| Il17f | 5.70E+00 | 1.229995 | 6.086704 | 0.031631 | 0.085684 | Infection | Intestine |
| Il17ra | 6.13E-01 | 5.921955 | 0.78085 | 0.386172 | 0.615857 | Infection | Intestine |
| Il17rb | 1.16E+00 | 3.89766 | 0.806747 | 0.377752 | 0.605323 | Infection | Intestine |
| Il17rc | -1.32E+00 | 9.15774 | 20.06685 | 0.000189 | 0.00108 | Healthy Control | Intestine |
| Il18 | -5.53E+00 | 9.027619 | 45.76585 | 8.14E-07 | 9.43E-06 | Healthy Control | Intestine |
| Il18r1 | -2.86E-01 | 3.687715 | 0.021036 | 0.885984 | 0.96867 | Healthy Control | Intestine |
| Il18rap | 2.08E+00 | 4.109717 | 5.486248 | 0.02813 | 0.078126 | Infection | Intestine |
| Il1a | 6.67E+00 | 2.23001 | 110.0451 | 9.97E-12 | 7.56E-10 | Infection | Intestine |
| Il1f10 | -9.77E-01 | 1.31873 | 0.021036 | 0.885979 | 0.96867 | Healthy Control | Intestine |
| Il1f5 | -1.58E+00 | 1.274873 | 0.042519 | 0.839052 | 0.954066 | Healthy Control | Intestine |

| | | | | | | | |
|---|---|---|---|---|---|---|---|
| Il1r1 | 1.53E+00 | 8.010402 | 68.71552 | 3.49E-08 | 6.52E-07 | Infection | Intestine |
| Il1rap | 9.38E-01 | 6.260598 | 5.901379 | 0.023732 | 0.068278 | Infection | Intestine |
| Il1rapl2 | -4.94E-01 | 1.22813 | 0.009389 | 0.924171 | 0.969735 | Healthy Control | Intestine |
| Il1rl1 | 3.65E+00 | 3.84955 | 44.81058 | 1.01E-06 | 1.14E-05 | Infection | Intestine |
| Il1rn | 2.76E+00 | 7.828094 | 72.07018 | 7.65E-08 | 1.27E-06 | Infection | Intestine |
| Il2 | -2.66E+00 | 1.402243 | 0.111769 | 0.741032 | 0.921009 | Healthy Control | Intestine |
| Il21 | -1.55E+00 | 1.44274 | 0.031773 | 0.859744 | 0.956553 | Healthy Control | Intestine |
| Il21r | -6.29E+00 | 3.998573 | 1.397805 | 0.251153 | 0.444841 | Healthy Control | Intestine |
| Il22 | 4.97E+00 | 1.319895 | 4.295583 | 0.052108 | 0.12952 | Infection | Intestine |
| Il22ra1 | -1.05E+01 | 7.625828 | 44.26831 | 1.86E-06 | 1.86E-05 | Healthy Control | Intestine |
| Il22ra2 | -6.13E+00 | 3.780169 | 0.112652 | 0.742352 | 0.92188 | Healthy Control | Intestine |
| Il23a | -2.87E+00 | 1.522139 | 0.135634 | 0.715055 | 0.901639 | Healthy Control | Intestine |
| Il23r | -2.67E+00 | 1.52468 | 0.121097 | 0.730424 | 0.91327 | Healthy Control | Intestine |
| Il25 | 5.33E+00 | 1.275264 | 4.750983 | 0.046013 | 0.117162 | Infection | Intestine |
| Il27 | 3.42E+00 | 1.666774 | 2.080351 | 0.157963 | 0.30986 | Infection | Intestine |
| Il27ra | 2.22E+00 | 3.662272 | 5.796673 | 0.024129 | 0.06914 | Infection | Intestine |
| Il2ra | 8.62E-01 | 3.628558 | 0.354713 | 0.556985 | 0.784985 | Infection | Intestine |
| Il2rb | -1.34E+00 | 4.761145 | 0.149899 | 0.703844 | 0.895866 | Healthy Control | Intestine |
| Il2rg | 1.39E+00 | 6.178828 | 3.178888 | 0.091928 | 0.203086 | Infection | Intestine |
| Il3 | -3.57E-01 | 1.227067 | 0.006351 | 0.937596 | 0.969735 | Healthy Control | Intestine |
| Il31 | -6.42E-01 | 1.273555 | 0.012735 | 0.91133 | 0.969735 | Healthy Control | Intestine |
| Il33 | 5.51E+00 | 6.442263 | 184.1103 | 2.72E-10 | 1.12E-08 | Infection | Intestine |
| Il34 | -1.18E+00 | 7.112377 | 3.302368 | 0.081973 | 0.186131 | Healthy Control | Intestine |
| Il3ra | -3.09E+00 | 1.914577 | 0.237828 | 0.630186 | 0.84554 | Healthy Control | Intestine |
| Il4 | 5.70E+00 | 1.81167 | 41.16398 | 1.34E-07 | 2.02E-06 | Infection | Intestine |
| Il4ra | 2.45E+00 | 6.546322 | 56.31708 | 1.79E-07 | 2.57E-06 | Infection | Intestine |
| Il5 | -2.63E+00 | 1.363598 | 0.155625 | 0.697584 | 0.892753 | Healthy Control | Intestine |
| Il5ra | 1.65E+00 | 1.930703 | 0.628337 | 0.433833 | 0.666066 | Infection | Intestine |
| Il6ra | -3.42E-01 | 6.184808 | 0.231398 | 0.634937 | 0.850547 | Healthy Control | Intestine |
| Il6st | -5.57E-01 | 8.388807 | 2.670058 | 0.116621 | 0.24468 | Healthy Control | Intestine |
| Il7 | -1.29E+00 | 5.808596 | 1.393012 | 0.249741 | 0.443139 | Healthy Control | Intestine |
| Il7r | 5.92E-01 | 4.971 | 0.348249 | 0.560602 | 0.789416 | Infection | Intestine |
| Il9 | -4.95E-01 | 1.228692 | 0.008683 | 0.927069 | 0.969735 | Healthy Control | Intestine |
| Ilf2 | -1.79E+00 | 7.692615 | 17.74319 | 0.000319 | 0.001722 | Healthy Control | Intestine |
| Ilk | 1.45E+00 | 7.227662 | 20.46946 | 0.000171 | 0.00099 | Infection | Intestine |
| Ilkap | 3.68E-01 | 8.326096 | 3.90572 | 0.060924 | 0.147266 | Infection | Intestine |
| Immt | -1.42E+00 | 10.04943 | 64.21328 | 6.18E-08 | 1.08E-06 | Healthy Control | Intestine |
| Impact | 1.32E+00 | 7.241796 | 15.70788 | 0.00067 | 0.003277 | Infection | Intestine |
| Impad1 | -6.36E-01 | 8.390797 | 3.64584 | 0.069403 | 0.163539 | Healthy Control | Intestine |
| Ina | -3.35E+00 | 1.561689 | 0.169098 | 0.683865 | 0.88417 | Healthy Control | Intestine |
| Ing3 | -1.17E+00 | 6.864458 | 5.104915 | 0.033367 | 0.089276 | Healthy Control | Intestine |
| Inha | 4.22E-01 | 3.058339 | 0.073919 | 0.788261 | 0.941774 | Infection | Intestine |
| Inhba | 4.86E+00 | 3.599031 | 92.402 | 2.90E-09 | 7.94E-08 | Infection | Intestine |
| Inhbb | 6.00E+00 | 1.872277 | 23.64858 | 2.58E-05 | 0.000189 | Infection | Intestine |
| Inka2 | -5.07E+00 | 2.720734 | 0.887397 | 0.357462 | 0.582126 | Healthy Control | Intestine |
| Ino80d | 3.44E-01 | 8.814485 | 3.305809 | 0.082812 | 0.187413 | Infection | Intestine |
| Inpp1 | -1.47E+00 | 6.70164 | 5.755487 | 0.024418 | 0.069808 | Healthy Control | Intestine |
| Inpp4b | -1.20E+00 | 7.643041 | 7.282293 | 0.012765 | 0.04052 | Healthy Control | Intestine |
| Inpp5b | 3.89E-02 | 7.382798 | 0.014645 | 0.904778 | 0.969735 | Infection | Intestine |
| Inpp5d | 1.38E+00 | 6.635918 | 11.22264 | 0.002923 | 0.011443 | Infection | Intestine |

| | | | | | | | |
|---|---|---|---|---|---|---|---|
| Inpp5e | 1.27E+00 | 6.048273 | 7.349345 | 0.012744 | 0.04049 | Infection | Intestine |
| Ins1 | -3.57E-01 | 1.228299 | 0.00618 | 0.938438 | 0.969735 | Healthy Control | Intestine |
| Insm1 | 7.54E-01 | 2.948539 | 0.095723 | 0.759864 | 0.928591 | Infection | Intestine |
| Insr | -2.69E+00 | 9.14407 | 69.31527 | 2.50E-08 | 4.88E-07 | Healthy Control | Intestine |
| Insrr | -5.50E-01 | 5.097976 | 0.177335 | 0.67726 | 0.880469 | Healthy Control | Intestine |
| Ints11 | -1.53E+00 | 7.394953 | 11.13856 | 0.002789 | 0.011004 | Healthy Control | Intestine |
| Ints4 | -2.10E-01 | 7.44892 | 0.387474 | 0.540018 | 0.772705 | Healthy Control | Intestine |
| Invs | -6.64E-01 | 5.650208 | 0.773895 | 0.387559 | 0.61689 | Healthy Control | Intestine |
| Ipmk | -7.73E-01 | 9.01172 | 6.696144 | 0.016865 | 0.051198 | Healthy Control | Intestine |
| Ipo9 | 6.81E-01 | 6.174265 | 2.147178 | 0.156835 | 0.308252 | Infection | Intestine |
| Iqgap1 | -1.27E+00 | 10.16919 | 84.7459 | 5.84E-09 | 1.41E-07 | Healthy Control | Intestine |
| Iqgap2 | -1.96E+00 | 8.74458 | 21.81599 | 0.000112 | 0.000686 | Healthy Control | Intestine |
| Iqgap3 | -7.69E+00 | 4.872579 | 4.277979 | 0.051926 | 0.129155 | Healthy Control | Intestine |
| Irak1 | 1.43E-01 | 7.943841 | 0.273404 | 0.606329 | 0.829653 | Infection | Intestine |
| Irak2 | -2.10E+00 | 7.042484 | 12.18335 | 0.001848 | 0.007728 | Healthy Control | Intestine |
| Irak3 | 1.28E+00 | 5.843291 | 9.689424 | 0.00504 | 0.018374 | Infection | Intestine |
| Ireb2 | -4.57E-01 | 8.53134 | 2.933081 | 0.100997 | 0.218586 | Healthy Control | Intestine |
| Irf1 | -3.35E-01 | 9.289791 | 2.876928 | 0.104107 | 0.223629 | Healthy Control | Intestine |
| Irf3 | -5.39E-01 | 8.808107 | 6.50949 | 0.01828 | 0.054962 | Healthy Control | Intestine |
| Irf4 | -1.83E+00 | 6.90813 | 0.661771 | 0.428986 | 0.661421 | Healthy Control | Intestine |
| Irf5 | 1.63E+00 | 6.096388 | 9.034546 | 0.006526 | 0.023038 | Infection | Intestine |
| Irf7 | -1.24E+00 | 4.109963 | 0.506082 | 0.484668 | 0.718888 | Healthy Control | Intestine |
| Irf8 | 5.97E-01 | 8.58211 | 4.549821 | 0.044439 | 0.11385 | Infection | Intestine |
| Irf9 | -8.96E-01 | 8.168432 | 3.662045 | 0.069229 | 0.163473 | Healthy Control | Intestine |
| Irs1 | 1.04E-01 | 6.533196 | 0.034946 | 0.85335 | 0.956553 | Infection | Intestine |
| Irs2 | -7.51E-01 | 7.609358 | 3.145012 | 0.089909 | 0.199383 | Healthy Control | Intestine |
| Irx2 | -2.55E+00 | 1.597321 | 0.130038 | 0.721018 | 0.906205 | Healthy Control | Intestine |
| Irx3 | 5.78E+00 | 1.607147 | 22.34866 | 6.02E-05 | 0.000395 | Infection | Intestine |
| Irx4 | -3.57E-01 | 1.227509 | 0.006273 | 0.937979 | 0.969735 | Healthy Control | Intestine |
| Irx5 | -3.57E-01 | 1.227067 | 0.006351 | 0.937596 | 0.969735 | Healthy Control | Intestine |
| Irx6 | 5.06E+00 | 1.3199 | 4.309684 | 0.051206 | 0.127658 | Infection | Intestine |
| Isca1 | -1.33E+00 | 7.99702 | 14.68948 | 0.000871 | 0.004098 | Healthy Control | Intestine |
| Isg15 | -9.79E+00 | 6.426523 | 5.114782 | 0.039822 | 0.103613 | Healthy Control | Intestine |
| Isg20 | -3.38E+00 | 7.999028 | 30.72777 | 1.05E-05 | 8.56E-05 | Healthy Control | Intestine |
| Isl1 | -7.25E+00 | 4.835864 | 2.890248 | 0.104061 | 0.223629 | Healthy Control | Intestine |
| Isl2 | 2.16E+00 | 3.468132 | 4.203949 | 0.051936 | 0.129155 | Infection | Intestine |
| Itch | -2.59E-01 | 9.092505 | 2.080945 | 0.163379 | 0.318364 | Healthy Control | Intestine |
| Itga1 | 2.36E+00 | 7.613095 | 85.99237 | 5.08E-09 | 1.27E-07 | Infection | Intestine |
| Itga11 | -1.48E+00 | 4.959471 | 0.824568 | 0.373288 | 0.59932 | Healthy Control | Intestine |
| Itga2 | -1.36E+00 | 7.498164 | 5.520054 | 0.027694 | 0.077277 | Healthy Control | Intestine |
| Itga2b | 3.82E+00 | 3.76006 | 31.58532 | 1.25E-05 | 0.0001 | Infection | Intestine |
| Itga3 | -3.28E+00 | 8.513127 | 73.88694 | 8.44E-09 | 1.93E-07 | Healthy Control | Intestine |
| Itga4 | 1.91E+00 | 6.578749 | 21.34727 | 0.000136 | 0.000804 | Infection | Intestine |
| Itga5 | 2.54E+00 | 8.049867 | 129.1401 | 1.25E-10 | 6.40E-09 | Infection | Intestine |
| Itga6 | -3.40E+00 | 7.768656 | 27.16303 | 2.54E-05 | 0.000187 | Healthy Control | Intestine |
| Itga7 | 7.72E-01 | 5.972865 | 1.861298 | 0.186148 | 0.351644 | Infection | Intestine |
| Itga8 | 8.30E-01 | 5.425501 | 0.961948 | 0.336771 | 0.556274 | Infection | Intestine |
| Itga9 | 1.92E+00 | 5.800421 | 18.83383 | 0.000268 | 0.001471 | Infection | Intestine |
| Itgae | -3.21E+00 | 6.539936 | 3.675535 | 0.069885 | 0.164247 | Healthy Control | Intestine |
| Itgal | 5.51E-01 | 5.889605 | 0.636382 | 0.433326 | 0.66602 | Infection | Intestine |

| | | | | | | | |
|---|---|---|---|---|---|---|---|
| Itgam | 3.39E+00 | 6.537702 | 122.8428 | 2.01E-10 | 8.74E-09 | Infection | Intestine |
| Itgav | 3.15E-02 | 7.756989 | 0.008981 | 0.925361 | 0.969735 | Infection | Intestine |
| Itgax | 3.53E+00 | 4.541312 | 39.1782 | 2.80E-06 | 2.67E-05 | Infection | Intestine |
| Itgb1 | 2.19E+00 | 11.39668 | 443.8988 | 5.50E-16 | 1.72E-13 | Infection | Intestine |
| Itgb2 | 4.17E+00 | 6.697222 | 311.637 | 2.13E-14 | 4.24E-12 | Infection | Intestine |
| Itgb3 | 2.42E+00 | 6.285792 | 23.13901 | 8.56E-05 | 0.000542 | Infection | Intestine |
| Itgb4 | -3.77E+00 | 9.817002 | 228.1969 | 1.45E-13 | 2.34E-11 | Healthy Control | Intestine |
| Itgb5 | 8.71E-01 | 7.774463 | 7.170486 | 0.013821 | 0.043162 | Infection | Intestine |
| Itgb6 | -2.71E+00 | 6.813698 | 12.1351 | 0.001983 | 0.008205 | Healthy Control | Intestine |
| Itgb7 | 1.13E+00 | 6.271011 | 1.236395 | 0.283349 | 0.487555 | Infection | Intestine |
| Itgb8 | 3.30E+00 | 3.663602 | 16.36612 | 0.000506 | 0.002557 | Infection | Intestine |
| Itk | -4.23E-01 | 3.773878 | 0.062671 | 0.804628 | 0.947891 | Healthy Control | Intestine |
| Itln1 | 4.55E-01 | 8.32116 | 0.385884 | 0.543908 | 0.775484 | Infection | Intestine |
| Itpa | 3.41E+00 | 3.71703 | 29.79665 | 1.67E-05 | 0.00013 | Infection | Intestine |
| Itpkb | 1.03E+00 | 6.858342 | 8.317708 | 0.008672 | 0.029134 | Infection | Intestine |
| Itpr1 | 1.49E+00 | 6.959942 | 4.966203 | 0.040409 | 0.104866 | Infection | Intestine |
| Itpr2 | -2.76E-02 | 7.014852 | 0.004211 | 0.948842 | 0.977346 | Healthy Control | Intestine |
| Itpr3 | -3.41E+00 | 8.356018 | 50.06164 | 2.67E-07 | 3.59E-06 | Healthy Control | Intestine |
| Itsn1 | 1.09E+00 | 8.597641 | 37.94119 | 3.51E-06 | 3.26E-05 | Infection | Intestine |
| Itsn2 | 7.12E-01 | 8.731742 | 19.19318 | 0.000242 | 0.001353 | Infection | Intestine |
| Ivd | -1.69E+00 | 8.232036 | 13.41669 | 0.001334 | 0.00588 | Healthy Control | Intestine |
| Izumo1r | 2.32E+00 | 1.92385 | 1.086381 | 0.305635 | 0.516547 | Infection | Intestine |
| Jade1 | 8.10E-02 | 6.770413 | 0.024586 | 0.87682 | 0.965968 | Infection | Intestine |
| Jade2 | 5.11E-01 | 6.858626 | 2.259638 | 0.146942 | 0.293649 | Infection | Intestine |
| Jade3 | -2.18E+00 | 5.658314 | 4.156395 | 0.053267 | 0.131812 | Healthy Control | Intestine |
| Jag1 | 8.47E-01 | 7.658917 | 8.670339 | 0.007551 | 0.025996 | Infection | Intestine |
| Jag2 | -8.22E+00 | 5.328277 | 2.701275 | 0.117246 | 0.245374 | Healthy Control | Intestine |
| Jak1 | 1.16E+00 | 9.338677 | 76.60047 | 1.39E-08 | 2.91E-07 | Infection | Intestine |
| Jak2 | 1.94E+00 | 8.691727 | 86.99334 | 4.58E-09 | 1.17E-07 | Infection | Intestine |
| Jak3 | 5.42E-01 | 6.918102 | 2.118551 | 0.159677 | 0.312609 | Infection | Intestine |
| Jam2 | 1.28E+00 | 5.452839 | 2.793941 | 0.108631 | 0.231308 | Infection | Intestine |
| Jam3 | 1.55E+00 | 5.364044 | 5.269113 | 0.031582 | 0.085684 | Infection | Intestine |
| Jaml | -9.33E-01 | 7.210297 | 1.635064 | 0.214663 | 0.392765 | Healthy Control | Intestine |
| Jarid2 | -7.37E-01 | 8.139112 | 7.24513 | 0.013313 | 0.041864 | Healthy Control | Intestine |
| Jazf1 | 5.15E-01 | 3.317853 | 0.090817 | 0.765851 | 0.931224 | Infection | Intestine |
| Jchain | -1.61E+00 | 10.64951 | 5.025944 | 0.044668 | 0.114378 | Healthy Control | Intestine |
| Jmjd6 | 5.93E-01 | 7.231864 | 4.000841 | 0.058011 | 0.141246 | Infection | Intestine |
| Jph2 | 1.49E+00 | 6.201928 | 6.58898 | 0.017649 | 0.053416 | Infection | Intestine |
| Jpt1 | -1.76E+00 | 10.06118 | 57.43829 | 1.50E-07 | 2.21E-06 | Healthy Control | Intestine |
| Jup | -2.95E+00 | 10.13242 | 140.7959 | 4.66E-11 | 2.65E-09 | Healthy Control | Intestine |
| Kansl2 | -1.42E+00 | 8.193421 | 16.53321 | 0.000498 | 0.002528 | Healthy Control | Intestine |
| Kat14 | -1.04E-01 | 8.409546 | 0.168002 | 0.685897 | 0.885076 | Healthy Control | Intestine |
| Kat2a | -1.55E-01 | 7.822958 | 0.303257 | 0.587421 | 0.813227 | Healthy Control | Intestine |
| Kat2b | -1.06E+00 | 7.125926 | 4.269677 | 0.05 | 0.124962 | Healthy Control | Intestine |
| Kat5 | -8.40E-01 | 7.780291 | 5.214787 | 0.032301 | 0.086934 | Healthy Control | Intestine |
| Kat6a | -4.31E-01 | 8.351791 | 2.578798 | 0.122676 | 0.253872 | Healthy Control | Intestine |
| Kat6b | 1.13E+00 | 6.335884 | 5.35853 | 0.030376 | 0.082978 | Infection | Intestine |
| Kat8 | -6.71E-01 | 7.708631 | 2.851074 | 0.105372 | 0.225811 | Healthy Control | Intestine |
| Katna1 | -9.24E-01 | 7.528759 | 6.984059 | 0.014713 | 0.045521 | Healthy Control | Intestine |
| Katnal2 | 1.22E+00 | 2.884319 | 0.618031 | 0.43957 | 0.672726 | Infection | Intestine |

| | | | | | | | |
|---|---|---|---|---|---|---|---|
| Kcna10 | -3.57E-01 | 1.227417 | 0.006288 | 0.937908 | 0.969735 | Healthy Control | Intestine |
| Kcna4 | -2.92E+00 | 1.525094 | 0.145767 | 0.705395 | 0.896371 | Healthy Control | Intestine |
| Kcnab2 | -4.05E-01 | 4.797574 | 0.105713 | 0.747886 | 0.924698 | Healthy Control | Intestine |
| Kcnab3 | 1.55E+00 | 3.039633 | 1.699367 | 0.204977 | 0.379059 | Infection | Intestine |
| Kcnc1 | 2.04E+00 | 2.096527 | 0.696794 | 0.410043 | 0.642263 | Infection | Intestine |
| Kcnc2 | -2.44E+00 | 1.444577 | 0.094891 | 0.760095 | 0.928591 | Healthy Control | Intestine |
| Kcnc4 | -2.82E+00 | 1.63048 | 0.165961 | 0.686349 | 0.885076 | Healthy Control | Intestine |
| Kcnf1 | -1.52E+00 | 4.931185 | 1.113777 | 0.302553 | 0.51307 | Healthy Control | Intestine |
| Kcng1 | -3.69E+00 | 1.94834 | 0.491028 | 0.489667 | 0.722876 | Healthy Control | Intestine |
| Kcng2 | -7.00E+00 | 4.166733 | 0.871459 | 0.364212 | 0.588714 | Healthy Control | Intestine |
| Kcnh1 | -6.08E+00 | 3.40534 | 1.358382 | 0.257805 | 0.453216 | Healthy Control | Intestine |
| Kcnh2 | 1.11E+00 | 3.877571 | 0.696247 | 0.412051 | 0.644199 | Infection | Intestine |
| Kcnh5 | -6.42E-01 | 1.273434 | 0.012853 | 0.910919 | 0.969735 | Healthy Control | Intestine |
| Kcnh8 | -6.40E+00 | 3.765748 | 1.624301 | 0.217334 | 0.396492 | Healthy Control | Intestine |
| Kcnj10 | -1.38E+00 | 4.969652 | 0.99693 | 0.328539 | 0.545676 | Healthy Control | Intestine |
| Kcnj11 | -4.44E+00 | 2.138432 | 0.48811 | 0.491439 | 0.724424 | Healthy Control | Intestine |
| Kcnj2 | 2.64E+00 | 3.401559 | 5.32748 | 0.029888 | 0.081803 | Infection | Intestine |
| Kcnj3 | -2.81E+00 | 1.483809 | 0.104142 | 0.749076 | 0.924753 | Healthy Control | Intestine |
| Kcnj5 | -5.29E+00 | 2.79795 | 1.25599 | 0.275032 | 0.47669 | Healthy Control | Intestine |
| Kcnj6 | -9.77E-01 | 1.318419 | 0.021469 | 0.884821 | 0.96867 | Healthy Control | Intestine |
| Kcnj8 | 1.12E+00 | 4.865289 | 1.94562 | 0.17609 | 0.337095 | Infection | Intestine |
| Kcnj9 | -2.13E+00 | 1.483101 | 0.10645 | 0.746466 | 0.924073 | Healthy Control | Intestine |
| Kcnk10 | -5.03E+00 | 2.710333 | 0.629979 | 0.436205 | 0.668803 | Healthy Control | Intestine |
| Kcnk18 | -9.71E-16 | 1.180616 | 0 | 1 | 1 | Healthy Control | Intestine |
| Kcnk2 | 2.36E+00 | 4.186403 | 9.249922 | 0.00539 | 0.019511 | Infection | Intestine |
| Kcnk5 | -3.43E+00 | 8.232866 | 41.36687 | 1.33E-06 | 1.43E-05 | Healthy Control | Intestine |
| Kcnma1 | -2.05E+00 | 7.643464 | 13.63356 | 0.00113 | 0.005109 | Healthy Control | Intestine |
| Kcnmb2 | 2.56E+00 | 2.026799 | 1.98035 | 0.172433 | 0.331617 | Infection | Intestine |
| Kcnn2 | 2.29E+00 | 1.951674 | 1.111692 | 0.300229 | 0.509645 | Infection | Intestine |
| Kcnn4 | -3.12E+00 | 8.284497 | 28.07934 | 1.86E-05 | 0.000143 | Healthy Control | Intestine |
| Kcnq1 | -2.34E+00 | 9.610527 | 41.21736 | 1.91E-06 | 1.90E-05 | Healthy Control | Intestine |
| Kcnq3 | -4.39E+00 | 1.867123 | 0.416852 | 0.523225 | 0.7574 | Healthy Control | Intestine |
| Kcns2 | -9.71E-16 | 1.180616 | 0 | 1 | 1 | Healthy Control | Intestine |
| Kcns3 | 8.80E-01 | 3.837032 | 0.407175 | 0.529282 | 0.763504 | Infection | Intestine |
| Kcnt2 | 2.88E+00 | 4.584804 | 14.13345 | 0.001086 | 0.004955 | Infection | Intestine |
| Kctd15 | 6.54E-02 | 4.190159 | 0.003858 | 0.950995 | 0.978831 | Infection | Intestine |
| Kdf1 | -8.14E+00 | 5.395529 | 6.897753 | 0.016261 | 0.049607 | Healthy Control | Intestine |
| Kdm1a | -1.94E+00 | 7.555166 | 15.2095 | 0.000684 | 0.003334 | Healthy Control | Intestine |
| Kdm1b | -1.03E+00 | 7.593207 | 8.140829 | 0.008997 | 0.030068 | Healthy Control | Intestine |
| Kdm2a | -2.19E-01 | 8.205903 | 0.901636 | 0.352721 | 0.576845 | Healthy Control | Intestine |
| Kdm2b | -1.20E+00 | 6.19758 | 3.735484 | 0.064976 | 0.155039 | Healthy Control | Intestine |
| Kdm4a | -2.82E-01 | 8.507866 | 1.457258 | 0.240315 | 0.429188 | Healthy Control | Intestine |
| Kdm4b | -1.88E+00 | 7.174769 | 10.60952 | 0.003319 | 0.012799 | Healthy Control | Intestine |
| Kdm4c | 5.64E-01 | 6.943535 | 2.424388 | 0.133733 | 0.27192 | Infection | Intestine |
| Kdm5a | 6.71E-01 | 8.433192 | 11.51694 | 0.002637 | 0.010493 | Infection | Intestine |
| Kdm5b | -6.76E-01 | 9.336602 | 5.492109 | 0.02865 | 0.079108 | Healthy Control | Intestine |
| Kdm5c | 4.38E-01 | 9.090363 | 7.91481 | 0.01018 | 0.033438 | Infection | Intestine |
| Kdm6a | 2.88E-01 | 7.661319 | 0.860143 | 0.363863 | 0.588676 | Infection | Intestine |
| Kdm6b | 1.25E+00 | 8.324518 | 33.79157 | 7.83E-06 | 6.70E-05 | Infection | Intestine |
| Kdm7a | -3.72E-01 | 7.529138 | 1.069309 | 0.312257 | 0.525608 | Healthy Control | Intestine |

| | | | | | | | |
|---|---|---|---|---|---|---|---|
| Kdm8 | -7.20E-01 | 5.030321 | 0.440821 | 0.513057 | 0.748008 | Healthy Control | Intestine |
| Kdr | -9.74E+00 | 6.671199 | 5.091313 | 0.040347 | 0.104813 | Healthy Control | Intestine |
| Keap1 | -1.74E+00 | 7.072044 | 7.303153 | 0.012384 | 0.039564 | Healthy Control | Intestine |
| Khdrbs1 | -5.20E-01 | 9.13316 | 7.799647 | 0.010672 | 0.03469 | Healthy Control | Intestine |
| Khdrbs2 | -3.39E+00 | 1.731886 | 0.212521 | 0.647991 | 0.859523 | Healthy Control | Intestine |
| Khsrp | -1.60E+00 | 8.518748 | 31.47024 | 1.15E-05 | 9.34E-05 | Healthy Control | Intestine |
| Kif19a | 3.90E-01 | 3.314805 | 0.061743 | 0.806006 | 0.948042 | Infection | Intestine |
| Kif1b | -7.36E-01 | 8.772692 | 10.61738 | 0.003628 | 0.013822 | Healthy Control | Intestine |
| Kif1c | -1.54E+00 | 10.29932 | 63.19275 | 7.06E-08 | 1.20E-06 | Healthy Control | Intestine |
| Kif20a | -2.29E-01 | 6.597944 | 0.151658 | 0.700609 | 0.89357 | Healthy Control | Intestine |
| Kif20b | 1.65E-01 | 5.399081 | 0.042381 | 0.838632 | 0.954066 | Infection | Intestine |
| Kif21b | 1.15E+00 | 7.421981 | 14.09442 | 0.001112 | 0.00505 | Infection | Intestine |
| Kif24 | 1.33E+00 | 4.593902 | 3.213564 | 0.085872 | 0.192338 | Infection | Intestine |
| Kif2b | 6.70E+00 | 1.322684 | 12.4169 | 0.003235 | 0.012486 | Infection | Intestine |
| Kif2c | -6.89E-01 | 5.259115 | 0.290522 | 0.59461 | 0.820046 | Healthy Control | Intestine |
| Kif3a | 2.34E+00 | 4.339551 | 14.10054 | 0.001057 | 0.004852 | Infection | Intestine |
| Kif3b | -5.73E-02 | 7.872429 | 0.052472 | 0.820939 | 0.953035 | Healthy Control | Intestine |
| Kif4 | -2.30E+00 | 5.732416 | 2.860933 | 0.104344 | 0.223895 | Healthy Control | Intestine |
| Kif5a | -6.51E+00 | 3.962454 | 1.534193 | 0.230036 | 0.416328 | Healthy Control | Intestine |
| Kif5b | -6.83E-01 | 9.990003 | 18.12817 | 0.000327 | 0.001761 | Healthy Control | Intestine |
| Kif5c | 3.45E-01 | 3.913783 | 0.077112 | 0.783719 | 0.940206 | Infection | Intestine |
| Kifap3 | -3.98E-01 | 5.985661 | 0.509582 | 0.482218 | 0.716316 | Healthy Control | Intestine |
| Kirrel | 2.01E+00 | 6.103772 | 19.87758 | 0.000201 | 0.001147 | Infection | Intestine |
| Kiss1 | -1.65E+00 | 1.318052 | 0.039297 | 0.844667 | 0.954066 | Healthy Control | Intestine |
| Kiss1r | -2.72E-01 | 4.06485 | 0.038576 | 0.846033 | 0.954066 | Healthy Control | Intestine |
| Kit | -3.54E+00 | 9.388234 | 45.31219 | 1.03E-06 | 1.15E-05 | Healthy Control | Intestine |
| Kitl | -1.83E+00 | 9.089882 | 55.2117 | 1.89E-07 | 2.69E-06 | Healthy Control | Intestine |
| Kl | -3.48E+00 | 1.952989 | 0.239012 | 0.628456 | 0.843445 | Healthy Control | Intestine |
| Klb | -7.70E+00 | 5.326341 | 4.255838 | 0.0525 | 0.130236 | Healthy Control | Intestine |
| Klc1 | -1.48E-01 | 9.040565 | 0.498638 | 0.487587 | 0.721551 | Healthy Control | Intestine |
| Klc2 | -1.35E-01 | 7.626291 | 0.219206 | 0.644277 | 0.857095 | Healthy Control | Intestine |
| Klc3 | -1.27E+00 | 4.596429 | 0.68724 | 0.41586 | 0.648534 | Healthy Control | Intestine |
| Klf1 | -2.87E+00 | 1.825985 | 0.177775 | 0.676744 | 0.8804 | Healthy Control | Intestine |
| Klf10 | 3.91E-01 | 6.178902 | 0.25315 | 0.619577 | 0.838379 | Infection | Intestine |
| Klf11 | 6.64E-01 | 6.476939 | 1.200377 | 0.285137 | 0.489505 | Infection | Intestine |
| Klf12 | -6.06E-01 | 6.97267 | 1.272548 | 0.270712 | 0.470386 | Healthy Control | Intestine |
| Klf15 | -5.94E+00 | 3.264039 | 1.317149 | 0.264932 | 0.463237 | Healthy Control | Intestine |
| Klf17 | -3.81E+00 | 2.224309 | 0.299623 | 0.589151 | 0.814946 | Healthy Control | Intestine |
| Klf4 | -4.49E+00 | 10.07018 | 73.87387 | 3.19E-08 | 6.02E-07 | Healthy Control | Intestine |
| Klf5 | -4.31E+00 | 12.37383 | 508.7353 | 1.33E-16 | 6.07E-14 | Healthy Control | Intestine |
| Klhl18 | 2.14E+00 | 6.730731 | 50.53298 | 4.20E-07 | 5.34E-06 | Infection | Intestine |
| Klhl21 | 6.91E-01 | 6.896219 | 3.829724 | 0.063166 | 0.151586 | Infection | Intestine |
| Klhl22 | -4.79E-01 | 6.534209 | 0.587659 | 0.450972 | 0.684875 | Healthy Control | Intestine |
| Klhl3 | 2.79E+00 | 3.966007 | 12.73925 | 0.001685 | 0.007132 | Infection | Intestine |
| Klhl36 | -1.26E+00 | 6.270874 | 2.725959 | 0.111782 | 0.23651 | Healthy Control | Intestine |
| Klhl41 | 2.12E+00 | 2.32014 | 1.2922 | 0.265819 | 0.464007 | Infection | Intestine |
| Klk1b4 | -2.56E+00 | 1.661688 | 0.097083 | 0.757608 | 0.928591 | Healthy Control | Intestine |
| Klk5 | -1.58E+00 | 1.274099 | 0.047248 | 0.830479 | 0.954066 | Healthy Control | Intestine |
| Klk6 | -9.71E-16 | 1.180616 | 0 | 1 | 1 | Healthy Control | Intestine |
| Klra8 | 3.01E+00 | 1.670202 | 2.158434 | 0.150473 | 0.299117 | Infection | Intestine |

| | | | | | | | |
|---|---|---|---|---|---|---|---|
| Klrb1 | -1.65E+00 | 1.31693 | 0.054107 | 0.818198 | 0.952901 | Healthy Control | Intestine |
| Klrb1c | -2.97E+00 | 1.562495 | 0.185453 | 0.669551 | 0.876052 | Healthy Control | Intestine |
| Klrc1 | -4.59E+00 | 2.527916 | 0.332146 | 0.570456 | 0.799164 | Healthy Control | Intestine |
| Klrd1 | -3.45E+00 | 1.76632 | 0.153022 | 0.698394 | 0.893044 | Healthy Control | Intestine |
| Klrg1 | -6.16E+00 | 3.362502 | 1.028463 | 0.322885 | 0.538967 | Healthy Control | Intestine |
| Klrk1 | -5.58E-01 | 3.999087 | 0.099973 | 0.754771 | 0.926983 | Healthy Control | Intestine |
| Kmt2a | 1.26E+00 | 8.072492 | 40.34705 | 2.26E-06 | 2.21E-05 | Infection | Intestine |
| Kmt2b | 1.26E+00 | 8.019301 | 40.28998 | 2.28E-06 | 2.22E-05 | Infection | Intestine |
| Kmt2c | -6.25E-01 | 8.100825 | 4.883471 | 0.037868 | 0.099354 | Healthy Control | Intestine |
| Kmt2d | 5.98E-01 | 9.047395 | 15.3204 | 0.000754 | 0.003594 | Infection | Intestine |
| Kmt2e | 6.33E-01 | 9.115659 | 18.37333 | 0.000305 | 0.001656 | Infection | Intestine |
| Kmt5b | 5.72E-01 | 8.228671 | 8.114137 | 0.009403 | 0.031194 | Infection | Intestine |
| Kmt5c | -4.55E-01 | 7.533989 | 2.337076 | 0.140442 | 0.283147 | Healthy Control | Intestine |
| Kndc1 | 4.33E+00 | 1.362935 | 3.534117 | 0.072724 | 0.169486 | Infection | Intestine |
| Knl1 | -1.83E+00 | 5.174977 | 1.654415 | 0.211398 | 0.388067 | Healthy Control | Intestine |
| Knstrn | 4.92E-01 | 7.479442 | 1.345292 | 0.258667 | 0.454346 | Infection | Intestine |
| Kpna2 | -7.08E-01 | 7.054856 | 1.760802 | 0.1977 | 0.368209 | Healthy Control | Intestine |
| Kpnb1 | -5.96E-01 | 10.22147 | 21.68381 | 0.000124 | 0.000743 | Healthy Control | Intestine |
| Kras | 5.66E-01 | 9.364692 | 8.635389 | 0.007647 | 0.02629 | Infection | Intestine |
| Kremen1 | -2.06E+00 | 5.459436 | 2.487295 | 0.12869 | 0.263701 | Healthy Control | Intestine |
| Kremen2 | -6.73E+00 | 4.337179 | 2.341588 | 0.141838 | 0.285502 | Healthy Control | Intestine |
| Krit1 | -3.28E-01 | 6.61981 | 0.373857 | 0.54689 | 0.777534 | Healthy Control | Intestine |
| Krt2 | -3.57E-01 | 1.228299 | 0.00618 | 0.938438 | 0.969735 | Healthy Control | Intestine |
| Krt20 | -4.16E+00 | 10.79128 | 47.38259 | 5.47E-06 | 4.82E-05 | Healthy Control | Intestine |
| Krt28 | 4.15E+00 | 1.405975 | 3.342181 | 0.078553 | 0.180053 | Infection | Intestine |
| Krt40 | -9.71E-16 | 1.180616 | 0 | 1 | 1 | Healthy Control | Intestine |
| Krt71 | -1.91E+00 | 1.522092 | 0.074183 | 0.787326 | 0.941331 | Healthy Control | Intestine |
| Krt76 | -9.71E-16 | 1.180616 | 0 | 1 | 1 | Healthy Control | Intestine |
| Krt78 | -3.10E+00 | 1.696777 | 0.164799 | 0.688304 | 0.887139 | Healthy Control | Intestine |
| Krt79 | -6.42E-01 | 1.273598 | 0.012729 | 0.91135 | 0.969735 | Healthy Control | Intestine |
| Krt82 | -9.71E-16 | 1.180616 | 0 | 1 | 1 | Healthy Control | Intestine |
| Ksr2 | -4.75E+00 | 2.449283 | 0.796932 | 0.381706 | 0.610486 | Healthy Control | Intestine |
| Kyat1 | -1.14E+00 | 6.255328 | 2.56641 | 0.122291 | 0.253179 | Healthy Control | Intestine |
| L1cam | -3.24E+00 | 8.635765 | 77.46603 | 5.44E-09 | 1.32E-07 | Healthy Control | Intestine |
| L3hypdh | -5.62E+00 | 3.140921 | 1.293367 | 0.26911 | 0.468415 | Healthy Control | Intestine |
| L3mbtl2 | -7.76E-01 | 6.656598 | 1.637902 | 0.213096 | 0.390467 | Healthy Control | Intestine |
| Lacc1 | 1.11E+00 | 4.204596 | 1.204425 | 0.283161 | 0.48745 | Infection | Intestine |
| Lag3 | 1.32E+00 | 3.889825 | 1.361538 | 0.254661 | 0.449202 | Infection | Intestine |
| Lama1 | 2.02E+00 | 3.263883 | 2.900108 | 0.101264 | 0.219069 | Infection | Intestine |
| Lama2 | 1.50E+00 | 7.931729 | 11.61715 | 0.003469 | 0.013309 | Infection | Intestine |
| Lama3 | -8.83E+00 | 5.751644 | 7.541965 | 0.012521 | 0.039898 | Healthy Control | Intestine |
| Lama4 | 3.37E-01 | 7.743813 | 0.432608 | 0.518907 | 0.754254 | Infection | Intestine |
| Lama5 | -3.79E-01 | 7.00223 | 0.726029 | 0.403162 | 0.633866 | Healthy Control | Intestine |
| Lamb1 | 2.27E+00 | 8.392819 | 89.76634 | 4.76E-09 | 1.21E-07 | Infection | Intestine |
| Lamb2 | 1.97E-01 | 8.16391 | 0.22541 | 0.640173 | 0.854589 | Infection | Intestine |
| Lamb3 | -4.42E+00 | 8.788128 | 45.38591 | 5.86E-07 | 7.14E-06 | Healthy Control | Intestine |
| Lamc1 | 9.70E-01 | 9.483279 | 18.53724 | 0.000331 | 0.001774 | Infection | Intestine |
| Lamp1 | -1.25E+00 | 10.82186 | 93.80845 | 2.35E-09 | 6.80E-08 | Healthy Control | Intestine |
| Lamp2 | 1.70E+00 | 9.349722 | 184.2314 | 4.12E-12 | 3.61E-10 | Infection | Intestine |
| Lamp3 | -1.92E+00 | 1.59086 | 0.043659 | 0.836661 | 0.954066 | Healthy Control | Intestine |

| | | | | | | | |
|---|---|---|---|---|---|---|---|
| Lamp5 | -9.71E-16 | 1.180616 | 0 | 1 | 1 | Healthy Control | Intestine |
| Lamtor1 | -1.43E-01 | 8.39257 | 0.369419 | 0.549609 | 0.778975 | Healthy Control | Intestine |
| Lamtor2 | -1.40E+00 | 9.707776 | 44.33408 | 1.13E-06 | 1.25E-05 | Healthy Control | Intestine |
| Lamtor3 | 1.35E+00 | 7.237525 | 23.04136 | 8.77E-05 | 0.000552 | Infection | Intestine |
| Lamtor4 | -1.44E+00 | 9.917082 | 33.22995 | 8.74E-06 | 7.35E-05 | Healthy Control | Intestine |
| Lamtor5 | -2.17E+00 | 7.519976 | 18.01113 | 0.000284 | 0.001548 | Healthy Control | Intestine |
| Lancl2 | 3.01E-02 | 6.121584 | 0.003136 | 0.955824 | 0.981696 | Infection | Intestine |
| Laptm4b | -4.00E+00 | 9.070365 | 126.3581 | 3.32E-11 | 2.02E-09 | Healthy Control | Intestine |
| Large1 | 1.60E+00 | 5.938241 | 12.31774 | 0.001999 | 0.008251 | Infection | Intestine |
| Large2 | -2.65E+00 | 5.822952 | 5.518677 | 0.02816 | 0.078126 | Healthy Control | Intestine |
| Lasp1 | 7.00E-01 | 9.178871 | 17.7629 | 0.000363 | 0.001928 | Infection | Intestine |
| Lat | 7.95E-01 | 4.815317 | 0.463626 | 0.502434 | 0.736079 | Infection | Intestine |
| Lats1 | -1.13E+00 | 8.571388 | 24.05521 | 6.70E-05 | 0.000436 | Healthy Control | Intestine |
| Lats2 | -9.06E-02 | 6.168399 | 0.019905 | 0.889027 | 0.969599 | Healthy Control | Intestine |
| Lbr | -1.82E+00 | 8.079531 | 29.45006 | 1.69E-05 | 0.000131 | Healthy Control | Intestine |
| Lbx1 | -1.52E+00 | 1.275203 | 0.037966 | 0.847784 | 0.954066 | Healthy Control | Intestine |
| Lck | -1.67E+00 | 5.023589 | 1.003713 | 0.327018 | 0.544104 | Healthy Control | Intestine |
| Lcp2 | 2.42E+00 | 6.55122 | 49.05065 | 5.29E-07 | 6.53E-06 | Infection | Intestine |
| Ldah | -1.56E-01 | 8.215461 | 0.282853 | 0.600207 | 0.824505 | Healthy Control | Intestine |
| Ldb1 | 3.41E-02 | 8.907749 | 0.032456 | 0.858696 | 0.956553 | Infection | Intestine |
| Ldb2 | 2.85E-01 | 4.980172 | 0.061128 | 0.806781 | 0.948508 | Infection | Intestine |
| Ldha | -3.61E+00 | 10.56778 | 123.1039 | 1.64E-10 | 7.61E-09 | Healthy Control | Intestine |
| Ldhb | -4.21E-01 | 5.996032 | 0.409926 | 0.528094 | 0.762294 | Healthy Control | Intestine |
| Ldlr | -2.55E+00 | 7.987404 | 24.3862 | 4.76E-05 | 0.000321 | Healthy Control | Intestine |
| Ldlrad4 | 9.86E-01 | 6.288919 | 3.483579 | 0.075423 | 0.174329 | Infection | Intestine |
| Lect2 | 4.84E+00 | 1.362372 | 4.414994 | 0.046119 | 0.117176 | Infection | Intestine |
| Lef1 | -6.11E+00 | 3.234028 | 1.555165 | 0.22671 | 0.411051 | Healthy Control | Intestine |
| Lefty1 | -4.74E-01 | 4.015832 | 0.117912 | 0.734511 | 0.915806 | Healthy Control | Intestine |
| Lefty2 | -2.88E+00 | 1.362623 | 0.145164 | 0.708506 | 0.897011 | Healthy Control | Intestine |
| Leo1 | -6.17E-01 | 6.972819 | 1.652382 | 0.211606 | 0.388307 | Healthy Control | Intestine |
| Lep | -9.71E-16 | 1.180616 | 0 | 1 | 1 | Healthy Control | Intestine |
| Lepr | 7.84E-02 | 7.142754 | 0.007259 | 0.933231 | 0.969735 | Infection | Intestine |
| Leprot | -9.83E-01 | 8.625188 | 14.18405 | 0.001071 | 0.004897 | Healthy Control | Intestine |
| Letmd1 | -1.63E+00 | 6.970681 | 10.1829 | 0.003919 | 0.014818 | Healthy Control | Intestine |
| Lfng | -2.65E-01 | 6.982788 | 0.267529 | 0.609989 | 0.83218 | Healthy Control | Intestine |
| Lgals9 | -2.25E+00 | 9.222373 | 25.58086 | 6.02E-05 | 0.000395 | Healthy Control | Intestine |
| Lgi3 | -3.43E+00 | 2.125318 | 0.266329 | 0.610804 | 0.832929 | Healthy Control | Intestine |
| Lgr4 | -2.37E+00 | 9.096673 | 48.11893 | 4.41E-07 | 5.57E-06 | Healthy Control | Intestine |
| Lgr5 | -8.22E+00 | 5.618518 | 6.327903 | 0.020635 | 0.060728 | Healthy Control | Intestine |
| Lgr6 | 4.53E+00 | 1.40414 | 3.537373 | 0.070584 | 0.165423 | Infection | Intestine |
| Lhb | -9.76E-01 | 1.317411 | 0.023087 | 0.880593 | 0.96842 | Healthy Control | Intestine |
| Lhcgr | 4.73E+00 | 1.44612 | 3.66116 | 0.067616 | 0.160495 | Infection | Intestine |
| Lhfpl5 | -7.56E-01 | 1.274081 | 0.014574 | 0.905288 | 0.969735 | Healthy Control | Intestine |
| Lhx1 | -6.42E-01 | 1.273689 | 0.012641 | 0.911654 | 0.969735 | Healthy Control | Intestine |
| Lhx2 | -3.57E-01 | 1.228299 | 0.00618 | 0.938438 | 0.969735 | Healthy Control | Intestine |
| Lhx3 | -9.76E-01 | 1.317411 | 0.023087 | 0.880593 | 0.96842 | Healthy Control | Intestine |
| Lhx4 | -2.09E+00 | 1.444926 | 0.08175 | 0.776822 | 0.935926 | Healthy Control | Intestine |
| Lhx5 | 5.70E+00 | 1.229995 | 6.086704 | 0.031631 | 0.085684 | Infection | Intestine |
| Lhx6 | 9.93E-01 | 2.428232 | 0.151803 | 0.700412 | 0.89357 | Infection | Intestine |
| Lhx8 | -3.57E-01 | 1.228299 | 0.00618 | 0.938438 | 0.969735 | Healthy Control | Intestine |

| | | | | | | | |
|---|---|---|---|---|---|---|---|
| Lhx9 | 4.53E+00 | 1.704061 | 8.225267 | 0.006842 | 0.023917 | Infection | Intestine |
| Lif | 3.46E+00 | 3.996959 | 46.3088 | 7.83E-07 | 9.12E-06 | Infection | Intestine |
| Lifr | -1.26E+00 | 7.337812 | 3.530393 | 0.072793 | 0.169568 | Healthy Control | Intestine |
| Lig4 | -2.33E+00 | 5.57769 | 4.387597 | 0.047735 | 0.120323 | Healthy Control | Intestine |
| Lilra5 | -1.79E+00 | 5.384473 | 1.036986 | 0.319109 | 0.53368 | Healthy Control | Intestine |
| Lima1 | -1.69E+00 | 10.89858 | 88.45008 | 4.01E-09 | 1.03E-07 | Healthy Control | Intestine |
| Limd1 | -1.02E+00 | 8.424948 | 16.75125 | 0.000485 | 0.002473 | Healthy Control | Intestine |
| Limk1 | 1.79E+00 | 5.720118 | 14.45051 | 0.000982 | 0.004553 | Infection | Intestine |
| Limk2 | -1.65E+00 | 9.550984 | 41.71996 | 1.72E-06 | 1.75E-05 | Healthy Control | Intestine |
| Lin28a | -1.49E+00 | 1.44069 | 0.042708 | 0.837815 | 0.954066 | Healthy Control | Intestine |
| Lin28b | -9.71E-16 | 1.180616 | 0 | 1 | 1 | Healthy Control | Intestine |
| Lin7c | -1.73E+00 | 9.563656 | 89.25894 | 3.52E-09 | 9.24E-08 | Healthy Control | Intestine |
| Lingo1 | 2.37E+00 | 2.228173 | 1.280231 | 0.267417 | 0.466279 | Infection | Intestine |
| Lipe | -1.95E+00 | 6.330814 | 4.92638 | 0.035991 | 0.095177 | Healthy Control | Intestine |
| Lipg | 1.62E+00 | 8.452473 | 25.18845 | 8.50E-05 | 0.000539 | Infection | Intestine |
| Lipm | 5.58E+00 | 1.322434 | 9.473288 | 0.008234 | 0.027928 | Infection | Intestine |
| Litaf | 4.58E-01 | 9.192459 | 5.243209 | 0.032086 | 0.086589 | Infection | Intestine |
| Llgl2 | -3.63E+00 | 8.898868 | 87.88979 | 1.60E-09 | 4.97E-08 | Healthy Control | Intestine |
| Lmna | -3.20E-01 | 10.47902 | 6.469848 | 0.018587 | 0.055682 | Healthy Control | Intestine |
| Lmnb1 | 1.11E-01 | 8.442964 | 0.192691 | 0.665011 | 0.871934 | Infection | Intestine |
| Lmnb2 | -1.32E+00 | 7.064015 | 5.343946 | 0.029796 | 0.08164 | Healthy Control | Intestine |
| Lmo2 | 8.62E-01 | 5.018168 | 0.904668 | 0.351188 | 0.575088 | Infection | Intestine |
| Lmo4 | 6.41E-01 | 8.736734 | 13.04945 | 0.001562 | 0.006701 | Infection | Intestine |
| Lmx1a | 1.28E+00 | 2.477796 | 0.311045 | 0.582448 | 0.808579 | Infection | Intestine |
| Lmx1b | -2.76E+00 | 1.561884 | 0.138741 | 0.712379 | 0.900306 | Healthy Control | Intestine |
| Lnpk | -3.94E-01 | 6.717003 | 0.659122 | 0.425113 | 0.657841 | Healthy Control | Intestine |
| Lox | 1.66E+00 | 5.531688 | 7.574665 | 0.01158 | 0.037303 | Infection | Intestine |
| Loxl1 | 2.91E+00 | 5.219655 | 23.30641 | 8.21E-05 | 0.000522 | Infection | Intestine |
| Lpar1 | 6.36E-01 | 6.722735 | 2.10647 | 0.16084 | 0.314395 | Infection | Intestine |
| Lpar2 | -6.37E+00 | 3.723311 | 2.476502 | 0.131492 | 0.268017 | Healthy Control | Intestine |
| Lpar5 | -8.17E+00 | 5.525541 | 6.209997 | 0.021698 | 0.0633 | Healthy Control | Intestine |
| Lpar6 | -1.35E+00 | 7.667889 | 15.21616 | 0.000724 | 0.003484 | Healthy Control | Intestine |
| Lpin1 | -1.83E+00 | 5.154378 | 2.341512 | 0.139885 | 0.282137 | Healthy Control | Intestine |
| Lpin2 | 4.16E-01 | 6.798512 | 1.088502 | 0.308194 | 0.520169 | Infection | Intestine |
| Lpin3 | -1.39E+00 | 6.263241 | 4.095584 | 0.054173 | 0.133789 | Healthy Control | Intestine |
| Lpl | 6.68E-01 | 7.666863 | 2.056848 | 0.168177 | 0.324555 | Infection | Intestine |
| Lpp | -8.71E-01 | 9.722809 | 27.83043 | 2.80E-05 | 0.000202 | Healthy Control | Intestine |
| Lratd2 | -7.49E-01 | 7.782809 | 6.948818 | 0.015047 | 0.046439 | Healthy Control | Intestine |
| Lrig1 | -1.93E+00 | 7.972796 | 21.74072 | 0.000102 | 0.00063 | Healthy Control | Intestine |
| Lrp1 | 1.09E+00 | 10.49356 | 31.08662 | 2.43E-05 | 0.00018 | Infection | Intestine |
| Lrp12 | 6.88E-01 | 5.83784 | 1.830811 | 0.189466 | 0.355987 | Infection | Intestine |
| Lrp2 | -3.57E-01 | 1.227067 | 0.006351 | 0.937596 | 0.969735 | Healthy Control | Intestine |
| Lrp5 | -6.20E-01 | 7.741997 | 4.799807 | 0.039263 | 0.102424 | Healthy Control | Intestine |
| Lrp6 | 7.34E-02 | 8.902648 | 0.095322 | 0.760451 | 0.928591 | Infection | Intestine |
| Lrp8 | 9.88E-02 | 4.031591 | 0.007223 | 0.933018 | 0.969735 | Infection | Intestine |
| Lrpprc | -4.44E+00 | 7.7792 | 42.06361 | 1.35E-06 | 1.44E-05 | Healthy Control | Intestine |
| Lrrc15 | -1.56E+00 | 1.480659 | 0.085533 | 0.772673 | 0.934526 | Healthy Control | Intestine |
| Lrrc52 | -7.56E-01 | 1.273813 | 0.014863 | 0.90436 | 0.969735 | Healthy Control | Intestine |
| Lrrc6 | -9.71E-16 | 1.180616 | 0 | 1 | 1 | Healthy Control | Intestine |
| Lrrk1 | -8.78E-01 | 7.740877 | 7.982425 | 0.00976 | 0.03223 | Healthy Control | Intestine |

| | | | | | | | |
|---|---|---|---|---|---|---|---|
| Lrrk2 | 1.95E+00 | 6.794143 | 28.80828 | 2.21E-05 | 0.000166 | Infection | Intestine |
| Lrrtm1 | -8.35E-01 | 4.749435 | 0.205756 | 0.653923 | 0.864186 | Healthy Control | Intestine |
| Lrrtm3 | -2.34E+00 | 1.523299 | 0.082351 | 0.776111 | 0.935745 | Healthy Control | Intestine |
| Lsm14a | 1.65E+00 | 8.840411 | 131.2369 | 1.08E-10 | 5.66E-09 | Infection | Intestine |
| Lsr | -2.71E+00 | 9.622152 | 55.37651 | 1.59E-07 | 2.31E-06 | Healthy Control | Intestine |
| Lss | -4.96E-01 | 7.654736 | 1.773359 | 0.196434 | 0.366648 | Healthy Control | Intestine |
| Lta | -4.40E+00 | 2.15193 | 0.34411 | 0.562837 | 0.790783 | Healthy Control | Intestine |
| Lta4h | 3.34E-01 | 9.075851 | 3.613659 | 0.070618 | 0.165426 | Infection | Intestine |
| Ltb4r1 | 1.83E+00 | 3.515403 | 3.248758 | 0.084255 | 0.18965 | Infection | Intestine |
| Ltbp1 | 1.75E+00 | 7.514552 | 20.6459 | 0.000212 | 0.001197 | Infection | Intestine |
| Ltbr | -8.10E-01 | 8.203329 | 8.74273 | 0.007278 | 0.025178 | Healthy Control | Intestine |
| Luc7l3 | 5.14E-01 | 9.037694 | 6.923629 | 0.015319 | 0.047192 | Infection | Intestine |
| Lum | 8.65E-01 | 9.264621 | 2.343729 | 0.150653 | 0.299273 | Infection | Intestine |
| Lurap1 | -6.00E+00 | 3.391511 | 1.752053 | 0.200849 | 0.372803 | Healthy Control | Intestine |
| Lvrn | 3.00E+00 | 1.947572 | 2.086138 | 0.159954 | 0.313014 | Infection | Intestine |
| Ly6g | 1.29E+00 | 3.63523 | 0.516086 | 0.480718 | 0.714724 | Infection | Intestine |
| Ly6g5b | -3.55E+00 | 2.176929 | 0.245463 | 0.625248 | 0.841396 | Healthy Control | Intestine |
| Ly6g6d | -7.19E+00 | 4.883873 | 3.455133 | 0.078057 | 0.179163 | Healthy Control | Intestine |
| Ly6g6e | -3.68E+00 | 8.09292 | 32.27655 | 8.77E-06 | 7.37E-05 | Healthy Control | Intestine |
| Ly6h | 2.61E+00 | 1.833549 | 2.603208 | 0.116573 | 0.24468 | Infection | Intestine |
| Ly6i | -3.57E-01 | 1.227509 | 0.006273 | 0.937979 | 0.969735 | Healthy Control | Intestine |
| Ly6k | 2.56E+00 | 1.832145 | 1.560193 | 0.220538 | 0.401459 | Infection | Intestine |
| Ly6m | -3.40E+00 | 5.543303 | 8.414219 | 0.00892 | 0.029829 | Healthy Control | Intestine |
| Ly75 | -4.14E+00 | 7.482423 | 22.30083 | 9.68E-05 | 0.000602 | Healthy Control | Intestine |
| Ly86 | 2.71E+00 | 5.401741 | 18.17584 | 0.000322 | 0.001735 | Infection | Intestine |
| Ly96 | -2.93E-01 | 6.109228 | 0.183523 | 0.672224 | 0.877942 | Healthy Control | Intestine |
| Lyar | -3.27E+00 | 6.492525 | 12.96622 | 0.001555 | 0.006695 | Healthy Control | Intestine |
| Lyg1 | -3.57E-01 | 1.227067 | 0.006351 | 0.937596 | 0.969735 | Healthy Control | Intestine |
| Lyn | 5.29E-01 | 7.18342 | 2.778969 | 0.109742 | 0.232856 | Infection | Intestine |
| Lypd1 | 2.74E+00 | 2.3888 | 3.048549 | 0.092271 | 0.20359 | Infection | Intestine |
| Lypd6 | 3.75E+00 | 1.559869 | 3.151485 | 0.08494 | 0.190849 | Infection | Intestine |
| Lypd6b | -2.83E+00 | 5.55105 | 4.684007 | 0.042202 | 0.108954 | Healthy Control | Intestine |
| Lyplal1 | -8.68E+00 | 6.068713 | 6.374002 | 0.020234 | 0.059794 | Healthy Control | Intestine |
| Lyst | 3.63E+00 | 7.5846 | 538.3879 | 7.32E-17 | 4.07E-14 | Infection | Intestine |
| Lyve1 | -3.53E-01 | 6.840724 | 0.030455 | 0.86418 | 0.959859 | Healthy Control | Intestine |
| Lyzl6 | -3.57E-01 | 1.228299 | 0.00618 | 0.938438 | 0.969735 | Healthy Control | Intestine |
| Lztr1 | 4.25E-01 | 8.058692 | 3.255415 | 0.085043 | 0.190909 | Infection | Intestine |
| Mad1l1 | -2.06E-01 | 6.414899 | 0.107053 | 0.746494 | 0.924073 | Healthy Control | Intestine |
| Mad2l1 | -4.65E-01 | 5.470952 | 0.236332 | 0.631143 | 0.846597 | Healthy Control | Intestine |
| Mad2l2 | 2.16E-01 | 6.31277 | 0.207397 | 0.653156 | 0.863896 | Infection | Intestine |
| Madcam1 | -9.05E-01 | 4.015128 | 0.085335 | 0.773916 | 0.934669 | Healthy Control | Intestine |
| Mael | 5.06E+00 | 1.320064 | 4.285086 | 0.051812 | 0.12904 | Infection | Intestine |
| Maf | 1.07E+00 | 8.112493 | 15.6248 | 0.000687 | 0.003334 | Infection | Intestine |
| Maf1 | -1.51E+00 | 7.954049 | 16.42769 | 0.000501 | 0.00254 | Healthy Control | Intestine |
| Mafa | -4.26E+00 | 2.267894 | 0.324957 | 0.574345 | 0.802221 | Healthy Control | Intestine |
| Mafb | 3.31E-01 | 7.103572 | 0.312774 | 0.582225 | 0.808493 | Infection | Intestine |
| Maff | 1.10E+00 | 5.929797 | 4.318124 | 0.049364 | 0.123744 | Infection | Intestine |
| Mafk | -2.59E+00 | 5.771816 | 5.984502 | 0.022823 | 0.066156 | Healthy Control | Intestine |
| Mag | 4.07E+00 | 2.211924 | 9.609242 | 0.00439 | 0.016332 | Infection | Intestine |
| Mageb4 | 6.00E+00 | 1.276247 | 5.575755 | 0.032212 | 0.086743 | Infection | Intestine |

| | | | | | | | |
|---|---|---|---|---|---|---|---|
| Magoh | -2.33E+00 | 8.158055 | 37.59848 | 2.65E-06 | 2.54E-05 | Healthy Control | Intestine |
| Maip1 | -3.02E-01 | 7.099221 | 0.492246 | 0.490175 | 0.723199 | Healthy Control | Intestine |
| Malt1 | 6.21E-01 | 6.093156 | 1.218835 | 0.281095 | 0.484726 | Infection | Intestine |
| Maml1 | 2.12E-01 | 6.75897 | 0.316736 | 0.579224 | 0.806375 | Infection | Intestine |
| Man2b2 | 8.34E-01 | 8.165626 | 12.6791 | 0.00177 | 0.007463 | Infection | Intestine |
| Maoa | -3.93E+00 | 11.79226 | 303.704 | 2.87E-14 | 5.32E-12 | Healthy Control | Intestine |
| Maob | -8.75E-01 | 6.867782 | 1.762823 | 0.196808 | 0.367071 | Healthy Control | Intestine |
| Map1b | 1.74E-01 | 6.637087 | 0.117029 | 0.735488 | 0.916538 | Infection | Intestine |
| Map1lc3a | 3.55E-01 | 8.425415 | 2.878576 | 0.104015 | 0.223629 | Infection | Intestine |
| Map2 | -2.22E+00 | 5.534815 | 2.198199 | 0.152147 | 0.301403 | Healthy Control | Intestine |
| Map2k1 | -2.25E+00 | 10.09563 | 113.1643 | 3.75E-10 | 1.47E-08 | Healthy Control | Intestine |
| Map2k2 | -1.19E+00 | 8.283498 | 17.44744 | 0.000392 | 0.002058 | Healthy Control | Intestine |
| Map2k3 | -2.27E+00 | 7.112284 | 13.06272 | 0.001385 | 0.006056 | Healthy Control | Intestine |
| Map2k4 | 1.12E+00 | 6.512417 | 8.855479 | 0.006998 | 0.024413 | Infection | Intestine |
| Map2k5 | -3.93E-01 | 6.59915 | 0.503326 | 0.485199 | 0.719463 | Healthy Control | Intestine |
| Map3k1 | 4.97E-01 | 7.062911 | 2.437001 | 0.132865 | 0.270595 | Infection | Intestine |
| Map3k11 | -8.09E-01 | 7.658043 | 3.890654 | 0.061061 | 0.147453 | Healthy Control | Intestine |
| Map3k14 | 2.57E+00 | 6.077545 | 72.99503 | 2.12E-08 | 4.22E-07 | Infection | Intestine |
| Map3k15 | 1.71E+00 | 3.829797 | 2.884998 | 0.102299 | 0.220832 | Infection | Intestine |
| Map3k5 | 1.40E-01 | 8.023007 | 0.315178 | 0.580241 | 0.807306 | Infection | Intestine |
| Map3k7 | -2.27E-01 | 8.121695 | 0.804299 | 0.379602 | 0.607703 | Healthy Control | Intestine |
| Map3k8 | 1.38E+00 | 4.948599 | 4.687292 | 0.040936 | 0.106123 | Infection | Intestine |
| Map4 | 1.95E-03 | 8.773382 | 8.41E-05 | 0.992767 | 1 | Infection | Intestine |
| Map4k4 | 2.44E+00 | 8.396795 | 135.425 | 7.99E-11 | 4.30E-09 | Infection | Intestine |
| Map6 | 5.29E-01 | 4.083328 | 0.114564 | 0.737992 | 0.919002 | Infection | Intestine |
| Map7 | -3.93E+00 | 10.06995 | 177.3795 | 2.02E-12 | 2.07E-10 | Healthy Control | Intestine |
| Map9 | 2.49E-01 | 4.058573 | 0.059988 | 0.808657 | 0.94887 | Infection | Intestine |
| Mapk1 | -1.30E+00 | 9.257169 | 45.63865 | 9.03E-07 | 1.03E-05 | Healthy Control | Intestine |
| Mapk10 | -4.62E+00 | 2.067336 | 0.325377 | 0.573093 | 0.80173 | Healthy Control | Intestine |
| Mapk11 | 2.45E+00 | 3.369355 | 8.618783 | 0.007171 | 0.024894 | Infection | Intestine |
| Mapk14 | -1.22E+00 | 7.374867 | 7.72823 | 0.010568 | 0.034421 | Healthy Control | Intestine |
| Mapk3 | -3.01E+00 | 10.11166 | 163.9049 | 1.09E-11 | 7.83E-10 | Healthy Control | Intestine |
| Mapk7 | 8.49E-01 | 5.77045 | 2.930008 | 0.100669 | 0.218158 | Infection | Intestine |
| Mapk8 | 9.70E-01 | 6.681949 | 8.266372 | 0.008819 | 0.029551 | Infection | Intestine |
| Mapk8ip1 | 1.10E+00 | 5.908701 | 3.325153 | 0.081785 | 0.185928 | Infection | Intestine |
| Mapk8ip2 | -6.36E+00 | 3.337039 | 1.25071 | 0.276954 | 0.478979 | Healthy Control | Intestine |
| Mapk9 | -7.37E-01 | 8.042534 | 5.540545 | 0.027894 | 0.077663 | Healthy Control | Intestine |
| Mapkap1 | -2.27E+00 | 8.391335 | 57.17782 | 1.07E-07 | 1.68E-06 | Healthy Control | Intestine |
| Mapkapk2 | 9.13E-01 | 8.643688 | 28.54296 | 2.38E-05 | 0.000177 | Infection | Intestine |
| Mapre1 | -1.23E+00 | 8.645443 | 19.93973 | 0.000194 | 0.001106 | Healthy Control | Intestine |
| Mapre2 | -7.12E-01 | 9.019799 | 12.1814 | 0.002096 | 0.008587 | Healthy Control | Intestine |
| Mapt | -8.79E+00 | 6.040471 | 9.606809 | 0.005698 | 0.020461 | Healthy Control | Intestine |
| 2-Mar | -8.05E-01 | 7.554591 | 3.804156 | 0.063841 | 0.152914 | Healthy Control | Intestine |
| 5-Mar | -1.84E+00 | 6.711107 | 10.40257 | 0.003599 | 0.013738 | Healthy Control | Intestine |
| Marcks | 4.52E-01 | 9.252803 | 8.739725 | 0.007343 | 0.025387 | Infection | Intestine |
| Marcksl1 | 2.39E+00 | 6.124223 | 56.06984 | 1.85E-07 | 2.64E-06 | Infection | Intestine |
| Marco | -7.56E-01 | 1.27399 | 0.0146 | 0.905204 | 0.969735 | Healthy Control | Intestine |
| Mark2 | 4.08E-01 | 9.17118 | 6.524099 | 0.018162 | 0.05468 | Infection | Intestine |
| Mark3 | -4.53E-01 | 7.84841 | 2.188561 | 0.153242 | 0.302975 | Healthy Control | Intestine |
| Marveld2 | -3.29E+00 | 7.542118 | 33.20069 | 6.35E-06 | 5.56E-05 | Healthy Control | Intestine |

| | | | | | | | |
|---|---|---|---|---|---|---|---|
| Mas1 | -9.71E-16 | 1.180616 | 0 | 1 | 1 | Healthy Control | Intestine |
| Mast2 | -1.24E+00 | 9.675358 | 41.7564 | 1.77E-06 | 1.79E-05 | Healthy Control | Intestine |
| Mast3 | 4.92E-01 | 6.819322 | 1.764233 | 0.197685 | 0.368209 | Infection | Intestine |
| Matn1 | -9.71E-16 | 1.180616 | 0 | 1 | 1 | Healthy Control | Intestine |
| Mavs | -9.31E-01 | 7.252871 | 5.698117 | 0.025509 | 0.072145 | Healthy Control | Intestine |
| Max | 8.93E-01 | 9.474235 | 20.33551 | 0.000177 | 0.001018 | Infection | Intestine |
| Maz | 1.36E-01 | 7.393711 | 0.174322 | 0.680332 | 0.882694 | Infection | Intestine |
| Mbd2 | -1.42E-01 | 10.41778 | 1.318121 | 0.26337 | 0.460989 | Healthy Control | Intestine |
| Mbd3 | 2.57E-01 | 8.518463 | 1.645601 | 0.213045 | 0.390467 | Infection | Intestine |
| Mbd5 | -2.35E-01 | 5.618569 | 0.085015 | 0.773137 | 0.934635 | Healthy Control | Intestine |
| Mbl1 | -7.56E-01 | 1.274081 | 0.014574 | 0.905288 | 0.969735 | Healthy Control | Intestine |
| Mbl2 | -2.16E+00 | 1.3196 | 0.08693 | 0.771266 | 0.933501 | Healthy Control | Intestine |
| Mbnl1 | -3.99E-01 | 9.097561 | 2.955902 | 0.09975 | 0.216261 | Healthy Control | Intestine |
| Mboat1 | -2.48E+00 | 10.34757 | 88.90231 | 3.49E-09 | 9.20E-08 | Healthy Control | Intestine |
| Mboat7 | -1.12E-01 | 8.668286 | 0.347687 | 0.561494 | 0.789783 | Healthy Control | Intestine |
| Mbtd1 | 4.73E-01 | 7.905781 | 2.535113 | 0.125749 | 0.258627 | Infection | Intestine |
| Mc1r | 5.33E+00 | 1.276041 | 4.71979 | 0.046639 | 0.118156 | Infection | Intestine |
| Mc3r | -3.57E-01 | 1.228299 | 0.00618 | 0.938438 | 0.969735 | Healthy Control | Intestine |
| Mc4r | -2.72E+00 | 1.445279 | 0.169643 | 0.683589 | 0.88417 | Healthy Control | Intestine |
| Mc5r | 4.04E+00 | 1.445532 | 2.731342 | 0.108336 | 0.230828 | Infection | Intestine |
| Mcam | 1.82E+00 | 6.750912 | 8.553184 | 0.009701 | 0.032075 | Infection | Intestine |
| Mcl1 | -4.62E-01 | 11.06224 | 13.41978 | 0.001381 | 0.00605 | Healthy Control | Intestine |
| Mcm2 | -7.59E-01 | 6.445206 | 0.919935 | 0.347132 | 0.570217 | Healthy Control | Intestine |
| Mcph1 | 3.40E-01 | 5.935683 | 0.222812 | 0.64137 | 0.855379 | Infection | Intestine |
| Mctp2 | -5.45E-01 | 7.168384 | 1.511359 | 0.231313 | 0.417689 | Healthy Control | Intestine |
| Mcu | -2.96E+00 | 9.885805 | 108.6076 | 5.50E-10 | 2.01E-08 | Healthy Control | Intestine |
| Mcub | 6.14E-02 | 4.942664 | 0.005875 | 0.939522 | 0.970342 | Infection | Intestine |
| Mcur1 | -3.32E+00 | 6.53918 | 10.92985 | 0.003172 | 0.012279 | Healthy Control | Intestine |
| Mdk | 3.42E+00 | 4.628777 | 26.76745 | 3.60E-05 | 0.000252 | Infection | Intestine |
| Mdm2 | 2.13E+00 | 7.24978 | 80.32016 | 9.25E-09 | 2.08E-07 | Infection | Intestine |
| Mdm4 | 1.18E+00 | 9.017462 | 39.74609 | 2.51E-06 | 2.42E-05 | Infection | Intestine |
| Mdp1 | -2.07E+00 | 8.634461 | 36.31954 | 4.22E-06 | 3.85E-05 | Healthy Control | Intestine |
| Me1 | -9.77E-01 | 4.498773 | 0.511845 | 0.481638 | 0.715666 | Healthy Control | Intestine |
| Meaf6 | 1.58E+00 | 5.74387 | 13.38451 | 0.00139 | 0.006071 | Infection | Intestine |
| Mecom | -2.98E+00 | 8.786545 | 99.87524 | 6.40E-10 | 2.24E-08 | Healthy Control | Intestine |
| Mecp2 | -4.79E-01 | 7.579259 | 1.513219 | 0.231598 | 0.417689 | Healthy Control | Intestine |
| Med1 | -2.64E+00 | 8.274572 | 40.79334 | 1.38E-06 | 1.46E-05 | Healthy Control | Intestine |
| Med12 | -1.16E-01 | 7.420772 | 0.130997 | 0.720858 | 0.906205 | Healthy Control | Intestine |
| Med14 | -2.07E+00 | 7.03482 | 10.49431 | 0.003439 | 0.013222 | Healthy Control | Intestine |
| Med15 | -5.79E-01 | 8.698168 | 6.251939 | 0.020436 | 0.060328 | Healthy Control | Intestine |
| Med21 | -1.98E+00 | 7.860792 | 26.75283 | 2.86E-05 | 0.000206 | Healthy Control | Intestine |
| Med24 | -1.15E+00 | 7.072438 | 4.635897 | 0.041882 | 0.108296 | Healthy Control | Intestine |
| Med6 | 1.07E+00 | 7.130539 | 14.0028 | 0.001144 | 0.005155 | Infection | Intestine |
| Medag | 3.30E+00 | 5.229538 | 26.82886 | 3.53E-05 | 0.000249 | Infection | Intestine |
| Mef2a | 5.91E-02 | 8.009141 | 0.043732 | 0.836297 | 0.954066 | Infection | Intestine |
| Mef2c | -8.28E-01 | 4.136572 | 0.258601 | 0.616097 | 0.836864 | Healthy Control | Intestine |
| Mef2d | -2.84E+00 | 8.859141 | 68.52756 | 1.99E-08 | 3.98E-07 | Healthy Control | Intestine |
| Megf11 | 7.45E+00 | 1.608146 | 35.38756 | 2.10E-06 | 2.07E-05 | Infection | Intestine |
| Mei4 | -3.37E+00 | 1.59824 | 0.27162 | 0.606413 | 0.829653 | Healthy Control | Intestine |
| Meiob | 5.39E+00 | 1.44736 | 9.048618 | 0.005358 | 0.019407 | Infection | Intestine |

| | | | | | | | |
|---|---|---|---|---|---|---|---|
| Meis1 | 5.28E+00 | 6.017083 | 191.1317 | 3.03E-11 | 1.92E-09 | Infection | Intestine |
| Meis2 | 2.63E+00 | 5.745871 | 43.3924 | 1.34E-06 | 1.43E-05 | Infection | Intestine |
| Meis3 | 1.86E+00 | 6.118565 | 15.1307 | 0.000803 | 0.003804 | Infection | Intestine |
| Memo1 | -8.33E-02 | 7.632096 | 0.075417 | 0.786165 | 0.940842 | Healthy Control | Intestine |
| Men1 | -1.47E+00 | 7.820358 | 14.10098 | 0.001022 | 0.004715 | Healthy Control | Intestine |
| Meox1 | 3.84E-01 | 5.04862 | 0.180187 | 0.674921 | 0.879171 | Infection | Intestine |
| Meox2 | 2.22E+00 | 3.815824 | 3.230452 | 0.08602 | 0.192497 | Infection | Intestine |
| Mep1a | -3.78E+00 | 9.209951 | 52.69656 | 1.62E-07 | 2.33E-06 | Healthy Control | Intestine |
| Mep1b | -2.76E+00 | 8.236923 | 13.40431 | 0.00156 | 0.006701 | Healthy Control | Intestine |
| Mepe | -4.94E-01 | 1.22813 | 0.009389 | 0.924171 | 0.969735 | Healthy Control | Intestine |
| Mertk | -2.39E+00 | 6.865263 | 9.557884 | 0.00492 | 0.018054 | Healthy Control | Intestine |
| Mesp1 | -1.89E+00 | 1.401345 | 0.052896 | 0.819819 | 0.953035 | Healthy Control | Intestine |
| Mesp2 | -3.98E+00 | 2.352629 | 0.424354 | 0.52046 | 0.755917 | Healthy Control | Intestine |
| Mest | 5.79E+00 | 2.823297 | 75.71691 | 8.42E-09 | 1.93E-07 | Infection | Intestine |
| Met | -1.95E+00 | 7.576144 | 21.99265 | 9.28E-05 | 0.00058 | Healthy Control | Intestine |
| Metrnl | 1.39E-01 | 5.41925 | 0.033462 | 0.856401 | 0.956553 | Infection | Intestine |
| Mettl14 | -3.54E-02 | 6.626254 | 0.005732 | 0.940322 | 0.970767 | Healthy Control | Intestine |
| Mettl21c | -3.57E-01 | 1.227417 | 0.006288 | 0.937908 | 0.969735 | Healthy Control | Intestine |
| Mettl3 | -1.15E+00 | 7.040206 | 6.144579 | 0.020752 | 0.061036 | Healthy Control | Intestine |
| Mettl7b | -2.68E+00 | 10.61133 | 60.87701 | 2.38E-07 | 3.24E-06 | Healthy Control | Intestine |
| Mfap5 | 3.36E+00 | 7.276424 | 84.72127 | 2.70E-08 | 5.21E-07 | Infection | Intestine |
| Mfn1 | -5.08E+00 | 8.358962 | 57.9949 | 1.10E-07 | 1.73E-06 | Healthy Control | Intestine |
| Mfn2 | -1.17E+00 | 8.469061 | 14.13571 | 0.001079 | 0.004927 | Healthy Control | Intestine |
| Mfrp | -2.16E+00 | 1.320374 | 0.060202 | 0.808769 | 0.94887 | Healthy Control | Intestine |
| Mfsd2a | -5.55E+00 | 2.622856 | 1.116034 | 0.30286 | 0.513242 | Healthy Control | Intestine |
| Mfsd2b | -5.19E-01 | 4.029721 | 0.123863 | 0.728157 | 0.912201 | Healthy Control | Intestine |
| Mia | -5.04E+00 | 2.507615 | 0.893779 | 0.355371 | 0.580421 | Healthy Control | Intestine |
| Mia2 | 7.74E-01 | 9.47982 | 29.88949 | 1.77E-05 | 0.000137 | Infection | Intestine |
| Mib1 | 1.49E-01 | 8.90776 | 0.861657 | 0.363449 | 0.58824 | Infection | Intestine |
| Micu1 | -8.30E-01 | 8.708395 | 11.81759 | 0.002369 | 0.00957 | Healthy Control | Intestine |
| Micu2 | -2.09E+00 | 8.825237 | 50.88613 | 3.45E-07 | 4.52E-06 | Healthy Control | Intestine |
| Miga1 | -7.41E+00 | 4.621648 | 3.08716 | 0.094374 | 0.207028 | Healthy Control | Intestine |
| Miga2 | -1.39E+00 | 7.494642 | 9.081126 | 0.006175 | 0.021953 | Healthy Control | Intestine |
| Minar2 | -4.36E+00 | 2.210348 | 0.356616 | 0.556055 | 0.784337 | Healthy Control | Intestine |
| Mindy1 | -4.45E-01 | 7.974582 | 1.625162 | 0.215698 | 0.394226 | Healthy Control | Intestine |
| Mip | -6.41E-01 | 1.272338 | 0.013961 | 0.90718 | 0.969735 | Healthy Control | Intestine |
| Misp | -3.74E+00 | 10.14541 | 157.3857 | 1.04E-11 | 7.79E-10 | Healthy Control | Intestine |
| Mitf | 3.05E+00 | 4.125199 | 21.59611 | 0.000123 | 0.000737 | Infection | Intestine |
| Mixl1 | -1.52E+00 | 1.273358 | 0.043368 | 0.83747 | 0.954066 | Healthy Control | Intestine |
| Mki67 | -1.03E+00 | 8.786027 | 6.971738 | 0.015022 | 0.046391 | Healthy Control | Intestine |
| Mkks | -1.01E+00 | 6.369159 | 2.19891 | 0.151247 | 0.300096 | Healthy Control | Intestine |
| Mkln1 | -8.15E-01 | 8.425435 | 8.714065 | 0.007388 | 0.025525 | Healthy Control | Intestine |
| Mknk1 | 1.22E+00 | 7.066325 | 12.8473 | 0.001672 | 0.007094 | Infection | Intestine |
| Mknk2 | -4.78E-01 | 10.22069 | 7.356493 | 0.012786 | 0.040535 | Healthy Control | Intestine |
| Mkx | -1.54E-01 | 6.020196 | 0.039306 | 0.84455 | 0.954066 | Healthy Control | Intestine |
| Mlc1 | -2.96E+00 | 1.525051 | 0.146511 | 0.706088 | 0.896379 | Healthy Control | Intestine |
| Mlec | -3.21E-01 | 9.707503 | 3.181087 | 0.088437 | 0.196937 | Healthy Control | Intestine |
| Mlf1 | -4.76E+00 | 2.214685 | 0.678833 | 0.417163 | 0.649959 | Healthy Control | Intestine |
| Mlh1 | -8.67E-03 | 6.484095 | 0.00037 | 0.984822 | 1 | Healthy Control | Intestine |
| Mlkl | -3.06E-01 | 7.091426 | 0.544535 | 0.468256 | 0.703509 | Healthy Control | Intestine |

| | | | | | | | |
|---|---|---|---|---|---|---|---|
| Mllt10 | -2.72E+00 | 7.580043 | 28.42067 | 1.80E-05 | 0.000139 | Healthy Control | Intestine |
| Mllt6 | -7.06E-01 | 8.771964 | 8.837663 | 0.007067 | 0.024585 | Healthy Control | Intestine |
| Mlph | -3.90E+00 | 8.852273 | 44.3782 | 6.48E-07 | 7.76E-06 | Healthy Control | Intestine |
| Mlst8 | 1.01E+00 | 6.718959 | 10.20204 | 0.004214 | 0.015765 | Infection | Intestine |
| Mlx | -2.68E+00 | 8.600528 | 42.66592 | 1.00E-06 | 1.13E-05 | Healthy Control | Intestine |
| Mlxipl | -1.45E+00 | 6.998572 | 4.081751 | 0.054603 | 0.134585 | Healthy Control | Intestine |
| Mlycd | -1.72E+00 | 6.541593 | 6.231911 | 0.019639 | 0.058393 | Healthy Control | Intestine |
| Mmd2 | -2.47E+00 | 1.444644 | 0.103292 | 0.75017 | 0.92508 | Healthy Control | Intestine |
| Mme | -6.81E+00 | 4.547072 | 1.918796 | 0.180944 | 0.344021 | Healthy Control | Intestine |
| Mmp12 | 4.36E+00 | 3.89926 | 32.70449 | 1.09E-05 | 8.85E-05 | Infection | Intestine |
| Mmp13 | 1.76E-01 | 5.90425 | 0.02656 | 0.872054 | 0.962625 | Infection | Intestine |
| Mmp14 | 1.79E+00 | 9.441156 | 127.2383 | 1.44E-10 | 7.05E-09 | Infection | Intestine |
| Mmp15 | -5.75E+00 | 8.878324 | 76.8583 | 1.12E-08 | 2.46E-07 | Healthy Control | Intestine |
| Mmp16 | 2.12E+00 | 3.324469 | 3.249035 | 0.083833 | 0.189041 | Infection | Intestine |
| Mmp2 | 7.43E-01 | 8.265681 | 3.814379 | 0.066267 | 0.157969 | Infection | Intestine |
| Mmp25 | 5.48E+00 | 3.791568 | 106.6362 | 8.31E-10 | 2.79E-08 | Infection | Intestine |
| Mmp28 | 6.79E-01 | 5.722223 | 0.86227 | 0.362721 | 0.587498 | Infection | Intestine |
| Mmp3 | 3.22E+00 | 2.063656 | 6.346113 | 0.01717 | 0.051998 | Infection | Intestine |
| Mmp7 | -6.81E+00 | 3.880068 | 1.472546 | 0.237033 | 0.424388 | Healthy Control | Intestine |
| Mmp9 | 4.00E+00 | 4.65516 | 35.81002 | 6.02E-06 | 5.28E-05 | Infection | Intestine |
| Mmrn1 | -5.00E-01 | 5.94922 | 0.040147 | 0.843874 | 0.954066 | Healthy Control | Intestine |
| Mms19 | -4.37E-01 | 7.638657 | 0.888363 | 0.356119 | 0.580694 | Healthy Control | Intestine |
| Mmut | -1.39E+00 | 8.539592 | 29.52471 | 1.86E-05 | 0.000143 | Healthy Control | Intestine |
| Mnx1 | -5.54E+00 | 3.636959 | 0.477367 | 0.4968 | 0.72966 | Healthy Control | Intestine |
| Mob2 | 2.30E-03 | 8.324796 | 6.69E-05 | 0.993546 | 1 | Infection | Intestine |
| Mog | 4.53E+00 | 1.404669 | 3.002159 | 0.094308 | 0.206973 | Infection | Intestine |
| Mos | -1.65E+00 | 1.317249 | 0.052832 | 0.820314 | 0.953035 | Healthy Control | Intestine |
| Mov10l1 | -1.71E+00 | 1.319252 | 0.050134 | 0.825039 | 0.953685 | Healthy Control | Intestine |
| Moxd1 | -4.66E+00 | 2.185675 | 0.377861 | 0.546141 | 0.776921 | Healthy Control | Intestine |
| Mpc1 | -3.56E+00 | 8.40745 | 33.63837 | 5.29E-06 | 4.69E-05 | Healthy Control | Intestine |
| Mpc2 | -2.23E+00 | 8.707041 | 36.96236 | 3.63E-06 | 3.37E-05 | Healthy Control | Intestine |
| Mpi | -1.40E+00 | 7.930992 | 16.1516 | 0.000538 | 0.002696 | Healthy Control | Intestine |
| Mpl | -2.36E+00 | 1.403253 | 0.09047 | 0.765843 | 0.931224 | Healthy Control | Intestine |
| Mplkip | 8.86E-02 | 6.148692 | 0.031093 | 0.861558 | 0.957372 | Infection | Intestine |
| Mpp2 | 4.69E-01 | 4.414613 | 0.205176 | 0.654534 | 0.864378 | Infection | Intestine |
| Mpp3 | -8.74E+00 | 5.917405 | 8.573127 | 0.008375 | 0.028347 | Healthy Control | Intestine |
| Mprip | 9.53E-02 | 10.268 | 0.383307 | 0.542253 | 0.774913 | Infection | Intestine |
| Mrc1 | -2.03E+00 | 7.566531 | 12.68129 | 0.001578 | 0.006758 | Healthy Control | Intestine |
| Mre11a | -1.21E-02 | 7.187415 | 0.001263 | 0.971974 | 0.993765 | Healthy Control | Intestine |
| Mrgprf | 1.37E+00 | 5.758233 | 4.626483 | 0.042747 | 0.110249 | Infection | Intestine |
| Mrm3 | -1.22E+00 | 5.585783 | 1.534413 | 0.22734 | 0.412043 | Healthy Control | Intestine |
| Mrnip | 1.44E+00 | 4.284673 | 2.24571 | 0.147056 | 0.293761 | Infection | Intestine |
| Mrtfa | 8.06E-01 | 3.792276 | 0.451021 | 0.508402 | 0.743084 | Infection | Intestine |
| Ms4a1 | -5.82E+00 | 3.322872 | 0.203164 | 0.658376 | 0.867142 | Healthy Control | Intestine |
| Ms4a2 | 5.33E+00 | 1.276175 | 4.716338 | 0.046709 | 0.118213 | Infection | Intestine |
| Ms4a7 | 1.72E+00 | 5.280004 | 4.821146 | 0.038678 | 0.101055 | Infection | Intestine |
| Msh2 | -1.57E-01 | 7.227566 | 0.227257 | 0.638218 | 0.853116 | Healthy Control | Intestine |
| Msi1 | 1.39E+00 | 4.72054 | 2.639172 | 0.117854 | 0.246336 | Infection | Intestine |
| Msln | 2.19E+00 | 1.975831 | 0.857366 | 0.362282 | 0.5873 | Infection | Intestine |
| Msmo1 | -2.50E+00 | 9.091594 | 40.02227 | 1.78E-06 | 1.79E-05 | Healthy Control | Intestine |

| | | | | | | | |
|---|---|---|---|---|---|---|---|
| Msn | 2.28E+00 | 8.739466 | 124.9416 | 1.70E-10 | 7.76E-09 | Infection | Intestine |
| Msr1 | 7.36E-01 | 4.929899 | 0.680417 | 0.417629 | 0.650153 | Infection | Intestine |
| Msra | -3.06E+00 | 7.93785 | 37.6998 | 2.09E-06 | 2.07E-05 | Healthy Control | Intestine |
| Msrb1 | -2.01E+00 | 6.848904 | 11.0847 | 0.002751 | 0.010869 | Healthy Control | Intestine |
| Mst1 | -3.64E+00 | 1.668462 | 0.336619 | 0.565761 | 0.794001 | Healthy Control | Intestine |
| Mst1r | -1.78E+00 | 7.084959 | 9.402665 | 0.005289 | 0.019171 | Healthy Control | Intestine |
| Mstn | -2.47E+00 | 1.446302 | 0.089576 | 0.766806 | 0.932159 | Healthy Control | Intestine |
| Msx1 | 5.70E+00 | 2.496779 | 68.45872 | 8.91E-09 | 2.02E-07 | Infection | Intestine |
| Msx2 | -1.34E+00 | 1.402605 | 0.026184 | 0.872638 | 0.962844 | Healthy Control | Intestine |
| Mta1 | 4.78E-02 | 7.873048 | 0.035161 | 0.85299 | 0.956553 | Infection | Intestine |
| Mta2 | 2.59E-01 | 9.3565 | 2.914721 | 0.101982 | 0.220338 | Infection | Intestine |
| Mta3 | -1.53E+00 | 8.37719 | 23.06463 | 8.16E-05 | 0.000521 | Healthy Control | Intestine |
| Mtch1 | -4.06E-01 | 9.78504 | 8.389795 | 0.008423 | 0.028453 | Healthy Control | Intestine |
| Mtch2 | -1.41E+00 | 9.228819 | 35.29455 | 5.85E-06 | 5.14E-05 | Healthy Control | Intestine |
| Mtcl1 | -6.86E-01 | 3.907979 | 0.168027 | 0.685877 | 0.885076 | Healthy Control | Intestine |
| Mtf1 | -1.44E+00 | 7.046132 | 6.165845 | 0.020445 | 0.060328 | Healthy Control | Intestine |
| Mtf2 | -1.34E+00 | 7.882491 | 12.1083 | 0.002036 | 0.008384 | Healthy Control | Intestine |
| Mtfp1 | -2.17E+00 | 6.459023 | 6.400346 | 0.018465 | 0.055383 | Healthy Control | Intestine |
| Mthfr | -6.56E-01 | 5.135169 | 0.453785 | 0.506827 | 0.741648 | Healthy Control | Intestine |
| Mtif2 | -5.24E-01 | 7.778259 | 3.094747 | 0.092409 | 0.203774 | Healthy Control | Intestine |
| Mtmr12 | -9.32E-01 | 7.566142 | 6.221558 | 0.020451 | 0.060328 | Healthy Control | Intestine |
| Mtnr1a | -1.52E+00 | 1.274908 | 0.03863 | 0.846477 | 0.954066 | Healthy Control | Intestine |
| Mtnr1b | -1.37E+00 | 1.229023 | 0.032546 | 0.859435 | 0.956553 | Healthy Control | Intestine |
| Mtor | 8.96E-01 | 8.077919 | 17.20617 | 0.000428 | 0.002223 | Infection | Intestine |
| Mtrex | -9.08E-01 | 7.171886 | 5.861583 | 0.023784 | 0.068386 | Healthy Control | Intestine |
| Mttp | 6.67E-01 | 3.24687 | 0.163031 | 0.690145 | 0.888369 | Infection | Intestine |
| Mturn | 1.24E+00 | 5.682568 | 3.810213 | 0.063735 | 0.152806 | Infection | Intestine |
| Mtus1 | -1.56E+00 | 9.285332 | 41.67216 | 1.76E-06 | 1.79E-05 | Healthy Control | Intestine |
| Mtx1 | 1.19E-01 | 6.564984 | 0.056219 | 0.814739 | 0.951358 | Infection | Intestine |
| Muc1 | -3.34E-01 | 3.226053 | 0.053162 | 0.819786 | 0.953035 | Healthy Control | Intestine |
| Muc2 | -2.83E+00 | 14.64845 | 24.26449 | 0.000355 | 0.001889 | Healthy Control | Intestine |
| Muc20 | -5.98E-01 | 4.309689 | 0.256198 | 0.617916 | 0.837736 | Healthy Control | Intestine |
| Muc5ac | -3.57E-01 | 1.228165 | 0.006192 | 0.93838 | 0.969735 | Healthy Control | Intestine |
| Mus81 | 5.39E-01 | 6.020095 | 0.888816 | 0.355829 | 0.580457 | Infection | Intestine |
| Musk | -1.15E+00 | 4.595912 | 0.430058 | 0.518568 | 0.75398 | Healthy Control | Intestine |
| Mxd1 | -2.73E-01 | 9.532298 | 0.178927 | 0.679505 | 0.882616 | Healthy Control | Intestine |
| Mxd4 | -2.47E+00 | 7.887829 | 25.91916 | 3.19E-05 | 0.000227 | Healthy Control | Intestine |
| Myb | -2.72E+00 | 9.681382 | 57.24052 | 1.16E-07 | 1.80E-06 | Healthy Control | Intestine |
| Mybbp1a | -8.39E-01 | 8.104654 | 7.692777 | 0.011057 | 0.035779 | Healthy Control | Intestine |
| Mybl2 | 2.52E+00 | 3.602414 | 5.673328 | 0.02578 | 0.072789 | Infection | Intestine |
| Mybpc3 | 3.45E+00 | 1.669466 | 2.036435 | 0.162352 | 0.316855 | Infection | Intestine |
| Mybph | -2.93E+00 | 1.405615 | 0.188921 | 0.668856 | 0.8756 | Healthy Control | Intestine |
| Myc | -1.49E+00 | 8.061019 | 14.26993 | 0.000973 | 0.004513 | Healthy Control | Intestine |
| Mycbp2 | 3.79E-01 | 7.182043 | 1.501612 | 0.233417 | 0.419714 | Infection | Intestine |
| Mycl | -9.29E+00 | 6.721455 | 10.14944 | 0.004685 | 0.017295 | Healthy Control | Intestine |
| Mycn | 5.10E-01 | 5.023867 | 0.188684 | 0.667935 | 0.874622 | Infection | Intestine |
| Myd88 | 4.12E-01 | 7.531362 | 2.057553 | 0.165635 | 0.320638 | Infection | Intestine |
| Myf5 | 4.54E+00 | 1.523326 | 3.864055 | 0.058378 | 0.14207 | Infection | Intestine |
| Myf6 | -1.17E+00 | 1.359601 | 0.034947 | 0.853153 | 0.956553 | Healthy Control | Intestine |
| Myh1 | 4.03E+00 | 1.447988 | 2.203593 | 0.14763 | 0.294554 | Infection | Intestine |

| | | | | | | | |
|---|---|---|---|---|---|---|---|
| Myh10 | 7.53E-01 | 5.866126 | 0.958573 | 0.338079 | 0.558003 | Infection | Intestine |
| Myh11 | 1.29E+00 | 9.809788 | 15.62864 | 0.001394 | 0.006086 | Infection | Intestine |
| Myh2 | -2.33E+00 | 1.402041 | 0.094685 | 0.760589 | 0.928591 | Healthy Control | Intestine |
| Myh3 | -7.56E-01 | 1.273905 | 0.014836 | 0.904447 | 0.969735 | Healthy Control | Intestine |
| Myh4 | 6.12E+00 | 1.322644 | 9.606637 | 0.007516 | 0.025913 | Infection | Intestine |
| Myh7 | -4.95E-01 | 1.228692 | 0.008683 | 0.927069 | 0.969735 | Healthy Control | Intestine |
| Myh8 | -8.80E-01 | 1.318204 | 0.019988 | 0.888814 | 0.969599 | Healthy Control | Intestine |
| Myh9 | -5.95E-01 | 11.08495 | 27.83892 | 2.78E-05 | 0.000202 | Healthy Control | Intestine |
| Myl2 | -3.57E-01 | 1.227417 | 0.006288 | 0.937908 | 0.969735 | Healthy Control | Intestine |
| Mymk | -1.52E+00 | 1.273972 | 0.038768 | 0.846206 | 0.954066 | Healthy Control | Intestine |
| Myo10 | 5.75E-01 | 8.188262 | 6.394768 | 0.019211 | 0.057278 | Infection | Intestine |
| Myo16 | 8.32E-01 | 4.199887 | 0.397967 | 0.533985 | 0.765939 | Infection | Intestine |
| Myo18a | -2.96E+00 | 9.507218 | 118.7219 | 2.04E-10 | 8.82E-09 | Healthy Control | Intestine |
| Myo1c | -5.10E-01 | 9.468263 | 5.441229 | 0.029314 | 0.080615 | Healthy Control | Intestine |
| Myo1d | -3.02E+00 | 10.60702 | 196.5843 | 2.16E-12 | 2.16E-10 | Healthy Control | Intestine |
| Myo1e | -1.20E+00 | 9.636009 | 36.59251 | 4.54E-06 | 4.13E-05 | Healthy Control | Intestine |
| Myo5a | 2.37E+00 | 6.407536 | 31.54788 | 1.25E-05 | 1.00E-04 | Infection | Intestine |
| Myo6 | -3.38E+00 | 10.11596 | 139.8933 | 3.84E-11 | 2.26E-09 | Healthy Control | Intestine |
| Myo7a | 6.59E-01 | 3.289537 | 0.139622 | 0.712094 | 0.900212 | Infection | Intestine |
| Myo9a | 7.94E-01 | 6.459393 | 3.455833 | 0.07648 | 0.175865 | Infection | Intestine |
| Myocd | -7.93E-01 | 4.401645 | 0.248268 | 0.623027 | 0.839761 | Healthy Control | Intestine |
| Myod1 | -1.86E+00 | 1.555947 | 0.076036 | 0.78507 | 0.940206 | Healthy Control | Intestine |
| Myog | -6.42E-01 | 1.272459 | 0.013821 | 0.907644 | 0.969735 | Healthy Control | Intestine |
| Myrf | 4.14E+00 | 1.522564 | 3.345515 | 0.076058 | 0.175218 | Infection | Intestine |
| Myt1 | -6.13E+00 | 3.65257 | 1.129763 | 0.300693 | 0.510261 | Healthy Control | Intestine |
| Mzb1 | -9.64E-01 | 6.7899 | 0.380373 | 0.546501 | 0.777212 | Healthy Control | Intestine |
| Naa15 | -9.08E-01 | 8.067298 | 9.470358 | 0.005498 | 0.019816 | Healthy Control | Intestine |
| Naaa | -1.47E-01 | 7.130703 | 0.044095 | 0.835878 | 0.954066 | Healthy Control | Intestine |
| Nadk2 | -1.11E+00 | 6.834333 | 3.673363 | 0.067381 | 0.16009 | Healthy Control | Intestine |
| Nae1 | -3.56E-01 | 7.778199 | 1.412565 | 0.247288 | 0.439439 | Healthy Control | Intestine |
| Nagk | -2.81E+00 | 7.798044 | 35.79573 | 3.47E-06 | 3.23E-05 | Healthy Control | Intestine |
| Naip1 | -1.10E+01 | 8.160453 | 20.53461 | 0.000335 | 0.001793 | Healthy Control | Intestine |
| Naip5 | -4.25E+00 | 7.467465 | 25.83602 | 4.12E-05 | 0.000283 | Healthy Control | Intestine |
| Nampt | 1.08E+00 | 7.793749 | 18.88186 | 0.000265 | 0.00146 | Infection | Intestine |
| Nanog | -1.89E+00 | 1.400971 | 0.067967 | 0.796283 | 0.943769 | Healthy Control | Intestine |
| Nat1 | -3.57E-01 | 1.227387 | 0.006293 | 0.937883 | 0.969735 | Healthy Control | Intestine |
| Nbn | -2.83E+00 | 7.078106 | 12.71306 | 0.001562 | 0.006701 | Healthy Control | Intestine |
| Nbr1 | -2.33E+00 | 9.493513 | 99.46605 | 1.29E-09 | 4.07E-08 | Healthy Control | Intestine |
| Ncam1 | -1.30E+00 | 6.703221 | 1.86334 | 0.184858 | 0.349867 | Healthy Control | Intestine |
| Ncam2 | -3.87E+00 | 2.026481 | 0.184144 | 0.671425 | 0.877586 | Healthy Control | Intestine |
| Ncan | -4.95E-01 | 1.228692 | 0.008683 | 0.927069 | 0.969735 | Healthy Control | Intestine |
| Ncaph | -1.23E+00 | 6.309197 | 3.165401 | 0.087509 | 0.19527 | Healthy Control | Intestine |
| Ncbp2 | -6.09E-01 | 7.359091 | 3.023654 | 0.095734 | 0.20946 | Healthy Control | Intestine |
| Ncf1 | 5.97E+00 | 7.631637 | 804.0808 | 1.53E-17 | 1.27E-14 | Infection | Intestine |
| Ncf4 | 7.89E-01 | 5.394834 | 0.845257 | 0.367258 | 0.592872 | Infection | Intestine |
| Nck1 | -3.19E+00 | 7.362926 | 23.89132 | 5.76E-05 | 0.00038 | Healthy Control | Intestine |
| Ncl | -2.01E+00 | 8.658241 | 26.35765 | 3.37E-05 | 0.000238 | Healthy Control | Intestine |
| Ncoa1 | -1.38E+00 | 7.979322 | 13.53006 | 0.001255 | 0.005591 | Healthy Control | Intestine |
| Ncoa2 | -1.35E+00 | 8.418983 | 19.28015 | 0.00023 | 0.001291 | Healthy Control | Intestine |
| Ncoa3 | -2.07E-01 | 8.491627 | 0.415735 | 0.525802 | 0.759561 | Healthy Control | Intestine |

| | | | | | | | |
|---|---|---|---|---|---|---|---|
| Ncor1 | -1.01E+00 | 9.823872 | 42.91112 | 1.44E-06 | 1.51E-05 | Healthy Control | Intestine |
| Ncor2 | -2.96E-01 | 8.138879 | 1.163218 | 0.292571 | 0.499356 | Healthy Control | Intestine |
| Ncr1 | -6.92E+00 | 4.048768 | 1.405673 | 0.249896 | 0.443139 | Healthy Control | Intestine |
| Ncstn | 2.96E-01 | 8.442721 | 2.357239 | 0.139106 | 0.281019 | Infection | Intestine |
| Ndc1 | 1.23E+00 | 6.485809 | 8.379594 | 0.008465 | 0.028574 | Infection | Intestine |
| Ndc80 | 1.92E+00 | 4.416433 | 6.853296 | 0.015432 | 0.047483 | Infection | Intestine |
| Nde1 | 6.24E-01 | 6.994967 | 3.187615 | 0.088109 | 0.196382 | Infection | Intestine |
| Ndel1 | 1.44E+00 | 8.099962 | 42.82178 | 1.46E-06 | 1.53E-05 | Infection | Intestine |
| Ndn | 3.96E-01 | 5.745844 | 0.322193 | 0.575854 | 0.803435 | Infection | Intestine |
| Ndor1 | 3.41E-01 | 6.636559 | 0.715217 | 0.40674 | 0.637769 | Infection | Intestine |
| Ndrg2 | -2.75E+00 | 9.318947 | 46.9547 | 5.73E-07 | 7.02E-06 | Healthy Control | Intestine |
| Ndrg4 | 3.32E+00 | 4.644096 | 25.97386 | 4.25E-05 | 0.000291 | Infection | Intestine |
| Ndst1 | -2.48E+00 | 7.809244 | 22.00536 | 9.05E-05 | 0.000568 | Healthy Control | Intestine |
| Ndst2 | -1.29E-01 | 7.62227 | 0.197927 | 0.660757 | 0.868632 | Healthy Control | Intestine |
| Ndst3 | 1.36E+00 | 3.658475 | 1.13676 | 0.296733 | 0.505264 | Infection | Intestine |
| Ndufa12 | -7.18E+00 | 4.66498 | 3.220456 | 0.088029 | 0.196291 | Healthy Control | Intestine |
| Ndufa9 | -2.87E+00 | 9.339994 | 106.0014 | 5.63E-10 | 2.03E-08 | Healthy Control | Intestine |
| Ndufaf6 | 1.26E-01 | 6.578655 | 0.065703 | 0.800061 | 0.945939 | Infection | Intestine |
| Ndufs1 | -1.83E+00 | 9.899727 | 79.98764 | 9.66E-09 | 2.15E-07 | Healthy Control | Intestine |
| Ndufs4 | -1.20E+00 | 8.048294 | 14.06156 | 0.001095 | 0.004993 | Healthy Control | Intestine |
| Ndufv2 | -1.07E+01 | 7.94272 | 48.79431 | 9.35E-07 | 1.06E-05 | Healthy Control | Intestine |
| Neat1 | 1.73E+00 | 11.44871 | 172.155 | 1.21E-11 | 8.54E-10 | Infection | Intestine |
| Nectin2 | -2.75E+00 | 8.63601 | 54.7997 | 1.39E-07 | 2.09E-06 | Healthy Control | Intestine |
| Nectin3 | -1.10E+00 | 9.111145 | 21.51127 | 0.000129 | 0.000771 | Healthy Control | Intestine |
| Nedd4 | -1.27E+00 | 10.93877 | 80.62064 | 8.93E-09 | 2.02E-07 | Healthy Control | Intestine |
| Nedd4l | -8.81E-01 | 8.898119 | 13.00482 | 0.001589 | 0.006792 | Healthy Control | Intestine |
| Nefh | -5.44E+00 | 2.865243 | 1.106044 | 0.305572 | 0.516547 | Healthy Control | Intestine |
| Nefl | -6.35E+00 | 3.384238 | 1.170919 | 0.291673 | 0.498502 | Healthy Control | Intestine |
| Nefm | -6.08E+00 | 3.417922 | 0.976238 | 0.334935 | 0.554459 | Healthy Control | Intestine |
| Neil1 | -3.96E-01 | 6.850339 | 0.93534 | 0.343679 | 0.565383 | Healthy Control | Intestine |
| Nek2 | -2.57E+00 | 6.012612 | 4.664291 | 0.041524 | 0.107426 | Healthy Control | Intestine |
| Nek7 | -1.22E+00 | 7.965124 | 12.17927 | 0.002024 | 0.008347 | Healthy Control | Intestine |
| Nelfb | -8.24E-01 | 8.859489 | 11.90416 | 0.0023 | 0.009317 | Healthy Control | Intestine |
| Nelfe | 1.73E-01 | 7.58023 | 0.312429 | 0.58188 | 0.808462 | Infection | Intestine |
| Nell1 | -1.94E+00 | 1.402433 | 0.046915 | 0.830029 | 0.954066 | Healthy Control | Intestine |
| Neo1 | 2.17E-02 | 7.932565 | 0.009016 | 0.925216 | 0.969735 | Infection | Intestine |
| Nes | 3.17E+00 | 4.750692 | 29.35235 | 2.01E-05 | 0.000153 | Infection | Intestine |
| Neu1 | -9.81E-01 | 9.253146 | 7.931878 | 0.010726 | 0.034823 | Healthy Control | Intestine |
| Neurod1 | 4.94E-01 | 3.037329 | 0.076452 | 0.784725 | 0.940206 | Infection | Intestine |
| Neurod2 | -1.89E+00 | 1.402675 | 0.059275 | 0.809471 | 0.94887 | Healthy Control | Intestine |
| Neurod4 | -6.42E-01 | 1.273568 | 0.012758 | 0.911248 | 0.969735 | Healthy Control | Intestine |
| Neurod6 | -1.65E+00 | 1.318234 | 0.039104 | 0.845043 | 0.954066 | Healthy Control | Intestine |
| Neurog1 | -9.71E-16 | 1.180616 | 0 | 1 | 1 | Healthy Control | Intestine |
| Neurog2 | -3.57E-01 | 1.227067 | 0.006351 | 0.937596 | 0.969735 | Healthy Control | Intestine |
| Neurog3 | 1.21E+00 | 2.696533 | 0.369439 | 0.548891 | 0.778399 | Infection | Intestine |
| Nf1 | 9.28E-01 | 8.243542 | 23.17756 | 8.48E-05 | 0.000539 | Infection | Intestine |
| Nf2 | -2.35E+00 | 8.404319 | 36.12456 | 3.89E-06 | 3.57E-05 | Healthy Control | Intestine |
| Nfat5 | 8.86E-01 | 8.520291 | 26.69199 | 3.62E-05 | 0.000254 | Infection | Intestine |
| Nfatc1 | 6.78E-02 | 7.231214 | 0.045447 | 0.833146 | 0.954066 | Infection | Intestine |
| Nfatc2 | -2.02E+00 | 6.192038 | 3.4936 | 0.074176 | 0.172229 | Healthy Control | Intestine |

| | | | | | | | |
|---|---|---|---|---|---|---|---|
| Nfatc3 | -1.56E+00 | 8.304538 | 23.99214 | 6.40E-05 | 0.000417 | Healthy Control | Intestine |
| Nfe2 | 2.96E+00 | 3.045005 | 5.459354 | 0.027735 | 0.077306 | Infection | Intestine |
| Nfe2l1 | -2.80E-01 | 9.76548 | 3.978483 | 0.058739 | 0.14281 | Healthy Control | Intestine |
| Nfe2l2 | -2.12E+00 | 9.834235 | 85.74561 | 5.27E-09 | 1.29E-07 | Healthy Control | Intestine |
| Nfia | -2.38E+00 | 8.503365 | 48.68798 | 3.77E-07 | 4.88E-06 | Healthy Control | Intestine |
| Nfib | -7.01E-03 | 8.645376 | 0.000934 | 0.975899 | 0.996603 | Healthy Control | Intestine |
| Nfil3 | 1.73E+00 | 5.790974 | 11.36505 | 0.002762 | 0.010904 | Infection | Intestine |
| Nfix | 9.40E-01 | 8.678548 | 22.04969 | 0.000113 | 0.000689 | Infection | Intestine |
| Nfkb1 | -1.47E-01 | 8.546571 | 0.499938 | 0.487027 | 0.721412 | Healthy Control | Intestine |
| Nfkb2 | 8.95E-01 | 7.813573 | 13.61982 | 0.001296 | 0.005743 | Infection | Intestine |
| Nfkbib | -1.93E+00 | 7.751804 | 21.10755 | 0.000121 | 0.000732 | Healthy Control | Intestine |
| Nfkbie | 1.66E+00 | 6.081485 | 19.06432 | 0.000245 | 0.001364 | Infection | Intestine |
| Nfu1 | -7.19E+00 | 4.429359 | 3.436289 | 0.078753 | 0.180265 | Healthy Control | Intestine |
| Nfya | -3.42E-01 | 6.706821 | 0.490044 | 0.490926 | 0.72388 | Healthy Control | Intestine |
| Nfyc | -1.05E+00 | 7.129348 | 6.173085 | 0.020596 | 0.060648 | Healthy Control | Intestine |
| Ngb | -4.53E+00 | 2.560889 | 0.591729 | 0.450133 | 0.684289 | Healthy Control | Intestine |
| Ngf | -5.65E+00 | 3.259475 | 0.367703 | 0.552447 | 0.781891 | Healthy Control | Intestine |
| Ngfr | -5.94E+00 | 3.445009 | 0.633059 | 0.435782 | 0.66839 | Healthy Control | Intestine |
| Nhlh1 | -1.25E+00 | 1.360779 | 0.036513 | 0.850035 | 0.955127 | Healthy Control | Intestine |
| Nhlh2 | -2.47E+00 | 1.446099 | 0.078937 | 0.780694 | 0.938784 | Healthy Control | Intestine |
| Nhsl1 | -1.39E+00 | 8.839638 | 21.54849 | 0.000124 | 0.000743 | Healthy Control | Intestine |
| Nid1 | 3.17E-01 | 7.942644 | 0.704074 | 0.410542 | 0.642844 | Infection | Intestine |
| Nid2 | 3.29E-01 | 6.192138 | 0.243062 | 0.6268 | 0.842805 | Infection | Intestine |
| Ninl | 5.17E-01 | 4.057889 | 0.201577 | 0.657378 | 0.866693 | Infection | Intestine |
| Nipbl | -1.46E+00 | 8.296027 | 16.0845 | 0.000571 | 0.002849 | Healthy Control | Intestine |
| Nipsnap2 | -1.81E+00 | 6.68861 | 6.100815 | 0.020951 | 0.061506 | Healthy Control | Intestine |
| Nkain1 | -7.12E+00 | 4.432502 | 2.571113 | 0.124702 | 0.257214 | Healthy Control | Intestine |
| Nkain3 | -2.16E+00 | 1.661287 | 0.078229 | 0.782306 | 0.940206 | Healthy Control | Intestine |
| Nkd2 | 7.99E-01 | 2.867471 | 0.209854 | 0.651324 | 0.862346 | Infection | Intestine |
| Nkx2-1 | 5.33E+00 | 1.275264 | 4.750983 | 0.046013 | 0.117162 | Infection | Intestine |
| Nkx2-2 | -6.43E+00 | 3.61528 | 2.254028 | 0.149156 | 0.297126 | Healthy Control | Intestine |
| Nkx2-5 | 5.70E+00 | 1.229995 | 6.086704 | 0.031631 | 0.085684 | Infection | Intestine |
| Nkx3-1 | -1.71E+00 | 1.31855 | 0.04995 | 0.825355 | 0.953685 | Healthy Control | Intestine |
| Nkx3-2 | 1.99E+00 | 3.058418 | 3.24485 | 0.084459 | 0.190022 | Infection | Intestine |
| Nkx6-1 | 6.00E+00 | 1.275016 | 5.60432 | 0.031838 | 0.086058 | Infection | Intestine |
| Nkx6-2 | -1.32E+00 | 1.361836 | 0.035032 | 0.853503 | 0.956553 | Healthy Control | Intestine |
| Nlgn2 | 2.70E+00 | 5.897461 | 63.38558 | 6.91E-08 | 1.18E-06 | Infection | Intestine |
| Nlgn3 | 2.35E+00 | 3.99648 | 5.432214 | 0.028962 | 0.079882 | Infection | Intestine |
| Nlk | 2.93E-01 | 6.536774 | 0.3073 | 0.584802 | 0.810722 | Infection | Intestine |
| Nlrc4 | -2.64E+00 | 7.633507 | 14.55224 | 0.000823 | 0.003891 | Healthy Control | Intestine |
| Nlrp12 | 7.27E+00 | 1.685122 | 34.92478 | 4.95E-06 | 4.43E-05 | Infection | Intestine |
| Nlrp3 | -8.82E-03 | 3.708987 | 2.78E-05 | 0.995836 | 1 | Healthy Control | Intestine |
| Nlrp4e | -3.57E-01 | 1.227387 | 0.006293 | 0.937883 | 0.969735 | Healthy Control | Intestine |
| Nlrp4f | 4.49E+00 | 1.321338 | 3.124357 | 0.094157 | 0.206731 | Infection | Intestine |
| Nlrp6 | -4.68E+00 | 9.621102 | 84.29139 | 2.20E-09 | 6.43E-08 | Healthy Control | Intestine |
| Nlrp9b | -8.56E+00 | 5.518477 | 8.192161 | 0.009696 | 0.032075 | Healthy Control | Intestine |
| Nmb | 1.96E+00 | 4.134077 | 5.540796 | 0.027284 | 0.076389 | Infection | Intestine |
| Nme5 | 2.12E+00 | 2.411747 | 0.991553 | 0.328441 | 0.545676 | Infection | Intestine |
| Nmrk1 | 1.91E+00 | 4.484905 | 8.587464 | 0.00758 | 0.026078 | Infection | Intestine |
| Nmt1 | -1.65E+00 | 7.792913 | 16.06866 | 0.000551 | 0.00276 | Healthy Control | Intestine |

| | | | | | | | |
|---|---|---|---|---|---|---|---|
| Nmu | -2.73E+00 | 1.763335 | 0.143454 | 0.708157 | 0.897011 | Healthy Control | Intestine |
| Nmur1 | 5.08E-01 | 3.006813 | 0.087658 | 0.769925 | 0.932779 | Infection | Intestine |
| Nobox | -3.57E-01 | 1.228165 | 0.006192 | 0.93838 | 0.969735 | Healthy Control | Intestine |
| Noct | -9.66E-01 | 6.694976 | 1.614405 | 0.215901 | 0.394453 | Healthy Control | Intestine |
| Nod1 | -4.25E-01 | 7.107002 | 1.015353 | 0.324231 | 0.540673 | Healthy Control | Intestine |
| Nod2 | 4.98E-01 | 3.06243 | 0.094863 | 0.760982 | 0.928591 | Infection | Intestine |
| Nodal | -9.77E-01 | 1.317674 | 0.022422 | 0.882312 | 0.96867 | Healthy Control | Intestine |
| Nog | -5.19E+00 | 2.73639 | 0.738478 | 0.40039 | 0.6303 | Healthy Control | Intestine |
| Nos1ap | -1.46E+00 | 4.927576 | 1.303842 | 0.265375 | 0.463849 | Healthy Control | Intestine |
| Nos2 | 1.83E+00 | 2.369761 | 1.04704 | 0.315297 | 0.528947 | Infection | Intestine |
| Nos3 | -1.28E+00 | 4.394585 | 0.183397 | 0.674121 | 0.879138 | Healthy Control | Intestine |
| Nosip | -9.53E-02 | 8.97194 | 0.265982 | 0.611235 | 0.833064 | Healthy Control | Intestine |
| Notch1 | 1.20E+00 | 7.847958 | 15.83772 | 0.000644 | 0.003163 | Infection | Intestine |
| Notch2 | 3.57E-01 | 7.536008 | 0.762705 | 0.392006 | 0.621597 | Infection | Intestine |
| Notch3 | -4.80E-01 | 5.979026 | 0.141849 | 0.710388 | 0.898485 | Healthy Control | Intestine |
| Notch4 | 6.29E-01 | 4.777206 | 0.355152 | 0.556729 | 0.784845 | Infection | Intestine |
| Noto | -8.80E-01 | 1.31755 | 0.021349 | 0.885117 | 0.96867 | Healthy Control | Intestine |
| Nova2 | -6.66E+00 | 3.697677 | 2.374501 | 0.139262 | 0.281106 | Healthy Control | Intestine |
| Nox1 | -1.00E+01 | 7.25669 | 5.913606 | 0.029325 | 0.080615 | Healthy Control | Intestine |
| Nox3 | -9.71E-16 | 1.180616 | 0 | 1 | 1 | Healthy Control | Intestine |
| Nox4 | 3.03E+00 | 2.535118 | 4.519981 | 0.043284 | 0.111346 | Infection | Intestine |
| Noxo1 | -4.36E+00 | 8.704404 | 43.12603 | 9.27E-07 | 1.05E-05 | Healthy Control | Intestine |
| Npas1 | -9.71E-16 | 1.180616 | 0 | 1 | 1 | Healthy Control | Intestine |
| Npas2 | -1.33E+00 | 7.594399 | 6.963155 | 0.01466 | 0.045414 | Healthy Control | Intestine |
| Npas4 | 5.99E-01 | 3.061367 | 0.108256 | 0.745199 | 0.923383 | Infection | Intestine |
| Npbwr1 | -3.84E+00 | 2.522711 | 0.154284 | 0.698415 | 0.893044 | Healthy Control | Intestine |
| Npc1 | -8.32E-01 | 7.57807 | 5.504969 | 0.028195 | 0.078126 | Healthy Control | Intestine |
| Npc1l1 | -2.77E+00 | 1.701285 | 0.172397 | 0.680978 | 0.882694 | Healthy Control | Intestine |
| Npepps | -2.96E-01 | 8.908165 | 2.003297 | 0.171101 | 0.329689 | Healthy Control | Intestine |
| Npffr2 | -2.40E+00 | 1.405885 | 0.066648 | 0.798419 | 0.94489 | Healthy Control | Intestine |
| Nphs1 | -6.42E-01 | 1.273434 | 0.012853 | 0.910919 | 0.969735 | Healthy Control | Intestine |
| Nphs2 | -1.83E+00 | 1.361056 | 0.049243 | 0.826155 | 0.953685 | Healthy Control | Intestine |
| Nploc4 | -3.31E-01 | 8.161768 | 1.489435 | 0.235351 | 0.42272 | Healthy Control | Intestine |
| Nppa | -2.38E+00 | 1.557903 | 0.095653 | 0.759118 | 0.928591 | Healthy Control | Intestine |
| Nppb | -1.65E+00 | 1.318024 | 0.050317 | 0.824565 | 0.953685 | Healthy Control | Intestine |
| Npr1 | 1.35E+00 | 4.396141 | 1.571788 | 0.222012 | 0.403702 | Infection | Intestine |
| Npsr1 | -1.58E+00 | 1.275596 | 0.039413 | 0.844957 | 0.954066 | Healthy Control | Intestine |
| Nptx2 | 1.78E+00 | 3.617324 | 2.815261 | 0.106529 | 0.227801 | Infection | Intestine |
| Nptxr | 2.72E+00 | 3.156921 | 6.200565 | 0.020119 | 0.059544 | Infection | Intestine |
| Npy1r | -6.21E-01 | 5.437066 | 0.277581 | 0.603315 | 0.826665 | Healthy Control | Intestine |
| Npy2r | -3.16E-02 | 3.970566 | 0.000281 | 0.986767 | 1 | Healthy Control | Intestine |
| Npy4r | 2.08E+00 | 2.03777 | 0.845641 | 0.365629 | 0.590623 | Infection | Intestine |
| Nqo1 | -1.78E+00 | 7.99076 | 11.46847 | 0.00258 | 0.010284 | Healthy Control | Intestine |
| Nr0b1 | -1.37E+00 | 1.229023 | 0.032546 | 0.859435 | 0.956553 | Healthy Control | Intestine |
| Nr0b2 | -2.44E+00 | 1.524014 | 0.090272 | 0.767566 | 0.932403 | Healthy Control | Intestine |
| Nr1d1 | -1.55E+00 | 8.942271 | 12.07399 | 0.002228 | 0.009062 | Healthy Control | Intestine |
| Nr1h2 | 6.12E-01 | 8.323712 | 6.81023 | 0.016076 | 0.049132 | Infection | Intestine |
| Nr1h3 | 9.12E-02 | 6.694052 | 0.048671 | 0.827378 | 0.953685 | Infection | Intestine |
| Nr1h4 | -4.34E+00 | 7.751588 | 42.35602 | 1.22E-06 | 1.32E-05 | Healthy Control | Intestine |
| Nr1i2 | -4.13E+00 | 7.628736 | 27.40404 | 2.71E-05 | 0.000197 | Healthy Control | Intestine |

| | | | | | | | |
|---|---|---|---|---|---|---|---|
| Nr1i3 | -2.00E+00 | 1.445311 | 0.050183 | 0.824431 | 0.953685 | Healthy Control | Intestine |
| Nr2c2 | -5.72E-01 | 7.650417 | 3.482572 | 0.075326 | 0.174252 | Healthy Control | Intestine |
| Nr2e1 | -7.56E-01 | 1.274694 | 0.014662 | 0.905006 | 0.969735 | Healthy Control | Intestine |
| Nr2e3 | -3.60E+00 | 2.010515 | 0.370746 | 0.547039 | 0.777534 | Healthy Control | Intestine |
| Nr2f1 | 2.98E+00 | 4.896479 | 22.52085 | 0.0001 | 0.000623 | Infection | Intestine |
| Nr2f2 | 8.20E-01 | 7.945375 | 10.16026 | 0.004293 | 0.016022 | Infection | Intestine |
| Nr3c1 | 6.21E-01 | 7.861807 | 6.876886 | 0.015634 | 0.048043 | Infection | Intestine |
| Nr3c2 | -2.44E+00 | 9.396306 | 58.04691 | 1.25E-07 | 1.90E-06 | Healthy Control | Intestine |
| Nr4a1 | -2.49E+00 | 5.62868 | 5.534074 | 0.027921 | 0.077694 | Healthy Control | Intestine |
| Nr4a2 | -1.70E+00 | 4.817037 | 1.801963 | 0.193201 | 0.361691 | Healthy Control | Intestine |
| Nr4a3 | 1.26E+00 | 3.910917 | 1.318008 | 0.262012 | 0.458774 | Infection | Intestine |
| Nr5a1 | -1.52E+00 | 1.274134 | 0.042767 | 0.838583 | 0.954066 | Healthy Control | Intestine |
| Nr5a2 | -2.07E+00 | 7.090447 | 8.893682 | 0.006454 | 0.022849 | Healthy Control | Intestine |
| Nr6a1 | -1.99E+00 | 5.278091 | 2.949714 | 0.099705 | 0.216258 | Healthy Control | Intestine |
| Nrarp | -8.03E-01 | 5.364122 | 0.504523 | 0.484281 | 0.71874 | Healthy Control | Intestine |
| Nras | 1.87E+00 | 6.681835 | 38.13711 | 3.39E-06 | 3.18E-05 | Infection | Intestine |
| Nrcam | 3.39E+00 | 2.461178 | 8.214831 | 0.008042 | 0.027386 | Infection | Intestine |
| Nrep | 2.72E+00 | 5.882527 | 44.37536 | 1.13E-06 | 1.25E-05 | Infection | Intestine |
| Nrf1 | -1.50E+00 | 6.80669 | 7.321983 | 0.012313 | 0.039386 | Healthy Control | Intestine |
| Nrg1 | 1.89E+00 | 4.620242 | 8.517578 | 0.007782 | 0.026629 | Infection | Intestine |
| Nrip1 | -2.83E+00 | 8.789879 | 45.84766 | 5.93E-07 | 7.21E-06 | Healthy Control | Intestine |
| Nrip3 | 2.48E+00 | 1.774152 | 1.550636 | 0.222411 | 0.404282 | Infection | Intestine |
| Nrl | -8.79E-01 | 1.316883 | 0.01296 | 0.910361 | 0.969735 | Healthy Control | Intestine |
| Nrn1 | -7.36E-02 | 3.581598 | 0.00235 | 0.961766 | 0.986194 | Healthy Control | Intestine |
| Nrp1 | 7.41E-01 | 7.58732 | 6.040553 | 0.022427 | 0.065083 | Infection | Intestine |
| Nrp2 | -1.63E+00 | 8.80039 | 14.78044 | 0.000946 | 0.004406 | Healthy Control | Intestine |
| Nsf | -6.80E-01 | 7.602143 | 4.988069 | 0.035887 | 0.095002 | Healthy Control | Intestine |
| Nsmce3 | 3.05E-01 | 4.022158 | 0.061889 | 0.805736 | 0.947947 | Infection | Intestine |
| Nsmf | -9.87E-01 | 8.966877 | 22.36463 | 0.000104 | 0.000642 | Healthy Control | Intestine |
| Nsun2 | -4.11E-02 | 7.632696 | 0.022398 | 0.882404 | 0.96867 | Healthy Control | Intestine |
| Nt5c1b | -1.99E+00 | 1.442682 | 0.075459 | 0.785639 | 0.940437 | Healthy Control | Intestine |
| Nt5e | -1.28E+00 | 7.531728 | 3.315135 | 0.082291 | 0.186741 | Healthy Control | Intestine |
| Ntf3 | 1.07E+00 | 4.655447 | 1.106024 | 0.303569 | 0.514269 | Infection | Intestine |
| Ntf5 | -3.22E+00 | 1.671079 | 0.160313 | 0.692798 | 0.890181 | Healthy Control | Intestine |
| Nthl1 | -1.16E+00 | 4.659415 | 0.681653 | 0.417618 | 0.650153 | Healthy Control | Intestine |
| Ntmt1 | -3.51E-01 | 6.029679 | 0.379334 | 0.543804 | 0.775484 | Healthy Control | Intestine |
| Ntn1 | 3.01E-02 | 7.629902 | 0.00225 | 0.96271 | 0.986758 | Infection | Intestine |
| Ntn4 | -2.15E+00 | 7.406333 | 8.307776 | 0.008122 | 0.027602 | Healthy Control | Intestine |
| Ntng1 | -5.01E+00 | 2.582811 | 0.590614 | 0.451065 | 0.684875 | Healthy Control | Intestine |
| Ntng2 | 8.35E-01 | 6.13554 | 2.075411 | 0.163737 | 0.318667 | Infection | Intestine |
| Ntrk1 | -1.89E+00 | 1.40206 | 0.063389 | 0.803109 | 0.947264 | Healthy Control | Intestine |
| Ntrk2 | -1.42E+00 | 4.84489 | 0.836768 | 0.369406 | 0.595164 | Healthy Control | Intestine |
| Ntrk3 | 3.05E+00 | 2.611653 | 6.300984 | 0.01862 | 0.055749 | Infection | Intestine |
| Nts | 2.35E+00 | 3.621677 | 2.531258 | 0.123203 | 0.254751 | Infection | Intestine |
| Ntsr1 | -8.79E-01 | 1.317006 | 0.022441 | 0.882238 | 0.96867 | Healthy Control | Intestine |
| Ntsr2 | 3.89E+00 | 1.48538 | 3.591698 | 0.066461 | 0.158354 | Infection | Intestine |
| Nuak2 | -3.17E-01 | 7.043928 | 0.593757 | 0.448902 | 0.682909 | Healthy Control | Intestine |
| Nubpl | 1.37E+00 | 2.749092 | 0.567288 | 0.458734 | 0.693155 | Infection | Intestine |
| Numb | 1.15E-01 | 7.685921 | 0.153056 | 0.699429 | 0.89357 | Infection | Intestine |
| Nup155 | -1.89E-01 | 5.82031 | 0.101324 | 0.753011 | 0.925708 | Healthy Control | Intestine |

| | | | | | | | |
|---|---|---|---|---|---|---|---|
| Nup205 | 1.71E-01 | 6.796522 | 0.17545 | 0.679346 | 0.882616 | Infection | Intestine |
| Nup62 | -1.47E+00 | 5.93166 | 2.293639 | 0.143186 | 0.28746 | Healthy Control | Intestine |
| Nup98 | -4.12E-01 | 7.595544 | 1.518123 | 0.230901 | 0.41729 | Healthy Control | Intestine |
| Nxn | 2.25E+00 | 5.586872 | 16.80768 | 0.000482 | 0.00246 | Infection | Intestine |
| Nxph2 | 6.69E+00 | 1.230035 | 8.855416 | 0.012839 | 0.040654 | Infection | Intestine |
| Nxph4 | -2.89E+00 | 1.522689 | 0.184609 | 0.670216 | 0.876692 | Healthy Control | Intestine |
| Nyap1 | 2.00E-02 | 4.51077 | 0.000366 | 0.984897 | 1 | Infection | Intestine |
| Nyap2 | -2.81E+00 | 1.484128 | 0.124705 | 0.72637 | 0.910874 | Healthy Control | Intestine |
| Oasl2 | -8.98E-01 | 7.371077 | 0.909053 | 0.354109 | 0.578548 | Healthy Control | Intestine |
| Oc90 | -6.42E-01 | 1.272688 | 0.013552 | 0.908542 | 0.969735 | Healthy Control | Intestine |
| Oca2 | -3.57E-01 | 1.228165 | 0.006192 | 0.93838 | 0.969735 | Healthy Control | Intestine |
| Ocln | -1.12E+01 | 8.344525 | 59.92619 | 2.06E-07 | 2.88E-06 | Healthy Control | Intestine |
| Ocstamp | 2.51E+00 | 2.002022 | 2.074072 | 0.160588 | 0.314025 | Infection | Intestine |
| Odam | -1.88E+00 | 1.361618 | 0.069323 | 0.794701 | 0.943769 | Healthy Control | Intestine |
| Odc1 | -9.87E-01 | 9.262925 | 31.76933 | 1.19E-05 | 9.63E-05 | Healthy Control | Intestine |
| Odf2 | -7.48E-01 | 7.452273 | 3.159467 | 0.089057 | 0.197879 | Healthy Control | Intestine |
| Odr4 | -5.20E-01 | 7.346448 | 1.779077 | 0.19565 | 0.365456 | Healthy Control | Intestine |
| Ofd1 | 3.39E-01 | 5.446566 | 0.194758 | 0.662998 | 0.870207 | Infection | Intestine |
| Ogdh | -2.95E+00 | 10.15691 | 170.1177 | 8.29E-12 | 6.44E-10 | Healthy Control | Intestine |
| Ogg1 | -6.70E-01 | 5.821577 | 1.011305 | 0.324416 | 0.540801 | Healthy Control | Intestine |
| Ogn | -7.26E-03 | 8.618108 | 0.000444 | 0.983376 | 1 | Healthy Control | Intestine |
| Ogt | 4.63E-01 | 8.716891 | 7.543938 | 0.01184 | 0.038092 | Infection | Intestine |
| Olfm3 | -1.25E+00 | 1.361871 | 0.034074 | 0.855068 | 0.956553 | Healthy Control | Intestine |
| Olfr17 | -1.37E+00 | 1.229023 | 0.032546 | 0.859435 | 0.956553 | Healthy Control | Intestine |
| Olig1 | -9.71E-16 | 1.180616 | 0 | 1 | 1 | Healthy Control | Intestine |
| Olig2 | -3.57E-01 | 1.227509 | 0.006273 | 0.937979 | 0.969735 | Healthy Control | Intestine |
| Olig3 | -1.52E+00 | 1.275042 | 0.038572 | 0.846592 | 0.954066 | Healthy Control | Intestine |
| Olr1 | 8.13E+00 | 1.855257 | 77.80385 | 5.61E-10 | 2.03E-08 | Infection | Intestine |
| Omp | -6.40E-01 | 5.30233 | 0.400873 | 0.532736 | 0.765904 | Healthy Control | Intestine |
| Onecut1 | -4.95E-01 | 1.228692 | 0.008683 | 0.927069 | 0.969735 | Healthy Control | Intestine |
| Onecut2 | -1.61E+00 | 1.443626 | 0.043208 | 0.836855 | 0.954066 | Healthy Control | Intestine |
| Onecut3 | -1.94E+00 | 1.403583 | 0.067629 | 0.796646 | 0.943769 | Healthy Control | Intestine |
| Opa1 | -5.94E-02 | 8.484561 | 0.071533 | 0.791635 | 0.943104 | Healthy Control | Intestine |
| Opalin | -3.57E-01 | 1.228299 | 0.00618 | 0.938438 | 0.969735 | Healthy Control | Intestine |
| Opn1mw | -2.53E+00 | 1.320325 | 0.15293 | 0.700969 | 0.89357 | Healthy Control | Intestine |
| Opn1sw | 5.48E+00 | 1.566971 | 11.01811 | 0.0021 | 0.008594 | Infection | Intestine |
| Opn3 | -6.21E+00 | 3.767194 | 2.174774 | 0.155985 | 0.306822 | Healthy Control | Intestine |
| Opn4 | -1.71E+00 | 1.31808 | 0.052703 | 0.820694 | 0.953035 | Healthy Control | Intestine |
| Oprd1 | -5.74E+00 | 3.002759 | 0.702879 | 0.411776 | 0.643971 | Healthy Control | Intestine |
| Oprk1 | -7.56E-01 | 1.273784 | 0.014872 | 0.904331 | 0.969735 | Healthy Control | Intestine |
| Oprl1 | -6.42E-01 | 1.272658 | 0.010513 | 0.919403 | 0.969735 | Healthy Control | Intestine |
| Oprm1 | -2.46E+00 | 1.596956 | 0.121375 | 0.729896 | 0.91327 | Healthy Control | Intestine |
| Orai1 | -7.89E+00 | 5.073138 | 6.862773 | 0.016499 | 0.050213 | Healthy Control | Intestine |
| Orc1 | 2.63E-01 | 4.730984 | 0.101194 | 0.753092 | 0.925708 | Infection | Intestine |
| Orm1 | -1.17E+00 | 1.361027 | 0.025017 | 0.875542 | 0.965198 | Healthy Control | Intestine |
| Osgep | -2.44E-01 | 7.47973 | 0.433248 | 0.517224 | 0.752681 | Healthy Control | Intestine |
| Osm | 3.78E+00 | 2.304886 | 7.918663 | 0.008771 | 0.029409 | Infection | Intestine |
| Osmr | 3.45E+00 | 7.041304 | 73.75042 | 1.73E-07 | 2.49E-06 | Infection | Intestine |
| Osr1 | 1.53E+00 | 6.834055 | 4.294389 | 0.052564 | 0.13033 | Infection | Intestine |
| Osr2 | -1.81E+00 | 6.745704 | 2.664014 | 0.117585 | 0.245914 | Healthy Control | Intestine |

| | | | | | | | |
|---|---|---|---|---|---|---|---|
| Otc | -8.94E+00 | 6.263417 | 11.57589 | 0.002856 | 0.011248 | Healthy Control | Intestine |
| Otof | -3.51E+00 | 2.097851 | 0.070759 | 0.793472 | 0.943769 | Healthy Control | Intestine |
| Otp | -9.71E-16 | 1.180616 | 0 | 1 | 1 | Healthy Control | Intestine |
| Otud5 | -3.06E-01 | 8.942939 | 1.809926 | 0.192362 | 0.360661 | Healthy Control | Intestine |
| Otud7b | -1.47E+00 | 7.337339 | 12.26733 | 0.001865 | 0.007788 | Healthy Control | Intestine |
| Otx1 | -2.04E+00 | 1.442381 | 0.093894 | 0.761365 | 0.928591 | Healthy Control | Intestine |
| Otx2 | 4.79E+00 | 1.446062 | 5.335029 | 0.028143 | 0.078126 | Infection | Intestine |
| Ovol1 | -8.93E+00 | 6.133393 | 11.48663 | 0.002943 | 0.011503 | Healthy Control | Intestine |
| Oxt | -9.71E-16 | 1.180616 | 0 | 1 | 1 | Healthy Control | Intestine |
| Oxtr | -5.89E+00 | 3.217782 | 0.64532 | 0.430904 | 0.663316 | Healthy Control | Intestine |
| P2rx3 | 2.30E+00 | 2.783177 | 2.06709 | 0.162843 | 0.317567 | Infection | Intestine |
| P2rx4 | -2.19E+00 | 9.193962 | 57.91829 | 1.24E-07 | 1.89E-06 | Healthy Control | Intestine |
| P2rx7 | -2.97E-01 | 6.728645 | 0.222836 | 0.641375 | 0.855379 | Healthy Control | Intestine |
| P2ry1 | -1.60E+00 | 6.450357 | 7.256936 | 0.012593 | 0.040102 | Healthy Control | Intestine |
| P2ry13 | -8.26E+00 | 5.609672 | 5.05809 | 0.036056 | 0.095299 | Healthy Control | Intestine |
| P2ry2 | 8.96E-01 | 5.449082 | 1.67585 | 0.208205 | 0.383188 | Infection | Intestine |
| P3h1 | 3.36E+00 | 5.504492 | 48.65093 | 5.64E-07 | 6.92E-06 | Infection | Intestine |
| P3h3 | 8.27E-02 | 6.262316 | 0.018285 | 0.893625 | 0.969735 | Infection | Intestine |
| P4hb | -7.77E-01 | 11.4591 | 51.07838 | 3.83E-07 | 4.94E-06 | Healthy Control | Intestine |
| Pa2g4 | -1.64E+00 | 8.257991 | 20.84779 | 0.000141 | 0.000829 | Healthy Control | Intestine |
| Pacsin3 | 4.08E-01 | 6.164545 | 0.496615 | 0.488211 | 0.721792 | Infection | Intestine |
| Padi2 | -5.28E+00 | 2.672893 | 0.849258 | 0.367775 | 0.593515 | Healthy Control | Intestine |
| Padi4 | 4.91E+00 | 3.214291 | 38.60993 | 1.91E-06 | 1.90E-05 | Infection | Intestine |
| Padi6 | -9.71E-16 | 1.180616 | 0 | 1 | 1 | Healthy Control | Intestine |
| Pafah1b1 | -2.84E-01 | 8.444347 | 0.679216 | 0.41879 | 0.650873 | Healthy Control | Intestine |
| Pafah1b2 | -8.04E-01 | 8.027914 | 7.870436 | 0.010259 | 0.033631 | Healthy Control | Intestine |
| Pafah1b3 | -2.65E+00 | 8.871763 | 76.41328 | 7.53E-09 | 1.79E-07 | Healthy Control | Intestine |
| Pah | 4.39E+00 | 1.445991 | 2.478875 | 0.125229 | 0.257876 | Infection | Intestine |
| Paip1 | 8.10E-01 | 9.427616 | 39.33232 | 2.71E-06 | 2.59E-05 | Infection | Intestine |
| Pak1 | -4.56E-01 | 9.388272 | 5.842683 | 0.024474 | 0.069908 | Healthy Control | Intestine |
| Pak2 | -8.93E-01 | 9.302893 | 21.80396 | 0.000121 | 0.000729 | Healthy Control | Intestine |
| Pak3 | -4.07E+00 | 2.106729 | 0.366279 | 0.550552 | 0.779871 | Healthy Control | Intestine |
| Pakap | 1.91E+00 | 6.28245 | 24.47832 | 6.07E-05 | 0.000397 | Infection | Intestine |
| Pald1 | 2.04E+00 | 4.374548 | 6.802037 | 0.015724 | 0.048203 | Infection | Intestine |
| Palmd | -8.27E+00 | 5.194252 | 2.870265 | 0.106917 | 0.228534 | Healthy Control | Intestine |
| Panx1 | -2.04E+00 | 6.939352 | 8.353758 | 0.007907 | 0.026964 | Healthy Control | Intestine |
| Panx2 | 3.63E+00 | 1.59768 | 2.020658 | 0.163714 | 0.318667 | Infection | Intestine |
| Panx3 | -1.89E+00 | 1.401404 | 0.052728 | 0.820099 | 0.953035 | Healthy Control | Intestine |
| Pard3 | -1.41E+00 | 8.893732 | 26.29447 | 3.91E-05 | 0.00027 | Healthy Control | Intestine |
| Pard6a | 9.75E-02 | 4.361386 | 0.007917 | 0.929837 | 0.969735 | Infection | Intestine |
| Pard6b | -2.28E+00 | 6.616652 | 7.314772 | 0.012376 | 0.039563 | Healthy Control | Intestine |
| Park7 | -1.36E+00 | 8.921867 | 17.25237 | 0.00042 | 0.002188 | Healthy Control | Intestine |
| Parp1 | -5.27E-01 | 7.990268 | 3.064158 | 0.093949 | 0.206367 | Healthy Control | Intestine |
| Parp2 | -4.10E-01 | 7.082543 | 0.8758 | 0.359303 | 0.583415 | Healthy Control | Intestine |
| Parpbp | -9.58E-01 | 4.372222 | 0.400224 | 0.533348 | 0.765939 | Healthy Control | Intestine |
| Parva | -4.82E-01 | 8.804619 | 7.186053 | 0.013729 | 0.042928 | Healthy Control | Intestine |
| Parvb | 8.48E-01 | 5.770441 | 0.957791 | 0.338598 | 0.558492 | Infection | Intestine |
| Patj | -2.93E+00 | 7.833164 | 26.88998 | 2.59E-05 | 0.000189 | Healthy Control | Intestine |
| Patz1 | 5.93E-01 | 5.759959 | 0.818476 | 0.375004 | 0.601497 | Infection | Intestine |
| Pawr | -8.92E+00 | 6.003225 | 14.34178 | 0.001173 | 0.005267 | Healthy Control | Intestine |

| | | | | | | | |
|---|---|---|---|---|---|---|---|
| Pax1 | -9.71E-16 | 1.180616 | 0 | 1 | 1 | Healthy Control | Intestine |
| Pax2 | -3.57E-01 | 1.228299 | 0.00618 | 0.938438 | 0.969735 | Healthy Control | Intestine |
| Pax3 | -1.37E+00 | 1.228863 | 0.033199 | 0.858049 | 0.956553 | Healthy Control | Intestine |
| Pax4 | -4.31E+00 | 2.572996 | 0.418184 | 0.524284 | 0.758326 | Healthy Control | Intestine |
| Pax5 | -4.49E+00 | 2.544271 | 0.146427 | 0.705897 | 0.896379 | Healthy Control | Intestine |
| Pax6 | -3.95E+00 | 2.330236 | 0.345123 | 0.561926 | 0.790168 | Healthy Control | Intestine |
| Pax7 | -9.77E-01 | 1.318156 | 0.022077 | 0.883214 | 0.96867 | Healthy Control | Intestine |
| Pax8 | -3.86E+00 | 1.89774 | 0.346424 | 0.562795 | 0.790783 | Healthy Control | Intestine |
| Pax9 | -2.81E+00 | 1.7925 | 0.081714 | 0.777214 | 0.935948 | Healthy Control | Intestine |
| Paxbp1 | 5.62E-01 | 8.291186 | 8.205185 | 0.009068 | 0.030229 | Infection | Intestine |
| Paxip1 | 5.17E-01 | 7.331024 | 3.074221 | 0.093588 | 0.205663 | Infection | Intestine |
| Pbrm1 | 9.49E-02 | 9.079873 | 0.317733 | 0.57873 | 0.806375 | Infection | Intestine |
| Pbx2 | -1.16E+00 | 7.805135 | 8.984321 | 0.006503 | 0.022989 | Healthy Control | Intestine |
| Pbx3 | -1.96E+00 | 5.236425 | 2.357693 | 0.138706 | 0.280324 | Healthy Control | Intestine |
| Pcbp3 | -7.43E-01 | 4.17493 | 0.284489 | 0.598993 | 0.823556 | Healthy Control | Intestine |
| Pcdh10 | -4.47E+00 | 2.82897 | 0.544561 | 0.467213 | 0.702974 | Healthy Control | Intestine |
| Pcdh11x | -2.87E+00 | 1.824435 | 0.174153 | 0.679859 | 0.882617 | Healthy Control | Intestine |
| Pcdh20 | -3.18E+00 | 1.740239 | 0.197301 | 0.662207 | 0.869624 | Healthy Control | Intestine |
| Pcdh8 | 4.70E+00 | 1.363202 | 3.669157 | 0.068168 | 0.161653 | Infection | Intestine |
| Pcdha2 | -2.36E+00 | 1.405436 | 0.07655 | 0.784097 | 0.940206 | Healthy Control | Intestine |
| Pcdha7 | -2.83E+00 | 1.664313 | 0.15734 | 0.694561 | 0.891531 | Healthy Control | Intestine |
| Pcgf1 | -4.93E-01 | 4.016149 | 0.097768 | 0.757397 | 0.928591 | Healthy Control | Intestine |
| Pcgf2 | -1.55E-01 | 7.567041 | 0.20007 | 0.659027 | 0.867142 | Healthy Control | Intestine |
| Pcif1 | -2.73E-01 | 6.641805 | 0.260085 | 0.61496 | 0.835865 | Healthy Control | Intestine |
| Pclaf | -8.01E-01 | 4.260261 | 0.216984 | 0.645785 | 0.858189 | Healthy Control | Intestine |
| Pcm1 | -1.01E+00 | 7.817463 | 7.325584 | 0.012747 | 0.04049 | Healthy Control | Intestine |
| Pcmtd2 | -1.09E+00 | 8.349091 | 14.00662 | 0.001126 | 0.005095 | Healthy Control | Intestine |
| Pcna | 2.02E+00 | 7.641584 | 72.55733 | 2.20E-08 | 4.36E-07 | Infection | Intestine |
| Pcnt | 8.11E-01 | 7.799768 | 7.998791 | 0.009849 | 0.03247 | Infection | Intestine |
| Pcp2 | -3.57E-01 | 1.227509 | 0.006273 | 0.937979 | 0.969735 | Healthy Control | Intestine |
| Pcsk1 | -3.90E+00 | 6.923531 | 14.05617 | 0.001119 | 0.005071 | Healthy Control | Intestine |
| Pcsk4 | -1.13E+00 | 4.592162 | 0.633338 | 0.434467 | 0.666751 | Healthy Control | Intestine |
| Pcsk6 | -1.61E+00 | 7.59424 | 7.060789 | 0.01411 | 0.043927 | Healthy Control | Intestine |
| Pcsk9 | -2.67E+00 | 5.709845 | 3.967217 | 0.059156 | 0.143406 | Healthy Control | Intestine |
| Pcx | -4.53E+00 | 8.869134 | 50.93668 | 3.19E-07 | 4.21E-06 | Healthy Control | Intestine |
| Pdap1 | -2.57E-01 | 8.427166 | 1.105359 | 0.30462 | 0.515702 | Healthy Control | Intestine |
| Pdc | -9.71E-16 | 1.180616 | 0 | 1 | 1 | Healthy Control | Intestine |
| Pdcd1 | -3.57E+00 | 1.734929 | 0.246789 | 0.622781 | 0.839656 | Healthy Control | Intestine |
| Pdcd1lg2 | 3.30E+00 | 3.477612 | 16.00688 | 0.000588 | 0.002917 | Infection | Intestine |
| Pdcd4 | -2.32E+00 | 10.78652 | 97.41113 | 1.69E-09 | 5.24E-08 | Healthy Control | Intestine |
| Pde10a | 2.97E+00 | 3.593415 | 18.53582 | 0.000276 | 0.001507 | Infection | Intestine |
| Pde12 | 3.37E-02 | 5.523815 | 0.001869 | 0.965872 | 0.988784 | Infection | Intestine |
| Pde1b | 1.95E+00 | 5.998853 | 22.04499 | 0.000113 | 0.000689 | Infection | Intestine |
| Pde1c | 2.21E-01 | 4.765458 | 0.05793 | 0.811857 | 0.950013 | Infection | Intestine |
| Pde3b | 2.61E+00 | 5.309865 | 39.55575 | 2.57E-06 | 2.47E-05 | Infection | Intestine |
| Pde4c | -2.43E-02 | 3.556499 | 0.000225 | 0.988158 | 1 | Healthy Control | Intestine |
| Pde6b | -9.71E-16 | 1.180616 | 0 | 1 | 1 | Healthy Control | Intestine |
| Pde6c | -2.06E+00 | 1.275767 | 0.073089 | 0.790362 | 0.942036 | Healthy Control | Intestine |
| Pde6g | -4.78E+00 | 2.522541 | 0.77701 | 0.388266 | 0.61782 | Healthy Control | Intestine |
| Pde7b | -1.79E+00 | 5.236514 | 2.13852 | 0.157253 | 0.308831 | Healthy Control | Intestine |

| | | | | | | | |
|---|---|---|---|---|---|---|---|
| Pde8a | 2.50E-01 | 6.652577 | 0.180185 | 0.675312 | 0.879372 | Infection | Intestine |
| Pde8b | -1.43E+00 | 6.65402 | 3.97909 | 0.057194 | 0.139462 | Healthy Control | Intestine |
| Pdgfa | -3.61E-01 | 7.258974 | 1.073621 | 0.311193 | 0.524347 | Healthy Control | Intestine |
| Pdgfb | -1.47E-01 | 5.703845 | 0.046649 | 0.830825 | 0.954066 | Healthy Control | Intestine |
| Pdgfra | 1.49E+00 | 8.396804 | 34.23321 | 7.15E-06 | 6.14E-05 | Infection | Intestine |
| Pdgfrb | 2.10E+00 | 8.573417 | 67.74655 | 9.63E-08 | 1.54E-06 | Infection | Intestine |
| Pdia3 | -5.55E-01 | 12.21067 | 27.81266 | 2.80E-05 | 0.000202 | Healthy Control | Intestine |
| Pdk1 | -2.86E-01 | 7.593571 | 0.573318 | 0.456712 | 0.690725 | Healthy Control | Intestine |
| Pdk4 | -1.72E-01 | 4.58138 | 0.021259 | 0.885316 | 0.96867 | Healthy Control | Intestine |
| Pdlim1 | -6.50E+00 | 3.694288 | 1.48093 | 0.238027 | 0.42571 | Healthy Control | Intestine |
| Pdlim4 | 1.79E+00 | 6.464421 | 16.62517 | 0.000508 | 0.002569 | Infection | Intestine |
| Pdpk1 | -7.89E-01 | 8.068365 | 8.593907 | 0.007711 | 0.026441 | Healthy Control | Intestine |
| Pdpn | 5.42E-01 | 6.979712 | 0.865337 | 0.362708 | 0.587498 | Infection | Intestine |
| Pdx1 | -6.51E+00 | 4.013822 | 1.859011 | 0.188046 | 0.354038 | Healthy Control | Intestine |
| Pdxk | -7.21E-01 | 6.965032 | 1.929258 | 0.177958 | 0.339791 | Healthy Control | Intestine |
| Pdyn | -2.30E+00 | 1.521429 | 0.093731 | 0.76146 | 0.928591 | Healthy Control | Intestine |
| Pdzd7 | -1.43E+00 | 7.13786 | 3.899306 | 0.061447 | 0.148171 | Healthy Control | Intestine |
| Pdzk1 | 4.43E+00 | 1.794481 | 6.13813 | 0.020538 | 0.060515 | Infection | Intestine |
| Pecam1 | 1.31E+00 | 7.946893 | 2.833347 | 0.117156 | 0.245288 | Infection | Intestine |
| Peg10 | 2.10E+00 | 2.734861 | 1.877455 | 0.18313 | 0.347517 | Infection | Intestine |
| Peg3 | 8.27E-01 | 4.90052 | 1.009287 | 0.325038 | 0.541581 | Infection | Intestine |
| Per1 | 2.46E-01 | 5.95346 | 0.105102 | 0.748653 | 0.924698 | Infection | Intestine |
| Per2 | -9.17E-01 | 5.060905 | 0.687043 | 0.415686 | 0.648465 | Healthy Control | Intestine |
| Per3 | 4.50E-01 | 5.435212 | 0.386187 | 0.540285 | 0.772763 | Infection | Intestine |
| Pex3 | 2.51E-01 | 6.689003 | 0.382788 | 0.542428 | 0.774941 | Infection | Intestine |
| Pfas | -4.35E-01 | 6.14982 | 0.648844 | 0.428506 | 0.661048 | Healthy Control | Intestine |
| Pfkfb1 | 2.00E+00 | 3.757728 | 3.45455 | 0.075429 | 0.174329 | Infection | Intestine |
| Pfkfb2 | -1.01E+00 | 7.807848 | 9.607427 | 0.005145 | 0.018732 | Healthy Control | Intestine |
| Pfkl | -5.47E-01 | 9.329299 | 4.298248 | 0.050186 | 0.125304 | Healthy Control | Intestine |
| Pfkp | 6.53E-01 | 9.499259 | 17.66367 | 0.000374 | 0.001973 | Infection | Intestine |
| Pfn4 | 3.27E+00 | 1.736677 | 2.030763 | 0.163269 | 0.318273 | Infection | Intestine |
| Pgam5 | -2.21E+00 | 7.827391 | 18.24281 | 0.000268 | 0.001472 | Healthy Control | Intestine |
| Pgf | 1.25E+00 | 3.910728 | 0.838604 | 0.368749 | 0.594404 | Infection | Intestine |
| Pgk1 | -1.86E+00 | 10.16087 | 78.17638 | 1.17E-08 | 2.54E-07 | Healthy Control | Intestine |
| Pgk2 | -3.57E-01 | 1.227387 | 0.006293 | 0.937883 | 0.969735 | Healthy Control | Intestine |
| Pgpep1 | -9.06E-01 | 6.340079 | 1.663512 | 0.209559 | 0.385256 | Healthy Control | Intestine |
| Pgr | -3.08E+00 | 1.79687 | 0.144925 | 0.706216 | 0.896379 | Healthy Control | Intestine |
| Phb | -3.87E+00 | 7.242325 | 21.14296 | 0.000129 | 0.000771 | Healthy Control | Intestine |
| Phb2 | 3.75E-02 | 10.0096 | 0.093653 | 0.762487 | 0.929438 | Infection | Intestine |
| Phc1 | -6.79E-01 | 6.483477 | 0.954744 | 0.338487 | 0.558492 | Healthy Control | Intestine |
| Phc2 | 4.24E-01 | 7.738896 | 2.997721 | 0.097515 | 0.212151 | Infection | Intestine |
| Phex | -1.41E+00 | 1.403795 | 0.042964 | 0.837282 | 0.954066 | Healthy Control | Intestine |
| Phf10 | 4.22E-01 | 8.35194 | 4.483639 | 0.045882 | 0.117078 | Infection | Intestine |
| Phf3 | -5.77E-03 | 6.423048 | 0.000141 | 0.990619 | 1 | Healthy Control | Intestine |
| Phf6 | 2.68E-01 | 6.709929 | 0.369399 | 0.549488 | 0.778975 | Infection | Intestine |
| Phlda2 | 2.23E+00 | 2.43324 | 0.888718 | 0.354025 | 0.578548 | Infection | Intestine |
| Phox2a | -2.57E+00 | 1.484578 | 0.159665 | 0.692378 | 0.890099 | Healthy Control | Intestine |
| Phox2b | 1.85E+00 | 2.062715 | 0.821572 | 0.37266 | 0.598504 | Infection | Intestine |
| Phtf1 | 1.32E+00 | 6.816065 | 13.41876 | 0.001385 | 0.006056 | Infection | Intestine |
| Phykpl | 8.41E-01 | 5.95683 | 3.073484 | 0.093063 | 0.204946 | Infection | Intestine |

| | | | | | | | |
|---|---|---|---|---|---|---|---|
| Pi4ka | -4.69E-01 | 8.618145 | 4.720193 | 0.040968 | 0.106153 | Healthy Control | Intestine |
| Pianp | 2.12E+00 | 6.149773 | 26.59561 | 3.72E-05 | 0.000259 | Infection | Intestine |
| Pias1 | 4.05E-01 | 7.083713 | 1.441743 | 0.242675 | 0.432171 | Infection | Intestine |
| Pias2 | -5.25E-02 | 8.485365 | 0.055185 | 0.816464 | 0.952516 | Healthy Control | Intestine |
| Pias3 | -6.57E-01 | 6.343272 | 1.221891 | 0.28008 | 0.483309 | Healthy Control | Intestine |
| Picalm | -3.25E-01 | 10.01827 | 5.318463 | 0.030996 | 0.084375 | Healthy Control | Intestine |
| Pick1 | -1.94E+00 | 8.139899 | 24.32473 | 5.44E-05 | 0.000361 | Healthy Control | Intestine |
| Piezo1 | -9.13E-02 | 6.039571 | 0.022323 | 0.882509 | 0.96867 | Healthy Control | Intestine |
| Piezo2 | 4.38E+00 | 4.628916 | 123.8905 | 1.91E-10 | 8.54E-09 | Infection | Intestine |
| Pifo | -1.83E+00 | 1.361458 | 0.057213 | 0.812869 | 0.950244 | Healthy Control | Intestine |
| Pik3c3 | -1.96E+00 | 8.504214 | 29.00618 | 1.81E-05 | 0.00014 | Healthy Control | Intestine |
| Pik3ca | -2.33E-01 | 7.087173 | 0.37267 | 0.547677 | 0.778138 | Healthy Control | Intestine |
| Pik3cb | 1.24E+00 | 6.435082 | 9.301338 | 0.005909 | 0.021115 | Infection | Intestine |
| Pik3cd | 1.60E+00 | 5.86968 | 7.990686 | 0.009863 | 0.032485 | Infection | Intestine |
| Pik3cg | -2.66E+00 | 6.727529 | 9.611889 | 0.005023 | 0.018341 | Healthy Control | Intestine |
| Pik3r1 | -1.06E+00 | 9.601506 | 21.19967 | 0.000141 | 0.00083 | Healthy Control | Intestine |
| Pik3r2 | -1.51E+00 | 8.633724 | 34.24356 | 6.59E-06 | 5.74E-05 | Healthy Control | Intestine |
| Pim1 | 3.28E+00 | 3.902882 | 24.52362 | 5.88E-05 | 0.000387 | Infection | Intestine |
| Pim2 | 6.79E-01 | 5.729699 | 1.0992 | 0.305362 | 0.516433 | Infection | Intestine |
| Pin1 | 2.72E+00 | 2.300879 | 4.323596 | 0.047718 | 0.120323 | Infection | Intestine |
| Pink1 | 4.17E-02 | 8.522077 | 0.028208 | 0.868171 | 0.960668 | Infection | Intestine |
| Pinlyp | -2.65E+00 | 1.559349 | 0.132714 | 0.718072 | 0.903864 | Healthy Control | Intestine |
| Pip | 6.49E+00 | 1.277874 | 12.12344 | 0.005264 | 0.019124 | Infection | Intestine |
| Pip4k2c | -3.60E-02 | 8.454111 | 0.028227 | 0.868128 | 0.960668 | Healthy Control | Intestine |
| Pip4p1 | -2.42E-01 | 7.975829 | 0.68924 | 0.415426 | 0.648463 | Healthy Control | Intestine |
| Pip5k1c | -9.63E-01 | 7.755509 | 6.519597 | 0.018007 | 0.054304 | Healthy Control | Intestine |
| Pirb | 3.51E+00 | 5.79488 | 86.03212 | 5.12E-09 | 1.28E-07 | Infection | Intestine |
| Pisd | -2.87E-01 | 8.529921 | 1.360469 | 0.256079 | 0.451066 | Healthy Control | Intestine |
| Pithd1 | -3.10E+00 | 6.280326 | 8.854227 | 0.006923 | 0.024169 | Healthy Control | Intestine |
| Pitpnm1 | 1.18E+00 | 7.308722 | 13.44342 | 0.001373 | 0.006017 | Infection | Intestine |
| Pitrm1 | 2.56E-01 | 7.833309 | 0.910529 | 0.350444 | 0.574247 | Infection | Intestine |
| Pitx1 | 2.78E+00 | 7.514853 | 62.58757 | 2.55E-07 | 3.45E-06 | Infection | Intestine |
| Pitx2 | 1.34E+00 | 2.801029 | 0.180747 | 0.674643 | 0.879138 | Infection | Intestine |
| Piwil1 | 6.00E+00 | 1.275016 | 5.60432 | 0.031838 | 0.086058 | Infection | Intestine |
| Piwil2 | 2.48E+00 | 2.084299 | 1.210033 | 0.280425 | 0.483738 | Infection | Intestine |
| Piwil4 | -1.57E+00 | 4.650083 | 1.480494 | 0.236756 | 0.424245 | Healthy Control | Intestine |
| Pja1 | -6.74E-01 | 7.824553 | 4.020345 | 0.057341 | 0.139751 | Healthy Control | Intestine |
| Pja2 | 1.11E-01 | 8.405401 | 0.263791 | 0.612702 | 0.833799 | Infection | Intestine |
| Pkd1 | 1.29E+00 | 8.136285 | 33.26776 | 8.70E-06 | 7.34E-05 | Infection | Intestine |
| Pkd2 | 2.18E+00 | 7.953047 | 86.58753 | 4.78E-09 | 1.21E-07 | Infection | Intestine |
| Pkd2l1 | -1.78E+00 | 1.360984 | 0.060162 | 0.808174 | 0.94887 | Healthy Control | Intestine |
| Pkia | -1.79E-01 | 4.754974 | 0.027737 | 0.869087 | 0.960878 | Healthy Control | Intestine |
| Pkib | 1.47E+00 | 4.609706 | 3.245091 | 0.084507 | 0.190045 | Infection | Intestine |
| Pkig | -3.56E-01 | 6.897733 | 0.778783 | 0.386889 | 0.616217 | Healthy Control | Intestine |
| Pklr | -3.41E+00 | 7.763434 | 27.08183 | 2.46E-05 | 0.000181 | Healthy Control | Intestine |
| Pkm | -8.65E-01 | 9.765933 | 16.41766 | 0.00054 | 0.002708 | Healthy Control | Intestine |
| Pkn1 | 9.21E-01 | 8.452525 | 24.34497 | 6.35E-05 | 0.000415 | Infection | Intestine |
| Pknox2 | 1.29E+00 | 5.046644 | 2.614805 | 0.11962 | 0.249092 | Infection | Intestine |
| Pkp4 | -3.23E+00 | 8.042187 | 37.57055 | 2.21E-06 | 2.16E-05 | Healthy Control | Intestine |
| Pla2g10 | -1.18E+01 | 8.940201 | 75.02234 | 3.61E-08 | 6.70E-07 | Healthy Control | Intestine |

| | | | | | | | |
|---|---|---|---|---|---|---|---|
| Pla2g2e | 4.31E+00 | 2.587797 | 20.05043 | 0.000143 | 0.000841 | Infection | Intestine |
| Pla2g2f | -9.37E+00 | 6.490027 | 11.23459 | 0.0031 | 0.012048 | Healthy Control | Intestine |
| Pla2g4a | -6.53E-01 | 8.816448 | 5.788276 | 0.025068 | 0.071221 | Healthy Control | Intestine |
| Pla2g6 | -1.18E+00 | 5.536936 | 1.731338 | 0.200344 | 0.372142 | Healthy Control | Intestine |
| Pla2r1 | 1.59E+00 | 6.49088 | 15.59131 | 0.000695 | 0.003364 | Infection | Intestine |
| Plaat1 | -3.85E+00 | 1.860842 | 0.322645 | 0.574503 | 0.802221 | Healthy Control | Intestine |
| Plac9a | 1.35E+00 | 2.870008 | 0.618804 | 0.439022 | 0.672093 | Infection | Intestine |
| Plagl1 | 3.97E-01 | 3.090738 | 0.052339 | 0.821103 | 0.953035 | Infection | Intestine |
| Plagl2 | -1.58E+00 | 8.168778 | 15.13782 | 0.000738 | 0.003533 | Healthy Control | Intestine |
| Plau | -2.69E-01 | 8.74584 | 0.449501 | 0.51084 | 0.745993 | Healthy Control | Intestine |
| Plaur | 1.91E+00 | 5.440388 | 13.45725 | 0.001355 | 0.005957 | Infection | Intestine |
| Plcb2 | 2.81E+00 | 5.057207 | 21.87091 | 0.000118 | 0.00072 | Infection | Intestine |
| Plcb4 | -2.73E-02 | 6.442993 | 0.002096 | 0.963881 | 0.987554 | Healthy Control | Intestine |
| Plcg1 | 1.34E+00 | 7.233878 | 21.82542 | 0.00012 | 0.000728 | Infection | Intestine |
| Plcg2 | 1.38E+00 | 5.767219 | 5.003697 | 0.035598 | 0.094387 | Infection | Intestine |
| Pld1 | -2.28E+00 | 8.371566 | 45.77254 | 6.44E-07 | 7.73E-06 | Healthy Control | Intestine |
| Pld2 | 6.31E-01 | 4.826896 | 0.59218 | 0.449032 | 0.682909 | Infection | Intestine |
| Pld5 | -3.55E+00 | 1.703605 | 0.275329 | 0.603413 | 0.826665 | Healthy Control | Intestine |
| Plec | -2.65E-01 | 11.25948 | 5.183104 | 0.032983 | 0.088486 | Healthy Control | Intestine |
| Plekha1 | -3.10E-01 | 8.277149 | 1.132901 | 0.298803 | 0.507645 | Healthy Control | Intestine |
| Plekha2 | -1.12E+00 | 7.698013 | 8.408882 | 0.008106 | 0.027568 | Healthy Control | Intestine |
| Plekha5 | 5.75E-01 | 8.519486 | 9.235485 | 0.00607 | 0.021612 | Infection | Intestine |
| Plekhg2 | -7.49E-02 | 7.278552 | 0.053923 | 0.818513 | 0.952901 | Healthy Control | Intestine |
| Plekhg4 | -3.56E+00 | 1.857854 | 0.262668 | 0.612336 | 0.833657 | Healthy Control | Intestine |
| Plg | -8.61E-01 | 1.274702 | 0.021049 | 0.886432 | 0.96867 | Healthy Control | Intestine |
| Plgrkt | -1.20E+00 | 8.28527 | 8.732928 | 0.007271 | 0.025172 | Healthy Control | Intestine |
| Plin1 | -2.46E+00 | 1.630945 | 0.102991 | 0.750779 | 0.92508 | Healthy Control | Intestine |
| Plin2 | -2.27E+00 | 8.975425 | 49.85769 | 3.71E-07 | 4.81E-06 | Healthy Control | Intestine |
| Plin5 | -3.63E+00 | 2.003446 | 0.401207 | 0.532042 | 0.765638 | Healthy Control | Intestine |
| Plk1 | -5.64E-01 | 5.447803 | 0.25158 | 0.620394 | 0.838493 | Healthy Control | Intestine |
| Plk2 | -1.66E+00 | 7.706162 | 17.0518 | 0.000369 | 0.001953 | Healthy Control | Intestine |
| Plk3 | 1.40E+00 | 6.429021 | 7.165989 | 0.013778 | 0.043055 | Infection | Intestine |
| Plk4 | -7.09E+00 | 4.343754 | 2.531873 | 0.127428 | 0.261329 | Healthy Control | Intestine |
| Pln | 5.77E-01 | 4.724522 | 0.089439 | 0.767877 | 0.932538 | Infection | Intestine |
| Plod1 | 4.07E-01 | 7.360947 | 0.759837 | 0.39288 | 0.622391 | Infection | Intestine |
| Plod3 | 8.18E-01 | 7.806184 | 14.92684 | 0.000854 | 0.00402 | Infection | Intestine |
| Plp2 | -2.10E+00 | 10.03792 | 77.01474 | 1.15E-08 | 2.51E-07 | Healthy Control | Intestine |
| Plpp3 | -1.12E+00 | 9.260356 | 6.595398 | 0.019679 | 0.058418 | Healthy Control | Intestine |
| Plppr3 | 1.43E-01 | 4.254951 | 0.011806 | 0.914364 | 0.969735 | Infection | Intestine |
| Plppr4 | -4.69E+00 | 2.274307 | 0.401275 | 0.532997 | 0.765939 | Healthy Control | Intestine |
| Pls1 | -4.14E+00 | 9.779334 | 122.2962 | 7.20E-11 | 3.96E-09 | Healthy Control | Intestine |
| Pltp | -1.65E-01 | 9.583969 | 0.280731 | 0.602633 | 0.826406 | Healthy Control | Intestine |
| Plvap | -1.43E+00 | 7.5651 | 0.631012 | 0.440513 | 0.673346 | Healthy Control | Intestine |
| Plxdc2 | 1.05E+00 | 5.260787 | 1.730642 | 0.201462 | 0.373694 | Infection | Intestine |
| Plxna1 | 1.01E+00 | 6.210646 | 6.0366 | 0.022363 | 0.064972 | Infection | Intestine |
| Plxna2 | -1.79E+00 | 5.964027 | 5.328729 | 0.030013 | 0.082056 | Healthy Control | Intestine |
| Plxna3 | -8.33E+00 | 5.635957 | 8.531402 | 0.00851 | 0.028707 | Healthy Control | Intestine |
| Plxna4 | 2.73E-01 | 4.046582 | 0.033905 | 0.855462 | 0.956553 | Infection | Intestine |
| Plxnb2 | -4.09E-01 | 10.10035 | 5.589312 | 0.027406 | 0.076687 | Healthy Control | Intestine |
| Plxnc1 | 4.70E-02 | 4.225131 | 0.001135 | 0.973407 | 0.99487 | Infection | Intestine |

| | | | | | | | |
|---|---|---|---|---|---|---|---|
| Plxnd1 | 2.63E+00 | 6.628985 | 66.75872 | 4.46E-08 | 8.09E-07 | Infection | Intestine |
| Pmaip1 | -2.50E-01 | 5.644988 | 0.105794 | 0.747908 | 0.924698 | Healthy Control | Intestine |
| Pmch | -1.78E+00 | 1.361253 | 0.04884 | 0.826829 | 0.953685 | Healthy Control | Intestine |
| Pml | -6.68E-01 | 8.007229 | 3.85419 | 0.062432 | 0.150329 | Healthy Control | Intestine |
| Pmm2 | -1.98E+00 | 8.324394 | 29.65853 | 1.59E-05 | 0.000125 | Healthy Control | Intestine |
| Pms1 | -2.17E-01 | 4.765285 | 0.036878 | 0.849287 | 0.954543 | Healthy Control | Intestine |
| Pms2 | 1.58E+00 | 3.026676 | 1.251092 | 0.274557 | 0.476092 | Infection | Intestine |
| Pnkd | 5.34E-01 | 3.134797 | 0.079551 | 0.780509 | 0.938784 | Infection | Intestine |
| Pnoc | -1.65E+00 | 1.317863 | 0.048881 | 0.827044 | 0.953685 | Healthy Control | Intestine |
| Pnpla1 | -7.26E+00 | 4.835737 | 3.889263 | 0.062754 | 0.151033 | Healthy Control | Intestine |
| Pnpla2 | -1.39E+00 | 9.643265 | 47.70998 | 6.54E-07 | 7.80E-06 | Healthy Control | Intestine |
| Pnpla3 | -4.76E+00 | 2.52454 | 0.544843 | 0.468838 | 0.703959 | Healthy Control | Intestine |
| Podxl | -5.45E-01 | 7.903684 | 0.224062 | 0.644073 | 0.857095 | Healthy Control | Intestine |
| Polb | -1.66E+00 | 6.861155 | 8.609899 | 0.007188 | 0.024937 | Healthy Control | Intestine |
| Polg | 1.02E-01 | 8.678547 | 0.259269 | 0.615744 | 0.836703 | Infection | Intestine |
| Polk | -1.54E+00 | 6.346904 | 5.876316 | 0.023113 | 0.066804 | Healthy Control | Intestine |
| Polr1e | -1.13E+00 | 6.070591 | 2.767434 | 0.109078 | 0.232059 | Healthy Control | Intestine |
| Polr2a | -2.23E+00 | 9.124715 | 53.04376 | 2.56E-07 | 3.45E-06 | Healthy Control | Intestine |
| Polr2b | 4.94E-01 | 7.451998 | 3.017308 | 0.096467 | 0.210437 | Infection | Intestine |
| Polr2e | -1.61E+00 | 8.522312 | 35.29032 | 5.43E-06 | 4.79E-05 | Healthy Control | Intestine |
| Polr2f | 6.32E-02 | 8.270778 | 0.052987 | 0.820095 | 0.953035 | Infection | Intestine |
| Polr2h | -1.16E+00 | 6.932457 | 5.796896 | 0.02419 | 0.069275 | Healthy Control | Intestine |
| Polr2j | -1.74E+00 | 7.14973 | 10.04491 | 0.004133 | 0.015499 | Healthy Control | Intestine |
| Polr3e | 9.33E-01 | 6.085796 | 3.61581 | 0.070298 | 0.164908 | Infection | Intestine |
| Pom121 | -4.80E-01 | 7.096902 | 1.499904 | 0.233326 | 0.419702 | Healthy Control | Intestine |
| Pomc | 1.18E+00 | 2.858037 | 0.414575 | 0.525897 | 0.759561 | Infection | Intestine |
| Pon1 | -2.27E+00 | 1.554885 | 0.08794 | 0.769348 | 0.932758 | Healthy Control | Intestine |
| Pon3 | 5.53E-02 | 8.447432 | 0.057514 | 0.812713 | 0.950244 | Infection | Intestine |
| Por | -5.41E-02 | 7.704226 | 0.023773 | 0.878884 | 0.967178 | Healthy Control | Intestine |
| Postn | 6.38E-01 | 9.490873 | 2.164298 | 0.163618 | 0.318667 | Infection | Intestine |
| Pou1f1 | -7.56E-01 | 1.27364 | 0.014714 | 0.904837 | 0.969735 | Healthy Control | Intestine |
| Pou2af1 | -3.28E+00 | 6.448083 | 0.85885 | 0.37116 | 0.59686 | Healthy Control | Intestine |
| Pou2f1 | -1.44E+00 | 6.451736 | 5.131481 | 0.032644 | 0.087669 | Healthy Control | Intestine |
| Pou2f3 | -6.68E+00 | 3.514309 | 1.523763 | 0.231544 | 0.417689 | Healthy Control | Intestine |
| Pou3f1 | -8.61E-01 | 1.274141 | 0.016052 | 0.900735 | 0.969735 | Healthy Control | Intestine |
| Pou3f2 | -1.37E+00 | 1.228863 | 0.033199 | 0.858049 | 0.956553 | Healthy Control | Intestine |
| Pou3f3 | -3.56E+00 | 1.601122 | 0.233258 | 0.632714 | 0.848478 | Healthy Control | Intestine |
| Pou3f4 | -3.57E-01 | 1.228165 | 0.006192 | 0.93838 | 0.969735 | Healthy Control | Intestine |
| Pou4f1 | -3.48E+00 | 1.891923 | 0.287365 | 0.59617 | 0.82193 | Healthy Control | Intestine |
| Pou4f2 | -1.52E+00 | 1.275042 | 0.038572 | 0.846592 | 0.954066 | Healthy Control | Intestine |
| Pou4f3 | 4.04E+00 | 1.559978 | 3.068404 | 0.088892 | 0.197776 | Infection | Intestine |
| Pou5f1 | -4.95E-01 | 1.228515 | 0.008877 | 0.926263 | 0.969735 | Healthy Control | Intestine |
| Pou6f1 | 1.32E-01 | 6.387841 | 0.053238 | 0.819593 | 0.953035 | Infection | Intestine |
| Pou6f2 | -1.65E+00 | 1.318249 | 0.049681 | 0.825659 | 0.953685 | Healthy Control | Intestine |
| Ppara | -3.45E+00 | 6.932232 | 18.17284 | 0.00029 | 0.00158 | Healthy Control | Intestine |
| Pparg | -2.86E+00 | 8.168828 | 37.76096 | 2.31E-06 | 2.24E-05 | Healthy Control | Intestine |
| Ppargc1a | -4.22E+00 | 7.666714 | 25.33078 | 4.48E-05 | 0.000306 | Healthy Control | Intestine |
| Ppargc1b | -1.82E+00 | 9.186393 | 40.17295 | 2.27E-06 | 2.21E-05 | Healthy Control | Intestine |
| Ppfibp1 | 1.46E+00 | 7.995929 | 46.76002 | 7.57E-07 | 8.84E-06 | Infection | Intestine |
| Ppif | -4.44E+00 | 9.778398 | 108.0835 | 1.96E-10 | 8.68E-09 | Healthy Control | Intestine |

| | | | | | | | |
|---|---|---|---|---|---|---|---|
| Ppl | -8.19E+00 | 5.653295 | 6.902053 | 0.016232 | 0.049548 | Healthy Control | Intestine |
| Ppm1d | 1.21E+00 | 4.813914 | 2.416709 | 0.133557 | 0.271672 | Infection | Intestine |
| Ppm1h | -1.28E+00 | 7.320725 | 8.736748 | 0.007017 | 0.024458 | Healthy Control | Intestine |
| Ppp1cc | -2.20E+00 | 9.249503 | 79.24307 | 7.87E-09 | 1.83E-07 | Healthy Control | Intestine |
| Ppp1r10 | -1.08E+00 | 8.373531 | 9.010716 | 0.006593 | 0.023225 | Healthy Control | Intestine |
| Ppp1r16b | -4.43E-01 | 5.543302 | 0.146487 | 0.705188 | 0.896371 | Healthy Control | Intestine |
| Ppp1r1b | -4.89E+00 | 10.5474 | 225.7562 | 1.54E-13 | 2.41E-11 | Healthy Control | Intestine |
| Ppp1r1c | -1.25E+00 | 1.360602 | 0.037471 | 0.848104 | 0.954066 | Healthy Control | Intestine |
| Ppp1r2 | 4.70E-01 | 7.962016 | 2.921913 | 0.101607 | 0.219717 | Infection | Intestine |
| Ppp1r3a | -9.71E-16 | 1.180616 | 0 | 1 | 1 | Healthy Control | Intestine |
| Ppp1r7 | -1.88E+00 | 7.777545 | 15.13293 | 0.000721 | 0.00347 | Healthy Control | Intestine |
| Ppp2ca | -9.06E-01 | 10.22164 | 30.80533 | 1.45E-05 | 0.000115 | Healthy Control | Intestine |
| Ppp2r1a | -1.76E+00 | 9.431121 | 58.39061 | 1.32E-07 | 2.00E-06 | Healthy Control | Intestine |
| Ppp2r1b | -2.56E+00 | 6.606903 | 11.07894 | 0.002902 | 0.011405 | Healthy Control | Intestine |
| Ppp2r2a | -1.69E+00 | 6.032469 | 4.020008 | 0.056537 | 0.138275 | Healthy Control | Intestine |
| Ppp2r2b | 4.67E-01 | 4.010279 | 0.098381 | 0.75652 | 0.928448 | Infection | Intestine |
| Ppp2r5c | -1.07E+00 | 9.698557 | 27.23051 | 3.21E-05 | 0.000228 | Healthy Control | Intestine |
| Ppp3cb | -2.08E+00 | 8.531843 | 47.82972 | 4.96E-07 | 6.15E-06 | Healthy Control | Intestine |
| Ppp3r1 | -1.37E+00 | 8.117951 | 14.84171 | 0.000834 | 0.00394 | Healthy Control | Intestine |
| Ppp3r2 | -1.07E+00 | 1.318776 | 0.021583 | 0.884702 | 0.96867 | Healthy Control | Intestine |
| Ppp4r3a | 6.20E-01 | 9.058429 | 13.19903 | 0.001486 | 0.006425 | Infection | Intestine |
| Ppp5c | -1.98E+00 | 9.381086 | 71.93797 | 2.15E-08 | 4.27E-07 | Healthy Control | Intestine |
| Ppp6c | -7.91E-01 | 8.630968 | 10.28299 | 0.004097 | 0.01541 | Healthy Control | Intestine |
| Ppp6r3 | -1.42E+00 | 8.718542 | 17.15374 | 0.000428 | 0.002223 | Healthy Control | Intestine |
| Ppt1 | -1.26E+00 | 9.490672 | 20.38173 | 0.000175 | 0.00101 | Healthy Control | Intestine |
| Praf2 | 8.94E-01 | 6.673135 | 4.021463 | 0.057463 | 0.13998 | Infection | Intestine |
| Prap1 | -6.93E+00 | 4.644048 | 3.280159 | 0.082499 | 0.187043 | Healthy Control | Intestine |
| Prdm1 | 7.16E-03 | 5.441363 | 4.00E-05 | 0.995008 | 1 | Infection | Intestine |
| Prdm13 | -6.42E-01 | 1.272658 | 0.010513 | 0.919403 | 0.969735 | Healthy Control | Intestine |
| Prdm14 | -3.46E+00 | 2.304038 | 0.248589 | 0.624996 | 0.841282 | Healthy Control | Intestine |
| Prdm16 | 9.24E-01 | 3.001189 | 0.240597 | 0.628392 | 0.843445 | Infection | Intestine |
| Prdm9 | -4.21E-01 | 5.532104 | 0.276739 | 0.603753 | 0.826692 | Healthy Control | Intestine |
| Preb | -1.11E+00 | 8.94169 | 23.95882 | 6.96E-05 | 0.000451 | Healthy Control | Intestine |
| Prelp | 3.69E-02 | 8.073556 | 0.004167 | 0.949323 | 0.97748 | Infection | Intestine |
| Prep | -1.29E+00 | 8.163875 | 21.91171 | 0.000111 | 0.000678 | Healthy Control | Intestine |
| Prf1 | -3.57E+00 | 1.803076 | 0.229209 | 0.635657 | 0.851056 | Healthy Control | Intestine |
| Prickle3 | -1.94E-01 | 5.936383 | 0.08581 | 0.772115 | 0.934096 | Healthy Control | Intestine |
| Primpol | -5.37E-02 | 6.31975 | 0.009119 | 0.924745 | 0.969735 | Healthy Control | Intestine |
| Prkaa1 | -1.37E+00 | 8.644456 | 26.27886 | 3.97E-05 | 0.000274 | Healthy Control | Intestine |
| Prkaa2 | -2.56E+00 | 9.23412 | 70.84181 | 2.28E-08 | 4.49E-07 | Healthy Control | Intestine |
| Prkab1 | -1.66E+00 | 9.808126 | 80.11795 | 9.50E-09 | 2.12E-07 | Healthy Control | Intestine |
| Prkaca | -5.08E-01 | 8.97963 | 5.421637 | 0.029582 | 0.081233 | Healthy Control | Intestine |
| Prkacb | -1.19E-01 | 8.497951 | 0.150359 | 0.701881 | 0.894505 | Healthy Control | Intestine |
| Prkar1a | 4.54E-02 | 9.717099 | 0.097372 | 0.757971 | 0.928591 | Infection | Intestine |
| Prkar2b | 9.25E-01 | 7.441362 | 5.39705 | 0.029771 | 0.081617 | Infection | Intestine |
| Prkca | 2.66E-02 | 7.329566 | 0.004702 | 0.945956 | 0.975194 | Infection | Intestine |
| Prkcd | -7.74E-01 | 10.17707 | 21.13899 | 0.000143 | 0.000841 | Healthy Control | Intestine |
| Prkce | -1.02E+00 | 7.597464 | 6.218834 | 0.020479 | 0.060376 | Healthy Control | Intestine |
| Prkcg | 3.26E-01 | 4.002434 | 0.066984 | 0.798051 | 0.944678 | Infection | Intestine |
| Prkci | -2.23E+00 | 8.502796 | 44.86148 | 7.25E-07 | 8.50E-06 | Healthy Control | Intestine |

| | | | | | | | |
|---|---|---|---|---|---|---|---|
| Prkcq | 3.63E+00 | 4.290971 | 37.58528 | 3.69E-06 | 3.40E-05 | Infection | Intestine |
| Prkcz | -1.07E+01 | 7.838061 | 38.49373 | 4.84E-06 | 4.36E-05 | Healthy Control | Intestine |
| Prkd1 | 1.47E+00 | 4.435552 | 2.764383 | 0.10963 | 0.232829 | Infection | Intestine |
| Prkd3 | 2.54E+00 | 7.500378 | 186.9109 | 3.59E-12 | 3.21E-10 | Infection | Intestine |
| Prkdc | -1.53E+00 | 5.973251 | 3.020239 | 0.094828 | 0.207749 | Healthy Control | Intestine |
| Prkg2 | -3.00E+00 | 7.604759 | 16.04124 | 0.000584 | 0.002905 | Healthy Control | Intestine |
| Prkn | 3.00E+00 | 2.675222 | 6.567817 | 0.01666 | 0.050637 | Infection | Intestine |
| Prl | 4.33E+00 | 1.362162 | 3.913842 | 0.059876 | 0.145011 | Infection | Intestine |
| Prl2c2 | 4.77E+00 | 1.363978 | 3.659273 | 0.067738 | 0.16071 | Infection | Intestine |
| Prl3a1 | -9.71E-16 | 1.180616 | 0 | 1 | 1 | Healthy Control | Intestine |
| Prl3b1 | -3.57E-01 | 1.228299 | 0.00618 | 0.938438 | 0.969735 | Healthy Control | Intestine |
| Prl3c1 | 5.33E+00 | 1.276041 | 4.71979 | 0.046639 | 0.118156 | Infection | Intestine |
| Prl3d1 | -1.56E+00 | 1.480294 | 0.041444 | 0.840554 | 0.954066 | Healthy Control | Intestine |
| Prl7a1 | -1.17E+00 | 1.361479 | 0.029597 | 0.864735 | 0.96005 | Healthy Control | Intestine |
| Prl8a1 | 5.04E+00 | 1.406576 | 3.950598 | 0.056542 | 0.138275 | Infection | Intestine |
| Prl8a2 | -9.71E-16 | 1.180616 | 0 | 1 | 1 | Healthy Control | Intestine |
| Prl8a6 | 4.75E+00 | 1.447061 | 6.065801 | 0.021198 | 0.061947 | Infection | Intestine |
| Prm1 | -2.06E+00 | 1.275606 | 0.054031 | 0.819151 | 0.953035 | Healthy Control | Intestine |
| Prmt1 | 1.87E-01 | 3.282302 | 0.010806 | 0.918155 | 0.969735 | Infection | Intestine |
| Prmt3 | 6.83E-01 | 6.858727 | 2.780354 | 0.109671 | 0.232829 | Infection | Intestine |
| Prmt5 | -1.33E+00 | 7.486826 | 9.611474 | 0.005014 | 0.018321 | Healthy Control | Intestine |
| Prmt7 | 1.41E-01 | 6.18808 | 0.062193 | 0.805278 | 0.947891 | Infection | Intestine |
| Prodh | 9.72E-01 | 5.194333 | 2.286661 | 0.144148 | 0.288988 | Infection | Intestine |
| Prok1 | -3.57E-01 | 1.227387 | 0.006293 | 0.937883 | 0.969735 | Healthy Control | Intestine |
| Prokr2 | 5.33E+00 | 1.276779 | 3.834058 | 0.069519 | 0.163539 | Infection | Intestine |
| Prom1 | -4.40E+00 | 9.249405 | 87.55794 | 3.03E-09 | 8.23E-08 | Healthy Control | Intestine |
| Prom2 | -6.42E-01 | 1.272459 | 0.013821 | 0.907644 | 0.969735 | Healthy Control | Intestine |
| Prop1 | -1.52E+00 | 1.274134 | 0.042767 | 0.838583 | 0.954066 | Healthy Control | Intestine |
| Prorp | -1.43E-01 | 5.578368 | 0.037588 | 0.847916 | 0.954066 | Healthy Control | Intestine |
| Prox1 | -1.50E+00 | 5.742446 | 1.407414 | 0.247371 | 0.439439 | Healthy Control | Intestine |
| Prpf19 | -1.73E+00 | 8.326859 | 32.96306 | 8.03E-06 | 6.85E-05 | Healthy Control | Intestine |
| Prpf3 | -2.31E-01 | 5.432762 | 0.101579 | 0.752696 | 0.925708 | Healthy Control | Intestine |
| Prpf31 | -2.11E-01 | 6.56618 | 0.14687 | 0.705107 | 0.896371 | Healthy Control | Intestine |
| Prpf4b | -1.48E-02 | 8.331324 | 0.004127 | 0.949363 | 0.97748 | Healthy Control | Intestine |
| Prpf8 | -1.98E+00 | 9.73981 | 101.007 | 1.19E-09 | 3.83E-08 | Healthy Control | Intestine |
| Prph | 1.55E-01 | 2.702776 | 0.003087 | 0.956132 | 0.981696 | Infection | Intestine |
| Prph2 | -4.95E-01 | 1.228692 | 0.008683 | 0.927069 | 0.969735 | Healthy Control | Intestine |
| Prps1 | 1.26E+00 | 5.637783 | 5.035273 | 0.035002 | 0.092904 | Infection | Intestine |
| Prpsap2 | -1.67E+00 | 7.653176 | 18.23761 | 0.000275 | 0.001505 | Healthy Control | Intestine |
| Prr5 | -9.10E+00 | 6.371092 | 11.9744 | 0.002499 | 0.010015 | Healthy Control | Intestine |
| Prrc2b | -1.05E+00 | 8.719036 | 13.70536 | 0.001254 | 0.005588 | Healthy Control | Intestine |
| Prrx1 | 5.05E+00 | 3.296994 | 45.41736 | 8.15E-07 | 9.43E-06 | Infection | Intestine |
| Prrxl1 | -2.24E+00 | 1.362135 | 0.077367 | 0.783259 | 0.940206 | Healthy Control | Intestine |
| Prss1 | 4.70E+00 | 1.364293 | 3.106409 | 0.09151 | 0.20234 | Infection | Intestine |
| Prss21 | -9.71E-16 | 1.180616 | 0 | 1 | 1 | Healthy Control | Intestine |
| Prss41 | 9.80E-01 | 3.867715 | 0.565269 | 0.459517 | 0.693919 | Infection | Intestine |
| Prss42 | 7.06E+00 | 1.277946 | 14.29634 | 0.003136 | 0.012161 | Infection | Intestine |
| Prss50 | 4.77E+00 | 1.362815 | 4.078549 | 0.054712 | 0.134788 | Infection | Intestine |
| Prune1 | 1.44E-01 | 6.67791 | 0.090871 | 0.765846 | 0.931224 | Infection | Intestine |
| Psat1 | 1.43E+00 | 4.27998 | 3.336487 | 0.080247 | 0.18318 | Infection | Intestine |

| | | | | | | | |
|---|---|---|---|---|---|---|---|
| Psd2 | 1.89E+00 | 2.167225 | 1.066443 | 0.310365 | 0.523372 | Infection | Intestine |
| Psen1 | -1.78E+00 | 9.970499 | 89.39031 | 3.63E-09 | 9.40E-08 | Healthy Control | Intestine |
| Psen2 | -2.96E+00 | 6.270446 | 7.363794 | 0.012502 | 0.039864 | Healthy Control | Intestine |
| Psma4 | -1.12E+00 | 9.36763 | 31.65021 | 1.22E-05 | 9.82E-05 | Healthy Control | Intestine |
| Psmb11 | -1.52E+00 | 1.274413 | 0.038404 | 0.84692 | 0.954066 | Healthy Control | Intestine |
| Psmb6 | -1.07E+00 | 8.682397 | 13.57401 | 0.001311 | 0.005789 | Healthy Control | Intestine |
| Psmb9 | 4.01E+00 | 1.949907 | 5.626326 | 0.024118 | 0.06914 | Infection | Intestine |
| Psmc2 | -3.54E-01 | 8.323805 | 2.046124 | 0.166786 | 0.322466 | Healthy Control | Intestine |
| Psmd1 | -1.59E+00 | 8.727829 | 34.11178 | 7.09E-06 | 6.10E-05 | Healthy Control | Intestine |
| Psmd10 | -2.09E+00 | 6.376801 | 6.960746 | 0.014462 | 0.044885 | Healthy Control | Intestine |
| Psmd11 | -3.23E+00 | 6.466367 | 10.51604 | 0.003682 | 0.014007 | Healthy Control | Intestine |
| Psmd12 | 7.59E-02 | 7.765347 | 0.076292 | 0.784984 | 0.940206 | Infection | Intestine |
| Psmd13 | -3.70E-01 | 8.47934 | 2.474009 | 0.130162 | 0.265956 | Healthy Control | Intestine |
| Psmd14 | -8.30E-01 | 8.175587 | 7.807274 | 0.01061 | 0.034512 | Healthy Control | Intestine |
| Psmd3 | -1.38E+00 | 8.620957 | 25.27305 | 5.00E-05 | 0.000335 | Healthy Control | Intestine |
| Psmd5 | -1.47E+00 | 7.60532 | 13.62792 | 0.001189 | 0.005321 | Healthy Control | Intestine |
| Psmd6 | -8.27E-01 | 8.109887 | 8.245805 | 0.00887 | 0.0297 | Healthy Control | Intestine |
| Psmd9 | -1.44E+00 | 7.61207 | 12.09626 | 0.002037 | 0.008384 | Healthy Control | Intestine |
| Ptafr | 2.26E-01 | 4.350489 | 0.040133 | 0.842904 | 0.954066 | Infection | Intestine |
| Ptar1 | -3.37E-01 | 7.198056 | 0.805179 | 0.379166 | 0.607199 | Healthy Control | Intestine |
| Ptbp1 | 3.53E-01 | 10.30618 | 8.812186 | 0.007137 | 0.024793 | Infection | Intestine |
| Ptch1 | -1.44E+00 | 8.400573 | 14.54065 | 0.000946 | 0.004406 | Healthy Control | Intestine |
| Ptch2 | 3.36E-01 | 3.373751 | 0.037433 | 0.848308 | 0.954066 | Infection | Intestine |
| Ptchd1 | 3.02E+00 | 2.163576 | 3.060124 | 0.091306 | 0.202068 | Infection | Intestine |
| Pten | -1.92E+00 | 9.301822 | 64.58331 | 5.36E-08 | 9.46E-07 | Healthy Control | Intestine |
| Ptf1a | -1.77E+00 | 1.319084 | 0.064605 | 0.802043 | 0.946717 | Healthy Control | Intestine |
| Ptgdr | 5.06E+00 | 1.319246 | 4.42779 | 0.048408 | 0.121834 | Infection | Intestine |
| Ptger1 | 3.17E+00 | 4.107738 | 15.94544 | 0.000617 | 0.003045 | Infection | Intestine |
| Ptger2 | 2.34E+00 | 4.009259 | 8.45744 | 0.007933 | 0.027032 | Infection | Intestine |
| Ptger3 | -4.97E-02 | 3.551578 | 0.00108 | 0.974077 | 0.995352 | Healthy Control | Intestine |
| Ptger4 | -1.02E+01 | 7.103513 | 22.04289 | 0.000142 | 0.000836 | Healthy Control | Intestine |
| Ptges | 1.78E+00 | 6.563619 | 19.31117 | 0.000234 | 0.001311 | Infection | Intestine |
| Ptgfr | -5.89E+00 | 2.987455 | 1.503404 | 0.232352 | 0.418552 | Healthy Control | Intestine |
| Ptgir | 4.70E+00 | 3.759695 | 57.95623 | 1.50E-07 | 2.21E-06 | Infection | Intestine |
| Ptgr1 | -1.33E+00 | 8.487357 | 17.17919 | 0.00042 | 0.002188 | Healthy Control | Intestine |
| Ptgs1 | 1.82E+00 | 7.389041 | 60.77813 | 9.65E-08 | 1.54E-06 | Infection | Intestine |
| Ptgs2 | -5.99E-01 | 5.544481 | 0.218847 | 0.644063 | 0.857095 | Healthy Control | Intestine |
| Pth | -1.37E+00 | 1.228088 | 0.037791 | 0.848672 | 0.954066 | Healthy Control | Intestine |
| Pth1r | 1.00E+00 | 5.42311 | 2.067556 | 0.164244 | 0.31918 | Infection | Intestine |
| Pth2r | -1.65E+00 | 1.31693 | 0.054107 | 0.818198 | 0.952901 | Healthy Control | Intestine |
| Pthlh | -5.75E+00 | 2.989772 | 1.171859 | 0.292171 | 0.498843 | Healthy Control | Intestine |
| Ptk2 | -4.04E-01 | 7.940671 | 1.73031 | 0.201997 | 0.374444 | Healthy Control | Intestine |
| Ptk2b | 8.16E-01 | 7.907305 | 14.66985 | 0.000925 | 0.004337 | Infection | Intestine |
| Ptk6 | 1.85E-01 | 4.626426 | 0.02542 | 0.87469 | 0.964471 | Infection | Intestine |
| Ptp4a1 | -2.62E+00 | 8.993219 | 61.01356 | 6.45E-08 | 1.12E-06 | Healthy Control | Intestine |
| Ptpa | -1.12E+00 | 9.429461 | 25.67741 | 4.61E-05 | 0.000313 | Healthy Control | Intestine |
| Ptpn1 | 1.06E+00 | 7.592279 | 15.70456 | 0.000671 | 0.003277 | Infection | Intestine |
| Ptpn11 | -3.19E-01 | 8.150965 | 1.485848 | 0.23589 | 0.423249 | Healthy Control | Intestine |
| Ptpn2 | 1.04E+00 | 7.893672 | 19.63905 | 0.000215 | 0.001213 | Infection | Intestine |
| Ptpn22 | 2.46E+00 | 3.112959 | 4.725484 | 0.039788 | 0.103577 | Infection | Intestine |

| | | | | | | | |
|---|---|---|---|---|---|---|---|
| Ptpn23 | 1.24E+00 | 7.694889 | 22.35617 | 0.000105 | 0.000644 | Infection | Intestine |
| Ptpn5 | -4.24E+00 | 2.088351 | 0.315665 | 0.57904 | 0.806375 | Healthy Control | Intestine |
| Ptpn6 | 6.12E-01 | 7.131041 | 2.550876 | 0.12457 | 0.257153 | Infection | Intestine |
| Ptpra | -1.20E+00 | 7.879881 | 10.1508 | 0.004206 | 0.015761 | Healthy Control | Intestine |
| Ptprc | 6.37E-01 | 6.824537 | 1.403252 | 0.248857 | 0.441923 | Infection | Intestine |
| Ptprf | -1.68E+00 | 7.628785 | 17.11133 | 0.000385 | 0.002029 | Healthy Control | Intestine |
| Ptprk | -2.43E+00 | 9.702333 | 112.1412 | 4.06E-10 | 1.56E-08 | Healthy Control | Intestine |
| Ptprn2 | -4.08E+00 | 9.684586 | 45.63318 | 1.85E-06 | 1.84E-05 | Healthy Control | Intestine |
| Ptprq | -2.78E+00 | 1.482752 | 0.145268 | 0.705866 | 0.896379 | Healthy Control | Intestine |
| Ptpru | -3.54E+00 | 7.243384 | 9.843233 | 0.004751 | 0.01749 | Healthy Control | Intestine |
| Ptprz1 | -5.97E+00 | 3.033318 | 0.913302 | 0.350879 | 0.574771 | Healthy Control | Intestine |
| Pttg1 | -7.04E-01 | 6.186344 | 0.809939 | 0.37706 | 0.604407 | Healthy Control | Intestine |
| Ptx3 | 3.19E+00 | 1.447394 | 2.118135 | 0.162332 | 0.316855 | Infection | Intestine |
| Pvalb | -9.71E-16 | 1.180616 | 0 | 1 | 1 | Healthy Control | Intestine |
| Pwp1 | -1.58E+00 | 6.456518 | 5.649856 | 0.025716 | 0.07265 | Healthy Control | Intestine |
| Pxn | -2.00E-01 | 6.115521 | 0.115762 | 0.7367 | 0.917621 | Healthy Control | Intestine |
| Pycard | -1.10E+00 | 7.316927 | 4.599555 | 0.042692 | 0.110164 | Healthy Control | Intestine |
| Pycrl | -4.50E+00 | 8.010402 | 31.73065 | 1.04E-05 | 8.54E-05 | Healthy Control | Intestine |
| Pygb | -9.77E-01 | 8.911482 | 17.07481 | 0.000441 | 0.002276 | Healthy Control | Intestine |
| Pygl | 4.84E+00 | 6.126071 | 440.5758 | 5.98E-16 | 1.76E-13 | Infection | Intestine |
| Pyroxd1 | 2.47E-01 | 7.36409 | 0.423575 | 0.521948 | 0.756696 | Infection | Intestine |
| Pzp | -7.56E-01 | 1.273429 | 0.015546 | 0.902199 | 0.969735 | Healthy Control | Intestine |
| Qrfpr | 4.48E+00 | 1.32044 | 3.384918 | 0.082418 | 0.186943 | Infection | Intestine |
| Qsox1 | 1.37E+00 | 7.504112 | 18.94487 | 0.00026 | 0.001437 | Infection | Intestine |
| Rab10 | -1.60E+00 | 9.400113 | 50.01766 | 4.47E-07 | 5.62E-06 | Healthy Control | Intestine |
| Rab11a | -1.66E+00 | 7.49288 | 10.57744 | 0.003418 | 0.013154 | Healthy Control | Intestine |
| Rab1a | -1.11E+00 | 9.613021 | 40.35496 | 2.27E-06 | 2.21E-05 | Healthy Control | Intestine |
| Rab20 | -8.76E+00 | 6.090421 | 9.298781 | 0.00638 | 0.022602 | Healthy Control | Intestine |
| Rab21 | 2.83E-01 | 8.488819 | 1.606241 | 0.218407 | 0.39787 | Infection | Intestine |
| Rab23 | -3.08E-01 | 6.250437 | 0.163162 | 0.689938 | 0.888369 | Healthy Control | Intestine |
| Rab24 | 5.92E-02 | 6.926794 | 0.019763 | 0.88947 | 0.969599 | Infection | Intestine |
| Rab27a | 6.28E-01 | 7.69326 | 3.274874 | 0.084187 | 0.189583 | Infection | Intestine |
| Rab27b | -3.97E+00 | 8.89962 | 48.86834 | 2.67E-07 | 3.59E-06 | Healthy Control | Intestine |
| Rab28 | -1.77E+00 | 8.332129 | 31.68658 | 1.08E-05 | 8.82E-05 | Healthy Control | Intestine |
| Rab29 | 4.99E-01 | 7.197134 | 2.140175 | 0.157707 | 0.309479 | Infection | Intestine |
| Rab32 | 6.79E-01 | 7.101369 | 3.606093 | 0.070664 | 0.165457 | Infection | Intestine |
| Rab38 | 1.87E+00 | 4.810209 | 6.29443 | 0.019674 | 0.058418 | Infection | Intestine |
| Rab39 | -5.23E+00 | 3.016133 | 1.061148 | 0.315236 | 0.528947 | Healthy Control | Intestine |
| Rab3a | -6.70E-02 | 6.617379 | 0.018466 | 0.893088 | 0.969735 | Healthy Control | Intestine |
| Rab3b | -6.74E+00 | 3.596724 | 1.727268 | 0.203763 | 0.377152 | Healthy Control | Intestine |
| Rab3d | -2.42E+00 | 8.827842 | 37.59341 | 3.03E-06 | 2.87E-05 | Healthy Control | Intestine |
| Rab3gap2 | 1.07E+00 | 7.42778 | 22.9341 | 9.02E-05 | 0.000567 | Infection | Intestine |
| Rab43 | -1.14E+00 | 8.473313 | 12.8293 | 0.001674 | 0.007096 | Healthy Control | Intestine |
| Rab4b | -8.53E-01 | 6.584081 | 2.527487 | 0.125041 | 0.257733 | Healthy Control | Intestine |
| Rab5a | -3.73E+00 | 7.915861 | 25.59978 | 3.75E-05 | 0.000261 | Healthy Control | Intestine |
| Rab5b | -1.44E+00 | 8.862837 | 27.44224 | 2.98E-05 | 0.000214 | Healthy Control | Intestine |
| Rab5c | -5.82E-01 | 10.10954 | 13.02901 | 0.001572 | 0.006739 | Healthy Control | Intestine |
| Rab6a | -2.04E+00 | 9.515485 | 88.40697 | 3.61E-09 | 9.40E-08 | Healthy Control | Intestine |
| Rab7 | -2.43E-02 | 10.81677 | 0.046751 | 0.830826 | 0.954066 | Healthy Control | Intestine |
| Rab7b | 6.42E-01 | 5.880873 | 0.711735 | 0.407738 | 0.638853 | Infection | Intestine |

| | | | | | | | |
|---|---|---|---|---|---|---|---|
| Rab8a | -6.84E-01 | 8.971983 | 7.997608 | 0.009853 | 0.03247 | Healthy Control | Intestine |
| Rab8b | -1.58E-01 | 6.568045 | 0.081145 | 0.778316 | 0.936824 | Healthy Control | Intestine |
| Rabep1 | 7.26E-02 | 7.918245 | 0.083871 | 0.774861 | 0.934687 | Infection | Intestine |
| Rabep2 | -1.89E+00 | 6.14093 | 5.036554 | 0.034345 | 0.091551 | Healthy Control | Intestine |
| Rac1 | -1.52E+00 | 10.94286 | 120.4654 | 2.41E-10 | 1.01E-08 | Healthy Control | Intestine |
| Racgap1 | 3.80E-01 | 7.096809 | 0.682515 | 0.417677 | 0.650153 | Infection | Intestine |
| Rack1 | -1.51E+00 | 8.472067 | 17.20232 | 0.000404 | 0.002111 | Healthy Control | Intestine |
| Rad21l | -1.65E+00 | 1.318958 | 0.045773 | 0.832543 | 0.954066 | Healthy Control | Intestine |
| Rad23b | -4.03E-01 | 9.673648 | 8.569204 | 0.00785 | 0.026788 | Healthy Control | Intestine |
| Rad50 | -4.67E-01 | 6.401284 | 0.662273 | 0.423951 | 0.656651 | Healthy Control | Intestine |
| Rad51 | 1.75E+00 | 5.327207 | 8.654077 | 0.007547 | 0.025996 | Infection | Intestine |
| Rad51b | 1.89E+00 | 5.943173 | 29.58002 | 1.91E-05 | 0.000146 | Infection | Intestine |
| Rad51d | -7.40E-01 | 7.190288 | 2.568391 | 0.122823 | 0.254071 | Healthy Control | Intestine |
| Rad54l | 1.22E+00 | 5.269271 | 3.145149 | 0.089728 | 0.199104 | Infection | Intestine |
| Rad9a | 4.34E-02 | 6.346372 | 0.008756 | 0.926272 | 0.969735 | Infection | Intestine |
| Rae1 | 7.39E-02 | 7.001939 | 0.04873 | 0.827304 | 0.953685 | Infection | Intestine |
| Raf1 | -4.22E-02 | 9.936344 | 0.103418 | 0.750828 | 0.92508 | Healthy Control | Intestine |
| Rag1 | 3.76E+00 | 1.525001 | 3.075484 | 0.088294 | 0.196707 | Infection | Intestine |
| Rag2 | -2.16E+00 | 1.319306 | 0.078848 | 0.781855 | 0.939954 | Healthy Control | Intestine |
| Rai14 | 2.04E+00 | 7.012727 | 57.38206 | 1.54E-07 | 2.26E-06 | Infection | Intestine |
| Rala | -1.73E+00 | 8.576306 | 25.4714 | 4.49E-05 | 0.000306 | Healthy Control | Intestine |
| Ramp3 | -5.40E-01 | 4.940494 | 0.196447 | 0.661563 | 0.869462 | Healthy Control | Intestine |
| Ranbp1 | -1.05E+00 | 7.350196 | 4.710198 | 0.040584 | 0.105265 | Healthy Control | Intestine |
| Ranbp2 | 9.32E-01 | 8.545691 | 29.8438 | 1.79E-05 | 0.000138 | Infection | Intestine |
| Rangap1 | -3.97E-01 | 8.036685 | 1.762149 | 0.198082 | 0.368761 | Healthy Control | Intestine |
| Rap1a | -2.23E+00 | 8.612916 | 54.99276 | 1.60E-07 | 2.32E-06 | Healthy Control | Intestine |
| Rap1gap | -3.51E+00 | 9.720973 | 46.81353 | 1.21E-06 | 1.31E-05 | Healthy Control | Intestine |
| Rapgef1 | 1.05E+00 | 7.862459 | 20.44872 | 0.000172 | 0.000994 | Infection | Intestine |
| Rapgef3 | -2.31E+00 | 6.186121 | 4.556307 | 0.043954 | 0.11278 | Healthy Control | Intestine |
| Rapsn | 3.27E+00 | 2.445608 | 3.61827 | 0.068276 | 0.161712 | Infection | Intestine |
| Rara | -1.76E+00 | 8.66501 | 19.07274 | 0.000238 | 0.00133 | Healthy Control | Intestine |
| Rarb | 8.62E-01 | 3.062685 | 0.26427 | 0.612063 | 0.833585 | Infection | Intestine |
| Rarg | 2.03E+00 | 6.54817 | 33.107 | 9.03E-06 | 7.53E-05 | Infection | Intestine |
| Rasd1 | -3.81E+00 | 6.749312 | 10.06249 | 0.004503 | 0.016697 | Healthy Control | Intestine |
| Rasgrf1 | 4.04E+00 | 2.354151 | 13.05127 | 0.00115 | 0.005178 | Infection | Intestine |
| Rassf5 | 7.84E-01 | 5.794365 | 1.019199 | 0.323334 | 0.539358 | Infection | Intestine |
| Rax | -3.57E-01 | 1.227417 | 0.006288 | 0.937908 | 0.969735 | Healthy Control | Intestine |
| Rb1 | 5.77E-01 | 8.367916 | 8.961627 | 0.006741 | 0.023615 | Infection | Intestine |
| Rb1cc1 | -8.55E+00 | 5.846171 | 9.335143 | 0.006295 | 0.022333 | Healthy Control | Intestine |
| Rbbp4 | -1.08E+00 | 9.544321 | 35.86622 | 5.22E-06 | 4.63E-05 | Healthy Control | Intestine |
| Rbbp7 | -9.40E-01 | 8.894659 | 15.37872 | 0.00074 | 0.003539 | Healthy Control | Intestine |
| Rbck1 | -1.53E+00 | 8.008354 | 17.8449 | 0.00033 | 0.001773 | Healthy Control | Intestine |
| Rbfox2 | 5.33E-01 | 6.378138 | 1.27957 | 0.269995 | 0.469369 | Infection | Intestine |
| Rbfox3 | -3.53E+00 | 1.973089 | 0.307589 | 0.583841 | 0.810063 | Healthy Control | Intestine |
| Rbl1 | -1.20E+00 | 6.97391 | 4.024175 | 0.056467 | 0.138226 | Healthy Control | Intestine |
| Rbl2 | 1.12E+00 | 6.842231 | 12.02658 | 0.002207 | 0.008988 | Infection | Intestine |
| Rbm8a | -6.20E-01 | 5.018771 | 0.376061 | 0.545563 | 0.776921 | Healthy Control | Intestine |
| Rbms1 | 2.37E+00 | 8.597618 | 252.1108 | 1.82E-13 | 2.77E-11 | Infection | Intestine |
| Rbp1 | 4.33E+00 | 5.937757 | 126.025 | 1.58E-10 | 7.48E-09 | Infection | Intestine |
| Rbp3 | -9.71E-16 | 1.180616 | 0 | 1 | 1 | Healthy Control | Intestine |

| | | | | | | | |
|---|---|---|---|---|---|---|---|
| Rbpj | 1.64E+00 | 8.525215 | 92.06545 | 2.78E-09 | 7.76E-08 | Infection | Intestine |
| Rbpjl | -2.32E+00 | 1.402452 | 0.096713 | 0.758122 | 0.928591 | Healthy Control | Intestine |
| Rcan1 | -8.84E-01 | 6.53673 | 2.170942 | 0.153717 | 0.303553 | Healthy Control | Intestine |
| Rcc2 | -7.93E-01 | 8.007496 | 5.240399 | 0.0319 | 0.086181 | Healthy Control | Intestine |
| Rcn2 | -1.65E-01 | 7.023387 | 0.143584 | 0.708344 | 0.897011 | Healthy Control | Intestine |
| Rcor1 | -1.50E+00 | 8.662493 | 32.17764 | 1.02E-05 | 8.44E-05 | Healthy Control | Intestine |
| Rcor2 | 3.26E+00 | 3.655705 | 17.50748 | 0.000366 | 0.00194 | Infection | Intestine |
| Rdx | -1.78E-01 | 8.309778 | 0.229336 | 0.636769 | 0.852316 | Healthy Control | Intestine |
| Rec114 | -1.52E+00 | 1.273358 | 0.043368 | 0.83747 | 0.954066 | Healthy Control | Intestine |
| Rec8 | -3.70E+00 | 1.759714 | 0.14427 | 0.7075 | 0.897011 | Healthy Control | Intestine |
| Rel | -2.26E-02 | 5.777465 | 0.001418 | 0.970277 | 0.992279 | Healthy Control | Intestine |
| Rela | 5.74E-01 | 7.605853 | 4.432637 | 0.04701 | 0.118796 | Infection | Intestine |
| Relb | -7.28E-01 | 8.279853 | 5.561282 | 0.027668 | 0.077277 | Healthy Control | Intestine |
| Reln | 4.26E-01 | 5.216995 | 0.046417 | 0.832108 | 0.954066 | Infection | Intestine |
| Ren1 | -3.18E+00 | 1.565404 | 0.12542 | 0.725517 | 0.91026 | Healthy Control | Intestine |
| Rest | 5.65E-01 | 7.183331 | 3.073645 | 0.093546 | 0.205662 | Infection | Intestine |
| Ret | 1.08E-01 | 4.151145 | 0.006614 | 0.935865 | 0.969735 | Infection | Intestine |
| Retn | 4.20E+00 | 1.407171 | 2.649563 | 0.114873 | 0.242027 | Infection | Intestine |
| Retreg3 | -1.94E+00 | 8.393871 | 28.25765 | 2.20E-05 | 0.000166 | Healthy Control | Intestine |
| Rev1 | 1.71E-01 | 6.52147 | 0.17122 | 0.682883 | 0.883793 | Infection | Intestine |
| Rflna | -4.34E+00 | 2.318025 | 0.509161 | 0.483086 | 0.717392 | Healthy Control | Intestine |
| Rfx2 | 5.48E-01 | 4.346858 | 0.248339 | 0.622651 | 0.839656 | Infection | Intestine |
| Rfx3 | 1.25E+00 | 5.589922 | 6.35012 | 0.019387 | 0.057767 | Infection | Intestine |
| Rfx4 | -9.77E-01 | 1.319329 | 0.020434 | 0.88761 | 0.969534 | Healthy Control | Intestine |
| Rfx5 | -1.22E+00 | 5.630606 | 1.855286 | 0.185699 | 0.350928 | Healthy Control | Intestine |
| Rfx6 | -5.19E+00 | 3.021804 | 0.838634 | 0.369975 | 0.595529 | Healthy Control | Intestine |
| Rgcc | -2.84E+00 | 7.364112 | 7.873108 | 0.009927 | 0.03263 | Healthy Control | Intestine |
| Rgmb | -5.83E-02 | 9.099964 | 0.087407 | 0.7703 | 0.932782 | Healthy Control | Intestine |
| Rgr | -3.57E-01 | 1.227509 | 0.006273 | 0.937979 | 0.969735 | Healthy Control | Intestine |
| Rgs18 | 2.57E+00 | 3.793312 | 10.13375 | 0.004112 | 0.015442 | Infection | Intestine |
| Rgs3 | -4.91E-01 | 7.632018 | 1.353323 | 0.257126 | 0.452752 | Healthy Control | Intestine |
| Rgs6 | -5.96E-01 | 5.09136 | 0.205135 | 0.654586 | 0.864378 | Healthy Control | Intestine |
| Rgs9 | -8.19E-02 | 3.585165 | 0.002499 | 0.960577 | 0.985267 | Healthy Control | Intestine |
| Rhbdf2 | 1.49E+00 | 6.46852 | 18.78462 | 0.000269 | 0.001477 | Infection | Intestine |
| Rhbdl3 | -6.55E+00 | 3.65399 | 1.97519 | 0.175533 | 0.336416 | Healthy Control | Intestine |
| Rheb | -9.31E-02 | 8.069977 | 0.136531 | 0.715317 | 0.901639 | Healthy Control | Intestine |
| Rho | 5.39E+00 | 1.448992 | 7.091406 | 0.012454 | 0.039736 | Infection | Intestine |
| Rhoa | -1.16E+00 | 10.58918 | 64.89501 | 5.63E-08 | 9.88E-07 | Healthy Control | Intestine |
| Rhoc | 1.13E-02 | 9.968226 | 0.002491 | 0.960665 | 0.985267 | Infection | Intestine |
| Rhod | 1.15E+00 | 5.06672 | 2.508287 | 0.126617 | 0.259985 | Infection | Intestine |
| Rhog | -2.98E+00 | 7.970492 | 33.03837 | 6.52E-06 | 5.70E-05 | Healthy Control | Intestine |
| Rhoj | 3.10E-01 | 7.06739 | 0.315678 | 0.580121 | 0.807306 | Infection | Intestine |
| Rhot2 | -5.77E-01 | 8.371103 | 4.944846 | 0.036854 | 0.0971 | Healthy Control | Intestine |
| Rhox13 | -3.57E-01 | 1.227417 | 0.006288 | 0.937908 | 0.969735 | Healthy Control | Intestine |
| Rhox5 | 2.87E+00 | 1.951247 | 1.498478 | 0.230596 | 0.41719 | Infection | Intestine |
| Rhpn1 | -3.60E+00 | 1.739685 | 0.214246 | 0.646675 | 0.858916 | Healthy Control | Intestine |
| Rhpn2 | -2.13E+00 | 7.68027 | 16.35116 | 0.000475 | 0.002435 | Healthy Control | Intestine |
| Ric8a | -2.90E-01 | 8.228511 | 1.289112 | 0.268522 | 0.467555 | Healthy Control | Intestine |
| Ric8b | -1.28E+00 | 6.88906 | 5.175327 | 0.03218 | 0.086743 | Healthy Control | Intestine |
| Rictor | 9.51E-01 | 7.390905 | 12.07663 | 0.002172 | 0.00886 | Infection | Intestine |

| | | | | | | | |
|---|---|---|---|---|---|---|---|
| Rif1 | -4.96E-01 | 7.431026 | 1.746055 | 0.199739 | 0.371155 | Healthy Control | Intestine |
| Rims2 | -7.73E-01 | 5.410864 | 0.370514 | 0.548436 | 0.778194 | Healthy Control | Intestine |
| Ring1 | -6.20E-01 | 7.467477 | 3.014407 | 0.096347 | 0.210434 | Healthy Control | Intestine |
| Riok3 | -5.44E-01 | 9.180766 | 7.639839 | 0.011389 | 0.036736 | Healthy Control | Intestine |
| Riox1 | 1.22E-01 | 5.734554 | 0.035208 | 0.852754 | 0.956553 | Infection | Intestine |
| Riox2 | -1.57E-01 | 6.082079 | 0.058331 | 0.811234 | 0.949838 | Healthy Control | Intestine |
| Ripk1 | -1.04E+00 | 7.305086 | 7.792487 | 0.010347 | 0.033811 | Healthy Control | Intestine |
| Ripk2 | -1.46E+00 | 6.607597 | 5.960628 | 0.022225 | 0.064648 | Healthy Control | Intestine |
| Ripk3 | -1.06E+00 | 6.345065 | 3.607616 | 0.069486 | 0.163539 | Healthy Control | Intestine |
| Rita1 | -1.09E+00 | 5.361344 | 1.476185 | 0.23647 | 0.423987 | Healthy Control | Intestine |
| Rlim | -4.30E-01 | 8.2808 | 2.701807 | 0.114543 | 0.241432 | Healthy Control | Intestine |
| Rln1 | -1.99E+00 | 1.404908 | 0.067884 | 0.796375 | 0.943769 | Healthy Control | Intestine |
| Rln3 | -2.99E+00 | 1.80088 | 0.122178 | 0.729259 | 0.912896 | Healthy Control | Intestine |
| Rmdn3 | 1.41E+00 | 7.087198 | 28.79638 | 2.26E-05 | 0.000169 | Infection | Intestine |
| Rnase1 | -2.67E+00 | 1.730702 | 0.122999 | 0.728773 | 0.912515 | Healthy Control | Intestine |
| Rnaseh2b | -6.86E-01 | 6.292622 | 1.297505 | 0.265958 | 0.46406 | Healthy Control | Intestine |
| Rnaset2a | -1.10E+00 | 11.35287 | 107.1846 | 7.04E-10 | 2.43E-08 | Healthy Control | Intestine |
| Rnd2 | 1.19E+00 | 4.660894 | 2.10588 | 0.160009 | 0.313014 | Infection | Intestine |
| Rnf10 | -5.42E-02 | 10.14789 | 0.16895 | 0.685058 | 0.885006 | Healthy Control | Intestine |
| Rnf13 | 9.35E-01 | 8.771618 | 36.29165 | 4.79E-06 | 4.32E-05 | Infection | Intestine |
| Rnf130 | -1.33E+00 | 8.521299 | 28.47806 | 2.32E-05 | 0.000173 | Healthy Control | Intestine |
| Rnf14 | 1.67E-01 | 8.862835 | 0.746349 | 0.397049 | 0.626197 | Infection | Intestine |
| Rnf144a | 3.86E-02 | 5.33961 | 0.001627 | 0.968153 | 0.990714 | Infection | Intestine |
| Rnf19a | 1.37E+00 | 7.073671 | 27.49858 | 3.02E-05 | 0.000216 | Infection | Intestine |
| Rnf2 | -6.79E-03 | 7.231225 | 0.000339 | 0.985468 | 1 | Healthy Control | Intestine |
| Rnf20 | 1.32E-01 | 6.63184 | 0.082098 | 0.777115 | 0.935948 | Infection | Intestine |
| Rnf219 | 8.04E-01 | 6.087198 | 2.721116 | 0.113113 | 0.238821 | Infection | Intestine |
| Rnf220 | 1.08E+00 | 8.178706 | 31.80501 | 1.18E-05 | 9.52E-05 | Infection | Intestine |
| Rnf31 | -5.35E-01 | 6.663099 | 0.867297 | 0.36125 | 0.586019 | Healthy Control | Intestine |
| Rnh1 | -1.81E+00 | 9.488702 | 69.35132 | 3.13E-08 | 5.94E-07 | Healthy Control | Intestine |
| Robo1 | 9.45E-01 | 6.025639 | 2.483956 | 0.129141 | 0.264301 | Infection | Intestine |
| Robo2 | 9.97E-01 | 6.088903 | 3.042175 | 0.095097 | 0.208249 | Infection | Intestine |
| Robo3 | -2.93E+00 | 1.405749 | 0.188549 | 0.669161 | 0.87577 | Healthy Control | Intestine |
| Rock1 | -5.20E-01 | 8.093133 | 3.138541 | 0.090401 | 0.20033 | Healthy Control | Intestine |
| Rock2 | 1.27E-01 | 9.438028 | 0.755721 | 0.394142 | 0.623258 | Infection | Intestine |
| Ror1 | 4.90E-01 | 5.845255 | 0.64972 | 0.428307 | 0.661048 | Infection | Intestine |
| Ror2 | 2.26E+00 | 5.051232 | 16.74697 | 0.000482 | 0.00246 | Infection | Intestine |
| Rora | 1.02E-01 | 5.547047 | 0.014172 | 0.906234 | 0.969735 | Infection | Intestine |
| Rorb | 2.34E+00 | 2.946488 | 2.326678 | 0.141676 | 0.285345 | Infection | Intestine |
| Rorc | -2.27E+00 | 6.490063 | 6.689722 | 0.016178 | 0.049411 | Healthy Control | Intestine |
| Ros1 | 3.47E+00 | 1.524934 | 2.507126 | 0.123455 | 0.255063 | Infection | Intestine |
| Rp1l1 | -3.57E-01 | 1.228299 | 0.00618 | 0.938438 | 0.969735 | Healthy Control | Intestine |
| Rp2 | -4.71E-01 | 8.529759 | 4.164774 | 0.053564 | 0.13248 | Healthy Control | Intestine |
| Rpa1 | -2.29E-02 | 7.316232 | 0.003086 | 0.956202 | 0.981696 | Healthy Control | Intestine |
| Rpe65 | -1.34E+00 | 1.402691 | 0.033202 | 0.856755 | 0.956553 | Healthy Control | Intestine |
| Rph3a | -2.54E+00 | 1.484383 | 0.113453 | 0.738358 | 0.919229 | Healthy Control | Intestine |
| Rph3al | -1.77E+00 | 7.351212 | 15.50648 | 0.000624 | 0.003079 | Healthy Control | Intestine |
| Rpn1 | -1.76E+00 | 9.973259 | 106.4594 | 7.64E-10 | 2.58E-08 | Healthy Control | Intestine |
| Rprml | 4.91E-01 | 3.200112 | 0.092512 | 0.76372 | 0.929989 | Infection | Intestine |
| Rptor | -1.59E+00 | 7.493126 | 12.1625 | 0.001959 | 0.008119 | Healthy Control | Intestine |

| | | | | | | | |
|---|---|---|---|---|---|---|---|
| Rragd | 1.43E+00 | 3.894633 | 1.463129 | 0.237972 | 0.42571 | Infection | Intestine |
| Rras2 | -1.23E+00 | 7.748455 | 12.61916 | 0.001712 | 0.00724 | Healthy Control | Intestine |
| Rsad1 | -1.83E+00 | 6.360767 | 2.648653 | 0.116392 | 0.244609 | Healthy Control | Intestine |
| Rsbn1 | -6.14E-01 | 7.482143 | 3.269981 | 0.084091 | 0.189451 | Healthy Control | Intestine |
| Rsf1 | -1.58E-01 | 8.191837 | 0.38138 | 0.543265 | 0.775107 | Healthy Control | Intestine |
| Rsph1 | -4.88E-01 | 3.928773 | 0.094765 | 0.761051 | 0.928591 | Healthy Control | Intestine |
| Rsph10b | -4.34E+00 | 2.138468 | 0.521667 | 0.477359 | 0.712267 | Healthy Control | Intestine |
| Rsph9 | 6.22E-01 | 4.30161 | 0.386071 | 0.540091 | 0.772705 | Infection | Intestine |
| Rspo1 | -3.99E+00 | 2.002836 | 0.226058 | 0.638599 | 0.853397 | Healthy Control | Intestine |
| Rspo2 | -3.84E+00 | 2.007715 | 0.355066 | 0.55583 | 0.784337 | Healthy Control | Intestine |
| Rspo3 | 1.87E+00 | 4.558506 | 3.226086 | 0.08616 | 0.192638 | Infection | Intestine |
| Rsrp1 | 2.21E+00 | 11.42943 | 1007.011 | 9.58E-20 | 1.20E-16 | Infection | Intestine |
| Rtbdn | -1.58E+00 | 1.274661 | 0.044183 | 0.835981 | 0.954066 | Healthy Control | Intestine |
| Rtcb | -1.75E+00 | 8.887041 | 54.23228 | 2.08E-07 | 2.90E-06 | Healthy Control | Intestine |
| Rtf1 | 3.90E-01 | 7.147415 | 1.48584 | 0.235813 | 0.423249 | Infection | Intestine |
| Rtkn2 | -4.37E+00 | 2.460991 | 0.58422 | 0.452891 | 0.686424 | Healthy Control | Intestine |
| Rtn2 | -6.13E+00 | 3.519479 | 1.018731 | 0.3251 | 0.541581 | Healthy Control | Intestine |
| Rtn3 | 1.32E-01 | 10.39125 | 1.260853 | 0.273701 | 0.47525 | Infection | Intestine |
| Rtn4 | 3.14E-01 | 9.439895 | 2.640781 | 0.118521 | 0.24742 | Infection | Intestine |
| Rtn4r | -1.94E+00 | 4.761687 | 2.384468 | 0.137265 | 0.277972 | Healthy Control | Intestine |
| Rubcn | -1.10E+00 | 7.407504 | 5.851387 | 0.023914 | 0.068721 | Healthy Control | Intestine |
| Rundc1 | -1.46E+00 | 6.674584 | 6.689993 | 0.016075 | 0.049132 | Healthy Control | Intestine |
| Runx1 | 2.04E+00 | 6.794624 | 38.07437 | 3.43E-06 | 3.21E-05 | Infection | Intestine |
| Runx2 | 2.84E+00 | 6.237428 | 75.9318 | 1.50E-08 | 3.10E-07 | Infection | Intestine |
| Runx3 | 3.50E-01 | 4.628713 | 0.105017 | 0.748662 | 0.924698 | Infection | Intestine |
| Ruvbl1 | -1.01E+00 | 7.285949 | 7.198141 | 0.013314 | 0.041864 | Healthy Control | Intestine |
| Ruvbl2 | -1.43E+00 | 8.132362 | 15.02081 | 0.000797 | 0.003785 | Healthy Control | Intestine |
| Rxfp1 | -1.32E+00 | 1.363378 | 0.032817 | 0.858154 | 0.956553 | Healthy Control | Intestine |
| Rxfp2 | -1.52E+00 | 1.273038 | 0.043716 | 0.836829 | 0.954066 | Healthy Control | Intestine |
| Rxfp3 | 5.30E+00 | 1.486298 | 5.784486 | 0.022761 | 0.066015 | Infection | Intestine |
| Rxra | -2.02E+00 | 8.641046 | 30.0841 | 1.57E-05 | 0.000124 | Healthy Control | Intestine |
| Rxrb | -7.75E-01 | 7.98519 | 5.354355 | 0.030383 | 0.082978 | Healthy Control | Intestine |
| Rxrg | 1.90E+00 | 2.295972 | 0.994369 | 0.328206 | 0.545666 | Infection | Intestine |
| Rybp | 6.69E-01 | 7.076788 | 3.964056 | 0.059148 | 0.143406 | Infection | Intestine |
| Ryk | -1.27E+00 | 8.463926 | 21.42741 | 0.000129 | 0.000771 | Healthy Control | Intestine |
| Ryr1 | -2.36E+00 | 1.403891 | 0.087022 | 0.770213 | 0.932782 | Healthy Control | Intestine |
| Ryr2 | 3.19E+00 | 2.780724 | 8.22593 | 0.008399 | 0.02839 | Infection | Intestine |
| Ryr3 | 2.28E+00 | 2.495757 | 2.439891 | 0.130808 | 0.266948 | Infection | Intestine |
| S100a1 | -1.61E+00 | 9.503418 | 41.17725 | 1.94E-06 | 1.92E-05 | Healthy Control | Intestine |
| S100b | 2.64E+00 | 2.583449 | 2.391614 | 0.133443 | 0.271661 | Infection | Intestine |
| S1pr1 | -1.69E+00 | 6.363649 | 1.387145 | 0.252913 | 0.446747 | Healthy Control | Intestine |
| S1pr2 | 2.10E+00 | 6.354211 | 34.32113 | 7.06E-06 | 6.10E-05 | Infection | Intestine |
| S1pr3 | 4.57E+00 | 4.200478 | 19.03602 | 0.000474 | 0.002434 | Infection | Intestine |
| S1pr5 | -1.15E+00 | 1.319509 | 0.027734 | 0.869856 | 0.961258 | Healthy Control | Intestine |
| Saa1 | -3.51E+00 | 10.76723 | 16.31723 | 0.001317 | 0.00581 | Healthy Control | Intestine |
| Saa2 | -3.62E+00 | 1.889 | 0.199115 | 0.658936 | 0.867142 | Healthy Control | Intestine |
| Sacm1l | -1.24E+00 | 8.34608 | 20.03332 | 0.000187 | 0.00107 | Healthy Control | Intestine |
| Sag | -4.92E+00 | 2.869647 | 0.748317 | 0.396963 | 0.626197 | Healthy Control | Intestine |
| Sall1 | -1.26E+00 | 5.676368 | 1.243721 | 0.275633 | 0.47728 | Healthy Control | Intestine |
| Sall3 | 1.03E+00 | 2.417892 | 0.395526 | 0.53527 | 0.767123 | Infection | Intestine |

| | | | | | | | |
|---|---|---|---|---|---|---|---|
| Sall4 | 4.33E+00 | 1.363163 | 3.487122 | 0.074523 | 0.172793 | Infection | Intestine |
| Samd11 | -2.08E+00 | 1.445347 | 0.049325 | 0.825683 | 0.953685 | Healthy Control | Intestine |
| Samd14 | 2.20E+00 | 4.696861 | 13.14185 | 0.001471 | 0.006383 | Infection | Intestine |
| Samd8 | -4.26E-01 | 6.497697 | 0.652175 | 0.427388 | 0.660341 | Healthy Control | Intestine |
| Samhd1 | 6.13E-01 | 9.165523 | 7.654409 | 0.011315 | 0.03652 | Infection | Intestine |
| Sap130 | -1.34E+00 | 8.501208 | 17.87796 | 0.000344 | 0.001835 | Healthy Control | Intestine |
| Sap18 | -1.05E+00 | 6.964159 | 4.864931 | 0.037428 | 0.098405 | Healthy Control | Intestine |
| Saraf | -1.50E+00 | 9.184792 | 37.36311 | 3.88E-06 | 3.57E-05 | Healthy Control | Intestine |
| Sardh | 2.31E+00 | 5.196779 | 19.2221 | 0.00024 | 0.001341 | Infection | Intestine |
| Sarm1 | 1.63E+00 | 2.285399 | 0.633105 | 0.433736 | 0.666066 | Infection | Intestine |
| Sart3 | -6.20E-01 | 7.325762 | 2.588008 | 0.121656 | 0.252388 | Healthy Control | Intestine |
| Sass6 | -8.75E+00 | 5.860552 | 11.31315 | 0.003122 | 0.012126 | Healthy Control | Intestine |
| Satb2 | -3.43E+00 | 9.448994 | 132.911 | 3.68E-11 | 2.19E-09 | Healthy Control | Intestine |
| Saysd1 | -7.52E+00 | 4.884498 | 3.932759 | 0.061404 | 0.14814 | Healthy Control | Intestine |
| Scamp4 | -1.66E+00 | 7.915579 | 18.63211 | 0.000255 | 0.001414 | Healthy Control | Intestine |
| Scarb1 | -8.72E+00 | 5.912855 | 14.52937 | 0.001108 | 0.005042 | Healthy Control | Intestine |
| Scarb2 | 1.08E+00 | 9.809285 | 69.67964 | 3.10E-08 | 5.89E-07 | Infection | Intestine |
| Scgb1b27 | -1.99E+00 | 1.403389 | 0.07464 | 0.786729 | 0.941182 | Healthy Control | Intestine |
| Scimp | -5.21E+00 | 2.722747 | 0.766943 | 0.391669 | 0.621456 | Healthy Control | Intestine |
| Scin | -4.94E+00 | 9.304352 | 65.20255 | 5.33E-08 | 9.43E-07 | Healthy Control | Intestine |
| Scmh1 | -4.37E-02 | 7.034799 | 0.015019 | 0.903564 | 0.969735 | Healthy Control | Intestine |
| Scn10a | -3.57E-01 | 1.228299 | 0.00618 | 0.938438 | 0.969735 | Healthy Control | Intestine |
| Scn1a | -9.77E-01 | 1.318851 | 0.020879 | 0.886403 | 0.96867 | Healthy Control | Intestine |
| Scn2a | 1.80E+00 | 2.190572 | 0.759662 | 0.391364 | 0.621456 | Infection | Intestine |
| Scn3b | -4.65E+00 | 2.514601 | 0.349151 | 0.560904 | 0.78962 | Healthy Control | Intestine |
| Scn4a | -3.57E-01 | 1.227509 | 0.006273 | 0.937979 | 0.969735 | Healthy Control | Intestine |
| Scn4b | 1.39E+00 | 3.455579 | 0.542978 | 0.468189 | 0.703509 | Infection | Intestine |
| Scn5a | -2.51E+00 | 1.445836 | 0.11032 | 0.742636 | 0.92188 | Healthy Control | Intestine |
| Scn8a | -2.25E+00 | 1.483781 | 0.088216 | 0.768794 | 0.932538 | Healthy Control | Intestine |
| Scn9a | -1.88E+00 | 1.362915 | 0.051158 | 0.823084 | 0.953685 | Healthy Control | Intestine |
| Scnn1a | -4.35E+00 | 8.378989 | 34.79475 | 5.11E-06 | 4.55E-05 | Healthy Control | Intestine |
| Scnn1b | -1.12E+00 | 4.667809 | 0.453851 | 0.507285 | 0.741883 | Healthy Control | Intestine |
| Scnn1g | -3.87E+00 | 2.265289 | 0.245426 | 0.624443 | 0.84099 | Healthy Control | Intestine |
| Scrib | 3.30E-01 | 8.657722 | 3.335237 | 0.081552 | 0.185567 | Infection | Intestine |
| Scrt1 | -8.62E-01 | 1.275087 | 0.020018 | 0.889226 | 0.969599 | Healthy Control | Intestine |
| Scrt2 | -1.65E+00 | 1.318143 | 0.048108 | 0.828394 | 0.954066 | Healthy Control | Intestine |
| Sct | -2.12E+00 | 6.406883 | 3.098988 | 0.091554 | 0.202349 | Healthy Control | Intestine |
| Scx | -4.56E+00 | 2.40199 | 0.682837 | 0.41782 | 0.650173 | Healthy Control | Intestine |
| Sdc1 | 3.85E-01 | 10.21627 | 5.879459 | 0.024064 | 0.069035 | Infection | Intestine |
| Sdc2 | -4.58E-01 | 8.192543 | 1.824194 | 0.190684 | 0.357917 | Healthy Control | Intestine |
| Sdcbp | -2.13E+00 | 9.975386 | 115.1905 | 3.67E-10 | 1.44E-08 | Healthy Control | Intestine |
| Sdccag8 | 2.31E+00 | 6.469454 | 58.10758 | 1.40E-07 | 2.09E-06 | Infection | Intestine |
| Sdha | -2.62E+00 | 10.59634 | 191.8918 | 2.80E-12 | 2.65E-10 | Healthy Control | Intestine |
| Sdhb | -2.55E+00 | 10.85449 | 182.3596 | 4.68E-12 | 3.90E-10 | Healthy Control | Intestine |
| Sdhc | -1.32E+00 | 10.62855 | 63.18985 | 7.01E-08 | 1.20E-06 | Healthy Control | Intestine |
| Sdk1 | 3.93E-01 | 5.743955 | 0.371957 | 0.54786 | 0.778138 | Infection | Intestine |
| Sdk2 | 1.91E+00 | 3.392544 | 2.97977 | 0.096888 | 0.211063 | Infection | Intestine |
| Sdr16c5 | -2.68E+00 | 1.879291 | 0.110812 | 0.742697 | 0.92188 | Healthy Control | Intestine |
| Sec1 | -2.86E+00 | 1.484994 | 0.119362 | 0.732724 | 0.914945 | Healthy Control | Intestine |
| Sec22b | -4.69E-01 | 8.397438 | 3.368202 | 0.080147 | 0.183038 | Healthy Control | Intestine |

| | | | | | | | |
|---|---|---|---|---|---|---|---|
| Sec23a | -4.79E-02 | 7.824144 | 0.027912 | 0.868851 | 0.960878 | Healthy Control | Intestine |
| Sec24c | -1.47E+00 | 9.021953 | 33.42192 | 8.29E-06 | 7.06E-05 | Healthy Control | Intestine |
| Sec61a1 | -2.75E+00 | 9.627397 | 145.3051 | 3.30E-11 | 2.02E-09 | Healthy Control | Intestine |
| Sel1l | -2.54E-02 | 9.191704 | 0.013901 | 0.907225 | 0.969735 | Healthy Control | Intestine |
| Sele | 3.89E+00 | 1.634449 | 1.678034 | 0.205764 | 0.380095 | Infection | Intestine |
| Selenof | 1.19E+00 | 9.491076 | 74.36636 | 1.78E-08 | 3.60E-07 | Infection | Intestine |
| Selenoh | 3.32E-01 | 5.405442 | 0.18385 | 0.671939 | 0.877942 | Infection | Intestine |
| Selenoi | -4.12E-01 | 6.826622 | 0.790189 | 0.383163 | 0.612229 | Healthy Control | Intestine |
| Selenok | -4.07E+00 | 2.061853 | 0.404474 | 0.530765 | 0.764827 | Healthy Control | Intestine |
| Selenon | 6.31E-01 | 6.229368 | 1.444703 | 0.242064 | 0.431543 | Infection | Intestine |
| Selenoo | -9.75E-01 | 8.113508 | 11.59245 | 0.002543 | 0.010159 | Healthy Control | Intestine |
| Selenos | -2.94E+00 | 8.602491 | 55.30791 | 1.13E-07 | 1.77E-06 | Healthy Control | Intestine |
| Selenov | -1.52E+00 | 1.273811 | 0.0394 | 0.844976 | 0.954066 | Healthy Control | Intestine |
| Sell | 3.16E+00 | 3.497101 | 6.152544 | 0.022062 | 0.064209 | Infection | Intestine |
| Selp | -1.75E-01 | 3.518315 | 0.004943 | 0.944618 | 0.974399 | Healthy Control | Intestine |
| Selplg | 1.80E+00 | 6.035182 | 10.63467 | 0.003601 | 0.013738 | Infection | Intestine |
| Sema3a | 2.47E+00 | 5.924333 | 17.45024 | 0.000478 | 0.002447 | Infection | Intestine |
| Sema3b | -2.68E+00 | 8.163935 | 45.05387 | 5.96E-07 | 7.23E-06 | Healthy Control | Intestine |
| Sema3c | -2.75E+00 | 8.63981 | 37.12207 | 3.11E-06 | 2.93E-05 | Healthy Control | Intestine |
| Sema3d | 1.95E+00 | 5.151184 | 2.270922 | 0.152353 | 0.301693 | Infection | Intestine |
| Sema3e | 1.11E+00 | 2.582792 | 0.338194 | 0.566562 | 0.794649 | Infection | Intestine |
| Sema3f | -1.85E+00 | 7.052423 | 8.64868 | 0.007037 | 0.024496 | Healthy Control | Intestine |
| Sema4a | -3.11E+00 | 7.922704 | 22.66592 | 7.43E-05 | 0.000479 | Healthy Control | Intestine |
| Sema4d | -2.13E-01 | 6.965849 | 0.270143 | 0.608326 | 0.831483 | Healthy Control | Intestine |
| Sema4g | -4.15E+00 | 11.11057 | 356.1906 | 2.40E-15 | 6.69E-13 | Healthy Control | Intestine |
| Sema5a | -2.10E+00 | 8.484415 | 26.9369 | 2.83E-05 | 0.000204 | Healthy Control | Intestine |
| Sema5b | -3.55E+00 | 2.039006 | 0.204315 | 0.655299 | 0.865091 | Healthy Control | Intestine |
| Sema6a | -1.54E+00 | 7.892496 | 12.82496 | 0.001559 | 0.006701 | Healthy Control | Intestine |
| Sema6c | 1.69E+00 | 3.907699 | 2.796858 | 0.107415 | 0.229304 | Infection | Intestine |
| Sema6d | -8.88E-01 | 7.075689 | 1.938714 | 0.177201 | 0.338575 | Healthy Control | Intestine |
| Senp3 | -4.83E-01 | 6.629464 | 1.205866 | 0.283417 | 0.487555 | Healthy Control | Intestine |
| 10-Sep | -1.41E+00 | 6.928051 | 5.645905 | 0.025934 | 0.073183 | Healthy Control | Intestine |
| 11-Sep | 1.29E+00 | 8.901856 | 50.88141 | 3.95E-07 | 5.08E-06 | Infection | Intestine |
| 2-Sep | 8.45E-01 | 9.641918 | 61.23294 | 9.04E-08 | 1.47E-06 | Infection | Intestine |
| 3-Sep | -4.09E+00 | 2.082164 | 0.363653 | 0.552197 | 0.781891 | Healthy Control | Intestine |
| 4-Sep | 1.55E+00 | 7.227016 | 19.79922 | 0.000205 | 0.001167 | Infection | Intestine |
| 5-Sep | -1.56E+00 | 8.51321 | 20.8009 | 0.000155 | 0.000901 | Healthy Control | Intestine |
| 6-Sep | 9.50E-01 | 6.524422 | 6.722247 | 0.016651 | 0.050637 | Infection | Intestine |
| 7-Sep | 2.03E-01 | 8.701697 | 0.878167 | 0.358981 | 0.58313 | Infection | Intestine |
| 8-Sep | 1.01E+00 | 8.044689 | 24.12172 | 6.70E-05 | 0.000436 | Infection | Intestine |
| 9-Sep | 9.55E-01 | 8.357713 | 20.98851 | 0.000149 | 0.000872 | Infection | Intestine |
| Serpina3a | -2.06E+00 | 1.275767 | 0.073089 | 0.790362 | 0.942036 | Healthy Control | Intestine |
| Serpina3n | 4.44E+00 | 6.644974 | 158.2325 | 1.37E-10 | 6.87E-09 | Infection | Intestine |
| Serpinb13 | -1.52E+00 | 1.274292 | 0.038495 | 0.846741 | 0.954066 | Healthy Control | Intestine |
| Serpinb3b | -2.09E+00 | 1.446014 | 0.055559 | 0.815193 | 0.95147 | Healthy Control | Intestine |
| Serpinb5 | -3.24E+00 | 8.53222 | 51.1395 | 2.27E-07 | 3.13E-06 | Healthy Control | Intestine |
| Serpinc1 | 3.91E-01 | 3.369242 | 0.058743 | 0.810673 | 0.949736 | Infection | Intestine |
| Serpine1 | 5.36E+00 | 2.983734 | 63.5054 | 4.99E-08 | 8.99E-07 | Infection | Intestine |
| Serpine2 | 4.09E+00 | 5.655007 | 66.5225 | 1.49E-07 | 2.21E-06 | Infection | Intestine |
| Serpinf2 | -4.43E+00 | 2.280011 | 0.541798 | 0.469083 | 0.704117 | Healthy Control | Intestine |

| | | | | | | | |
|---|---|---|---|---|---|---|---|
| Serpini1 | -6.99E+00 | 4.316846 | 2.695083 | 0.11648 | 0.24468 | Healthy Control | Intestine |
| Sertad1 | 9.70E-01 | 5.047738 | 1.944463 | 0.176217 | 0.33721 | Infection | Intestine |
| Sesn1 | -6.15E-02 | 6.970923 | 0.016991 | 0.897472 | 0.969735 | Healthy Control | Intestine |
| Sesn2 | 1.33E+00 | 6.776408 | 13.79539 | 0.001222 | 0.005454 | Infection | Intestine |
| Set | 1.97E-01 | 7.923016 | 0.675922 | 0.419879 | 0.651756 | Infection | Intestine |
| Setbp1 | -1.25E+00 | 7.732483 | 9.519807 | 0.005282 | 0.019159 | Healthy Control | Intestine |
| Setd1a | 7.61E-02 | 8.248594 | 0.089655 | 0.767457 | 0.932403 | Infection | Intestine |
| Setd1b | -5.09E-01 | 8.447859 | 5.179129 | 0.033028 | 0.088559 | Healthy Control | Intestine |
| Setd5 | -1.93E-01 | 9.027989 | 1.386841 | 0.251649 | 0.445166 | Healthy Control | Intestine |
| Setd6 | 1.64E+00 | 6.238833 | 14.64642 | 0.000932 | 0.004364 | Infection | Intestine |
| Setdb1 | -1.21E-01 | 7.269408 | 0.105263 | 0.748625 | 0.924698 | Healthy Control | Intestine |
| Setx | -7.13E-01 | 7.505202 | 4.065848 | 0.055903 | 0.137181 | Healthy Control | Intestine |
| Sez6 | -2.45E+00 | 1.56114 | 0.117754 | 0.733932 | 0.915572 | Healthy Control | Intestine |
| Sez6l2 | 2.21E-01 | 4.261052 | 0.028553 | 0.867229 | 0.96049 | Infection | Intestine |
| Sf1 | -6.37E-01 | 9.950889 | 13.68852 | 0.001266 | 0.005629 | Healthy Control | Intestine |
| Sf3a1 | -1.29E+00 | 8.371161 | 19.01774 | 0.000248 | 0.001377 | Healthy Control | Intestine |
| Sf3b6 | -2.11E+00 | 8.364069 | 33.32232 | 6.90E-06 | 5.97E-05 | Healthy Control | Intestine |
| Sfpq | 6.06E-01 | 10.28268 | 32.53856 | 1.01E-05 | 8.37E-05 | Infection | Intestine |
| Sfrp1 | 1.03E-02 | 8.434941 | 0.000521 | 0.982049 | 1 | Infection | Intestine |
| Sfrp2 | -4.69E+00 | 2.929203 | 0.480481 | 0.494941 | 0.727774 | Healthy Control | Intestine |
| Sfrp4 | 2.36E+00 | 5.754487 | 2.547403 | 0.133535 | 0.271672 | Infection | Intestine |
| Sfxn1 | -1.12E+00 | 8.728609 | 15.3795 | 0.000731 | 0.003505 | Healthy Control | Intestine |
| Sfxn3 | -9.83E-01 | 8.556488 | 16.66248 | 0.000482 | 0.00246 | Healthy Control | Intestine |
| Sgca | -2.91E+00 | 1.69893 | 0.179583 | 0.674682 | 0.879138 | Healthy Control | Intestine |
| Sgcd | 2.45E+00 | 5.945522 | 16.69047 | 0.000559 | 0.002791 | Infection | Intestine |
| Sgms2 | -1.25E+00 | 8.359766 | 13.83122 | 0.001187 | 0.005321 | Healthy Control | Intestine |
| Sgo1 | 6.64E-01 | 5.47 | 0.692466 | 0.413967 | 0.646791 | Infection | Intestine |
| Sgo2a | -3.24E-01 | 5.401088 | 0.190643 | 0.66617 | 0.872997 | Healthy Control | Intestine |
| Sgpl1 | -4.62E-01 | 7.564224 | 1.712816 | 0.203997 | 0.377388 | Healthy Control | Intestine |
| Sgsh | -7.44E-01 | 6.640943 | 2.390296 | 0.135459 | 0.274984 | Healthy Control | Intestine |
| Sgta | -7.96E-02 | 8.436201 | 0.099842 | 0.755028 | 0.927071 | Healthy Control | Intestine |
| Sh2d1a | -3.78E+00 | 2.055744 | 0.221417 | 0.642014 | 0.855904 | Healthy Control | Intestine |
| Sh3bp1 | -1.39E-01 | 7.840035 | 0.160845 | 0.692267 | 0.890099 | Healthy Control | Intestine |
| Sh3bp2 | -9.31E-01 | 6.338034 | 1.674648 | 0.208099 | 0.383135 | Healthy Control | Intestine |
| Sh3bp5 | -2.83E+00 | 7.26107 | 10.48833 | 0.003481 | 0.013342 | Healthy Control | Intestine |
| Sh3gl1 | 1.46E+00 | 7.77417 | 44.08226 | 1.18E-06 | 1.29E-05 | Infection | Intestine |
| Sh3gl2 | -1.97E+00 | 6.401806 | 7.187038 | 0.012901 | 0.040822 | Healthy Control | Intestine |
| Sh3gl3 | 1.17E+00 | 3.466527 | 0.606662 | 0.443586 | 0.677217 | Infection | Intestine |
| Sh3rf2 | -2.87E+00 | 6.802531 | 6.466163 | 0.018739 | 0.056037 | Healthy Control | Intestine |
| Sh3tc2 | 1.89E+00 | 3.065731 | 1.680059 | 0.207305 | 0.382086 | Infection | Intestine |
| Shank3 | -6.48E-01 | 4.800669 | 0.255118 | 0.618178 | 0.837736 | Healthy Control | Intestine |
| Sharpin | -1.85E+00 | 8.292202 | 29.42816 | 1.71E-05 | 0.000133 | Healthy Control | Intestine |
| Shc1 | -2.99E-02 | 8.122112 | 0.014621 | 0.904864 | 0.969735 | Healthy Control | Intestine |
| Shcbp1l | -1.49E+00 | 1.442309 | 0.047535 | 0.829036 | 0.954066 | Healthy Control | Intestine |
| She | 4.64E-01 | 6.102433 | 0.397863 | 0.534585 | 0.76658 | Infection | Intestine |
| Shh | -6.06E+00 | 2.929932 | 1.372334 | 0.253217 | 0.447127 | Healthy Control | Intestine |
| Shisa3 | -1.16E+00 | 4.747232 | 0.269391 | 0.608975 | 0.831569 | Healthy Control | Intestine |
| Shld1 | 2.95E-01 | 4.023473 | 0.054263 | 0.817837 | 0.952901 | Infection | Intestine |
| Shmt1 | -1.43E+00 | 8.59407 | 16.21408 | 0.000554 | 0.002769 | Healthy Control | Intestine |
| Shoc2 | -8.56E-01 | 8.520728 | 11.92088 | 0.002271 | 0.009221 | Healthy Control | Intestine |

| | | | | | | | |
|---|---|---|---|---|---|---|---|
| Shox2 | -2.78E+00 | 1.447783 | 0.101197 | 0.753083 | 0.925708 | Healthy Control | Intestine |
| Shroom3 | -2.73E+00 | 10.10511 | 143.317 | 4.33E-11 | 2.49E-09 | Healthy Control | Intestine |
| Shtn1 | -2.19E+00 | 7.558189 | 20.90784 | 0.000122 | 0.000734 | Healthy Control | Intestine |
| Siah2 | -1.67E+00 | 6.738225 | 8.39564 | 0.007752 | 0.026542 | Healthy Control | Intestine |
| Sidt1 | -9.71E+00 | 6.464191 | 15.40758 | 0.000851 | 0.00401 | Healthy Control | Intestine |
| Sidt2 | 1.78E+00 | 6.903628 | 37.54809 | 3.79E-06 | 3.49E-05 | Infection | Intestine |
| Siglec1 | -2.12E+00 | 5.79094 | 2.908837 | 0.101733 | 0.219895 | Healthy Control | Intestine |
| Sigmar1 | 4.67E-01 | 7.965716 | 2.604454 | 0.120905 | 0.251245 | Infection | Intestine |
| Sim1 | -2.28E+00 | 1.363513 | 0.069906 | 0.793871 | 0.943769 | Healthy Control | Intestine |
| Sin3a | -7.19E-01 | 8.098666 | 3.702726 | 0.067317 | 0.160015 | Healthy Control | Intestine |
| Sin3b | 6.27E-01 | 6.530882 | 2.849937 | 0.105478 | 0.225941 | Infection | Intestine |
| Sinhcaf | 1.54E-01 | 6.90267 | 0.093383 | 0.762711 | 0.929438 | Infection | Intestine |
| Sirpa | 2.31E+00 | 7.8283 | 125.8933 | 1.59E-10 | 7.48E-09 | Infection | Intestine |
| Sirt1 | 9.49E-02 | 6.41765 | 0.038142 | 0.846911 | 0.954066 | Infection | Intestine |
| Sirt2 | -2.52E-01 | 8.205141 | 1.237237 | 0.278128 | 0.480769 | Healthy Control | Intestine |
| Sirt3 | -3.11E+00 | 6.454824 | 9.419743 | 0.005487 | 0.019789 | Healthy Control | Intestine |
| Sirt4 | -5.03E-01 | 5.969196 | 0.534764 | 0.471713 | 0.706752 | Healthy Control | Intestine |
| Sirt5 | 4.28E-01 | 6.07443 | 0.66229 | 0.424155 | 0.656764 | Infection | Intestine |
| Sirt6 | -1.79E+00 | 7.046134 | 9.803205 | 0.004524 | 0.016752 | Healthy Control | Intestine |
| Sirt7 | -3.91E-01 | 8.316492 | 2.654556 | 0.117603 | 0.245914 | Healthy Control | Intestine |
| Six1 | -2.04E+00 | 1.443618 | 0.068002 | 0.79602 | 0.943769 | Healthy Control | Intestine |
| Six2 | -4.52E+00 | 1.83708 | 0.527845 | 0.473511 | 0.708216 | Healthy Control | Intestine |
| Six3 | -1.65E+00 | 1.319049 | 0.045517 | 0.833004 | 0.954066 | Healthy Control | Intestine |
| Six6 | -1.77E+00 | 1.320018 | 0.057114 | 0.81363 | 0.950532 | Healthy Control | Intestine |
| Ska1 | 1.75E+00 | 3.877802 | 2.371039 | 0.13674 | 0.277021 | Infection | Intestine |
| Ska2 | 6.40E-01 | 6.548857 | 2.158796 | 0.155953 | 0.306822 | Infection | Intestine |
| Skp1a | 7.35E-01 | 7.211945 | 5.105966 | 0.034185 | 0.091173 | Infection | Intestine |
| Skp2 | 8.66E-01 | 4.71117 | 1.133358 | 0.297659 | 0.506487 | Infection | Intestine |
| Slamf6 | -1.63E-01 | 3.421932 | 0.005931 | 0.939327 | 0.97034 | Healthy Control | Intestine |
| Slamf9 | -7.37E+00 | 4.712293 | 2.342683 | 0.141704 | 0.285345 | Healthy Control | Intestine |
| Slc11a1 | -2.06E+00 | 5.60095 | 2.036094 | 0.167165 | 0.322849 | Healthy Control | Intestine |
| Slc11a2 | -3.47E-01 | 6.636319 | 0.316326 | 0.57925 | 0.806375 | Healthy Control | Intestine |
| Slc12a2 | -3.47E+00 | 10.18734 | 90.45675 | 2.77E-09 | 7.76E-08 | Healthy Control | Intestine |
| Slc12a5 | -4.99E+00 | 2.472872 | 0.542498 | 0.469595 | 0.704674 | Healthy Control | Intestine |
| Slc12a7 | -1.13E+00 | 8.358597 | 12.78849 | 0.001683 | 0.007126 | Healthy Control | Intestine |
| Slc13a4 | -9.71E-16 | 1.180616 | 0 | 1 | 1 | Healthy Control | Intestine |
| Slc14a1 | -5.26E+00 | 3.066941 | 0.669487 | 0.422853 | 0.655154 | Healthy Control | Intestine |
| Slc15a1 | -3.38E+00 | 8.360887 | 13.53569 | 0.001749 | 0.007382 | Healthy Control | Intestine |
| Slc16a1 | -2.57E+00 | 9.84928 | 59.14715 | 9.94E-08 | 1.58E-06 | Healthy Control | Intestine |
| Slc16a2 | 2.93E+00 | 4.097442 | 11.46398 | 0.002656 | 0.010535 | Infection | Intestine |
| Slc16a3 | 3.59E+00 | 5.235135 | 77.92781 | 1.22E-08 | 2.61E-07 | Infection | Intestine |
| Slc16a8 | -2.84E+00 | 1.485906 | 0.146458 | 0.704685 | 0.896371 | Healthy Control | Intestine |
| Slc17a2 | -5.15E+00 | 2.702174 | 0.910923 | 0.348596 | 0.572154 | Healthy Control | Intestine |
| Slc17a6 | -3.57E-01 | 1.227417 | 0.006288 | 0.937908 | 0.969735 | Healthy Control | Intestine |
| Slc17a7 | -4.41E+00 | 2.207113 | 0.586441 | 0.451772 | 0.685325 | Healthy Control | Intestine |
| Slc17a8 | -3.48E+00 | 1.913645 | 0.203196 | 0.655781 | 0.865113 | Healthy Control | Intestine |
| Slc18a2 | 4.87E+00 | 1.962442 | 23.76894 | 2.21E-05 | 0.000166 | Infection | Intestine |
| Slc18a3 | -3.57E-01 | 1.227067 | 0.006351 | 0.937596 | 0.969735 | Healthy Control | Intestine |
| Slc19a1 | -1.77E+00 | 6.113879 | 4.3854 | 0.046999 | 0.118796 | Healthy Control | Intestine |
| Slc19a2 | -3.83E-01 | 5.326716 | 0.249746 | 0.621811 | 0.839021 | Healthy Control | Intestine |

| | | | | | | | |
|---|---|---|---|---|---|---|---|
| Slc19a3 | -1.80E+00 | 1.522322 | 0.059129 | 0.809622 | 0.94887 | Healthy Control | Intestine |
| Slc1a1 | -4.18E+00 | 7.846371 | 25.92612 | 4.30E-05 | 0.000294 | Healthy Control | Intestine |
| Slc1a3 | -4.16E-02 | 4.120212 | 0.001248 | 0.972127 | 0.993765 | Healthy Control | Intestine |
| Slc20a2 | -7.05E-01 | 6.141836 | 1.098124 | 0.305077 | 0.516288 | Healthy Control | Intestine |
| Slc22a18 | -3.30E+00 | 6.68432 | 7.817025 | 0.010303 | 0.03371 | Healthy Control | Intestine |
| Slc22a2 | -1.65E+00 | 1.318582 | 0.047143 | 0.830096 | 0.954066 | Healthy Control | Intestine |
| Slc22a5 | -2.88E+00 | 7.195326 | 17.32282 | 0.000345 | 0.001838 | Healthy Control | Intestine |
| Slc22a6 | -9.71E-16 | 1.180616 | 0 | 1 | 1 | Healthy Control | Intestine |
| Slc22a8 | -1.07E+00 | 1.319243 | 0.019177 | 0.891274 | 0.969735 | Healthy Control | Intestine |
| Slc24a3 | 1.36E+00 | 6.176755 | 9.701563 | 0.005033 | 0.018365 | Infection | Intestine |
| Slc24a4 | -1.71E+00 | 1.318343 | 0.050771 | 0.823952 | 0.953685 | Healthy Control | Intestine |
| Slc25a17 | -1.36E-01 | 8.130577 | 0.256643 | 0.617523 | 0.837736 | Healthy Control | Intestine |
| Slc25a20 | -2.15E+00 | 9.499195 | 31.3885 | 1.41E-05 | 0.000112 | Healthy Control | Intestine |
| Slc25a22 | -4.82E-02 | 7.019164 | 0.014219 | 0.906154 | 0.969735 | Healthy Control | Intestine |
| Slc25a28 | 4.34E-01 | 7.959471 | 2.5296 | 0.126116 | 0.25917 | Infection | Intestine |
| Slc25a31 | -2.22E+00 | 1.520275 | 0.083542 | 0.774579 | 0.934669 | Healthy Control | Intestine |
| Slc25a51 | -1.63E+00 | 7.217417 | 8.548173 | 0.007447 | 0.025692 | Healthy Control | Intestine |
| Slc26a3 | -7.27E+00 | 10.04795 | 56.09038 | 1.72E-06 | 1.75E-05 | Healthy Control | Intestine |
| Slc26a4 | -3.14E+00 | 1.666423 | 0.143137 | 0.707606 | 0.897011 | Healthy Control | Intestine |
| Slc26a5 | -2.81E+00 | 1.485159 | 0.121274 | 0.730004 | 0.91327 | Healthy Control | Intestine |
| Slc2a1 | -3.17E-01 | 8.209197 | 0.999248 | 0.328445 | 0.545676 | Healthy Control | Intestine |
| Slc2a2 | -1.97E+00 | 1.594621 | 0.069901 | 0.793788 | 0.943769 | Healthy Control | Intestine |
| Slc2a3 | 4.53E+00 | 4.230294 | 77.2572 | 1.35E-08 | 2.86E-07 | Infection | Intestine |
| Slc2a4 | 1.37E+00 | 5.187177 | 4.320532 | 0.049339 | 0.123744 | Infection | Intestine |
| Slc2a5 | -6.00E+00 | 3.745741 | 1.448999 | 0.242861 | 0.432347 | Healthy Control | Intestine |
| Slc30a10 | -1.69E+00 | 6.622726 | 1.047623 | 0.319185 | 0.53368 | Healthy Control | Intestine |
| Slc30a3 | -9.77E-01 | 1.318411 | 0.021477 | 0.884798 | 0.96867 | Healthy Control | Intestine |
| Slc30a8 | -1.88E+00 | 1.362179 | 0.06514 | 0.800847 | 0.946422 | Healthy Control | Intestine |
| Slc31a1 | -3.15E+00 | 8.047696 | 33.35983 | 6.71E-06 | 5.81E-05 | Healthy Control | Intestine |
| Slc32a1 | -4.95E-01 | 1.228692 | 0.008683 | 0.927069 | 0.969735 | Healthy Control | Intestine |
| Slc39a5 | -4.00E+00 | 7.435677 | 28.00555 | 2.31E-05 | 0.000173 | Healthy Control | Intestine |
| Slc39a8 | -2.63E+00 | 5.918513 | 4.300728 | 0.049722 | 0.124393 | Healthy Control | Intestine |
| Slc3a1 | -2.94E+00 | 5.552577 | 4.510719 | 0.045886 | 0.117078 | Healthy Control | Intestine |
| Slc40a1 | -1.01E+00 | 9.081244 | 3.966895 | 0.063234 | 0.151676 | Healthy Control | Intestine |
| Slc44a5 | -3.93E+00 | 1.804089 | 0.191375 | 0.664826 | 0.87192 | Healthy Control | Intestine |
| Slc45a1 | -1.94E+00 | 1.402999 | 0.05698 | 0.813003 | 0.950244 | Healthy Control | Intestine |
| Slc45a2 | -8.79E-01 | 1.316883 | 0.01296 | 0.910361 | 0.969735 | Healthy Control | Intestine |
| Slc47a1 | -5.06E+00 | 2.975909 | 0.533477 | 0.472269 | 0.707202 | Healthy Control | Intestine |
| Slc4a1 | -3.24E+00 | 1.482968 | 0.159329 | 0.693499 | 0.890686 | Healthy Control | Intestine |
| Slc4a10 | -2.25E+00 | 1.363548 | 0.059617 | 0.80916 | 0.94887 | Healthy Control | Intestine |
| Slc4a7 | -1.43E+00 | 9.194047 | 28.99442 | 2.10E-05 | 0.000159 | Healthy Control | Intestine |
| Slc4a9 | -7.56E-01 | 1.273399 | 0.015556 | 0.902167 | 0.969735 | Healthy Control | Intestine |
| Slc52a3 | -4.04E+00 | 7.321692 | 21.45499 | 0.000122 | 0.000735 | Healthy Control | Intestine |
| Slc5a12 | -1.37E+00 | 1.228088 | 0.037791 | 0.848672 | 0.954066 | Healthy Control | Intestine |
| Slc5a2 | -3.57E-01 | 1.227067 | 0.006351 | 0.937596 | 0.969735 | Healthy Control | Intestine |
| Slc5a7 | -2.04E+00 | 1.444306 | 0.064937 | 0.800565 | 0.946311 | Healthy Control | Intestine |
| Slc6a11 | 5.80E-01 | 5.057834 | 0.360904 | 0.553677 | 0.782305 | Infection | Intestine |
| Slc6a14 | -3.90E+00 | 11.2779 | 187.4089 | 2.43E-12 | 2.34E-10 | Healthy Control | Intestine |
| Slc6a19 | -5.14E+00 | 2.521308 | 0.722758 | 0.404096 | 0.634936 | Healthy Control | Intestine |
| Slc6a3 | -3.57E-01 | 1.227387 | 0.006293 | 0.937883 | 0.969735 | Healthy Control | Intestine |

| | | | | | | | |
|---|---|---|---|---|---|---|---|
| Slc6a4 | -2.52E+00 | 6.309227 | 5.566575 | 0.027718 | 0.077302 | Healthy Control | Intestine |
| Slc6a5 | -6.42E-01 | 1.272688 | 0.013552 | 0.908542 | 0.969735 | Healthy Control | Intestine |
| Slc6a8 | -3.15E+00 | 9.545364 | 41.17613 | 2.17E-06 | 2.13E-05 | Healthy Control | Intestine |
| Slc6a9 | -3.90E-01 | 5.765217 | 0.358027 | 0.555186 | 0.783995 | Healthy Control | Intestine |
| Slc7a10 | -1.83E+00 | 1.360646 | 0.061908 | 0.805499 | 0.947891 | Healthy Control | Intestine |
| Slc8a1 | 7.41E-01 | 8.987635 | 8.923914 | 0.006835 | 0.023911 | Infection | Intestine |
| Slc8a3 | -5.42E+00 | 2.846884 | 1.411247 | 0.24895 | 0.441931 | Healthy Control | Intestine |
| Slc8b1 | -1.88E+00 | 8.846954 | 34.20878 | 5.42E-06 | 4.79E-05 | Healthy Control | Intestine |
| Slc9a1 | -1.75E-01 | 7.585182 | 0.342761 | 0.564243 | 0.792537 | Healthy Control | Intestine |
| Slc9a3 | -5.97E+00 | 8.533407 | 15.1044 | 0.001997 | 0.008248 | Healthy Control | Intestine |
| Slc9a3r1 | -2.83E+00 | 8.622886 | 26.07163 | 3.32E-05 | 0.000235 | Healthy Control | Intestine |
| Slc9a4 | -8.62E-01 | 1.275264 | 0.013825 | 0.907842 | 0.969735 | Healthy Control | Intestine |
| Slc9a8 | -1.08E+00 | 7.611398 | 8.920818 | 0.00668 | 0.023433 | Healthy Control | Intestine |
| Slc9a9 | 7.92E-01 | 6.007782 | 1.45832 | 0.239904 | 0.428608 | Infection | Intestine |
| Slco1a1 | -6.42E-01 | 1.273464 | 0.012824 | 0.911022 | 0.969735 | Healthy Control | Intestine |
| Slco3a1 | 1.20E+00 | 5.926213 | 3.03762 | 0.095338 | 0.208685 | Infection | Intestine |
| Slco4c1 | 5.85E+00 | 1.365128 | 10.25294 | 0.004583 | 0.016958 | Infection | Intestine |
| Slit1 | -4.45E+00 | 2.133544 | 0.319916 | 0.578308 | 0.806375 | Healthy Control | Intestine |
| Slit2 | -6.94E-01 | 3.370872 | 0.308706 | 0.58417 | 0.810295 | Healthy Control | Intestine |
| Slit3 | 2.93E+00 | 7.067415 | 64.5696 | 1.14E-07 | 1.78E-06 | Infection | Intestine |
| Slitrk1 | -2.68E+00 | 1.559149 | 0.130128 | 0.720501 | 0.906205 | Healthy Control | Intestine |
| Slitrk3 | -3.87E+00 | 1.925661 | 0.324076 | 0.573631 | 0.80173 | Healthy Control | Intestine |
| Slk | -4.17E-01 | 10.24263 | 8.142591 | 0.00929 | 0.030839 | Healthy Control | Intestine |
| Slurp2 | -1.65E+00 | 1.318742 | 0.046529 | 0.831187 | 0.954066 | Healthy Control | Intestine |
| Smad1 | -1.44E+00 | 9.006316 | 43.51641 | 1.22E-06 | 1.33E-05 | Healthy Control | Intestine |
| Smad2 | -7.04E-01 | 8.123502 | 6.159279 | 0.021178 | 0.061943 | Healthy Control | Intestine |
| Smad3 | 1.66E+00 | 7.795329 | 44.11994 | 1.17E-06 | 1.28E-05 | Infection | Intestine |
| Smad4 | -6.89E-01 | 9.653119 | 11.44548 | 0.002701 | 0.010708 | Healthy Control | Intestine |
| Smad5 | -1.86E+00 | 7.446115 | 14.6806 | 0.000816 | 0.003866 | Healthy Control | Intestine |
| Smad6 | -1.63E+00 | 6.541947 | 6.405041 | 0.018165 | 0.05468 | Healthy Control | Intestine |
| Smad7 | -1.28E+00 | 8.409077 | 12.54596 | 0.001816 | 0.00762 | Healthy Control | Intestine |
| Smad9 | -5.93E-01 | 4.076508 | 0.123317 | 0.728727 | 0.912515 | Healthy Control | Intestine |
| Smap1 | -8.35E-02 | 8.383244 | 0.093079 | 0.763197 | 0.929804 | Healthy Control | Intestine |
| Smap2 | 1.72E-01 | 7.223966 | 0.310624 | 0.582932 | 0.809026 | Infection | Intestine |
| Smarca1 | -3.23E-01 | 3.743975 | 0.041658 | 0.840132 | 0.954066 | Healthy Control | Intestine |
| Smarca2 | -2.55E-01 | 8.108958 | 0.990443 | 0.330556 | 0.548586 | Healthy Control | Intestine |
| Smarca4 | -6.79E-01 | 9.411454 | 15.54173 | 0.000705 | 0.003403 | Healthy Control | Intestine |
| Smarca5 | -2.47E+00 | 8.049251 | 43.25358 | 8.58E-07 | 9.87E-06 | Healthy Control | Intestine |
| Smarcb1 | -3.63E-01 | 7.85251 | 1.291254 | 0.268117 | 0.467337 | Healthy Control | Intestine |
| Smarcc1 | -9.42E-01 | 7.975807 | 8.790208 | 0.00713 | 0.024788 | Healthy Control | Intestine |
| Smarcc2 | 8.64E-01 | 8.985824 | 38.17137 | 3.35E-06 | 3.15E-05 | Infection | Intestine |
| Smarcd1 | 3.61E-01 | 6.547854 | 0.570511 | 0.457998 | 0.692251 | Infection | Intestine |
| Smarcd2 | -7.05E-01 | 8.616884 | 9.425935 | 0.00562 | 0.020195 | Healthy Control | Intestine |
| Smarcd3 | 1.16E+00 | 5.425158 | 3.29118 | 0.083114 | 0.187837 | Infection | Intestine |
| Smarce1 | -1.04E+00 | 7.234926 | 5.578224 | 0.02698 | 0.075707 | Healthy Control | Intestine |
| Smc1a | 8.34E-01 | 8.764005 | 21.37558 | 0.000134 | 0.000796 | Infection | Intestine |
| Smc3 | -6.48E-01 | 7.864327 | 5.321055 | 0.030853 | 0.084032 | Healthy Control | Intestine |
| Smc5 | -9.95E-01 | 7.501637 | 5.92047 | 0.023344 | 0.067355 | Healthy Control | Intestine |
| Smcp | -1.37E+00 | 1.228088 | 0.037791 | 0.848672 | 0.954066 | Healthy Control | Intestine |
| Smdt1 | -3.05E+00 | 10.38639 | 159.6402 | 1.61E-11 | 1.07E-09 | Healthy Control | Intestine |

| | | | | | | | |
|---|---|---|---|---|---|---|---|
| Smg9 | -2.79E-01 | 7.198569 | 0.455472 | 0.506706 | 0.741648 | Healthy Control | Intestine |
| Smgc | 4.08E+00 | 1.443564 | 3.618532 | 0.066762 | 0.158996 | Infection | Intestine |
| Smim7 | 7.40E-01 | 8.356047 | 14.2642 | 0.001052 | 0.004836 | Infection | Intestine |
| Smn1 | -2.45E-02 | 6.937043 | 0.003336 | 0.954459 | 0.981314 | Healthy Control | Intestine |
| Smo | -2.27E-01 | 6.978389 | 0.301576 | 0.588302 | 0.814156 | Healthy Control | Intestine |
| Smoc1 | 4.64E-01 | 4.210159 | 0.076933 | 0.783777 | 0.940206 | Infection | Intestine |
| Smoc2 | -2.75E+00 | 6.298961 | 5.427338 | 0.029156 | 0.080287 | Healthy Control | Intestine |
| Smpd1 | -1.61E+00 | 7.633988 | 14.17422 | 0.000993 | 0.004599 | Healthy Control | Intestine |
| Smpd3 | -4.77E+00 | 8.145306 | 35.25184 | 5.02E-06 | 4.47E-05 | Healthy Control | Intestine |
| Smpd4 | 8.78E-02 | 5.65214 | 0.01504 | 0.903431 | 0.969735 | Infection | Intestine |
| Smtn | 9.15E-01 | 8.628406 | 11.49747 | 0.002653 | 0.010531 | Infection | Intestine |
| Smurf1 | 1.67E+00 | 7.629628 | 69.33829 | 3.25E-08 | 6.12E-07 | Infection | Intestine |
| Smurf2 | -1.83E-01 | 7.986622 | 0.381268 | 0.543318 | 0.775107 | Healthy Control | Intestine |
| Smyd2 | -5.82E-01 | 7.954223 | 3.474405 | 0.075731 | 0.174704 | Healthy Control | Intestine |
| Smyd3 | 5.65E-01 | 6.283923 | 1.204367 | 0.284212 | 0.488755 | Infection | Intestine |
| Snai1 | 6.39E-01 | 6.043269 | 0.883421 | 0.357204 | 0.581895 | Infection | Intestine |
| Snai2 | -1.39E+00 | 5.665629 | 1.331453 | 0.259752 | 0.455772 | Healthy Control | Intestine |
| Snai3 | -4.82E+00 | 2.504522 | 0.650883 | 0.429086 | 0.661421 | Healthy Control | Intestine |
| Snap25 | -7.07E+00 | 4.085664 | 2.828279 | 0.108359 | 0.230828 | Healthy Control | Intestine |
| Snap29 | -2.36E+00 | 7.255104 | 14.19474 | 0.000959 | 0.004455 | Healthy Control | Intestine |
| Snap91 | -3.51E+00 | 1.893923 | 0.332115 | 0.568816 | 0.797393 | Healthy Control | Intestine |
| Snapin | -7.05E-01 | 7.532675 | 3.248706 | 0.084987 | 0.190867 | Healthy Control | Intestine |
| Sncb | -4.52E+00 | 2.057666 | 0.512742 | 0.4804 | 0.714463 | Healthy Control | Intestine |
| Sncg | -1.88E+00 | 4.977578 | 0.954638 | 0.339234 | 0.559172 | Healthy Control | Intestine |
| Snf8 | -2.33E-01 | 8.856135 | 1.102784 | 0.305173 | 0.516288 | Healthy Control | Intestine |
| Snph | -6.20E-01 | 4.221856 | 0.160884 | 0.692116 | 0.890099 | Healthy Control | Intestine |
| Snrk | -2.34E+00 | 7.913468 | 19.90159 | 0.000165 | 0.000954 | Healthy Control | Intestine |
| Snrpa | -2.62E-02 | 7.760142 | 0.00896 | 0.925448 | 0.969735 | Healthy Control | Intestine |
| Snrpd1 | 1.12E-01 | 4.45906 | 0.008481 | 0.927383 | 0.969735 | Infection | Intestine |
| Snrpn | 1.43E-01 | 5.550978 | 0.028742 | 0.866816 | 0.96049 | Infection | Intestine |
| Snta1 | 1.16E+00 | 4.729057 | 1.60684 | 0.217326 | 0.396492 | Infection | Intestine |
| Sntb1 | 3.39E-01 | 5.54657 | 0.232849 | 0.633943 | 0.849742 | Infection | Intestine |
| Sntn | -1.15E+00 | 1.319686 | 0.040261 | 0.84354 | 0.954066 | Healthy Control | Intestine |
| Snu13 | 1.01E+00 | 5.667758 | 2.786069 | 0.108929 | 0.231844 | Infection | Intestine |
| Snx27 | 6.12E-02 | 6.68341 | 0.022449 | 0.882235 | 0.96867 | Infection | Intestine |
| Snx30 | 2.69E-01 | 6.092271 | 0.249843 | 0.621975 | 0.839021 | Infection | Intestine |
| Snx4 | -2.91E-02 | 7.614275 | 0.012023 | 0.913687 | 0.969735 | Healthy Control | Intestine |
| Snx5 | 1.12E+00 | 10.17791 | 130.2777 | 1.15E-10 | 5.94E-09 | Infection | Intestine |
| Snx6 | -4.00E-01 | 8.033692 | 2.191634 | 0.152986 | 0.302708 | Healthy Control | Intestine |
| Socs1 | 5.07E+00 | 2.537547 | 37.66988 | 2.06E-06 | 2.04E-05 | Infection | Intestine |
| Socs2 | 8.32E-01 | 7.554258 | 5.03593 | 0.035335 | 0.09374 | Infection | Intestine |
| Socs3 | 4.30E+00 | 6.046046 | 163.1863 | 1.35E-11 | 9.38E-10 | Infection | Intestine |
| Socs6 | -5.65E-01 | 7.388871 | 2.144365 | 0.157166 | 0.308781 | Healthy Control | Intestine |
| Sod1 | -3.48E+00 | 10.57885 | 189.5086 | 2.97E-12 | 2.76E-10 | Healthy Control | Intestine |
| Sod2 | -1.86E+00 | 9.16406 | 64.86834 | 5.33E-08 | 9.43E-07 | Healthy Control | Intestine |
| Sohlh2 | 5.06E+00 | 1.319125 | 3.647154 | 0.070835 | 0.165701 | Infection | Intestine |
| Sorbs1 | 1.11E+00 | 8.125851 | 7.292397 | 0.015056 | 0.046439 | Infection | Intestine |
| Sorcs3 | -2.24E+00 | 1.631783 | 0.102505 | 0.751359 | 0.925207 | Healthy Control | Intestine |
| Sorl1 | -1.28E+00 | 8.776118 | 11.88541 | 0.0023 | 0.009317 | Healthy Control | Intestine |
| Sort1 | -3.27E+00 | 10.12402 | 177.1987 | 4.18E-12 | 3.61E-10 | Healthy Control | Intestine |

| | | | | | | | |
|---|---|---|---|---|---|---|---|
| Sos1 | -9.87E-01 | 7.58279 | 10.35156 | 0.003876 | 0.014676 | Healthy Control | Intestine |
| Sost | -3.57E-01 | 1.228165 | 0.006192 | 0.93838 | 0.969735 | Healthy Control | Intestine |
| Sostdc1 | -2.04E+00 | 1.444075 | 0.0667 | 0.797937 | 0.944678 | Healthy Control | Intestine |
| Sox1 | -3.72E+00 | 1.599309 | 0.250044 | 0.621911 | 0.839021 | Healthy Control | Intestine |
| Sox10 | -5.40E+00 | 3.06348 | 0.378779 | 0.54587 | 0.776921 | Healthy Control | Intestine |
| Sox11 | 3.99E+00 | 1.448466 | 2.461965 | 0.127006 | 0.260678 | Infection | Intestine |
| Sox13 | -1.36E+00 | 7.822012 | 10.01907 | 0.004389 | 0.016332 | Healthy Control | Intestine |
| Sox14 | -1.65E+00 | 1.318115 | 0.050007 | 0.825097 | 0.953685 | Healthy Control | Intestine |
| Sox17 | -9.14E-01 | 4.374604 | 0.109496 | 0.744372 | 0.923113 | Healthy Control | Intestine |
| Sox18 | -1.21E+00 | 6.531209 | 0.494129 | 0.492025 | 0.725074 | Healthy Control | Intestine |
| Sox2 | -2.22E+00 | 1.520017 | 0.076966 | 0.783391 | 0.940206 | Healthy Control | Intestine |
| Sox21 | -8.38E+00 | 5.673374 | 6.28169 | 0.021043 | 0.061711 | Healthy Control | Intestine |
| Sox3 | -9.71E-16 | 1.180616 | 0 | 1 | 1 | Healthy Control | Intestine |
| Sox30 | -2.69E+00 | 1.405444 | 0.121421 | 0.730741 | 0.91327 | Healthy Control | Intestine |
| Sox4 | -2.25E-01 | 8.066021 | 0.442915 | 0.512702 | 0.74784 | Healthy Control | Intestine |
| Sox5 | -1.13E+00 | 4.413816 | 0.768206 | 0.390139 | 0.620012 | Healthy Control | Intestine |
| Sox6 | -1.55E+00 | 6.005786 | 2.074888 | 0.162429 | 0.316882 | Healthy Control | Intestine |
| Sox8 | -4.12E+00 | 2.227748 | 0.282154 | 0.600465 | 0.824505 | Healthy Control | Intestine |
| Sox9 | -1.97E+00 | 9.756817 | 27.60151 | 4.14E-05 | 0.000284 | Healthy Control | Intestine |
| Sp1 | 1.16E-01 | 8.242662 | 0.297524 | 0.590983 | 0.816577 | Infection | Intestine |
| Sp3 | 2.33E-01 | 9.015078 | 1.925187 | 0.17932 | 0.341841 | Infection | Intestine |
| Sp5 | -3.90E+00 | 1.866238 | 0.27763 | 0.602168 | 0.826104 | Healthy Control | Intestine |
| Sp7 | -1.74E+00 | 1.484747 | 0.048773 | 0.826757 | 0.953685 | Healthy Control | Intestine |
| Sp8 | -6.42E-01 | 1.273568 | 0.012758 | 0.911248 | 0.969735 | Healthy Control | Intestine |
| Sp9 | 4.39E+00 | 1.444841 | 3.581712 | 0.067506 | 0.160312 | Infection | Intestine |
| Spaca6 | 1.52E-01 | 4.405546 | 0.011461 | 0.915652 | 0.969735 | Infection | Intestine |
| Spaca9 | 4.12E+00 | 2.110121 | 10.44715 | 0.002973 | 0.0116 | Infection | Intestine |
| Spag9 | -6.71E-01 | 7.684751 | 4.318994 | 0.049453 | 0.123904 | Healthy Control | Intestine |
| Spast | 6.72E-02 | 8.698718 | 0.084352 | 0.774233 | 0.934669 | Infection | Intestine |
| Spata25 | -2.24E+00 | 1.361189 | 0.097495 | 0.757531 | 0.928591 | Healthy Control | Intestine |
| Spata4 | -9.71E-16 | 1.180616 | 0 | 1 | 1 | Healthy Control | Intestine |
| Spatc1l | -2.25E+00 | 1.362616 | 0.087691 | 0.769659 | 0.932779 | Healthy Control | Intestine |
| Spdef | -3.69E+00 | 8.784153 | 27.48993 | 4.92E-05 | 0.00033 | Healthy Control | Intestine |
| Spdl1 | 1.41E+00 | 3.809329 | 1.618743 | 0.215267 | 0.393581 | Infection | Intestine |
| Spdye4a | 4.42E+00 | 1.320565 | 3.866097 | 0.064723 | 0.154583 | Infection | Intestine |
| Speg | 2.04E+00 | 4.934678 | 7.344139 | 0.012781 | 0.040535 | Infection | Intestine |
| Sphk1 | 3.53E+00 | 3.649738 | 20.04723 | 0.000182 | 0.001042 | Infection | Intestine |
| Sphk2 | -1.44E+00 | 8.137531 | 21.13536 | 0.000133 | 0.000791 | Healthy Control | Intestine |
| Spi1 | -9.54E-01 | 6.949491 | 2.89185 | 0.102222 | 0.220761 | Healthy Control | Intestine |
| Spib | -1.86E+00 | 5.158271 | 0.25783 | 0.61955 | 0.838379 | Healthy Control | Intestine |
| Spink1 | -1.32E+01 | 9.822217 | 87.49592 | 5.12E-08 | 9.15E-07 | Healthy Control | Intestine |
| Spink12 | -3.57E-01 | 1.227509 | 0.006273 | 0.937979 | 0.969735 | Healthy Control | Intestine |
| Spink8 | -2.16E+00 | 1.318372 | 0.088982 | 0.768665 | 0.932538 | Healthy Control | Intestine |
| Spn | -7.41E-01 | 4.260004 | 0.173462 | 0.680947 | 0.882694 | Healthy Control | Intestine |
| Spo11 | -2.64E+00 | 1.524872 | 0.124028 | 0.726956 | 0.91138 | Healthy Control | Intestine |
| Spock3 | -4.95E+00 | 2.365402 | 0.587604 | 0.45168 | 0.685325 | Healthy Control | Intestine |
| Spon1 | 1.49E+00 | 7.583944 | 17.98586 | 0.000341 | 0.001824 | Infection | Intestine |
| Spop | -3.32E-01 | 9.48918 | 4.208069 | 0.052448 | 0.130171 | Healthy Control | Intestine |
| Spopl | -8.39E-01 | 7.08793 | 4.317953 | 0.048983 | 0.123158 | Healthy Control | Intestine |
| Spred1 | 2.61E-01 | 8.35065 | 1.679749 | 0.208528 | 0.383643 | Infection | Intestine |

| | | | | | | | |
|---|---|---|---|---|---|---|---|
| Sprr2d | -9.71E-16 | 1.180616 | 0 | 1 | 1 | Healthy Control | Intestine |
| Sprr2f | 4.59E+00 | 1.406773 | 3.51378 | 0.071119 | 0.166268 | Infection | Intestine |
| Spry1 | -1.62E-01 | 6.469568 | 0.075951 | 0.785333 | 0.940295 | Healthy Control | Intestine |
| Spry2 | -9.20E-01 | 7.722663 | 5.426928 | 0.029064 | 0.080118 | Healthy Control | Intestine |
| Spry4 | 1.46E+00 | 5.381147 | 5.793688 | 0.024564 | 0.070028 | Infection | Intestine |
| Spsb1 | 1.24E+00 | 5.936831 | 3.973695 | 0.05878 | 0.14284 | Infection | Intestine |
| Spsb2 | -1.73E+00 | 7.58788 | 11.04172 | 0.002884 | 0.01134 | Healthy Control | Intestine |
| Spta1 | -3.59E+00 | 1.484327 | 0.162871 | 0.692774 | 0.890181 | Healthy Control | Intestine |
| Sptan1 | -3.11E-01 | 10.34967 | 7.907152 | 0.01021 | 0.033494 | Healthy Control | Intestine |
| Sptb | 1.98E+00 | 1.979032 | 0.855704 | 0.362756 | 0.587498 | Infection | Intestine |
| Sptbn2 | -6.34E+00 | 3.690721 | 1.192113 | 0.288092 | 0.493225 | Healthy Control | Intestine |
| Sptlc2 | -1.48E+00 | 9.716428 | 71.96289 | 2.38E-08 | 4.67E-07 | Healthy Control | Intestine |
| Sqstm1 | -1.22E+00 | 9.350339 | 31.65867 | 1.22E-05 | 9.82E-05 | Healthy Control | Intestine |
| Sra1 | -2.60E+00 | 8.43882 | 45.99749 | 5.77E-07 | 7.05E-06 | Healthy Control | Intestine |
| Src | -6.73E-01 | 6.006624 | 0.935785 | 0.342967 | 0.564581 | Healthy Control | Intestine |
| Srd5a3 | 6.52E-01 | 7.430157 | 4.974568 | 0.036358 | 0.095907 | Infection | Intestine |
| Srebf1 | -2.02E-01 | 7.602789 | 0.256311 | 0.617702 | 0.837736 | Healthy Control | Intestine |
| Srebf2 | -1.37E+00 | 10.16649 | 81.89907 | 7.87E-09 | 1.83E-07 | Healthy Control | Intestine |
| Srf | -3.85E-01 | 6.973352 | 0.744552 | 0.397284 | 0.626197 | Healthy Control | Intestine |
| Srgap1 | -8.48E-01 | 6.69189 | 2.375754 | 0.136467 | 0.276581 | Healthy Control | Intestine |
| Srp72 | -7.33E-01 | 7.994633 | 5.051032 | 0.034947 | 0.092858 | Healthy Control | Intestine |
| Srpx2 | 2.11E+00 | 6.790027 | 43.66493 | 1.27E-06 | 1.37E-05 | Infection | Intestine |
| Srr | -1.58E+00 | 5.941364 | 3.081369 | 0.091984 | 0.203119 | Healthy Control | Intestine |
| Srrm1 | -2.58E-01 | 9.557828 | 3.511745 | 0.074407 | 0.172648 | Healthy Control | Intestine |
| Srrm4 | -4.38E+00 | 2.203897 | 0.398916 | 0.533973 | 0.765939 | Healthy Control | Intestine |
| Srsf1 | -2.30E+00 | 9.549473 | 116.5118 | 2.87E-10 | 1.17E-08 | Healthy Control | Intestine |
| Srsf2 | -6.21E-03 | 10.30582 | 0.002263 | 0.962489 | 0.986734 | Healthy Control | Intestine |
| Srsf4 | 5.16E-01 | 8.802647 | 10.65847 | 0.003579 | 0.013686 | Infection | Intestine |
| Sry | -3.57E-01 | 1.228299 | 0.00618 | 0.938438 | 0.969735 | Healthy Control | Intestine |
| Ss18l1 | 7.10E-01 | 5.539242 | 1.210385 | 0.282517 | 0.486508 | Infection | Intestine |
| Ssbp2 | -5.35E-01 | 5.851264 | 0.517211 | 0.478882 | 0.713187 | Healthy Control | Intestine |
| Ssr2 | -2.68E+00 | 8.281261 | 40.69847 | 1.39E-06 | 1.47E-05 | Healthy Control | Intestine |
| Ssrp1 | -9.06E-02 | 8.433571 | 0.182586 | 0.673356 | 0.878962 | Healthy Control | Intestine |
| Sst | 6.62E-01 | 5.107976 | 0.065279 | 0.801679 | 0.94651 | Infection | Intestine |
| Sstr1 | -7.33E+00 | 4.256185 | 3.516949 | 0.075611 | 0.174669 | Healthy Control | Intestine |
| Sstr2 | -6.08E+00 | 3.530644 | 1.441607 | 0.244125 | 0.434135 | Healthy Control | Intestine |
| Sstr3 | 2.09E+00 | 3.10329 | 3.623778 | 0.069437 | 0.163539 | Infection | Intestine |
| Sstr4 | -3.05E+00 | 1.737239 | 0.209728 | 0.650162 | 0.861262 | Healthy Control | Intestine |
| St6galnac5 | 5.92E-01 | 3.757249 | 0.213972 | 0.647912 | 0.859523 | Infection | Intestine |
| Stac | 2.99E+00 | 1.86208 | 2.015042 | 0.165426 | 0.320511 | Infection | Intestine |
| Stag2 | -3.85E-01 | 8.20222 | 1.670052 | 0.209757 | 0.385478 | Healthy Control | Intestine |
| Stam2 | -1.28E+00 | 8.065029 | 19.58573 | 0.000205 | 0.001166 | Healthy Control | Intestine |
| Star | 1.32E+00 | 3.372084 | 0.961573 | 0.336732 | 0.556274 | Infection | Intestine |
| Stat1 | -1.23E+00 | 7.735626 | 4.73387 | 0.040201 | 0.104489 | Healthy Control | Intestine |
| Stat2 | -2.12E-01 | 7.70348 | 0.286973 | 0.597554 | 0.822945 | Healthy Control | Intestine |
| Stat3 | -6.14E-01 | 8.795677 | 7.97867 | 0.009919 | 0.03263 | Healthy Control | Intestine |
| Stat4 | -5.94E+00 | 3.312298 | 0.974077 | 0.335706 | 0.555369 | Healthy Control | Intestine |
| Stat5a | -2.79E-01 | 5.967337 | 0.26567 | 0.610999 | 0.832968 | Healthy Control | Intestine |
| Stat5b | -8.65E-01 | 7.210955 | 3.480425 | 0.074976 | 0.173547 | Healthy Control | Intestine |
| Stat6 | -5.60E-02 | 8.038181 | 0.046048 | 0.832084 | 0.954066 | Healthy Control | Intestine |

| | | | | | | | |
|---|---|---|---|---|---|---|---|
| Stau2 | -8.21E-01 | 6.061207 | 1.620466 | 0.215208 | 0.393581 | Healthy Control | Intestine |
| Steap1 | 2.13E+00 | 5.814451 | 24.65838 | 5.29E-05 | 0.000352 | Infection | Intestine |
| Stim1 | -1.49E+00 | 8.668842 | 22.38388 | 9.76E-05 | 0.000607 | Healthy Control | Intestine |
| Stim2 | -9.83E-01 | 7.952372 | 5.92059 | 0.023473 | 0.067648 | Healthy Control | Intestine |
| Stimate | -1.37E+00 | 6.382766 | 3.47017 | 0.074648 | 0.172924 | Healthy Control | Intestine |
| Stip1 | -2.39E+00 | 7.621264 | 30.72721 | 1.02E-05 | 8.44E-05 | Healthy Control | Intestine |
| Stk11 | 1.30E+00 | 8.37547 | 52.33729 | 3.18E-07 | 4.21E-06 | Infection | Intestine |
| Stk3 | 8.83E-01 | 7.110748 | 6.940852 | 0.015215 | 0.0469 | Infection | Intestine |
| Stk32a | 8.90E-01 | 2.414953 | 0.271835 | 0.606918 | 0.830118 | Infection | Intestine |
| Stk38 | -7.40E-01 | 8.176186 | 8.141724 | 0.009225 | 0.030684 | Healthy Control | Intestine |
| Stk38l | -1.48E+00 | 8.210307 | 17.79524 | 0.000342 | 0.001825 | Healthy Control | Intestine |
| Stk39 | -2.71E+00 | 8.249347 | 25.41425 | 3.78E-05 | 0.000262 | Healthy Control | Intestine |
| Stk4 | 9.82E-01 | 7.863255 | 15.79218 | 0.000653 | 0.003202 | Infection | Intestine |
| Stmn2 | 4.24E-01 | 6.944918 | 0.302773 | 0.588417 | 0.814156 | Infection | Intestine |
| Stom | 1.14E+00 | 7.750284 | 25.56428 | 4.74E-05 | 0.00032 | Infection | Intestine |
| Stoml3 | -3.57E-01 | 1.228299 | 0.00618 | 0.938438 | 0.969735 | Healthy Control | Intestine |
| Stra6l | -2.09E+00 | 6.337597 | 3.638234 | 0.06841 | 0.16192 | Healthy Control | Intestine |
| Stra8 | -4.95E-01 | 1.228692 | 0.008683 | 0.927069 | 0.969735 | Healthy Control | Intestine |
| Strn3 | -1.10E+00 | 8.778014 | 23.76512 | 7.25E-05 | 0.000468 | Healthy Control | Intestine |
| Stub1 | 3.66E-01 | 8.550385 | 2.34823 | 0.13982 | 0.282121 | Infection | Intestine |
| Stx1a | 2.82E+00 | 4.363618 | 28.34811 | 2.44E-05 | 0.00018 | Infection | Intestine |
| Stx8 | 9.83E-01 | 7.164139 | 9.978498 | 0.004592 | 0.016977 | Infection | Intestine |
| Stxbp1 | -3.94E-01 | 7.460748 | 1.059986 | 0.313987 | 0.527558 | Healthy Control | Intestine |
| Stxbp4 | 1.80E+00 | 5.593895 | 8.629227 | 0.007661 | 0.026304 | Infection | Intestine |
| Stxbp5l | 2.04E+00 | 2.135805 | 0.955523 | 0.33692 | 0.556274 | Infection | Intestine |
| Sucnr1 | -1.40E+00 | 1.362959 | 0.050057 | 0.826006 | 0.953685 | Healthy Control | Intestine |
| Sufu | -8.45E-02 | 7.125769 | 0.048795 | 0.827186 | 0.953685 | Healthy Control | Intestine |
| Sulf1 | 9.24E-01 | 5.056007 | 0.964817 | 0.336142 | 0.555906 | Infection | Intestine |
| Sulf2 | -1.10E+00 | 7.490267 | 2.608033 | 0.120066 | 0.249741 | Healthy Control | Intestine |
| Sult2a8 | -1.37E+00 | 1.229023 | 0.032546 | 0.859435 | 0.956553 | Healthy Control | Intestine |
| Sumf1 | -7.27E+00 | 4.440693 | 3.827745 | 0.064677 | 0.154545 | Healthy Control | Intestine |
| Sumo1 | -1.58E+00 | 8.670238 | 21.49756 | 0.000125 | 0.000746 | Healthy Control | Intestine |
| Sumo3 | -2.39E+00 | 8.433957 | 49.52743 | 3.31E-07 | 4.36E-06 | Healthy Control | Intestine |
| Sun1 | -1.17E+00 | 8.500689 | 21.96577 | 0.000112 | 0.000687 | Healthy Control | Intestine |
| Supt16 | -7.82E-01 | 7.944359 | 7.812621 | 0.010547 | 0.03439 | Healthy Control | Intestine |
| Supt20 | -7.13E-01 | 7.916768 | 5.261414 | 0.031661 | 0.08572 | Healthy Control | Intestine |
| Supt3 | -9.83E-01 | 5.406525 | 0.922081 | 0.346532 | 0.569514 | Healthy Control | Intestine |
| Supt5 | -4.65E-01 | 8.733833 | 5.175409 | 0.033118 | 0.088752 | Healthy Control | Intestine |
| Supt6 | -1.22E+00 | 9.527291 | 47.67565 | 6.56E-07 | 7.80E-06 | Healthy Control | Intestine |
| Suv39h1 | -9.99E-01 | 5.490049 | 0.853001 | 0.36461 | 0.589166 | Healthy Control | Intestine |
| Suz12 | -1.10E+00 | 7.701747 | 10.91749 | 0.003164 | 0.012258 | Healthy Control | Intestine |
| Sv2a | 1.94E-01 | 3.440113 | 0.014946 | 0.903792 | 0.969735 | Infection | Intestine |
| Sv2c | -7.29E+00 | 4.114034 | 2.625362 | 0.121105 | 0.251453 | Healthy Control | Intestine |
| Svil | 2.68E+00 | 7.522317 | 216.0676 | 8.55E-13 | 9.96E-11 | Infection | Intestine |
| Syce2 | -7.24E+00 | 4.623992 | 3.918912 | 0.061831 | 0.149026 | Healthy Control | Intestine |
| Sycp1 | -1.81E+00 | 1.554798 | 0.063594 | 0.803453 | 0.947264 | Healthy Control | Intestine |
| Sycp3 | 3.30E+00 | 1.632339 | 2.429857 | 0.127681 | 0.261741 | Infection | Intestine |
| Syk | 2.10E+00 | 8.851113 | 142.2292 | 5.02E-11 | 2.82E-09 | Infection | Intestine |
| Syn1 | -5.78E+00 | 2.934892 | 1.096173 | 0.307722 | 0.519547 | Healthy Control | Intestine |
| Syn2 | -2.87E+00 | 6.658398 | 7.677562 | 0.011314 | 0.03652 | Healthy Control | Intestine |

| | | | | | | | |
|---|---|---|---|---|---|---|---|
| Syn3 | -7.90E+00 | 5.152544 | 5.493217 | 0.029645 | 0.081362 | Healthy Control | Intestine |
| Syncrip | -3.60E-01 | 8.610631 | 2.544244 | 0.125108 | 0.257733 | Healthy Control | Intestine |
| Syne1 | 2.48E+00 | 7.580062 | 87.31833 | 4.44E-09 | 1.14E-07 | Infection | Intestine |
| Syne4 | 6.12E-01 | 3.091694 | 0.122025 | 0.730121 | 0.91327 | Infection | Intestine |
| Syngr3 | -4.83E+00 | 2.386061 | 0.441933 | 0.513316 | 0.748081 | Healthy Control | Intestine |
| Synj2bp | 7.10E-01 | 7.223697 | 4.901297 | 0.0376 | 0.098755 | Infection | Intestine |
| Synpo | 4.43E-01 | 6.735377 | 1.005369 | 0.326661 | 0.54382 | Infection | Intestine |
| Synpo2 | 9.21E-01 | 6.652188 | 4.808074 | 0.039298 | 0.102461 | Infection | Intestine |
| Syp | 9.08E-02 | 4.35162 | 0.004607 | 0.946441 | 0.975476 | Infection | Intestine |
| Sypl2 | -3.23E+00 | 1.803179 | 0.194959 | 0.661845 | 0.869498 | Healthy Control | Intestine |
| Syt2 | -3.75E+00 | 2.296412 | 0.215335 | 0.647334 | 0.859335 | Healthy Control | Intestine |
| Syt4 | -4.38E+00 | 2.015835 | 0.375849 | 0.54571 | 0.776921 | Healthy Control | Intestine |
| Syt6 | -1.61E+00 | 1.445682 | 0.039299 | 0.844305 | 0.954066 | Healthy Control | Intestine |
| Syt9 | -3.21E+00 | 1.917033 | 0.138718 | 0.712548 | 0.900306 | Healthy Control | Intestine |
| Sytl5 | 1.12E+00 | 3.296325 | 0.646947 | 0.429479 | 0.661686 | Infection | Intestine |
| Szt2 | -1.40E-02 | 6.569227 | 0.001005 | 0.974988 | 0.995876 | Healthy Control | Intestine |
| T | -3.57E-01 | 1.227387 | 0.006293 | 0.937883 | 0.969735 | Healthy Control | Intestine |
| Taar5 | -9.71E-16 | 1.180616 | 0 | 1 | 1 | Healthy Control | Intestine |
| Tab1 | -7.29E-01 | 7.178341 | 2.92629 | 0.100712 | 0.218158 | Healthy Control | Intestine |
| Tac1 | -9.28E+00 | 6.104512 | 16.14059 | 0.000686 | 0.003334 | Healthy Control | Intestine |
| Tac2 | -6.42E-01 | 1.273248 | 0.013089 | 0.910112 | 0.969735 | Healthy Control | Intestine |
| Tacr1 | 1.16E+00 | 2.485998 | 0.363826 | 0.552412 | 0.781891 | Infection | Intestine |
| Tacr2 | -5.11E+00 | 2.624151 | 0.679849 | 0.419201 | 0.65131 | Healthy Control | Intestine |
| Tacr3 | -2.84E+00 | 1.486307 | 0.176635 | 0.677324 | 0.880469 | Healthy Control | Intestine |
| Tacstd2 | -2.36E+00 | 1.40432 | 0.100434 | 0.753702 | 0.926115 | Healthy Control | Intestine |
| Taf11 | -1.13E+00 | 6.130507 | 1.962588 | 0.174026 | 0.334168 | Healthy Control | Intestine |
| Taf12 | 2.39E-01 | 7.69762 | 0.7906 | 0.383593 | 0.612525 | Infection | Intestine |
| Taf15 | 2.55E-01 | 8.508406 | 1.367154 | 0.254943 | 0.44954 | Infection | Intestine |
| Taf4 | -6.37E-01 | 8.382466 | 6.172438 | 0.021149 | 0.061913 | Healthy Control | Intestine |
| Taf5l | -1.67E+00 | 7.586442 | 16.46304 | 0.000469 | 0.002417 | Healthy Control | Intestine |
| Taf6l | -6.09E+00 | 3.51531 | 1.203364 | 0.285887 | 0.490456 | Healthy Control | Intestine |
| Taf9 | -9.23E-01 | 7.049142 | 5.399524 | 0.029157 | 0.080287 | Healthy Control | Intestine |
| Tagln | 2.45E+00 | 9.668202 | 27.10094 | 0.000207 | 0.001171 | Infection | Intestine |
| Tal1 | -4.13E+00 | 2.186606 | 0.306098 | 0.585285 | 0.810829 | Healthy Control | Intestine |
| Tap1 | 4.48E-01 | 7.458493 | 0.548274 | 0.468431 | 0.70356 | Infection | Intestine |
| Tardbp | -4.81E-01 | 10.1583 | 9.74275 | 0.005007 | 0.018314 | Healthy Control | Intestine |
| Tas1r1 | -1.92E+00 | 1.591279 | 0.103917 | 0.750616 | 0.92508 | Healthy Control | Intestine |
| Tas1r2 | -9.71E-16 | 1.180616 | 0 | 1 | 1 | Healthy Control | Intestine |
| Tas1r3 | -7.14E-01 | 5.120499 | 0.394166 | 0.536049 | 0.76802 | Healthy Control | Intestine |
| Tbc1d1 | 6.29E-01 | 6.814582 | 2.629447 | 0.119234 | 0.24857 | Infection | Intestine |
| Tbc1d22a | 1.53E-01 | 7.826286 | 0.381852 | 0.543003 | 0.775099 | Infection | Intestine |
| Tbck | 7.17E-02 | 7.554293 | 0.054883 | 0.816951 | 0.952856 | Infection | Intestine |
| Tbk1 | 2.26E-01 | 7.617761 | 0.589319 | 0.450908 | 0.684875 | Infection | Intestine |
| Tbl1xr1 | -1.00E+00 | 8.358513 | 10.15932 | 0.004232 | 0.01582 | Healthy Control | Intestine |
| Tbp | 1.07E+00 | 6.615316 | 10.57921 | 0.003666 | 0.013954 | Infection | Intestine |
| Tbpl1 | 9.19E-02 | 7.340767 | 0.084002 | 0.774656 | 0.934669 | Infection | Intestine |
| Tbpl2 | -9.77E-01 | 1.317881 | 0.017898 | 0.894771 | 0.969735 | Healthy Control | Intestine |
| Tbr1 | -2.33E+00 | 1.402227 | 0.123691 | 0.727708 | 0.912095 | Healthy Control | Intestine |
| Tbx1 | -6.71E-01 | 4.042403 | 0.032583 | 0.859128 | 0.956553 | Healthy Control | Intestine |
| Tbx18 | -1.93E+00 | 1.363852 | 0.077177 | 0.784183 | 0.940206 | Healthy Control | Intestine |

| | | | | | | | |
|---|---|---|---|---|---|---|---|
| Tbx2 | -1.01E+00 | 7.001325 | 3.08448 | 0.092279 | 0.20359 | Healthy Control | Intestine |
| Tbx20 | -9.71E-16 | 1.180616 | 0 | 1 | 1 | Healthy Control | Intestine |
| Tbx21 | -5.74E+00 | 3.170742 | 0.672125 | 0.422177 | 0.65431 | Healthy Control | Intestine |
| Tbx3 | -9.45E-01 | 6.969741 | 1.960612 | 0.174702 | 0.335337 | Healthy Control | Intestine |
| Tbx4 | -4.36E+00 | 2.721971 | 0.449201 | 0.51012 | 0.745159 | Healthy Control | Intestine |
| Tbx5 | -3.57E-01 | 1.228299 | 0.00618 | 0.938438 | 0.969735 | Healthy Control | Intestine |
| Tbx6 | -5.47E+00 | 3.0722 | 1.173447 | 0.291812 | 0.498571 | Healthy Control | Intestine |
| Tbxa2r | -6.96E+00 | 4.603016 | 3.047224 | 0.096425 | 0.210437 | Healthy Control | Intestine |
| Tcaf1 | -3.37E-01 | 7.230591 | 0.516627 | 0.479728 | 0.713676 | Healthy Control | Intestine |
| Tcam1 | -7.56E-01 | 1.273429 | 0.015546 | 0.902199 | 0.969735 | Healthy Control | Intestine |
| Tceal5 | 6.01E-01 | 4.384143 | 0.231272 | 0.634831 | 0.850547 | Infection | Intestine |
| Tceal9 | -5.72E-01 | 8.409911 | 3.68201 | 0.068211 | 0.161679 | Healthy Control | Intestine |
| Tcf12 | -9.78E-01 | 8.318034 | 12.13317 | 0.002119 | 0.008653 | Healthy Control | Intestine |
| Tcf21 | -1.32E+00 | 7.117386 | 3.938196 | 0.058879 | 0.142943 | Healthy Control | Intestine |
| Tcf3 | -2.75E-01 | 7.38198 | 0.772469 | 0.388883 | 0.618408 | Healthy Control | Intestine |
| Tcf7 | 1.89E-01 | 6.049368 | 0.085182 | 0.772975 | 0.934635 | Infection | Intestine |
| Tcf7l1 | -1.14E+00 | 8.216178 | 11.43562 | 0.002519 | 0.01008 | Healthy Control | Intestine |
| Tcf7l2 | -2.56E+00 | 8.79456 | 40.0353 | 1.76E-06 | 1.79E-05 | Healthy Control | Intestine |
| Tcof1 | -1.47E+00 | 6.974935 | 6.642475 | 0.0165 | 0.050213 | Healthy Control | Intestine |
| Tcp1 | -1.52E+00 | 9.44739 | 57.94837 | 1.41E-07 | 2.10E-06 | Healthy Control | Intestine |
| Tctn1 | 5.41E-01 | 5.193316 | 0.481322 | 0.494535 | 0.727488 | Infection | Intestine |
| Tdg | -1.72E+00 | 7.617543 | 16.47894 | 0.000472 | 0.002429 | Healthy Control | Intestine |
| Tdgf1 | -1.49E+00 | 1.441366 | 0.029364 | 0.86521 | 0.960364 | Healthy Control | Intestine |
| Tdh | -1.08E+00 | 1.359678 | 0.034948 | 0.85327 | 0.956553 | Healthy Control | Intestine |
| Tdrd1 | -3.34E+00 | 1.740306 | 0.203932 | 0.656119 | 0.865261 | Healthy Control | Intestine |
| Tdrkh | 1.79E+00 | 3.326339 | 2.301909 | 0.142025 | 0.285762 | Infection | Intestine |
| Tead1 | 7.22E-01 | 7.981878 | 6.908918 | 0.015422 | 0.047481 | Infection | Intestine |
| Tead2 | -1.11E+00 | 5.421786 | 1.33621 | 0.259032 | 0.454827 | Healthy Control | Intestine |
| Tead3 | 2.64E-01 | 7.240516 | 0.750647 | 0.395621 | 0.624757 | Infection | Intestine |
| Tec | 2.22E+00 | 7.733272 | 98.40059 | 1.53E-09 | 4.79E-08 | Infection | Intestine |
| Tecpr2 | -7.77E-01 | 6.589975 | 1.493137 | 0.233717 | 0.419953 | Healthy Control | Intestine |
| Tecr | -9.95E+00 | 7.203226 | 29.64756 | 2.58E-05 | 0.000189 | Healthy Control | Intestine |
| Tectb | -1.55E+00 | 1.443522 | 0.047207 | 0.82949 | 0.954066 | Healthy Control | Intestine |
| Tek | 8.31E-01 | 6.055078 | 0.48405 | 0.497079 | 0.72973 | Infection | Intestine |
| Tenm1 | 5.35E+00 | 1.362897 | 5.273059 | 0.030593 | 0.083504 | Infection | Intestine |
| Tenm4 | -6.75E+00 | 3.940966 | 2.032143 | 0.169628 | 0.327102 | Healthy Control | Intestine |
| Tent5c | 6.23E-01 | 6.283221 | 1.512865 | 0.231091 | 0.417482 | Infection | Intestine |
| Tepsin | -2.35E+00 | 7.584715 | 16.60197 | 0.000431 | 0.002232 | Healthy Control | Intestine |
| Terf1 | -5.44E-01 | 6.725811 | 0.941668 | 0.341916 | 0.563222 | Healthy Control | Intestine |
| Terf2 | 4.16E-01 | 7.702249 | 2.767499 | 0.110527 | 0.234152 | Infection | Intestine |
| Tert | 2.57E-01 | 4.490747 | 0.0482 | 0.828093 | 0.954066 | Infection | Intestine |
| Tesmin | 2.82E+00 | 2.082529 | 1.772221 | 0.195148 | 0.364655 | Infection | Intestine |
| Tespa1 | -2.46E+00 | 1.598359 | 0.088284 | 0.768347 | 0.932538 | Healthy Control | Intestine |
| Tet1 | -7.57E+00 | 4.868925 | 6.207406 | 0.021724 | 0.063338 | Healthy Control | Intestine |
| Tet2 | -2.06E+00 | 6.933615 | 12.81283 | 0.001524 | 0.006576 | Healthy Control | Intestine |
| Tet3 | -1.34E+00 | 8.399189 | 22.93449 | 8.51E-05 | 0.000539 | Healthy Control | Intestine |
| Tex101 | -3.57E-01 | 1.228299 | 0.00618 | 0.938438 | 0.969735 | Healthy Control | Intestine |
| Tex19.1 | -1.99E+00 | 1.443254 | 0.074468 | 0.787015 | 0.941184 | Healthy Control | Intestine |
| Tfam | -3.85E-01 | 7.504071 | 1.188135 | 0.28748 | 0.492514 | Healthy Control | Intestine |
| Tfap2a | 2.39E+00 | 1.892575 | 1.484482 | 0.232558 | 0.418772 | Infection | Intestine |

| | | | | | | | |
|---|---|---|---|---|---|---|---|
| Tfap2b | -1.71E+00 | 1.320187 | 0.045546 | 0.833104 | 0.954066 | Healthy Control | Intestine |
| Tfap2c | -9.71E-16 | 1.180616 | 0 | 1 | 1 | Healthy Control | Intestine |
| Tfap2d | -3.57E-01 | 1.228165 | 0.006192 | 0.93838 | 0.969735 | Healthy Control | Intestine |
| Tfeb | 3.08E-01 | 7.299084 | 0.985161 | 0.33174 | 0.55008 | Infection | Intestine |
| Tfpi | 1.88E+00 | 6.494078 | 27.70508 | 2.88E-05 | 0.000207 | Infection | Intestine |
| Tfrc | -1.12E+00 | 9.086512 | 14.67501 | 0.000918 | 0.004312 | Healthy Control | Intestine |
| Tgfa | -2.65E+00 | 6.859841 | 15.68295 | 0.000588 | 0.002916 | Healthy Control | Intestine |
| Tgfb1 | 6.49E-01 | 7.193932 | 2.316199 | 0.142338 | 0.286278 | Infection | Intestine |
| Tgfb1i1 | 1.33E+00 | 7.553903 | 13.35377 | 0.001413 | 0.006156 | Infection | Intestine |
| Tgfb2 | 1.18E-01 | 5.73701 | 0.02484 | 0.876118 | 0.96549 | Infection | Intestine |
| Tgfb3 | 1.51E+00 | 5.358768 | 3.506143 | 0.074426 | 0.172648 | Infection | Intestine |
| Tgfbr1 | 2.13E+00 | 7.18867 | 78.63846 | 1.11E-08 | 2.44E-07 | Infection | Intestine |
| Tgfbr2 | 1.86E+00 | 9.304653 | 70.29811 | 9.10E-08 | 1.47E-06 | Infection | Intestine |
| Tgfbr3 | 1.40E+00 | 8.73848 | 13.30773 | 0.002218 | 0.009027 | Infection | Intestine |
| Tgif2 | 1.63E+00 | 3.205825 | 1.161006 | 0.29159 | 0.498502 | Infection | Intestine |
| Tgm2 | 3.44E+00 | 9.998484 | 439.4406 | 2.20E-14 | 4.24E-12 | Infection | Intestine |
| Tgs1 | -9.37E-01 | 4.514781 | 0.49829 | 0.487379 | 0.721551 | Healthy Control | Intestine |
| Th | -1.56E+00 | 1.482166 | 0.032531 | 0.858518 | 0.956553 | Healthy Control | Intestine |
| Thap12 | 1.60E+00 | 8.565915 | 122.1714 | 2.10E-10 | 9.01E-09 | Infection | Intestine |
| Thbd | -1.57E+00 | 7.345601 | 3.826405 | 0.062911 | 0.151145 | Healthy Control | Intestine |
| Thbs1 | -8.21E-01 | 9.156581 | 3.160455 | 0.09472 | 0.207603 | Healthy Control | Intestine |
| Thbs2 | 1.43E+00 | 5.838245 | 6.404016 | 0.019084 | 0.056967 | Infection | Intestine |
| Thbs4 | 1.82E+00 | 10.28565 | 6.521829 | 0.02554 | 0.072194 | Infection | Intestine |
| Themis | -5.70E+00 | 3.098938 | 0.843688 | 0.369511 | 0.595164 | Healthy Control | Intestine |
| Thpo | -4.61E+00 | 2.811986 | 0.347003 | 0.561114 | 0.789692 | Healthy Control | Intestine |
| Thra | -1.68E-01 | 8.713995 | 0.451351 | 0.508759 | 0.743388 | Healthy Control | Intestine |
| Thrap3 | -3.48E-01 | 8.913272 | 2.870471 | 0.104477 | 0.224085 | Healthy Control | Intestine |
| Thrb | -1.01E+00 | 7.432893 | 5.784295 | 0.024697 | 0.070356 | Healthy Control | Intestine |
| Thumpd1 | -6.79E-01 | 7.35281 | 2.561146 | 0.123424 | 0.255063 | Healthy Control | Intestine |
| Tiam1 | 1.80E+00 | 5.692946 | 12.40246 | 0.001937 | 0.008045 | Infection | Intestine |
| Ticam1 | -5.46E-01 | 4.983211 | 0.367107 | 0.550347 | 0.779801 | Healthy Control | Intestine |
| Ticam2 | 3.47E+00 | 3.673465 | 15.63671 | 0.000602 | 0.002978 | Infection | Intestine |
| Tie1 | -3.62E-01 | 5.488706 | 0.094753 | 0.760798 | 0.928591 | Healthy Control | Intestine |
| Tigar | 5.25E-01 | 4.961763 | 0.581425 | 0.453203 | 0.686662 | Infection | Intestine |
| Tigit | -5.34E-01 | 3.865022 | 0.073709 | 0.788521 | 0.941823 | Healthy Control | Intestine |
| Timeless | 1.79E+00 | 4.515401 | 5.839153 | 0.023983 | 0.068881 | Infection | Intestine |
| Timm50 | 4.34E-01 | 7.38999 | 2.048554 | 0.166551 | 0.322162 | Infection | Intestine |
| Timmdc1 | -3.55E-01 | 7.042234 | 0.762797 | 0.391641 | 0.621456 | Healthy Control | Intestine |
| Timp1 | 6.42E+00 | 5.491693 | 511.4291 | 1.31E-16 | 6.07E-14 | Infection | Intestine |
| Timp4 | 3.87E-02 | 4.110921 | 0.000499 | 0.982349 | 1 | Infection | Intestine |
| Tiparp | 1.23E+00 | 6.446907 | 9.263219 | 0.005989 | 0.021356 | Infection | Intestine |
| Tirap | -1.99E-01 | 6.964284 | 0.284896 | 0.598728 | 0.823556 | Healthy Control | Intestine |
| Tjp1 | -2.16E-01 | 8.772139 | 1.228351 | 0.27982 | 0.483195 | Healthy Control | Intestine |
| Tjp2 | -2.81E+00 | 7.656795 | 18.69641 | 0.00023 | 0.00129 | Healthy Control | Intestine |
| Tjp3 | -3.89E+00 | 10.01088 | 182.3744 | 1.68E-12 | 1.75E-10 | Healthy Control | Intestine |
| Tlcd1 | -1.95E+00 | 6.8568 | 8.08595 | 0.009016 | 0.03011 | Healthy Control | Intestine |
| Tle1 | -1.39E+00 | 8.491744 | 28.86437 | 2.14E-05 | 0.000162 | Healthy Control | Intestine |
| Tle4 | -1.61E+00 | 9.074881 | 27.52745 | 2.83E-05 | 0.000204 | Healthy Control | Intestine |
| Tle6 | 3.68E-01 | 4.945291 | 0.174463 | 0.679857 | 0.882617 | Infection | Intestine |
| Tll1 | -6.62E+00 | 3.68579 | 1.208432 | 0.284936 | 0.489327 | Healthy Control | Intestine |

| | | | | | | | |
|---|---|---|---|---|---|---|---|
| Tln1 | 1.65E+00 | 8.060208 | 68.16818 | 3.73E-08 | 6.89E-07 | Infection | Intestine |
| Tlr1 | -1.03E+01 | 7.69319 | 49.00502 | 9.07E-07 | 1.03E-05 | Healthy Control | Intestine |
| Tlr11 | -6.85E+00 | 4.180751 | 1.178908 | 0.290656 | 0.497156 | Healthy Control | Intestine |
| Tlr12 | -4.87E+00 | 2.507066 | 0.738712 | 0.400083 | 0.630014 | Healthy Control | Intestine |
| Tlr13 | -7.72E+00 | 5.128891 | 3.636556 | 0.071144 | 0.166268 | Healthy Control | Intestine |
| Tlr2 | -2.08E+00 | 7.587226 | 12.67435 | 0.001597 | 0.006822 | Healthy Control | Intestine |
| Tlr3 | -3.51E+00 | 6.823888 | 11.67897 | 0.002393 | 0.009652 | Healthy Control | Intestine |
| Tlr4 | 4.87E-02 | 5.99294 | 0.006782 | 0.935061 | 0.969735 | Infection | Intestine |
| Tlr5 | -2.25E+00 | 5.842342 | 4.108029 | 0.05437 | 0.134084 | Healthy Control | Intestine |
| Tlr6 | 7.59E-01 | 5.446662 | 1.132607 | 0.298023 | 0.506761 | Infection | Intestine |
| Tlr7 | -6.87E+00 | 4.464727 | 2.357316 | 0.14057 | 0.28329 | Healthy Control | Intestine |
| Tlr8 | -6.00E+00 | 3.960019 | 0.967319 | 0.3369 | 0.556274 | Healthy Control | Intestine |
| Tlr9 | -7.73E+00 | 5.102386 | 2.56031 | 0.125415 | 0.258154 | Healthy Control | Intestine |
| Tlx1 | -1.37E+00 | 1.228863 | 0.033199 | 0.858049 | 0.956553 | Healthy Control | Intestine |
| Tlx3 | -3.57E-01 | 1.227067 | 0.006351 | 0.937596 | 0.969735 | Healthy Control | Intestine |
| Tmco5 | -1.52E+00 | 1.274268 | 0.042695 | 0.838717 | 0.954066 | Healthy Control | Intestine |
| Tmem106b | -4.46E-01 | 8.061384 | 2.179635 | 0.154072 | 0.304135 | Healthy Control | Intestine |
| Tmem108 | -2.20E+00 | 1.318658 | 0.07665 | 0.784843 | 0.940206 | Healthy Control | Intestine |
| Tmem109 | 9.66E-01 | 8.560105 | 24.00481 | 6.89E-05 | 0.000447 | Infection | Intestine |
| Tmem119 | 1.53E+00 | 8.108749 | 22.57558 | 0.000124 | 0.000743 | Infection | Intestine |
| Tmem120a | -1.55E+00 | 7.629174 | 7.797353 | 0.01029 | 0.033689 | Healthy Control | Intestine |
| Tmem163 | -3.59E+00 | 2.003887 | 0.162813 | 0.689919 | 0.888369 | Healthy Control | Intestine |
| Tmem173 | 5.43E-01 | 6.370964 | 1.256359 | 0.273954 | 0.475526 | Infection | Intestine |
| Tmem259 | -6.61E-01 | 8.535161 | 9.743269 | 0.005008 | 0.018314 | Healthy Control | Intestine |
| Tmem35a | -7.70E-01 | 5.311454 | 0.325769 | 0.573233 | 0.80173 | Healthy Control | Intestine |
| Tmem41b | -1.28E+00 | 6.564489 | 3.446887 | 0.075646 | 0.174669 | Healthy Control | Intestine |
| Tmem43 | -2.19E-02 | 6.28067 | 0.001843 | 0.966127 | 0.988843 | Healthy Control | Intestine |
| Tmem63b | -1.10E+00 | 8.858752 | 14.56093 | 0.000954 | 0.004439 | Healthy Control | Intestine |
| Tmem94 | -2.43E+00 | 9.277649 | 105.7163 | 7.19E-10 | 2.47E-08 | Healthy Control | Intestine |
| Tmem98 | -2.65E+00 | 9.244209 | 99.49964 | 1.04E-09 | 3.46E-08 | Healthy Control | Intestine |
| Tmigd1 | -2.93E+00 | 7.07411 | 3.230707 | 0.091195 | 0.202001 | Healthy Control | Intestine |
| Tmprss15 | -1.52E+00 | 1.273479 | 0.043253 | 0.837684 | 0.954066 | Healthy Control | Intestine |
| Tmprss2 | -4.96E+00 | 10.20513 | 234.2252 | 2.60E-13 | 3.82E-11 | Healthy Control | Intestine |
| Tmprss4 | -3.80E+00 | 8.716363 | 58.24703 | 7.43E-08 | 1.24E-06 | Healthy Control | Intestine |
| Tnc | 1.08E+00 | 8.684161 | 4.158096 | 0.062103 | 0.14961 | Infection | Intestine |
| Tnf | 1.08E+00 | 3.620848 | 0.451835 | 0.508005 | 0.742721 | Infection | Intestine |
| Tnfaip3 | -1.38E+00 | 7.466292 | 9.103428 | 0.006043 | 0.021529 | Healthy Control | Intestine |
| Tnfrsf11a | -4.72E+00 | 10.46877 | 300.9834 | 7.04E-15 | 1.68E-12 | Healthy Control | Intestine |
| Tnfrsf11b | 2.52E+00 | 5.010454 | 18.36861 | 0.000306 | 0.00166 | Infection | Intestine |
| Tnfrsf13b | 2.05E-01 | 4.679161 | 0.029312 | 0.86547 | 0.960439 | Infection | Intestine |
| Tnfrsf13c | -4.25E+00 | 2.363036 | 0.233058 | 0.633997 | 0.849742 | Healthy Control | Intestine |
| Tnfrsf14 | -2.43E-02 | 8.045002 | 0.006291 | 0.937507 | 0.969735 | Healthy Control | Intestine |
| Tnfrsf18 | 1.00E-01 | 3.128674 | 0.003606 | 0.952679 | 0.980088 | Infection | Intestine |
| Tnfrsf1a | 8.18E-01 | 9.187017 | 23.9563 | 6.97E-05 | 0.000451 | Infection | Intestine |
| Tnfrsf1b | -1.44E+00 | 7.828716 | 14.41355 | 0.000944 | 0.004402 | Healthy Control | Intestine |
| Tnfrsf4 | -6.75E+00 | 4.146538 | 2.625339 | 0.12103 | 0.251401 | Healthy Control | Intestine |
| Tnfrsf9 | 3.13E+00 | 2.58247 | 5.963322 | 0.021867 | 0.063692 | Infection | Intestine |
| Tnfsf10 | -8.92E+00 | 6.195394 | 6.290769 | 0.020961 | 0.061506 | Healthy Control | Intestine |
| Tnfsf11 | 1.87E+00 | 2.232688 | 0.873856 | 0.358747 | 0.583081 | Infection | Intestine |
| Tnfsf12 | 8.03E-01 | 5.574438 | 1.312428 | 0.263991 | 0.461914 | Infection | Intestine |

| | | | | | | | |
|---|---|---|---|---|---|---|---|
| Tnfsf13 | -1.45E+00 | 8.205765 | 17.48423 | 0.000369 | 0.001953 | Healthy Control | Intestine |
| Tnfsf13b | 5.61E-01 | 6.994742 | 1.251565 | 0.275196 | 0.47669 | Infection | Intestine |
| Tnfsf14 | 2.08E+00 | 2.681078 | 1.289695 | 0.267142 | 0.465963 | Infection | Intestine |
| Tnfsf4 | -2.62E+00 | 1.557689 | 0.102351 | 0.751476 | 0.925207 | Healthy Control | Intestine |
| Tnfsf9 | 4.58E+00 | 1.600028 | 8.691186 | 0.00541 | 0.019569 | Infection | Intestine |
| Tnks2 | 7.04E-01 | 9.178649 | 23.05657 | 8.72E-05 | 0.00055 | Infection | Intestine |
| Tnnc1 | -2.71E+00 | 1.596251 | 0.094942 | 0.760086 | 0.928591 | Healthy Control | Intestine |
| Tnni3k | 5.06E+00 | 1.319545 | 4.022353 | 0.058831 | 0.142896 | Infection | Intestine |
| Tnnt2 | 1.13E+00 | 3.65721 | 0.724743 | 0.402647 | 0.633454 | Infection | Intestine |
| Tnp1 | -2.24E+00 | 1.361417 | 0.095901 | 0.759455 | 0.928591 | Healthy Control | Intestine |
| Tnpo1 | 1.26E-01 | 9.066056 | 0.547255 | 0.467338 | 0.702974 | Infection | Intestine |
| Tns2 | -5.98E-01 | 7.550196 | 1.445764 | 0.241919 | 0.431543 | Healthy Control | Intestine |
| Tnxb | 1.64E-01 | 8.675687 | 0.069392 | 0.796169 | 0.943769 | Infection | Intestine |
| Togaram1 | 8.10E-03 | 7.220937 | 0.000507 | 0.982237 | 1 | Infection | Intestine |
| Top2a | 4.26E-01 | 7.738395 | 1.203796 | 0.284558 | 0.489014 | Infection | Intestine |
| Top2b | 3.34E-01 | 9.296384 | 4.617567 | 0.043016 | 0.110787 | Infection | Intestine |
| Topbp1 | -1.65E+00 | 7.036318 | 7.537121 | 0.011238 | 0.036341 | Healthy Control | Intestine |
| Tor1a | -8.04E-02 | 8.663482 | 0.185989 | 0.670516 | 0.876855 | Healthy Control | Intestine |
| Tor2a | -2.70E-01 | 7.598622 | 0.904742 | 0.351811 | 0.575545 | Healthy Control | Intestine |
| Tox | -1.04E+01 | 7.396647 | 33.21362 | 1.27E-05 | 0.000101 | Healthy Control | Intestine |
| Tox3 | -1.01E+00 | 7.538479 | 6.973408 | 0.0148 | 0.045762 | Healthy Control | Intestine |
| Tpbpa | 4.77E+00 | 1.364579 | 3.580507 | 0.07056 | 0.165423 | Infection | Intestine |
| Tpcn1 | -1.40E+00 | 9.964886 | 58.90401 | 1.23E-07 | 1.89E-06 | Healthy Control | Intestine |
| Tph1 | -6.01E-01 | 6.613951 | 0.545194 | 0.468012 | 0.703509 | Healthy Control | Intestine |
| Tph2 | -3.57E-01 | 1.227509 | 0.006273 | 0.937979 | 0.969735 | Healthy Control | Intestine |
| Tpi1 | -1.34E+00 | 10.12011 | 68.50324 | 3.61E-08 | 6.70E-07 | Healthy Control | Intestine |
| Tppp | -2.20E+00 | 7.634102 | 24.32704 | 4.79E-05 | 0.000323 | Healthy Control | Intestine |
| Tpr | 7.75E-01 | 9.20378 | 28.26281 | 2.53E-05 | 0.000186 | Infection | Intestine |
| Tpsab1 | 5.33E+00 | 1.276175 | 4.716338 | 0.046709 | 0.118213 | Infection | Intestine |
| Tpsb2 | 5.72E+00 | 1.44991 | 13.61391 | 0.001196 | 0.005346 | Infection | Intestine |
| Traf2 | 6.23E-01 | 7.625682 | 5.949959 | 0.023331 | 0.067355 | Infection | Intestine |
| Traf3 | 3.45E-01 | 7.010798 | 0.874243 | 0.359937 | 0.584256 | Infection | Intestine |
| Traf3ip2 | -2.54E+00 | 7.89641 | 22.68121 | 7.92E-05 | 0.000507 | Healthy Control | Intestine |
| Traf3ip3 | 1.96E+00 | 4.801577 | 5.217976 | 0.03219 | 0.086743 | Infection | Intestine |
| Traf6 | -1.80E+00 | 7.11178 | 14.17866 | 0.000943 | 0.004402 | Healthy Control | Intestine |
| Trappc10 | -4.02E-01 | 9.605225 | 6.428498 | 0.018929 | 0.056571 | Healthy Control | Intestine |
| Trdn | 5.26E+00 | 1.364474 | 4.916323 | 0.036363 | 0.095907 | Infection | Intestine |
| Treh | 2.92E+00 | 1.997129 | 1.582885 | 0.220901 | 0.401973 | Infection | Intestine |
| Trem1 | -1.65E+00 | 1.318143 | 0.048108 | 0.828394 | 0.954066 | Healthy Control | Intestine |
| Trem2 | 2.06E+00 | 4.914032 | 7.387227 | 0.012228 | 0.039139 | Infection | Intestine |
| Trex1 | -8.38E+00 | 5.6691 | 10.58661 | 0.004014 | 0.015144 | Healthy Control | Intestine |
| Trh | -1.52E+00 | 1.275069 | 0.038023 | 0.847672 | 0.954066 | Healthy Control | Intestine |
| Trhr | -2.77E+00 | 1.598115 | 0.142989 | 0.707681 | 0.897011 | Healthy Control | Intestine |
| Trhr2 | -4.95E-01 | 1.228515 | 0.008877 | 0.926263 | 0.969735 | Healthy Control | Intestine |
| Trib2 | -5.94E+00 | 2.909747 | 1.396605 | 0.251178 | 0.444841 | Healthy Control | Intestine |
| Trib3 | 3.32E+00 | 3.468392 | 17.30483 | 0.000386 | 0.002031 | Infection | Intestine |
| Trim17 | -1.69E+00 | 1.520247 | 0.059908 | 0.809014 | 0.94887 | Healthy Control | Intestine |
| Trim2 | -3.09E+00 | 10.06458 | 159.7473 | 1.40E-11 | 9.57E-10 | Healthy Control | Intestine |
| Trim21 | -2.03E-01 | 6.339269 | 0.153035 | 0.699248 | 0.89357 | Healthy Control | Intestine |
| Trim24 | -1.33E+00 | 7.969238 | 13.69964 | 0.001188 | 0.005321 | Healthy Control | Intestine |

| | | | | | | | |
|---|---|---|---|---|---|---|---|
| Trim27 | 2.59E-01 | 8.367973 | 1.350077 | 0.257842 | 0.453216 | Infection | Intestine |
| Trim28 | -8.30E-01 | 9.601671 | 18.65189 | 0.000283 | 0.001542 | Healthy Control | Intestine |
| Trim29 | 2.16E+00 | 2.224861 | 1.515639 | 0.228525 | 0.413891 | Infection | Intestine |
| Trim32 | -1.97E-01 | 5.86449 | 0.070969 | 0.79222 | 0.943577 | Healthy Control | Intestine |
| Trim33 | -6.80E-01 | 7.476867 | 3.345486 | 0.080783 | 0.183986 | Healthy Control | Intestine |
| Trim36 | -1.32E+00 | 4.810809 | 0.828191 | 0.372321 | 0.598401 | Healthy Control | Intestine |
| Trim63 | 4.44E+00 | 1.445982 | 4.026263 | 0.053613 | 0.132538 | Infection | Intestine |
| Trim65 | -3.46E-01 | 6.206752 | 0.221246 | 0.642425 | 0.856225 | Healthy Control | Intestine |
| Trim7 | -8.05E+00 | 5.237137 | 6.382612 | 0.020162 | 0.059617 | Healthy Control | Intestine |
| Trip13 | -8.27E+00 | 5.524768 | 8.315975 | 0.009243 | 0.030725 | Healthy Control | Intestine |
| Trmt13 | 6.68E-01 | 6.006982 | 1.613631 | 0.216854 | 0.395905 | Infection | Intestine |
| Trmt2a | -9.69E-01 | 6.243112 | 1.870845 | 0.183918 | 0.348484 | Healthy Control | Intestine |
| Trp53 | 5.33E-01 | 9.180328 | 7.251161 | 0.013356 | 0.04197 | Infection | Intestine |
| Trp53bp1 | 6.32E-02 | 6.985864 | 0.020987 | 0.886125 | 0.96867 | Infection | Intestine |
| Trp53cor1 | 3.81E+00 | 1.633526 | 2.661996 | 0.112297 | 0.237397 | Infection | Intestine |
| Trp63 | -9.76E-01 | 1.317006 | 0.018532 | 0.892934 | 0.969735 | Healthy Control | Intestine |
| Trp73 | -2.02E+00 | 1.592596 | 0.07945 | 0.780235 | 0.938784 | Healthy Control | Intestine |
| Trpa1 | -6.76E+00 | 3.798506 | 1.783165 | 0.197034 | 0.367356 | Healthy Control | Intestine |
| Trpc1 | 2.04E+00 | 3.75596 | 4.800269 | 0.038544 | 0.100757 | Infection | Intestine |
| Trpc3 | -5.81E+00 | 3.052258 | 1.07277 | 0.312922 | 0.526374 | Healthy Control | Intestine |
| Trpc4 | -4.07E+00 | 2.084666 | 0.42001 | 0.523121 | 0.7574 | Healthy Control | Intestine |
| Trpc5 | -1.52E+00 | 1.275042 | 0.038572 | 0.846592 | 0.954066 | Healthy Control | Intestine |
| Trpc6 | 2.99E+00 | 3.682531 | 8.577943 | 0.007402 | 0.025553 | Infection | Intestine |
| Trpc7 | -2.15E+00 | 1.317599 | 0.073256 | 0.78953 | 0.941823 | Healthy Control | Intestine |
| Trpm1 | -1.56E+00 | 1.478692 | 0.034469 | 0.854413 | 0.956553 | Healthy Control | Intestine |
| Trpm2 | 1.28E+00 | 3.343546 | 0.595962 | 0.447562 | 0.681623 | Infection | Intestine |
| Trpm5 | -7.26E+00 | 4.039983 | 2.189298 | 0.15477 | 0.30512 | Healthy Control | Intestine |
| Trpm6 | -3.96E+00 | 7.358584 | 13.18685 | 0.001412 | 0.006155 | Healthy Control | Intestine |
| Trpm7 | -5.83E-01 | 8.369475 | 5.534916 | 0.028067 | 0.078057 | Healthy Control | Intestine |
| Trpm8 | -9.71E-16 | 1.180616 | 0 | 1 | 1 | Healthy Control | Intestine |
| Trpv1 | -3.00E+00 | 1.885102 | 0.156001 | 0.696174 | 0.892492 | Healthy Control | Intestine |
| Trpv2 | 9.03E-01 | 5.595275 | 2.120268 | 0.158982 | 0.311371 | Infection | Intestine |
| Trpv3 | -9.19E+00 | 6.437318 | 4.66565 | 0.046038 | 0.117167 | Healthy Control | Intestine |
| Trpv4 | 2.05E+00 | 3.749659 | 4.149885 | 0.052865 | 0.131011 | Infection | Intestine |
| Trpv5 | -1.89E+00 | 1.402554 | 0.075976 | 0.784909 | 0.940206 | Healthy Control | Intestine |
| Trpv6 | -5.59E+00 | 3.542667 | 0.212027 | 0.653318 | 0.863896 | Healthy Control | Intestine |
| Trrap | -6.49E-01 | 8.454587 | 3.352477 | 0.080782 | 0.183986 | Healthy Control | Intestine |
| Tsc1 | -3.64E-01 | 7.201862 | 1.080133 | 0.309855 | 0.522795 | Healthy Control | Intestine |
| Tsc2 | -1.01E+00 | 8.481716 | 11.85136 | 0.002329 | 0.009417 | Healthy Control | Intestine |
| Tsc22d3 | -1.52E+00 | 7.986707 | 8.505836 | 0.007726 | 0.026474 | Healthy Control | Intestine |
| Tsg101 | -6.63E+00 | 3.926213 | 2.914493 | 0.103494 | 0.222643 | Healthy Control | Intestine |
| Tshb | 7.13E+00 | 1.321845 | 14.35483 | 0.001862 | 0.007782 | Infection | Intestine |
| Tshr | -3.11E+00 | 1.889697 | 0.177701 | 0.676956 | 0.880448 | Healthy Control | Intestine |
| Tshz1 | -3.37E+00 | 8.275761 | 71.55401 | 1.32E-08 | 2.82E-07 | Healthy Control | Intestine |
| Tslp | -5.91E+00 | 3.486232 | 0.917155 | 0.349853 | 0.573653 | Healthy Control | Intestine |
| Tspyl2 | 1.09E+00 | 6.227136 | 7.505823 | 0.01193 | 0.038331 | Infection | Intestine |
| Ttbk1 | -4.53E+00 | 2.226369 | 0.477476 | 0.496854 | 0.72966 | Healthy Control | Intestine |
| Ttc8 | 1.02E+00 | 5.004997 | 1.682397 | 0.207273 | 0.382086 | Infection | Intestine |
| Ttk | 1.49E+00 | 5.719279 | 5.750575 | 0.025445 | 0.072046 | Infection | Intestine |
| Ttll3 | 2.42E+00 | 5.003143 | 21.75869 | 0.00012 | 0.000728 | Infection | Intestine |

| | | | | | | | |
|---|---|---|---|---|---|---|---|
| Ttll6 | -1.03E+01 | 7.657096 | 39.27908 | 4.23E-06 | 3.86E-05 | Healthy Control | Intestine |
| Ttn | 3.61E+00 | 1.485549 | 2.708317 | 0.109532 | 0.232829 | Infection | Intestine |
| Tuba3a | -1.41E+00 | 1.401951 | 0.03921 | 0.844453 | 0.954066 | Healthy Control | Intestine |
| Tubb1 | -1.65E+00 | 1.31734 | 0.05249 | 0.820884 | 0.953035 | Healthy Control | Intestine |
| Tubb2b | 2.14E+00 | 4.540449 | 5.742514 | 0.025183 | 0.071469 | Infection | Intestine |
| Tubb3 | 2.29E+00 | 2.738684 | 1.531972 | 0.22827 | 0.41358 | Infection | Intestine |
| Tubg1 | 2.93E-01 | 6.081135 | 0.285137 | 0.598529 | 0.823556 | Infection | Intestine |
| Tubg2 | 4.92E-02 | 3.602267 | 0.000831 | 0.977262 | 0.997385 | Infection | Intestine |
| Tubgcp2 | 4.62E-01 | 6.518728 | 1.385071 | 0.251751 | 0.445166 | Infection | Intestine |
| Tufm | -3.87E+00 | 8.244386 | 42.4788 | 8.69E-07 | 9.97E-06 | Healthy Control | Intestine |
| Twf1 | -2.20E+00 | 8.852617 | 58.60061 | 1.06E-07 | 1.67E-06 | Healthy Control | Intestine |
| Twf2 | -3.34E-01 | 7.958932 | 0.975687 | 0.334041 | 0.553346 | Healthy Control | Intestine |
| Twist1 | 1.90E+00 | 3.192806 | 2.023463 | 0.166837 | 0.322466 | Infection | Intestine |
| Twist2 | 1.08E+00 | 2.587294 | 0.34961 | 0.560282 | 0.789187 | Infection | Intestine |
| Twnk | -1.27E+00 | 5.755635 | 1.984512 | 0.171856 | 0.330761 | Healthy Control | Intestine |
| Txnip | -1.31E+00 | 11.70557 | 39.55528 | 4.73E-06 | 4.29E-05 | Healthy Control | Intestine |
| Tyk2 | -3.57E-01 | 7.137148 | 0.714902 | 0.406791 | 0.637769 | Healthy Control | Intestine |
| Tyr | -1.65E+00 | 1.318742 | 0.046529 | 0.831187 | 0.954066 | Healthy Control | Intestine |
| Tyro3 | -3.28E+00 | 6.609295 | 11.53466 | 0.002525 | 0.010096 | Healthy Control | Intestine |
| Tyrp1 | 5.70E+00 | 1.229995 | 6.086704 | 0.031631 | 0.085684 | Infection | Intestine |
| Tyw5 | 2.05E+00 | 5.245449 | 12.34477 | 0.001967 | 0.008145 | Infection | Intestine |
| Uba1y | -3.57E-01 | 1.227387 | 0.006293 | 0.937883 | 0.969735 | Healthy Control | Intestine |
| Uba6 | -1.55E-03 | 7.102613 | 1.81E-05 | 0.996642 | 1 | Healthy Control | Intestine |
| Ubac1 | -1.49E+00 | 8.088683 | 11.66662 | 0.002407 | 0.009695 | Healthy Control | Intestine |
| Ubap1 | -2.65E-01 | 7.880513 | 0.889669 | 0.355835 | 0.580457 | Healthy Control | Intestine |
| Ubash3b | 2.64E+00 | 4.603158 | 22.28975 | 0.000104 | 0.00064 | Infection | Intestine |
| Ube2b | 8.24E-01 | 8.961332 | 28.18303 | 2.58E-05 | 0.000189 | Infection | Intestine |
| Ube2d1 | 3.91E-01 | 6.719442 | 0.760574 | 0.39241 | 0.622041 | Infection | Intestine |
| Ube2g1 | -3.10E+00 | 7.904768 | 31.55379 | 8.64E-06 | 7.31E-05 | Healthy Control | Intestine |
| Ube2g2 | -6.79E-01 | 7.396468 | 3.641388 | 0.069131 | 0.163319 | Healthy Control | Intestine |
| Ube2i | 4.73E-01 | 8.553943 | 4.397413 | 0.047831 | 0.120504 | Infection | Intestine |
| Ube2m | -1.06E+00 | 8.727992 | 17.56469 | 0.000384 | 0.002022 | Healthy Control | Intestine |
| Ube2n | -9.02E+00 | 6.15256 | 14.36279 | 0.001166 | 0.005238 | Healthy Control | Intestine |
| Ube2r2 | -4.60E-01 | 8.689791 | 3.085385 | 0.092982 | 0.204871 | Healthy Control | Intestine |
| Ube2u | -6.42E-01 | 1.272459 | 0.013821 | 0.907644 | 0.969735 | Healthy Control | Intestine |
| Ube3a | -1.95E-01 | 8.932998 | 1.476978 | 0.237262 | 0.424646 | Healthy Control | Intestine |
| Ublcp1 | -4.10E-01 | 6.326 | 0.53279 | 0.472664 | 0.707583 | Healthy Control | Intestine |
| Ubr5 | -2.97E-01 | 9.209308 | 3.475583 | 0.075819 | 0.174827 | Healthy Control | Intestine |
| Ubr7 | 7.06E-01 | 7.476881 | 6.553836 | 0.017927 | 0.054126 | Infection | Intestine |
| Ubtf | 5.85E-01 | 9.105926 | 14.29601 | 0.00104 | 0.004793 | Infection | Intestine |
| Ubxn7 | 6.75E-02 | 7.836718 | 0.062586 | 0.804796 | 0.947891 | Infection | Intestine |
| Uchl1 | -1.23E+00 | 4.014242 | 0.138497 | 0.71529 | 0.901639 | Healthy Control | Intestine |
| Ucn3 | -1.07E+00 | 1.318803 | 0.019312 | 0.890895 | 0.969735 | Healthy Control | Intestine |
| Ucp1 | -1.37E+00 | 1.228088 | 0.037791 | 0.848672 | 0.954066 | Healthy Control | Intestine |
| Ucp3 | -2.55E+00 | 1.596656 | 0.100783 | 0.753178 | 0.925708 | Healthy Control | Intestine |
| Ufd1 | 4.62E-01 | 7.993671 | 5.251297 | 0.031982 | 0.086354 | Infection | Intestine |
| Ufl1 | -7.05E-01 | 7.562629 | 3.450494 | 0.076417 | 0.175801 | Healthy Control | Intestine |
| Ugt8a | 7.30E-01 | 5.098373 | 0.40837 | 0.529111 | 0.763504 | Infection | Intestine |
| Uhrf1 | -1.09E+00 | 6.482344 | 1.991019 | 0.171002 | 0.329624 | Healthy Control | Intestine |
| Uhrf2 | 1.05E-01 | 8.004099 | 0.173826 | 0.680806 | 0.882694 | Infection | Intestine |

| | | | | | | | |
|---|---|---|---|---|---|---|---|
| Ulk1 | -6.39E-01 | 7.827767 | 3.794926 | 0.064226 | 0.15376 | Healthy Control | Intestine |
| Unc13b | -2.66E+00 | 7.154544 | 13.19306 | 0.001364 | 0.005988 | Healthy Control | Intestine |
| Unc5c | 1.49E+00 | 3.112269 | 1.633573 | 0.213709 | 0.391305 | Infection | Intestine |
| Unc93b1 | -3.37E+00 | 8.326609 | 24.55551 | 4.53E-05 | 0.000308 | Healthy Control | Intestine |
| Uncx | 4.66E+00 | 1.276901 | 3.484567 | 0.083149 | 0.187837 | Infection | Intestine |
| Ung | -9.32E+00 | 6.428347 | 13.56653 | 0.001493 | 0.006447 | Healthy Control | Intestine |
| Upk1b | -5.87E+00 | 3.399737 | 1.405994 | 0.249873 | 0.443139 | Healthy Control | Intestine |
| Upk3b | 2.35E+00 | 2.294671 | 1.432577 | 0.24201 | 0.431543 | Infection | Intestine |
| Uri1 | -2.12E+00 | 8.427064 | 43.25757 | 1.04E-06 | 1.15E-05 | Healthy Control | Intestine |
| Urm1 | -1.37E+00 | 4.879472 | 1.131233 | 0.298667 | 0.507645 | Healthy Control | Intestine |
| Usp14 | -3.42E-01 | 7.673287 | 0.943774 | 0.341898 | 0.563222 | Healthy Control | Intestine |
| Usp18 | -6.40E+00 | 3.512801 | 2.242072 | 0.150224 | 0.298896 | Healthy Control | Intestine |
| Usp2 | -8.20E-01 | 5.921669 | 0.787897 | 0.383403 | 0.612417 | Healthy Control | Intestine |
| Usp22 | -1.67E+00 | 8.177326 | 27.0149 | 2.96E-05 | 0.000213 | Healthy Control | Intestine |
| Usp25 | -6.76E-01 | 8.15212 | 7.414249 | 0.012441 | 0.03972 | Healthy Control | Intestine |
| Usp3 | -1.10E+00 | 7.954041 | 5.78447 | 0.024874 | 0.07075 | Healthy Control | Intestine |
| Usp32 | 7.92E-01 | 7.555838 | 7.507336 | 0.012022 | 0.038579 | Infection | Intestine |
| Usp47 | -7.49E-01 | 8.860992 | 12.51117 | 0.00187 | 0.007799 | Healthy Control | Intestine |
| Usp50 | -2.04E+00 | 1.443505 | 0.068427 | 0.795399 | 0.943769 | Healthy Control | Intestine |
| Usp54 | -1.97E+00 | 5.208051 | 2.686064 | 0.115264 | 0.242645 | Healthy Control | Intestine |
| Usp7 | -2.81E-01 | 8.983782 | 1.704643 | 0.20531 | 0.37949 | Healthy Control | Intestine |
| Utp25 | -2.76E-01 | 6.729024 | 0.420882 | 0.523015 | 0.7574 | Healthy Control | Intestine |
| Utrn | -1.97E+00 | 9.557294 | 43.96332 | 1.17E-06 | 1.28E-05 | Healthy Control | Intestine |
| Uts2 | -1.17E+00 | 1.361294 | 0.024754 | 0.876193 | 0.96549 | Healthy Control | Intestine |
| Uts2b | -4.30E+00 | 2.242195 | 0.531128 | 0.472994 | 0.707865 | Healthy Control | Intestine |
| Vamp2 | 9.07E-01 | 8.153057 | 17.51473 | 0.000391 | 0.002053 | Infection | Intestine |
| Vamp3 | 3.71E-02 | 9.154778 | 0.042212 | 0.839124 | 0.954066 | Infection | Intestine |
| Vangl1 | 1.00E+00 | 6.549987 | 4.516568 | 0.045134 | 0.115376 | Infection | Intestine |
| Vangl2 | -9.44E-01 | 4.450591 | 0.517955 | 0.479017 | 0.713187 | Healthy Control | Intestine |
| Vasp | -6.43E-01 | 7.127571 | 2.010801 | 0.169406 | 0.326801 | Healthy Control | Intestine |
| Vat1 | 6.34E-01 | 8.569005 | 5.863265 | 0.024242 | 0.069386 | Infection | Intestine |
| Vav1 | -1.30E+00 | 6.780678 | 4.672256 | 0.041096 | 0.106375 | Healthy Control | Intestine |
| Vav2 | -7.81E-01 | 4.234893 | 0.250689 | 0.621413 | 0.839021 | Healthy Control | Intestine |
| Vax1 | -8.79E-01 | 1.31714 | 0.022114 | 0.883093 | 0.96867 | Healthy Control | Intestine |
| Vax2 | -9.71E-16 | 1.180616 | 0 | 1 | 1 | Healthy Control | Intestine |
| Vcam1 | -5.93E-01 | 6.038714 | 0.566787 | 0.458961 | 0.693289 | Healthy Control | Intestine |
| Vcan | 1.93E+00 | 7.249158 | 21.24668 | 0.000176 | 0.00101 | Infection | Intestine |
| Vcl | -1.28E-01 | 9.267222 | 0.446329 | 0.511099 | 0.746154 | Healthy Control | Intestine |
| Vcp | -2.10E+00 | 7.922579 | 28.17553 | 2.05E-05 | 0.000156 | Healthy Control | Intestine |
| Vdac1 | -2.76E+00 | 10.44759 | 235.4039 | 3.32E-13 | 4.61E-11 | Healthy Control | Intestine |
| Vdac2 | -2.81E+00 | 10.16509 | 150.0812 | 2.61E-11 | 1.68E-09 | Healthy Control | Intestine |
| Vdr | -4.93E+00 | 9.895721 | 163.3915 | 2.37E-12 | 2.33E-10 | Healthy Control | Intestine |
| Vegfa | 2.19E+00 | 7.281887 | 58.60184 | 1.30E-07 | 1.96E-06 | Infection | Intestine |
| Vegfb | -1.26E+00 | 7.894522 | 9.085261 | 0.006285 | 0.022329 | Healthy Control | Intestine |
| Vegfc | -1.29E+00 | 5.66375 | 2.361982 | 0.13744 | 0.278102 | Healthy Control | Intestine |
| Vegfd | 2.83E+00 | 4.120698 | 13.64946 | 0.001263 | 0.005621 | Infection | Intestine |
| Vgll2 | -9.76E-01 | 1.317411 | 0.023087 | 0.880593 | 0.96842 | Healthy Control | Intestine |
| Vhl | -7.36E-01 | 5.69172 | 0.76187 | 0.391415 | 0.621456 | Healthy Control | Intestine |
| Vil1 | -4.81E+00 | 11.36177 | 357.8768 | 2.69E-15 | 7.07E-13 | Healthy Control | Intestine |
| Vip | -9.06E+00 | 5.189029 | 0.368338 | 0.558029 | 0.786235 | Healthy Control | Intestine |

| | | | | | | | |
|---|---|---|---|---|---|---|---|
| Vipas39 | -3.36E+00 | 6.717632 | 15.94131 | 0.000586 | 0.002911 | Healthy Control | Intestine |
| Vipr1 | -2.85E+00 | 9.067257 | 58.85617 | 7.92E-08 | 1.31E-06 | Healthy Control | Intestine |
| Vipr2 | 9.40E-01 | 3.946469 | 0.585579 | 0.451573 | 0.685325 | Infection | Intestine |
| Virma | -1.22E+00 | 7.740703 | 12.05532 | 0.00211 | 0.008629 | Healthy Control | Intestine |
| Vit | -6.39E+00 | 3.909098 | 1.446732 | 0.243306 | 0.432948 | Healthy Control | Intestine |
| Vldlr | -4.43E-01 | 6.632049 | 0.753478 | 0.394174 | 0.623258 | Healthy Control | Intestine |
| Vps13a | -9.64E-01 | 7.81358 | 3.96849 | 0.058586 | 0.142509 | Healthy Control | Intestine |
| Vps25 | -5.48E-01 | 7.636018 | 1.755625 | 0.198717 | 0.369805 | Healthy Control | Intestine |
| Vps26b | -8.60E-01 | 7.379985 | 5.115934 | 0.033578 | 0.089792 | Healthy Control | Intestine |
| Vps28 | -2.60E+00 | 9.524907 | 108.9796 | 5.26E-10 | 1.94E-08 | Healthy Control | Intestine |
| Vps51 | 4.86E-01 | 7.797045 | 4.762996 | 0.04016 | 0.104436 | Infection | Intestine |
| Vps54 | -7.48E-01 | 7.876247 | 6.456395 | 0.018532 | 0.055551 | Healthy Control | Intestine |
| Vps72 | -1.17E+00 | 8.142133 | 11.64007 | 0.002473 | 0.009919 | Healthy Control | Intestine |
| Vps9d1 | -2.41E-01 | 6.599594 | 0.257207 | 0.616883 | 0.837115 | Healthy Control | Intestine |
| Vsig1 | -7.99E+00 | 5.563729 | 4.392792 | 0.050661 | 0.126418 | Healthy Control | Intestine |
| Vsir | -5.83E-01 | 7.154802 | 0.89945 | 0.352994 | 0.577103 | Healthy Control | Intestine |
| Vstm2a | -3.16E+00 | 1.669315 | 0.168792 | 0.683881 | 0.88417 | Healthy Control | Intestine |
| Vstm4 | 1.41E+00 | 7.447971 | 16.09421 | 0.000595 | 0.002949 | Infection | Intestine |
| Vstm5 | -8.31E+00 | 5.937583 | 7.975499 | 0.010552 | 0.03439 | Healthy Control | Intestine |
| Vsx1 | -4.95E-01 | 1.228692 | 0.008683 | 0.927069 | 0.969735 | Healthy Control | Intestine |
| Vsx2 | -6.42E-01 | 1.273568 | 0.012758 | 0.911248 | 0.969735 | Healthy Control | Intestine |
| Vtcn1 | -6.62E+00 | 4.663537 | 1.910899 | 0.180278 | 0.343015 | Healthy Control | Intestine |
| Vtn | 8.94E-01 | 2.897769 | 0.241931 | 0.627514 | 0.843311 | Infection | Intestine |
| Vwa8 | -1.72E+00 | 9.169368 | 54.69952 | 2.20E-07 | 3.06E-06 | Healthy Control | Intestine |
| Vwc2l | 5.33E+00 | 1.275264 | 4.750983 | 0.046013 | 0.117162 | Infection | Intestine |
| Vwf | 4.24E-01 | 6.083087 | 0.073656 | 0.789997 | 0.942036 | Infection | Intestine |
| Vxn | 5.99E+00 | 1.276046 | 5.400686 | 0.034639 | 0.092169 | Infection | Intestine |
| Wap | 4.59E+00 | 1.40501 | 4.006319 | 0.054913 | 0.135199 | Infection | Intestine |
| Wapl | -1.61E+00 | 9.466596 | 45.10813 | 9.75E-07 | 1.10E-05 | Healthy Control | Intestine |
| Was | 2.56E+00 | 3.166389 | 5.778994 | 0.024316 | 0.069558 | Infection | Intestine |
| Washc1 | -9.60E-01 | 7.501605 | 4.764453 | 0.039646 | 0.103262 | Healthy Control | Intestine |
| Washc2 | 2.01E+00 | 8.332905 | 240.7243 | 2.91E-13 | 4.16E-11 | Infection | Intestine |
| Washc3 | -7.12E-01 | 6.961675 | 2.74103 | 0.1114 | 0.235901 | Healthy Control | Intestine |
| Washc4 | 6.01E-02 | 7.143766 | 0.034634 | 0.854059 | 0.956553 | Infection | Intestine |
| Washc5 | 9.13E-01 | 7.146748 | 8.967637 | 0.006725 | 0.023576 | Infection | Intestine |
| Wasl | -1.34E+00 | 8.524772 | 19.64131 | 0.000207 | 0.001171 | Healthy Control | Intestine |
| Wdr34 | -8.00E+00 | 5.400678 | 8.52941 | 0.008518 | 0.028713 | Healthy Control | Intestine |
| Wdr36 | -4.59E-01 | 7.439954 | 1.281047 | 0.269817 | 0.469369 | Healthy Control | Intestine |
| Wdr37 | -2.56E-01 | 6.189688 | 0.227397 | 0.637837 | 0.853063 | Healthy Control | Intestine |
| Wdr4 | -6.54E-01 | 5.471042 | 0.553846 | 0.46385 | 0.699619 | Healthy Control | Intestine |
| Wdr45 | -9.02E-01 | 7.806417 | 7.110405 | 0.014016 | 0.043689 | Healthy Control | Intestine |
| Wdr5 | 9.08E-02 | 7.09953 | 0.069727 | 0.794182 | 0.943769 | Infection | Intestine |
| Wdr59 | -1.61E+00 | 6.978304 | 7.163908 | 0.013168 | 0.041497 | Healthy Control | Intestine |
| Wdr61 | -2.24E+00 | 7.622403 | 16.5607 | 0.00044 | 0.002275 | Healthy Control | Intestine |
| Wdr62 | -2.15E+00 | 6.338823 | 6.652056 | 0.016452 | 0.050151 | Healthy Control | Intestine |
| Wdr82 | 3.51E-01 | 7.685317 | 1.870851 | 0.185299 | 0.350438 | Infection | Intestine |
| Wdr83 | 1.04E-01 | 6.322757 | 0.04075 | 0.841795 | 0.954066 | Infection | Intestine |
| Wdtc1 | 1.84E-01 | 8.861156 | 0.679455 | 0.418713 | 0.650873 | Infection | Intestine |
| Wee1 | 9.51E-01 | 6.499634 | 4.359739 | 0.048644 | 0.122368 | Infection | Intestine |
| Wfdc18 | -4.28E+00 | 7.732139 | 12.72448 | 0.002907 | 0.011406 | Healthy Control | Intestine |

| | | | | | | | |
|---|---|---|---|---|---|---|---|
| Whrn | -5.89E-02 | 6.717762 | 0.012098 | 0.9134 | 0.969735 | Healthy Control | Intestine |
| Wif1 | -4.96E+00 | 2.711983 | 0.579399 | 0.455323 | 0.689399 | Healthy Control | Intestine |
| Wipf1 | 1.15E+00 | 5.930859 | 6.090221 | 0.021871 | 0.063692 | Infection | Intestine |
| Wiz | -3.90E-01 | 8.102411 | 1.951502 | 0.176452 | 0.337532 | Healthy Control | Intestine |
| Wls | 2.83E-01 | 8.910115 | 2.129082 | 0.158797 | 0.311253 | Infection | Intestine |
| Wnt1 | -3.57E-01 | 1.228165 | 0.006192 | 0.93838 | 0.969735 | Healthy Control | Intestine |
| Wnt10a | -3.58E+00 | 1.765276 | 0.321858 | 0.574697 | 0.802268 | Healthy Control | Intestine |
| Wnt10b | -3.79E+00 | 2.062698 | 0.239775 | 0.628056 | 0.843445 | Healthy Control | Intestine |
| Wnt11 | 3.87E+00 | 2.916232 | 22.79835 | 7.26E-05 | 0.000468 | Infection | Intestine |
| Wnt16 | -4.47E+00 | 2.538943 | 0.52054 | 0.478404 | 0.713187 | Healthy Control | Intestine |
| Wnt2 | -4.62E+00 | 2.793777 | 0.719284 | 0.405136 | 0.63637 | Healthy Control | Intestine |
| Wnt2b | 1.45E+00 | 7.55288 | 7.519507 | 0.014635 | 0.045365 | Infection | Intestine |
| Wnt3 | -1.25E+00 | 1.362034 | 0.033795 | 0.855655 | 0.956553 | Healthy Control | Intestine |
| Wnt3a | -3.57E-01 | 1.228299 | 0.00618 | 0.938438 | 0.969735 | Healthy Control | Intestine |
| Wnt4 | -8.78E+00 | 6.42442 | 6.6443 | 0.018063 | 0.054439 | Healthy Control | Intestine |
| Wnt5a | -1.44E+00 | 6.65139 | 2.787612 | 0.107729 | 0.229876 | Healthy Control | Intestine |
| Wnt5b | -6.94E+00 | 4.390176 | 2.733313 | 0.114078 | 0.240655 | Healthy Control | Intestine |
| Wnt6 | -1.58E-01 | 4.089455 | 0.007496 | 0.931781 | 0.969735 | Healthy Control | Intestine |
| Wnt7a | -3.57E-01 | 1.227417 | 0.006288 | 0.937908 | 0.969735 | Healthy Control | Intestine |
| Wnt7b | -3.57E-01 | 1.228299 | 0.00618 | 0.938438 | 0.969735 | Healthy Control | Intestine |
| Wnt8a | -1.58E+00 | 1.274873 | 0.042519 | 0.839052 | 0.954066 | Healthy Control | Intestine |
| Wnt8b | -9.71E-16 | 1.180616 | 0 | 1 | 1 | Healthy Control | Intestine |
| Wnt9a | 3.69E+00 | 2.847131 | 19.26401 | 0.000206 | 0.001169 | Infection | Intestine |
| Wnt9b | -6.42E-01 | 1.273248 | 0.013089 | 0.910112 | 0.969735 | Healthy Control | Intestine |
| Wrap53 | 1.18E+00 | 5.870162 | 6.322218 | 0.019698 | 0.058418 | Infection | Intestine |
| Wrap73 | 4.46E-01 | 6.249243 | 0.74899 | 0.395959 | 0.625094 | Infection | Intestine |
| Wrnip1 | 1.83E-01 | 8.205414 | 0.492154 | 0.490399 | 0.723317 | Infection | Intestine |
| Wsb1 | 8.16E-01 | 7.521173 | 6.406614 | 0.019116 | 0.05703 | Infection | Intestine |
| Wt1 | 5.34E+00 | 1.741643 | 15.87374 | 0.00037 | 0.001955 | Infection | Intestine |
| Wtip | -5.81E-01 | 7.039836 | 1.140978 | 0.296739 | 0.505264 | Healthy Control | Intestine |
| Wwox | 5.12E-01 | 7.181585 | 2.666302 | 0.116847 | 0.24493 | Infection | Intestine |
| Wwtr1 | 2.39E+00 | 5.980759 | 39.10016 | 2.85E-06 | 2.71E-05 | Infection | Intestine |
| Xcl1 | -6.57E+00 | 3.852001 | 0.538001 | 0.474729 | 0.709562 | Healthy Control | Intestine |
| Xcr1 | -1.96E+00 | 5.217098 | 0.482919 | 0.496886 | 0.72966 | Healthy Control | Intestine |
| Xiap | -1.32E+00 | 8.464105 | 15.53759 | 0.000692 | 0.003357 | Healthy Control | Intestine |
| Xist | 2.59E-01 | 9.602763 | 0.121475 | 0.73321 | 0.915324 | Infection | Intestine |
| Xkr6 | -4.17E+00 | 2.060188 | 0.401553 | 0.532246 | 0.765638 | Healthy Control | Intestine |
| Xpa | 2.90E-01 | 6.788535 | 0.472069 | 0.499173 | 0.732165 | Infection | Intestine |
| Xpo1 | 6.10E-01 | 8.082296 | 7.290845 | 0.013145 | 0.041489 | Infection | Intestine |
| Xrcc1 | -5.21E-01 | 6.912772 | 1.002812 | 0.327236 | 0.544233 | Healthy Control | Intestine |
| Xrcc2 | -5.77E-01 | 3.997903 | 0.199882 | 0.659104 | 0.867142 | Healthy Control | Intestine |
| Xrcc4 | 1.14E-02 | 6.333473 | 0.000404 | 0.984138 | 1 | Infection | Intestine |
| Xrcc5 | -9.02E+00 | 6.343139 | 12.05397 | 0.002434 | 0.009777 | Healthy Control | Intestine |
| Xrcc6 | -7.78E-01 | 7.30966 | 3.067596 | 0.093344 | 0.205399 | Healthy Control | Intestine |
| Yap1 | -4.67E-01 | 6.839551 | 1.043936 | 0.317457 | 0.531679 | Healthy Control | Intestine |
| Yars | -9.93E-02 | 8.430323 | 0.21771 | 0.645411 | 0.857921 | Healthy Control | Intestine |
| Ybx3 | -2.38E+00 | 10.29603 | 203.6515 | 1.52E-12 | 1.62E-10 | Healthy Control | Intestine |
| Yeats2 | -1.80E-01 | 6.550575 | 0.097609 | 0.757545 | 0.928591 | Healthy Control | Intestine |
| Yeats4 | 6.65E-01 | 7.921131 | 5.921805 | 0.023631 | 0.068065 | Infection | Intestine |
| Yes1 | -1.58E+00 | 7.995124 | 17.97315 | 0.000309 | 0.001673 | Healthy Control | Intestine |

| | | | | | | | |
|---|---|---|---|---|---|---|---|
| Yif1a | -1.47E-01 | 6.087102 | 0.079538 | 0.780387 | 0.938784 | Healthy Control | Intestine |
| Yipf1 | -1.64E+00 | 8.057244 | 15.90994 | 0.000578 | 0.002881 | Healthy Control | Intestine |
| Ywhae | -9.36E-01 | 9.875725 | 19.33796 | 0.000233 | 0.001307 | Healthy Control | Intestine |
| Ywhag | 3.26E-01 | 8.684297 | 1.846885 | 0.188052 | 0.354038 | Infection | Intestine |
| Ywhaz | 4.13E-02 | 11.17267 | 0.127516 | 0.724454 | 0.909154 | Infection | Intestine |
| Yy1 | 6.84E-01 | 9.486687 | 33.18811 | 8.83E-06 | 7.41E-05 | Infection | Intestine |
| Zap70 | 2.37E+00 | 3.338047 | 3.603637 | 0.069817 | 0.164163 | Infection | Intestine |
| Zbp1 | 1.92E-01 | 5.353404 | 0.031856 | 0.859866 | 0.956553 | Infection | Intestine |
| Zbtb16 | 8.52E-01 | 7.221409 | 5.104206 | 0.033996 | 0.090766 | Infection | Intestine |
| Zbtb4 | 2.85E-01 | 5.509272 | 0.153374 | 0.698825 | 0.893339 | Infection | Intestine |
| Zbtb46 | 3.40E-01 | 4.600893 | 0.069009 | 0.794974 | 0.943769 | Infection | Intestine |
| Zbtb7a | -2.36E+00 | 9.474173 | 88.67943 | 3.11E-09 | 8.42E-08 | Healthy Control | Intestine |
| Zbtb7b | -2.14E+00 | 7.702337 | 17.60485 | 0.000323 | 0.00174 | Healthy Control | Intestine |
| Zbtb8b | -2.18E+00 | 1.520191 | 0.062354 | 0.804844 | 0.947891 | Healthy Control | Intestine |
| Zc3h11a | -5.30E-01 | 9.01189 | 8.071615 | 0.009566 | 0.031694 | Healthy Control | Intestine |
| Zc3h12a | 1.26E+00 | 6.511527 | 12.8736 | 0.00165 | 0.007016 | Infection | Intestine |
| Zc3h14 | -5.80E-01 | 7.122028 | 2.267427 | 0.145913 | 0.291943 | Healthy Control | Intestine |
| Zc3hc1 | 1.66E-01 | 6.515155 | 0.105479 | 0.748377 | 0.924698 | Infection | Intestine |
| Zcchc12 | 1.99E+00 | 1.954296 | 0.830257 | 0.369658 | 0.59521 | Infection | Intestine |
| Zdhhc18 | -1.72E+00 | 7.266105 | 7.088274 | 0.013623 | 0.042704 | Healthy Control | Intestine |
| Zdhhc2 | -2.30E+00 | 7.27052 | 11.85189 | 0.002189 | 0.008922 | Healthy Control | Intestine |
| Zdhhc6 | -1.40E+00 | 7.673046 | 12.47794 | 0.00178 | 0.007495 | Healthy Control | Intestine |
| Zdhhc7 | -2.94E+00 | 8.216873 | 52.13343 | 1.81E-07 | 2.58E-06 | Healthy Control | Intestine |
| Zdhhc8 | 5.49E-01 | 7.117781 | 1.688054 | 0.207382 | 0.382086 | Infection | Intestine |
| Zeb1 | 1.38E+00 | 6.538846 | 12.87989 | 0.001656 | 0.007037 | Infection | Intestine |
| Zeb2 | -5.56E-02 | 8.237163 | 0.017177 | 0.897021 | 0.969735 | Healthy Control | Intestine |
| Zfand2a | 1.80E-01 | 6.360341 | 0.135288 | 0.716391 | 0.902657 | Infection | Intestine |
| Zfand2b | -4.55E-01 | 7.667548 | 1.860935 | 0.186271 | 0.351743 | Healthy Control | Intestine |
| Zfhx3 | -1.63E+00 | 8.239207 | 15.30391 | 0.000705 | 0.003403 | Healthy Control | Intestine |
| Zfp106 | -1.88E+00 | 8.986197 | 31.26502 | 1.25E-05 | 0.0001 | Healthy Control | Intestine |
| Zfp111 | -9.29E+00 | 6.595837 | 17.07567 | 0.000525 | 0.002641 | Healthy Control | Intestine |
| Zfp148 | -1.04E+00 | 7.925245 | 11.32444 | 0.002736 | 0.010818 | Healthy Control | Intestine |
| Zfp219 | -4.42E-01 | 7.163248 | 0.932169 | 0.344565 | 0.566654 | Healthy Control | Intestine |
| Zfp24 | 1.48E+00 | 6.617303 | 13.59687 | 0.001303 | 0.005765 | Infection | Intestine |
| Zfp281 | -7.77E-01 | 5.674059 | 0.825817 | 0.372357 | 0.598401 | Healthy Control | Intestine |
| Zfp42 | -3.57E-01 | 1.227417 | 0.006288 | 0.937908 | 0.969735 | Healthy Control | Intestine |
| Zfp503 | 1.66E+00 | 6.368015 | 10.82007 | 0.00338 | 0.013028 | Infection | Intestine |
| Zfp536 | -8.99E-01 | 4.7033 | 0.26818 | 0.609536 | 0.832108 | Healthy Control | Intestine |
| Zfp57 | 2.29E-01 | 3.379934 | 0.020185 | 0.888288 | 0.969599 | Infection | Intestine |
| Zfp750 | -1.73E+00 | 6.241142 | 3.312447 | 0.081933 | 0.186131 | Healthy Control | Intestine |
| Zfp819 | 4.84E+00 | 1.362495 | 3.49247 | 0.073689 | 0.171337 | Infection | Intestine |
| Zfp979 | 2.07E+00 | 2.939577 | 2.848189 | 0.10431 | 0.223895 | Infection | Intestine |
| Zfpl1 | 1.39E-01 | 4.852557 | 0.028172 | 0.8681 | 0.960668 | Infection | Intestine |
| Zfpm2 | 2.86E+00 | 3.667388 | 13.47688 | 0.001284 | 0.0057 | Infection | Intestine |
| Zfyve9 | 2.78E-01 | 6.575753 | 0.340726 | 0.565328 | 0.793616 | Infection | Intestine |
| Zic1 | -6.42E-01 | 1.273689 | 0.012641 | 0.911654 | 0.969735 | Healthy Control | Intestine |
| Zic2 | -9.77E-01 | 1.317623 | 0.02244 | 0.882264 | 0.96867 | Healthy Control | Intestine |
| Zic3 | -1.65E+00 | 1.318958 | 0.045773 | 0.832543 | 0.954066 | Healthy Control | Intestine |
| Zic4 | 4.59E+00 | 1.405504 | 3.316998 | 0.079059 | 0.180716 | Infection | Intestine |
| Zkscan1 | -7.65E-01 | 8.315968 | 7.189669 | 0.013677 | 0.042791 | Healthy Control | Intestine |

| | | | | | | | |
|---|---|---|---|---|---|---|---|
| Zmpste24 | -2.49E-01 | 7.839874 | 0.738674 | 0.399416 | 0.629161 | Healthy Control | Intestine |
| Zmynd10 | -1.81E+00 | 1.556082 | 0.066193 | 0.799567 | 0.945927 | Healthy Control | Intestine |
| Zmynd19 | 1.07E+00 | 5.829699 | 4.90101 | 0.037335 | 0.098212 | Infection | Intestine |
| Zp1 | -3.75E+00 | 1.79832 | 0.283237 | 0.598995 | 0.823556 | Healthy Control | Intestine |
| Zp3 | -2.63E+00 | 1.365078 | 0.074391 | 0.787973 | 0.941655 | Healthy Control | Intestine |
| Zscan2 | 6.95E-01 | 5.442809 | 0.684113 | 0.416547 | 0.649201 | Infection | Intestine |
| Zscan21 | -7.63E-01 | 6.96403 | 2.237611 | 0.14822 | 0.295614 | Healthy Control | Intestine |
| Zswim5 | -1.31E+00 | 5.599732 | 2.142967 | 0.156231 | 0.307184 | Healthy Control | Intestine |
| Zswim9 | 1.82E-01 | 4.901326 | 0.031509 | 0.860569 | 0.95691 | Infection | Intestine |
| Zup1 | -4.41E-01 | 6.132143 | 0.55312 | 0.46434 | 0.699958 | Healthy Control | Intestine |
| Zyx | 7.25E-01 | 7.54305 | 7.716573 | 0.011039 | 0.035746 | Infection | Intestine |
| Zzef1 | 3.28E-01 | 8.746859 | 3.056492 | 0.094496 | 0.207204 | Infection | Intestine |
| a | 2.52E+00 | 2.108758 | 1.628138 | 0.212316 | 0.389467 | Infection | Intestine |
