## Supplementary Table 3 for "Mapping fibrotic microenvironments: single-cell and spatial profiling of Schistosoma mansoni-induced tissue fibrosis"

| Tissue | Term | Overlap | P-value | Adjusted P-value | Old P-value | Old Adjusted P-value | Odds Ratio | Combined Score | Genes |
|---|---|---|---|---|---|---|---|---|---|
| Intestine | Interleukin-4 and Interleukin-13 Signaling | 12/112 | 1.66E-14 | 6.16E-12 | 0 | 0 | 36.04 | 1143.588761 | ITGB1;IL4;IL1A;SOCS3;ITGAM;COL1A2;ALOX5;ITGB2;ALOX15;HMOX1;TIMP1;IL13RA2 |
| Intestine | Signaling by Interleukins | 16/452 | 1.69E-11 | 3.15E-09 | 0 | 0 | 11.53358982 | 286.048773 | ITGB1;DUSP4;IL33;ITGAM;ITGB2;ALOX15;IL4;IL1A;HCK;SOCS3;COL1A2;ALOX5;HMOX1;TIMP1;IL13RA2;CCR5 |
| Intestine | Extracellular Matrix Organization | 13/300 | 1.38E-10 | 1.72E-08 | 0 | 0 | 13.68292683 | 310.6176677 | ITGB1;ITGAM;COL24A1;ITGB2;SERPINE1;DMP1;MFAP5;MMP25;COL1A2;BMP1;COL6A1;TIMP1;ITGA5 |
| Intestine | Immune System | 28/2150 | 3.58E-09 | 3.33E-07 | 0 | 0 | 4.697455231 | 91.35666887 | ITGB1;CD274;ITGAM;NCF1;ITGB2;ALOX15;SLC2A3;PYGL;SOCS3;MMP25;ALOX5;OLR1;SIRPA;HMOX1;TIMP1;IL13RA2;CCR5;CD300LF;DUSP4;IL33;ARG1;ENAH;IL4;IL1A;HCK;CD200R1;COL1A2;FCGR2B |
| Intestine | Cytokine Signaling in Immune System | 16/776 | 4.10E-08 | 3.05E-06 | 0 | 0 | 6.506621392 | 110.6708316 | ITGB1;DUSP4;IL33;ITGAM;ITGB2;ALOX15;IL4;IL1A;HCK;SOCS3;COL1A2;ALOX5;HMOX1;TIMP1;IL13RA2;CCR5 |
| Intestine | Cell Surface Interactions at the Vascular Wall | 9/233 | 3.07E-07 | 1.91E-05 | 0 | 0 | 11.4701087 | 171.9987978 | ITGB1;CD84;ITGAM;COL1A2;ITGB2;OLR1;SIRPA;ITGA5;SLC16A3 |
| Intestine | Integrin Cell Surface Interactions | 6/85 | 9.91E-07 | 5.27E-05 | 0 | 0 | 20.9314346 | 289.3722246 | ITGB1;ITGAM;COL1A2;ITGB2;COL6A1;ITGA5 |
| Intestine | Hemostasis | 13/707 | 3.20E-06 | 1.49E-04 | 0 | 0 | 5.541210375 | 70.11171566 | ITGB1;CD84;ITGAM;ITGB2;SERPINE1;ARRB2;IGF1;COL1A2;OLR1;SIRPA;TIMP1;ITGA5;SLC16A3 |
| Intestine | ECM Proteoglycans | 5/76 | 1.18E-05 | 4.87E-04 | 0 | 0 | 19.15010612 | 217.3206379 | ITGB1;COL1A2;SERPINE1;COL6A1;DMP1 |
| Intestine | Signaling by TGFB Family Members | 6/161 | 3.95E-05 | 0.001467729 | 0 | 0 | 10.62741935 | 107.7657262 | ITGB1;COL1A2;SERPINE1;ARRB2;TIMP1;INHBA |
| Intestine | Assembly of Collagen Fibrils and Other Multimeric Structures | 4/61 | 9.44E-05 | 0.003038322 | 0 | 0 | 18.838312 | 174.6013575 | BMP1;COL1A2;COL24A1;COL6A1 |
| Intestine | Neutrophil Degranulation | 9/478 | 9.98E-05 | 0.003038322 | 0 | 0 | 5.410123297 | 49.84076466 | MMP25;ITGAM;ARG1;ALOX5;ITGB2;OLR1;SIRPA;SLC2A3;PYGL |
| Intestine | Signal Transduction | 23/2613 | 1.06E-04 | 0.003038322 | 0 | 0 | 2.798427518 | 25.60671525 | ITGB1;DUSP4;ABCA1;CD274;IL33;NCF1;COL24A1;SERPINE1;ARRB2;INHBA;IGF1;SOCS3;COL1A2;CCL7;ALDH1A2;COL6A1;GAS1;ARHGEF1;TIMP1;ITGA5;CCR5;CCR3;ARHGEF6 |
| Intestine | Collagen Biosynthesis and Modifying Enzymes | 4/67 | 1.36E-04 | 0.003617907 | 0 | 0 | 17.03903904 | 151.6763484 | BMP1;COL1A2;COL24A1;COL6A1 |
| Intestine | Biosynthesis of E-series 18(S)-resolvins | 2/5 | 1.49E-04 | 0.003695543 | 0 | 0 | 174.7280702 | 1539.61138 | ALOX5;ALOX15 |
| Intestine | MET Activates PTK2 Signaling | 3/30 | 2.15E-04 | 0.004365237 | 0 | 0 | 29.47407407 | 248.9400929 | ITGB1;COL1A2;COL24A1 |
| Intestine | Degradation of the Extracellular Matrix | 5/140 | 2.19E-04 | 0.004365237 | 0 | 0 | 10.03906646 | 84.58618666 | MMP25;COL1A2;BMP1;COL6A1;TIMP1 |
| Intestine | Fibronectin Matrix Formation | 2/6 | 2.23E-04 | 0.004365237 | 0 | 0 | 131.0394737 | 1101.850345 | ITGB1;ITGA5 |
| Intestine | Biosynthesis of EPA-derived SPMs | 2/6 | 2.23E-04 | 0.004365237 | 0 | 0 | 131.0394737 | 1101.850345 | ALOX5;ALOX15 |
| Intestine | Immunoregulatory Interactions Between a Lymphoid and a non-Lymphoid Cell | 6/223 | 2.36E-04 | 0.004396687 | 0 | 0 | 7.567204301 | 63.18665163 | ITGB1;CD200R1;COL1A2;ITGB2;CD300LF;FCGR2B |
| Intestine | Cell-Cell Communication | 5/157 | 3.72E-04 | 0.006574645 | 0 | 0 | 8.908615717 | 70.34439095 | ITGB1;CDH6;SIRPA;ANGPTL4;ARHGEF6 |
| Intestine | HDL Assembly | 2/8 | 4.14E-04 | 0.006574645 | 0 | 0 | 87.35087719 | 680.4149259 | ABCA1;BMP1 |
| Intestine | Interleukin-18 Signaling | 2/8 | 4.14E-04 | 0.006574645 | 0 | 0 | 87.35087719 | 680.4149259 | IL4;ALOX5 |
| Intestine | Collagen Formation | 4/90 | 4.24E-04 | 0.006574645 | 0 | 0 | 12.46763042 | 96.81582205 | COL1A2;BMP1;COL24A1;COL6A1 |
| Intestine | Innate Immune System | 13/1149 | 4.68E-04 | 0.006959638 | 0 | 0 | 3.307394366 | 25.35992888 | DUSP4;ITGAM;NCF1;ARG1;ITGB2;SLC2A3;PYGL;HCK;MMP25;ALOX5;OLR1;SIRPA;HMOX1 |
| Intestine | MET Promotes Cell Motility | 3/41 | 5.47E-04 | 0.007822222 | 0 | 0 | 20.93052632 | 157.221406 | ITGB1;COL1A2;COL24A1 |
| Intestine | Elastic Fibre Formation | 3/44 | 6.74E-04 | 0.008948693 | 0 | 0 | 19.39609756 | 141.6484818 | ITGB1;MFAP5;ITGA5 |
| Intestine | Collagen Chain Trimerization | 3/44 | 6.74E-04 | 0.008948693 | 0 | 0 | 19.39609756 | 141.6484818 | COL1A2;COL24A1;COL6A1 |
| Intestine | Interleukin-10 Signaling | 3/46 | 7.68E-04 | 0.009847846 | 0 | 0 | 18.49209302 | 132.62715 | IL1A;TIMP1;CCR5 |
| Intestine | Signaling by TGFBR3 | 3/49 | 9.24E-04 | 0.011457374 | 0 | 0 | 17.28347826 | 120.7565207 | ARRB2;TIMP1;INHBA |
| Intestine | Signaling by Receptor Tyrosine Kinases | 8/536 | 0.001143 | 0.013713348 | 0 | 0 | 4.197835498 | 28.43736446 | ITGB1;DUSP4;CD274;COL1A2;NCF1;COL24A1;COL6A1;IGF1 |
| Intestine | Cell Junction Organization | 4/119 | 0.001211 | 0.014072093 | 0 | 0 | 9.309988249 | 62.53258054 | ITGB1;CDH6;ANGPTL4;ARHGEF6 |
| Intestine | Chemokine Receptors Bind Chemokines | 3/57 | 0.001435 | 0.016175986 | 0 | 0 | 14.71703704 | 96.346757 | CCL7;CCR5;CCR3 |
| Intestine | Anchoring Fibril Formation | 2/15 | 0.001526 | 0.016691416 | 0 | 0 | 40.30161943 | 261.3718812 | BMP1;COL1A2 |
| Intestine | Adaptive Immune System | 10/854 | 0.001735 | 0.017970667 | 0 | 0 | 3.324156677 | 21.13004431 | ITGB1;ENAH;CD274;SOCS3;CD200R1;COL1A2;NCF1;ITGB2;CD300LF;FCGR2B |
| Intestine | Platelet Adhesion to Exposed Collagen | 2/16 | 0.001739 | 0.017970667 | 0 | 0 | 37.42105263 | 237.7879384 | ITGB1;COL1A2 |
| Intestine | Biosynthesis of DHA-derived SPMs | 2/17 | 0.001966 | 0.019766309 | 0 | 0 | 34.9245614 | 217.6412156 | ALOX5;ALOX15 |
| Intestine | Interleukin-1 Family Signaling | 4/140 | 0.002197 | 0.02001701 | 0 | 0 | 7.864069952 | 48.13213229 | IL4;IL1A;IL33;ALOX5 |
| Intestine | Activation of SMO | 2/18 | 0.002206 | 0.02001701 | 0 | 0 | 32.74013158 | 200.2548405 | GAS1;ARRB2 |
| Intestine | Cell-extracellular Matrix Interactions | 2/18 | 0.002206 | 0.02001701 | 0 | 0 | 32.74013158 | 200.2548405 | ITGB1;ARHGEF6 |
| Intestine | Crosslinking of Collagen Fibrils | 2/18 | 0.002206 | 0.02001701 | 0 | 0 | 32.74013158 | 200.2548405 | BMP1;COL1A2 |
| Intestine | Biosynthesis of Specialized Proresolving Mediators (SPMs) | 2/19 | 0.00246 | 0.021277607 | 0 | 0 | 30.8126935 | 185.1163087 | ALOX5;ALOX15 |
| Intestine | Plasma Lipoprotein Assembly | 2/19 | 0.00246 | 0.021277607 | 0 | 0 | 30.8126935 | 185.1163087 | ABCA1;BMP1 |
| Intestine | Synthesis of Leukotrienes (LT) and Eoxins (EX) | 2/20 | 0.002726 | 0.023046229 | 0 | 0 | 29.0994152 | 171.8308057 | ALOX5;ALOX15 |
| Intestine | Signal Transduction by L1 | 2/21 | 0.003005 | 0.024843331 | 0 | 0 | 27.56648199 | 160.0895126 | ITGB1;ITGA5 |
| Intestine | Plasma Lipoprotein Assembly, Remodeling, and Clearance | 3/75 | 0.003149 | 0.025468846 | 0 | 0 | 11.02777778 | 63.52608468 | ABCA1;BMP1;ANGPTL4 |
| Intestine | Platelet Activation, Signaling and Aggregation | 5/262 | 0.003602 | 0.028267323 | 0 | 0 | 5.240925324 | 29.48721883 | COL1A2;SERPINE1;ARRB2;TIMP1;IGF1 |
| Intestine | Signaling by MET | 3/79 | 0.003647 | 0.028267323 | 0 | 0 | 10.44526316 | 58.63700899 | ITGB1;COL1A2;COL24A1 |

| | | | | | | | | | |
|---|---|---|---|---|---|---|---|---|---|
| Intestine | Basigin Interactions | 2/25 | 0.00425 | 0.031620863 | 0 | 0 | 22.76773455 | 124.3302497 | ITGB1;SLC16A3 |
| Intestine | Inactivation of CSF3 (G-CSF) Signaling | 2/25 | 0.00425 | 0.031620863 | 0 | 0 | 22.76773455 | 124.3302497 | SOCS3;HCK |
| Intestine | Cell Recruitment (Pro-Inflammatory Response) | 2/27 | 0.004948 | 0.034726838 | 0 | 0 | 20.94421053 | 111.1895545 | IL1A;HMOX1 |
| Intestine | Syndecan Interactions | 2/27 | 0.004948 | 0.034726838 | 0 | 0 | 20.94421053 | 111.1895545 | ITGB1;COL1A2 |
| Intestine | Purinergic Signaling in Leishmaniasis Infection | 2/27 | 0.004948 | 0.034726838 | 0 | 0 | 20.94421053 | 111.1895545 | IL1A;HMOX1 |
| Intestine | Signaling by TGF-beta Receptor Complex | 3/94 | 0.00593 | 0.040851171 | 0 | 0 | 8.716923077 | 44.69802447 | ITGB1;COL1A2;SERPINE1 |
| Intestine | Signaling by CSF3 (G-CSF) | 2/30 | 0.006086 | 0.041044526 | 0 | 0 | 18.69736842 | 95.39094459 | SOCS3;HCK |
| Intestine | Signaling by GPCR | 8/706 | 0.006179 | 0.041044526 | 0 | 0 | 3.147605403 | 16.01073547 | ITGB1;CCL7;ARHGEF1;ARRB2;ITGA5;CCR5;CCR3;ARHGEF6 |
| Intestine | Turbulent Flow Shear Stress Activates Signaling by PIEZO1 and Integrins in Endothelial Cells | 2/31 | 0.006489 | 0.042348579 | 0 | 0 | 18.05172414 | 90.93850284 | ITGB1;ITGA5 |
| Intestine | Activation of Matrix Metalloproteinases | 2/33 | 0.007331 | 0.047019921 | 0 | 0 | 16.88539898 | 83.00245407 | MMP25;TIMP1 |
| Intestine | SMAD2 SMAD3 SMAD4 Heterotrimer Regulates Transcription | 2/36 | 0.008682 | 0.054738095 | 0 | 0 | 15.39318885 | 73.06456812 | COL1A2;SERPINE1 |
| Intestine | Molecules Associated With Elastic Fibres | 2/37 | 0.009155 | 0.056758791 | 0 | 0 | 14.95263158 | 70.18008698 | ITGB1;MFAP5 |
| Intestine | GPCR Downstream Signalling | 7/633 | 0.011693 | 0.071306203 | 0 | 0 | 3.039013635 | 13.51993937 | ITGB1;ARHGEF1;ARRB2;ITGA5;CCR5;CCR3;ARHGEF6 |
| Intestine | Regulation of IGF Transport and Uptake by Insulin-like Growth Factor Binding Proteins (IGFBPs) | 3/124 | 0.012626 | 0.075753167 | 0 | 0 | 6.545785124 | 28.61839837 | DMP1;TIMP1;IGF1 |
| Intestine | GPER1 Signaling | 2/45 | 0.013338 | 0.078756829 | 0 | 0 | 12.16585067 | 52.52179508 | ITGB1;ITGA5 |
| Intestine | Platelet Degranulation | 3/128 | 0.013745 | 0.07989348 | 0 | 0 | 6.33504 | 27.15877136 | SERPINE1;TIMP1;IGF1 |
| Intestine | Response to Elevated Platelet Cytosolic Ca2+ | 3/133 | 0.015222 | 0.087115757 | 0 | 0 | 6.089846154 | 25.48615286 | SERPINE1;TIMP1;IGF1 |
| Intestine | Transcriptional Activity of SMAD2 SMAD3 SMAD4 Heterotrimer | 2/51 | 0.016921 | 0.095375245 | 0 | 0 | 10.67293233 | 43.53676225 | COL1A2;SERPINE1 |
| Intestine | MET Interacts With TNS Proteins | 1/5 | 0.01935 | 0.105857324 | 0 | 0 | 64.66883117 | 255.1217235 | ITGB1 |
| Intestine | Regulation of HMOX1 Expression and Activity | 1/5 | 0.01935 | 0.105857324 | 0 | 0 | 64.66883117 | 255.1217235 | HMOX1 |
| Intestine | Toll Like Receptor 4 (TLR4) Cascade | 3/148 | 0.020172 | 0.108754019 | 0 | 0 | 5.455724138 | 21.29616825 | DUSP4;ITGAM;ITGB2 |
| Intestine | NRAGE Signals Death Through JNK | 2/59 | 0.022258 | 0.110530444 | 0 | 0 | 9.171283472 | 34.89703029 | ARHGEF1;ARHGEF6 |
| Intestine | Adherens Junctions Interactions | 2/59 | 0.022258 | 0.110530444 | 0 | 0 | 9.171283472 | 34.89703029 | CDH6;ANGPTL4 |
| Intestine | Non-integrin membrane-ECM Interactions | 2/59 | 0.022258 | 0.110530444 | 0 | 0 | 9.171283472 | 34.89703029 | ITGB1;COL1A2 |
| Intestine | Arachidonate Metabolism | 2/59 | 0.022258 | 0.110530444 | 0 | 0 | 9.171283472 | 34.89703029 | ALOX5;ALOX15 |
| Intestine | Transport of Small Molecules | 7/724 | 0.022733 | 0.110530444 | 0 | 0 | 2.640795961 | 9.992635631 | ABCA1;ABCD2;BMP1;HMOX1;SLC2A3;ANGPTL4;SLC16A3 |
| Intestine | FLT3 Signaling Through SRC Family Kinases | 1/6 | 0.023176 | 0.110530444 | 0 | 0 | 51.73246753 | 194.7545981 | HCK |
| Intestine | Biosynthesis of Lipoxins (LX) | 1/6 | 0.023176 | 0.110530444 | 0 | 0 | 51.73246753 | 194.7545981 | ALOX5 |
| Intestine | Synthesis of 15-Eicosatetraenoic Acid Derivatives | 1/6 | 0.023176 | 0.110530444 | 0 | 0 | 51.73246753 | 194.7545981 | ALOX15 |
| Intestine | Proton-coupled Monocarboxylate Transport | 1/6 | 0.023176 | 0.110530444 | 0 | 0 | 51.73246753 | 194.7545981 | SLC16A3 |
| Intestine | G Alpha (S) Signalling Events | 3/158 | 0.023909 | 0.112583727 | 0 | 0 | 5.10116129 | 19.04520658 | ITGB1;ARRB2;ITGA5 |
| Intestine | Collagen Degradation | 2/64 | 0.025901 | 0.120440814 | 0 | 0 | 8.429541596 | 30.79702707 | COL1A2;COL6A1 |
| Intestine | Enhanced Binding of GP1BA Variant to VWF Multimer Collagen | 1/7 | 0.026986 | 0.120951462 | 0 | 0 | 43.10822511 | 155.72498 | COL1A2 |
| Intestine | Synthesis of 12-Eicosatetraenoic Acid Derivatives | 1/7 | 0.026986 | 0.120951462 | 0 | 0 | 43.10822511 | 155.72498 | ALOX15 |
| Intestine | Defective Binding of VWF Variant to GPIb IX V | 1/7 | 0.026986 | 0.120951462 | 0 | 0 | 43.10822511 | 155.72498 | COL1A2 |
| Intestine | Toll-like Receptor Cascades | 3/174 | 0.030615 | 0.123130203 | 0 | 0 | 4.620116959 | 16.10688892 | DUSP4;ITGAM;ITGB2 |
| Intestine | Biosynthesis of Maresins | 1/8 | 0.030783 | 0.123130203 | 0 | 0 | 36.94805195 | 128.6090483 | ALOX5 |
| Intestine | TGFBR3 Regulates TGF-beta Signaling | 1/8 | 0.030783 | 0.123130203 | 0 | 0 | 36.94805195 | 128.6090483 | ARRB2 |
| Intestine | Cross-presentation of Particulate Exogenous Antigens (Phagosomes) | 1/8 | 0.030783 | 0.123130203 | 0 | 0 | 36.94805195 | 128.6090483 | NCF1 |
| Intestine | RUNX1 Regulates Transcription of Genes Involved in Differentiation of Keratinocytes | 1/8 | 0.030783 | 0.123130203 | 0 | 0 | 36.94805195 | 128.6090483 | SOCS3 |
| Intestine | Ligand-receptor Interactions | 1/8 | 0.030783 | 0.123130203 | 0 | 0 | 36.94805195 | 128.6090483 | GAS1 |
| Intestine | RUNX2 Regulates Genes Involved in Cell Migration | 1/8 | 0.030783 | 0.123130203 | 0 | 0 | 36.94805195 | 128.6090483 | ITGA5 |
| Intestine | Defects of Platelet Adhesion to Exposed Collagen | 1/8 | 0.030783 | 0.123130203 | 0 | 0 | 36.94805195 | 128.6090483 | COL1A2 |
| Intestine | Regulation of Cytoskeletal Remodeling and Cell Spreading by IPP Complex Components | 1/8 | 0.030783 | 0.123130203 | 0 | 0 | 36.94805195 | 128.6090483 | ARHGEF6 |

| | | | | | | | | | |
|---|---|---|---|---|---|---|---|---|---|
| Intestine | Vitamin C (Ascorbate) Metabolism | 1/8 | 0.030783 | 0.123130203 | 0 | 0 | 36.94805195 | 128.6090483 | SLC2A3 |
| Intestine | Hedgehog 'On' State | 2/73 | 0.033016 | 0.129876847 | 0 | 0 | 7.35767235 | 25.09520887 | GAS1;ARRB2 |
| Intestine | SHC-related Events Triggered by IGF1R | 1/9 | 0.034564 | 0.129876847 | 0 | 0 | 32.32792208 | 108.7816026 | IGF1 |
| Intestine | Nef and Signal Transduction | 1/9 | 0.034564 | 0.129876847 | 0 | 0 | 32.32792208 | 108.7816026 | HCK |
| Intestine | CHL1 Interactions | 1/9 | 0.034564 | 0.129876847 | 0 | 0 | 32.32792208 | 108.7816026 | ITGB1 |
| Intestine | Synthesis of 5-Eicosatetraenoic Acids | 1/9 | 0.034564 | 0.129876847 | 0 | 0 | 32.32792208 | 108.7816026 | ALOX5 |
| Intestine | Interleukin-1 Processing | 1/9 | 0.034564 | 0.129876847 | 0 | 0 | 32.32792208 | 108.7816026 | IL1A |
| Intestine | RAC1 GTPase Cycle | 3/184 | 0.035259 | 0.130897989 | 0 | 0 | 4.362651934 | 14.59321317 | ITGB1;NCF1;ARHGEF6 |
| Intestine | Cell Death Signalling via NRAGE, NRIF and NADE | 2/76 | 0.03554 | 0.130897989 | 0 | 0 | 7.058321479 | 23.55439737 | ARHGEF1;ARHGEF6 |
| Intestine | Glycoprotein Hormones | 1/10 | 0.038331 | 0.138437756 | 0 | 0 | 28.73448773 | 93.71751272 | INHBA |
| Intestine | Urea Cycle | 1/10 | 0.038331 | 0.138437756 | 0 | 0 | 28.73448773 | 93.71751272 | ARG1 |
| Intestine | G Alpha (12 13) Signalling Events | 2/80 | 0.039016 | 0.139557248 | 0 | 0 | 6.695006748 | 21.71715132 | ARHGEF1;ARHGEF6 |
| Intestine | STAT3 Nuclear Events Downstream of ALK Signaling | 1/11 | 0.042083 | 0.141884113 | 0 | 0 | 25.85974026 | 81.92637909 | CD274 |
| Intestine | Signaling by Leptin | 1/11 | 0.042083 | 0.141884113 | 0 | 0 | 25.85974026 | 81.92637909 | SOCS3 |
| Intestine | TGFBR3 PTM Regulation | 1/11 | 0.042083 | 0.141884113 | 0 | 0 | 25.85974026 | 81.92637909 | TIMP1 |
| Intestine | Interleukin-6 Signaling | 1/11 | 0.042083 | 0.141884113 | 0 | 0 | 25.85974026 | 81.92637909 | SOCS3 |
| Intestine | Regulation of CDH11 Function | 1/11 | 0.042083 | 0.141884113 | 0 | 0 | 25.85974026 | 81.92637909 | ANGPTL4 |
| Intestine | Peptide Ligand-Binding Receptors | 3/198 | 0.042336 | 0.141884113 | 0 | 0 | 4.046564103 | 12.795674 | CCL7;CCR5;CCR3 |
| Intestine | Metabolism of Vitamins and Cofactors | 3/198 | 0.042336 | 0.141884113 | 0 | 0 | 4.046564103 | 12.795674 | RBP1;SLC2A3;ALDH1L2 |
| Intestine | Peptide Hormone Metabolism | 2/86 | 0.044462 | 0.146943786 | 0 | 0 | 6.214912281 | 19.34780389 | INHBA;IGF1 |
| Intestine | GP1b-IX-V Activation Signalling | 1/12 | 0.045821 | 0.146943786 | 0 | 0 | 23.50767414 | 72.47436707 | COL1A2 |
| Intestine | Binding and Entry of HIV Virion | 1/12 | 0.045821 | 0.146943786 | 0 | 0 | 23.50767414 | 72.47436707 | CCR5 |
| Intestine | Peptide Hormone Biosynthesis | 1/12 | 0.045821 | 0.146943786 | 0 | 0 | 23.50767414 | 72.47436707 | INHBA |
| Intestine | Presynaptic Depolarization and Calcium Channel Opening | 1/12 | 0.045821 | 0.146943786 | 0 | 0 | 23.50767414 | 72.47436707 | CACNA2D1 |
| Intestine | RAC2 GTPase Cycle | 2/88 | 0.046336 | 0.147325772 | 0 | 0 | 6.069767442 | 18.64528721 | ITGB1;NCF1 |
| Intestine | Retinoid Cycle Disease Events | 1/13 | 0.049545 | 0.151070733 | 0 | 0 | 21.54761905 | 64.74800798 | RBP1 |
| Intestine | Diseases Associated With Visual Transduction | 1/13 | 0.049545 | 0.151070733 | 0 | 0 | 21.54761905 | 64.74800798 | RBP1 |
| Intestine | Diseases of the Neuronal System | 1/13 | 0.049545 | 0.151070733 | 0 | 0 | 21.54761905 | 64.74800798 | RBP1 |
| Intestine | Dissolution of Fibrin Clot | 1/13 | 0.049545 | 0.151070733 | 0 | 0 | 21.54761905 | 64.74800798 | SERPINE1 |
| Intestine | ERKs Are Inactivated | 1/13 | 0.049545 | 0.151070733 | 0 | 0 | 21.54761905 | 64.74800798 | DUSP4 |
| Intestine | Cell-cell Junction Organization | 2/92 | 0.050171 | 0.151737051 | 0 | 0 | 5.798830409 | 17.35193118 | CDH6;ANGPTL4 |
| Intestine | Response of Endothelial Cells to Shear Stress | 2/93 | 0.051147 | 0.152214257 | 0 | 0 | 5.734817814 | 17.04987877 | ITGB1;ITGA5 |
| Intestine | Cellular Responses to Mechanical Stimuli | 2/93 | 0.051147 | 0.152214257 | 0 | 0 | 5.734817814 | 17.04987877 | ITGB1;ITGA5 |
| Intestine | RAC3 GTPase Cycle | 2/94 | 0.05213 | 0.153908355 | 0 | 0 | 5.672196796 | 16.75572549 | ITGB1;NCF1 |
| Intestine | Glycogen Breakdown (Glycogenolysis) | 1/14 | 0.053254 | 0.154769091 | 0 | 0 | 19.88911089 | 58.32848923 | PYGL |
| Intestine | Regulation of IFNG Signaling | 1/14 | 0.053254 | 0.154769091 | 0 | 0 | 19.88911089 | 58.32848923 | SOCS3 |
| Intestine | P75 NTR Receptor-Mediated Signalling | 2/98 | 0.056129 | 0.16186122 | 0 | 0 | 5.434758772 | 15.65263476 | ARHGEF1;ARHGEF6 |
| Intestine | WNT5A-dependent Internalization of FZD4 | 1/15 | 0.056949 | 0.162961091 | 0 | 0 | 18.46753247 | 52.9206204 | ARRB2 |
| Intestine | Signal Regulatory Protein Family Interactions | 1/16 | 0.060629 | 0.170864731 | 0 | 0 | 17.23549784 | 48.31066981 | SIRPA |
| Intestine | Heme Degradation | 1/16 | 0.060629 | 0.170864731 | 0 | 0 | 17.23549784 | 48.31066981 | HMOX1 |
| Intestine | Signaling by Activin | 1/17 | 0.064296 | 0.175868173 | 0 | 0 | 16.15746753 | 44.34028458 | INHBA |
| Intestine | Specification of the Neural Plate Border | 1/17 | 0.064296 | 0.175868173 | 0 | 0 | 16.15746753 | 44.34028458 | MSX1 |
| Intestine | The NLRP3 Inflammasome | 1/17 | 0.064296 | 0.175868173 | 0 | 0 | 16.15746753 | 44.34028458 | HMOX1 |
| Intestine | Metabolism of Folate and Pterines | 1/17 | 0.064296 | 0.175868173 | 0 | 0 | 16.15746753 | 44.34028458 | ALDH1L2 |
| Intestine | Post-translational Protein Phosphorylation | 2/107 | 0.065502 | 0.177859421 | 0 | 0 | 4.966666667 | 13.53751788 | DMP1;TIMP1 |
| Intestine | ABC Transporters in Lipid Homeostasis | 1/18 | 0.067948 | 0.179933643 | 0 | 0 | 15.20626432 | 40.88978354 | ABCD2 |
| Intestine | MECP2 Regulates Neuronal Receptors and Channels | 1/18 | 0.067948 | 0.179933643 | 0 | 0 | 15.20626432 | 40.88978354 | SLC2A3 |
| Intestine | Parasitic Infection Pathways | 3/245 | 0.070805 | 0.179933643 | 0 | 0 | 3.252892562 | 8.613107214 | IL1A;HCK;HMOX1 |
| Intestine | Leishmania Infection | 3/245 | 0.070805 | 0.179933643 | 0 | 0 | 3.252892562 | 8.613107214 | IL1A;HCK;HMOX1 |
| Intestine | NFE2L2 Regulating Anti-Oxidant Detoxification Enzymes | 1/19 | 0.071587 | 0.179933643 | 0 | 0 | 14.36075036 | 37.86712629 | HMOX1 |
| Intestine | SARS-CoV-1 Targets Host Intracellular Signalling and Regulatory Pathways | 1/19 | 0.071587 | 0.179933643 | 0 | 0 | 14.36075036 | 37.86712629 | SERPINE1 |
| Intestine | Scavenging by Class A Receptors | 1/19 | 0.071587 | 0.179933643 | 0 | 0 | 14.36075036 | 37.86712629 | COL1A2 |
| Intestine | Assembly of Active LPL and LIPC Lipase Complexes | 1/19 | 0.071587 | 0.179933643 | 0 | 0 | 14.36075036 | 37.86712629 | ANGPTL4 |

| | | | | | | | | | |
|---|---|---|---|---|---|---|---|---|---|
| Intestine | Other Semaphorin Interactions | 1/19 | 0.071587 | 0.179933643 | 0 | 0 | 14.36075036 | 37.86712629 | ITGB1 |
| Intestine | Synthesis, Secretion, and Deacylation of Ghrelin | 1/19 | 0.071587 | 0.179933643 | 0 | 0 | 14.36075036 | 37.86712629 | IGF1 |
| Intestine | Diseases of Hemostasis | 1/19 | 0.071587 | 0.179933643 | 0 | 0 | 14.36075036 | 37.86712629 | COL1A2 |
| Intestine | G Beta Gamma Signalling Through CDC42 | 1/20 | 0.075211 | 0.186522664 | 0 | 0 | 13.60423787 | 35.20043604 | ARHGEF6 |
| Intestine | Class I Peroxisomal Membrane Protein Import | 1/20 | 0.075211 | 0.186522664 | 0 | 0 | 13.60423787 | 35.20043604 | ABCD2 |
| Intestine | PPARA Activates Gene Expression | 2/117 | 0.076479 | 0.188412506 | 0 | 0 | 4.532494279 | 11.6518442 | ABCA1;ANGPTL4 |
| Intestine | Regulation of Lipid Metabolism by PPARalpha | 2/119 | 0.078741 | 0.190430267 | 0 | 0 | 4.454565902 | 11.32169678 | ABCA1;ANGPTL4 |
| Intestine | Formation of the Ureteric Bud | 1/21 | 0.078821 | 0.190430267 | 0 | 0 | 12.92337662 | 32.83281271 | ITGB1 |
| Intestine | L1CAM Interactions | 2/120 | 0.079879 | 0.190430267 | 0 | 0 | 4.416592328 | 11.16177353 | ITGB1;ITGA5 |
| Intestine | Cellular Hexose Transport | 1/22 | 0.082417 | 0.190430267 | 0 | 0 | 12.30735931 | 30.71866043 | SLC2A3 |
| Intestine | RA Biosynthesis Pathway | 1/22 | 0.082417 | 0.190430267 | 0 | 0 | 12.30735931 | 30.71866043 | ALDH1A2 |
| Intestine | Insertion of Tail-Anchored Proteins Into the Endoplasmic Reticulum Membrane | 1/22 | 0.082417 | 0.190430267 | 0 | 0 | 12.30735931 | 30.71866043 | HMOX1 |
| Intestine | The Canonical Retinoid Cycle in Rods (Twilight Vision) | 1/22 | 0.082417 | 0.190430267 | 0 | 0 | 12.30735931 | 30.71866043 | RBP1 |
| Intestine | Regulation of Signaling by CBL | 1/22 | 0.082417 | 0.190430267 | 0 | 0 | 12.30735931 | 30.71866043 | HCK |
| Intestine | ERK MAPK Targets | 1/22 | 0.082417 | 0.190430267 | 0 | 0 | 12.30735931 | 30.71866043 | DUSP4 |
| Intestine | Early Phase of HIV Life Cycle | 1/22 | 0.082417 | 0.190430267 | 0 | 0 | 12.30735931 | 30.71866043 | CCR5 |
| Intestine | RAF-independent MAPK1 3 Activation | 1/23 | 0.086 | 0.197481268 | 0 | 0 | 11.74734357 | 28.82103918 | DUSP4 |
| Intestine | Metabolism of Water-Soluble Vitamins and Cofactors | 2/127 | 0.087992 | 0.200720007 | 0 | 0 | 4.167789474 | 10.12985659 | SLC2A3;ALDH1L2 |
| Intestine | Growth Hormone Receptor Signaling | 1/24 | 0.089569 | 0.200720007 | 0 | 0 | 11.23602484 | 27.10972356 | SOCS3 |
| Intestine | RHO GTPases Activate NADPH Oxidases | 1/24 | 0.089569 | 0.200720007 | 0 | 0 | 11.23602484 | 27.10972356 | NCF1 |
| Intestine | Interleukin-6 Family Signaling | 1/24 | 0.089569 | 0.200720007 | 0 | 0 | 11.23602484 | 27.10972356 | SOCS3 |
| Intestine | Glycogen Metabolism | 1/25 | 0.093124 | 0.204982009 | 0 | 0 | 10.76731602 | 25.55975844 | PYGL |
| Intestine | Inflammasomes | 1/25 | 0.093124 | 0.204982009 | 0 | 0 | 10.76731602 | 25.55975844 | HMOX1 |
| Intestine | Regulation of IFNA IFNB Signaling | 1/25 | 0.093124 | 0.204982009 | 0 | 0 | 10.76731602 | 25.55975844 | SOCS3 |
| Intestine | Gluconeogenesis | 1/26 | 0.096665 | 0.210288306 | 0 | 0 | 10.3361039 | 24.15036983 | ENO2 |
| Intestine | Interleukin-20 Family Signaling | 1/26 | 0.096665 | 0.210288306 | 0 | 0 | 10.3361039 | 24.15036983 | SOCS3 |
| Intestine | Signaling by Hedgehog | 2/136 | 0.098765 | 0.212838393 | 0 | 0 | 3.886095837 | 8.996372354 | GAS1;ARRB2 |
| Intestine | RHO GTPase Cycle | 4/450 | 0.098981 | 0.212838393 | 0 | 0 | 2.360441159 | 5.459285854 | ITGB1;NCF1;ARHGEF1;ARHGEF6 |
| Intestine | Pyroptosis | 1/27 | 0.100192 | 0.214204384 | 0 | 0 | 9.938061938 | 22.86413349 | IL1A |
| Intestine | BMAL1 CLOCK,NPAS2 Activates Circadian Gene Expression | 1/28 | 0.103706 | 0.219197535 | 0 | 0 | 9.56950457 | 21.68633305 | SERPINE1 |
| Intestine | Metabolism of Porphyrins | 1/28 | 0.103706 | 0.219197535 | 0 | 0 | 9.56950457 | 21.68633305 | HMOX1 |
| Intestine | The Role of Nef in HIV-1 Replication and Disease Pathogenesis | 1/29 | 0.107207 | 0.225315988 | 0 | 0 | 9.227272727 | 20.60445933 | HCK |
| Intestine | Signaling by ALK | 1/30 | 0.110694 | 0.227503166 | 0 | 0 | 8.908643081 | 19.60781605 | CD274 |
| Intestine | PD-1 Signaling | 1/30 | 0.110694 | 0.227503166 | 0 | 0 | 8.908643081 | 19.60781605 | CD274 |
| Intestine | Laminin Interactions | 1/30 | 0.110694 | 0.227503166 | 0 | 0 | 8.908643081 | 19.60781605 | ITGB1 |
| Intestine | Regulation of CDH11 Expression and Function | 1/30 | 0.110694 | 0.227503166 | 0 | 0 | 8.908643081 | 19.60781605 | ANGPTL4 |
| Intestine | Activated NOTCH1 Transmits Signal to the Nucleus | 1/31 | 0.114167 | 0.228334487 | 0 | 0 | 8.611255411 | 18.68720662 | ARRB2 |
| Intestine | Signaling by BMP | 1/31 | 0.114167 | 0.228334487 | 0 | 0 | 8.611255411 | 18.68720662 | INHBA |
| Intestine | Signaling by CSF1 (M-CSF) in Myeloid Cells | 1/31 | 0.114167 | 0.228334487 | 0 | 0 | 8.611255411 | 18.68720662 | HCK |
| Intestine | MAPK Targets Nuclear Events Mediated by MAP Kinases | 1/31 | 0.114167 | 0.228334487 | 0 | 0 | 8.611255411 | 18.68720662 | DUSP4 |
| Intestine | Downregulation of SMAD2 3 SMAD4 Transcriptional Activity | 1/31 | 0.114167 | 0.228334487 | 0 | 0 | 8.611255411 | 18.68720662 | SERPINE1 |
| Intestine | G-protein Beta Gamma Signalling | 1/32 | 0.117627 | 0.232751989 | 0 | 0 | 8.333054043 | 17.83468317 | ARHGEF6 |
| Intestine | Thrombin Signalling Through Proteinase Activated Receptors (PARs) | 1/32 | 0.117627 | 0.232751989 | 0 | 0 | 8.333054043 | 17.83468317 | ARRB2 |
| Intestine | Regulation of Expression and Function of Type II Classical Cadherins | 1/33 | 0.121074 | 0.237050366 | 0 | 0 | 8.07224026 | 17.04334406 | ANGPTL4 |
| Intestine | Regulation of Homotypic Cell-Cell Adhesion | 1/33 | 0.121074 | 0.237050366 | 0 | 0 | 8.07224026 | 17.04334406 | ANGPTL4 |
| Intestine | Plasma Lipoprotein Remodeling | 1/34 | 0.124508 | 0.242496444 | 0 | 0 | 7.827233373 | 16.30716925 | ANGPTL4 |
| Intestine | Death Receptor Signaling | 2/157 | 0.125199 | 0.242573245 | 0 | 0 | 3.356027165 | 6.973321237 | ARHGEF1;ARHGEF6 |
| Intestine | GPVI-mediated Activation Cascade | 1/35 | 0.127928 | 0.246575887 | 0 | 0 | 7.596638655 | 15.62088558 | COL1A2 |

| | | | | | | | | | |
|---|---|---|---|---|---|---|---|---|---|
| Intestine | Signaling by High-Kinase Activity BRAF Mutants | 1/36 | 0.131335 | 0.249268186 | 0 | 0 | 7.379220779 | 14.97985584 | ARRB2 |
| Intestine | ROS and RNS Production in Phagocytes | 1/36 | 0.131335 | 0.249268186 | 0 | 0 | 7.379220779 | 14.97985584 | NCF1 |
| Intestine | Detoxification of Reactive Oxygen Species | 1/36 | 0.131335 | 0.249268186 | 0 | 0 | 7.379220779 | 14.97985584 | NCF1 |
| Intestine | Cellular Responses to Stimuli | 6/887 | 0.13206 | 0.249372287 | 0 | 0 | 1.80107832 | 3.646280347 | ITGB1;IL1A;NCF1;HMOX1;ITGA5;MAP4K4 |
| Intestine | Protein Localization | 2/164 | 0.134356 | 0.251854747 | 0 | 0 | 3.209876543 | 6.44305737 | ABCD2;HMOX1 |
| Intestine | Sphingolipid De Novo Biosynthesis | 1/37 | 0.134729 | 0.251854747 | 0 | 0 | 7.173881674 | 14.37998692 | PRKD3 |
| Intestine | FLT3 Signaling | 1/38 | 0.13811 | 0.254468901 | 0 | 0 | 6.97964198 | 13.81765323 | HCK |
| Intestine | Defective B3GALTL Causes PpS | 1/38 | 0.13811 | 0.254468901 | 0 | 0 | 6.97964198 | 13.81765323 | ADAMTS6 |
| Intestine | Cellular Senescence | 2/167 | 0.138327 | 0.254468901 | 0 | 0 | 3.151036683 | 6.233181779 | IL1A;MAP4K4 |
| Intestine | Metabolism | 12/2181 | 0.138863 | 0.254468901 | 0 | 0 | 1.488159604 | 2.938020713 | ABCA1;ARG1;PRKD3;ALOX5;RBP1;ALOX15;HMOX1;SLC2A3;PYGL;ANGPTL4;ENO2;ALDH1L2 |
| Intestine | Class A 1 (Rhodopsin-like Receptors) | 3/333 | 0.140896 | 0.25548332 | 0 | 0 | 2.374787879 | 4.653946788 | CCL7;CCR5;CCR3 |
| Intestine | O-glycosylation of TSR Domain-Containing Proteins | 1/39 | 0.141477 | 0.25548332 | 0 | 0 | 6.795625427 | 13.28963273 | ADAMTS6 |
| Intestine | Platelet Aggregation (Plug Formation) | 1/39 | 0.141477 | 0.25548332 | 0 | 0 | 6.795625427 | 13.28963273 | COL1A2 |
| Intestine | RHOU GTPase Cycle | 1/40 | 0.144832 | 0.259026636 | 0 | 0 | 6.621045621 | 12.79305295 | ARHGEF6 |
| Intestine | MAP2K and MAPK Activation | 1/40 | 0.144832 | 0.259026636 | 0 | 0 | 6.621045621 | 12.79305295 | ARRB2 |
| Intestine | Generation of Second Messenger Molecules | 1/41 | 0.148174 | 0.261235557 | 0 | 0 | 6.455194805 | 12.32534552 | ENAH |
| Intestine | Signaling by RAF1 Mutants | 1/41 | 0.148174 | 0.261235557 | 0 | 0 | 6.455194805 | 12.32534552 | ARRB2 |
| Intestine | Signaling by Retinoic Acid | 1/41 | 0.148174 | 0.261235557 | 0 | 0 | 6.455194805 | 12.32534552 | ALDH1A2 |
| Intestine | Fatty Acid Metabolism | 2/176 | 0.150389 | 0.263890031 | 0 | 0 | 2.986690865 | 5.658376619 | ALOX5;ALOX15 |
| Intestine | NCAM1 Interactions | 1/42 | 0.151503 | 0.264596566 | 0 | 0 | 6.297434273 | 11.88420746 | COL6A1 |
| Intestine | Negative Regulation of MAPK Pathway | 1/43 | 0.154819 | 0.269124565 | 0 | 0 | 6.147186147 | 11.46756826 | DUSP4 |
| Intestine | Retinoid Metabolism and Transport | 1/44 | 0.158122 | 0.271699426 | 0 | 0 | 6.003926306 | 11.07356161 | RBP1 |
| Intestine | NR1H3 & NR1H2 Regulate Gene Expression Linked to Cholesterol Transport and Efflux | 1/44 | 0.158122 | 0.271699426 | 0 | 0 | 6.003926306 | 11.07356161 | ABCA1 |
| Intestine | Axon Guidance | 4/541 | 0.160545 | 0.271699426 | 0 | 0 | 1.95128089 | 3.569252001 | ITGB1;ENAH;COL6A1;ITGA5 |
| Intestine | Paradoxical Activation of RAF Signaling by Kinase Inactive BRAF | 1/45 | 0.161413 | 0.271699426 | 0 | 0 | 5.867178276 | 10.70050124 | ARRB2 |
| Intestine | Signaling by RAS Mutants | 1/45 | 0.161413 | 0.271699426 | 0 | 0 | 5.867178276 | 10.70050124 | ARRB2 |
| Intestine | Signaling by Moderate Kinase Activity BRAF Mutants | 1/45 | 0.161413 | 0.271699426 | 0 | 0 | 5.867178276 | 10.70050124 | ARRB2 |
| Intestine | Signaling Downstream of RAS Mutants | 1/45 | 0.161413 | 0.271699426 | 0 | 0 | 5.867178276 | 10.70050124 | ARRB2 |
| Intestine | Kidney Development | 1/46 | 0.164691 | 0.275968185 | 0 | 0 | 5.736507937 | 10.34685999 | ITGB1 |
| Intestine | Heme Signaling | 1/47 | 0.167956 | 0.278926767 | 0 | 0 | 5.611518916 | 10.01125174 | HMOX1 |
| Intestine | TGF-beta Receptor Signaling Activates SMADs | 1/47 | 0.167956 | 0.278926767 | 0 | 0 | 5.611518916 | 10.01125174 | ITGB1 |
| Intestine | Ub-specific Processing Proteases | 2/190 | 0.169547 | 0.280316958 | 0 | 0 | 2.762318029 | 4.902086303 | IL33;ARRB2 |
| Intestine | Interleukin-3, Interleukin-5 and GM-CSF Signaling | 1/48 | 0.171209 | 0.28057078 | 0 | 0 | 5.491848577 | 9.692415749 | HCK |
| Intestine | Metabolism of Fat-Soluble Vitamins | 1/48 | 0.171209 | 0.28057078 | 0 | 0 | 5.491848577 | 9.692415749 | RBP1 |
| Intestine | Metabolism of Lipids | 5/758 | 0.173689 | 0.283387139 | 0 | 0 | 1.743619131 | 3.052187086 | ABCA1;PRKD3;ALOX5;ALOX15;ANGPTL4 |
| Intestine | Nervous System Development | 4/567 | 0.180285 | 0.292864464 | 0 | 0 | 1.858672171 | 3.184309147 | ITGB1;ENAH;COL6A1;ITGA5 |
| Intestine | IRS-related Events Triggered by IGF1R | 1/52 | 0.184094 | 0.297751886 | 0 | 0 | 5.060096766 | 8.563248721 | IGF1 |
| Intestine | Cellular Response to Chemical Stress | 2/202 | 0.186281 | 0.299984922 | 0 | 0 | 2.595 | 4.360895448 | NCF1;HMOX1 |
| Intestine | IGF1R Signaling Cascade | 1/53 | 0.187284 | 0.300300579 | 0 | 0 | 4.962537463 | 8.31288475 | IGF1 |
| Intestine | NR1H2 and NR1H3-mediated Signaling | 1/54 | 0.190462 | 0.302786126 | 0 | 0 | 4.868659642 | 8.073704687 | ABCA1 |
| Intestine | Signaling by Type 1 Insulin-like Growth Factor 1 Receptor (IGF1R) | 1/54 | 0.190462 | 0.302786126 | 0 | 0 | 4.868659642 | 8.073704687 | IGF1 |
| Intestine | Signaling by Non-Receptor Tyrosine Kinases | 1/55 | 0.193628 | 0.305210207 | 0 | 0 | 4.778258778 | 7.845024507 | SOCS3 |
| Intestine | Signaling by PTK6 | 1/55 | 0.193628 | 0.305210207 | 0 | 0 | 4.778258778 | 7.845024507 | SOCS3 |
| Intestine | Signaling by PDGF | 1/58 | 0.203052 | 0.318714622 | 0 | 0 | 4.526087947 | 7.215909839 | COL6A1 |
| Intestine | Disease | 11/2131 | 0.205266 | 0.319562244 | 0 | 0 | 1.37863982 | 2.183001952 | ITGB1;ABCA1;IL1A;HCK;COL1A2;SERPINE1;RBP1;HMOX1;ARRB2;CCR5;ADAMTS6 |
| Intestine | Cytoprotection by HMOX1 | 1/59 | 0.206169 | 0.319562244 | 0 | 0 | 4.447828034 | 7.02337905 | HMOX1 |
| Intestine | Iron Uptake and Transport | 1/59 | 0.206169 | 0.319562244 | 0 | 0 | 4.447828034 | 7.02337905 | HMOX1 |
| Intestine | Nuclear Events (Kinase and Transcription Factor Activation) | 1/61 | 0.212367 | 0.325105635 | 0 | 0 | 4.299134199 | 6.661239871 | DUSP4 |
| Intestine | Nucleotide-binding Domain, Leucine Rich Repeat Containing Receptor (NLR) Signaling Pathways | 1/61 | 0.212367 | 0.325105635 | 0 | 0 | 4.299134199 | 6.661239871 | HMOX1 |
| Intestine | Regulated Necrosis | 1/61 | 0.212367 | 0.325105635 | 0 | 0 | 4.299134199 | 6.661239871 | IL1A |

| | | | | | | | | | |
|---|---|---|---|---|---|---|---|---|---|
| Intestine | NCAM Signaling for Neurite Out-Growth | 1/63 | 0.218518 | 0.330441569 | 0 | 0 | 4.160033515 | 6.326944006 | COL6A1 |
| Intestine | Transcriptional Regulation by MECP2 | 1/63 | 0.218518 | 0.330441569 | 0 | 0 | 4.160033515 | 6.326944006 | SLC2A3 |
| Intestine | MAP Kinase Activation | 1/63 | 0.218518 | 0.330441569 | 0 | 0 | 4.160033515 | 6.326944006 | DUSP4 |
| Intestine | Semaphorin Interactions | 1/64 | 0.221575 | 0.332711169 | 0 | 0 | 4.093795094 | 6.169321198 | ITGB1 |
| Intestine | HIV Infection | 2/227 | 0.221807 | 0.332711169 | 0 | 0 | 2.30374269 | 3.469311246 | HCK;CCR5 |
| Intestine | ABC Transporter Disorders | 1/65 | 0.224621 | 0.334235776 | 0 | 0 | 4.029626623 | 6.017608891 | ABCA1 |
| Intestine | Signaling by BRAF and RAF1 Fusions | 1/65 | 0.224621 | 0.334235776 | 0 | 0 | 4.029626623 | 6.017608891 | ARRB2 |
| Intestine | Circadian Clock | 1/70 | 0.239673 | 0.353803363 | 0 | 0 | 3.736683606 | 5.337773161 | SERPINE1 |
| Intestine | RHOB GTPase Cycle | 1/70 | 0.239673 | 0.353803363 | 0 | 0 | 3.736683606 | 5.337773161 | ARHGEF1 |
| Intestine | Diseases Associated With O-glycosylation of Proteins | 1/71 | 0.242649 | 0.356780246 | 0 | 0 | 3.683116883 | 5.215807707 | ADAMTS6 |
| Intestine | Interleukin-17 Signaling | 1/72 | 0.245613 | 0.359716851 | 0 | 0 | 3.631059082 | 5.097998493 | DUSP4 |
| Intestine | Signaling by NOTCH1 | 1/74 | 0.251507 | 0.362638256 | 0 | 0 | 3.531222202 | 4.874088629 | ARRB2 |
| Intestine | Glycolysis | 1/74 | 0.251507 | 0.362638256 | 0 | 0 | 3.531222202 | 4.874088629 | ENO2 |
| Intestine | RHOC GTPase Cycle | 1/74 | 0.251507 | 0.362638256 | 0 | 0 | 3.531222202 | 4.874088629 | ARHGEF1 |
| Intestine | RHOG GTPase Cycle | 1/74 | 0.251507 | 0.362638256 | 0 | 0 | 3.531222202 | 4.874088629 | ITGB1 |
| Intestine | SLC-mediated Transmembrane Transport | 2/249 | 0.253508 | 0.36411206 | 0 | 0 | 2.096207117 | 2.876749528 | SLC2A3;SLC16A3 |
| Intestine | Interferon Alpha Beta Signaling | 1/78 | 0.263159 | 0.375077548 | 0 | 0 | 3.347107438 | 4.468374796 | SOCS3 |
| Intestine | Costimulation by the CD28 Family | 1/78 | 0.263159 | 0.375077548 | 0 | 0 | 3.347107438 | 4.468374796 | CD274 |
| Intestine | PCP CE Pathway | 1/79 | 0.266044 | 0.377517051 | 0 | 0 | 3.304029304 | 4.374841806 | ARRB2 |
| Intestine | Signaling by Rho GTPases | 4/672 | 0.2669 | 0.377517051 | 0 | 0 | 1.558019097 | 2.057955274 | ITGB1;NCF1;ARHGEF1;ARHGEF6 |
| Intestine | Senescence-Associated Secretory Phenotype (SASP) | 1/81 | 0.271781 | 0.382963593 | 0 | 0 | 3.221103896 | 4.196325643 | IL1A |
| Intestine | GPCR Ligand Binding | 3/467 | 0.274397 | 0.38407212 | 0 | 0 | 1.677413793 | 2.169195693 | CCL7;CCR5;CCR3 |
| Intestine | Oncogenic MAPK Signaling | 1/82 | 0.274632 | 0.38407212 | 0 | 0 | 3.181176848 | 4.111106345 | ARRB2 |
| Intestine | RAF MAP Kinase Cascade | 2/265 | 0.276653 | 0.385449812 | 0 | 0 | 1.967080248 | 2.527677425 | DUSP4;ARRB2 |
| Intestine | Transcriptional Regulation of White Adipocyte Differentiation | 1/84 | 0.280302 | 0.38571694 | 0 | 0 | 3.104209044 | 3.948200984 | ANGPTL4 |
| Intestine | Transport of Bile Salts and Organic Acids, Metal Ions and Amine Compounds | 1/84 | 0.280302 | 0.38571694 | 0 | 0 | 3.104209044 | 3.948200984 | SLC16A3 |
| Intestine | Signaling by Rho GTPases, Miro GTPases and RHOBTB3 | 4/688 | 0.280797 | 0.38571694 | 0 | 0 | 1.520309783 | 1.93098145 | ITGB1;NCF1;ARHGEF1;ARHGEF6 |
| Intestine | Deubiquitination | 2/268 | 0.280993 | 0.38571694 | 0 | 0 | 1.944598338 | 2.468524717 | IL33;ARRB2 |
| Intestine | Glucose Metabolism | 1/85 | 0.283121 | 0.386831891 | 0 | 0 | 3.067099567 | 3.870313956 | ENO2 |
| Intestine | Signaling by Nuclear Receptors | 2/270 | 0.283885 | 0.386831891 | 0 | 0 | 1.929890024 | 2.430092681 | ABCA1;ALDH1A2 |
| Intestine | MAPK1 MAPK3 Signaling | 2/271 | 0.28533 | 0.387382838 | 0 | 0 | 1.922617883 | 2.411169657 | DUSP4;ARRB2 |
| Intestine | Nuclear Events Mediated by NFE2L2 | 1/87 | 0.288726 | 0.390567125 | 0 | 0 | 2.995469647 | 3.721206598 | HMOX1 |
| Intestine | Developmental Biology | 7/1385 | 0.294878 | 0.39744367 | 0 | 0 | 1.326764652 | 1.620238432 | ITGB1;ENAH;MEIS1;COL6A1;ANGPTL4;ITGA5;MSX1 |
| Intestine | ABC-family Proteins Mediated Transport | 1/90 | 0.297052 | 0.398928599 | 0 | 0 | 2.894060995 | 3.512953554 | ABCD2 |
| Intestine | Activation of HOX Genes During Differentiation | 1/91 | 0.299806 | 0.399740745 | 0 | 0 | 2.861760462 | 3.447337186 | MEIS1 |
| Intestine | Activation of Anterior HOX Genes in Hindbrain Development During Early Embryogenesis | 1/91 | 0.299806 | 0.399740745 | 0 | 0 | 2.861760462 | 3.447337186 | MEIS1 |
| Intestine | FCGR Activation | 1/94 | 0.308003 | 0.407748373 | 0 | 0 | 2.769026672 | 3.260928233 | HCK |
| Intestine | Oxidative Stress Induced Senescence | 1/94 | 0.308003 | 0.407748373 | 0 | 0 | 2.769026672 | 3.260928233 | MAP4K4 |
| Intestine | Metabolism of Carbohydrates | 2/290 | 0.312732 | 0.411516989 | 0 | 0 | 1.794042398 | 2.085413178 | PYGL;ENO2 |
| Intestine | Antigen processing-Cross Presentation | 1/96 | 0.313416 | 0.411516989 | 0 | 0 | 2.710457963 | 3.144737764 | NCF1 |
| Intestine | RHO GTPase Effectors | 2/291 | 0.314169 | 0.411516989 | 0 | 0 | 1.78774358 | 2.06989348 | ITGB1;NCF1 |
| Intestine | MyD88 Cascade Initiated on Plasma Membrane | 1/98 | 0.318787 | 0.412351244 | 0 | 0 | 2.654304458 | 3.034487705 | DUSP4 |
| Intestine | Toll Like Receptor 10 (TLR10) Cascade | 1/98 | 0.318787 | 0.412351244 | 0 | 0 | 2.654304458 | 3.034487705 | DUSP4 |
| Intestine | Toll Like Receptor 5 (TLR5) Cascade | 1/98 | 0.318787 | 0.412351244 | 0 | 0 | 2.654304458 | 3.034487705 | DUSP4 |
| Intestine | Interferon Gamma Signaling | 1/99 | 0.321457 | 0.412351244 | 0 | 0 | 2.627087199 | 2.981462057 | SOCS3 |
| Intestine | VEGFA-VEGFR2 Pathway | 1/99 | 0.321457 | 0.412351244 | 0 | 0 | 2.627087199 | 2.981462057 | NCF1 |
| Intestine | Visual Phototransduction | 1/99 | 0.321457 | 0.412351244 | 0 | 0 | 2.627087199 | 2.981462057 | RBP1 |
| Intestine | Interleukin-1 Signaling | 1/102 | 0.329404 | 0.417284649 | 0 | 0 | 2.548669153 | 2.830219734 | IL1A |
| Intestine | TRAF6 Mediated Induction of NFkB and MAP Kinases Upon TLR7 8 or 9 Activation | 1/102 | 0.329404 | 0.417284649 | 0 | 0 | 2.548669153 | 2.830219734 | DUSP4 |

| | | | | | | | | | |
|---|---|---|---|---|---|---|---|---|---|
| Intestine | Epigenetic Regulation of Adipogenesis Genes by MLL3 and MLL4 Complexes | 1/103 | 0.332033 | 0.417284649 | 0 | 0 | 2.523554876 | 2.782272441 | ANGPTL4 |
| Intestine | Epigenetic Regulation of Gene Expression by MLL3 and MLL4 Complexes | 1/103 | 0.332033 | 0.417284649 | 0 | 0 | 2.523554876 | 2.782272441 | ANGPTL4 |
| Intestine | MyD88 Dependent Cascade Initiated on Endosome | 1/103 | 0.332033 | 0.417284649 | 0 | 0 | 2.523554876 | 2.782272441 | DUSP4 |
| Intestine | MLL4 and MLL3 Complexes Regulate Expression of PPARG Targets in Adipogenesis and Hepatic Steatosis | 1/103 | 0.332033 | 0.417284649 | 0 | 0 | 2.523554876 | 2.782272441 | ANGPTL4 |
| Intestine | Cargo Recognition for Clathrin-Mediated Endocytosis | 1/105 | 0.33726 | 0.422426564 | 0 | 0 | 2.474775225 | 2.689836595 | ARRB2 |
| Intestine | Toll Like Receptor 3 (TLR3) Cascade | 1/106 | 0.339858 | 0.422833665 | 0 | 0 | 2.451082251 | 2.645273372 | DUSP4 |
| Intestine | Transcriptional Regulation by RUNX2 | 1/106 | 0.339858 | 0.422833665 | 0 | 0 | 2.451082251 | 2.645273372 | ITGA5 |
| Intestine | Sphingolipid Metabolism | 1/107 | 0.342447 | 0.422910071 | 0 | 0 | 2.427836315 | 2.601766058 | PRKD3 |
| Intestine | Toll Like Receptor 7 8 (TLR7 8) Cascade | 1/107 | 0.342447 | 0.422910071 | 0 | 0 | 2.427836315 | 2.601766058 | DUSP4 |
| Intestine | Signaling by VEGF | 1/108 | 0.345025 | 0.422910071 | 0 | 0 | 2.405024882 | 2.559281033 | NCF1 |
| Intestine | MAPK Family Signaling Cascades | 2/314 | 0.347028 | 0.422910071 | 0 | 0 | 1.654014845 | 1.750525922 | DUSP4;ARRB2 |
| Intestine | G Alpha (I) Signalling Events | 2/316 | 0.349864 | 0.422910071 | 0 | 0 | 1.643312102 | 1.725824412 | CCR5;CCR3 |
| Intestine | MyD88-independent TLR4 Cascade | 1/110 | 0.350151 | 0.422910071 | 0 | 0 | 2.360657691 | 2.477250081 | DUSP4 |
| Intestine | Adipogenesis | 1/110 | 0.350151 | 0.422910071 | 0 | 0 | 2.360657691 | 2.477250081 | ANGPTL4 |
| Intestine | TRIF (TICAM1)-mediated TLR4 Signaling | 1/110 | 0.350151 | 0.422910071 | 0 | 0 | 2.360657691 | 2.477250081 | DUSP4 |
| Intestine | Toll Like Receptor 9 (TLR9) Cascade | 1/110 | 0.350151 | 0.422910071 | 0 | 0 | 2.360657691 | 2.477250081 | DUSP4 |
| Intestine | PI5P, PP2A and IER3 Regulate PI3K AKT Signaling | 1/111 | 0.3527 | 0.424609433 | 0 | 0 | 2.339079103 | 2.437643429 | IL33 |
| Intestine | SARS-CoV-1-host Interactions | 1/112 | 0.355238 | 0.426285992 | 0 | 0 | 2.317889318 | 2.398937497 | SERPINE1 |
| Intestine | TCR Signaling | 1/113 | 0.357767 | 0.427940013 | 0 | 0 | 2.297077922 | 2.361104815 | ENAH |
| Intestine | O-linked Glycosylation | 1/114 | 0.360286 | 0.428443823 | 0 | 0 | 2.27663487 | 2.324118975 | ADAMTS6 |
| Intestine | MyD88 MAL(TIRAP) Cascade Initiated on Plasma Membrane | 1/115 | 0.362795 | 0.428443823 | 0 | 0 | 2.256550467 | 2.287954578 | DUSP4 |
| Intestine | Signaling by NTRK1 (TRKA) | 1/115 | 0.362795 | 0.428443823 | 0 | 0 | 2.256550467 | 2.287954578 | DUSP4 |
| Intestine | Toll Like Receptor TLR6 TLR2 Cascade | 1/115 | 0.362795 | 0.428443823 | 0 | 0 | 2.256550467 | 2.287954578 | DUSP4 |
| Intestine | Cellular Responses to Stress | 4/787 | 0.368587 | 0.431781883 | 0 | 0 | 1.321252287 | 1.318712271 | IL1A;NCF1;HMOX1;MAP4K4 |
| Intestine | Negative Regulation of the PI3K AKT Network | 1/118 | 0.370265 | 0.431781883 | 0 | 0 | 2.198357198 | 2.184150248 | IL33 |
| Intestine | Toll Like Receptor 2 (TLR2) Cascade | 1/118 | 0.370265 | 0.431781883 | 0 | 0 | 2.198357198 | 2.184150248 | DUSP4 |
| Intestine | Toll Like Receptor TLR1 TLR2 Cascade | 1/118 | 0.370265 | 0.431781883 | 0 | 0 | 2.198357198 | 2.184150248 | DUSP4 |
| Intestine | KEAP1-NFE2L2 Pathway | 1/119 | 0.372735 | 0.433304568 | 0 | 0 | 2.179616993 | 2.151036258 | HMOX1 |
| Intestine | Gastrulation | 1/120 | 0.375196 | 0.434806676 | 0 | 0 | 2.161191749 | 2.118630321 | MSX1 |
| Intestine | FCGR3A-mediated IL10 Synthesis | 1/121 | 0.377648 | 0.436288443 | 0 | 0 | 2.143073593 | 2.0869122 | HCK |
| Intestine | Binding and Uptake of Ligands by Scavenger Receptors | 1/122 | 0.380089 | 0.437750098 | 0 | 0 | 2.12525491 | 2.055862392 | COL1A2 |
| Intestine | Host Interactions of HIV Factors | 1/126 | 0.389763 | 0.447505681 | 0 | 0 | 2.056831169 | 1.937980027 | HCK |
| Intestine | Infectious Disease | 6/1311 | 0.404473 | 0.462966151 | 0 | 0 | 1.188825032 | 1.076088764 | ITGB1;IL1A;HCK;SERPINE1;HMOX1;CCR5 |
| Intestine | Beta-catenin Independent WNT Signaling | 1/134 | 0.408665 | 0.46490356 | 0 | 0 | 1.932330827 | 1.729163584 | ARRB2 |
| Intestine | Signaling by NTRKs | 1/134 | 0.408665 | 0.46490356 | 0 | 0 | 1.932330827 | 1.729163584 | DUSP4 |
| Intestine | Epigenetic Regulation by WDR5-containing Histone Modifying Complexes | 1/135 | 0.410987 | 0.466119384 | 0 | 0 | 1.91781353 | 1.705307776 | ANGPTL4 |
| Intestine | RHO GTPases Activate Formins | 1/139 | 0.420184 | 0.475102073 | 0 | 0 | 1.861848297 | 1.614337371 | ITGB1 |
| Intestine | FCGR3A-mediated Phagocytosis | 1/142 | 0.426989 | 0.478433699 | 0 | 0 | 1.821958184 | 1.550480079 | HCK |
| Intestine | Parasite Infection | 1/142 | 0.426989 | 0.478433699 | 0 | 0 | 1.821958184 | 1.550480079 | HCK |
| Intestine | Leishmania Phagocytosis | 1/142 | 0.426989 | 0.478433699 | 0 | 0 | 1.821958184 | 1.550480079 | HCK |
| Intestine | Class I MHC Mediated Antigen Processing & Presentation | 2/373 | 0.428644 | 0.478845305 | 0 | 0 | 1.386792453 | 1.174792176 | SOCS3;NCF1 |
| Intestine | Clathrin-mediated Endocytosis | 1/145 | 0.433715 | 0.483060064 | 0 | 0 | 1.783730159 | 1.490069557 | ARRB2 |
| Intestine | Diseases of Glycosylation | 1/146 | 0.43594 | 0.484088435 | 0 | 0 | 1.771339006 | 1.470655972 | ADAMTS6 |
| Intestine | RHOA GTPase Cycle | 1/149 | 0.442562 | 0.489979494 | 0 | 0 | 1.735170235 | 1.414466412 | ARHGEF1 |
| Intestine | CDC42 GTPase Cycle | 1/155 | 0.455577 | 0.502892397 | 0 | 0 | 1.667060213 | 1.310626032 | ARHGEF6 |
| Intestine | SARS-CoV-1 Infection | 1/157 | 0.459849 | 0.506105565 | 0 | 0 | 1.645521146 | 1.278336206 | SERPINE1 |
| Intestine | HIV Life Cycle | 1/158 | 0.461972 | 0.506276588 | 0 | 0 | 1.634957399 | 1.262597981 | CCR5 |
| Intestine | Anti-inflammatory Response Favouring Leishmania Parasite Infection | 1/159 | 0.464087 | 0.506276588 | 0 | 0 | 1.624527371 | 1.24712289 | HCK |

| | | | | | | | | | |
|---|---|---|---|---|---|---|---|---|---|
| Intestine | Leishmania Parasite Growth and Survival | 1/159 | 0.464087 | 0.506276588 | 0 | 0 | 1.624527371 | 1.24712289 | HCK |
| Intestine | Potential Therapeutics for SARS | 1/164 | 0.474539 | 0.514660859 | 0 | 0 | 1.574296869 | 1.173497462 | ITGB1 |
| Intestine | Disorders of Transmembrane Transporters | 1/164 | 0.474539 | 0.514660859 | 0 | 0 | 1.574296869 | 1.173497462 | ABCA1 |
| Intestine | Fcgamma Receptor (FCGR) Dependent Phagocytosis | 1/168 | 0.482756 | 0.522050507 | 0 | 0 | 1.536278093 | 1.118783977 | HCK |
| Intestine | Post-translational Protein Modification | 6/1457 | 0.50659 | 0.546236434 | 0 | 0 | 1.060820124 | 0.721413701 | SOCS3;IL33;DMP1;ARRB2;TIMP1;ADAMTS6 |
| Intestine | Generic Transcription Pathway | 5/1230 | 0.52808 | 0.567761907 | 0 | 0 | 1.045401174 | 0.667497228 | SOCS3;COL1A2;SERPINE1;SLC2A3;ITGA5 |
| Intestine | Programmed Cell Death | 1/202 | 0.547668 | 0.58543863 | 0 | 0 | 1.274213349 | 0.767185117 | IL1A |
| Intestine | Transcriptional Regulation by RUNX1 | 1/202 | 0.547668 | 0.58543863 | 0 | 0 | 1.274213349 | 0.767185117 | SOCS3 |
| Intestine | HCMV Early Events | 1/205 | 0.552994 | 0.589437614 | 0 | 0 | 1.255283932 | 0.743640651 | ITGB1 |
| Intestine | Signaling by NOTCH | 1/209 | 0.559998 | 0.595198275 | 0 | 0 | 1.230894106 | 0.713698735 | ARRB2 |
| Intestine | Signaling by ROBO Receptors | 1/210 | 0.561733 | 0.595340428 | 0 | 0 | 1.224942522 | 0.706460513 | ENAH |
| Intestine | Metabolism of Proteins | 8/2067 | 0.563914 | 0.595954174 | 0 | 0 | 0.99149379 | 0.567981366 | SOCS3;IL33;DMP1;ARRB2;TIMP1;INHBA;IGF1;ADAMTS6 |
| Intestine | Viral Infection Pathways | 4/1029 | 0.574681 | 0.60561312 | 0 | 0 | 0.996545814 | 0.552026299 | ITGB1;HCK;SERPINE1;CCR5 |
| Intestine | SARS-CoV Infections | 2/498 | 0.581936 | 0.611526443 | 0 | 0 | 1.030666384 | 0.55799662 | ITGB1;SERPINE1 |
| Intestine | Neddylation | 1/234 | 0.601386 | 0.630184414 | 0 | 0 | 1.097430467 | 0.558064075 | SOCS3 |
| Intestine | Gene Expression (Transcription) | 6/1615 | 0.609725 | 0.637128122 | 0 | 0 | 0.948466957 | 0.469251783 | SOCS3;COL1A2;SERPINE1;SLC2A3;ANGPTL4;ITGA5 |
| Intestine | RNA Polymerase II Transcription | 5/1360 | 0.619101 | 0.645114146 | 0 | 0 | 0.938533084 | 0.450013549 | SOCS3;COL1A2;SERPINE1;SLC2A3;ITGA5 |
| Intestine | HCMV Infection | 1/259 | 0.638925 | 0.663911114 | 0 | 0 | 0.989831874 | 0.443412877 | ITGB1 |
| Intestine | Diseases of Metabolism | 1/264 | 0.646003 | 0.669395954 | 0 | 0 | 0.970766876 | 0.424177557 | ADAMTS6 |
| Intestine | Transmission Across Chemical Synapses | 1/269 | 0.652944 | 0.674708758 | 0 | 0 | 0.952413258 | 0.40597946 | CACNA2D1 |
| Intestine | Interferon Signaling | 1/280 | 0.667744 | 0.68754095 | 0 | 0 | 0.914350882 | 0.369260328 | SOCS3 |
| Intestine | PIP3 Activates AKT Signaling | 1/281 | 0.669059 | 0.68754095 | 0 | 0 | 0.911038961 | 0.366131553 | IL33 |
| Intestine | Signaling by WNT | 1/287 | 0.676837 | 0.693617805 | 0 | 0 | 0.891653801 | 0.348034951 | ARRB2 |
| Intestine | Epigenetic Regulation of Gene Expression | 1/291 | 0.681922 | 0.696908846 | 0 | 0 | 0.879175996 | 0.336584304 | ANGPTL4 |
| Intestine | Antigen Processing Ubiquitination & Proteasome Degradation | 1/296 | 0.688167 | 0.701364364 | 0 | 0 | 0.86405459 | 0.322918153 | SOCS3 |
| Intestine | Intracellular Signaling by Second Messengers | 1/321 | 0.717622 | 0.729385965 | 0 | 0 | 0.795535714 | 0.263968904 | IL33 |
| Intestine | Metabolism of Amino Acids and Derivatives | 1/358 | 0.756244 | 0.766546856 | 0 | 0 | 0.711739241 | 0.198853865 | ARG1 |
| Intestine | Vesicle-mediated Transport | 2/752 | 0.796918 | 0.805579801 | 0 | 0 | 0.672701754 | 0.15270594 | COL1A2;ARRB2 |
| Intestine | Neuronal System | 1/411 | 0.802645 | 0.809170711 | 0 | 0 | 0.618055116 | 0.135874831 | CACNA2D1 |
| Intestine | Diseases of Signal Transduction by Growth Factor Receptors and Second Messengers | 1/451 | 0.831783 | 0.836279187 | 0 | 0 | 0.561962482 | 0.103504281 | ARRB2 |
| Intestine | Membrane Trafficking | 1/633 | 0.919017 | 0.921272771 | 0 | 0 | 0.396391583 | 0.033475625 | ARRB2 |
| Intestine | Sensory Perception | 1/640 | 0.921273 | 0.921272771 | 0 | 0 | 0.391906998 | 0.032136028 | RBP1 |
| Liver | Extracellular Matrix Organization | 11/300 | 1.55E-09 | 2.95E-07 | 0 | 0 | 14.95847751 | 303.462555 | DDR1;MMP12;VCAN;COL1A2;COL12A1;TNC;ITGAX;ITGB8;MMP9;LOXL1;CD44 |
| Liver | Interleukin-4 and Interleukin-13 Signaling | 8/112 | 1.73E-09 | 2.95E-07 | 0 | 0 | 28.78809869 | 580.8493753 | CCL11;COL1A2;ALOX5;ALOX15;ITGAX;F13A1;IL2RG;MMP9 |
| Liver | Chemokine Receptors Bind Chemokines | 6/57 | 2.01E-08 | 2.29E-06 | 0 | 0 | 42.54117647 | 753.9187608 | CCR1;CCL11;CCL7;CXCR4;ACKR3;CX3CL1 |
| Liver | Cytokine Signaling in Immune System | 14/776 | 6.45E-08 | 5.51E-06 | 0 | 0 | 7.496453901 | 124.1184246 | CCR1;CIITA;IL1RN;CCL11;SPHK1;TNFRSF9;ALOX15;F13A1;IL2RG;MMP9;COL1A2;ALOX5;ITGAX;CD44 |
| Liver | Signaling by Interleukins | 10/452 | 1.01E-06 | 6.92E-05 | 0 | 0 | 8.649188182 | 119.3865121 | CCR1;IL1RN;CCL11;COL1A2;ALOX5;ALOX15;ITGAX;F13A1;IL2RG;MMP9 |
| Liver | Signal Transduction | 22/2613 | 4.31E-06 | 2.46E-04 | 0 | 0 | 3.7769399 | 46.66122835 | CCR1;PTGIR;CCL11;FZD2;EGR3;GPR35;SPHK1;FST;BUB1B;CXCR4;IL2RG;MMP9;CX3CL1;DNM1;EREG;ALDH1A3;COL1A 2;CCL7;PRKAR2B;ALDH1A2;ITGB8;ACKR3 |
| Liver | Integrin Cell Surface Interactions | 5/85 | 6.07E-06 | 2.97E-04 | 0 | 0 | 22.1640625 | 266.2240601 | COL1A2;TNC;ITGAX;ITGB8;CD44 |
| Liver | Class A 1 (Rhodopsin-like Receptors) | 8/333 | 7.42E-06 | 3.17E-04 | 0 | 0 | 9.109550073 | 107.5958592 | CCR1;PTGIR;CCL11;CCL7;GPR35;CXCR4;ACKR3;CX3CL1 |
| Liver | GPCR Ligand Binding | 9/467 | 1.13E-05 | 4.29E-04 | 0 | 0 | 7.361815586 | 83.86772887 | CCR1;PTGIR;CCL11;FZD2;CCL7;GPR35;CXCR4;ACKR3;CX3CL1 |
| Liver | Peptide Ligand-Binding Receptors | 6/198 | 3.08E-05 | 0.001053706 | 0 | 0 | 11.21988636 | 116.548444 | CCR1;CCL11;CCL7;CXCR4;ACKR3;CX3CL1 |
| Liver | Assembly of Collagen Fibrils and Other Multimeric Structures | 4/61 | 3.59E-05 | 0.001115584 | 0 | 0 | 24.47768544 | 250.5362714 | COL1A2;COL12A1;MMP9;LOXL1 |
| Liver | Collagen Degradation | 4/64 | 4.34E-05 | 0.001218994 | 0 | 0 | 23.2502924 | 233.5559463 | MMP12;COL1A2;COL12A1;MMP9 |
| Liver | Immune System | 18/2150 | 4.78E-05 | 0.001218994 | 0 | 0 | 3.496291287 | 34.78143327 | CCR1;CIITA;IL1RN;CCL11;SPHK1;TNFRSF9;ALOX15;F13A1;MYO5A;PDCD1LG2;IL2RG;MMP9;DNM1;ENAH;COL1A2;ALO X5;ITGAX;CD44 |
| Liver | Signaling by GPCR | 10/706 | 4.99E-05 | 0.001218994 | 0 | 0 | 5.421174217 | 53.69934719 | CCR1;PTGIR;CCL11;FZD2;CCL7;GPR35;PRKAR2B;CXCR4;ACKR3;CX3CL1 |
| Liver | Degradation of the Extracellular Matrix | 5/140 | 6.79E-05 | 0.001548671 | 0 | 0 | 13.0978836 | 125.701942 | MMP12;COL1A2;COL12A1;MMP9;CD44 |
| Liver | ECM Proteoglycans | 4/76 | 8.53E-05 | 0.00182233 | 0 | 0 | 19.36354776 | 181.4337586 | VCAN;COL1A2;TNC;ITGAX |
| Liver | Biosynthesis of E-series 18(S)-resolvins | 2/5 | 9.10E-05 | 0.001829942 | 0 | 0 | 225.2655367 | 2096.111197 | ALOX5;ALOX15 |
| Liver | Biosynthesis of EPA-derived SPMs | 2/6 | 1.36E-04 | 0.002587331 | 0 | 0 | 168.940678 | 1503.836809 | ALOX5;ALOX15 |
| Liver | Collagen Formation | 4/90 | 1.64E-04 | 0.002960607 | 0 | 0 | 16.1999184 | 141.1455611 | COL1A2;COL12A1;MMP9;LOXL1 |
| Liver | Non-integrin membrane-ECM Interactions | 3/59 | 7.77E-04 | 0.013288938 | 0 | 0 | 18.3648399 | 131.4904503 | DDR1;COL1A2;TNC |
| Liver | Signaling by Receptor Tyrosine Kinases | 7/536 | 0.001191 | 0.018783164 | 0 | 0 | 4.756353707 | 32.02290311 | EGR3;COL1A2;SPHK1;IL2RG;MMP9;DNM1;EREG |
| Liver | Biosynthesis of DHA-derived SPMs | 2/17 | 0.001208 | 0.018783164 | 0 | 0 | 45.0259887 | 302.5099237 | ALOX5;ALOX15 |

| | | | | | | | | | |
|---|---|---|---|---|---|---|---|---|---|
| Liver | Crosslinking of Collagen Fibrils | 2/18 | 0.001357 | 0.019771574 | 0 | 0 | 42.20974576 | 278.7000379 | COL1A2;LOXL1 |
| Liver | Signaling by Nuclear Receptors | 5/270 | 0.001387 | 0.019771574 | 0 | 0 | 6.628706199 | 43.6186564 | ALDH1A3;SPHK1;ALDH1A2;MMP9;EREG |
| Liver | Biosynthesis of Specialized Proresolving Mediators (SPMs) | 2/19 | 0.001513 | 0.020510766 | 0 | 0 | 39.72482552 | 257.9522541 | ALOX5;ALOX15 |
| Liver | Extra-nuclear Estrogen Signaling | 3/75 | 0.001559 | 0.020510766 | 0 | 0 | 14.27227011 | 92.24909678 | SPHK1;MMP9;EREG |
| Liver | Synthesis of Leukotrienes (LT) and Eoxins (EX) | 2/20 | 0.001678 | 0.021256254 | 0 | 0 | 37.51600753 | 239.7302132 | ALOX5;ALOX15 |
| Liver | RA Biosynthesis Pathway | 2/22 | 0.002032 | 0.024822637 | 0 | 0 | 33.76101695 | 209.2712255 | ALDH1A3;ALDH1A2 |
| Liver | G Alpha (I) Signalling Events | 5/316 | 0.002754 | 0.0324764 | 0 | 0 | 5.635048232 | 33.21723735 | CCR1;PRKAR2B;CXCR4;ACKR3;CX3CL1 |
| Liver | Syndecan Interactions | 2/27 | 0.003058 | 0.034859841 | 0 | 0 | 27.0020339 | 156.3426713 | COL1A2;TNC |
| Liver | Axon Guidance | 6/541 | 0.00601 | 0.066305058 | 0 | 0 | 3.956635514 | 20.23547098 | ENAH;SEMA4D;CXCR4;CACNA1C;MMP9;DNM1 |
| Liver | Signaling by Retinoic Acid | 2/41 | 0.006951 | 0.074287247 | 0 | 0 | 17.29682747 | 85.94604723 | ALDH1A3;ALDH1A2 |
| Liver | Nervous System Development | 6/567 | 0.007504 | 0.077771837 | 0 | 0 | 3.768206125 | 18.43511599 | ENAH;SEMA4D;CXCR4;CACNA1C;MMP9;DNM1 |
| Liver | Elastic Fibre Formation | 2/44 | 0.007972 | 0.077898726 | 0 | 0 | 16.05891848 | 77.59361647 | ITGB8;LOXL1 |
| Liver | Collagen Chain Trimerization | 2/44 | 0.007972 | 0.077898726 | 0 | 0 | 16.05891848 | 77.59361647 | COL1A2;COL12A1 |
| Liver | Interleukin-10 Signaling | 2/46 | 0.008688 | 0.082537365 | 0 | 0 | 15.32742681 | 72.74084051 | CCR1;IL1RN |
| Liver | EPH-ephrin Mediated Repulsion of Cells | 2/51 | 0.010599 | 0.097968871 | 0 | 0 | 13.75994466 | 62.56644514 | MMP9;DNM1 |
| Liver | GPCR Downstream Signalling | 6/633 | 0.012485 | 0.11236694 | 0 | 0 | 3.360069595 | 14.72789097 | CCR1;PTGIR;PRKAR2B;CXCR4;ACKR3;CX3CL1 |
| Liver | Signaling by TGFB Family Members | 3/161 | 0.013103 | 0.1149008 | 0 | 0 | 6.475665648 | 28.07159109 | COL1A2;FST;ITGB8 |
| Liver | Arachidonate Metabolism | 2/59 | 0.014004 | 0.119730884 | 0 | 0 | 11.8239667 | 50.46989209 | ALOX5;ALOX15 |
| Liver | Collagen Biosynthesis and Modifying Enzymes | 2/67 | 0.017817 | 0.139959885 | 0 | 0 | 10.36453716 | 41.74441134 | COL1A2;COL12A1 |
| Liver | Biosynthesis of Lipoxins (LX) | 1/6 | 0.018163 | 0.139959885 | 0 | 0 | 66.44666667 | 266.3422351 | ALOX5 |
| Liver | Synthesis of 15-Eicosatetraenoic Acid Derivatives | 1/6 | 0.018163 | 0.139959885 | 0 | 0 | 66.44666667 | 266.3422351 | ALOX15 |
| Liver | Hemostasis | 6/707 | 0.020443 | 0.139959885 | 0 | 0 | 2.993852937 | 11.64637194 | PTGIR;COL1A2;PRKAR2B;ITGAX;F13A1;CD44 |
| Liver | ESR-mediated Signaling | 3/192 | 0.020863 | 0.139959885 | 0 | 0 | 5.405035577 | 20.91641328 | SPHK1;MMP9;EREG |
| Liver | Enhanced Binding of GP1BA Variant to VWF Multimer Collagen | 1/7 | 0.021159 | 0.139959885 | 0 | 0 | 55.36944444 | 213.4883869 | COL1A2 |
| Liver | Synthesis of 12-Eicosatetraenoic Acid Derivatives | 1/7 | 0.021159 | 0.139959885 | 0 | 0 | 55.36944444 | 213.4883869 | ALOX15 |
| Liver | Defective Binding of VWF Variant to GPIb IX V | 1/7 | 0.021159 | 0.139959885 | 0 | 0 | 55.36944444 | 213.4883869 | COL1A2 |
| Liver | Regulation of Insulin Secretion | 2/78 | 0.023688 | 0.139959885 | 0 | 0 | 8.859500446 | 33.15919842 | PRKAR2B;CACNA1C |
| Liver | Sema4D Mediated Inhibition of Cell Attachment and Migration | 1/8 | 0.024145 | 0.139959885 | 0 | 0 | 47.45714286 | 176.7148687 | SEMA4D |
| Liver | Biosynthesis of Maresins | 1/8 | 0.024145 | 0.139959885 | 0 | 0 | 47.45714286 | 176.7148687 | ALOX5 |
| Liver | Signaling by Overexpressed Wild-Type EGFR in Cancer | 1/8 | 0.024145 | 0.139959885 | 0 | 0 | 47.45714286 | 176.7148687 | EREG |
| Liver | Inhibition of Signaling by Overexpressed EGFR | 1/8 | 0.024145 | 0.139959885 | 0 | 0 | 47.45714286 | 176.7148687 | EREG |
| Liver | Insulin-like Growth Factor-2 mRNA Binding Proteins (IGF2BPs IMPs VICKZs) Bind RNA | 1/8 | 0.024145 | 0.139959885 | 0 | 0 | 47.45714286 | 176.7148687 | CD44 |
| Liver | Interleukin-18 Signaling | 1/8 | 0.024145 | 0.139959885 | 0 | 0 | 47.45714286 | 176.7148687 | ALOX5 |
| Liver | Defective CHST14 Causes EDS, Musculocontractural Type | 1/8 | 0.024145 | 0.139959885 | 0 | 0 | 47.45714286 | 176.7148687 | VCAN |
| Liver | Defective CHST3 Causes SEDCJD | 1/8 | 0.024145 | 0.139959885 | 0 | 0 | 47.45714286 | 176.7148687 | VCAN |
| Liver | Defective CHSY1 Causes TPBS | 1/8 | 0.024145 | 0.139959885 | 0 | 0 | 47.45714286 | 176.7148687 | VCAN |
| Liver | Defects of Platelet Adhesion to Exposed Collagen | 1/8 | 0.024145 | 0.139959885 | 0 | 0 | 47.45714286 | 176.7148687 | COL1A2 |
| Liver | Synthesis of 5-Eicosatetraenoic Acids | 1/9 | 0.027123 | 0.147237318 | 0 | 0 | 41.52291667 | 149.7891714 | ALOX5 |
| Liver | Prostanoid Ligand Receptors | 1/9 | 0.027123 | 0.147237318 | 0 | 0 | 41.52291667 | 149.7891714 | PTGIR |
| Liver | Interleukin-9 Signaling | 1/9 | 0.027123 | 0.147237318 | 0 | 0 | 41.52291667 | 149.7891714 | IL2RG |
| Liver | EGFR Interacts With Phospholipase C-gamma | 1/9 | 0.027123 | 0.147237318 | 0 | 0 | 41.52291667 | 149.7891714 | EREG |
| Liver | PI3K Events in ERBB4 Signaling | 1/10 | 0.030091 | 0.155855747 | 0 | 0 | 36.90740741 | 129.3058522 | EREG |
| Liver | Interleukin-21 Signaling | 1/10 | 0.030091 | 0.155855747 | 0 | 0 | 36.90740741 | 129.3058522 | IL2RG |
| Liver | EPH-Ephrin Signaling | 2/92 | 0.032138 | 0.155855747 | 0 | 0 | 7.476082863 | 25.70071278 | MMP9;DNM1 |
| Liver | Activation of NMDA Receptors and Postsynaptic Events | 2/93 | 0.032781 | 0.155855747 | 0 | 0 | 7.393555597 | 25.27054507 | PRKAR2B;GRIA3 |
| Liver | STAT3 Nuclear Events Downstream of ALK Signaling | 1/11 | 0.033051 | 0.155855747 | 0 | 0 | 33.215 | 113.2533429 | IL2RG |

| | | | | | | | | | |
|---|---|---|---|---|---|---|---|---|---|
| Liver | Formation of Annular Gap Junctions | 1/11 | 0.033051 | 0.155855747 | 0 | 0 | 33.215 | 113.2533429 | DNM1 |
| Liver | Dermatan Sulfate Biosynthesis | 1/11 | 0.033051 | 0.155855747 | 0 | 0 | 33.215 | 113.2533429 | VCAN |
| Liver | Signaling by TGF-beta Receptor Complex | 2/94 | 0.033429 | 0.155855747 | 0 | 0 | 7.312822402 | 24.85144987 | COL1A2;ITGB8 |
| Liver | Cell Surface Interactions at the Vascular Wall | 3/233 | 0.034269 | 0.155855747 | 0 | 0 | 4.432308846 | 14.95250564 | COL1A2;ITGAX;CD44 |
| Liver | GP1b-IX-V Activation Signalling | 1/12 | 0.036002 | 0.155855747 | 0 | 0 | 30.19393939 | 100.3703093 | COL1A2 |
| Liver | Gap Junction Degradation | 1/12 | 0.036002 | 0.155855747 | 0 | 0 | 30.19393939 | 100.3703093 | DNM1 |
| Liver | Binding and Entry of HIV Virion | 1/12 | 0.036002 | 0.155855747 | 0 | 0 | 30.19393939 | 100.3703093 | CXCR4 |
| Liver | CREB1 Phosphorylation Through the Activation of Adenylate Cyclase | 1/12 | 0.036002 | 0.155855747 | 0 | 0 | 30.19393939 | 100.3703093 | PRKAR2B |
| Liver | Specification of Primordial Germ Cells | 1/12 | 0.036002 | 0.155855747 | 0 | 0 | 30.19393939 | 100.3703093 | CXCR4 |
| Liver | Hyaluronan Uptake and Degradation | 1/12 | 0.036002 | 0.155855747 | 0 | 0 | 30.19393939 | 100.3703093 | CD44 |
| Liver | Interleukin-2 Signaling | 1/12 | 0.036002 | 0.155855747 | 0 | 0 | 30.19393939 | 100.3703093 | IL2RG |
| Liver | Interferon Gamma Signaling | 2/99 | 0.036744 | 0.156691192 | 0 | 0 | 6.934125459 | 22.90889677 | CIITA;CD44 |
| Liver | WNT5A-dependent Internalization of FZD2, FZD5 and ROR2 | 1/13 | 0.038944 | 0.156691192 | 0 | 0 | 27.67638889 | 89.82753445 | FZD2 |
| Liver | Nephron Development | 1/13 | 0.038944 | 0.156691192 | 0 | 0 | 27.67638889 | 89.82753445 | WT1 |
| Liver | GRB2 Events in EGFR Signaling | 1/13 | 0.038944 | 0.156691192 | 0 | 0 | 27.67638889 | 89.82753445 | EREG |
| Liver | Transcriptional Regulation of Testis Differentiation | 1/13 | 0.038944 | 0.156691192 | 0 | 0 | 27.67638889 | 89.82753445 | WT1 |
| Liver | ERBB2 Activates PTK6 Signaling | 1/13 | 0.038944 | 0.156691192 | 0 | 0 | 27.67638889 | 89.82753445 | EREG |
| Liver | Retrograde Neurotrophin Signalling | 1/14 | 0.041877 | 0.158965705 | 0 | 0 | 25.54615385 | 81.05851862 | DNM1 |
| Liver | SHC1 Events in EGFR Signaling | 1/14 | 0.041877 | 0.158965705 | 0 | 0 | 25.54615385 | 81.05851862 | EREG |
| Liver | SHC1 Events in ERBB4 Signaling | 1/14 | 0.041877 | 0.158965705 | 0 | 0 | 25.54615385 | 81.05851862 | EREG |
| Liver | CS DS Degradation | 1/14 | 0.041877 | 0.158965705 | 0 | 0 | 25.54615385 | 81.05851862 | VCAN |
| Liver | Interleukin-15 Signaling | 1/14 | 0.041877 | 0.158965705 | 0 | 0 | 25.54615385 | 81.05851862 | IL2RG |
| Liver | Post-translational Protein Phosphorylation | 2/107 | 0.042298 | 0.158965705 | 0 | 0 | 6.40322841 | 20.25352686 | VCAN;TNC |
| Liver | Integration of Energy Metabolism | 2/108 | 0.043013 | 0.159604106 | 0 | 0 | 6.342500799 | 19.95509283 | PRKAR2B;CACNA1C |
| Liver | Eicosanoid Ligand-Binding Receptors | 1/15 | 0.044801 | 0.159604106 | 0 | 0 | 23.7202381 | 73.66370739 | PTGIR |
| Liver | Anchoring Fibril Formation | 1/15 | 0.044801 | 0.159604106 | 0 | 0 | 23.7202381 | 73.66370739 | COL1A2 |
| Liver | Phase 2 - Plateau Phase | 1/15 | 0.044801 | 0.159604106 | 0 | 0 | 23.7202381 | 73.66370739 | CACNA1C |
| Liver | ERBB2 Regulates Cell Motility | 1/15 | 0.044801 | 0.159604106 | 0 | 0 | 23.7202381 | 73.66370739 | EREG |
| Liver | GRB2 Events in ERBB2 Signaling | 1/16 | 0.047717 | 0.163529936 | 0 | 0 | 22.13777778 | 67.3536152 | EREG |
| Liver | PI3K Events in ERBB2 Signaling | 1/16 | 0.047717 | 0.163529936 | 0 | 0 | 22.13777778 | 67.3536152 | EREG |
| Liver | Platelet Adhesion to Exposed Collagen | 1/16 | 0.047717 | 0.163529936 | 0 | 0 | 22.13777778 | 67.3536152 | COL1A2 |
| Liver | Signaling by NTRK1 (TRKA) | 2/115 | 0.048145 | 0.163529936 | 0 | 0 | 5.947502625 | 18.04196078 | EGR3;DNM1 |
| Liver | Signaling by Activin | 1/17 | 0.050623 | 0.163529936 | 0 | 0 | 20.753125 | 61.91362722 | FST |
| Liver | GAB1 Signalosome | 1/17 | 0.050623 | 0.163529936 | 0 | 0 | 20.753125 | 61.91362722 | EREG |
| Liver | PKA Activation in Glucagon Signalling | 1/17 | 0.050623 | 0.163529936 | 0 | 0 | 20.753125 | 61.91362722 | PRKAR2B |
| Liver | Hyaluronan Metabolism | 1/17 | 0.050623 | 0.163529936 | 0 | 0 | 20.753125 | 61.91362722 | CD44 |
| Liver | Trafficking of GluR2-containing AMPA Receptors | 1/17 | 0.050623 | 0.163529936 | 0 | 0 | 20.753125 | 61.91362722 | GRIA3 |
| Liver | Formation of Definitive Endoderm | 1/18 | 0.053522 | 0.163529936 | 0 | 0 | 19.53137255 | 57.18144006 | CXCR4 |
| Liver | PKA Activation | 1/18 | 0.053522 | 0.163529936 | 0 | 0 | 19.53137255 | 57.18144006 | PRKAR2B |
| Liver | Interferon Signaling | 3/280 | 0.053966 | 0.163529936 | 0 | 0 | 3.671480144 | 10.71849447 | CIITA;SPHK1;CD44 |
| Liver | Regulation of IGF Transport and Uptake by Insulin-like Growth Factor Binding Proteins (IGFBPs) | 2/124 | 0.055052 | 0.163529936 | 0 | 0 | 5.506251737 | 15.96520741 | VCAN;TNC |
| Liver | Scavenging by Class A Receptors | 1/19 | 0.056411 | 0.163529936 | 0 | 0 | 18.44537037 | 53.03215997 | COL1A2 |
| Liver | Other Semaphorin Interactions | 1/19 | 0.056411 | 0.163529936 | 0 | 0 | 18.44537037 | 53.03215997 | SEMA4D |
| Liver | Prostacyclin Signalling Through Prostacyclin Receptor | 1/19 | 0.056411 | 0.163529936 | 0 | 0 | 18.44537037 | 53.03215997 | PTGIR |
| Liver | Diseases of Hemostasis | 1/19 | 0.056411 | 0.163529936 | 0 | 0 | 18.44537037 | 53.03215997 | COL1A2 |
| Liver | VEGFR2 Mediated Cell Proliferation | 1/19 | 0.056411 | 0.163529936 | 0 | 0 | 18.44537037 | 53.03215997 | SPHK1 |
| Liver | Glycosaminoglycan Metabolism | 2/127 | 0.057429 | 0.163529936 | 0 | 0 | 5.373288136 | 15.3526368 | VCAN;CD44 |
| Liver | Neutrophil Degranulation | 4/478 | 0.057938 | 0.163529936 | 0 | 0 | 2.881782515 | 8.208394447 | ALOX5;ITGAX;MMP9;CD44 |
| Liver | Developmental Biology | 8/1385 | 0.058425 | 0.163529936 | 0 | 0 | 2.034721366 | 5.778627238 | ENAH;SEMA4D;WT1;CXCR4;MYO5A;CACNA1C;MMP9;DNM1 |
| Liver | Metabolism of Carbohydrates | 3/290 | 0.058731 | 0.163529936 | 0 | 0 | 3.541751772 | 10.04013327 | VCAN;ENO2;CD44 |
| Liver | Sema4D Induced Cell Migration and Growth-Cone Collapse | 1/20 | 0.059292 | 0.163529936 | 0 | 0 | 17.47368421 | 49.36819704 | SEMA4D |
| Liver | PKA-mediated Phosphorylation of CREB | 1/20 | 0.059292 | 0.163529936 | 0 | 0 | 17.47368421 | 49.36819704 | PRKAR2B |

| | | | | | | | | | |
|---|---|---|---|---|---|---|---|---|---|
| Liver | Chondroitin Sulfate Biosynthesis | 1/20 | 0.059292 | 0.163529936 | 0 | 0 | 17.47368421 | 49.36819704 | VCAN |
| Liver | Defective B3GALT6 Causes EDSP2 and SEMDJL1 | 1/20 | 0.059292 | 0.163529936 | 0 | 0 | 17.47368421 | 49.36819704 | VCAN |
| Liver | Defective B3GAT3 Causes JDSSDHD | 1/20 | 0.059292 | 0.163529936 | 0 | 0 | 17.47368421 | 49.36819704 | VCAN |
| Liver | Defective B4GALT7 Causes EDS, Progeroid Type | 1/20 | 0.059292 | 0.163529936 | 0 | 0 | 17.47368421 | 49.36819704 | VCAN |
| Liver | Synaptic Adhesion-Like Molecules | 1/21 | 0.062164 | 0.166093321 | 0 | 0 | 16.59916667 | 46.11225173 | GRIA3 |
| Liver | Inactivation of APC C via Direct Inhibition of the APC C Complex | 1/21 | 0.062164 | 0.166093321 | 0 | 0 | 16.59916667 | 46.11225173 | BUB1B |
| Liver | Inhib of Proteolytic Activity of APC C Required for Onset of Anaphase by Mitotic Spindle Checkpoint | 1/21 | 0.062164 | 0.166093321 | 0 | 0 | 16.59916667 | 46.11225173 | BUB1B |
| Liver | Unblocking of NMDA Receptors, Glutamate Binding and Activation | 1/21 | 0.062164 | 0.166093321 | 0 | 0 | 16.59916667 | 46.11225173 | GRIA3 |
| Liver | Signaling by NTRKs | 2/134 | 0.063109 | 0.167212244 | 0 | 0 | 5.0865434 | 14.05356933 | EGR3;DNM1 |
| Liver | SHC1 Events in ERBB2 Signaling | 1/22 | 0.065027 | 0.167212244 | 0 | 0 | 15.80793651 | 43.20234689 | EREG |
| Liver | Signaling by ERBB2 TMD JMD Mutants | 1/22 | 0.065027 | 0.167212244 | 0 | 0 | 15.80793651 | 43.20234689 | EREG |
| Liver | Common Pathway of Fibrin Clot Formation | 1/22 | 0.065027 | 0.167212244 | 0 | 0 | 15.80793651 | 43.20234689 | F13A1 |
| Liver | Early Phase of HIV Life Cycle | 1/22 | 0.065027 | 0.167212244 | 0 | 0 | 15.80793651 | 43.20234689 | CXCR4 |
| Liver | Interleukin-1 Family Signaling | 2/140 | 0.068124 | 0.173867896 | 0 | 0 | 4.8639155 | 13.06657057 | IL1RN;ALOX5 |
| Liver | Estrogen-dependent Nuclear Events Downstream of ESR-membrane Signaling | 1/24 | 0.070728 | 0.175940329 | 0 | 0 | 14.43188406 | 38.22881323 | EREG |
| Liver | Sema4D in Semaphorin Signaling | 1/24 | 0.070728 | 0.175940329 | 0 | 0 | 14.43188406 | 38.22881323 | SEMA4D |
| Liver | DARPP-32 Events | 1/24 | 0.070728 | 0.175940329 | 0 | 0 | 14.43188406 | 38.22881323 | PRKAR2B |
| Liver | Clathrin-mediated Endocytosis | 2/145 | 0.0724 | 0.175940329 | 0 | 0 | 4.692663269 | 12.32078652 | DNM1;EREG |
| Liver | Diseases of Glycosylation | 2/146 | 0.073266 | 0.175940329 | 0 | 0 | 4.659839925 | 12.17921997 | SPON1;VCAN |
| Liver | Signaling by EGFR in Cancer | 1/25 | 0.073566 | 0.175940329 | 0 | 0 | 13.82986111 | 36.09008042 | EREG |
| Liver | Signaling by ERBB2 KD Mutants | 1/25 | 0.073566 | 0.175940329 | 0 | 0 | 13.82986111 | 36.09008042 | EREG |
| Liver | Insulin Processing | 1/25 | 0.073566 | 0.175940329 | 0 | 0 | 13.82986111 | 36.09008042 | MYO5A |
| Liver | Interleukin-7 Signaling | 1/25 | 0.073566 | 0.175940329 | 0 | 0 | 13.82986111 | 36.09008042 | IL2RG |
| Liver | A Tetrasaccharide Linker Sequence Is Required for GAG Synthesis | 1/26 | 0.076395 | 0.177734954 | 0 | 0 | 13.276 | 34.143748 | VCAN |
| Liver | APC-Cdc20 Mediated Degradation of Nek2A | 1/26 | 0.076395 | 0.177734954 | 0 | 0 | 13.276 | 34.143748 | BUB1B |
| Liver | Signaling by ERBB2 in Cancer | 1/26 | 0.076395 | 0.177734954 | 0 | 0 | 13.276 | 34.143748 | EREG |
| Liver | Gluconeogenesis | 1/26 | 0.076395 | 0.177734954 | 0 | 0 | 13.276 | 34.143748 | ENO2 |
| Liver | Interleukin Receptor SHC Signaling | 1/27 | 0.079216 | 0.183052047 | 0 | 0 | 12.76474359 | 32.36606964 | IL2RG |
| Liver | Adrenaline,noradrenaline Inhibits Insulin Secretion | 1/28 | 0.082028 | 0.188278321 | 0 | 0 | 12.29135802 | 30.73697946 | CACNA1C |
| Liver | G Alpha (S) Signalling Events | 2/158 | 0.08391 | 0.190870707 | 0 | 0 | 4.298783138 | 10.65241484 | PTGIR;PRKAR2B |
| Liver | TNFs Bind Their Physiological Receptors | 1/29 | 0.084831 | 0.190870707 | 0 | 0 | 11.85178571 | 29.23941281 | TNFRSF9 |
| Liver | Downregulation of ERBB2 Signaling | 1/29 | 0.084831 | 0.190870707 | 0 | 0 | 11.85178571 | 29.23941281 | EREG |
| Liver | Signaling by ALK | 1/30 | 0.087627 | 0.193344157 | 0 | 0 | 11.44252874 | 27.85877118 | IL2RG |
| Liver | PD-1 Signaling | 1/30 | 0.087627 | 0.193344157 | 0 | 0 | 11.44252874 | 27.85877118 | PDCD1LG2 |
| Liver | MET Activates PTK2 Signaling | 1/30 | 0.087627 | 0.193344157 | 0 | 0 | 11.44252874 | 27.85877118 | COL1A2 |
| Liver | Glutamate Binding, Activation of AMPA Receptors and Synaptic Plasticity | 1/31 | 0.090414 | 0.194474647 | 0 | 0 | 11.06055556 | 26.58249692 | GRIA3 |
| Liver | Trafficking of AMPA Receptors | 1/31 | 0.090414 | 0.194474647 | 0 | 0 | 11.06055556 | 26.58249692 | GRIA3 |
| Liver | Disassembly of the Destruction Complex and Recruitment of AXIN to the Membrane | 1/31 | 0.090414 | 0.194474647 | 0 | 0 | 11.06055556 | 26.58249692 | FZD2 |
| Liver | EGFR Downregulation | 1/31 | 0.090414 | 0.194474647 | 0 | 0 | 11.06055556 | 26.58249692 | EREG |
| Liver | Nuclear Signaling by ERBB4 | 1/32 | 0.093192 | 0.197961053 | 0 | 0 | 10.70322581 | 25.39973247 | EREG |
| Liver | Phase 0 - Rapid Depolarisation | 1/32 | 0.093192 | 0.197961053 | 0 | 0 | 10.70322581 | 25.39973247 | CACNA1C |
| Liver | Activation of Matrix Metalloproteinases | 1/33 | 0.095962 | 0.201344387 | 0 | 0 | 10.36822917 | 24.30104522 | MMP9 |
| Liver | Glucagon Signaling in Metabolic Regulation | 1/33 | 0.095962 | 0.201344387 | 0 | 0 | 10.36822917 | 24.30104522 | PRKAR2B |
| Liver | Fatty Acid Metabolism | 2/176 | 0.100688 | 0.207816848 | 0 | 0 | 3.850574713 | 8.839889377 | ALOX5;ALOX15 |
| Liver | GPVI-mediated Activation Cascade | 1/35 | 0.101478 | 0.207816848 | 0 | 0 | 9.757352941 | 22.32399464 | COL1A2 |
| Liver | CaM Pathway | 1/35 | 0.101478 | 0.207816848 | 0 | 0 | 9.757352941 | 22.32399464 | PRKAR2B |
| Liver | Calmodulin Induced Events | 1/35 | 0.101478 | 0.207816848 | 0 | 0 | 9.757352941 | 22.32399464 | PRKAR2B |
| Liver | SMAD2 SMAD3 SMAD4 Heterotrimer Regulates Transcription | 1/36 | 0.104223 | 0.212168474 | 0 | 0 | 9.478095238 | 21.43207168 | COL1A2 |

| | | | | | | | | | |
|---|---|---|---|---|---|---|---|---|---|
| Liver | Molecules Associated With Elastic Fibres | 1/37 | 0.10696 | 0.213920303 | 0 | 0 | 9.214351852 | 20.59683081 | ITGB8 |
| Liver | Ca-dependent Events | 1/37 | 0.10696 | 0.213920303 | 0 | 0 | 9.214351852 | 20.59683081 | PRKAR2B |
| Liver | Sphingolipid De Novo Biosynthesis | 1/37 | 0.10696 | 0.213920303 | 0 | 0 | 9.214351852 | 20.59683081 | SPHK1 |
| Liver | Defective B3GALTL Causes PpS | 1/38 | 0.109689 | 0.217198173 | 0 | 0 | 8.964864865 | 19.81330602 | SPON1 |
| Liver | NGF-stimulated Transcription | 1/39 | 0.11241 | 0.217198173 | 0 | 0 | 8.728508772 | 19.07708209 | EGR3 |
| Liver | Formation of Fibrin Clot (Clotting Cascade) | 1/39 | 0.11241 | 0.217198173 | 0 | 0 | 8.728508772 | 19.07708209 | F13A1 |
| Liver | O-glycosylation of TSR Domain-Containing Proteins | 1/39 | 0.11241 | 0.217198173 | 0 | 0 | 8.728508772 | 19.07708209 | SPON1 |
| Liver | Association of TriC CCT With Target Proteins During Biosynthesis | 1/39 | 0.11241 | 0.217198173 | 0 | 0 | 8.728508772 | 19.07708209 | SPHK1 |
| Liver | Platelet Aggregation (Plug Formation) | 1/39 | 0.11241 | 0.217198173 | 0 | 0 | 8.728508772 | 19.07708209 | COL1A2 |
| Liver | Generation of Second Messenger Molecules | 1/41 | 0.117826 | 0.222633116 | 0 | 0 | 8.29125 | 17.73120117 | ENAH |
| Liver | DAG and IP3 Signaling | 1/41 | 0.117826 | 0.222633116 | 0 | 0 | 8.29125 | 17.73120117 | PRKAR2B |
| Liver | MET Promotes Cell Motility | 1/41 | 0.117826 | 0.222633116 | 0 | 0 | 8.29125 | 17.73120117 | COL1A2 |
| Liver | Diseases Associated With Glycosaminoglycan Metabolism | 1/41 | 0.117826 | 0.222633116 | 0 | 0 | 8.29125 | 17.73120117 | VCAN |
| Liver | NCAM1 Interactions | 1/42 | 0.120522 | 0.224014556 | 0 | 0 | 8.088617886 | 17.11486211 | CACNA1C |
| Liver | Glucagon-like Peptide-1 (GLP1) Regulates Insulin Secretion | 1/42 | 0.120522 | 0.224014556 | 0 | 0 | 8.088617886 | 17.11486211 | PRKAR2B |
| Liver | Regulation of MITF-M-dependent Genes Involved in Pigmentation | 1/42 | 0.120522 | 0.224014556 | 0 | 0 | 8.088617886 | 17.11486211 | MYO5A |
| Liver | ADORA2B Mediated Anti-Inflammatory Cytokines Production | 1/43 | 0.12321 | 0.225336847 | 0 | 0 | 7.895634921 | 16.53236218 | PRKAR2B |
| Liver | Signaling by SCF-KIT | 1/43 | 0.12321 | 0.225336847 | 0 | 0 | 7.895634921 | 16.53236218 | MMP9 |
| Liver | Vasopressin Regulates Renal Water Homeostasis via Aquaporins | 1/43 | 0.12321 | 0.225336847 | 0 | 0 | 7.895634921 | 16.53236218 | PRKAR2B |
| Liver | Interleukin-2 Family Signaling | 1/44 | 0.12589 | 0.229013504 | 0 | 0 | 7.711627907 | 15.98113875 | IL2RG |
| Liver | Mitotic Prometaphase | 2/203 | 0.127392 | 0.230200687 | 0 | 0 | 3.32877983 | 6.858892975 | PRKAR2B;BUB1B |
| Liver | Neurotransmitter Receptors and Postsynaptic Signal Transmission | 2/204 | 0.128412 | 0.230200687 | 0 | 0 | 3.312132908 | 6.798189361 | PRKAR2B;GRIA3 |
| Liver | GPER1 Signaling | 1/45 | 0.128562 | 0.230200687 | 0 | 0 | 7.535984848 | 15.45887549 | PRKAR2B |
| Liver | Kidney Development | 1/46 | 0.131226 | 0.233746752 | 0 | 0 | 7.368148148 | 14.96347378 | WT1 |
| Liver | TGF-beta Receptor Signaling Activates SMADs | 1/47 | 0.133882 | 0.236126243 | 0 | 0 | 7.207608696 | 14.4930281 | ITGB8 |
| Liver | Signaling by ROBO Receptors | 2/210 | 0.13457 | 0.236126243 | 0 | 0 | 3.215612777 | 6.449457435 | ENAH;CXCR4 |
| Liver | Gap Junction Trafficking | 1/48 | 0.13653 | 0.236126243 | 0 | 0 | 7.053900709 | 14.0458046 | DNM1 |
| Liver | Interleukin-3, Interleukin-5 and GM-CSF Signaling | 1/48 | 0.13653 | 0.236126243 | 0 | 0 | 7.053900709 | 14.0458046 | IL2RG |
| Liver | Recycling Pathway of L1 | 1/48 | 0.13653 | 0.236126243 | 0 | 0 | 7.053900709 | 14.0458046 | DNM1 |
| Liver | Innate Immune System | 6/1149 | 0.136705 | 0.236126243 | 0 | 0 | 1.793939394 | 3.569818116 | ALOX5;ITGAX;MYO5A;MMP9;DNM1;CD44 |
| Liver | PLC Beta Mediated Events | 1/49 | 0.13917 | 0.239176419 | 0 | 0 | 6.906597222 | 13.62022245 | PRKAR2B |
| Liver | Signaling by ERBB2 | 1/50 | 0.141802 | 0.241274842 | 0 | 0 | 6.765306122 | 13.21483754 | EREG |
| Liver | Chondroitin Sulfate Dermatan Sulfate Metabolism | 1/50 | 0.141802 | 0.241274842 | 0 | 0 | 6.765306122 | 13.21483754 | VCAN |
| Liver | Asymmetric Localization of PCP Proteins | 1/51 | 0.144426 | 0.242125842 | 0 | 0 | 6.629666667 | 12.82832824 | FZD2 |
| Liver | Gap Junction Trafficking and Regulation | 1/51 | 0.144426 | 0.242125842 | 0 | 0 | 6.629666667 | 12.82832824 | DNM1 |
| Liver | Transcriptional Activity of SMAD2 SMAD3 SMAD4 Heterotrimer | 1/51 | 0.144426 | 0.242125842 | 0 | 0 | 6.629666667 | 12.82832824 | COL1A2 |
| Liver | Aquaporin-mediated Transport | 1/52 | 0.147042 | 0.245309275 | 0 | 0 | 6.499346405 | 12.45948282 | PRKAR2B |
| Liver | Signaling by EGFR | 1/53 | 0.14965 | 0.248448737 | 0 | 0 | 6.374038462 | 12.10718848 | EREG |
| Liver | G-protein Mediated Events | 1/54 | 0.152251 | 0.251544903 | 0 | 0 | 6.253459119 | 11.77042153 | PRKAR2B |
| Liver | Signaling by Non-Receptor Tyrosine Kinases | 1/55 | 0.154843 | 0.253380262 | 0 | 0 | 6.137345679 | 11.4482388 | EREG |
| Liver | Signaling by PTK6 | 1/55 | 0.154843 | 0.253380262 | 0 | 0 | 6.137345679 | 11.4482388 | EREG |
| Liver | Heparan Sulfate Heparin (HS-GAG) Metabolism | 1/57 | 0.160005 | 0.260580192 | 0 | 0 | 5.917559524 | 10.84421089 | VCAN |
| Liver | Signaling by ERBB4 | 1/58 | 0.162575 | 0.263509676 | 0 | 0 | 5.813450292 | 10.56081724 | EREG |
| Liver | Cdc20 Phospho-APC C Mediated Degradation of Cyclin A | 1/60 | 0.16769 | 0.270518967 | 0 | 0 | 5.615819209 | 10.02781753 | BUB1B |
| Liver | APC Cdc20 Mediated Degradation of Cell Cycle Proteins Prior to Satisfation of the Checkpoint | 1/61 | 0.170236 | 0.27182805 | 0 | 0 | 5.521944444 | 9.776977069 | BUB1B |

| | | | | | | | | | |
|---|---|---|---|---|---|---|---|---|---|
| Liver | Nuclear Events (Kinase and Transcription Factor Activation) | 1/61 | 0.170236 | 0.27182805 | 0 | 0 | 5.521944444 | 9.776977069 | EGR3 |
| Liver | Parasitic Infection Pathways | 2/245 | 0.171681 | 0.27182805 | 0 | 0 | 2.747576201 | 4.841553237 | PRKAR2B;MYO5A |
| Liver | Leishmania Infection | 2/245 | 0.171681 | 0.27182805 | 0 | 0 | 2.747576201 | 4.841553237 | PRKAR2B;MYO5A |
| Liver | APC C Cdc20 Mediated Degradation of Mitotic Proteins | 1/63 | 0.175306 | 0.273765061 | 0 | 0 | 5.34327957 | 9.30384657 | BUB1B |
| Liver | NCAM Signaling for Neurite Out-Growth | 1/63 | 0.175306 | 0.273765061 | 0 | 0 | 5.34327957 | 9.30384657 | CACNA1C |
| Liver | Ca2+ Pathway | 1/63 | 0.175306 | 0.273765061 | 0 | 0 | 5.34327957 | 9.30384657 | FZD2 |
| Liver | Activation of APC C and APC C Cdc20 Mediated Degradation of Mitotic Proteins | 1/64 | 0.177829 | 0.275192326 | 0 | 0 | 5.258201058 | 9.080561423 | BUB1B |
| Liver | Semaphorin Interactions | 1/64 | 0.177829 | 0.275192326 | 0 | 0 | 5.258201058 | 9.080561423 | SEMA4D |
| Liver | Regulation of APC C Activators Between G1 S and Early Anaphase | 1/68 | 0.187846 | 0.28802424 | 0 | 0 | 4.943283582 | 8.265818938 | BUB1B |
| Liver | Platelet Activation, Signaling and Aggregation | 2/262 | 0.190281 | 0.28802424 | 0 | 0 | 2.565710561 | 4.257161131 | COL1A2;F13A1 |
| Liver | High Laminar Flow Shear Stress Activates Signaling by PIEZO1 and PECAM1 CDH5 KDR in Endothelial Cell | 1/69 | 0.190332 | 0.28802424 | 0 | 0 | 4.870343137 | 8.079832893 | PRKAR2B |
| Liver | Loss of Nlp From Mitotic Centrosomes | 1/69 | 0.190332 | 0.28802424 | 0 | 0 | 4.870343137 | 8.079832893 | PRKAR2B |
| Liver | Loss of Proteins Required for Interphase Microtubule Organization From the Centrosome | 1/69 | 0.190332 | 0.28802424 | 0 | 0 | 4.870343137 | 8.079832893 | PRKAR2B |
| Liver | Diseases of Metabolism | 2/264 | 0.192488 | 0.290004527 | 0 | 0 | 2.545866218 | 4.194873328 | SPON1;VCAN |
| Liver | RAF MAP Kinase Cascade | 2/265 | 0.193593 | 0.290373487 | 0 | 0 | 2.536057228 | 4.164194949 | IL2RG;EREG |
| Liver | Translocation of SLC2A4 (GLUT4) to the Plasma Membrane | 1/71 | 0.19528 | 0.290373487 | 0 | 0 | 4.730714286 | 7.726764237 | MYO5A |
| Liver | Diseases Associated With O-glycosylation of Proteins | 1/71 | 0.19528 | 0.290373487 | 0 | 0 | 4.730714286 | 7.726764237 | SPON1 |
| Liver | Vesicle-mediated Transport | 4/752 | 0.196317 | 0.290650735 | 0 | 0 | 1.800450324 | 2.931179905 | COL1A2;MYO5A;DNM1;EREG |
| Liver | AURKA Activation by TPX2 | 1/72 | 0.197744 | 0.290658256 | 0 | 0 | 4.663849765 | 7.55909442 | PRKAR2B |
| Liver | Transmission Across Chemical Synapses | 2/269 | 0.198022 | 0.290658256 | 0 | 0 | 2.497556021 | 4.044490682 | PRKAR2B;GRIA3 |
| Liver | Metabolism of Lipids | 4/758 | 0.200078 | 0.291413979 | 0 | 0 | 1.785564708 | 2.873055812 | SPHK1;ALOX5;ALOX15;FHL2 |
| Liver | MAPK1 MAPK3 Signaling | 2/271 | 0.200241 | 0.291413979 | 0 | 0 | 2.478734799 | 3.986389645 | IL2RG;EREG |
| Liver | Glycolysis | 1/74 | 0.202648 | 0.293667277 | 0 | 0 | 4.535616438 | 7.240144628 | ENO2 |
| Liver | APC C-mediated Degradation of Cell Cycle Proteins | 1/75 | 0.205089 | 0.294707073 | 0 | 0 | 4.474099099 | 7.088375395 | BUB1B |
| Liver | Regulation of Mitotic Cell Cycle | 1/75 | 0.205089 | 0.294707073 | 0 | 0 | 4.474099099 | 7.088375395 | BUB1B |
| Liver | Costimulation by the CD28 Family | 1/78 | 0.212367 | 0.303889739 | 0 | 0 | 4.299134199 | 6.661239871 | PDCD1LG2 |
| Liver | PCP CE Pathway | 1/79 | 0.214779 | 0.304412925 | 0 | 0 | 4.243803419 | 6.527586495 | FZD2 |
| Liver | Signaling by MET | 1/79 | 0.214779 | 0.304412925 | 0 | 0 | 4.243803419 | 6.527586495 | COL1A2 |
| Liver | Metabolism | 9/2181 | 0.216578 | 0.304412925 | 0 | 0 | 1.415772418 | 2.165856771 | VCAN;PRKAR2B;SPHK1;ALOX5;ALOX15;FHL2;CACNA1C;ENO2;CD44 |
| Liver | PKR-mediated Signaling | 1/80 | 0.217183 | 0.304412925 | 0 | 0 | 4.189873418 | 6.397989946 | SPHK1 |
| Liver | Post NMDA Receptor Activation Events | 1/80 | 0.217183 | 0.304412925 | 0 | 0 | 4.189873418 | 6.397989946 | PRKAR2B |
| Liver | Centrosome Maturation | 1/81 | 0.219581 | 0.305270682 | 0 | 0 | 4.137291667 | 6.272281522 | PRKAR2B |
| Liver | Recruitment of Mitotic Centrosome Proteins and Complexes | 1/81 | 0.219581 | 0.305270682 | 0 | 0 | 4.137291667 | 6.272281522 | PRKAR2B |
| Liver | Constitutive Signaling by Aberrant PI3K in Cancer | 1/83 | 0.224353 | 0.310643136 | 0 | 0 | 4.03597561 | 6.031898326 | EREG |
| Liver | Glucose Metabolism | 1/85 | 0.229097 | 0.315932669 | 0 | 0 | 3.939484127 | 5.805255921 | ENO2 |
| Liver | Peptide Hormone Metabolism | 1/86 | 0.231459 | 0.317907074 | 0 | 0 | 3.892941176 | 5.69675104 | MYO5A |
| Liver | Platelet Homeostasis | 1/87 | 0.233813 | 0.318581625 | 0 | 0 | 3.84748062 | 5.591291022 | PTGIR |
| Liver | Regulation of PLK1 Activity at G2 M Transition | 1/87 | 0.233813 | 0.318581625 | 0 | 0 | 3.84748062 | 5.591291022 | PRKAR2B |
| Liver | Protein-protein Interactions at Synapses | 1/88 | 0.23616 | 0.319235922 | 0 | 0 | 3.803065134 | 5.488759041 | GRIA3 |
| Liver | TNFR2 Non-Canonical NF-kB Pathway | 1/88 | 0.23616 | 0.319235922 | 0 | 0 | 3.803065134 | 5.488759041 | TNFRSF9 |
| Liver | Opioid Signalling | 1/90 | 0.240833 | 0.324271024 | 0 | 0 | 3.717228464 | 5.292040073 | PRKAR2B |
| Liver | Chaperonin-mediated Protein Folding | 1/91 | 0.243159 | 0.325218051 | 0 | 0 | 3.675740741 | 5.197646636 | SPHK1 |
| Liver | Response of Endothelial Cells to Shear Stress | 1/93 | 0.24779 | 0.325218051 | 0 | 0 | 3.595471014 | 5.016311897 | PRKAR2B |
| Liver | Cellular Responses to Mechanical Stimuli | 1/93 | 0.24779 | 0.325218051 | 0 | 0 | 3.595471014 | 5.016311897 | PRKAR2B |
| Liver | MAPK Family Signaling Cascades | 2/314 | 0.248519 | 0.325218051 | 0 | 0 | 2.132442416 | 2.968860681 | IL2RG;EREG |

| | | | | | | | | | |
|---|---|---|---|---|---|---|---|---|---|
| Liver | Amplification of Signal From Unattached Kinetochores via a MAD2 Inhibitory Signal | 1/94 | 0.250095 | 0.325218051 | 0 | 0 | 3.556630824 | 4.929191928 | BUB1B |
| Liver | Amplification of Signal From the Kinetochores | 1/94 | 0.250095 | 0.325218051 | 0 | 0 | 3.556630824 | 4.929191928 | BUB1B |
| Liver | Class B 2 (Secretin Family Receptors) | 1/94 | 0.250095 | 0.325218051 | 0 | 0 | 3.556630824 | 4.929191928 | FZD2 |
| Liver | Recruitment of NuMA to Mitotic Centrosomes | 1/94 | 0.250095 | 0.325218051 | 0 | 0 | 3.556630824 | 4.929191928 | PRKAR2B |
| Liver | MITF-M-dependent Gene Expression | 1/94 | 0.250095 | 0.325218051 | 0 | 0 | 3.556630824 | 4.929191928 | MYO5A |
| Liver | Intracellular Signaling by Second Messengers | 2/321 | 0.256442 | 0.330387234 | 0 | 0 | 2.084905159 | 2.837249583 | PRKAR2B;EREG |
| Liver | Anchoring of the Basal Body to the Plasma Membrane | 1/97 | 0.256968 | 0.330387234 | 0 | 0 | 3.444965278 | 4.681033646 | PRKAR2B |
| Liver | Protein Folding | 1/97 | 0.256968 | 0.330387234 | 0 | 0 | 3.444965278 | 4.681033646 | SPHK1 |
| Liver | Hedgehog 'Off' State | 1/99 | 0.261516 | 0.333725083 | 0 | 0 | 3.374319728 | 4.52584509 | PRKAR2B |
| Liver | VEGFA-VEGFR2 Pathway | 1/99 | 0.261516 | 0.333725083 | 0 | 0 | 3.374319728 | 4.52584509 | SPHK1 |
| Liver | Adaptive Immune System | 4/854 | 0.262934 | 0.334287508 | 0 | 0 | 1.575975232 | 2.105271575 | ENAH;COL1A2;PDCD1LG2;DNM1 |
| Liver | Interleukin-1 Signaling | 1/102 | 0.268286 | 0.339828737 | 0 | 0 | 3.27359736 | 4.307079516 | IL1RN |
| Liver | Cargo Recognition for Clathrin-Mediated Endocytosis | 1/105 | 0.274995 | 0.347041745 | 0 | 0 | 3.178685897 | 4.103690234 | EREG |
| Liver | Sphingolipid Metabolism | 1/107 | 0.279434 | 0.351347447 | 0 | 0 | 3.118396226 | 3.975918826 | SPHK1 |
| Liver | Signaling by VEGF | 1/108 | 0.281644 | 0.352828459 | 0 | 0 | 3.089096573 | 3.914232059 | SPHK1 |
| Liver | PI3K AKT Signaling in Cancer | 1/110 | 0.286043 | 0.357031627 | 0 | 0 | 3.032110092 | 3.795030047 | EREG |
| Liver | Mitotic Spindle Checkpoint | 1/111 | 0.288232 | 0.357157652 | 0 | 0 | 3.004393939 | 3.737429597 | BUB1B |
| Liver | PI5P, PP2A and IER3 Regulate PI3K AKT Signaling | 1/111 | 0.288232 | 0.357157652 | 0 | 0 | 3.004393939 | 3.737429597 | EREG |
| Liver | TCR Signaling | 1/113 | 0.292592 | 0.361250664 | 0 | 0 | 2.950446429 | 3.626029128 | ENAH |
| Liver | O-linked Glycosylation | 1/114 | 0.294762 | 0.362620586 | 0 | 0 | 2.924188791 | 3.572153447 | SPON1 |
| Liver | EML4 and NUDC in Mitotic Spindle Formation | 1/116 | 0.299082 | 0.366616454 | 0 | 0 | 2.873043478 | 3.467872707 | BUB1B |
| Liver | PPARA Activates Gene Expression | 1/117 | 0.301232 | 0.367933509 | 0 | 0 | 2.848132184 | 3.417400253 | FHL2 |
| Liver | Negative Regulation of the PI3K AKT Network | 1/118 | 0.303376 | 0.368909715 | 0 | 0 | 2.823646724 | 3.367996891 | EREG |
| Liver | Membrane Trafficking | 3/633 | 0.304189 | 0.368909715 | 0 | 0 | 1.585303777 | 1.88668113 | MYO5A;DNM1;EREG |
| Liver | Regulation of Lipid Metabolism by PPARalpha | 1/119 | 0.305513 | 0.369172875 | 0 | 0 | 2.799576271 | 3.319631958 | FHL2 |
| Liver | Gastrulation | 1/120 | 0.307644 | 0.369172875 | 0 | 0 | 2.775910364 | 3.272275909 | CXCR4 |
| Liver | L1CAM Interactions | 1/120 | 0.307644 | 0.369172875 | 0 | 0 | 2.775910364 | 3.272275909 | DNM1 |
| Liver | FCGR3A-mediated IL10 Synthesis | 1/121 | 0.309768 | 0.370422462 | 0 | 0 | 2.752638889 | 3.225900266 | PRKAR2B |
| Liver | Binding and Uptake of Ligands by Scavenger Receptors | 1/122 | 0.311887 | 0.3716557 | 0 | 0 | 2.729752066 | 3.180477573 | COL1A2 |
| Liver | M Phase | 2/372 | 0.314136 | 0.373036302 | 0 | 0 | 1.792853871 | 2.075998917 | PRKAR2B;BUB1B |
| Liver | Resolution of Sister Chromatid Cohesion | 1/126 | 0.320295 | 0.377727011 | 0 | 0 | 2.641866667 | 3.007800471 | BUB1B |
| Liver | MHC Class II Antigen Presentation | 1/126 | 0.320295 | 0.377727011 | 0 | 0 | 2.641866667 | 3.007800471 | DNM1 |
| Liver | Reproduction | 1/127 | 0.322381 | 0.378880851 | 0 | 0 | 2.620767196 | 2.966763425 | CXCR4 |
| Liver | Platelet Degranulation | 1/128 | 0.324461 | 0.38001941 | 0 | 0 | 2.6 | 2.926533696 | F13A1 |
| Liver | Cardiac Conduction | 1/130 | 0.328602 | 0.383556011 | 0 | 0 | 2.559431525 | 2.848411166 | CACNA1C |
| Liver | Response to Elevated Platelet Cytosolic Ca2+ | 1/133 | 0.334767 | 0.389422719 | 0 | 0 | 2.500883838 | 2.736769237 | F13A1 |
| Liver | Beta-catenin Independent WNT Signaling | 1/134 | 0.336809 | 0.390470604 | 0 | 0 | 2.481954887 | 2.700957549 | FZD2 |
| Liver | Signaling by Hedgehog | 1/136 | 0.340876 | 0.392523916 | 0 | 0 | 2.444938272 | 2.63133159 | PRKAR2B |
| Liver | MITF-M-regulated Melanocyte Development | 1/136 | 0.340876 | 0.392523916 | 0 | 0 | 2.444938272 | 2.63133159 | MYO5A |
| Liver | RHO GTPases Activate Formins | 1/139 | 0.34693 | 0.398154533 | 0 | 0 | 2.391425121 | 2.531639937 | BUB1B |
| Liver | FCGR3A-mediated Phagocytosis | 1/142 | 0.352929 | 0.401002628 | 0 | 0 | 2.340189125 | 2.437278355 | MYO5A |
| Liver | Parasite Infection | 1/142 | 0.352929 | 0.401002628 | 0 | 0 | 2.340189125 | 2.437278355 | MYO5A |
| Liver | Leishmania Phagocytosis | 1/142 | 0.352929 | 0.401002628 | 0 | 0 | 2.340189125 | 2.437278355 | MYO5A |
| Liver | Regulation of Actin Dynamics for Phagocytic Cup Formation | 1/143 | 0.354917 | 0.401925748 | 0 | 0 | 2.323591549 | 2.406942575 | MYO5A |
| Liver | Neuronal System | 2/411 | 0.357699 | 0.403739742 | 0 | 0 | 1.618664788 | 1.664088961 | PRKAR2B;GRIA3 |
| Liver | Toll Like Receptor 4 (TLR4) Cascade | 1/148 | 0.364766 | 0.410361222 | 0 | 0 | 2.24399093 | 2.263066007 | DNM1 |
| Liver | Factors Involved in Megakaryocyte Development and Platelet Production | 1/158 | 0.384021 | 0.427129101 | 0 | 0 | 2.1 | 2.009820806 | PRKAR2B |
| Liver | Antiviral Mechanism by IFN-stimulated Genes | 1/158 | 0.384021 | 0.427129101 | 0 | 0 | 2.1 | 2.009820806 | SPHK1 |
| Liver | HIV Life Cycle | 1/158 | 0.384021 | 0.427129101 | 0 | 0 | 2.1 | 2.009820806 | CXCR4 |

| | | | | | | | | | |
|---|---|---|---|---|---|---|---|---|---|
| Liver | Anti-inflammatory Response Favouring Leishmania Parasite Infection | 1/159 | 0.385915 | 0.427129101 | 0 | 0 | 2.086603376 | 1.986735256 | PRKAR2B |
| Liver | Leishmania Parasite Growth and Survival | 1/159 | 0.385915 | 0.427129101 | 0 | 0 | 2.086603376 | 1.986735256 | PRKAR2B |
| Liver | Fcgamma Receptor (FCGR) Dependent Phagocytosis | 1/168 | 0.402702 | 0.44427174 | 0 | 0 | 1.973253493 | 1.79478715 | MYO5A |
| Liver | Toll-like Receptor Cascades | 1/174 | 0.413643 | 0.454873774 | 0 | 0 | 1.904238921 | 1.680972892 | DNM1 |
| Liver | Separation of Sister Chromatids | 1/177 | 0.419038 | 0.459330564 | 0 | 0 | 1.871496212 | 1.627813734 | BUB1B |
| Liver | G2 M Transition | 1/184 | 0.43144 | 0.471413225 | 0 | 0 | 1.799271403 | 1.512517528 | PRKAR2B |
| Liver | Mitotic G2-G2 M Phases | 1/186 | 0.434935 | 0.473718691 | 0 | 0 | 1.77963964 | 1.481655646 | PRKAR2B |
| Liver | TCF Dependent Signaling in Response to WNT | 1/189 | 0.440138 | 0.477863884 | 0 | 0 | 1.750975177 | 1.436968327 | FZD2 |
| Liver | Cilium Assembly | 1/200 | 0.458816 | 0.496566368 | 0 | 0 | 1.653266332 | 1.288070805 | PRKAR2B |
| Liver | Muscle Contraction | 1/204 | 0.465455 | 0.502162353 | 0 | 0 | 1.620361248 | 1.239156298 | CACNA1C |
| Liver | Cell Cycle, Mitotic | 2/516 | 0.469286 | 0.504703276 | 0 | 0 | 1.281078942 | 0.969192522 | PRKAR2B;BUB1B |
| Liver | Disease | 7/2131 | 0.477902 | 0.512358703 | 0 | 0 | 1.087265467 | 0.802782394 | SPON1;VCAN;COL1A2;PRKAR2B;CXCR4;MYO5A;EREG |
| Liver | Mitotic Anaphase | 1/221 | 0.492787 | 0.52666646 | 0 | 0 | 1.493863636 | 1.057173796 | BUB1B |
| Liver | Mitotic Metaphase and Anaphase | 1/222 | 0.494352 | 0.526692371 | 0 | 0 | 1.487028658 | 1.047623959 | BUB1B |
| Liver | Immunoregulatory Interactions Between a Lymphoid and a non-Lymphoid Cell | 1/223 | 0.495911 | 0.526713081 | 0 | 0 | 1.480255255 | 1.038189619 | COL1A2 |
| Liver | HIV Infection | 1/227 | 0.502102 | 0.531637532 | 0 | 0 | 1.453761062 | 1.001571253 | CXCR4 |
| Liver | Cell Cycle Checkpoints | 1/261 | 0.551793 | 0.582448374 | 0 | 0 | 1.261474359 | 0.750049881 | BUB1B |
| Liver | PIP3 Activates AKT Signaling | 1/281 | 0.578709 | 0.608980386 | 0 | 0 | 1.170178571 | 0.640034756 | EREG |
| Liver | Signaling by WNT | 1/287 | 0.58647 | 0.615253427 | 0 | 0 | 1.14527972 | 0.611160628 | FZD2 |
| Liver | RHO GTPase Effectors | 1/291 | 0.591565 | 0.618542724 | 0 | 0 | 1.129252874 | 0.592839702 | BUB1B |
| Liver | Cell Cycle | 2/649 | 0.593222 | 0.618542724 | 0 | 0 | 1.010766772 | 0.527808402 | PRKAR2B;BUB1B |
| Liver | Organelle Biogenesis and Maintenance | 1/297 | 0.599092 | 0.622764353 | 0 | 0 | 1.106024775 | 0.566660804 | PRKAR2B |
| Liver | Diseases of Signal Transduction by Growth Factor Receptors and Second Messengers | 1/451 | 0.751772 | 0.77910918 | 0 | 0 | 0.721814815 | 0.205949771 | EREG |
| Liver | Metabolism of Proteins | 5/2067 | 0.769091 | 0.79464982 | 0 | 0 | 0.774083761 | 0.203232673 | SPON1;VCAN;SPHK1;TNC;MYO5A |
| Liver | Infectious Disease | 3/1311 | 0.771999 | 0.795251593 | 0 | 0 | 0.73675261 | 0.190651325 | PRKAR2B;CXCR4;MYO5A |
| Liver | Post-translational Protein Modification | 3/1457 | 0.831017 | 0.853476898 | 0 | 0 | 0.657579567 | 0.1217213 | SPON1;VCAN;TNC |
| Liver | Signaling by Rho GTPases | 1/672 | 0.876065 | 0.897049056 | 0 | 0 | 0.47858917 | 0.063324274 | BUB1B |
| Liver | Signaling by Rho GTPases, Miro GTPases and RHOBTB3 | 1/688 | 0.88218 | 0.900613602 | 0 | 0 | 0.467054828 | 0.058549607 | BUB1B |
| Liver | Transport of Small Molecules | 1/724 | 0.894873 | 0.910853336 | 0 | 0 | 0.44296911 | 0.049201892 | PRKAR2B |
| Liver | Metabolism of RNA | 1/761 | 0.906517 | 0.91996649 | 0 | 0 | 0.420592105 | 0.041279365 | CD44 |
| Liver | Cellular Responses to Stimuli | 1/887 | 0.937426 | 0.948519794 | 0 | 0 | 0.358408578 | 0.023159453 | PRKAR2B |
| Liver | Viral Infection Pathways | 1/1029 | 0.960323 | 0.968821907 | 0 | 0 | 0.306598573 | 0.012412675 | CXCR4 |
| Liver | Generic Transcription Pathway | 1/1230 | 0.979302 | 0.985062818 | 0 | 0 | 0.253729319 | 0.005306745 | COL1A2 |
| Liver | RNA Polymerase II Transcription | 1/1360 | 0.986463 | 0.989355484 | 0 | 0 | 0.227863625 | 0.003105743 | COL1A2 |
| Liver | Gene Expression (Transcription) | 1/1615 | 0.994164 | 0.994163799 | 0 | 0 | 0.189229657 | 0.001107618 | COL1A2 |
