## Supplementary Table 4 for "Mapping fibrotic microenvironments: single-cell and spatial profiling of Schistosoma mansoni-induced tissue fibrosis"

**Supplementary Table 4. Statistical summary of qPCR gene expression analysis.**

| **Gene** | **t-value** | **df** | **p-value** |
|---|---|---|---|
| *Ccl7* | 10.81 | 4 | < 0.001 |
| *Ccl8* | 19.96 | 4 | < 0.001 |
| *Cd163* | 3.97 | 4 | 0.017 |
| *Chil3* | 18.10 | 4 | < 0.001 |
| *F13a1* | 24.99 | 4 | < 0.001 |
| *Lox* | 9.52 | 4 | 0.001 |
| *Mmp12* | 16.45 | 4 | < 0.001 |
| *Vcan* | 11.24 | 4 | < 0.001 |
| *Bmp10* | -3.23 | 4 | 0.032 |
| *Trpc5* | 1.55 | 4 | 0.196 |
