## Supplementary Table 5 for "Mapping fibrotic microenvironments: single-cell and spatial profiling of Schistosoma mansoni-induced tissue fibrosis"

**Supplementary Table 5. List of primers used for qPCR.**

| Primer name | Sequence (5′ → 3′) | Amplicon size (bp) |
|---|---|---|
| GapDH_F | CATCACTGCCACCCAGAAGACTG | 153 |
| GapDH_R | ATGCCAGTGAGCTTCCCGTTCAG | |
| Ccl7_F | AGACATGAAAACCCCAACTCCA | 90 |
| Ccl7_R | CAGCGGTGAGGAATTTTGCTT | |
| Ccl8_F | AAAGCTGAAGATCCCCCTTCG | 96 |
| Ccl8_R | CCCTGCTTGGTCTGGAAAAC | |
| Cd163_F | CTGCTGTCACTAACGCTCCT | 140 |
| Cd163_R | TTCATTCATGCTCCAGCCGT | |
| Bmp10_F | GCTCCACAATCCAGTCCCATC | 115 |
| Bmp10_R | GGCTTGCTCTTTCTAAACTGCC | |
| Chil3_F | TCTATGCCTTTGCTGGAATGC | 106 |
| Chil3_R | AGCTCAGTGTTCTTGTCTTTCAG | |
| F13a1_F | CCAGAGCCACACTATCCCAG | 92 |
| F13a1_R | TGGCGAAGTGGGTCAAAGAC | |
| Lox_F | AACCCAAATTAGCGAAGCACA | 95 |
| Lox_R | GTGCGTGCTCCTTGGTTTTT | |
| Mmp12_F | ACCAGAGCCACACTATCCCA | 94 |
| Mmp12_R | TTGGCGAAGTGGGTCAAAGA | |
| Trpc5_F | ACCGGGCAGTCGGGC | 116 |
| Trpc5_R | CTGGGCTTCCTTCTTTGCTCTT | |
| Vcan_F | TGTTCCACATCACTGCTCCC | 95 |
| Vcan_R | AAGTTCTCCAACAGTCGCCA | |
